# *Eclipse*: A Python package for alignment of two or more nontargeted LC-MS metabolomics datasets

**DOI:** 10.1101/2023.06.09.544417

**Authors:** Daniel S. Hitchcock, Jesse N. Krejci, Chloe E. Sturgeon, Courtney A. Dennis, Sarah T. Jeanfavre, Julian R. Avila-Pacheco, Clary B. Clish

## Abstract

Nontargeted LC-MS metabolomics datasets contain a wealth of information but present many challenges during analysis and processing. Often, two or more independently processed datasets must be aligned to form a complete dataset, but no available software meets our needs. For this, we have created an open-source Python package called *Eclipse. Eclipse* uses a novel graph-based subalignment approach to handle complex matching scenarios that arise from *n>2* datasets. *Eclipse* is open source (https://github.com/broadinstitute/bmxp) and can be installed via “pip install bmxp”.

## INTRODUCTION

Nontargeted liquid-chromatography tandem mass-spectrometry (LC-MS) is a powerful methodology for inspecting the metabolic state of a biological specimen (Clish 2015). In a routine data processing workflow, feature extraction software converts raw instrument files to tabular datasets, and thousands of features are identified, integrated, and reported with their chromatographic retention time (**RT**) and mass-to-charge ratio (***m/z***) (Smith *et al*. 2006; Pluskal *et al*. 2010). While many features represent redundant ions or chemical contaminants (Mahieu and Patti 2017), a substantial portion arises from yet unannotated compounds of biological significance (Chen *et al*. 2022; Tahir *et al*. 2022; Vatanen *et al*. 2022). A challenge in analyzing nontargeted metabolomics is the concatenation of unknown features among datasets that have been acquired and processed separately, i.e. alignment (Smith, Ventura and Prince 2015). These challenges are exacerbated when *n>2* datasets are introduced, leading to complex matching that can’t be represented in tabular data (**Supplementary Figure 1**). Some solutions exist to align datasets based on feature descriptors (Brunius, Shi and Landberg 2016; Koch *et al*. 2016; Mak *et al*. 2020; Habra *et al*. 2021, 2024; Climaco Pinto *et al*. 2022), but none satisfy all our requirements, specifically it must run robustly in default settings, not produce multiple matches, be written in Python, and align *n>2* datasets, with results being independent of dataset order.

For this, we developed *Eclipse* (https://github.com/broadinstitute/bmxp). *Eclipse* uses a novel graph-based alignment strategy, and the results can be customized for variety of experimental use cases, such as generating a combined dataset for processing or identifying overlapping features in disparate matrices.

### ECLIPSE OVERVIEW

*Eclipse* aligns multiple datasets by running directed subalignments between dataset pairs (e.g., DS1->DS2, DS2->DS1), identifying feature pairs by comparing the descriptors **RT, *m/z***, and average **Intensity** (**Figure 1a**) and producing a feature match table. In each subalignment, one dataset acts as the Source (left of the arrow) and the other as the Target (right of the arrow). It is important to note that each subalignment is distinct, for example, DS1->DS2 is performed independently of the reverse, DS2->DS1. These match tables are then combined in a graph, and tabular outputs can then be generated using a customizable clustering algorithm.

**Figure 1.**
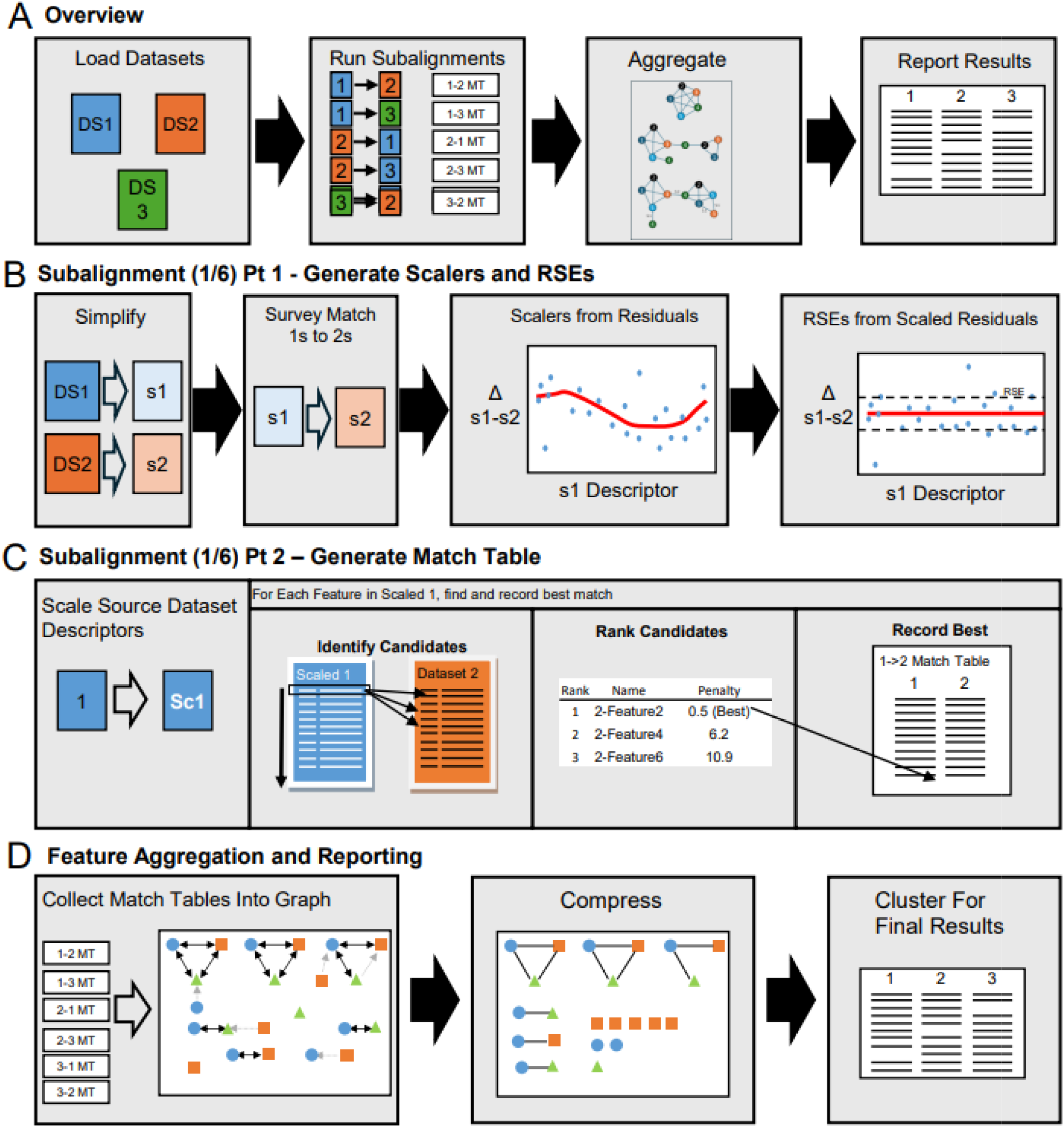
Overview of Eclipse. **A**) High level overview of the Eclipse algorithm with a three-dataset example. **B**) Generation and scalers in the DS1->DS2 subalignment, one of six to be run. Datasets are simplified (s1, s2), then survey matched. Scalers are generated from the residuals of each descriptor (**RT, *m/z*, Intensity**), then subtracted to reveal the residual square error. **C**) Match table generation of the DS1->DS2 subalignment. DS1 is scaled (1->Sc1), then each feature is queried to DS2. DS2 that fall within +/- 6 RSEs of all descriptors are ranked, and the best match is recorded in the DS1->DS2 match table. **D**) Aggregation and reporting of alignment results. Once all subalignments have been run, they are collected into a directed graph. The graph is compressed and clustered to reveal a results table.

#### Subalignment Matching

Subalignment match tables are generated in two steps: 1) surveying the subalignment pair to determine inter-dataset scalers and scoring parameters (**Figure 1b**), and 2) recording best matches (**Figure 1c**). All calculations for feature comparisons are done in a transformed space -- **RT** by addition/subtraction (*linear*), ***m/z*** by *ppm* difference, and **Intensity** by *log10* scaling. *Eclipse* otherwise treats all descriptors identically. Descriptors are customizable and may be removed or alternatives added.

First, each subalignment generates **RT, *m/z***, and **Intensity** scalers to account for inter-dataset trends. *Source* and *Target datasets* are simplified by removing features within a close proximity to neighbors (default thresholds: **RT** +/- 0.5 min, ***m/z*** +/- 15 ppm, **Intensity** +/- 2 log10) (**Supplementary Figure 2**). A survey match is then performed using the same thresholds to identify *Source*->*Target* match pairs. Residuals are calculated from these matches, plotted against the *Source* descriptor values, and modeled using a LOWESS smoothing curve. The scalers are then applied to the residuals to reveal the inter-dataset fluctuations that remain after scaling. Outliers are removed and the standard deviation of the scaled residuals is calculated as the Residual Standard Error (**RSE**), which is used during matching to properly weigh descriptor contributions and determine cutoffs during matching.

Next, the *Source* dataset is scaled to the *Target* dataset, and potential matches are found by identifying *Target* features that fall within ±6 RSE of the Source’s **RT, *m/z***, and **Intensity** (**Figure\ 1c**). Potential matches are ranked by a penalty (**Supplementary Equation 1**), and the best-ranked match, along with its penalty, is recorded in the subalignment match table.

#### Feature Aggregation and Clustering

Once all subalignments have been completed, a Combined Dataset may be produced. All subalignment match tables are loaded into a directed graph (**Figure 1d**), with features as nodes, matches as edges, and match penalties as edge weights. Next, the graph is compressed to an undirected graph, and only bidirectional matches are kept. Penalties from bidirectional edges are summed. A clustering algorithm then ranks valid groups (described below), records the best group, removes it from the graph, and repeats this until no valid groups remain (**Supplementary Figure 3a**). This process ensures that no redundant matches exist, and that dataset order does not influence the results. By default, valid groups are those which contain a member from all datasets and are fully interconnected, i.e. a clique; however, this may be customized. A user may specify the *minimum group size, minimum clique size*, and *diameter*, with 1 enforcing strict cliques (‘clique mode’), 2 allowing one node in a clique to have neighbors, and 3 surveying all clique members for neighbors (**Supplementary Figure 3b**). Groups are ranked by group size (higher is better), clique size (higher is better), number of edges (higher is better), total edge penalty (lower is better). An example can be seen in **Supplementary Figure 4**.

## METHODS

Samples within LC-MS datasets (denoted as DS1 through DS11) were acquired on instruments comprised of Shimadzu Nexera X2 U-HPLCs coupled to Thermo Exactive series orbitrap mass spectrometers. DS1-4 were created from pooled reference samples in distinctly processed human plasma datasets. DS5-9 were derived from datasets of various rodent tissues. All datasets were acquired using the same HILIC-pos method (Mascanfroni *et al*. 2015). Feature extraction was performed using Progenesis QI and features were annotated based on comparison to known LC-MS standards. Dataset information, including specific instrument models and acquisition dates, is summarized in **Supplementary Table 1**. All alignments were performed on an AMD Ryzen 5 3500x Windows 11 PC, running Python 3.12 and *BMXP* version 0.2.4 and *Eclipse* 0.2.3, using default settings unless otherwise noted. The benchmark times did not include file I/O. All data and scripts are available in the Supplementary Information. Spurious matches were reported as the total number of rows (matches) that contained an annotation mismatch.

## RESULTS AND DISCUSSION

### Multi-Batch Alignments – All-By-All

Our primary use case for *Eclipse* is for combining multiple datasets as part of our processing workflow, reporting only features that are found in all datasets and that form a clique. Four human plasma datasets (DS1, DS2, DS3, DS4) were aligned, running in 16 seconds (**Supplementary Code 1**). *Eclipse* identified 3861 features (29% of the smallest dataset, DS2) and correctly matched 95% (362 of 381) of overlapping annotated features. In addition to the overlapping annotations, there were 111 non-overlapping annotations which should not appear in the combined table. Most of these arose from missing features in the nontargeted data. *Eclipse* generated three spurious matches, i.e. rows where annotations were mismatched (including overlapping annotations). This is summarized in **Supplementary Table 4**.

Clustering settings can be relaxed if a user wishes to capture more features. We set the *diameter* to 2 and *minimum clique size* to 3. Compared to the strict clique mode, this matched 4,530 (+669, 34%) features and 96% (367 of 381) of overlapping annotated features, yielding 5 additional annotated features (**Supplementary Code 2**). There were 13 (+10) spurious matches. A visual representation of scaling results DS1->DS2 is explained in **Supplementary Figure 5 and 6**, and all Plasma subalignments can be viewed in **Supplementary Report 1**.

### Five Disparate-Matrix Datasets—One-By-All

*Eclipse* is also used to identify equivalent features across biospecimens of different origins, like tissue types or biological fluids, relative to a reference. To demonstrate, rat plasma (DS5) was aligned to rat gastrocnemius (DS6), rat liver (DS7), rat heart (DS8); and rat white adipose (DS9). We performed a One-By-All alignment (**Supplementary Code 3**), in which DS5 was aligned with all others, but DS6-DS9 were not aligned to each other. This generates hub-spoke type clusters (**Supplementary Figure 7**) which are captured by setting *minimum clique size* and *minimum group size* to 2 and *diameter* to 2. **Intensity** was disabled as well. The alignment finished in 10 seconds. Of the 12,461 features in DS5, 1,600 had matches in all datasets, 3,681 had partial matches, and 7,180 did not have a match in any other dataset. Out of the 140 overlapping features that were present in all datasets, 130 were fully matched and the remaining 10 were partially matched.

### Robust, Recurring Features – All-By-All

One potential use case for Eclipse is to better understand the robustness of feature detection across datasets over time, for example features in plasma datasets from different human cohorts, acquired months or years apart. For this, six plasma datasets (DS1-4, DS10, DS11), collected over 6 years on three different instruments were aligned allowing no minimum group size (*minimum group size* and *minimum clique size* to 1), but still enforcing the requirement of groups being cliques (**Supplementary Code 4**). This ran in 52 seconds. There were 1,435 common features found among all datasets. 6,376 features were found n>=4 datasets (i.e. >50% of datasets). 47,522 clusters of <4 were formed, which are likely fragmented groups left by the strict clique-based clustering. Subalignment scaling and matching results can be viewed in **Supplementary Report 1**.

A similar experiment was conducted using all eleven datasets, allowing for non-clique clustering (*diameter set* to 2) with **Intensity** disabled. This revealed 976 features found in all datasets, and a total of 5334 features found in n>=6 datasets. All results, including *diameter* set to 1, 2, and 3, can be seen in **Supplementary Table 2**. We expect that features found in many datasets are real and robust, and high priority targets for identification. The scaling and matching reports for all 110 subalignments can be viewed in **Supplementary Report 2**.

### Comparison to Other Tools

The most similar tools to *Eclipse* are M2S (Climaco Pinto *et al*. 2022), written in Matlab and *metabCombiner* (Habra *et al*. 2021, 2024), written in R. *metabCombiner* is capable of aligning *n>2* datasets using a step-wise approach, which differs from *Eclipse*’s graph based approach. We aligned DS1-4 using *metabCombiner* in *intersection* mode (**Supplementary Code 5 and 6**). Compared to *Eclipse*’s “clique” mode (*diameter*=1), *metabCombiner* identified the same number of overlapping annotations (362 of 381), but a yielded a much higher number of spurious hits, 26 vs *Eclipse*’s two. We also observed that the results were dependent on dataset order. The annotation-evaluation results, with the six-plasma datasets and alternate parameters, can be viewed in **Supplementary Tables 3 and 4**. We also ran plasma and all datasets in both *union* and *intersection* mode, and the feature counts are reported on **Supplementary Table 2**.

An interesting feature of metabCombiner is its robust handling of various column gradients, a use case distinct from *Eclipse*’s original design. For this we have implemented a “prescaling” option, where a user may provide known descriptor values, such as the retention times of reference metabolites, to correct for large retention time differences before *Eclipse* applies its data driven scaling.

### Benefits of Eclipse’s Graph Based Approach

*Eclipse*’s graph-based approach decouples the matching steps from the aggregation steps, introducing several advantages. For one, results are not dependent on insertion order. Second, feature matching can be visualized. For example, annotation *Sphingosine* was not aligned by either *Eclipse* or *metabCombiner*. Using *Eclipse*’s *explain* method, we can visualize the component and see that it was aligned in all datasets except DS2<->DS4 (**Supplementary Figure 8**). A final advantage is that we may re-cluster with new result params (*minimum clique size, minimum group size, diameter*) without rerunning the subalignments.

## Conclusion

We offer *Eclipse* as a means to combine *n>=2* datasets, especially if a workflow requires symmetrical results (independent of order) and a Python environment. *Eclipse* is critical to our workflow and we intend to support it indefinitely. *Eclipse* is open source, and we welcome feedback and new feature requests from the metabolomics community. The code, instructions, and examples can be found as part of a larger processing toolset used by the Metabolomics Platform at the Broad (including *Gravity* – feature clustering by RT/correlation, and *Blueshift* -- drift correction) at https://github.com/broadinstitute/bmxp.

## Supporting information

Supplementary Reports and Code

## Notes

### Competing Interest Statement

The authors have declared no competing interest.

### Summary of Updates

The manuscript has been revised to include a comparison to current state of the art, as well as a description of a new clustering algorithm.

