## Supplementary Information for "*Eclipse*: A Python package for alignment of two or more nontargeted LC-MS metabolomics datasets"

TOC

1. Pg 2. Supplementary Tables
2. Pg 5. Supplementary Figures
3. Pg 14. Supplementary Equation 1
4. Pg 15. Supplementary Codes
5. Pg 16. Supplementary Report 1 and 2

**Supplementary Table 1.** A list of datasets used in the manuscript. All datasets include a feature list with metadata and are attached as CSV files.

|  | Name | Sample Matrix | # Samples | System | Date | Features | Annotations |
| --- | --- | --- | --- | --- | --- | --- | --- |
| Cohort 1 | DS1 | Human Plasma | 1001 | Exactive Plus 1 | 10/07/19 | 13361 | 432 |
|  | DS2 | Human Plasma | 942 | Exactive Plus 1 | 10/31/19 | 14135 | 436 |
|  | DS3 | Human Plasma | 535 | Exactive Plus 1 | 12/03/19 | 13527 | 423 |
|  | DS4 | Human Plasma | 844 | Exactive Plus 1 | 01/10/20 | 15126 | 448 |
| Cohort 2 | DS5 | Rat Plasma | 150 | Q Exactive HF | 08/30/22 | 12459 | 193 |
|  | DS6 | Rat Gastrocnemius | 150 | Q Exactive HF | 09/06/22 | 9413 | 175 |
|  | DS7 | Rat Liver | 150 | Q Exactive HF | 09/27/22 | 13451 | 196 |
|  | DS8 | Rat Heart | 150 | Q Exactive HF | 10/26/22 | 12647 | 186 |
|  | DS9 | Rat White Adipose | 150 | Q Exactive HF | 10/07/22 | 13012 | 177 |
| Cohort 3 | DS10 | Human Plasma | 1102 | Exactive Plus 2 | 02/03/17 | 9460 | 418 |
| Cohort 4 | DS11 | Human Plasma | 839 | Q Exactive HF | 08/08/23 | 18392 | 450 |

**Supplementary Table 2.** A summary of All-By-All runs of Plasma and All datasets. Each column lists the number of features found and their cluster sizes (i.e. how many datasets a feature was found in), and the total number of reported features, including singletons on the bottom row. *Eclipse* was run at various diameters, and *metabCombiner* was run in Union and Intersection mode. In *Eclipse*, minimum group and clique size was set to 1.

|  | **Plasma** | | | | | **All Datasets** | | | | |
| --- | --- | --- | --- | --- | --- | --- | --- | --- | --- | --- |
|  | **Eclipse** | | | **metabCombiner** | | **Eclipse** | | | **metabCombiner** | |
| **Time (S)** | **40 s** | | | **18 s** | **2.8** | **92 s** |  |  | **37 s** | **5.9 s** |
| **Cluster Size** | **D=1** | **D=2** | **D=3** | **Union** | **Inter.** | **D=1** | **D=2** | **D=3** | **Union** | **Inter.** |
| 11 | - | - | - | - | - | 630 | 976 | 997 | 606 | 410 |
| 10 | - | - | - | - | - | 400 | 517 | 529 | 381 | - |
| 9 | - | - | - | - | - | 465 | 565 | 594 | 512 | - |
| 8 | - | - | - | - | - | 560 | 717 | 724 | 623 | - |
| 7 | - | - | - | - | - | 847 | 1004 | 1023 | 1018 | - |
| 6 | 2352 | 2964 | 2984 | 2422 | 2880 | 1411 | 1555 | 1536 | 1759 | - |
| 5 | 1578 | 1709 | 1733 | 1830 | - | 1862 | 2013 | 2020 | 2300 | - |
| 4 | 1814 | 1978 | 1982 | 2367 | - | 3365 | 3561 | 3526 | 4280 | - |
| 3 | 2819 | 2762 | 2694 | 3464 | - | 5156 | 4634 | 4461 | 5592 | - |
| 2 | 7333 | 5471 | 5452 | 6715 | - | 12735 | 9007 | 9024 | 10632 | - |
| 1 | 31620 | 30532 | 30518 | 21419 | - | 47285 | 45673 | 45576 | 31544 | - |
| **Total** | **47516** | **45416** | **45363** | **38217** | **2880** | **74716** | **70222** | **70010** | **59247** | **410** |

**Supplementary Table 4.** An evaluation of alignment quality of four plasma datasets, DS1-4. Both *Eclipse* and *metabCombiner* were evaluated. *Eclipse* was run at various diameters and minimum clique sizes (*D=1*, *minimum clique size=4*; *D=2, minimum clique size=3*; *D=3*, *minimum clique size=2*). *metabCombiner* was run in *Intersection* and *Union* mode, and datasets were aligned in both ascending (DS1-4) or descending (DS4-1) order. Only complete groups, i.e. a match was found in all datasets, were evaluated. The benchmark time, number of features found, number of correct overlapping annotations, and number of spurious hits is shown.

| **Cluster Size** | **Eclipse** | **Eclipse** | **Eclipse** | **metabC. Ascending** | **metabC. Descending** | **metabC. Ascending** | **metabC. Descending** |
| --- | --- | --- | --- | --- | --- | --- | --- |
| **Option** | **D=1** | **D=2** | **D=3** | **Intersection** | **Intersection** | **Union** | **Union** |
| Time (s) | 16 | 16 | 17 | 2.8 | 7.8 | 18 | 5.2 |
| Features (*n=4*) | 3861 | 4530 | 4633 | 4622 | 4647 | 4295 | 4396 |
| Annotations (381) | 362 | 367 | 369 | 362 | 361 | 351 | 357 |
| Spurious Hits | 3 | 13 | 14 | 26 | 26 | 20 | 20 |

**Supplementary Table 5.** An evaluation of alignment quality of six plasma datasets, DS1-4, 10, and 11. Both *Eclipse* and *metabCombiner* were evaluated. *Eclipse* was run at various diameters and minimum clique sizes (*D=1*, *minimum clique size=6*; *D=2, minimum clique size=5*; *D=3*, *minimum clique size=4*). *metabCombiner* was run in *Intersection* and *Union* mode, and datasets were aligned in both ascending (DS1, 2, 3, 4, 10, 11) or descending (DS11, 10, 4, 3, 2, 1) order. Only complete groups, i.e. a match was found in all datasets, were evaluated. The benchmark time, number of features found, number of correct overlapping annotations, and number of spurious hits is shown.

| **Cluster Size** | **Eclipse** | **Eclipse** | **Eclipse** | **metabC. Ascending** | **metabC. Descending** | **metabC. Ascending** | **metabC. Descending** |
| --- | --- | --- | --- | --- | --- | --- | --- |
| **Option** | **D=1** | **D=2** | **D=3** | **Intersection** | **Intersection** | **Union** | **Union** |
| Time (s) | 35 | 35 | 35 | 3.6 | 3.2 | 23 | 11 |
| Features (*n=6*) | 2352 | 2807 | 2952 | 2879 | 1836 | 2422 | 2381 |
| Annotations (336) | 309 | 325 | 328 | 308 | 112 | 279 | 283 |
| Spurious Hits | 2 | 11 | 15 | 36 | 32 | 25 | 22 |

**Supplementary Figure 1**

**A**

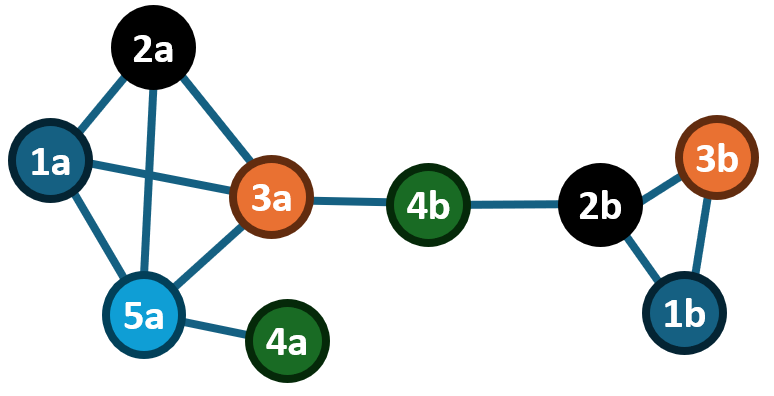

**B. Record All Maximal Cliques**

| DS1 | DS2 | DS3 | DS4 | DS5 |
| --- | --- | --- | --- | --- |
| 1a | 2a | 3a | - | 5a |
| - | - | - | 4a | 5a |
| - | - | 3a | 4b | - |
| 1b | 2b | 3b | - | - |
| - | 2b | - | 4b | - |

**C. Record Maximal Cliques w/o duplicates**

| DS1 |  | DS2 | DS3 | DS4 | DS5 |
| --- | --- | --- | --- | --- | --- |
| 1a |  | 2a | 3a | - | 5a |
| - |  | - | - | 4a | - |
| - |  | - | - | 4b | - |
| 1b |  | 2b | 3b | - | - |

**D. Allow non-clique groups**

| DS1 | DS2 | DS3 | DS4 | DS5 |
| --- | --- | --- | --- | --- |
| 1a | 2a | 3a | 4a | 5a |
| 1b | 2b | 3b | 4b | - |

**Supplementary Figure 1**. Various outcomes of creating tabular data from 5 aligned datasets. **A)** A graph representing the matches of nine features among five datasets, represented as a graph. Each node represents a unique feature and is labeled by its dataset number (1-5). An edge indidcates the features correspond with each other. **B-D)** Results tables produced using various approaches, where a row represents an aligned feature. **B)** All maximal cliques are recorded, preserving all edges, however the table contains duplicated features, such as 3a, 4b, 2b. C) All maximal cliques are recorded, and duplicates are resolved by a ranking algorithm. Some connection information is lost, such as 5a<->4a. **D)** Largest groups are ranked and recorded, resulting in fewer, but more populated, rows. Some connection information is lost, such as 3a<->4b, and the table suggests that some connections exist that did not, such as 4a<->3a

**Supplementary Figure 2**

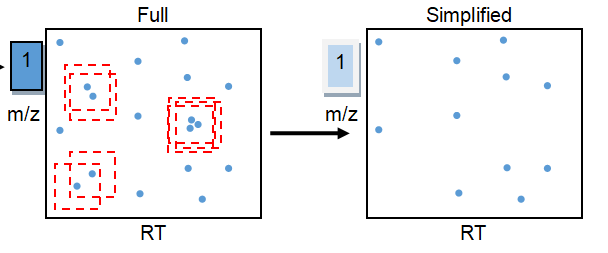

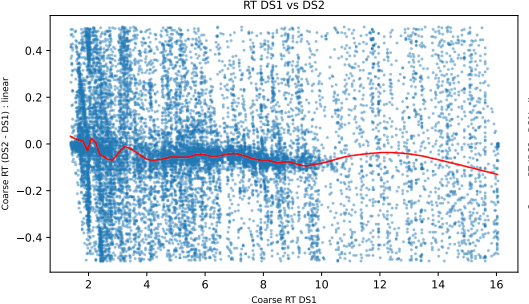

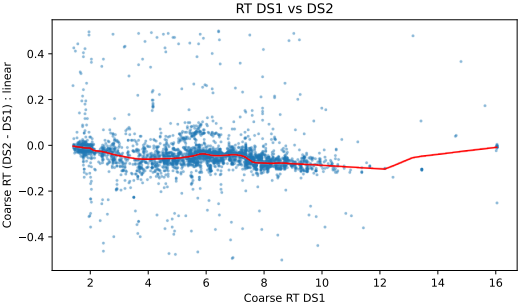

**Supplementary Figure 2**. Dataset simplification. **Top Row:** Datasets are simplified by removing all features (blue dots) that fall within a threshold (red dashed square) of another feature. **Bottom Row:** The survey match results of DS1->DS2 without and with simplification. Simplification is required since the unsimplified results are noisy and difficult to model (left), however the simplified results reveal a strong trend (right).

**Supplementary Figure 3**

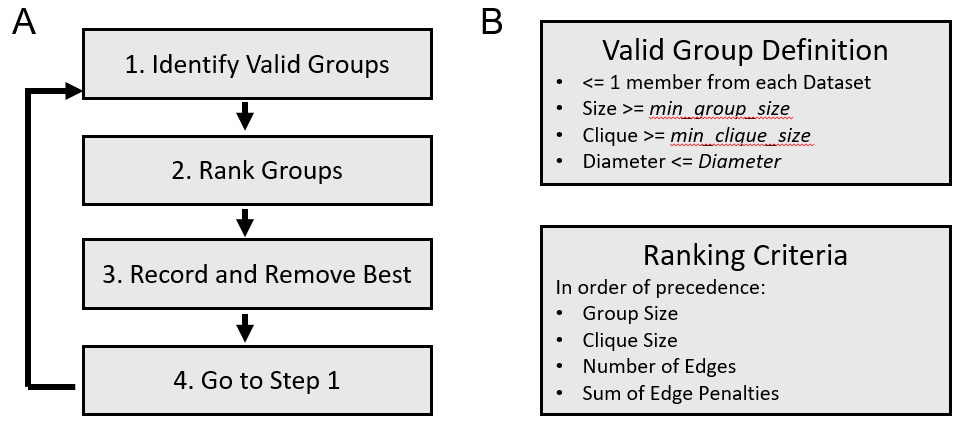

**Supplementary Figure 3**. Diagram of the clustering algorithm to report a dataset from the graph. **A)** A flow chart showing the clustering algorithm. Valid groups are ranked and removed until none remain. **B)** Definitions for a Valid Group and the Ranking Criteria.

**Supplementary Figure 4**

**A**

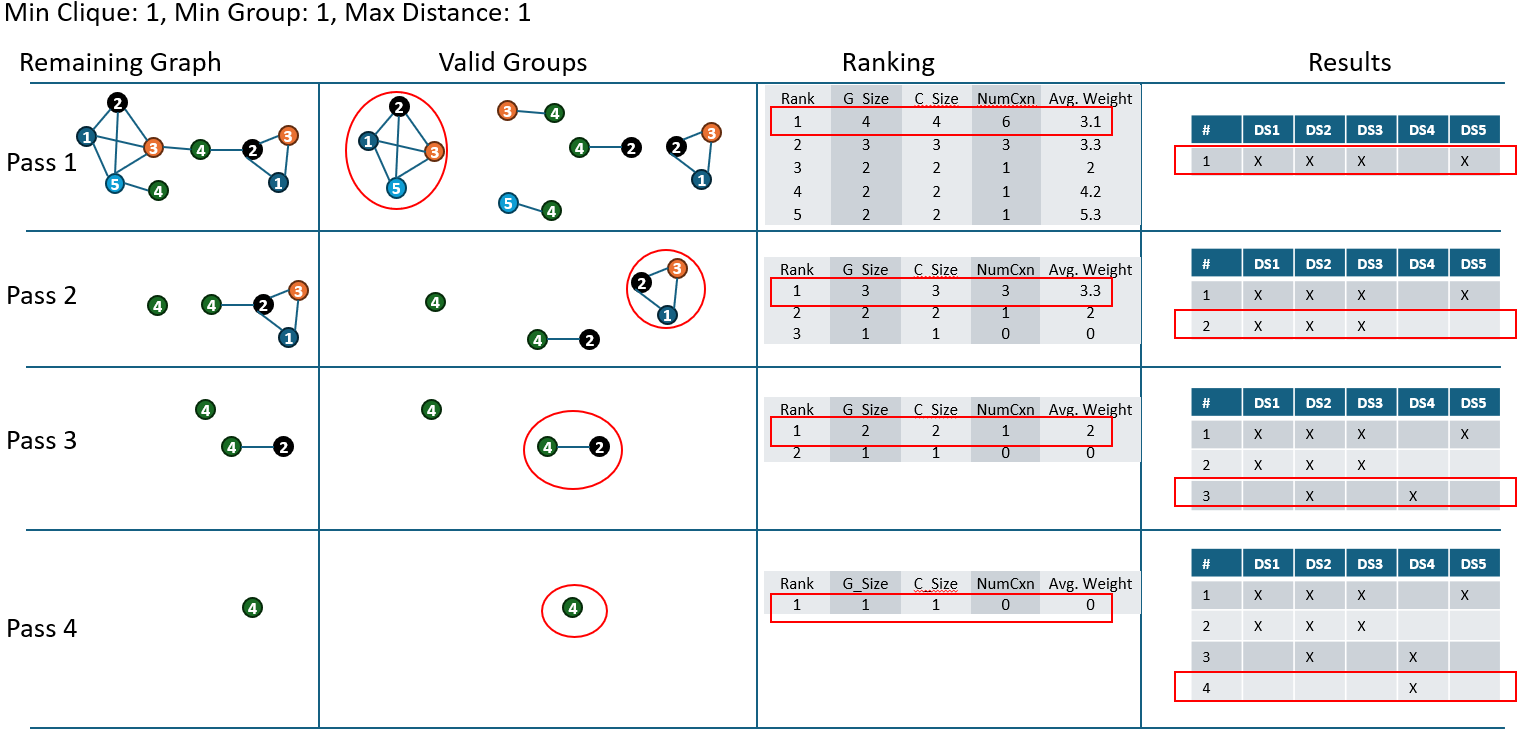

**B**

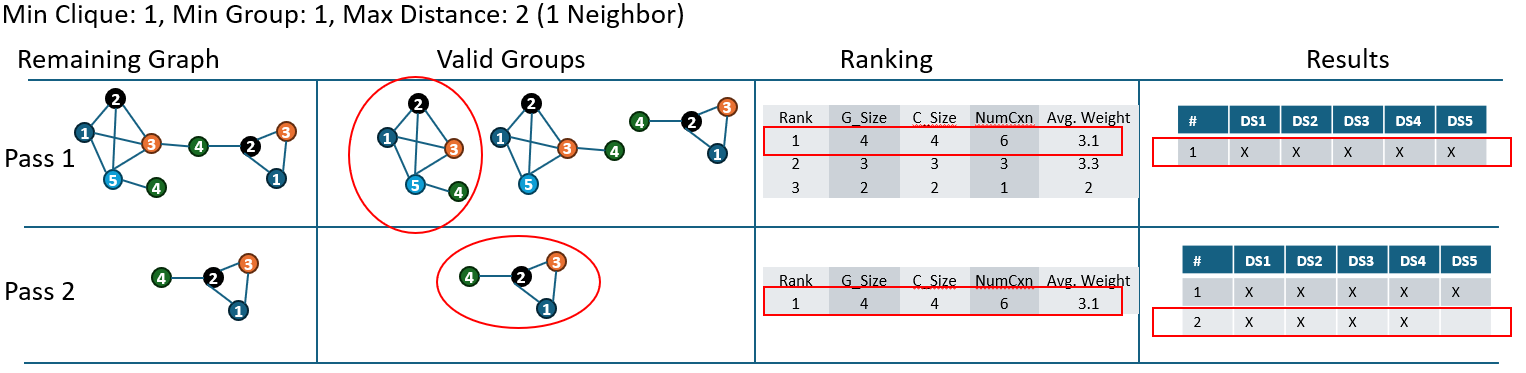

**Supplementary Figure 4**. An example of the clustering algorithm, showing how features would be reported under two settings. A table displays the clustering algorithm, with the columns being the steps in the algorithm, and the rows each being an iteration. **Column 1** shows the component, where each node represents a feature, and a number denotes its corresponding dataset. Edges weights are omitted for clarity. First, valid subgraphs are identified based on the settings (**Column 2**). Next, they are ranked (**Column 3**), and the top-ranking group recorded into the growing feature table (**Column 4**). The nodes are removed from the graph and the remaining subgraph is shown. **Caption A** shows the algorithm with *minimum clique size* and *minimum group size* to 1 and diameter left at 1. **Caption B** shows with the algorithm with a diameter of 2, allowing for one member to have non-clique neighbors.

**Supplementary Figure 5**

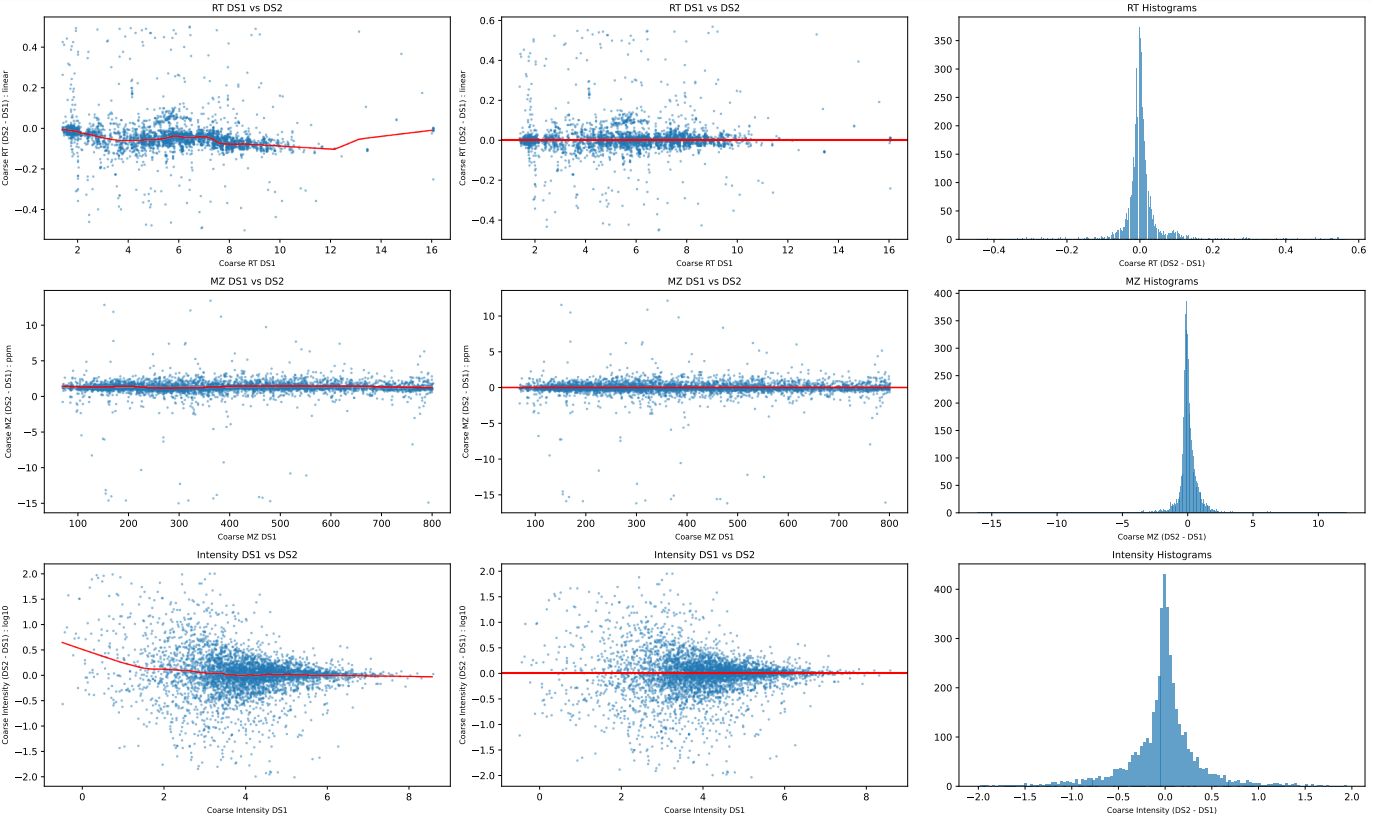

**Supplementary Figure 5**. Survey (coarse) matching results from subalignment DS1->DS2 used for determining scalers, and RSEs. This is the output of *Eclipse*’s report function. Each descriptor (RT, *m/z*, intensity) is plotted as three panels comprising a row. Blue dots represent a coarse feature match, with the source dataset descriptor value on the X axis and the difference (residual) on the Y axis. RT is modelled in *linear* mode, *m/z* in *ppm* and intensity in *log*. A red line represents the scalers, determined from a LOWESS smoothing curve. The first column shows unscaled feature pair differences, the second shows the differences after applying scalers, and the third shows a histogram of the second column. These values are used to calculate the descriptor RSEs. They are RT: 0.020 min, *m/z*: 0.39 ppm, intensity: 0.23 log10

**Supplementary Figure 6**

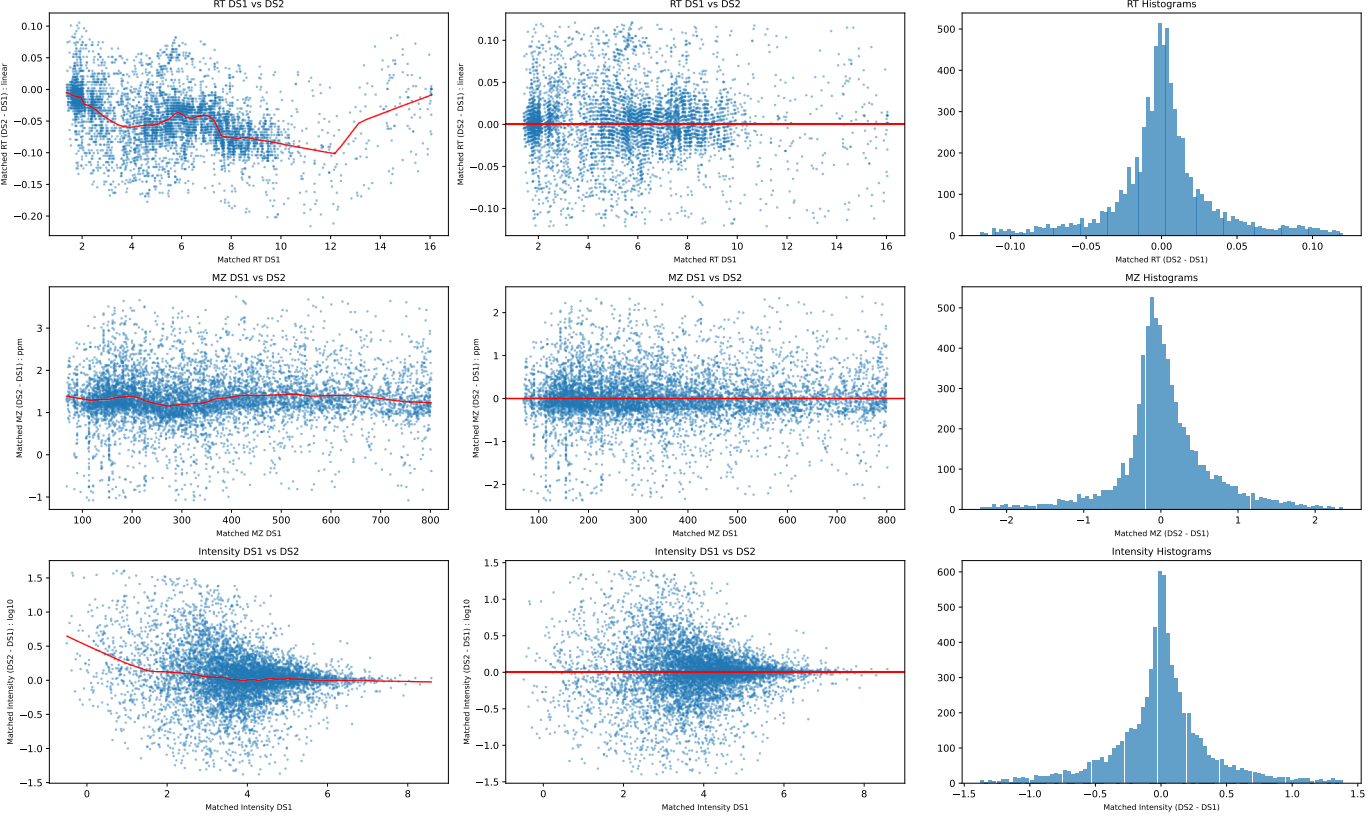

**Supplementary Figure 6.** Subalignment DS1->DS2 match results. Like **Supplementary Figure 4**, each descriptor (RT, *m/z*, intensity) is plotted as three panels comprising a row. Blue dots represent a one-way feature match, with the source dataset descriptor values on the X axis and the difference (residual) on the Y axis. The first column shows unscaled feature pair differences, the second shows the differences after applying scalers, and the third shows a histogram of the second column. The cutoffs shown in the second column are calculated from 6x (default) the descriptor’s RSE, which are RT: +/-0.12 min, *m/z*: +/-2.4 ppm, intensity: +/-1.39 log10.

**Supplementary Figure 7**

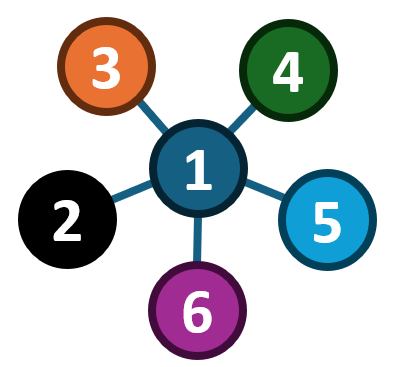

**Supplementary Figure 7.** The hub-spoke type groups that form from One-By-All alignments. Each node represents a feature from a Dataset, denoted by the number. In this example, Dataset1 has a matching feature in all datasets, however Datasets 2-6 were not aligned to each other. For this to be a valid group, the diameter must be set to 2.

**Supplementary Figure 8**

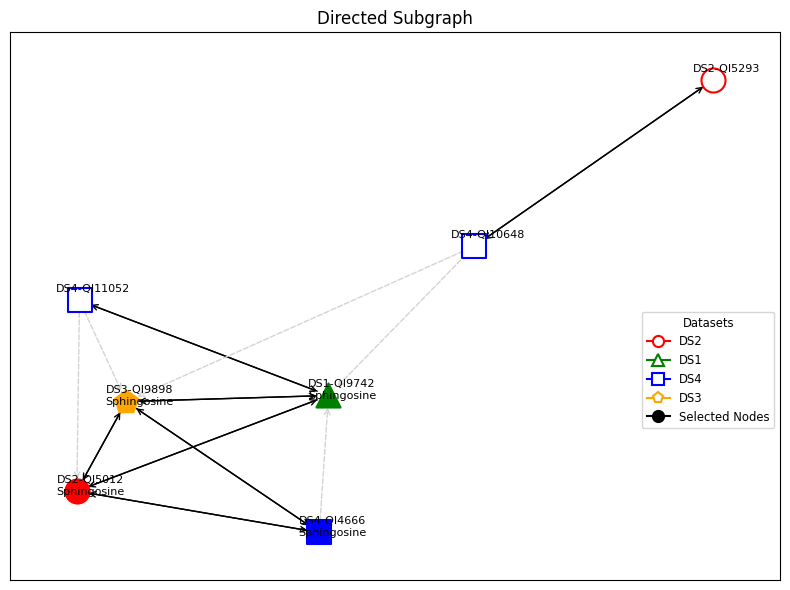

**Supplementary Figure 8**. The output of *Eclipse*’s *explain* function, displaying a visual explanation of why a specific feature, Spingosine, was not reported during the alignment of DS1-4. Each node is a feature, labeled by its name and dataset and colored/shaped by its dataset. Bidirectional matches are solid lines, and directed matches are dashed gray lines. In this example, There was no DS1->DS4 alignment of Sphingosine. Instead, DS1 points to the incorrect feature in DS4, QI11052, forming a bidrectional match. Given the necessity to form a clique (*minimum clique size = 4*), this would not be recorded. However, if the diameter is set to 2 and *minimum clique size* to 3, this group would be valid.

**Supplementary Equation 1. The scoring function used when descriptors are RT, m/z, and Intensity**

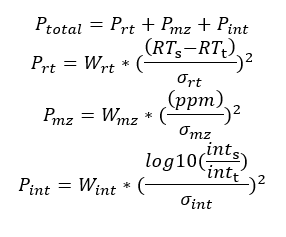

*P_total_* is the total penalty, *P_rt_*, *P_mz_*, and *P_int_* are the individual penalties for each feature descriptor, *W_rt_*, *W_mz_*, and *W_int_* are the user-selected weights of a term (default 1), *RT_s_* and *RT_t_* are the retention times for the features being compared, *ppm* is the *m/z* difference in parts-per-million, *int_s_* and *int_t_* are the intensities, and *σ_rt_*, *σ_mz_*, *σ_in_*_t_ are the RSEs of the scaled residuals between the datasets.

**Supplementary Code 1**

Attached as “Supplementary Code 1.ipynb”. This file contains Python code to execute and evaluate an All-by-All alignment of DS1-4. This experiment is for combining batched studies to produce a single dataset. Groups must be cliques (*diameter*=1),and must contain a member from all datasets (*minimum group size* and *minimum clique size* = 4).

**Supplementary Code 2**

Attached as “Supplementary Code 2.ipynb”. This file contains Python code to execute and evaluate an All-by-All alignment of DS1-4. Like Supplementary Code 1, the purpose is to combine multi-batch datasets to produce a single dataset, however this has less stringent clustering conditions. Groups may be non-cliques (*diameter*=2), but must contain a member from all datasets (*minimum group size* = 4, *minimum clique size* = 3).

**Supplementary Code 3**

Attached as “Supplementary Code 3.ipynb”. This file contains Python code to execute and evaluate a One-by-all alignment of DS5 to DS6-9. The purpose of this is to identify features in DS6-9 (rodent tissue) that correspond to those in DS5 (plasma). This produces hub-spoke type groups. Non clique groups are allowed (*diameter* = 2) and the *minimum group size* is 2.

**Supplementary Code 4**

Attached as “Supplementary Code 4.ipynb”. This file contains Python code to execute and evaluate All-by-All alignments of the following combinations and conditions:

- Six plasma datasets or all 11 datasets.
- Diameter=1, 2, or 3
- No minimum group or clique size
- Intensity disabled for Plasma

**Supplementary Code 5**

Attached as “Supplementary Code 5.r”. This file contains R code to combine 4 batches into single dataset, as comparison to Supplementary Code 1. Default settings were used, and “Intersection” mode was specified to simulate Eclipse’s “clique-mode” (*diameter* = 1). Additionally, this code was modified to run a comparison for the plasma/all dataset all-by-all alignments, summarized in **Supplementary Table 2**. This specific code is not reported.

**Supplementary Code 6**

Attached as “Supplementary Code 6.ipynb”. This file contains Python code to evaluate the output of “Supplementary Code 5.r”.

**Supplementary Reports 1 and 2**

Attached as “Supplementary Report 1.pdf” and “Supplementary Report 2.pdf”. These are the outputs of the subalignment scaling and matching reports, used to quickly evaluate scaling and matching performance. **Supplementary Report 1** is the results of matching Plasma datasets DS1-4, DS10, and DS11. **Supplementary Report 2** is the alignment of all datasets. Each subalignment has two pages: odd pages contain the survey (denoted as “coarse”) alignment results to determine scalers and RSEs for each feature. Even pages contain the actual matching results. Each row on a page is a descriptor. The first column contains the unscaled results matching results. Each dot represents a matched pair. They are plotted with their Source value on the X axis, and the difference on the Y axis. The red line denotes the scalers, as determined in the survey scan (odd pages). The second column contains the scaled data. The third column is a histogram of the scaled data (second column).

It is important to note that these are not the final reported feature results, rather the individual subalignments. There will be features represented in these reports that are not present in the final data, given that a two way match is needed.
