## Supplementary Reports and Code for "*Eclipse*: A Python package for alignment of two or more nontargeted LC-MS metabolomics datasets": Supplementary Code 1.ipynb

```
# %% [code]
"""
Aligns all features
"""
import time
import logging
import pandas as pd
import numpy as np
from matplotlib import pyplot as plt
%matplotlib inline
import bmxp.eclipse as eclipse

# load files
eclipse.LOGGER.setLevel(logging.WARNING)
files = ['DS1', 'DS2', "DS3", "DS4"]
dfs = [pd.read_csv(f + '.csv') for f in files]
t0 = time.time()

# run alignment
aligner = eclipse.MSAligner(*dfs, names=files)
aligner.align()

# benchmark network deconvolution
results = aligner.results()
print(time.time()-t0)
aligner.to_csv(filepath='DS1,DS2,DS3,DS4-cliques.csv')

# evaluation code
df = pd.read_csv('DS1,DS2,DS3,DS4-cliques.csv')
columns =['Annotation_ID']+ ["Annotation_ID_"+name for name in files[1:]]
matched_ann = 0
matched_ann_set = set()
spurious_ann = 0
for row in df[columns].iterrows():
    results = set(row[1][column] for column in columns)
    if results == {np.nan}:
        continue
    if len(results)==1:
        matched_ann +=1
        matched_ann_set.update(results)
    elif len(results)>1:
        spurious_ann +=1
        
ds1_annotations = dfs[0].loc[pd.notnull(dfs[0]['Annotation_ID']), 'Annotation_ID']
ds2_annotations = dfs[1].loc[pd.notnull(dfs[1]['Annotation_ID']), 'Annotation_ID']
ds3_annotations = dfs[2].loc[pd.notnull(dfs[2]['Annotation_ID']), 'Annotation_ID']
ds4_annotations = dfs[3].loc[pd.notnull(dfs[3]['Annotation_ID']), 'Annotation_ID']
overlapping = set(ds1_annotations) & set(ds2_annotations) & set(ds3_annotations) & set(ds4_annotations)
non_overlapping = (set(ds1_annotations) | set(ds2_annotations) | set(ds3_annotations) | set(ds4_annotations)) - overlapping
print("DS2 Features, Annotations: ", len(dfs[0]), len(ds1_annotations))
print("DS3 Features, Annotations: ", len(dfs[1]), len(ds2_annotations))
print("DS4 Features, Annotations: ", len(dfs[2]), len(ds3_annotations))
print("DS5 Features, Annotations: ", len(dfs[3]), len(ds4_annotations))
print("Total Features: ", len(df))
print("Annotations: ", matched_ann, " out of ", len(overlapping), matched_ann/len(overlapping)*100, "%")
print("Spurious: ", spurious_ann, ", including ", len(non_overlapping), " non-overlapping.")
print('Misaligned overlapping annotations: ', overlapping - matched_ann_set)
sg = aligner.explain(annotation_ids = "Sphingosine", compress=False)

# %% [code]
```
