## Supplementary Reports and Code for "*Eclipse*: A Python package for alignment of two or more nontargeted LC-MS metabolomics datasets": Supplementary Code 3.ipynb

```
# %% [code]
"""
All-by-all
Keep All Plasma
"""

import time
import logging
import pandas as pd
import bmxp.eclipse as eclipse
# silence the info messages
eclipse.LOGGER.setLevel(logging.WARNING)
files = ['DS5','DS6', 'DS7', 'DS8',  'DS9']
dfs = [pd.read_csv(f + '.csv') for f in files]

t0 = time.time()
aligner = eclipse.MSAligner(*dfs, names=files)

# disable intensity
aligner.descriptors = {"RT": "linear", "MZ": "ppm"}
aligner.align(files[1:], files[:1])
aligner.align(files[:1], files[1:])

# just for benchmarking
results = aligner.results('DS5', c_size_or_loss=2, g_size_or_loss=2, diameter=2)
print(time.time()-t0)
aligner.to_csv('DS5', filepath='DS5_one_by_all.csv',  c_size_or_loss=2, g_size_or_loss=2, diameter=2)

# Evaluation
ds5_annotations = dfs[0].loc[pd.notnull(dfs[0]['Annotation_ID']), 'Annotation_ID']
ds6_annotations = dfs[1].loc[pd.notnull(dfs[1]['Annotation_ID']), 'Annotation_ID']
ds7_annotations = dfs[2].loc[pd.notnull(dfs[2]['Annotation_ID']), 'Annotation_ID']
ds8_annotations = dfs[3].loc[pd.notnull(dfs[3]['Annotation_ID']), 'Annotation_ID']
ds9_annotations = dfs[4].loc[pd.notnull(dfs[4]['Annotation_ID']), 'Annotation_ID']
overlapping = set(ds5_annotations) & set(ds6_annotations) & set(ds7_annotations) & set(ds8_annotations)& set(ds9_annotations)


df = pd.read_csv('DS5_one_by_all.csv')
ann_columns =['Annotation_ID']+ ["Annotation_ID_"+name for name in files[1:]]
cmp_columns = ['Compound_ID']+ ["Compound_ID_"+name for name in files[1:]]
whole_found = []
partial_found = []
whole_ann = []
partial_ann = []
for row in df.iterrows():
    # figure out if feature is whole or partial match
    features = row[1].loc[cmp_columns]
    features = features.loc[pd.notnull(features)]
    if len(features) == len(dfs):
        whole_found.append(features.iloc[0])
    elif len(features) >1:
        partial_found.append(features.iloc[0])

    # find correctly matched annotations
    annotations = row[1].loc[ann_columns]
    annotations = annotations.loc[pd.notnull(annotations)]
    annotation_set = set(annotations)
    if len(annotation_set) == 1 and annotations.iloc[0] in overlapping:
        if len(annotations) == len(dfs):
            whole_ann.append(annotations.iloc[0])
        elif len(annotations) > 1:
            partial_ann.append(annotations.iloc[0])

whole_found_set = set(whole_found)
partial_found_set = set(partial_found)

whole_ann_set = set(whole_ann)
partial_ann_set = set(partial_ann)


print("Total Features: ", len(dfs[0]))
print("Results: ", len(df))
print("All features")
print("Whole Matches: ", len(whole_found_set))
print("Partial Matches: ", len(partial_found_set))
print("Zero Matches: ", len(dfs[0])-len(whole_found_set)-len(partial_found_set))
print("Overlapping Annotations")
print("Complete matches: ", len(whole_ann_set))
print("Partial (at least one) matches: ", len(partial_ann_set))
print("Metabolites not found: ", len(overlapping) - len(whole_ann_set)- len(partial_ann_set))


# %% [code]
```
