## Supplementary Reports and Code for "*Eclipse*: A Python package for alignment of two or more nontargeted LC-MS metabolomics datasets": Supplementary Code 4.ipynb

```
# %% [code]
"""
Run all nine datasets
"""
import time
import logging
import pandas as pd
import bmxp.eclipse as eclipse
# silence the info messages
eclipse.LOGGER.setLevel(logging.WARNING)
files = ["DS%s"%i for i in range(1, 12)]
columns = ['Compound_ID', 'RT', 'MZ', "Intensity"]
dfs = [pd.read_csv(f + '.csv')[columns] for f in files]

# fill in a few missing values for intensity
for df in dfs:
    df['Intensity'] = df['Intensity'].replace('#DIV/0!', 1).astype(float)

"""
Plasma Datasets
"""
t0 = time.time()
datasets = [*dfs[:4], dfs[9], dfs[10]]
p_files=[*files[:4], files[9], files[10]]
aligner = eclipse.MSAligner(*datasets, names=p_files)
aligner.align()
aligner.report(filepath='Supplementary Report 1.pdf')
results = aligner.results(c_size_or_loss=1, g_size_or_loss=1)
print(time.time()-t0)
# aligner.report()


# evaluate
results = aligner.results(c_size_or_loss=1, g_size_or_loss=1)
just_table = results.loc[:, p_files]
group_counts = just_table.count(axis=1).value_counts().reset_index()
group_counts.columns = ['Number Present', 'Count']
print("Plasma, Cliques")
print(len(results))
print(group_counts.sort_values(by='Number Present', ascending=False))
results.to_csv('Plasma D1.csv')

results = aligner.results(c_size_or_loss=1, g_size_or_loss=1, diameter=2)
just_table = results.loc[:, p_files]
group_counts = just_table.count(axis=1).value_counts().reset_index()
group_counts.columns = ['Number Present', 'Count']
print("Plasma D2")
print(len(results))
print(group_counts.sort_values(by='Number Present', ascending=False))
results.to_csv('Plasma D2.csv')

results = aligner.results(c_size_or_loss=1, g_size_or_loss=1, diameter=3)
print(len(results))
just_table = results.loc[:, p_files]
group_counts = just_table.count(axis=1).value_counts().reset_index()
group_counts.columns = ['Number Present', 'Count']
print("Plasma D3")
print(len(results))
print(group_counts.sort_values(by='Number Present', ascending=False))
results.to_csv('Plasma D3.csv')

"""
All Datasets
"""
t0 = time.time()
aligner = eclipse.MSAligner(*dfs, names=files)
aligner.descriptors = {"RT": "linear", "MZ": "ppm"}
aligner.align()
aligner.report(filepath='Supplementary Report 2.pdf')
results = aligner.results(c_size_or_loss=1, g_size_or_loss=1, diameter =1)

print(time.time()-t0)
# aligner.report("All.pdf")

# evaluate
just_table = results.loc[:, files]
group_counts = just_table.count(axis=1).value_counts().reset_index()
group_counts.columns = ['Number Present', 'Count']
print("All D1")
print(len(results))
print(group_counts.sort_values(by='Number Present', ascending=False))
results.to_csv('All D1.csv')

results = aligner.results(c_size_or_loss=1, g_size_or_loss=1, diameter =2)
just_table = results.loc[:, files]
group_counts = just_table.count(axis=1).value_counts().reset_index()
group_counts.columns = ['Number Present', 'Count']
print("All D2")
print(len(results))
print(group_counts.sort_values(by='Number Present', ascending=False))
results.to_csv('All D2.csv')

results = aligner.results(c_size_or_loss=1, g_size_or_loss=1, diameter =3)
just_table = results.loc[:, files]
group_counts = just_table.count(axis=1).value_counts().reset_index()
group_counts.columns = ['Number Present', 'Count']
print("All D3")
print(len(results))
print(group_counts.sort_values(by='Number Present', ascending=False))
results.to_csv('All D3.csv')


# %% [code]
```
