## Supplementary Reports and Code for "*Eclipse*: A Python package for alignment of two or more nontargeted LC-MS metabolomics datasets": Supplementary Code 5.R

```
library(metabCombiner)

# set batch options
union<-FALSE
datasetStart<-1
datasetEnd<-4

uOrI<-"Intersection"
if (union == TRUE){
  uOrI <- "Union"
}

datasets <- lapply(paste0("./DS", datasetStart:datasetEnd, ".csv"), function(file) {
  read.delim(file, sep = ",", stringsAsFactors = FALSE)
})

# Process each dataset with metabData and store in 'batches' list
t0 <- Sys.time()
batches <- lapply(datasets, function(dataset) {
  metabData(
    dataset,
    mz = "MZ",
    rt = "RT",
    Q = NULL,
    id = "Compound_ID",
    adduct = NULL,
    samples = "In",
    extra = "Anno",

    misspc = 50,
    measure = "median",
    zero = TRUE,
    duplicate = opts.duplicate()
  )
})

saparam <- selectAnchorsParam(tolmz = 0.002, tolQ = 0.2, tolrtq = 0.1)
fgparam <- fitgamParam(k = 20, iterFilter = 1)
csparam <- calcScoresParam(A = 70, B = 35, C = 0)
rtparam <- reduceTableParam(minScore = 0.5, maxRTerr = 0.33)


combinedRes <- batchCombine(batches = batches, union = union,
                            anchorParam = saparam, fitParam = fgparam,
                            scoreParam = csparam, reduceParam = rtparam)
object <- combinedRes$object
new <- Sys.time() - t0 # calculate difference
print(new) # print in nice format


#if names were set earlier, the names should be returned by this
write2file(object, file = paste0("./Metab - DS", datasetStart, "-", datasetEnd, " - ", uOrI, ".csv"), sep = ",")
```
