## Supplementary Reports and Code for "*Eclipse*: A Python package for alignment of two or more nontargeted LC-MS metabolomics datasets": Supplementary Report 1.pdf

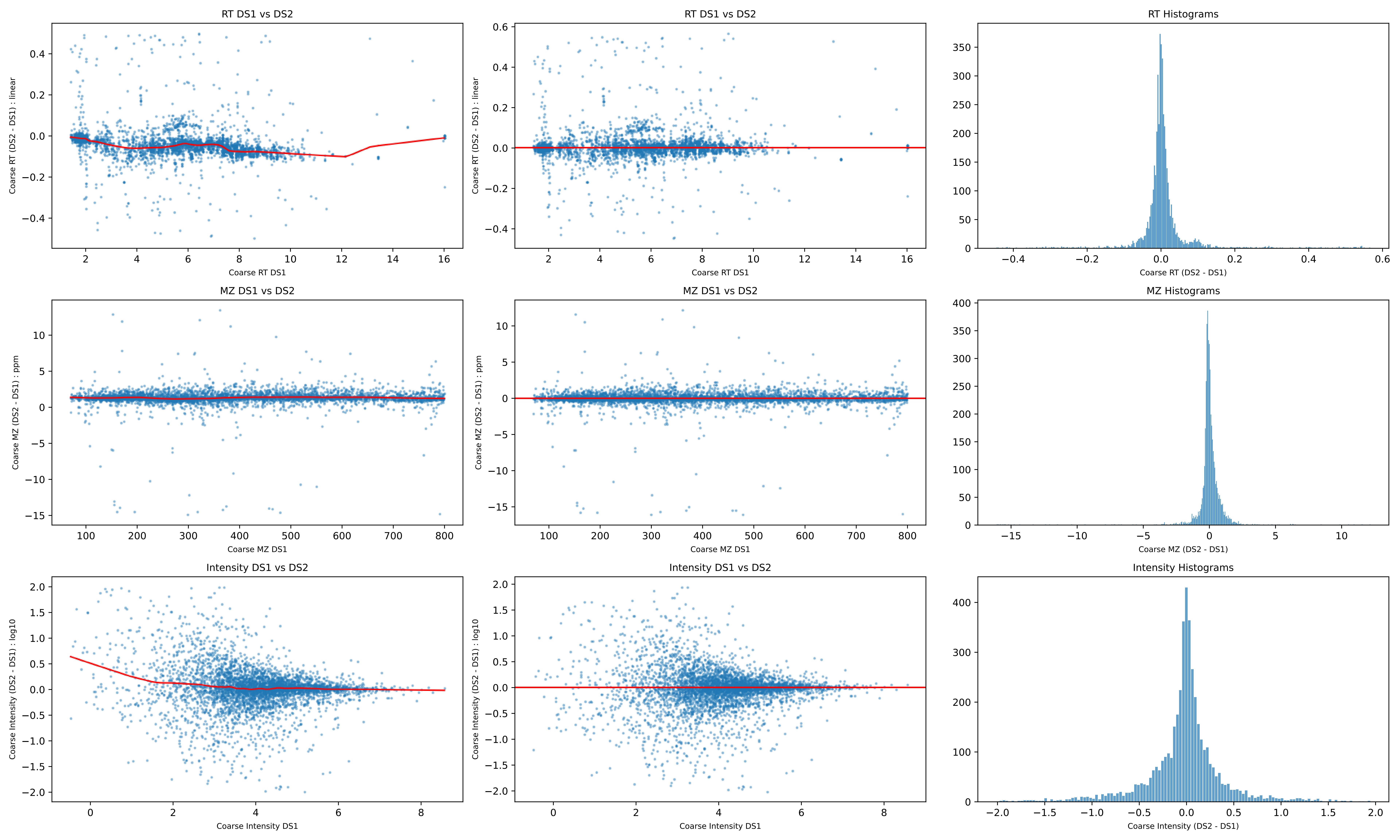
RT DS1 vs DS2
Coarse RT (DS2 - DS1) : linear
Coarse RT DS1
RT DS1 vs DS2
Coarse RT (DS2 - DS1) : linear
Coarse RT DS1
RT Histograms
Coarse RT (DS2 - DS1)
MZ DS1 vs DS2
Coarse MZ (DS2 - DS1) : ppm
Coarse MZ DS1
MZ DS1 vs DS2
Coarse MZ (DS2 - DS1) : ppm
Coarse MZ DS1
MZ Histograms
Coarse MZ (DS2 - DS1)
Intensity DS1 vs DS2
Coarse Intensity (DS2 - DS1) : log10
Coarse Intensity DS1
Intensity DS1 vs DS2
Coarse Intensity (DS2 - DS1) : log10
Coarse Intensity DS1
Intensity Histograms
Coarse Intensity (DS2 - DS1)

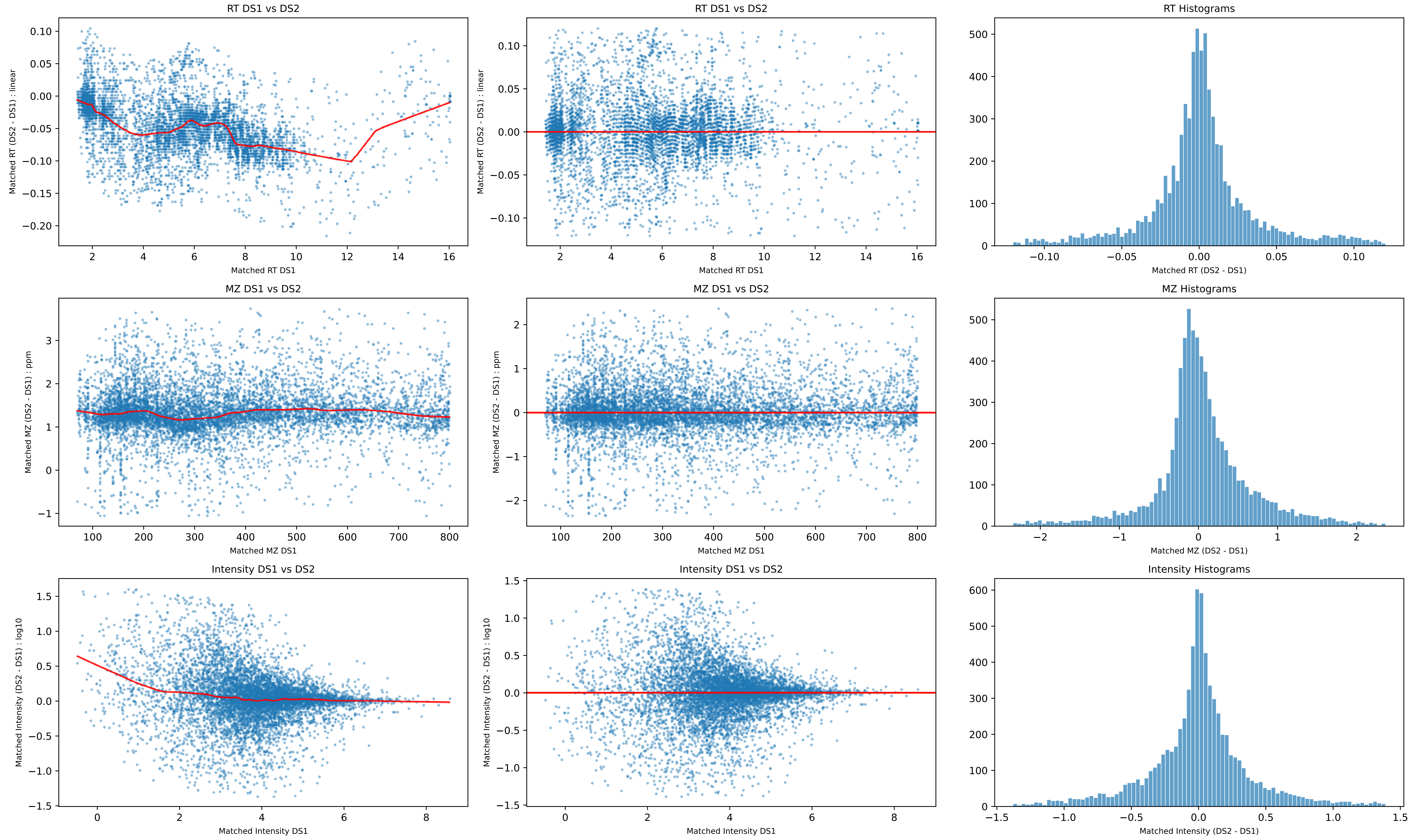
RT DS1 vs DS2
Matched RT (DS2 - DS1) : linear
Matched RT DS1
RT DS1 vs DS2
Matched RT (DS2 - DS1) : linear
Matched RT DS1
RT Histograms
Matched RT (DS2 - DS1)
MZ DS1 vs DS2
Matched MZ (DS2 - DS1) : ppm
Matched MZ DS1
MZ DS1 vs DS2
Matched MZ (DS2 - DS1) : ppm
Matched MZ DS1
MZ Histograms
Matched MZ (DS2 - DS1)
Intensity DS1 vs DS2
Matched Intensity (DS2 - DS1) : log10
Matched Intensity DS1
Intensity DS1 vs DS2
Matched Intensity (DS2 - DS1) : log10
Matched Intensity DS1
Intensity Histograms
Matched Intensity (DS2 - DS1)

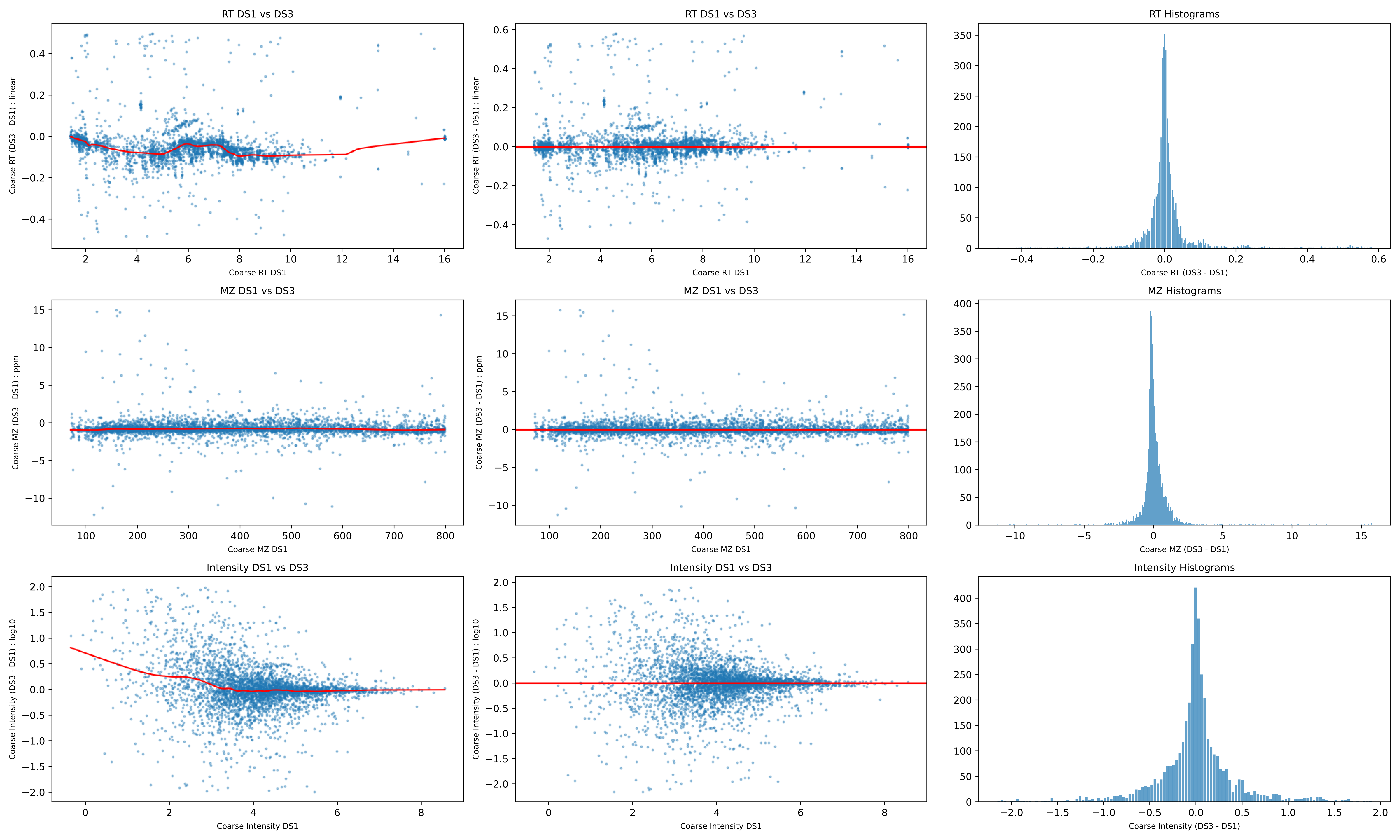
RT DS1 vs DS3
Coarse RT (DS3 - DS1) : linear
Coarse RT DS1
RT DS1 vs DS3
Coarse RT (DS3 - DS1) : linear
Coarse RT DS1
RT Histograms
Coarse RT (DS3 - DS1)
MZ DS1 vs DS3
Coarse MZ (DS3 - DS1) : ppm
Coarse MZ DS1
MZ DS1 vs DS3
Coarse MZ (DS3 - DS1) : ppm
Coarse MZ DS1
MZ Histograms
Coarse MZ (DS3 - DS1)
Intensity DS1 vs DS3
Coarse Intensity (DS3 - DS1) : log10
Coarse Intensity DS1
Intensity DS1 vs DS3
Coarse Intensity (DS3 - DS1) : log10
Coarse Intensity DS1
Intensity Histograms
Coarse Intensity (DS3 - DS1)

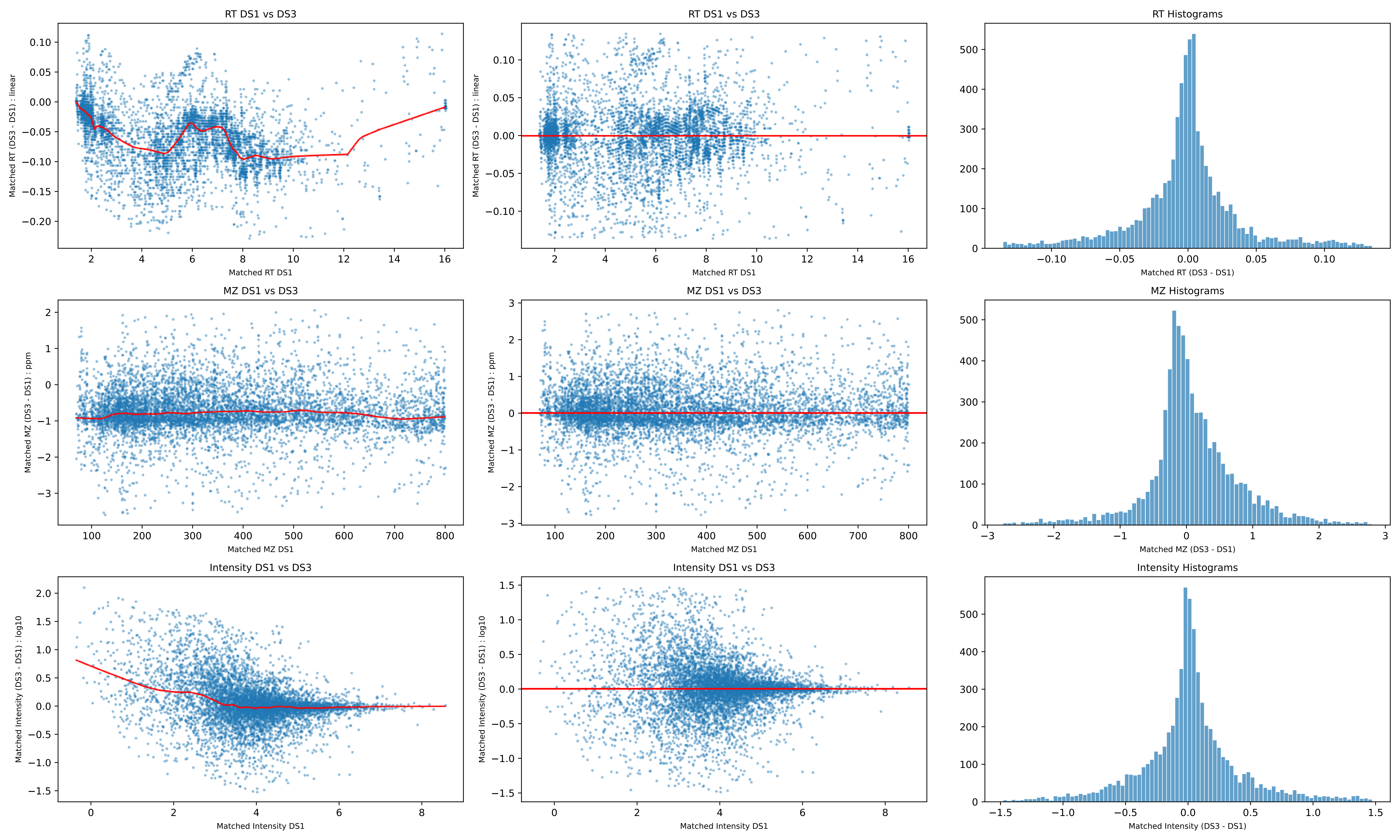
RT DS1 vs DS3
Matched RT (DS3 - DS1) : linear
Matched RT DS1
RT DS1 vs DS3
Matched RT (DS3 - DS1) : linear
Matched RT DS1
RT Histograms
Matched RT (DS3 - DS1)
MZ DS1 vs DS3
Matched MZ (DS3 - DS1) : ppm
Matched MZ DS1
MZ DS1 vs DS3
Matched MZ (DS3 - DS1) : ppm
Matched MZ DS1
MZ Histograms
Matched MZ (DS3 - DS1)
Intensity DS1 vs DS3
Matched Intensity (DS3 - DS1) : log10
Matched Intensity DS1
Intensity DS1 vs DS3
Matched Intensity (DS3 - DS1) : log10
Matched Intensity DS1
Intensity Histograms
Matched Intensity (DS3 - DS1)

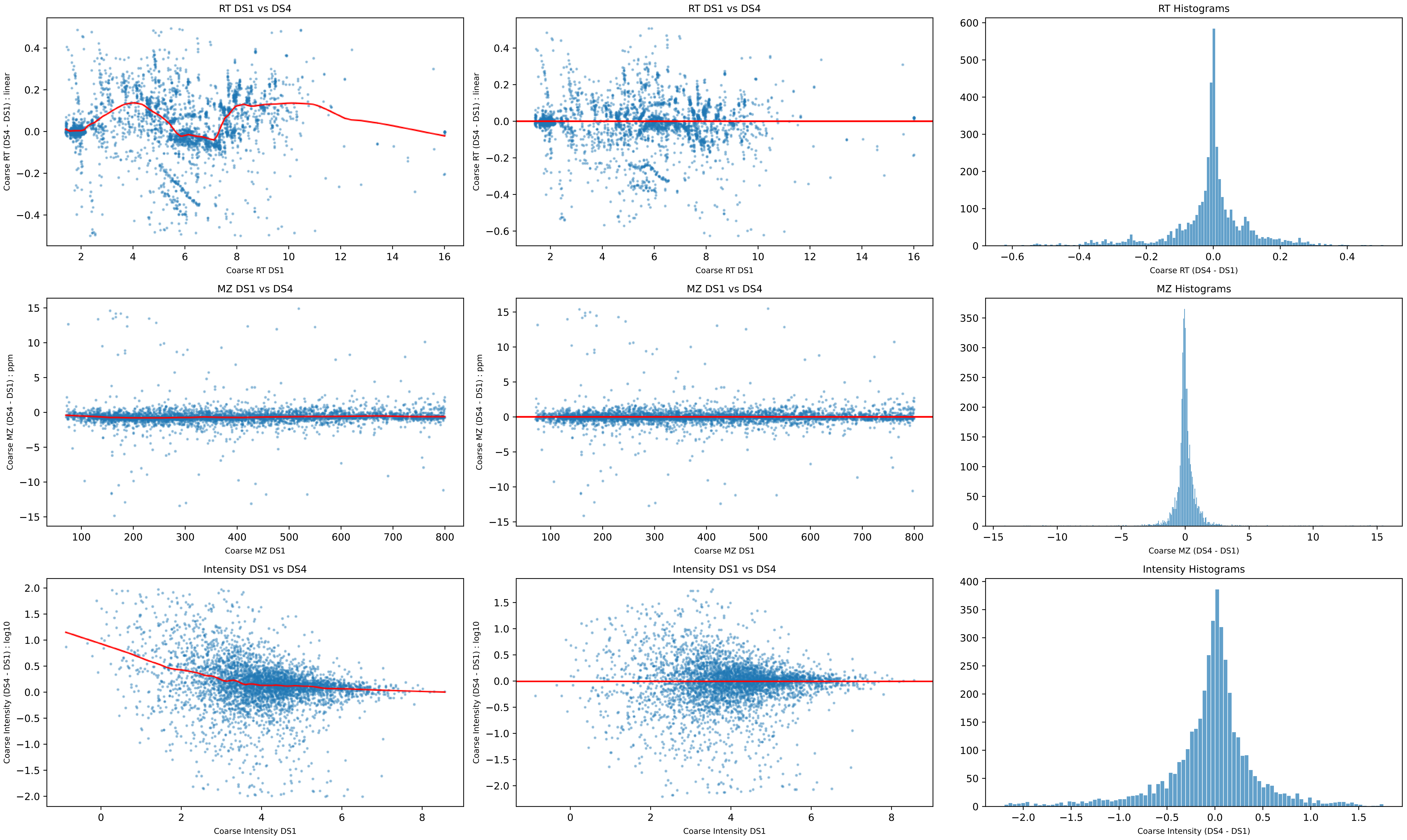
RT DS1 vs DS4
Coarse RT (DS4 - DS1) : linear
Coarse RT DS1
RT DS1 vs DS4
Coarse RT (DS4 - DS1) : linear
Coarse RT DS1
RT Histograms
Coarse RT (DS4 - DS1)
MZ DS1 vs DS4
Coarse MZ (DS4 - DS1) : ppm
Coarse MZ DS1
MZ DS1 vs DS4
Coarse MZ (DS4 - DS1) : ppm
Coarse MZ DS1
MZ Histograms
Coarse MZ (DS4 - DS1)
Intensity DS1 vs DS4
Coarse Intensity (DS4 - DS1) : log10
Coarse Intensity DS1
Intensity DS1 vs DS4
Coarse Intensity (DS4 - DS1) : log10
Coarse Intensity DS1
Intensity Histograms
Coarse Intensity (DS4 - DS1)

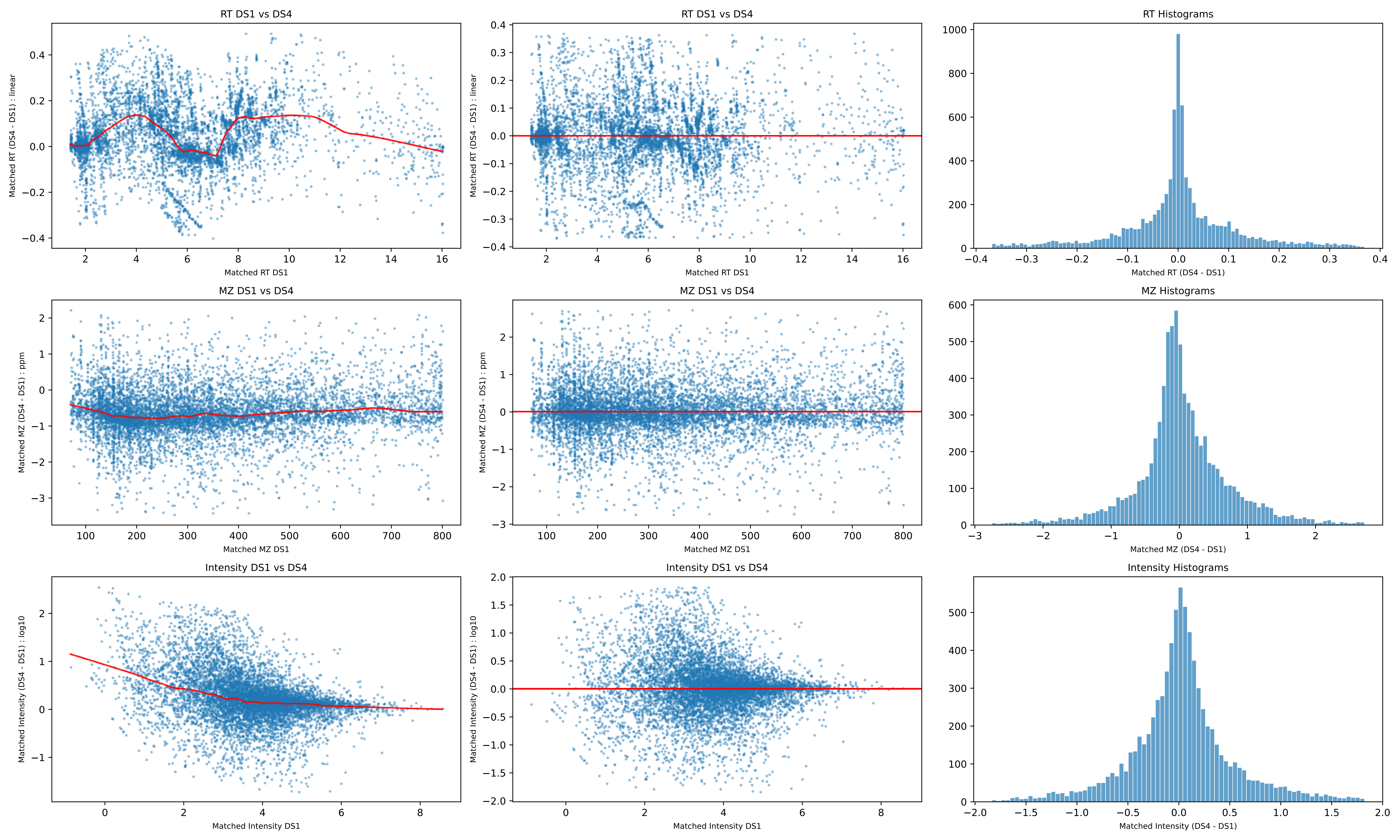
RT DS1 vs DS4
Matched RT (DS4 - DS1) : linear
Matched RT DS1
RT Histograms
Matched RT (DS4 - DS1)
MZ DS1 vs DS4
Matched MZ (DS4 - DS1) : ppm
Matched MZ DS1
MZ Histograms
Matched MZ (DS4 - DS1)
Intensity DS1 vs DS4
Matched Intensity (DS4 - DS1) : log10
Matched Intensity DS1
Intensity Histograms
Matched Intensity (DS4 - DS1)

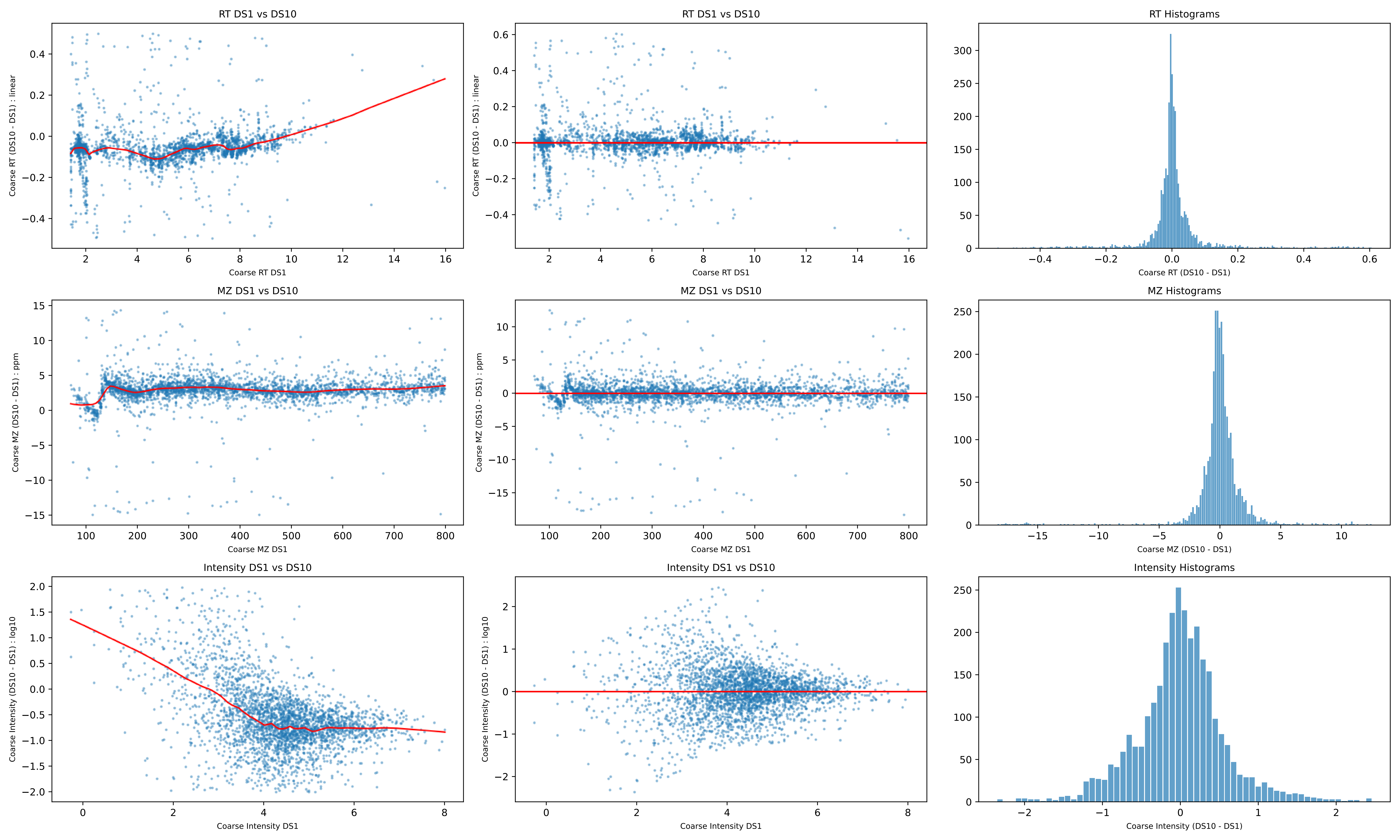
RT DS1 vs DS10
Coarse RT (DS10 - DS1) : linear
Coarse RT DS1
RT DS1 vs DS10
Coarse RT (DS10 - DS1) : linear
Coarse RT DS1
RT Histograms
Coarse RT (DS10 - DS1)
MZ DS1 vs DS10
Coarse MZ (DS10 - DS1) : ppm
Coarse MZ DS1
MZ DS1 vs DS10
Coarse MZ (DS10 - DS1) : ppm
Coarse MZ DS1
MZ Histograms
Coarse MZ (DS10 - DS1)
Intensity DS1 vs DS10
Coarse Intensity (DS10 - DS1) : log10
Coarse Intensity DS1
Intensity DS1 vs DS10
Coarse Intensity (DS10 - DS1) : log10
Coarse Intensity DS1
Intensity Histograms
Coarse Intensity (DS10 - DS1)

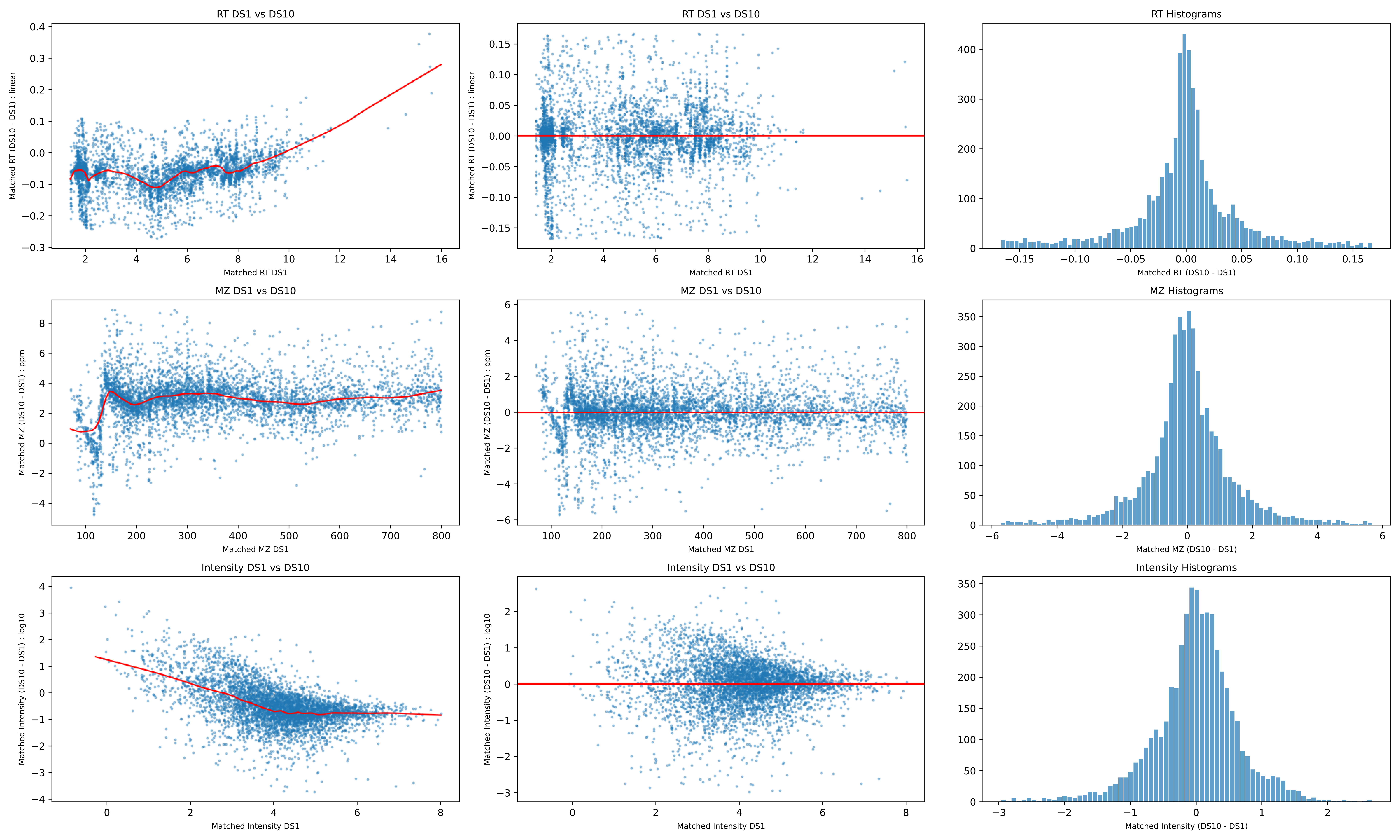
RT DS1 vs DS10
Matched RT (DS10 - DS1) : linear
Matched RT DS1
RT DS1 vs DS10
Matched RT (DS10 - DS1) : linear
Matched RT DS1
RT Histograms
Matched RT (DS10 - DS1)
MZ DS1 vs DS10
Matched MZ (DS10 - DS1) : ppm
Matched MZ DS1
MZ DS1 vs DS10
Matched MZ (DS10 - DS1) : ppm
Matched MZ DS1
MZ Histograms
Matched MZ (DS10 - DS1)
Intensity DS1 vs DS10
Matched Intensity (DS10 - DS1) : log10
Matched Intensity DS1
Intensity DS1 vs DS10
Matched Intensity (DS10 - DS1) : log10
Matched Intensity DS1
Intensity Histograms
Matched Intensity (DS10 - DS1)

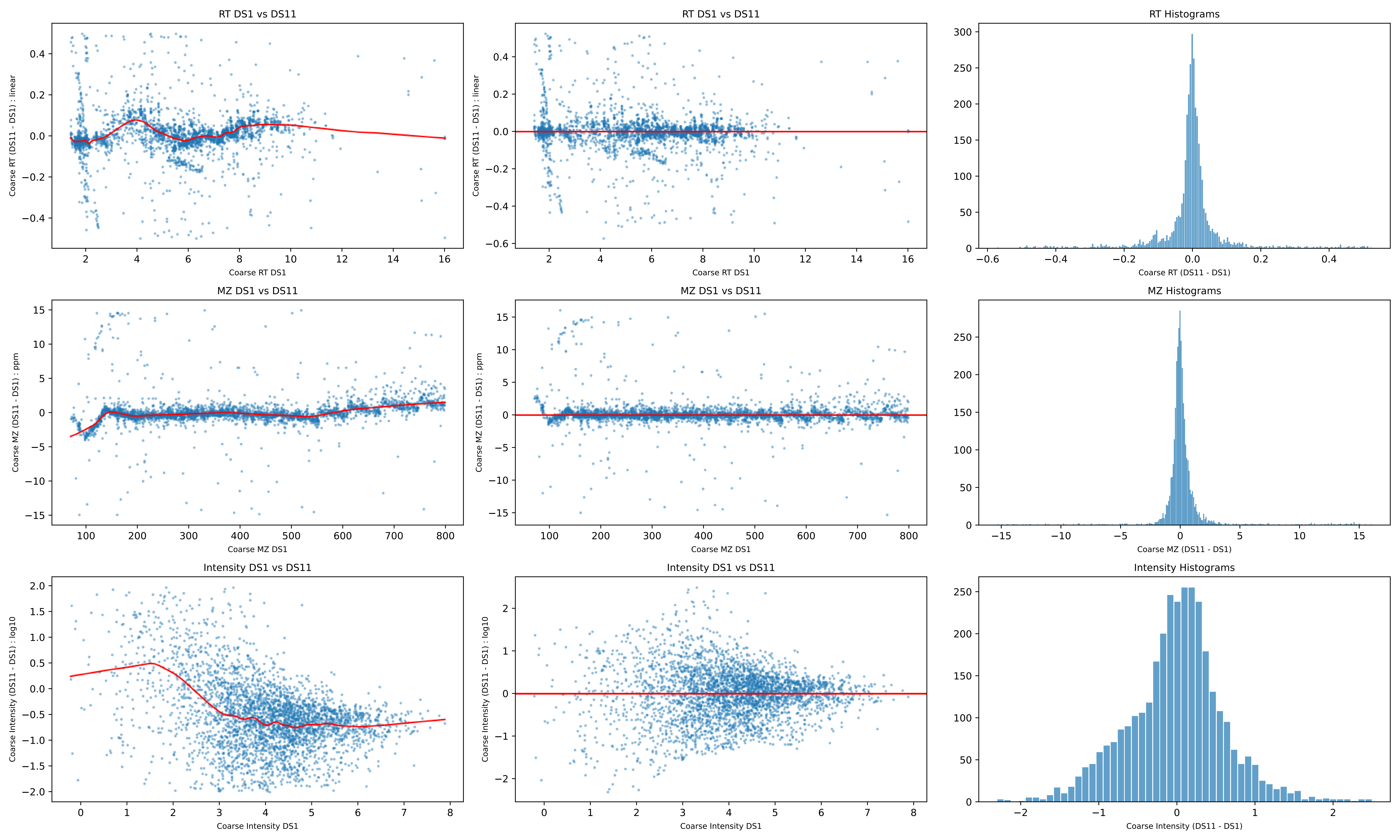

RT DS1 vs DS11
Coarse RT (DS11 - DS1) : linear
Coarse RT DS1
RT Histograms
Coarse RT (DS11 - DS1)
MZ DS1 vs DS11
Coarse MZ (DS11 - DS1) : ppm
Coarse MZ DS1
MZ Histograms
Coarse MZ (DS11 - DS1)
Intensity DS1 vs DS11
Coarse Intensity (DS11 - DS1) : log10
Coarse Intensity DS1
Intensity Histograms
Coarse Intensity (DS11 - DS1)

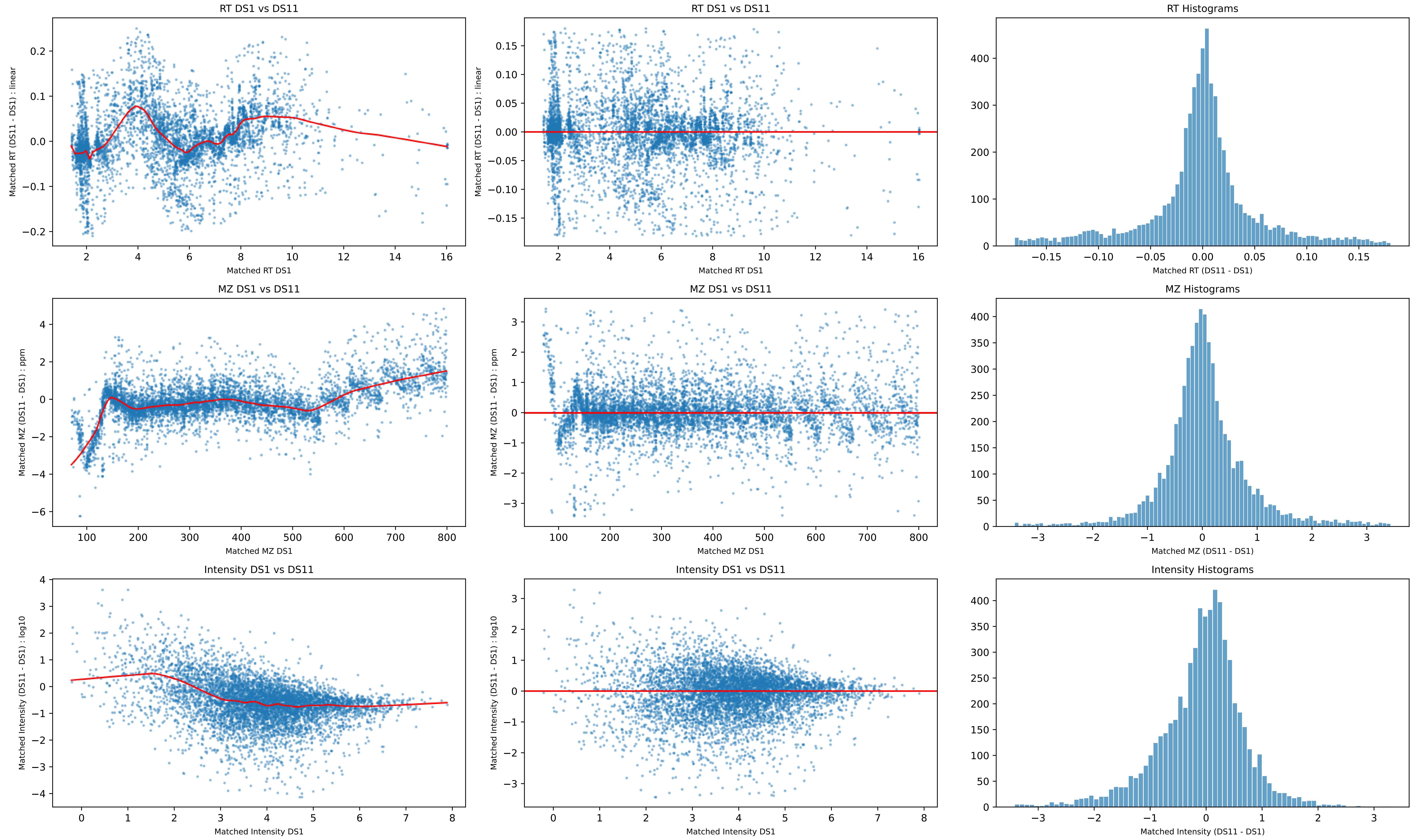
RT DS1 vs DS11
Matched RT (DS11 - DS1) : linear
Matched RT DS1
RT DS1 vs DS11
Matched RT (DS11 - DS1) : linear
Matched RT DS1
RT Histograms
Matched RT (DS11 - DS1)
MZ DS1 vs DS11
Matched MZ (DS11 - DS1) : ppm
Matched MZ DS1
MZ DS1 vs DS11
Matched MZ (DS11 - DS1) : ppm
Matched MZ DS1
MZ Histograms
Matched MZ (DS11 - DS1)
Intensity DS1 vs DS11
Matched Intensity (DS11 - DS1) : log10
Matched Intensity DS1
Intensity DS1 vs DS11
Matched Intensity (DS11 - DS1) : log10
Matched Intensity DS1
Intensity Histograms
Matched Intensity (DS11 - DS1)

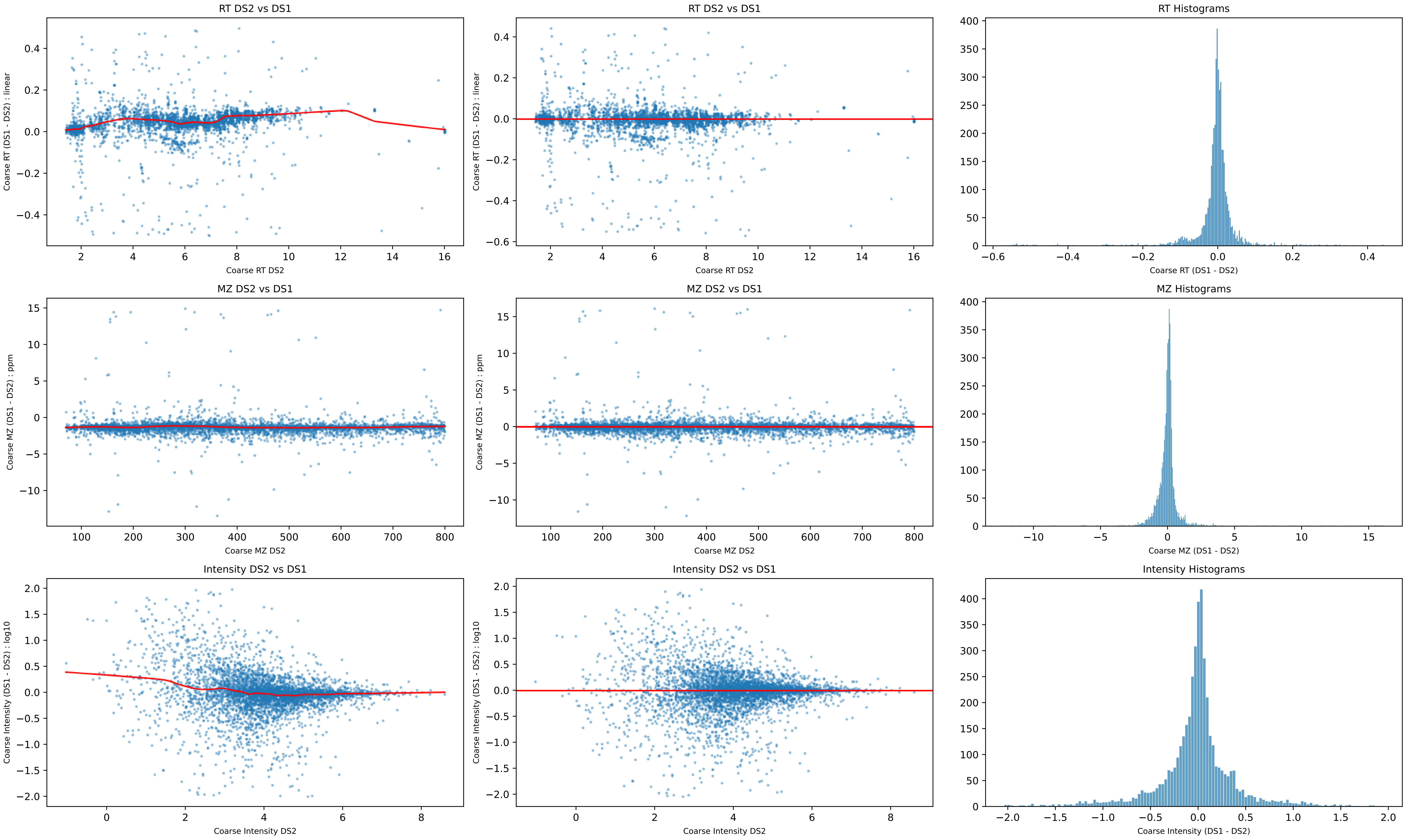
RT DS2 vs DS1
Coarse RT (DS1 - DS2) : linear
Coarse RT DS2
RT DS2 vs DS1
Coarse RT (DS1 - DS2) : linear
Coarse RT DS2
RT Histograms
Coarse RT (DS1 - DS2)
MZ DS2 vs DS1
Coarse MZ (DS1 - DS2) : ppm
Coarse MZ DS2
MZ DS2 vs DS1
Coarse MZ (DS1 - DS2) : ppm
Coarse MZ DS2
MZ Histograms
Coarse MZ (DS1 - DS2)
Intensity DS2 vs DS1
Coarse Intensity (DS1 - DS2) : log10
Coarse Intensity DS2
Intensity DS2 vs DS1
Coarse Intensity (DS1 - DS2) : log10
Coarse Intensity DS2
Intensity Histograms
Coarse Intensity (DS1 - DS2)

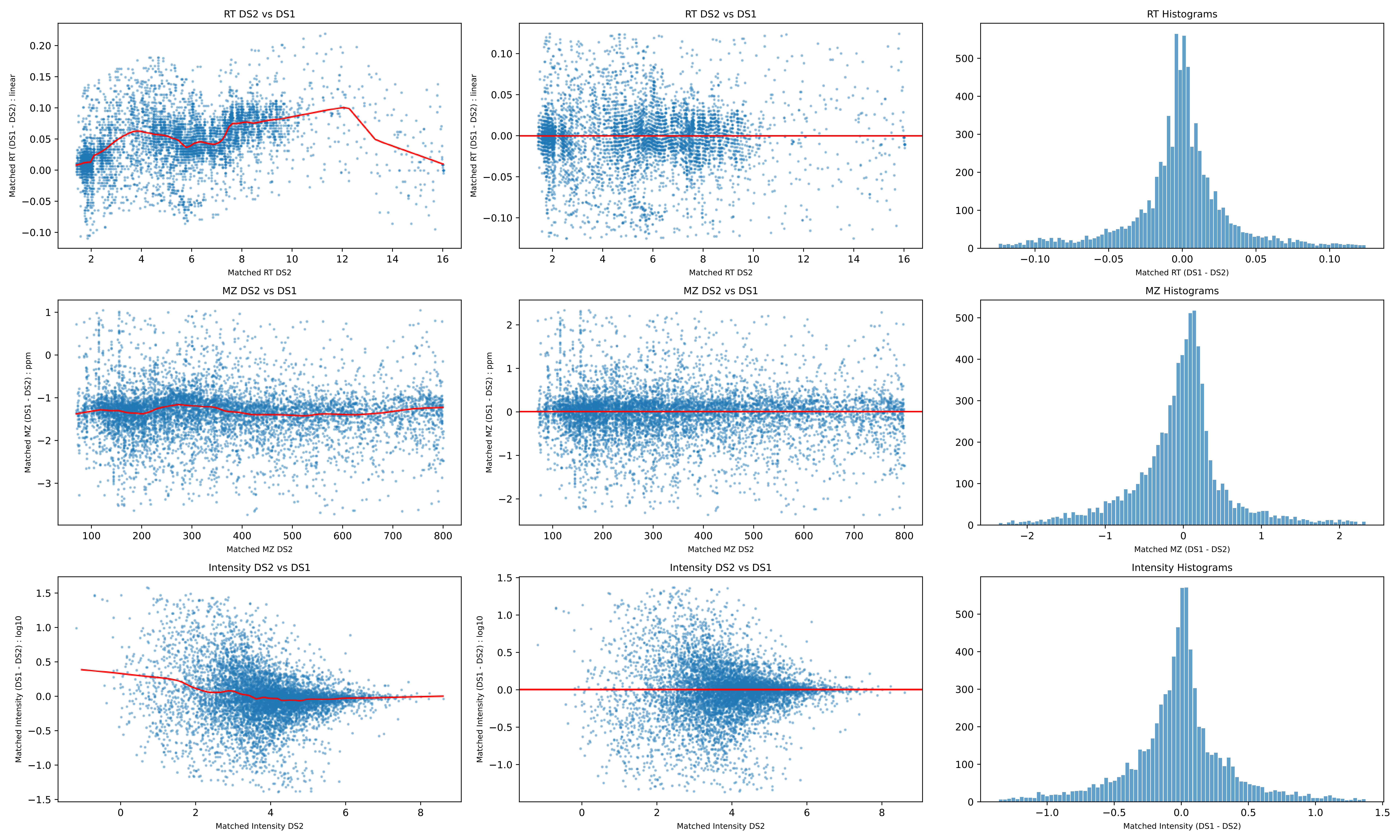
RT DS2 vs DS1
Matched RT (DS1 - DS2) : linear
Matched RT DS2
RT DS2 vs DS1
Matched RT (DS1 - DS2) : linear
Matched RT DS2
RT Histograms
Matched RT (DS1 - DS2)
MZ DS2 vs DS1
Matched MZ (DS1 - DS2) : ppm
Matched MZ DS2
MZ DS2 vs DS1
Matched MZ (DS1 - DS2) : ppm
Matched MZ DS2
MZ Histograms
Matched MZ (DS1 - DS2)
Intensity DS2 vs DS1
Matched Intensity (DS1 - DS2) : log10
Matched Intensity DS2
Intensity DS2 vs DS1
Matched Intensity (DS1 - DS2) : log10
Matched Intensity DS2
Intensity Histograms
Matched Intensity (DS1 - DS2)

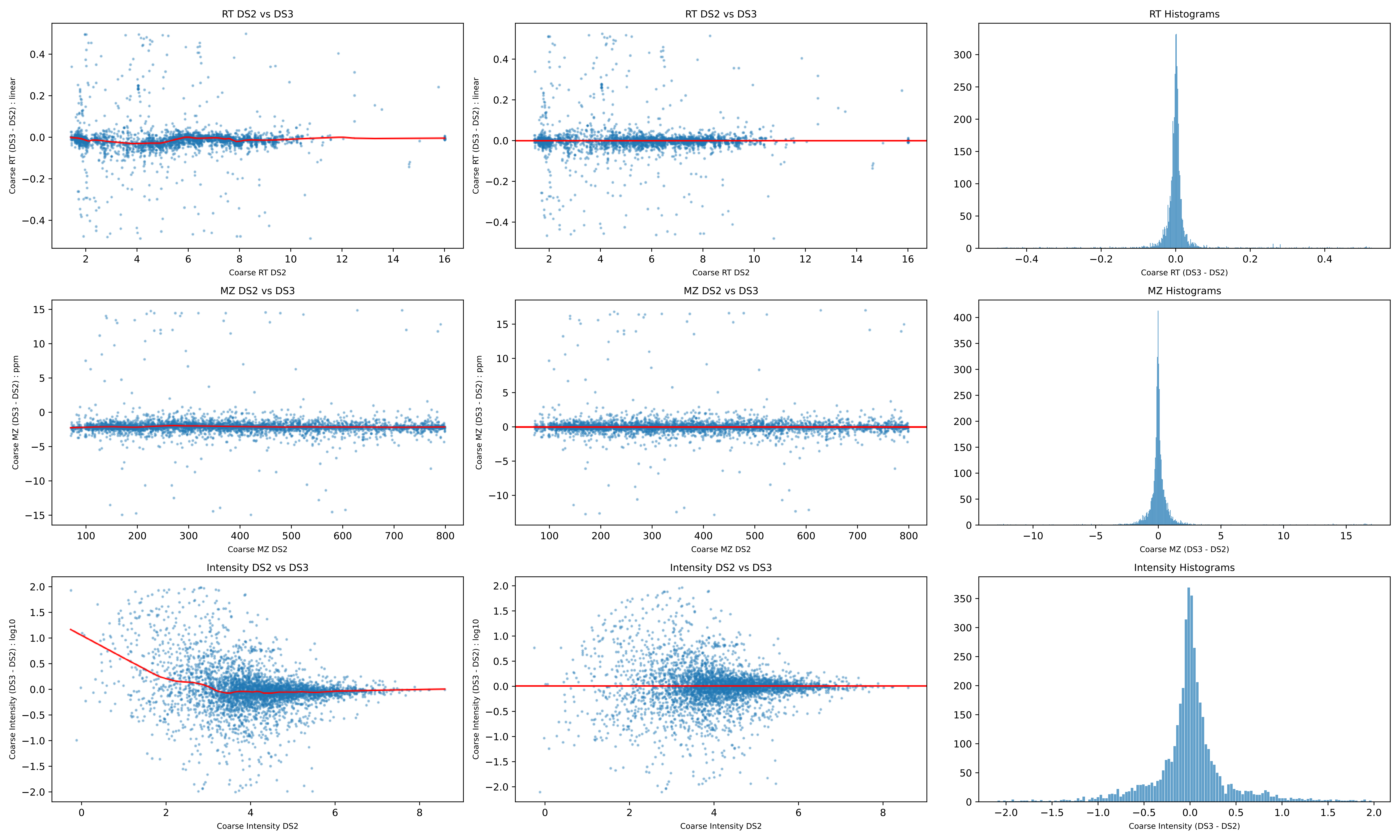

RT DS2 vs DS3
Coarse RT (DS3 - DS2) : linear
Coarse RT DS2
RT DS2 vs DS3
Coarse RT (DS3 - DS2) : linear
Coarse RT DS2
RT Histograms
Coarse RT (DS3 - DS2)
MZ DS2 vs DS3
Coarse MZ (DS3 - DS2) : ppm
Coarse MZ DS2
MZ DS2 vs DS3
Coarse MZ (DS3 - DS2) : ppm
Coarse MZ DS2
MZ Histograms
Coarse MZ (DS3 - DS2)
Intensity DS2 vs DS3
Coarse Intensity (DS3 - DS2) : log10
Coarse Intensity DS2
Intensity DS2 vs DS3
Coarse Intensity (DS3 - DS2) : log10
Coarse Intensity DS2
Intensity Histograms
Coarse Intensity (DS3 - DS2)

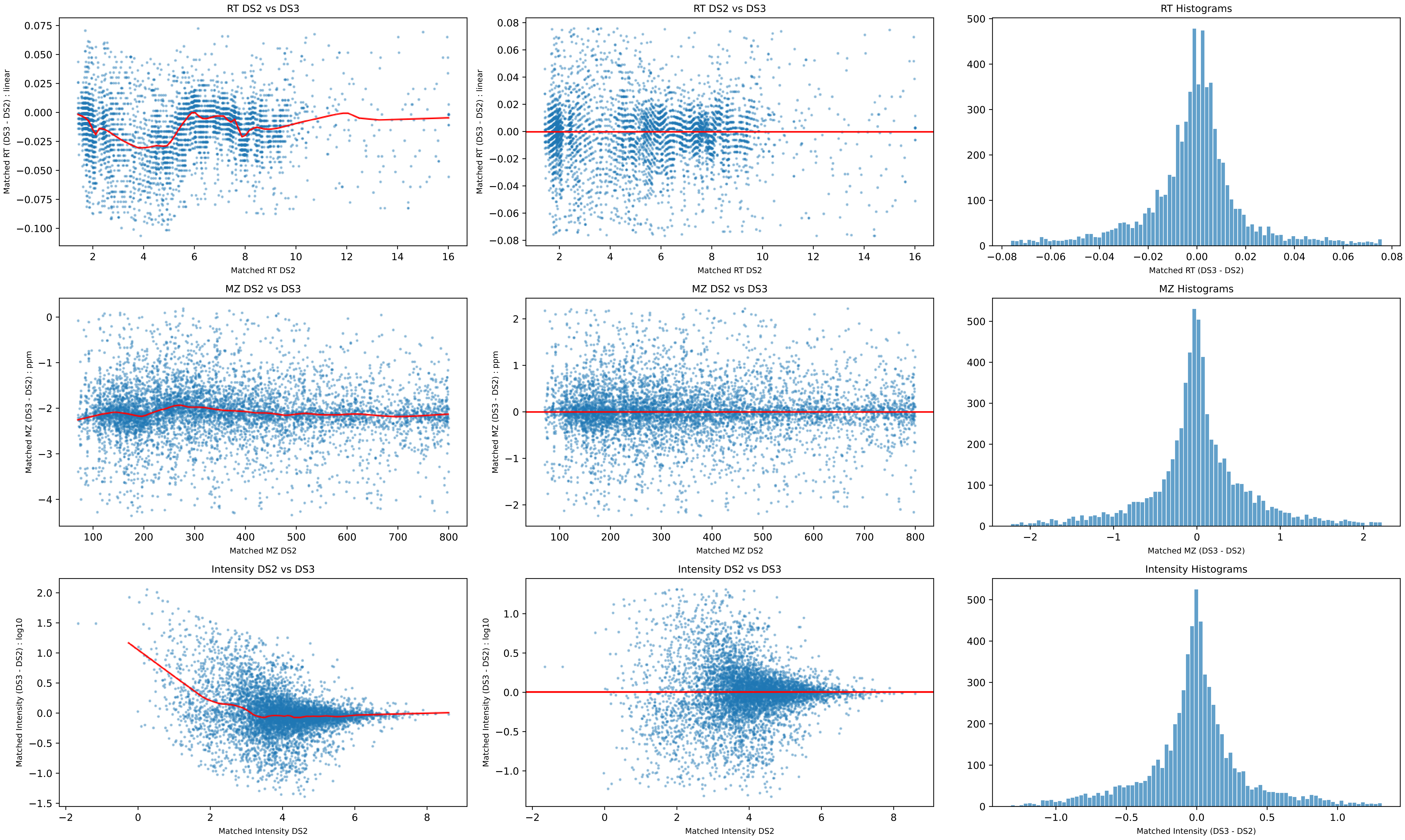
RT DS2 vs DS3
Matched RT (DS3 - DS2) : linear
Matched RT DS2
RT DS2 vs DS3
Matched RT (DS3 - DS2) : linear
Matched RT DS2
RT Histograms
Matched RT (DS3 - DS2)
MZ DS2 vs DS3
Matched MZ (DS3 - DS2) : ppm
Matched MZ DS2
MZ DS2 vs DS3
Matched MZ (DS3 - DS2) : ppm
Matched MZ DS2
MZ Histograms
Matched MZ (DS3 - DS2)
Intensity DS2 vs DS3
Matched Intensity (DS3 - DS2) : log10
Matched Intensity DS2
Intensity DS2 vs DS3
Matched Intensity (DS3 - DS2) : log10
Matched Intensity DS2
Intensity Histograms
Matched Intensity (DS3 - DS2)

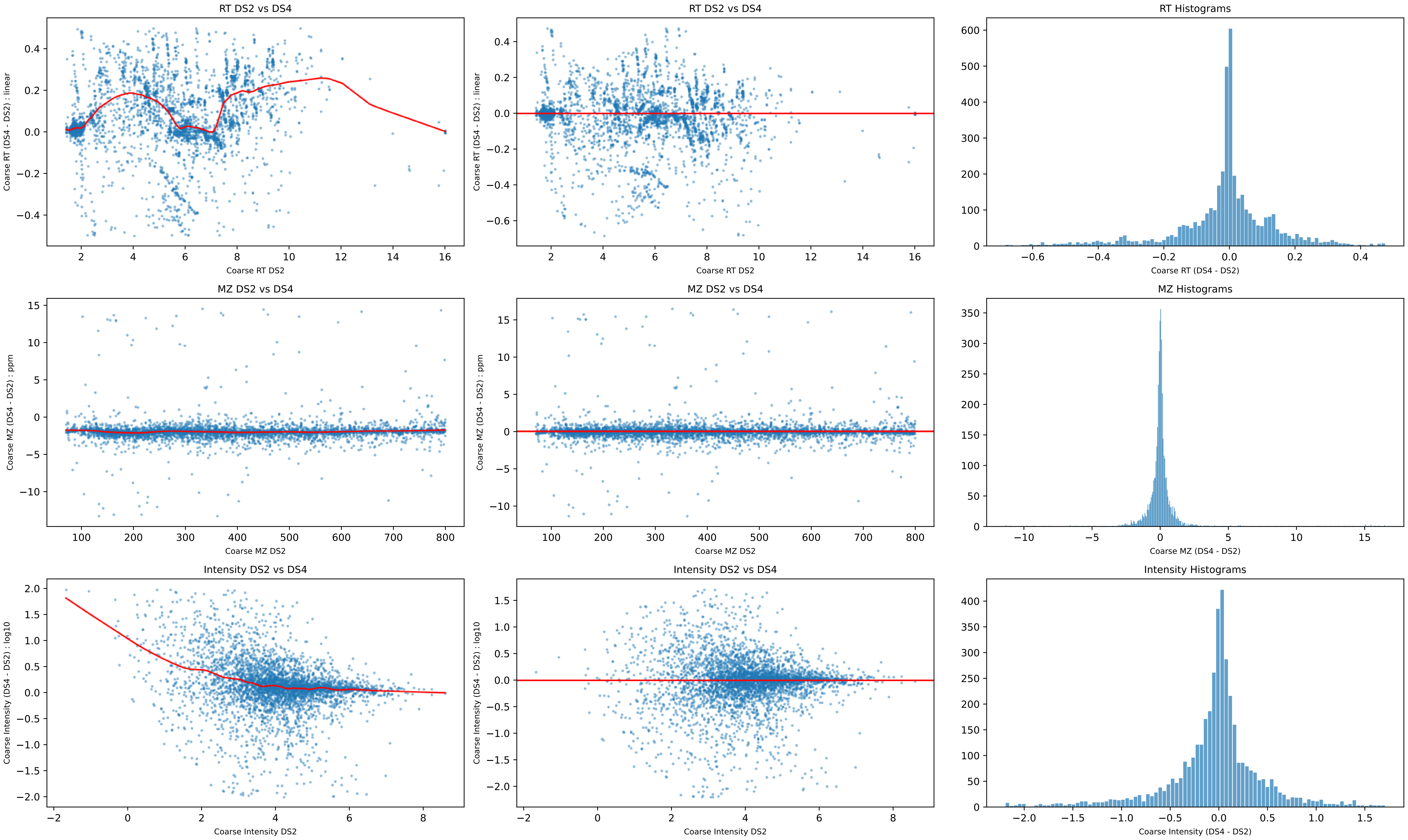
RT DS2 vs DS4
Coarse RT (DS4 - DS2) : linear
Coarse RT DS2
RT DS2 vs DS4
Coarse RT (DS4 - DS2) : linear
Coarse RT DS2
RT Histograms
Coarse RT (DS4 - DS2)
MZ DS2 vs DS4
Coarse MZ (DS4 - DS2) : ppm
Coarse MZ DS2
MZ DS2 vs DS4
Coarse MZ (DS4 - DS2) : ppm
Coarse MZ DS2
MZ Histograms
Coarse MZ (DS4 - DS2)
Intensity DS2 vs DS4
Coarse Intensity (DS4 - DS2) : log10
Coarse Intensity DS2
Intensity DS2 vs DS4
Coarse Intensity (DS4 - DS2) : log10
Coarse Intensity DS2
Intensity Histograms
Coarse Intensity (DS4 - DS2)

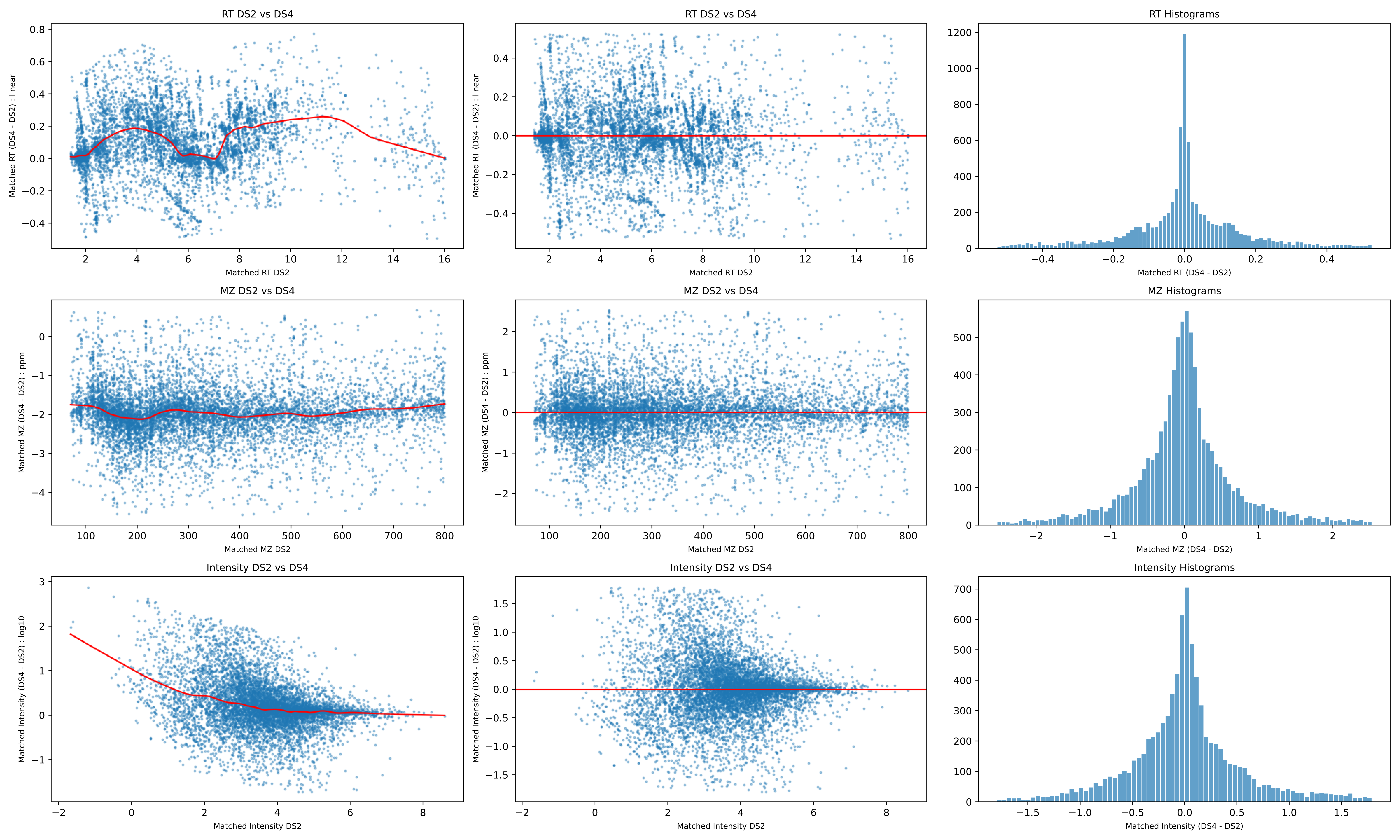
RT DS2 vs DS4
Matched RT (DS4 - DS2) : linear
Matched RT DS2
RT DS2 vs DS4
Matched RT (DS4 - DS2) : linear
Matched RT DS2
RT Histograms
Matched RT (DS4 - DS2)
MZ DS2 vs DS4
Matched MZ (DS4 - DS2) : ppm
Matched MZ DS2
MZ DS2 vs DS4
Matched MZ (DS4 - DS2) : ppm
Matched MZ DS2
MZ Histograms
Matched MZ (DS4 - DS2)
Intensity DS2 vs DS4
Matched Intensity (DS4 - DS2) : log10
Matched Intensity DS2
Intensity DS2 vs DS4
Matched Intensity (DS4 - DS2) : log10
Matched Intensity DS2
Intensity Histograms
Matched Intensity (DS4 - DS2)

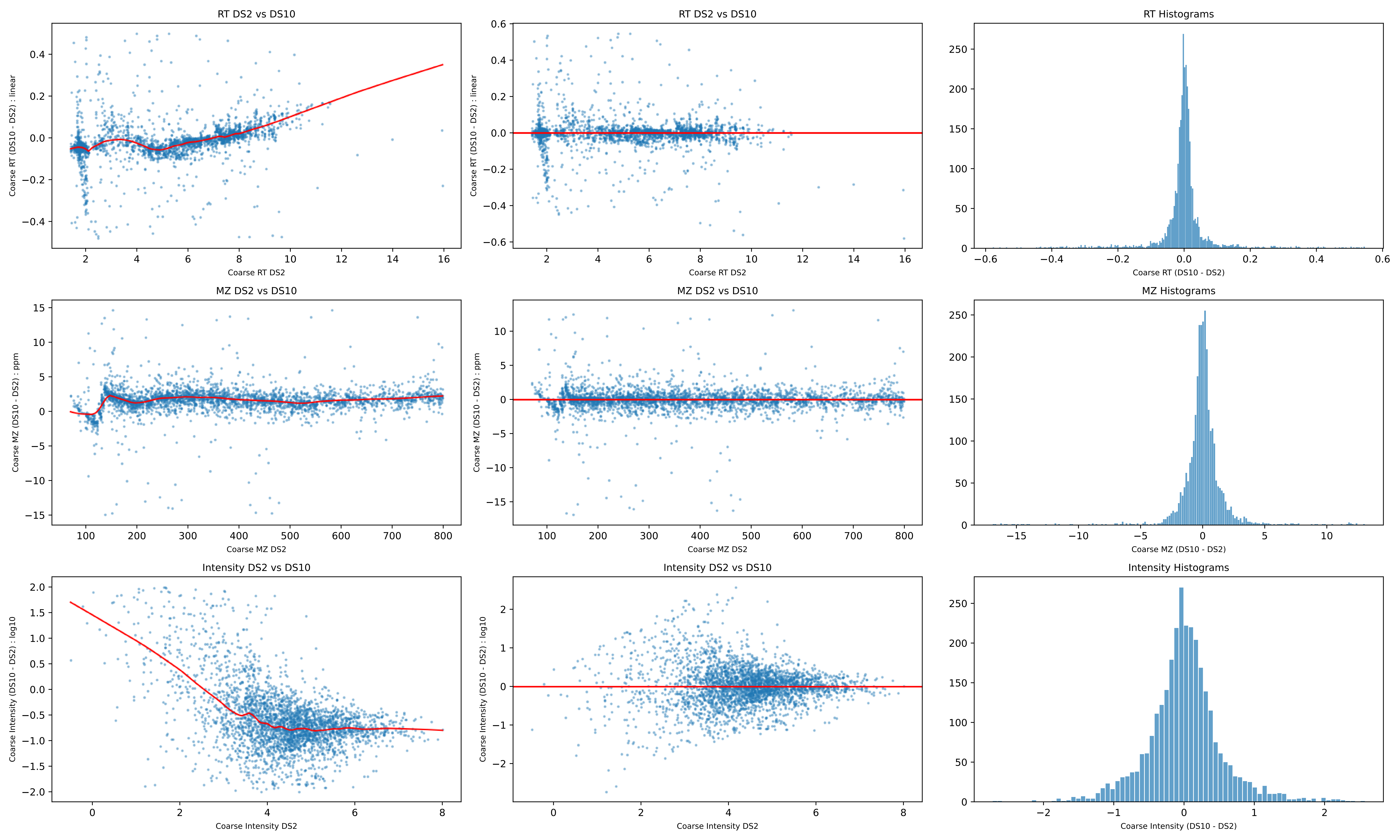
RT DS2 vs DS10
Coarse RT (DS10 - DS2) : linear
Coarse RT DS2
RT DS2 vs DS10
Coarse RT (DS10 - DS2) : linear
Coarse RT DS2
RT Histograms
Coarse RT (DS10 - DS2)
MZ DS2 vs DS10
Coarse MZ (DS10 - DS2) : ppm
Coarse MZ DS2
MZ DS2 vs DS10
Coarse MZ (DS10 - DS2) : ppm
Coarse MZ DS2
MZ Histograms
Coarse MZ (DS10 - DS2)
Intensity DS2 vs DS10
Coarse Intensity (DS10 - DS2) : log10
Coarse Intensity DS2
Intensity DS2 vs DS10
Coarse Intensity (DS10 - DS2) : log10
Coarse Intensity DS2
Intensity Histograms
Coarse Intensity (DS10 - DS2)

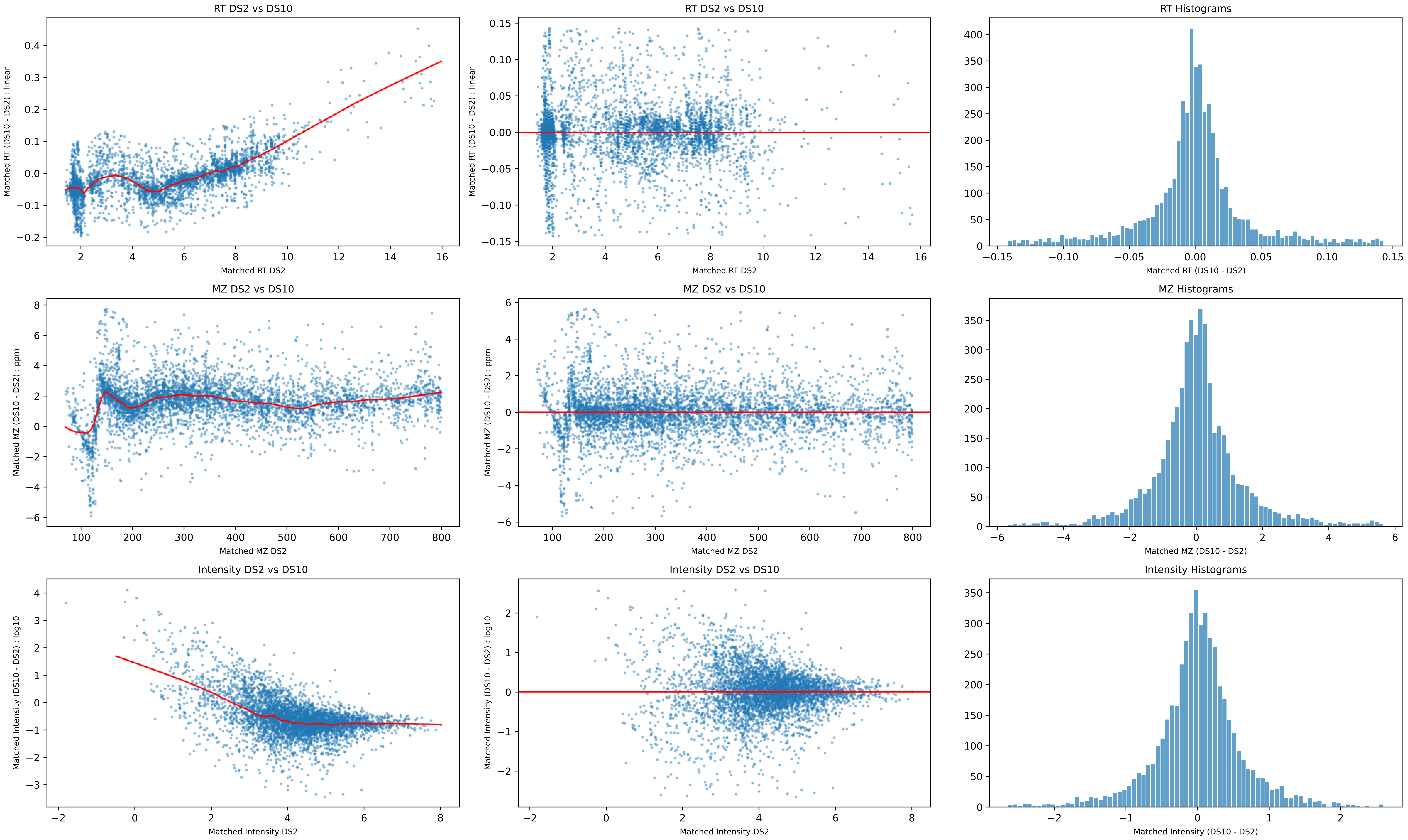
RT DS2 vs DS10
Matched RT (DS10 - DS2) : linear
Matched RT DS2
RT Histograms
Matched RT (DS10 - DS2)
MZ DS2 vs DS10
Matched MZ (DS10 - DS2) : ppm
Matched MZ DS2
MZ Histograms
Matched MZ (DS10 - DS2)
Intensity DS2 vs DS10
Matched Intensity (DS10 - DS2) : log10
Matched Intensity DS2
Intensity Histograms
Matched Intensity (DS10 - DS2)

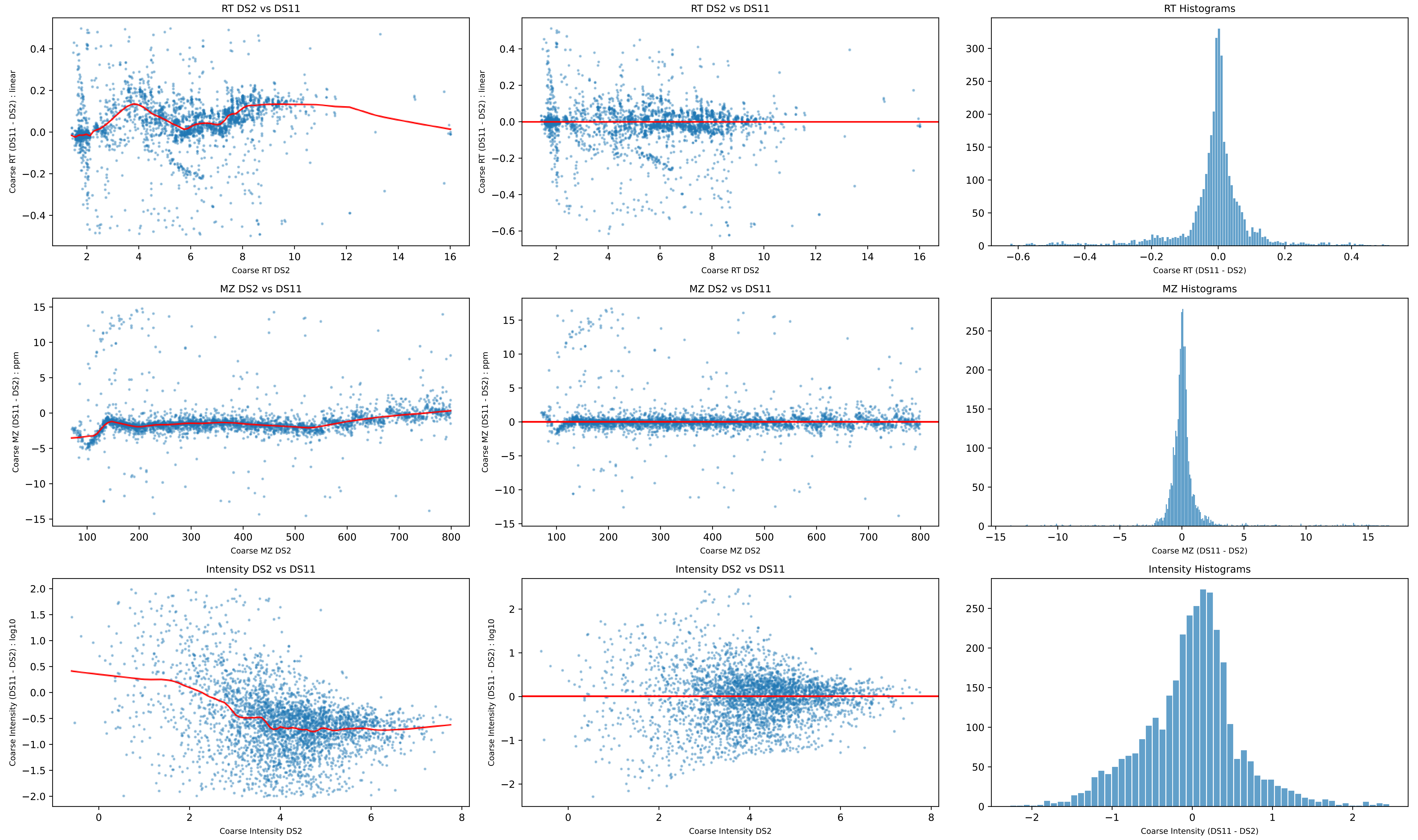
RT DS2 vs DS11
Coarse RT (DS11 - DS2) : linear
Coarse RT DS2
RT DS2 vs DS11
Coarse RT (DS11 - DS2) : linear
Coarse RT DS2
RT Histograms
Coarse RT (DS11 - DS2)
MZ DS2 vs DS11
Coarse MZ (DS11 - DS2) : ppm
Coarse MZ DS2
MZ DS2 vs DS11
Coarse MZ (DS11 - DS2) : ppm
Coarse MZ DS2
MZ Histograms
Coarse MZ (DS11 - DS2)
Intensity DS2 vs DS11
Coarse Intensity (DS11 - DS2) : log10
Coarse Intensity DS2
Intensity DS2 vs DS11
Coarse Intensity (DS11 - DS2) : log10
Coarse Intensity DS2
Intensity Histograms
Coarse Intensity (DS11 - DS2)

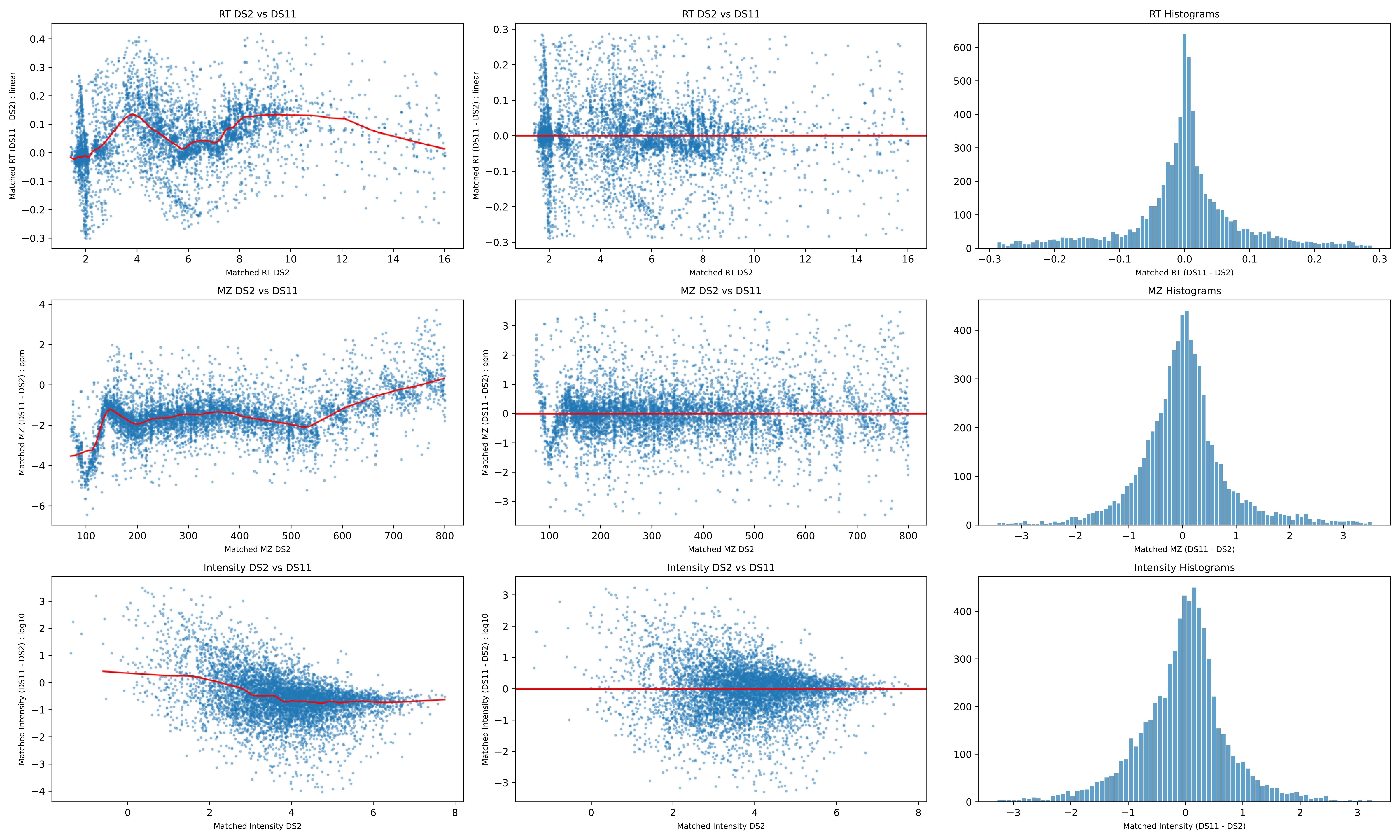
RT DS2 vs DS11
Matched RT (DS11 - DS2) : linear
Matched RT DS2
RT DS2 vs DS11
Matched RT (DS11 - DS2) : linear
Matched RT DS2
RT Histograms
Matched RT (DS11 - DS2)
MZ DS2 vs DS11
Matched MZ (DS11 - DS2) : ppm
Matched MZ DS2
MZ DS2 vs DS11
Matched MZ (DS11 - DS2) : ppm
Matched MZ DS2
MZ Histograms
Matched MZ (DS11 - DS2)
Intensity DS2 vs DS11
Matched Intensity (DS11 - DS2) : log10
Matched Intensity DS2
Intensity DS2 vs DS11
Matched Intensity (DS11 - DS2) : log10
Matched Intensity DS2
Intensity Histograms
Matched Intensity (DS11 - DS2)

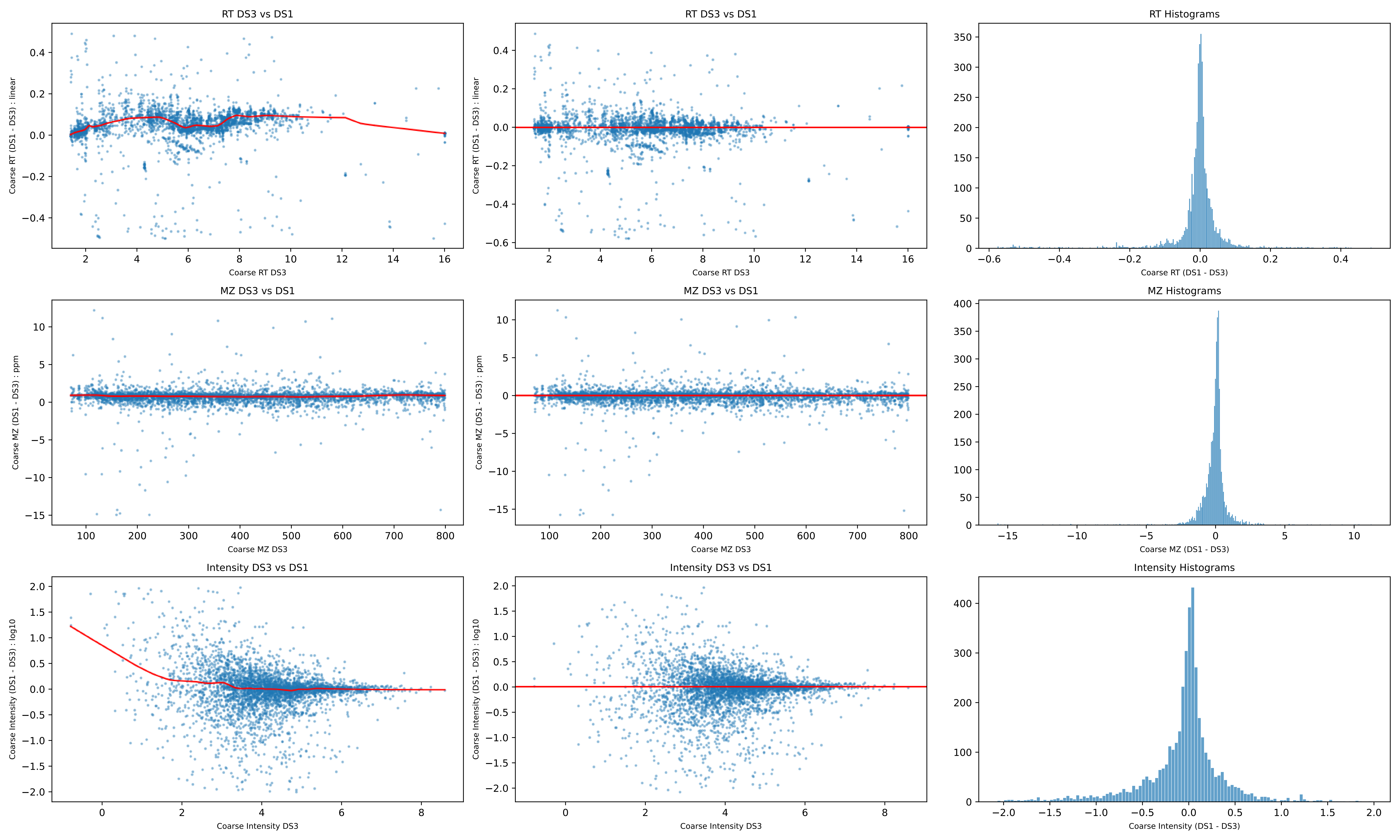

RT DS3 vs DS1
Coarse RT (DS1 - DS3) : linear
Coarse RT DS3
RT DS3 vs DS1
Coarse RT (DS1 - DS3) : linear
Coarse RT DS3
RT Histograms
Coarse RT (DS1 - DS3)
MZ DS3 vs DS1
Coarse MZ (DS1 - DS3) : ppm
Coarse MZ DS3
MZ DS3 vs DS1
Coarse MZ (DS1 - DS3) : ppm
Coarse MZ DS3
MZ Histograms
Coarse MZ (DS1 - DS3)
Intensity DS3 vs DS1
Coarse Intensity (DS1 - DS3) : log10
Coarse Intensity DS3
Intensity DS3 vs DS1
Coarse Intensity (DS1 - DS3) : log10
Coarse Intensity DS3
Intensity Histograms
Coarse Intensity (DS1 - DS3)

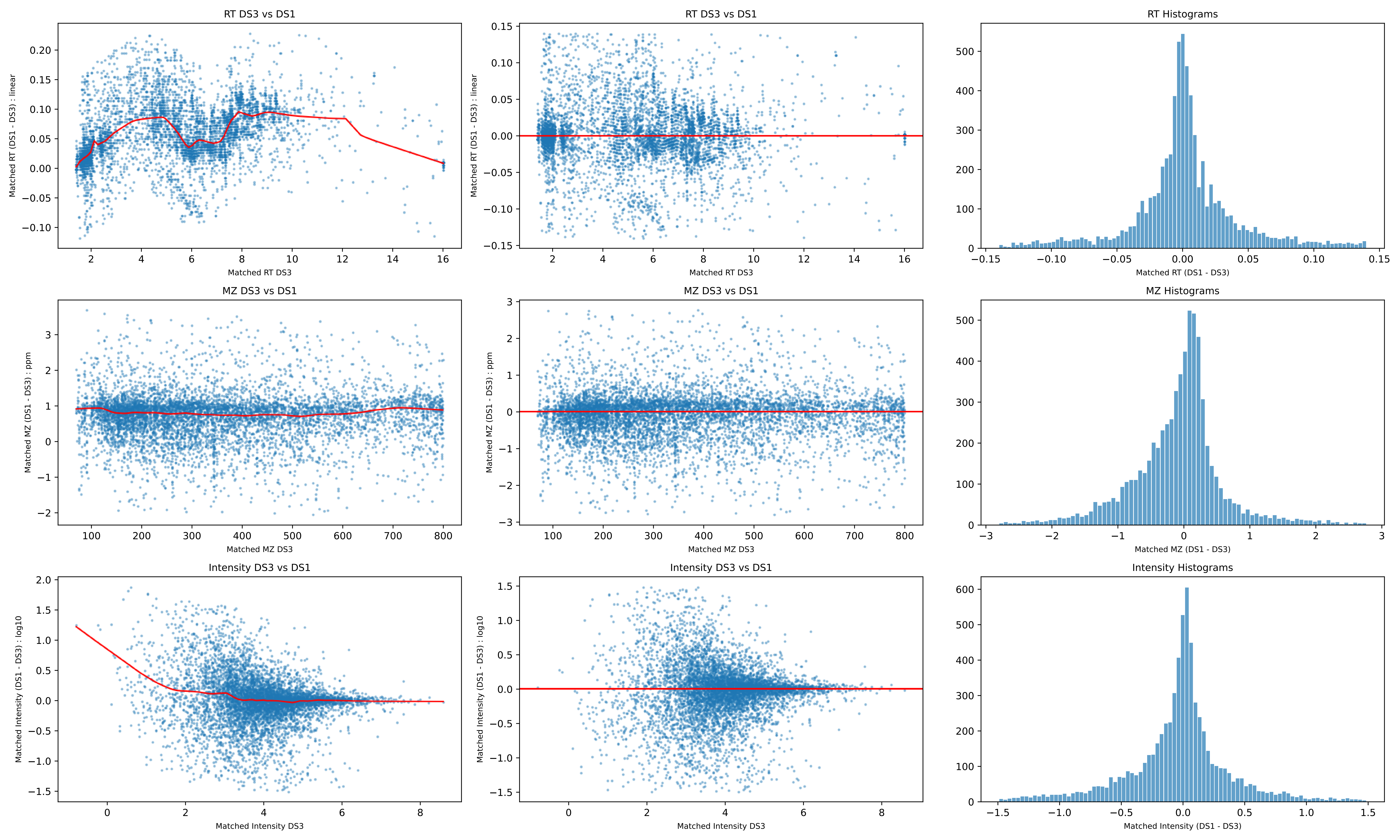
RT DS3 vs DS1
Matched RT (DS1 - DS3) : linear
Matched RT DS3
RT DS3 vs DS1
Matched RT (DS1 - DS3) : linear
Matched RT DS3
RT Histograms
Matched RT (DS1 - DS3)
MZ DS3 vs DS1
Matched MZ (DS1 - DS3) : ppm
Matched MZ DS3
MZ DS3 vs DS1
Matched MZ (DS1 - DS3) : ppm
Matched MZ DS3
MZ Histograms
Matched MZ (DS1 - DS3)
Intensity DS3 vs DS1
Matched Intensity (DS1 - DS3) : log10
Matched Intensity DS3
Intensity DS3 vs DS1
Matched Intensity (DS1 - DS3) : log10
Matched Intensity DS3
Intensity Histograms
Matched Intensity (DS1 - DS3)

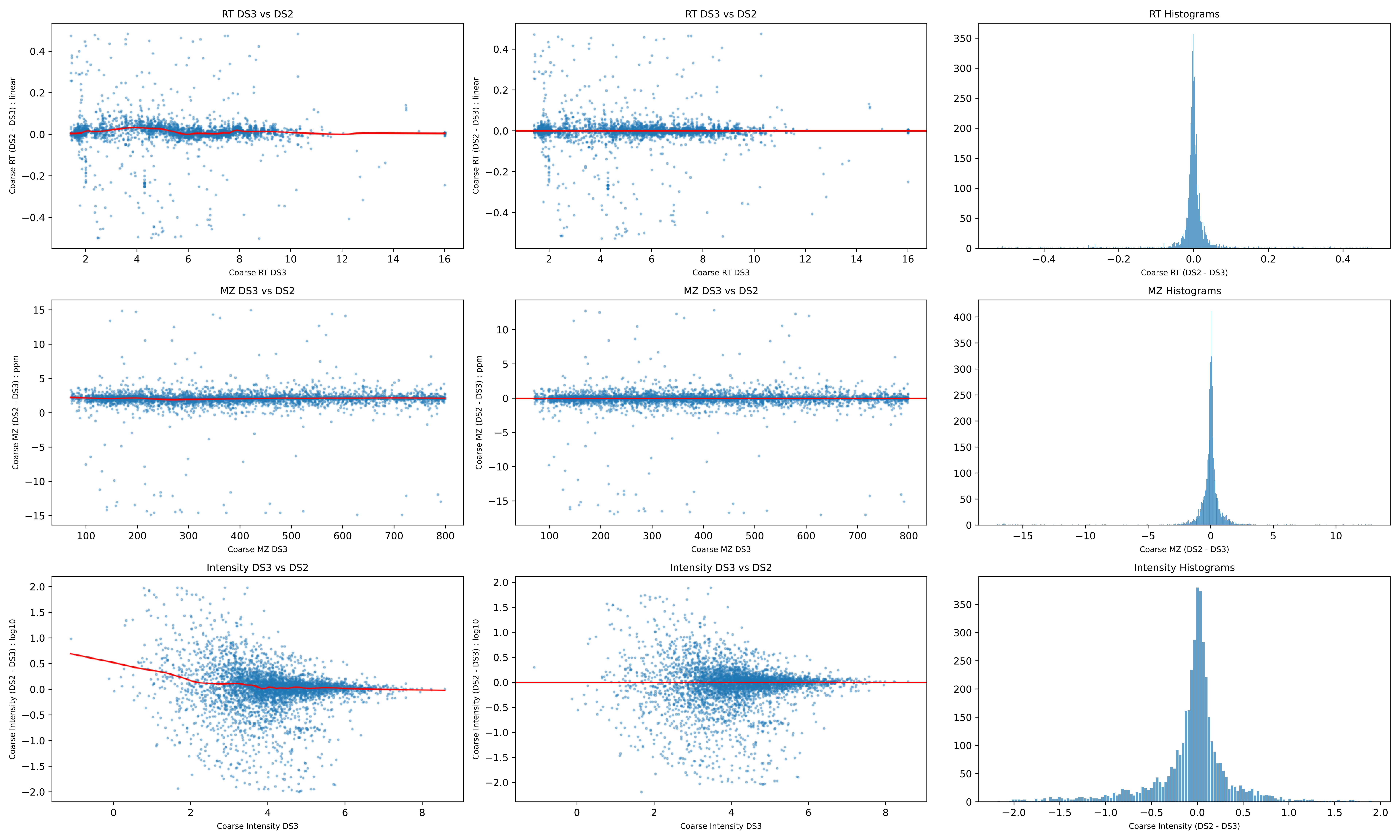
RT DS3 vs DS2
Coarse RT (DS2 - DS3) : linear
Coarse RT DS3
RT DS3 vs DS2
Coarse RT (DS2 - DS3) : linear
Coarse RT DS3
RT Histograms
Coarse RT (DS2 - DS3)
MZ DS3 vs DS2
Coarse MZ (DS2 - DS3) : ppm
Coarse MZ DS3
MZ DS3 vs DS2
Coarse MZ (DS2 - DS3) : ppm
Coarse MZ DS3
MZ Histograms
Coarse MZ (DS2 - DS3)
Intensity DS3 vs DS2
Coarse Intensity (DS2 - DS3) : log10
Coarse Intensity DS3
Intensity DS3 vs DS2
Coarse Intensity (DS2 - DS3) : log10
Coarse Intensity DS3
Intensity Histograms
Coarse Intensity (DS2 - DS3)

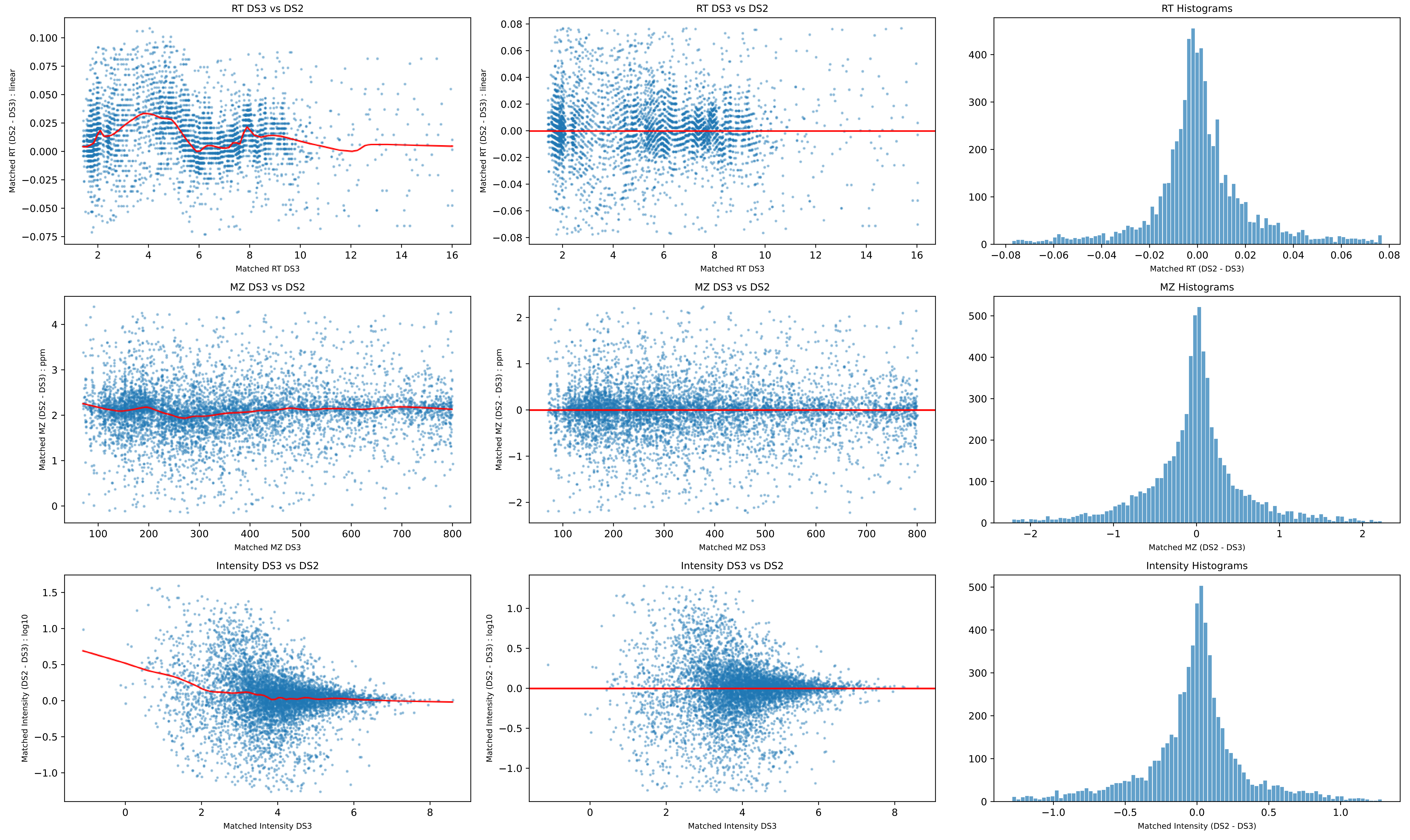
RT DS3 vs DS2
Matched RT (DS2 - DS3) : linear
Matched RT DS3
RT Histograms
Matched RT (DS2 - DS3)
MZ DS3 vs DS2
Matched MZ (DS2 - DS3) : ppm
Matched MZ DS3
MZ Histograms
Matched MZ (DS2 - DS3)
Intensity DS3 vs DS2
Matched Intensity (DS2 - DS3) : log10
Matched Intensity DS3
Intensity Histograms
Matched Intensity (DS2 - DS3)

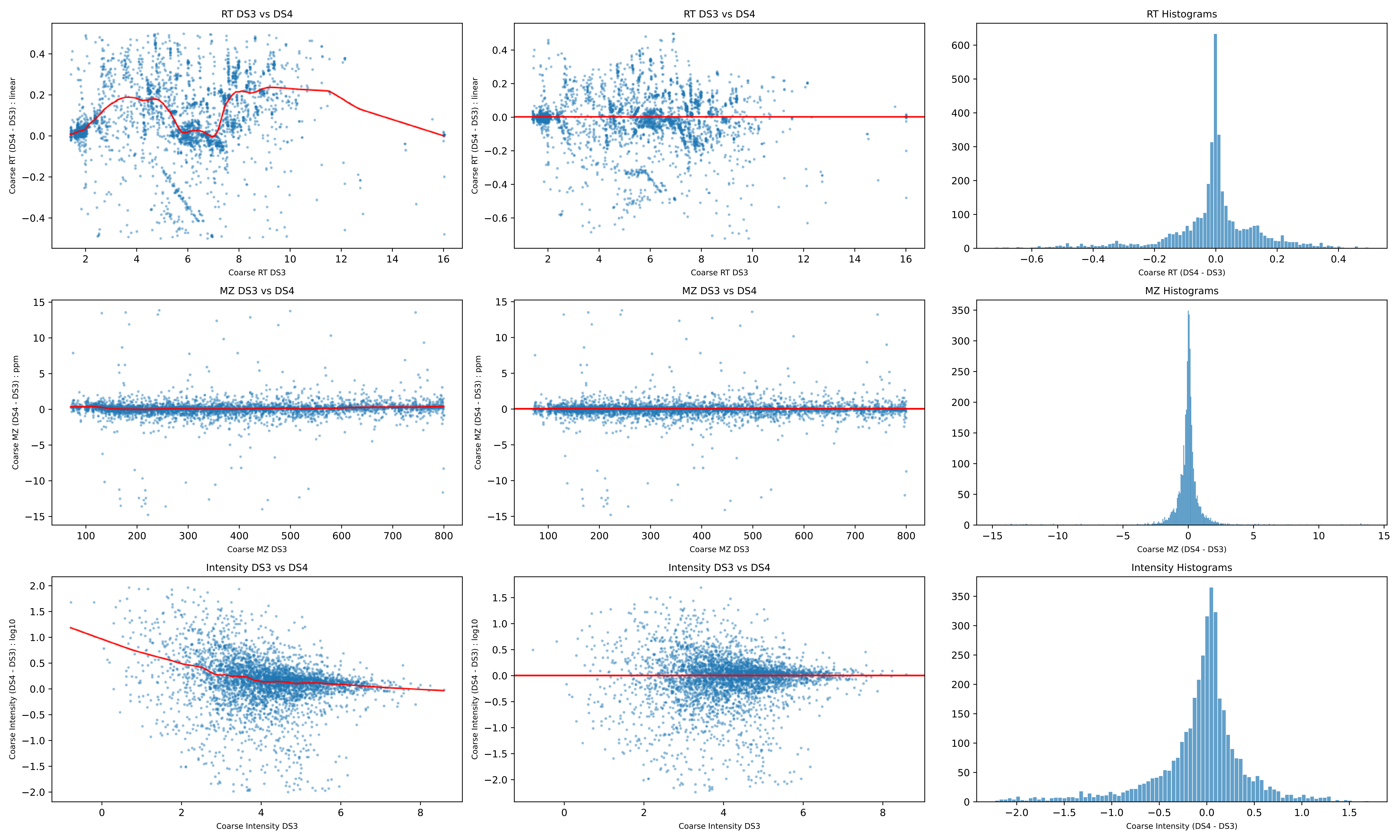
RT DS3 vs DS4
Coarse RT (DS4 - DS3) : linear
Coarse RT DS3
RT DS3 vs DS4
Coarse RT (DS4 - DS3) : linear
Coarse RT DS3
RT Histograms
Coarse RT (DS4 - DS3)
MZ DS3 vs DS4
Coarse MZ (DS4 - DS3) : ppm
Coarse MZ DS3
MZ DS3 vs DS4
Coarse MZ (DS4 - DS3) : ppm
Coarse MZ DS3
MZ Histograms
Coarse MZ (DS4 - DS3)
Intensity DS3 vs DS4
Coarse Intensity (DS4 - DS3) : log10
Coarse Intensity DS3
Intensity DS3 vs DS4
Coarse Intensity (DS4 - DS3) : log10
Coarse Intensity DS3
Intensity Histograms
Coarse Intensity (DS4 - DS3)

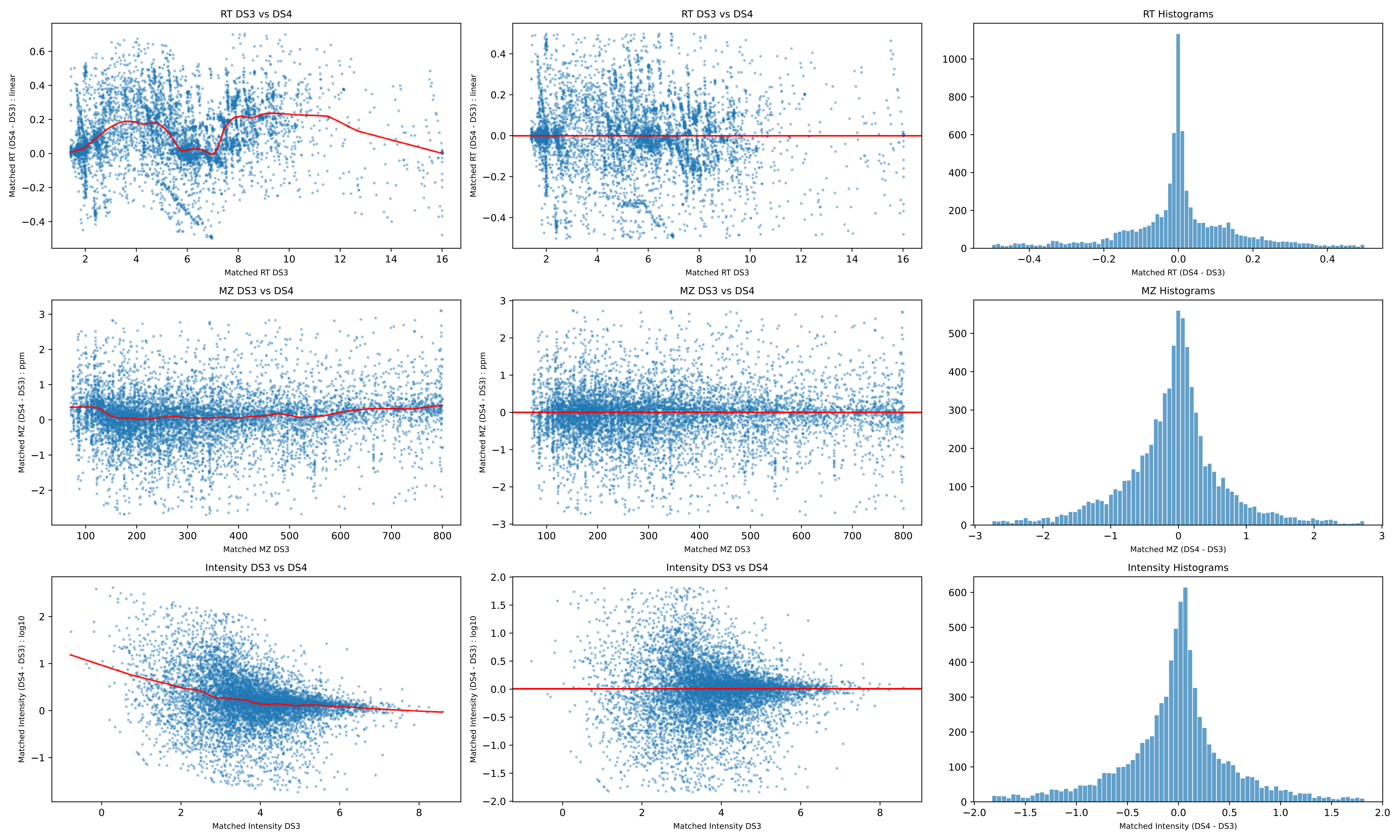
RT DS3 vs DS4
Matched RT (DS4 - DS3) : linear
Matched RT DS3
RT Histograms
Matched RT (DS4 - DS3)
MZ DS3 vs DS4
Matched MZ (DS4 - DS3) : ppm
Matched MZ DS3
MZ Histograms
Matched MZ (DS4 - DS3)
Intensity DS3 vs DS4
Matched Intensity (DS4 - DS3) : log10
Matched Intensity DS3
Intensity Histograms
Matched Intensity (DS4 - DS3)

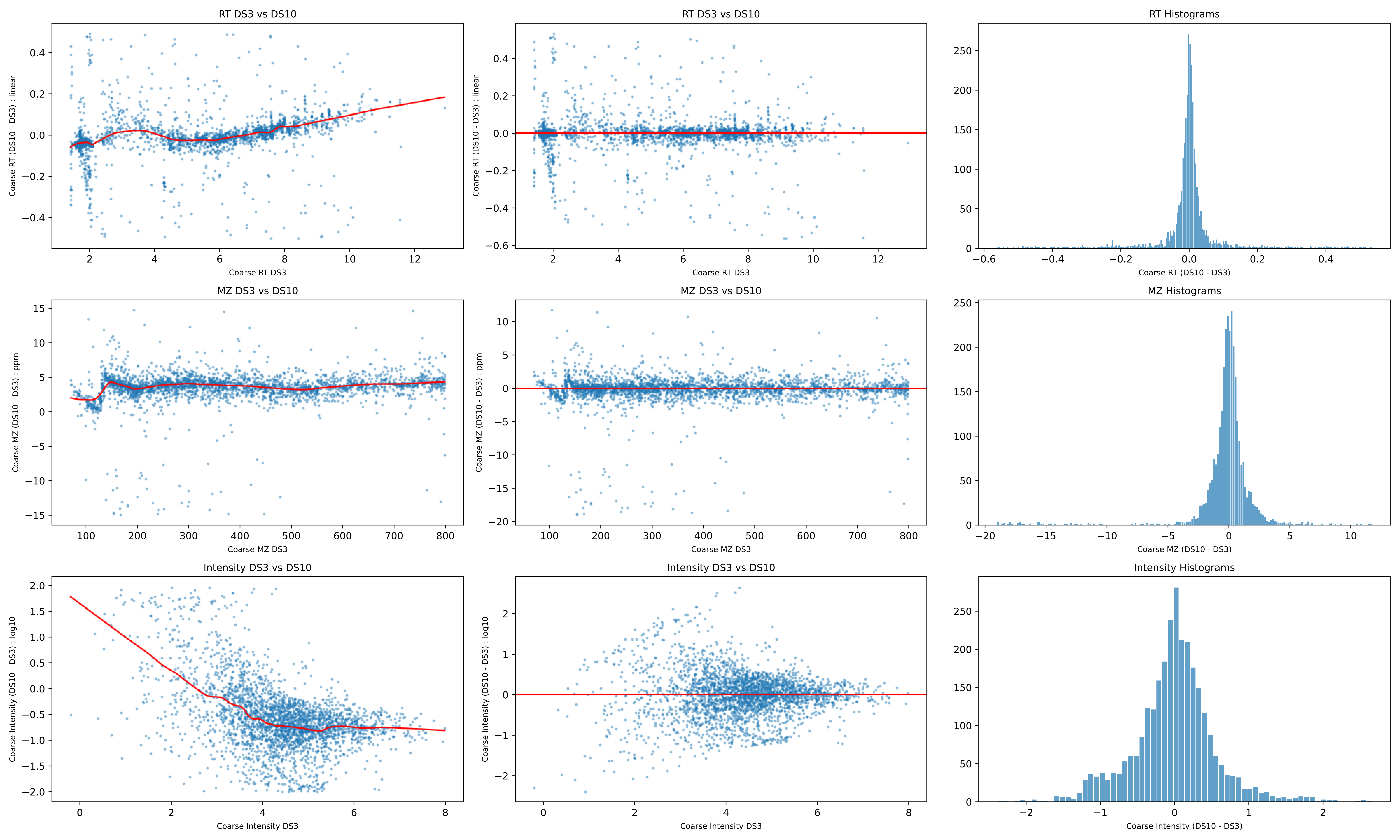
RT DS3 vs DS10
Coarse RT (DS10 - DS3) : linear
Coarse RT DS3
RT DS3 vs DS10
Coarse RT (DS10 - DS3) : linear
Coarse RT DS3
RT Histograms
Coarse RT (DS10 - DS3)
MZ DS3 vs DS10
Coarse MZ (DS10 - DS3) : ppm
Coarse MZ DS3
MZ DS3 vs DS10
Coarse MZ (DS10 - DS3) : ppm
Coarse MZ DS3
MZ Histograms
Coarse MZ (DS10 - DS3)
Intensity DS3 vs DS10
Coarse Intensity (DS10 - DS3) : log10
Coarse Intensity DS3
Intensity DS3 vs DS10
Coarse Intensity (DS10 - DS3) : log10
Coarse Intensity DS3
Intensity Histograms
Coarse Intensity (DS10 - DS3)

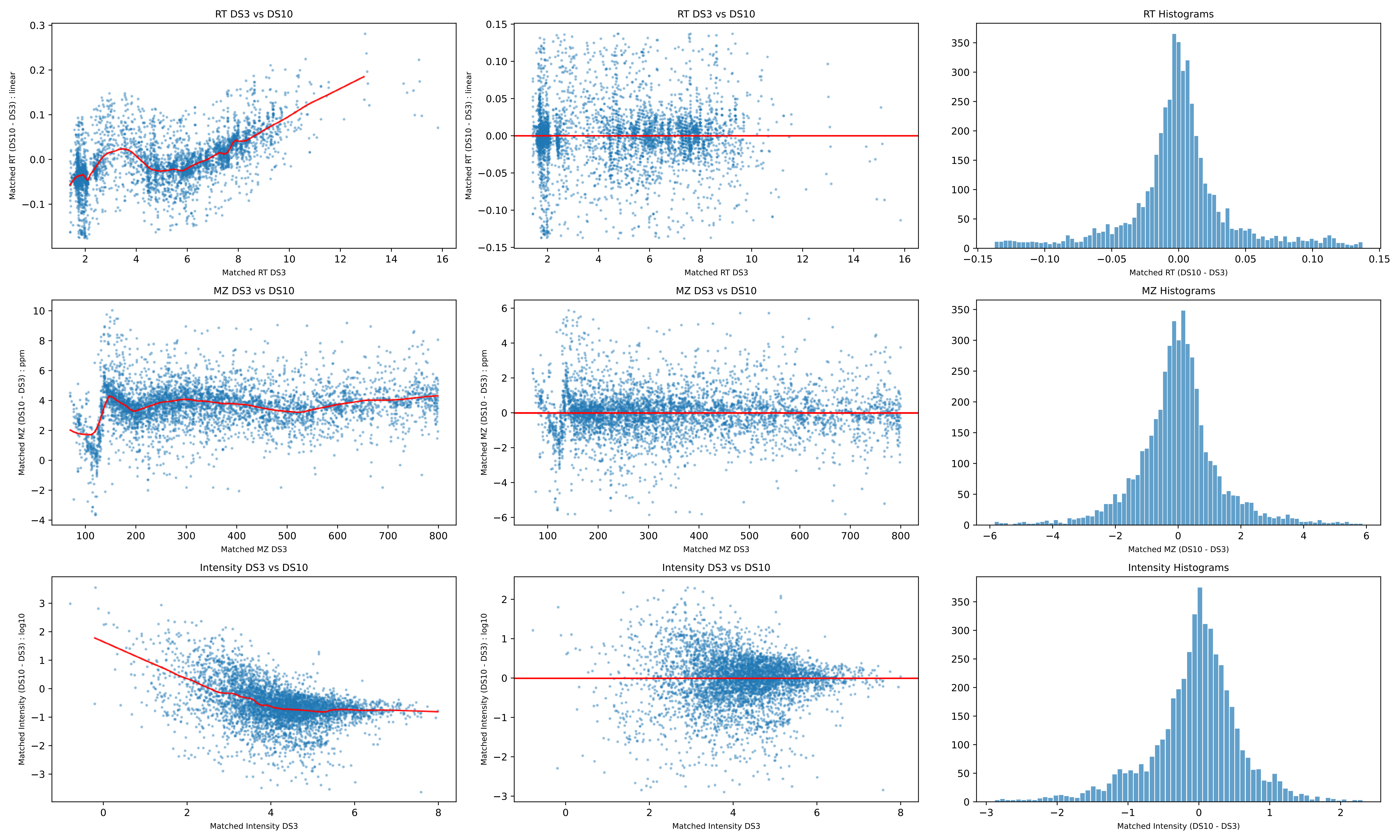
RT DS3 vs DS10
Matched RT (DS10 - DS3) : linear
Matched RT DS3
RT DS3 vs DS10
Matched RT (DS10 - DS3) : linear
Matched RT DS3
RT Histograms
Matched RT (DS10 - DS3)
MZ DS3 vs DS10
Matched MZ (DS10 - DS3) : ppm
Matched MZ DS3
MZ DS3 vs DS10
Matched MZ (DS10 - DS3) : ppm
Matched MZ DS3
MZ Histograms
Matched MZ (DS10 - DS3)
Intensity DS3 vs DS10
Matched Intensity (DS10 - DS3) : log10
Matched Intensity DS3
Intensity DS3 vs DS10
Matched Intensity (DS10 - DS3) : log10
Matched Intensity DS3
Intensity Histograms
Matched Intensity (DS10 - DS3)

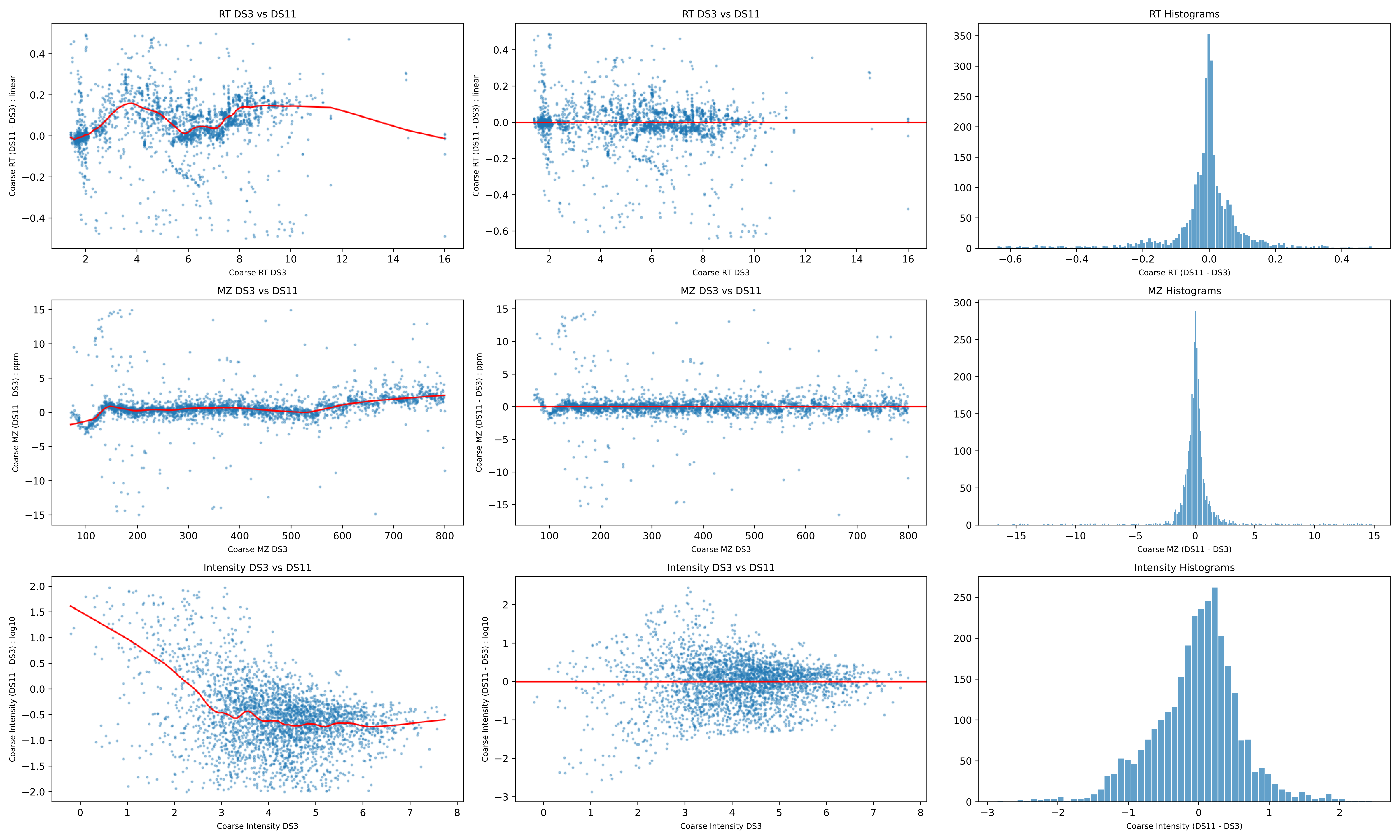
RT DS3 vs DS11
Coarse RT (DS11 - DS3) : linear
Coarse RT DS3
RT DS3 vs DS11
Coarse RT (DS11 - DS3) : linear
Coarse RT DS3
RT Histograms
Coarse RT (DS11 - DS3)
MZ DS3 vs DS11
Coarse MZ (DS11 - DS3) : ppm
Coarse MZ DS3
MZ DS3 vs DS11
Coarse MZ (DS11 - DS3) : ppm
Coarse MZ DS3
MZ Histograms
Coarse MZ (DS11 - DS3)
Intensity DS3 vs DS11
Coarse Intensity (DS11 - DS3) : log10
Coarse Intensity DS3
Intensity DS3 vs DS11
Coarse Intensity (DS11 - DS3) : log10
Coarse Intensity DS3
Intensity Histograms
Coarse Intensity (DS11 - DS3)

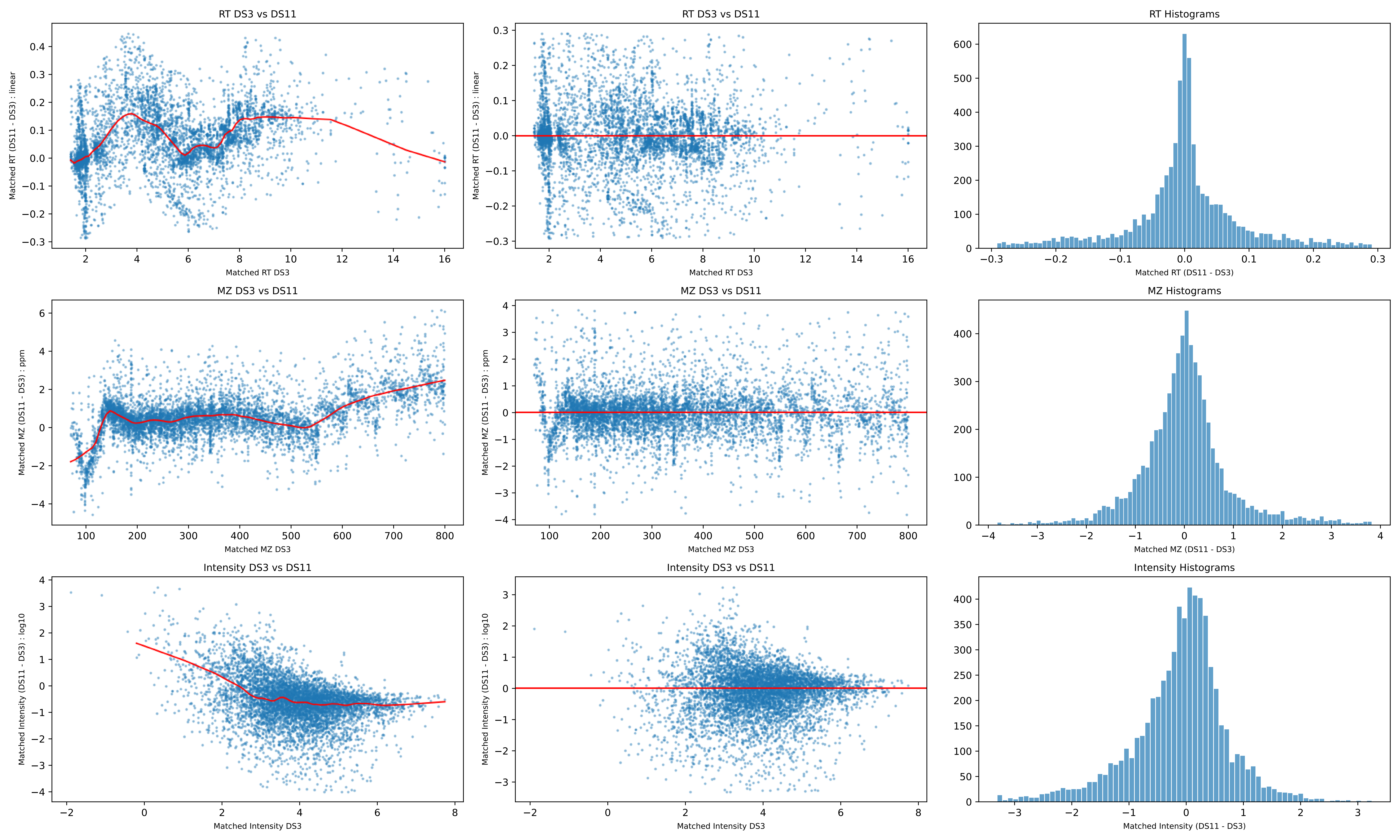
RT DS3 vs DS11
Matched RT (DS11 - DS3) : linear
Matched RT DS3
RT DS3 vs DS11
Matched RT (DS11 - DS3) : linear
Matched RT DS3
RT Histograms
Matched RT (DS11 - DS3)
MZ DS3 vs DS11
Matched MZ (DS11 - DS3) : ppm
Matched MZ DS3
MZ DS3 vs DS11
Matched MZ (DS11 - DS3) : ppm
Matched MZ DS3
MZ Histograms
Matched MZ (DS11 - DS3)
Intensity DS3 vs DS11
Matched Intensity (DS11 - DS3) : log10
Matched Intensity DS3
Intensity DS3 vs DS11
Matched Intensity (DS11 - DS3) : log10
Matched Intensity DS3
Intensity Histograms
Matched Intensity (DS11 - DS3)

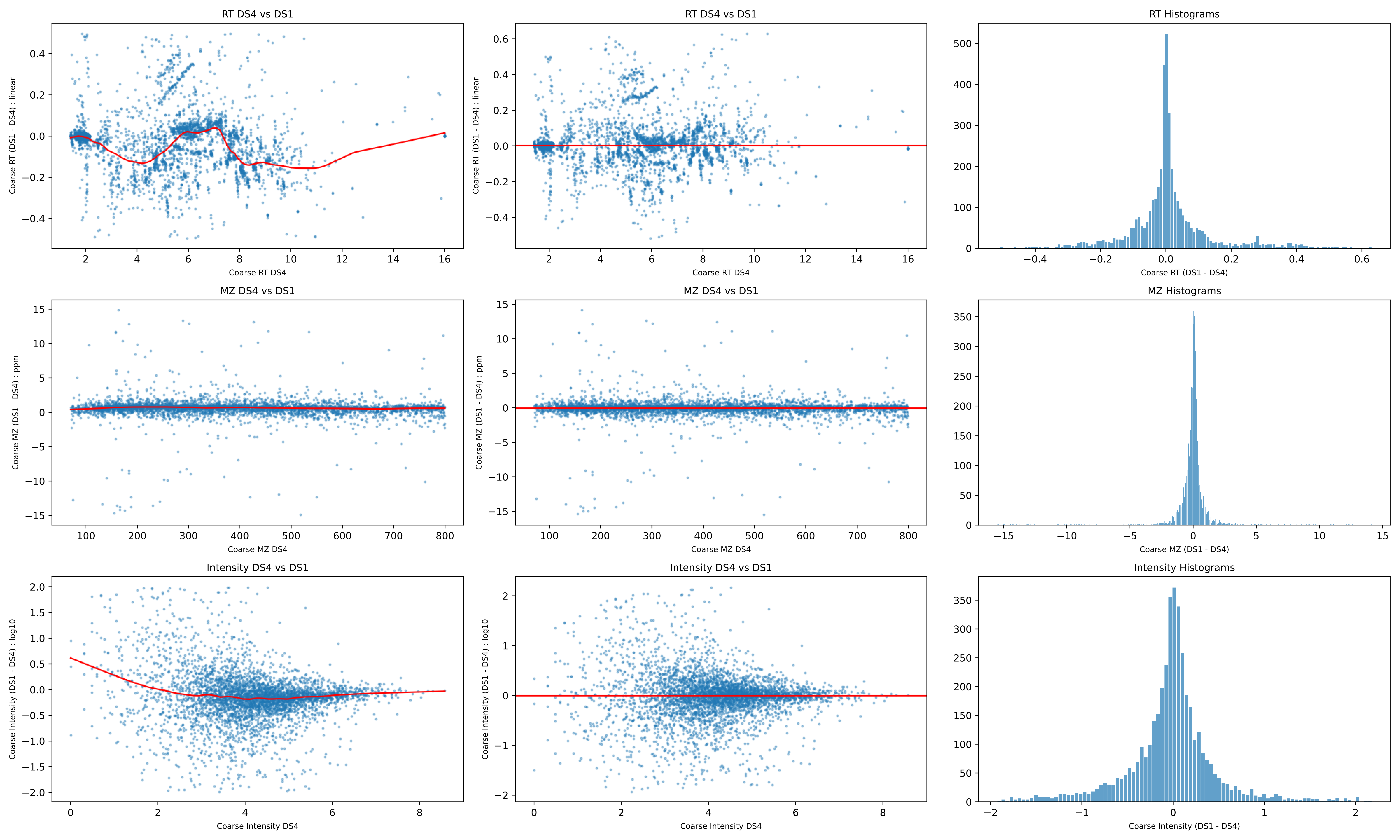
RT DS4 vs DS1
Coarse RT (DS1 - DS4) : linear
Coarse RT DS4
RT DS4 vs DS1
Coarse RT (DS1 - DS4) : linear
Coarse RT DS4
RT Histograms
Coarse RT (DS1 - DS4)
MZ DS4 vs DS1
Coarse MZ (DS1 - DS4) : ppm
Coarse MZ DS4
MZ DS4 vs DS1
Coarse MZ (DS1 - DS4) : ppm
Coarse MZ DS4
MZ Histograms
Coarse MZ (DS1 - DS4)
Intensity DS4 vs DS1
Coarse Intensity (DS1 - DS4) : log10
Coarse Intensity DS4
Intensity DS4 vs DS1
Coarse Intensity (DS1 - DS4) : log10
Coarse Intensity DS4
Intensity Histograms
Coarse Intensity (DS1 - DS4)

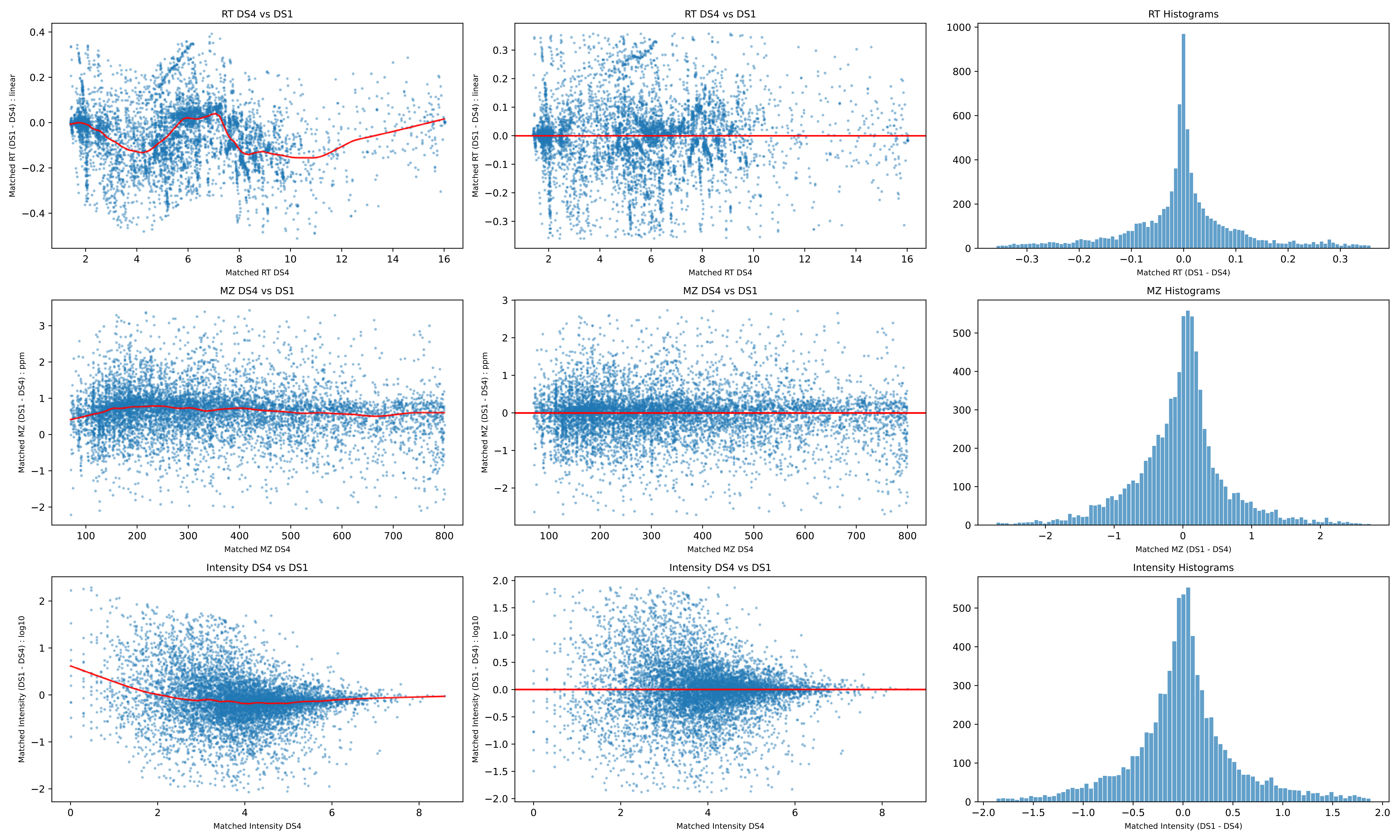
RT DS4 vs DS1
Matched RT (DS1 - DS4) : linear
Matched RT DS4
RT DS4 vs DS1
Matched RT (DS1 - DS4) : linear
Matched RT DS4
RT Histograms
Matched RT (DS1 - DS4)
MZ DS4 vs DS1
Matched MZ (DS1 - DS4) : ppm
Matched MZ DS4
MZ DS4 vs DS1
Matched MZ (DS1 - DS4) : ppm
Matched MZ DS4
MZ Histograms
Matched MZ (DS1 - DS4)
Intensity DS4 vs DS1
Matched Intensity (DS1 - DS4) : log10
Matched Intensity DS4
Intensity DS4 vs DS1
Matched Intensity (DS1 - DS4) : log10
Matched Intensity DS4
Intensity Histograms
Matched Intensity (DS1 - DS4)

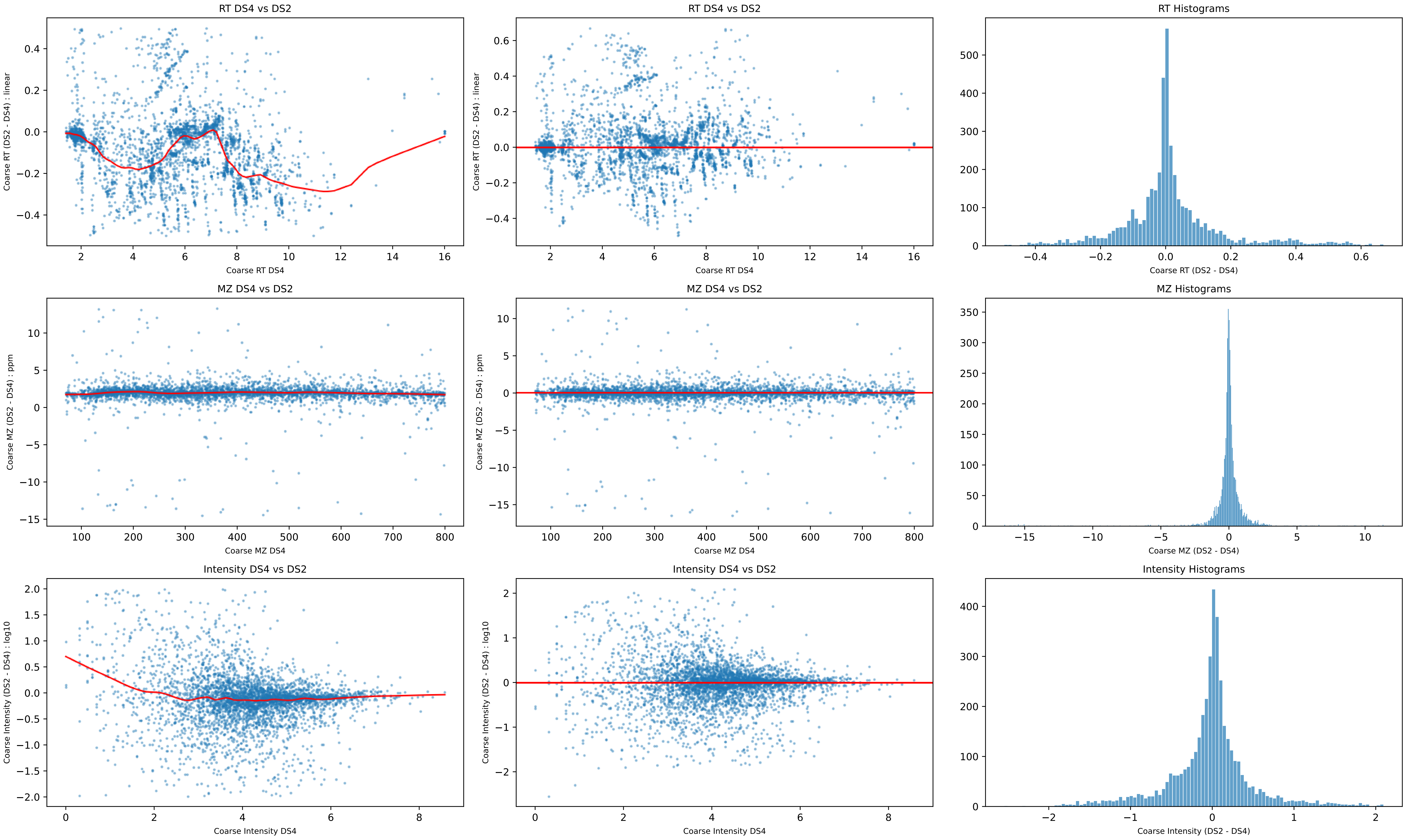
RT DS4 vs DS2
Coarse RT (DS2 - DS4) : linear
Coarse RT DS4
RT DS4 vs DS2
Coarse RT (DS2 - DS4) : linear
Coarse RT DS4
RT Histograms
Coarse RT (DS2 - DS4)
MZ DS4 vs DS2
Coarse MZ (DS2 - DS4) : ppm
Coarse MZ DS4
MZ DS4 vs DS2
Coarse MZ (DS2 - DS4) : ppm
Coarse MZ DS4
MZ Histograms
Coarse MZ (DS2 - DS4)
Intensity DS4 vs DS2
Coarse Intensity (DS2 - DS4) : log10
Coarse Intensity DS4
Intensity DS4 vs DS2
Coarse Intensity (DS2 - DS4) : log10
Coarse Intensity DS4
Intensity Histograms
Coarse Intensity (DS2 - DS4)

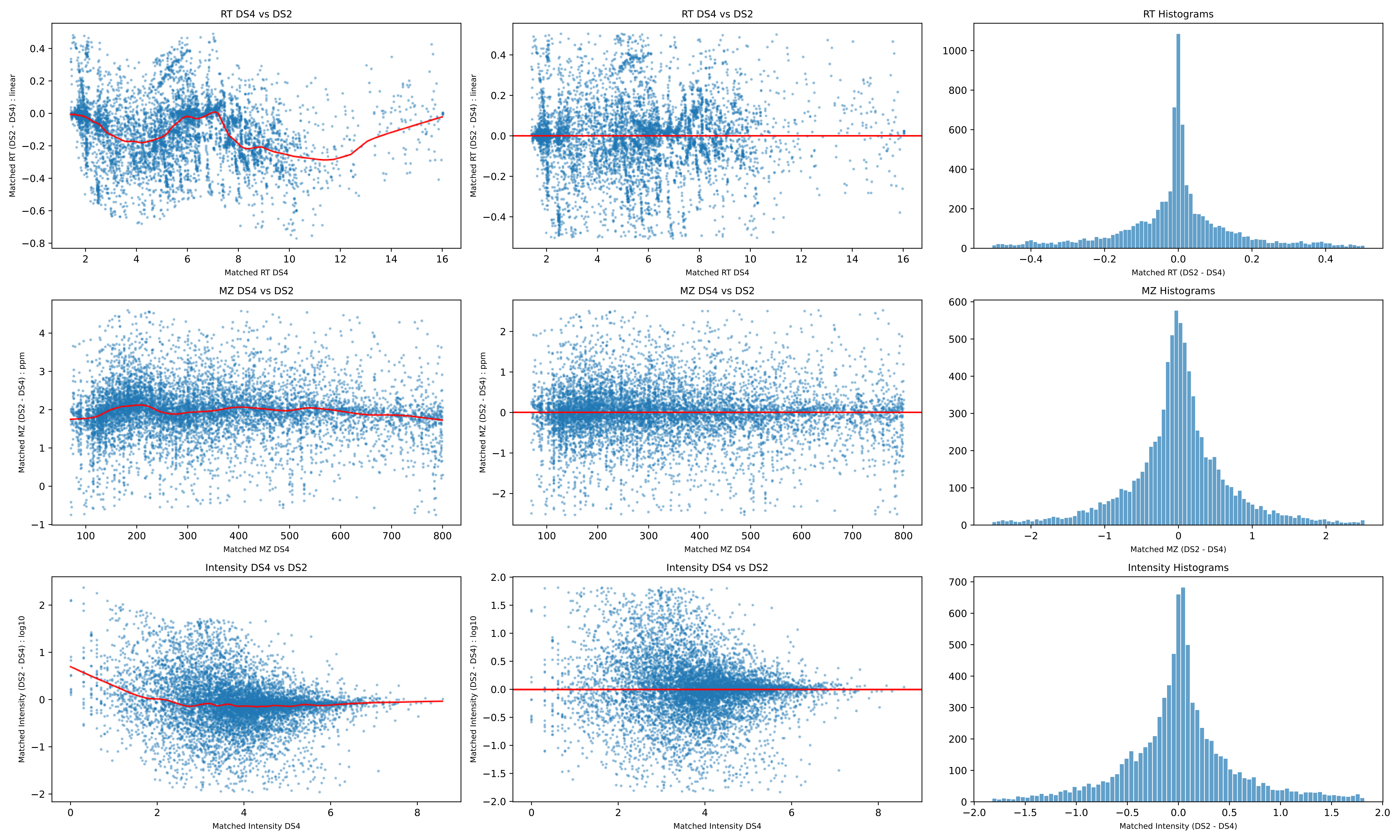
RT DS4 vs DS2
Matched RT (DS2 - DS4) : linear
Matched RT DS4
RT DS4 vs DS2
Matched RT (DS2 - DS4) : linear
Matched RT DS4
RT Histograms
Matched RT (DS2 - DS4)
MZ DS4 vs DS2
Matched MZ (DS2 - DS4) : ppm
Matched MZ DS4
MZ DS4 vs DS2
Matched MZ (DS2 - DS4) : ppm
Matched MZ DS4
MZ Histograms
Matched MZ (DS2 - DS4)
Intensity DS4 vs DS2
Matched Intensity (DS2 - DS4) : log10
Matched Intensity DS4
Intensity DS4 vs DS2
Matched Intensity (DS2 - DS4) : log10
Matched Intensity DS4
Intensity Histograms
Matched Intensity (DS2 - DS4)

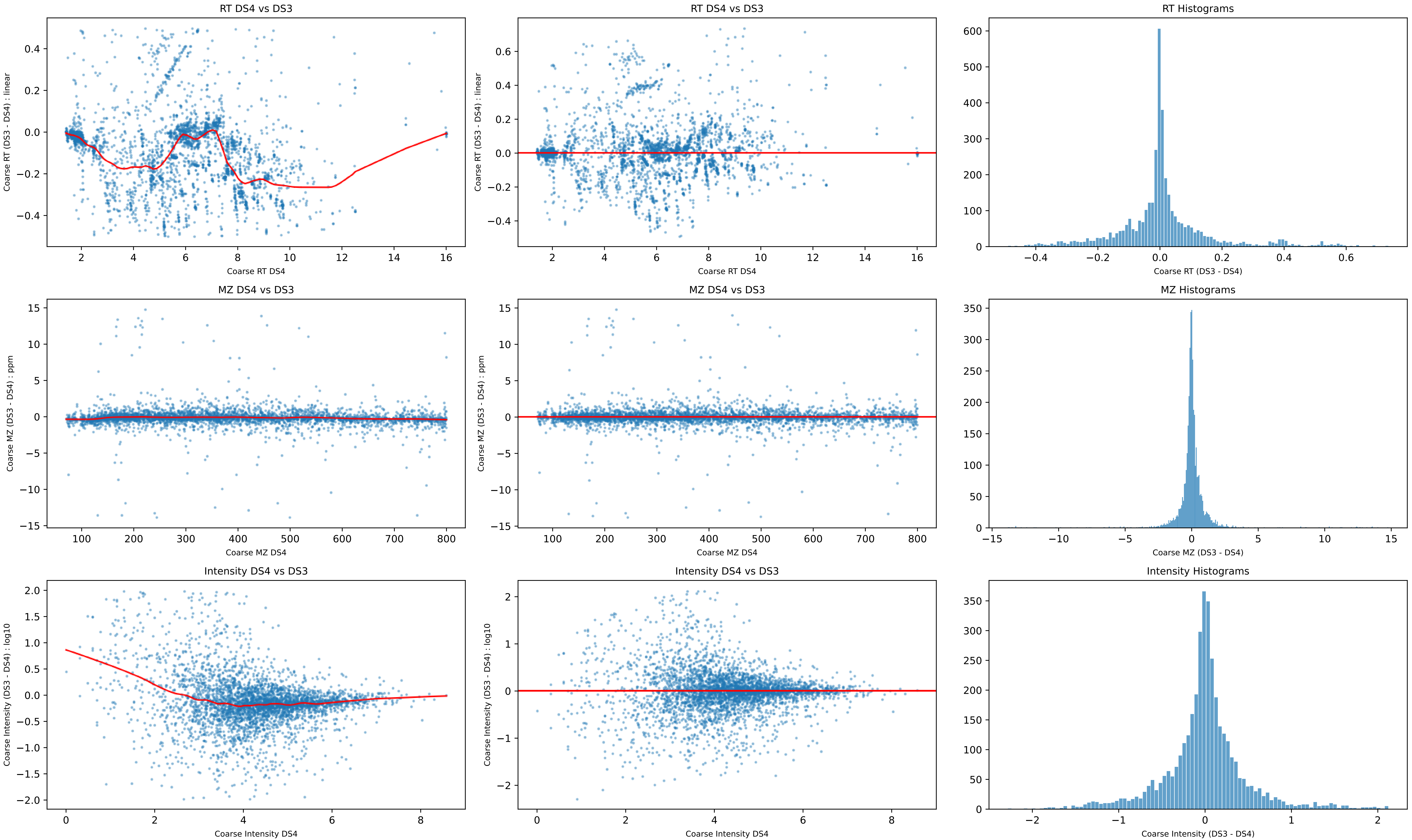
RT DS4 vs DS3
Coarse RT (DS3 - DS4) : linear
Coarse RT DS4
RT DS4 vs DS3
Coarse RT (DS3 - DS4) : linear
Coarse RT DS4
RT Histograms
Coarse RT (DS3 - DS4)
MZ DS4 vs DS3
Coarse MZ (DS3 - DS4) : ppm
Coarse MZ DS4
MZ DS4 vs DS3
Coarse MZ (DS3 - DS4) : ppm
Coarse MZ DS4
MZ Histograms
Coarse MZ (DS3 - DS4)
Intensity DS4 vs DS3
Coarse Intensity (DS3 - DS4) : log10
Coarse Intensity DS4
Intensity DS4 vs DS3
Coarse Intensity (DS3 - DS4) : log10
Coarse Intensity DS4
Intensity Histograms
Coarse Intensity (DS3 - DS4)

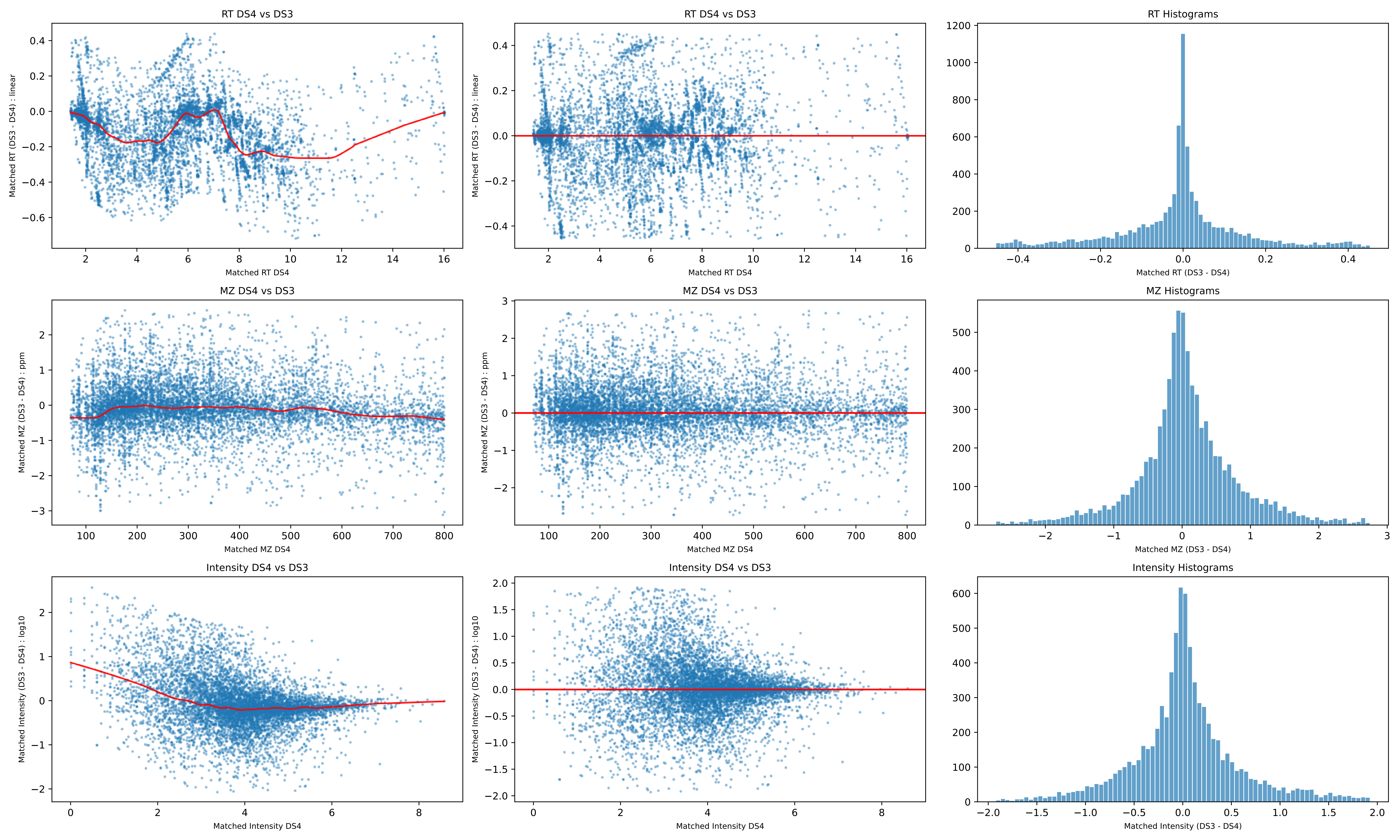
RT DS4 vs DS3
Matched RT (DS3 - DS4) : linear
Matched RT DS4
RT DS4 vs DS3
Matched RT (DS3 - DS4) : linear
Matched RT DS4
RT Histograms
Matched RT (DS3 - DS4)
MZ DS4 vs DS3
Matched MZ (DS3 - DS4) : ppm
Matched MZ DS4
MZ DS4 vs DS3
Matched MZ (DS3 - DS4) : ppm
Matched MZ DS4
MZ Histograms
Matched MZ (DS3 - DS4)
Intensity DS4 vs DS3
Matched Intensity (DS3 - DS4) : log10
Matched Intensity DS4
Intensity DS4 vs DS3
Matched Intensity (DS3 - DS4) : log10
Matched Intensity DS4
Intensity Histograms
Matched Intensity (DS3 - DS4)

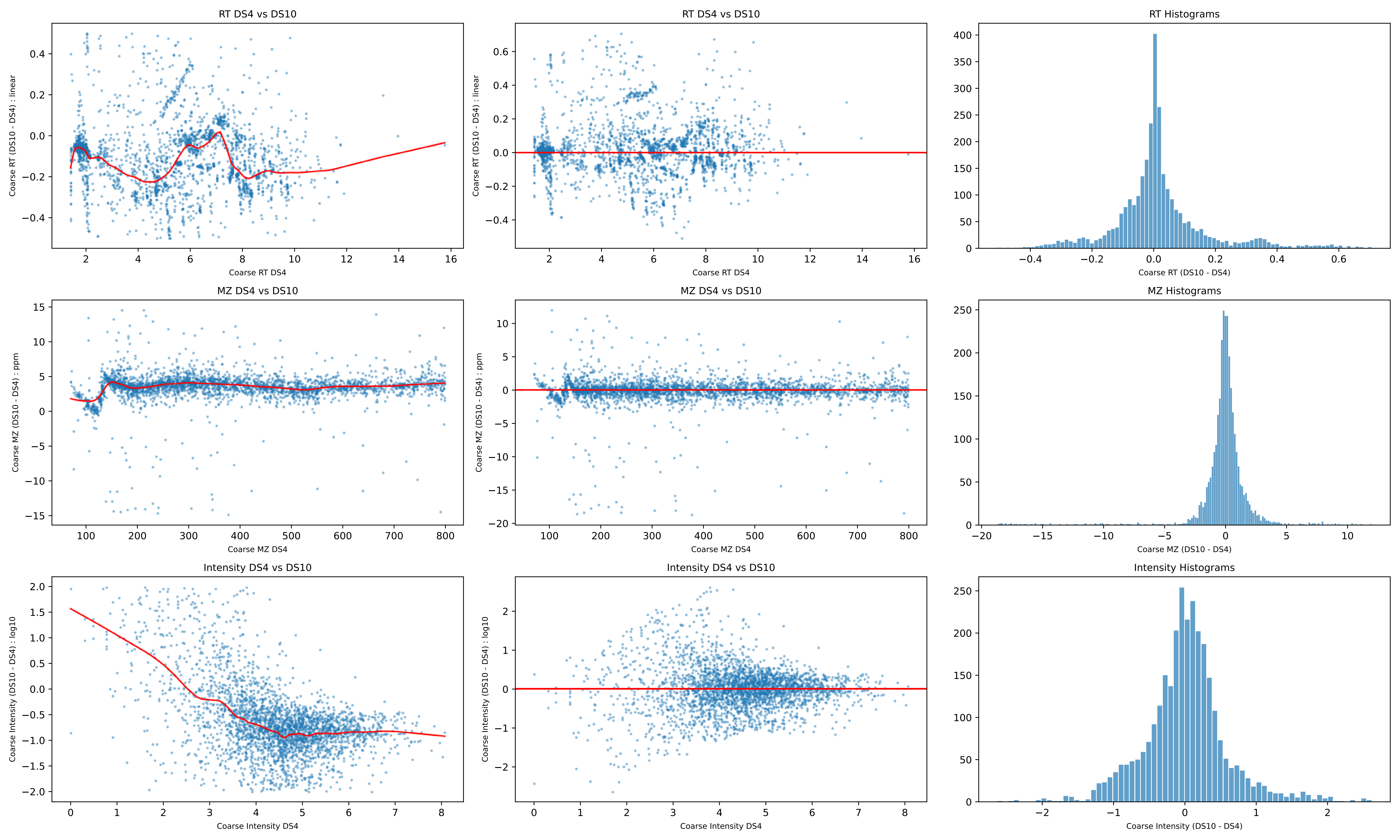
RT DS4 vs DS10
Coarse RT (DS10 - DS4) : linear
Coarse RT DS4
RT DS4 vs DS10
Coarse RT (DS10 - DS4) : linear
Coarse RT DS4
RT Histograms
Coarse RT (DS10 - DS4)
MZ DS4 vs DS10
Coarse MZ (DS10 - DS4) : ppm
Coarse MZ DS4
MZ DS4 vs DS10
Coarse MZ (DS10 - DS4) : ppm
Coarse MZ DS4
MZ Histograms
Coarse MZ (DS10 - DS4)
Intensity DS4 vs DS10
Coarse Intensity (DS10 - DS4) : log10
Coarse Intensity DS4
Intensity DS4 vs DS10
Coarse Intensity (DS10 - DS4) : log10
Coarse Intensity DS4
Intensity Histograms
Coarse Intensity (DS10 - DS4)

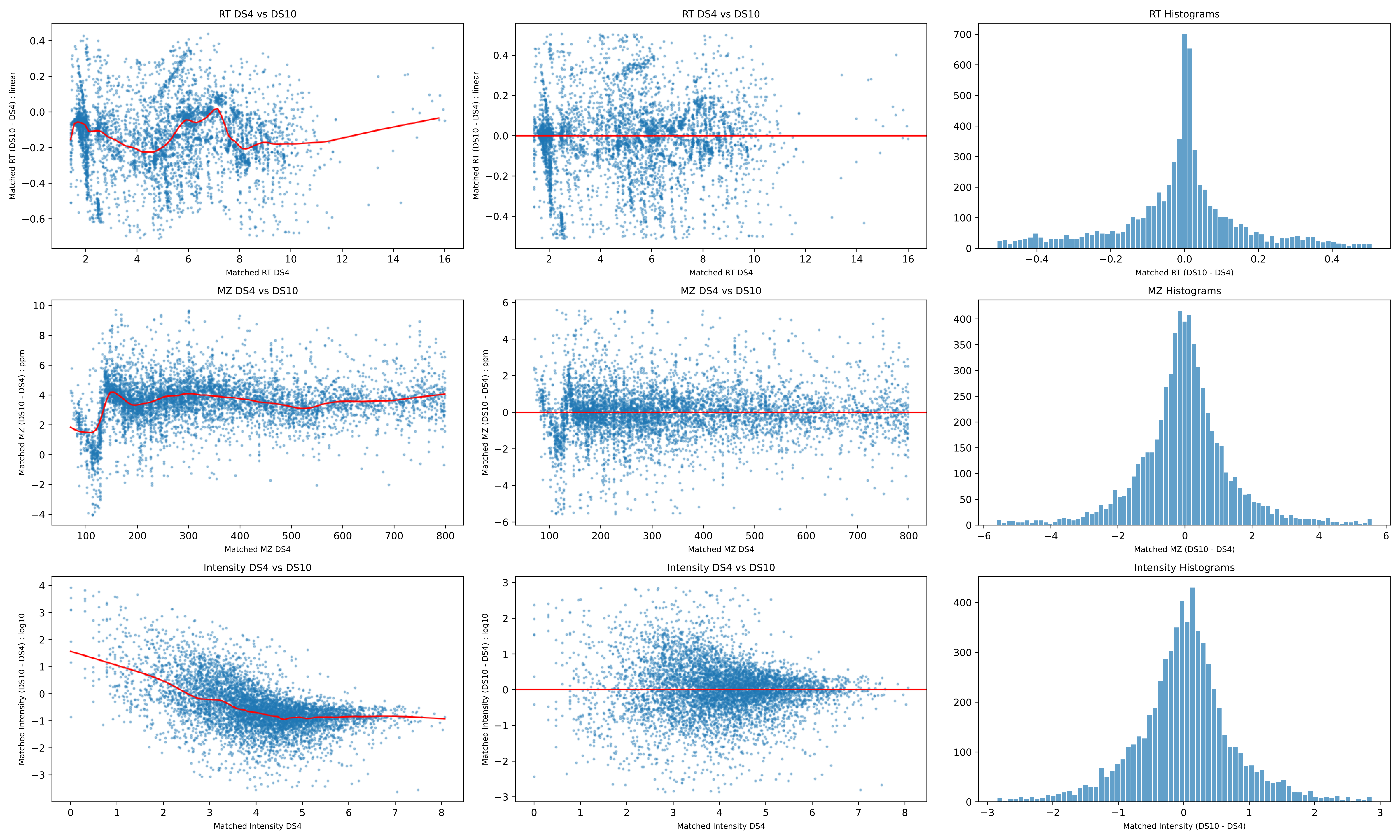
RT DS4 vs DS10
Matched RT (DS10 - DS4) : linear
Matched RT DS4
RT DS4 vs DS10
Matched RT (DS10 - DS4) : linear
Matched RT DS4
RT Histograms
Matched RT (DS10 - DS4)
MZ DS4 vs DS10
Matched MZ (DS10 - DS4) : ppm
Matched MZ DS4
MZ DS4 vs DS10
Matched MZ (DS10 - DS4) : ppm
Matched MZ DS4
MZ Histograms
Matched MZ (DS10 - DS4)
Intensity DS4 vs DS10
Matched Intensity (DS10 - DS4) : log10
Matched Intensity DS4
Intensity DS4 vs DS10
Matched Intensity (DS10 - DS4) : log10
Matched Intensity DS4
Intensity Histograms
Matched Intensity (DS10 - DS4)

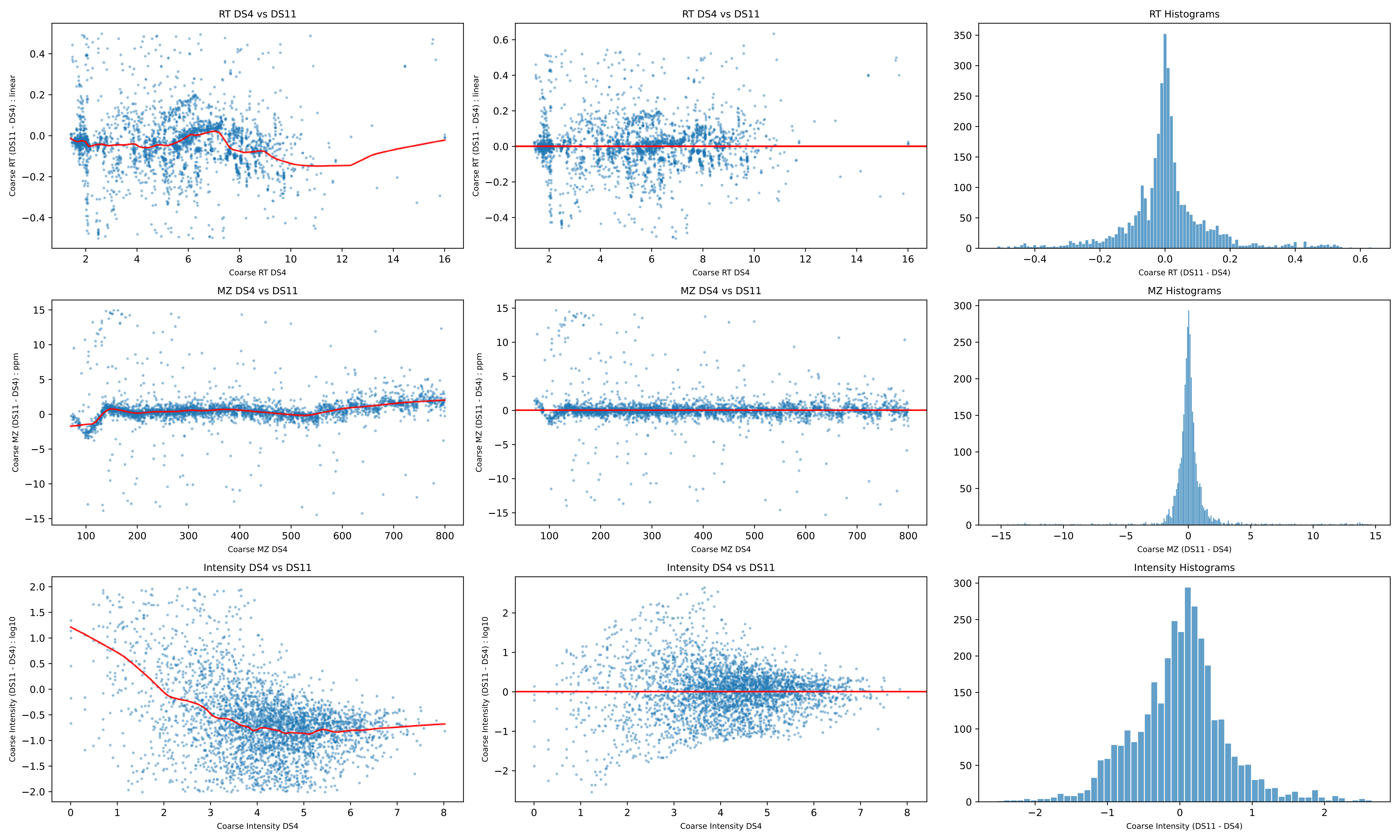

RT DS4 vs DS11
Coarse RT (DS11 - DS4) : linear
Coarse RT DS4
RT Histograms
Coarse RT (DS11 - DS4)
MZ DS4 vs DS11
Coarse MZ (DS11 - DS4) : ppm
Coarse MZ DS4
MZ Histograms
Coarse MZ (DS11 - DS4)
Intensity DS4 vs DS11
Coarse Intensity (DS11 - DS4) : log10
Coarse Intensity DS4
Intensity Histograms
Coarse Intensity (DS11 - DS4)

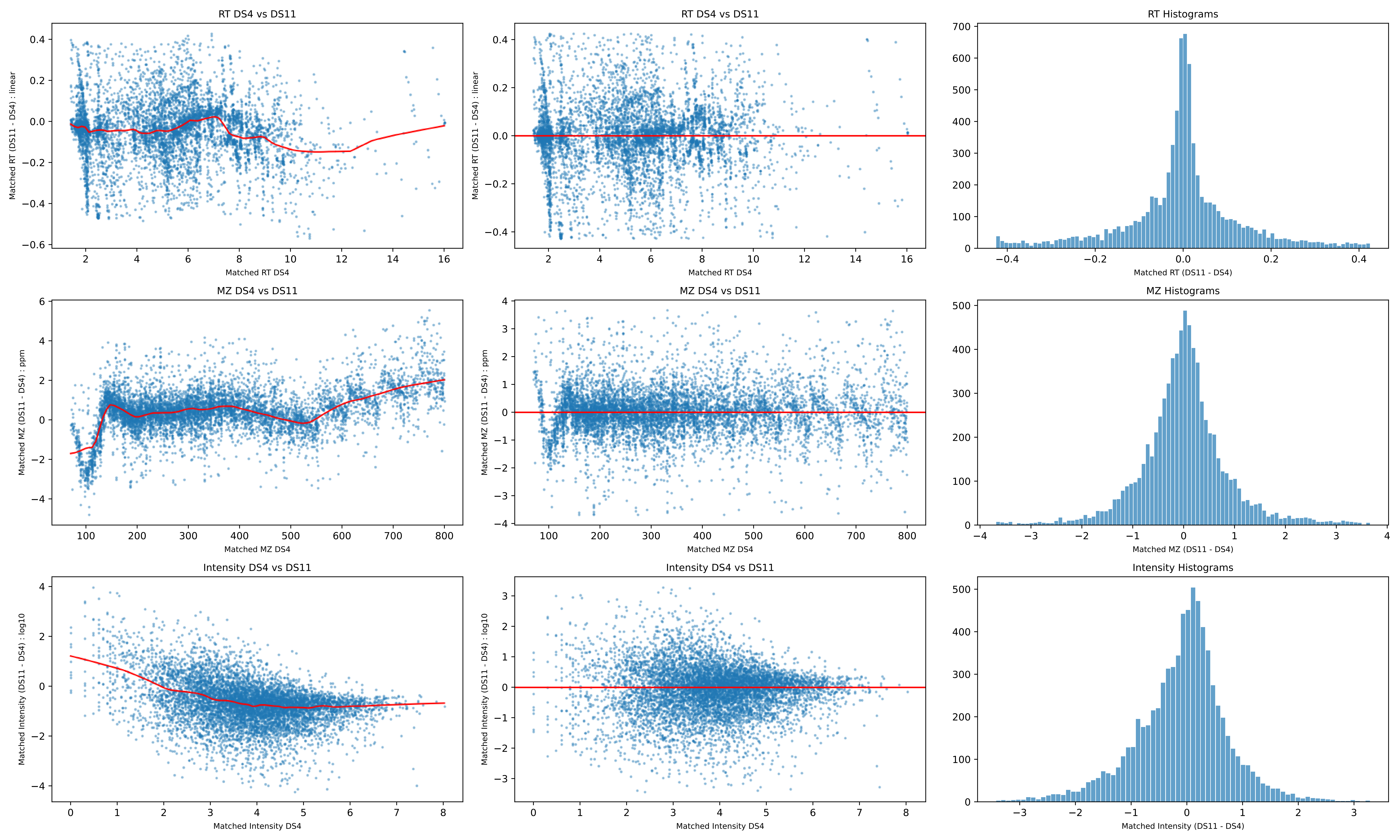
RT DS4 vs DS11
Matched RT (DS11 - DS4) : linear
Matched RT DS4
RT DS4 vs DS11
Matched RT (DS11 - DS4) : linear
Matched RT DS4
RT Histograms
Matched RT (DS11 - DS4)
MZ DS4 vs DS11
Matched MZ (DS11 - DS4) : ppm
Matched MZ DS4
MZ DS4 vs DS11
Matched MZ (DS11 - DS4) : ppm
Matched MZ DS4
MZ Histograms
Matched MZ (DS11 - DS4)
Intensity DS4 vs DS11
Matched Intensity (DS11 - DS4) : log10
Matched Intensity DS4
Intensity DS4 vs DS11
Matched Intensity (DS11 - DS4) : log10
Matched Intensity DS4
Intensity Histograms
Matched Intensity (DS11 - DS4)

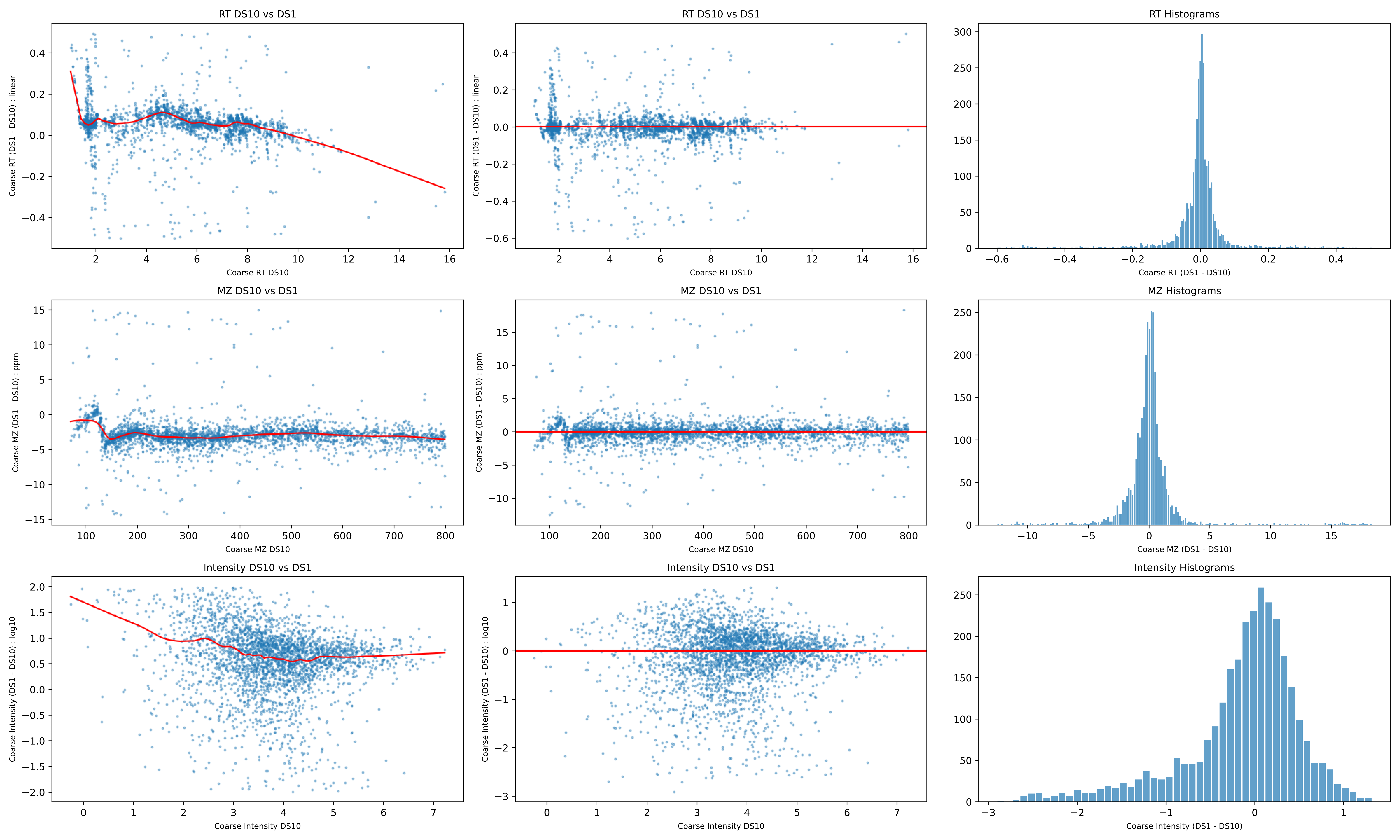
RT DS10 vs DS1
Coarse RT (DS1 - DS10) : linear
Coarse RT DS10
RT DS10 vs DS1
Coarse RT (DS1 - DS10) : linear
Coarse RT DS10
RT Histograms
Coarse RT (DS1 - DS10)
MZ DS10 vs DS1
Coarse MZ (DS1 - DS10) : ppm
Coarse MZ DS10
MZ DS10 vs DS1
Coarse MZ (DS1 - DS10) : ppm
Coarse MZ DS10
MZ Histograms
Coarse MZ (DS1 - DS10)
Intensity DS10 vs DS1
Coarse Intensity (DS1 - DS10) : log10
Coarse Intensity DS10
Intensity DS10 vs DS1
Coarse Intensity (DS1 - DS10) : log10
Coarse Intensity DS10
Intensity Histograms
Coarse Intensity (DS1 - DS10)

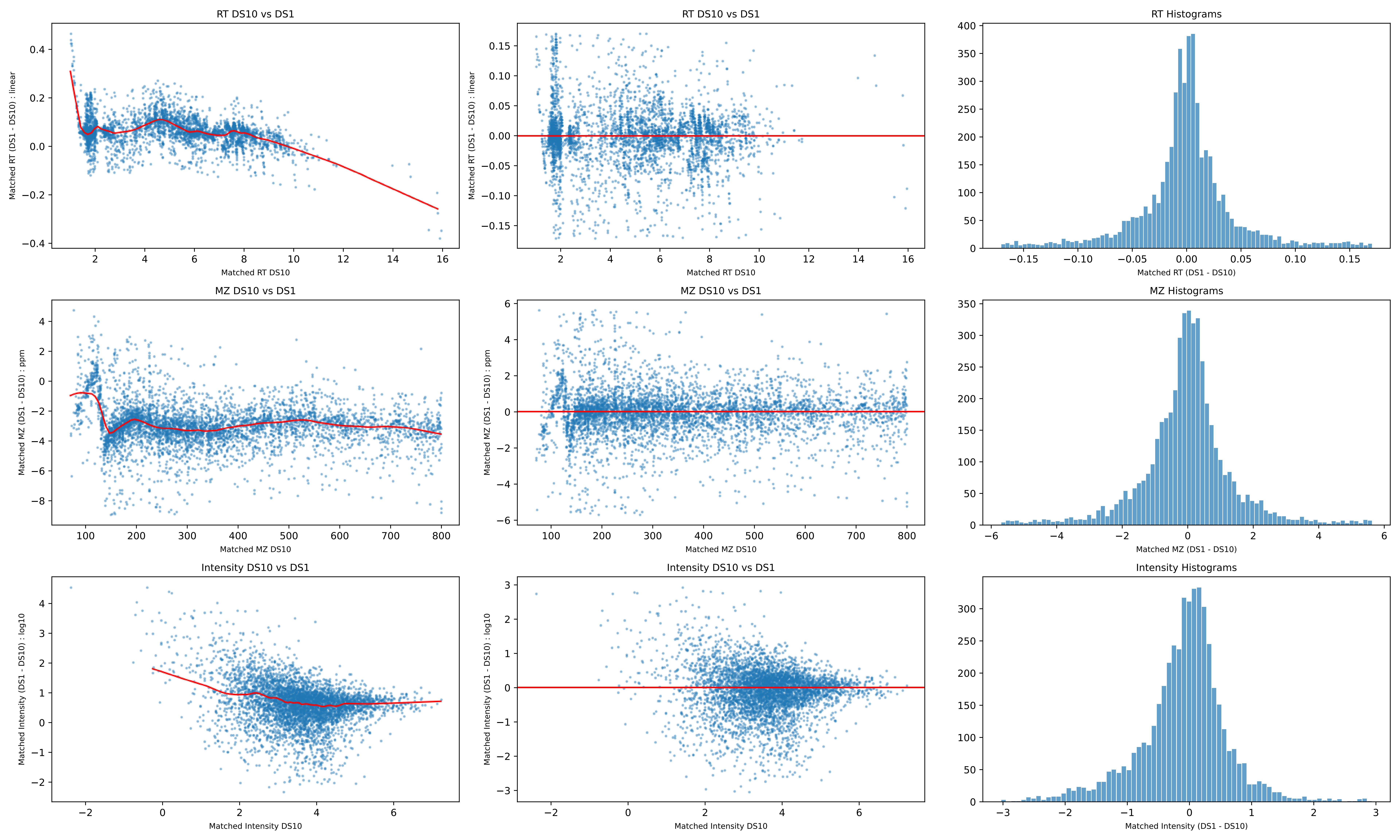
RT DS10 vs DS1
Matched RT (DS1 - DS10) : linear
Matched RT DS10
RT DS10 vs DS1
Matched RT (DS1 - DS10) : linear
Matched RT DS10
RT Histograms
Matched RT (DS1 - DS10)
MZ DS10 vs DS1
Matched MZ (DS1 - DS10) : ppm
Matched MZ DS10
MZ DS10 vs DS1
Matched MZ (DS1 - DS10) : ppm
Matched MZ DS10
MZ Histograms
Matched MZ (DS1 - DS10)
Intensity DS10 vs DS1
Matched Intensity (DS1 - DS10) : log10
Matched Intensity DS10
Intensity DS10 vs DS1
Matched Intensity (DS1 - DS10) : log10
Matched Intensity DS10
Intensity Histograms
Matched Intensity (DS1 - DS10)

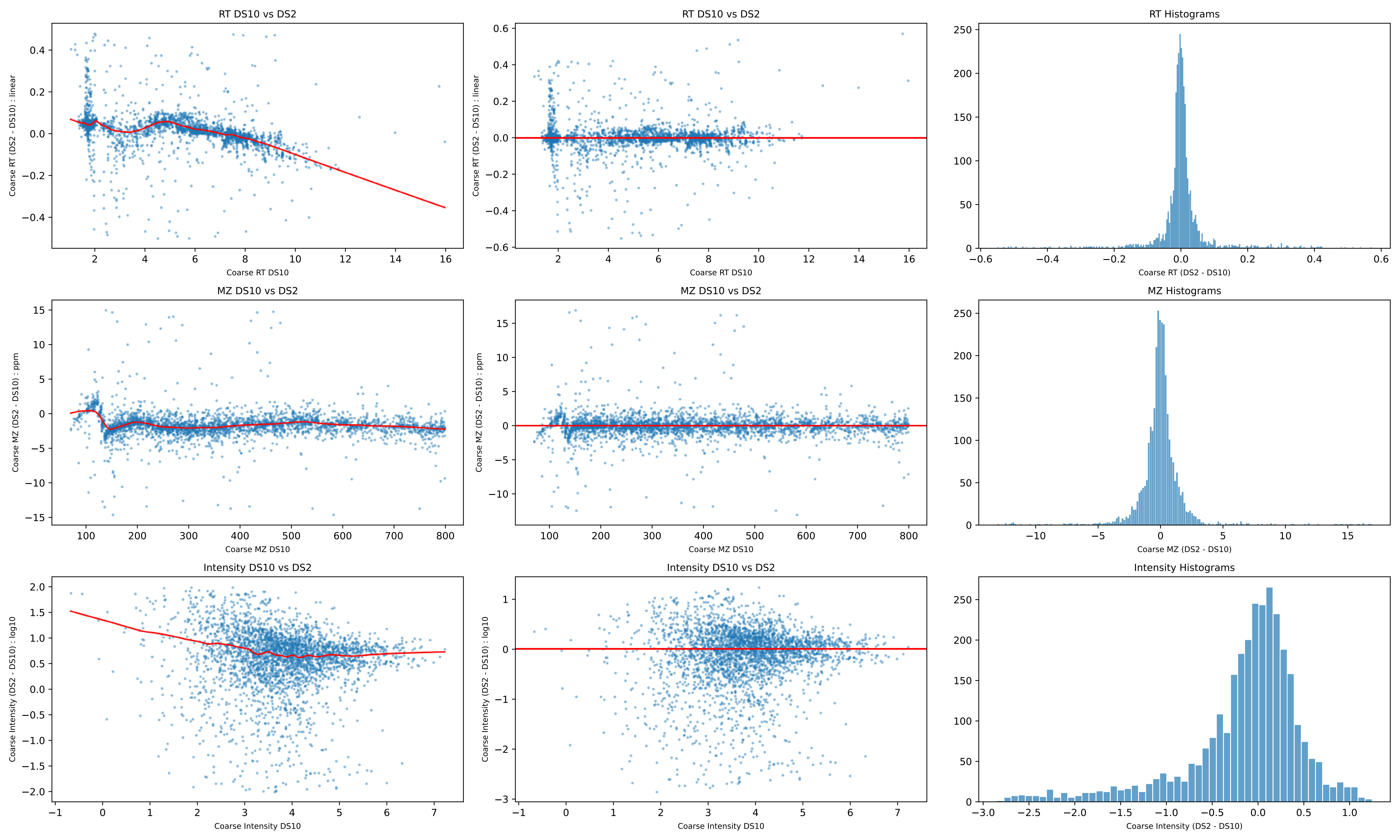

RT DS10 vs DS2
Coarse RT (DS2 - DS10) : linear
Coarse RT DS10
RT DS10 vs DS2
Coarse RT (DS2 - DS10) : linear
Coarse RT DS10
RT Histograms
Coarse RT (DS2 - DS10)
MZ DS10 vs DS2
Coarse MZ (DS2 - DS10) : ppm
Coarse MZ DS10
MZ DS10 vs DS2
Coarse MZ (DS2 - DS10) : ppm
Coarse MZ DS10
MZ Histograms
Coarse MZ (DS2 - DS10)
Intensity DS10 vs DS2
Coarse Intensity (DS2 - DS10) : log10
Coarse Intensity DS10
Intensity DS10 vs DS2
Coarse Intensity (DS2 - DS10) : log10
Coarse Intensity DS10
Intensity Histograms
Coarse Intensity (DS2 - DS10)

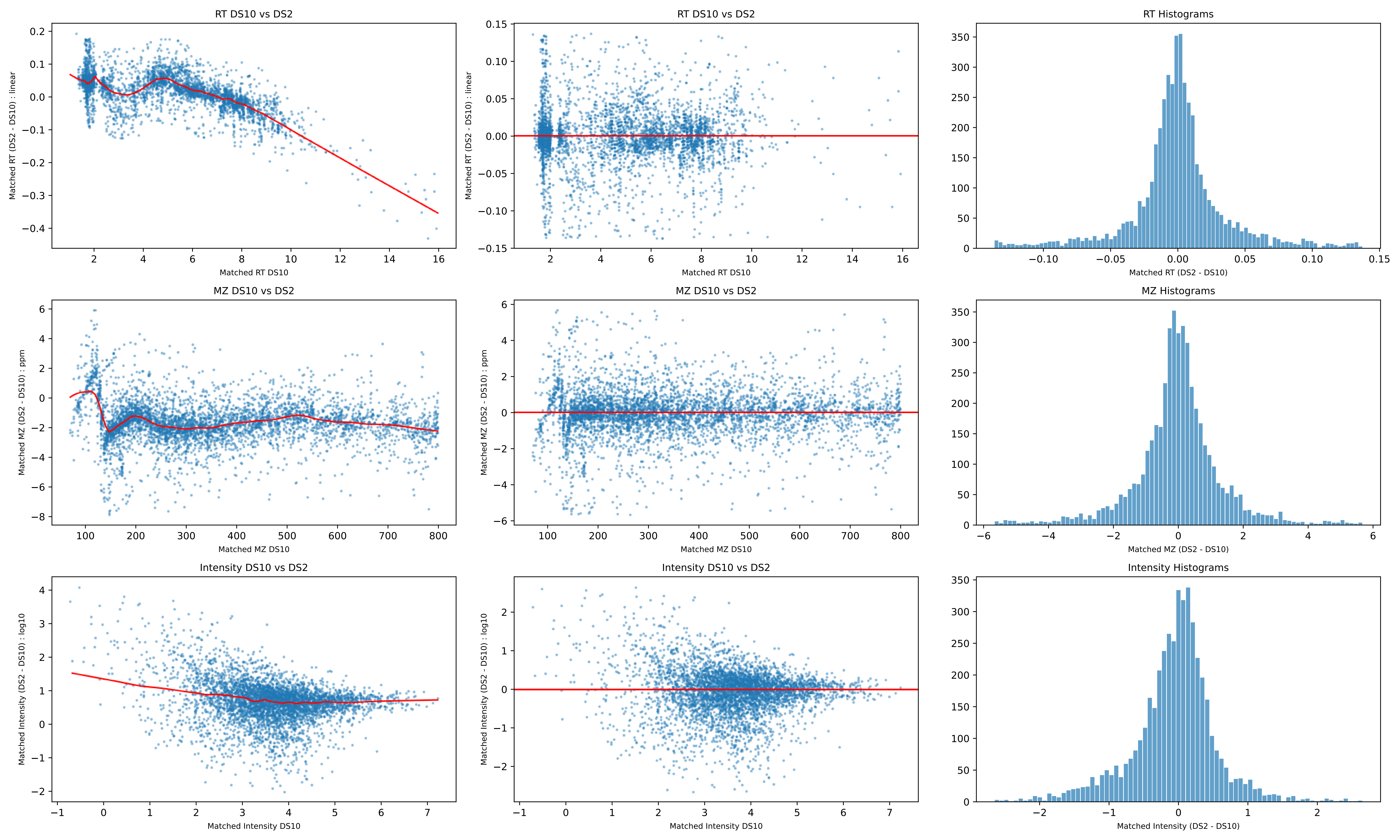
RT DS10 vs DS2
Matched RT (DS2 - DS10) : linear
Matched RT DS10
RT Histograms
Matched RT (DS2 - DS10)
MZ DS10 vs DS2
Matched MZ (DS2 - DS10) : ppm
Matched MZ DS10
MZ Histograms
Matched MZ (DS2 - DS10)
Intensity DS10 vs DS2
Matched Intensity (DS2 - DS10) : log10
Matched Intensity DS10
Intensity Histograms
Matched Intensity (DS2 - DS10)

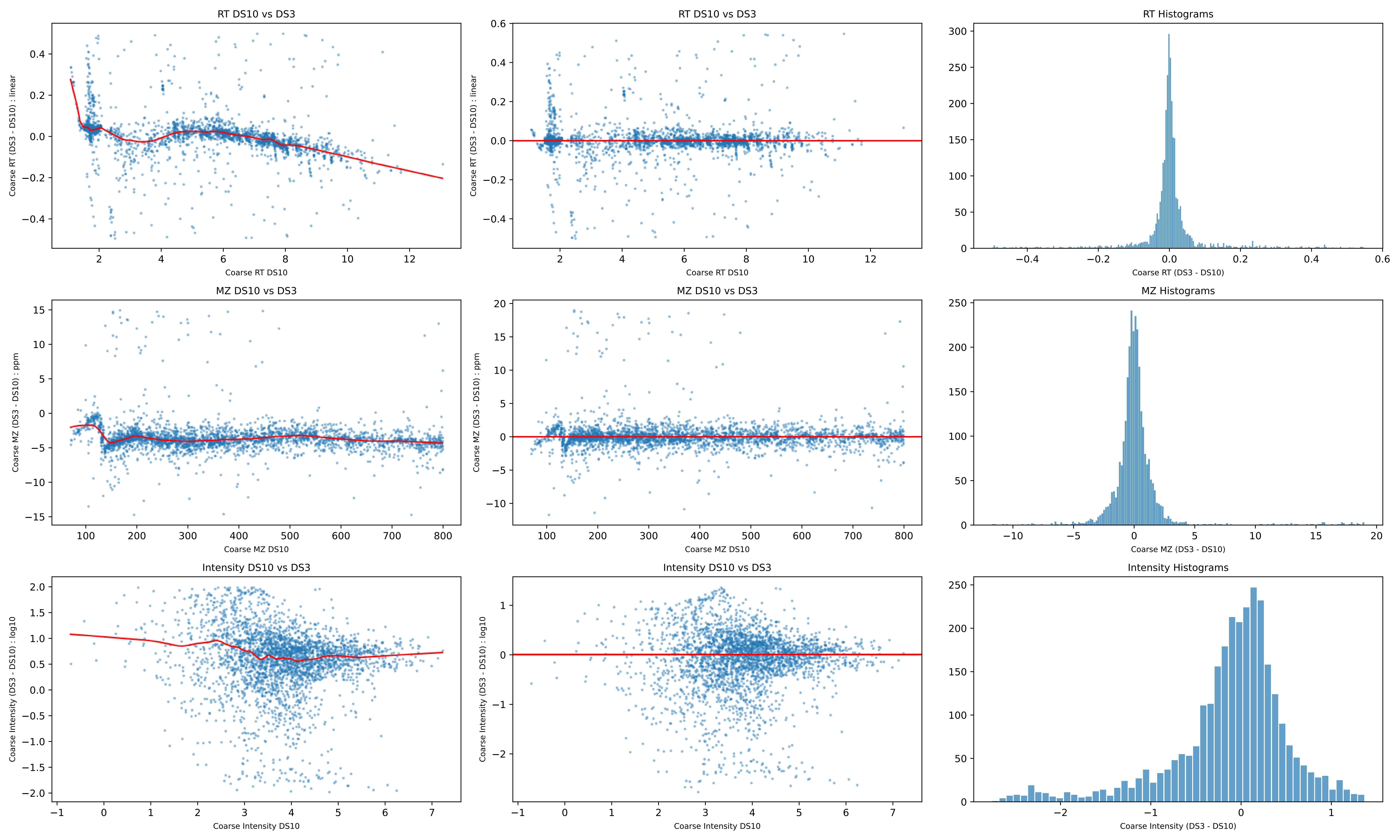
RT DS10 vs DS3
Coarse RT (DS3 - DS10) : linear
Coarse RT DS10
RT DS10 vs DS3
Coarse RT (DS3 - DS10) : linear
Coarse RT DS10
RT Histograms
Coarse RT (DS3 - DS10)
MZ DS10 vs DS3
Coarse MZ (DS3 - DS10) : ppm
Coarse MZ DS10
MZ DS10 vs DS3
Coarse MZ (DS3 - DS10) : ppm
Coarse MZ DS10
MZ Histograms
Coarse MZ (DS3 - DS10)
Intensity DS10 vs DS3
Coarse Intensity (DS3 - DS10) : log10
Coarse Intensity DS10
Intensity DS10 vs DS3
Coarse Intensity (DS3 - DS10) : log10
Coarse Intensity DS10
Intensity Histograms
Coarse Intensity (DS3 - DS10)

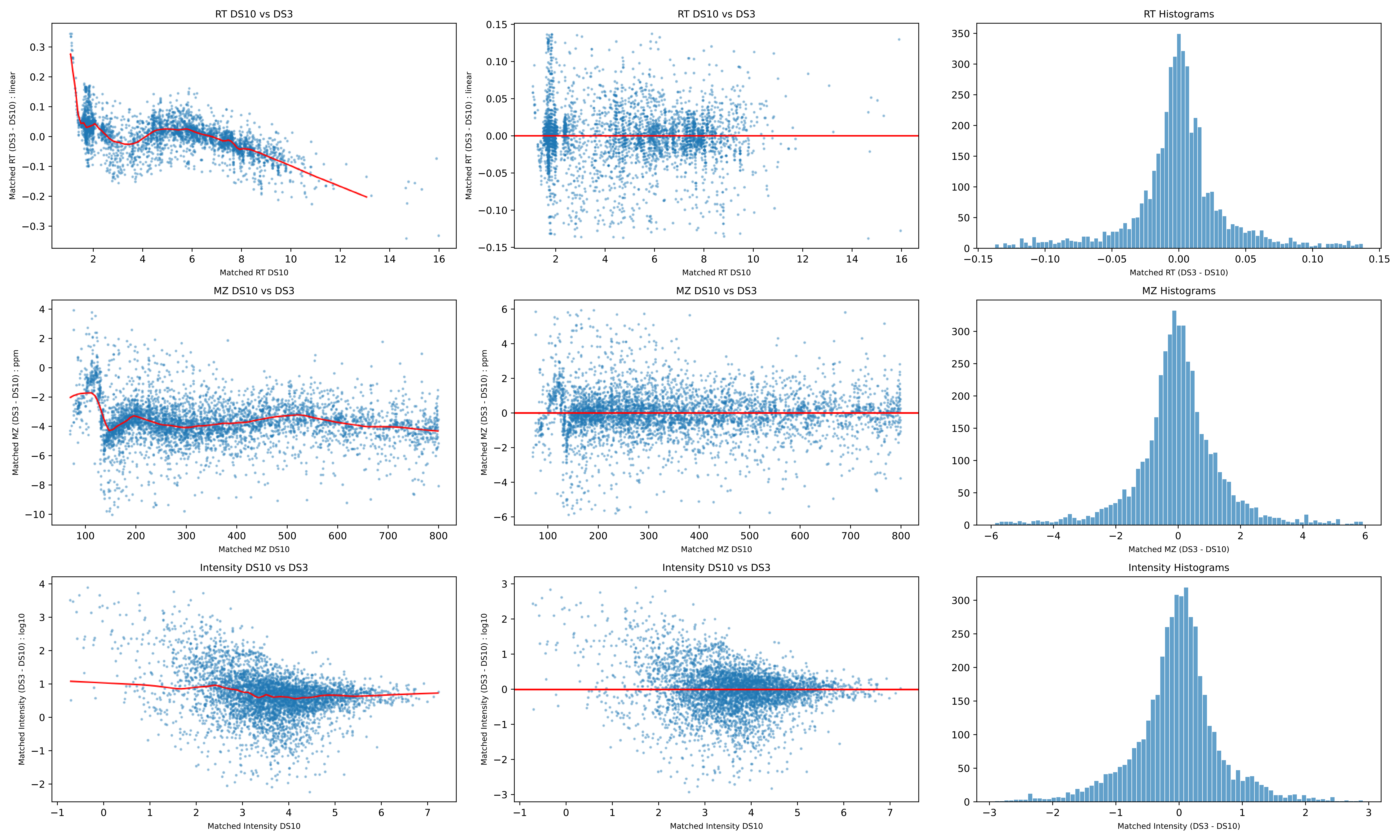
RT DS10 vs DS3
Matched RT (DS3 - DS10) : linear
Matched RT DS10
RT DS10 vs DS3
Matched RT (DS3 - DS10) : linear
Matched RT DS10
RT Histograms
Matched RT (DS3 - DS10)
MZ DS10 vs DS3
Matched MZ (DS3 - DS10) : ppm
Matched MZ DS10
MZ DS10 vs DS3
Matched MZ (DS3 - DS10) : ppm
Matched MZ DS10
MZ Histograms
Matched MZ (DS3 - DS10)
Intensity DS10 vs DS3
Matched Intensity (DS3 - DS10) : log10
Matched Intensity DS10
Intensity DS10 vs DS3
Matched Intensity (DS3 - DS10) : log10
Matched Intensity DS10
Intensity Histograms
Matched Intensity (DS3 - DS10)

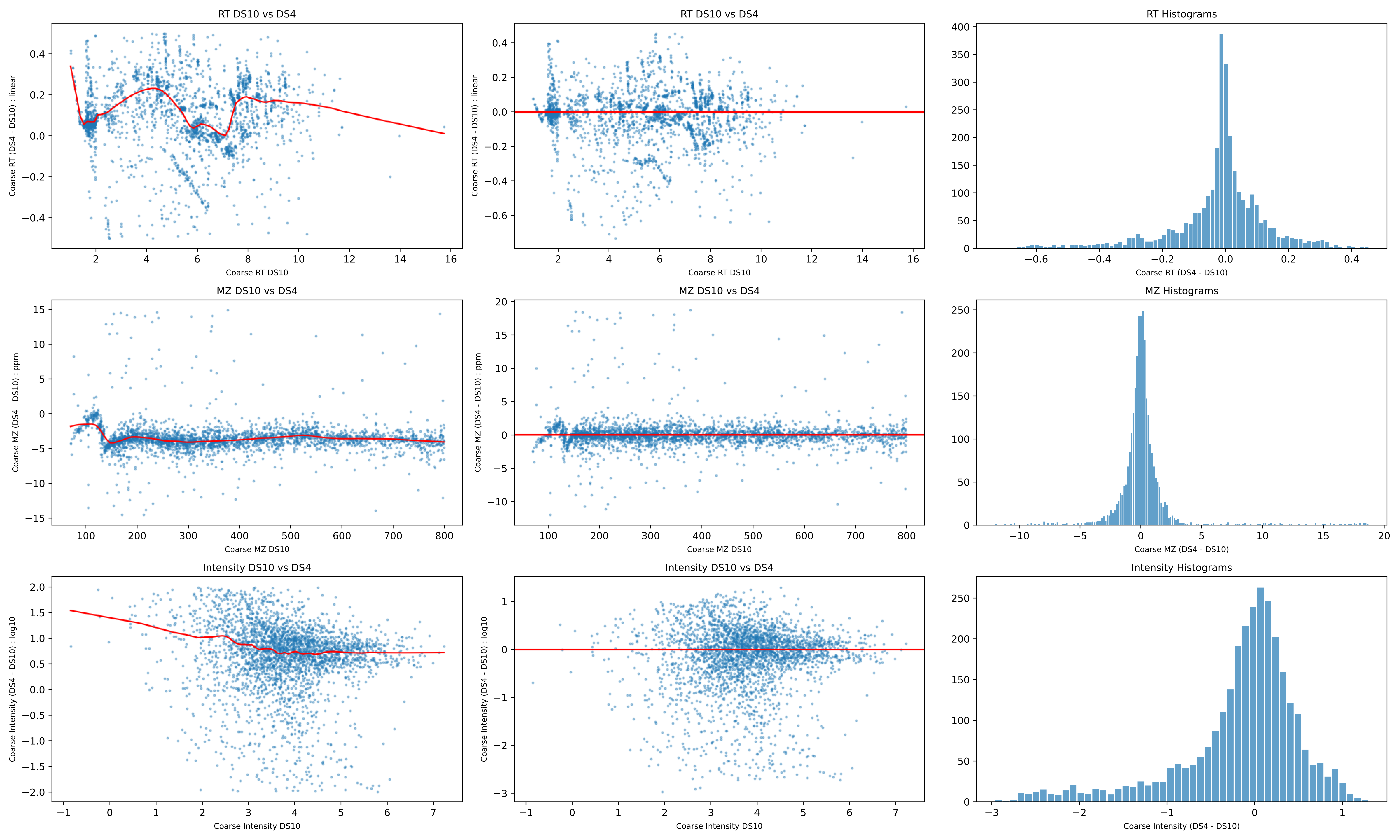
RT DS10 vs DS4
Coarse RT (DS4 - DS10) : linear
Coarse RT DS10
RT DS10 vs DS4
Coarse RT (DS4 - DS10) : linear
Coarse RT DS10
RT Histograms
Coarse RT (DS4 - DS10)
MZ DS10 vs DS4
Coarse MZ (DS4 - DS10) : ppm
Coarse MZ DS10
MZ DS10 vs DS4
Coarse MZ (DS4 - DS10) : ppm
Coarse MZ DS10
MZ Histograms
Coarse MZ (DS4 - DS10)
Intensity DS10 vs DS4
Coarse Intensity (DS4 - DS10) : log10
Coarse Intensity DS10
Intensity DS10 vs DS4
Coarse Intensity (DS4 - DS10) : log10
Coarse Intensity DS10
Intensity Histograms
Coarse Intensity (DS4 - DS10)

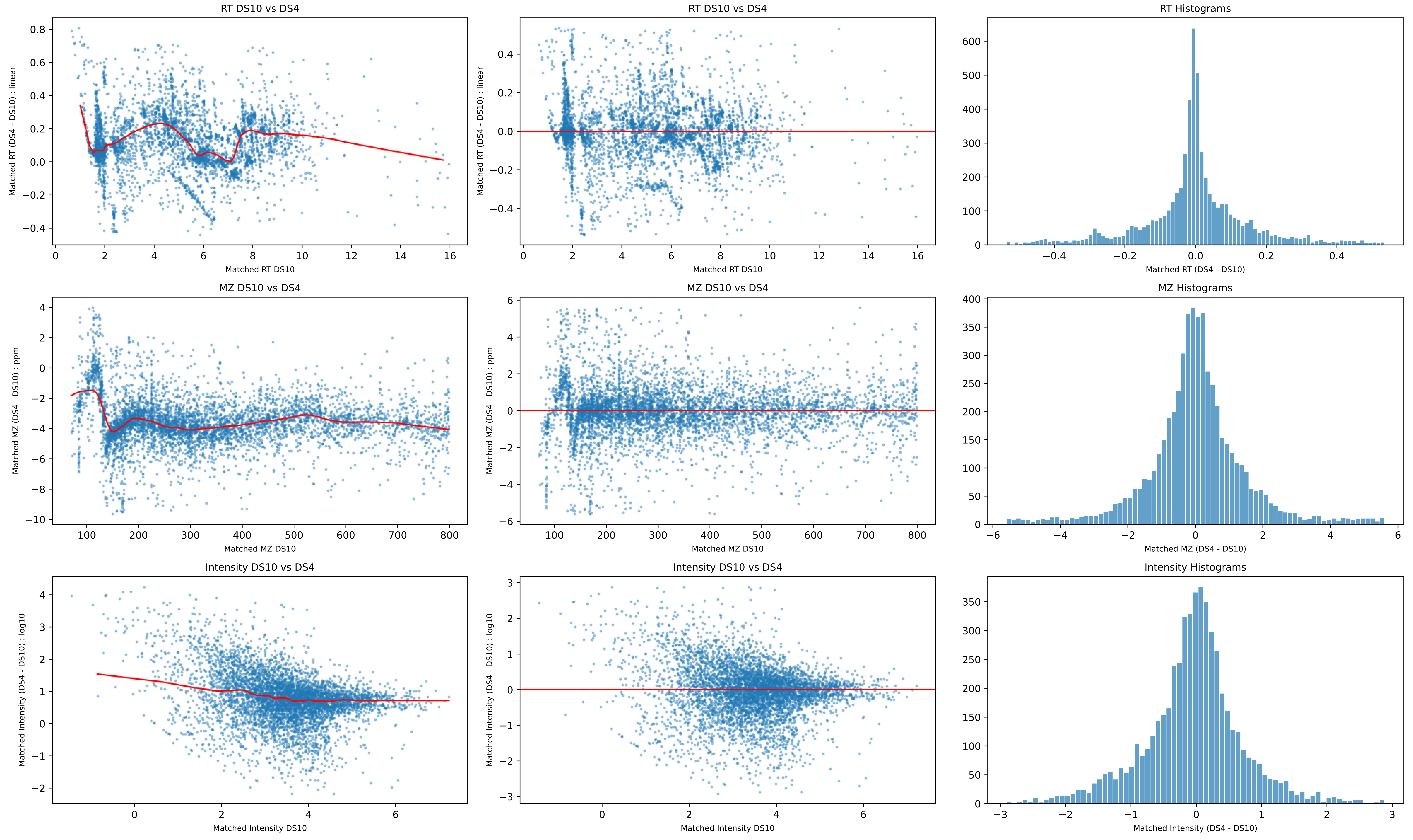
RT DS10 vs DS4
Matched RT (DS4 - DS10) : linear
Matched RT DS10
RT DS10 vs DS4
Matched RT (DS4 - DS10) : linear
Matched RT DS10
RT Histograms
Matched RT (DS4 - DS10)
MZ DS10 vs DS4
Matched MZ (DS4 - DS10) : ppm
Matched MZ DS10
MZ DS10 vs DS4
Matched MZ (DS4 - DS10) : ppm
Matched MZ DS10
MZ Histograms
Matched MZ (DS4 - DS10)
Intensity DS10 vs DS4
Matched Intensity (DS4 - DS10) : log10
Matched Intensity DS10
Intensity DS10 vs DS4
Matched Intensity (DS4 - DS10) : log10
Matched Intensity DS10
Intensity Histograms
Matched Intensity (DS4 - DS10)

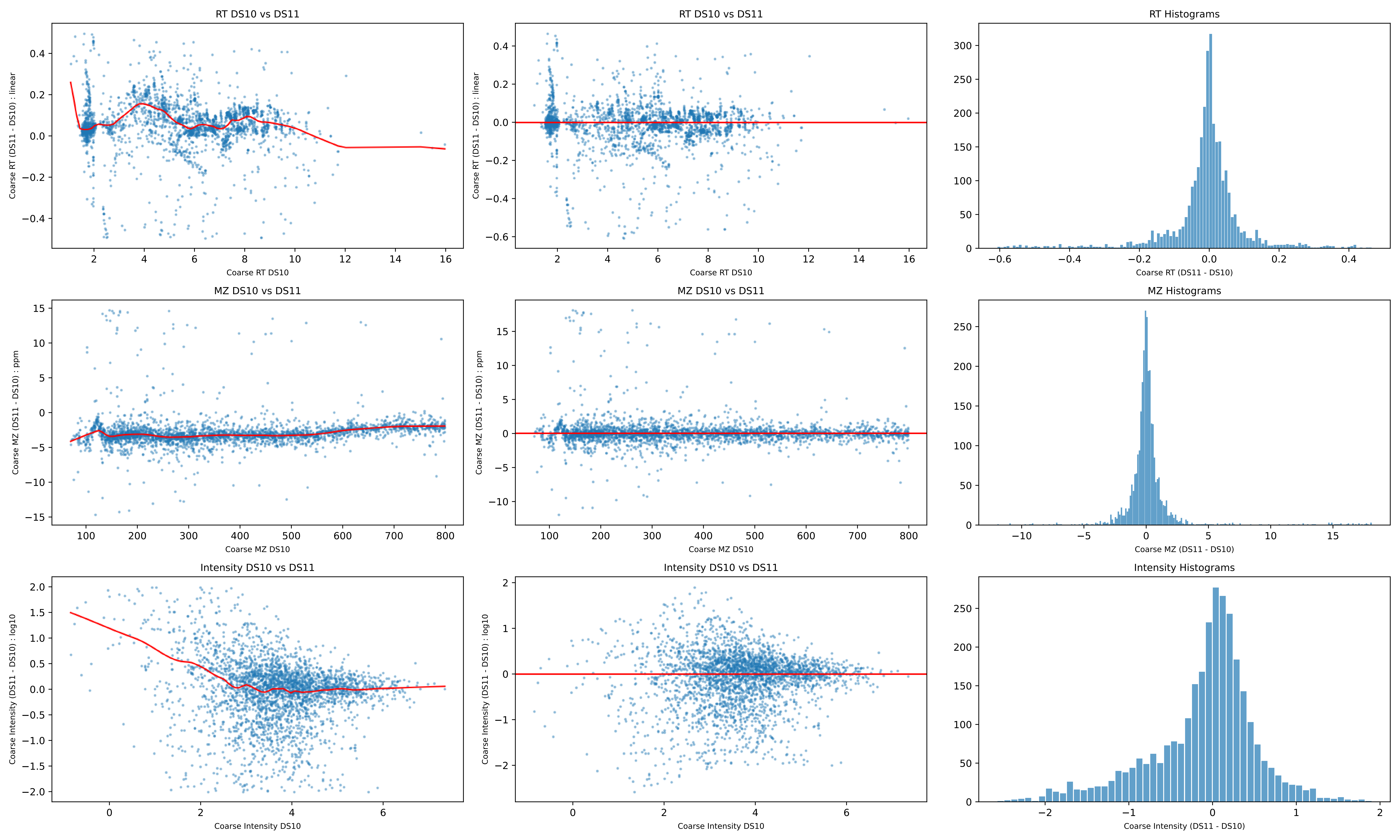
RT DS10 vs DS11
Coarse RT (DS11 - DS10) : linear
Coarse RT DS10
RT DS10 vs DS11
Coarse RT (DS11 - DS10) : linear
Coarse RT DS10
RT Histograms
Coarse RT (DS11 - DS10)
MZ DS10 vs DS11
Coarse MZ (DS11 - DS10) : ppm
Coarse MZ DS10
MZ DS10 vs DS11
Coarse MZ (DS11 - DS10) : ppm
Coarse MZ DS10
MZ Histograms
Coarse MZ (DS11 - DS10)
Intensity DS10 vs DS11
Coarse Intensity (DS11 - DS10) : log10
Coarse Intensity DS10
Intensity DS10 vs DS11
Coarse Intensity (DS11 - DS10) : log10
Coarse Intensity DS10
Intensity Histograms
Coarse Intensity (DS11 - DS10)

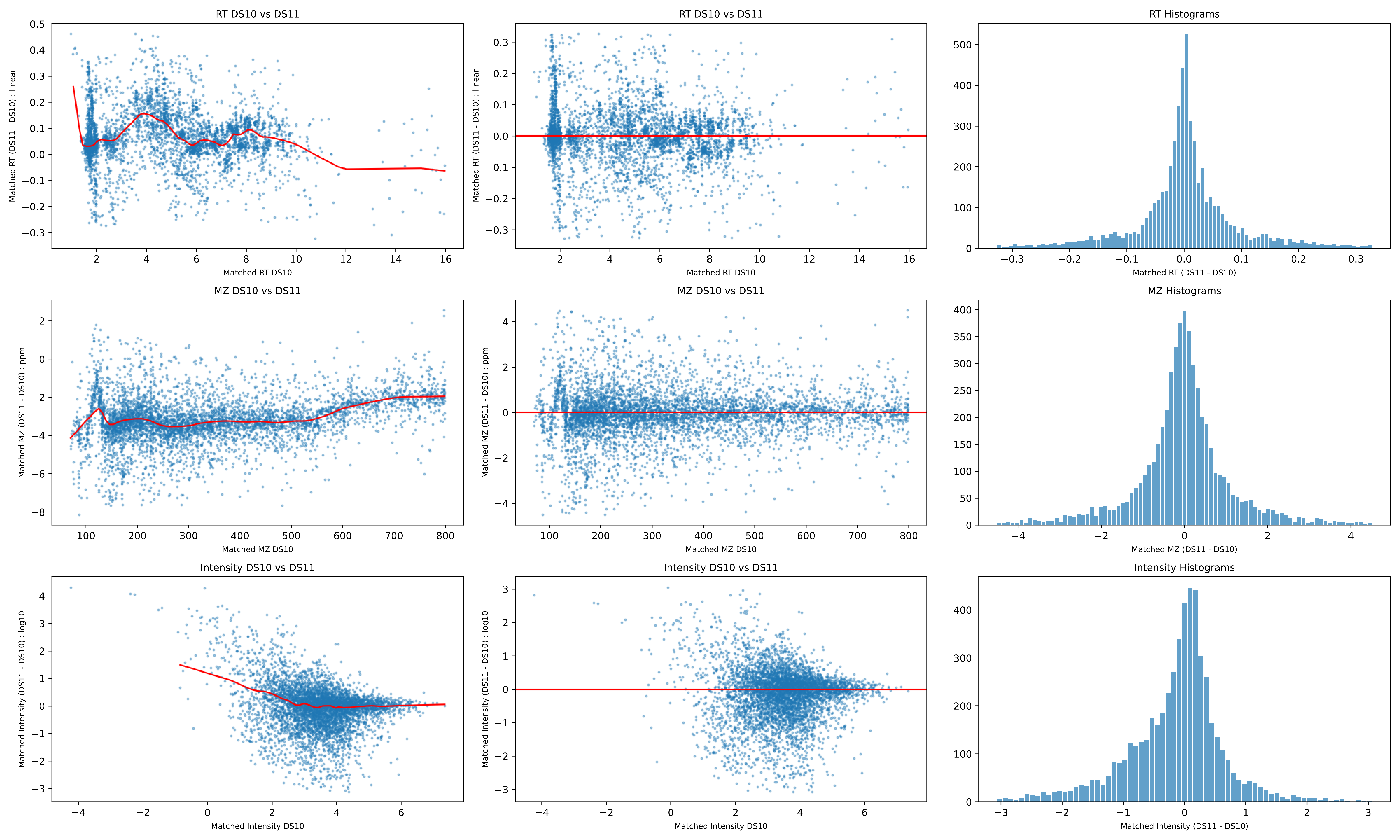
RT DS10 vs DS11
Matched RT (DS11 - DS10) : linear
Matched RT DS10
RT DS10 vs DS11
Matched RT (DS11 - DS10) : linear
Matched RT DS10
RT Histograms
Matched RT (DS11 - DS10)
MZ DS10 vs DS11
Matched MZ (DS11 - DS10) : ppm
Matched MZ DS10
MZ DS10 vs DS11
Matched MZ (DS11 - DS10) : ppm
Matched MZ DS10
MZ Histograms
Matched MZ (DS11 - DS10)
Intensity DS10 vs DS11
Matched Intensity (DS11 - DS10) : log10
Matched Intensity DS10
Intensity DS10 vs DS11
Matched Intensity (DS11 - DS10) : log10
Matched Intensity DS10
Intensity Histograms
Matched Intensity (DS11 - DS10)

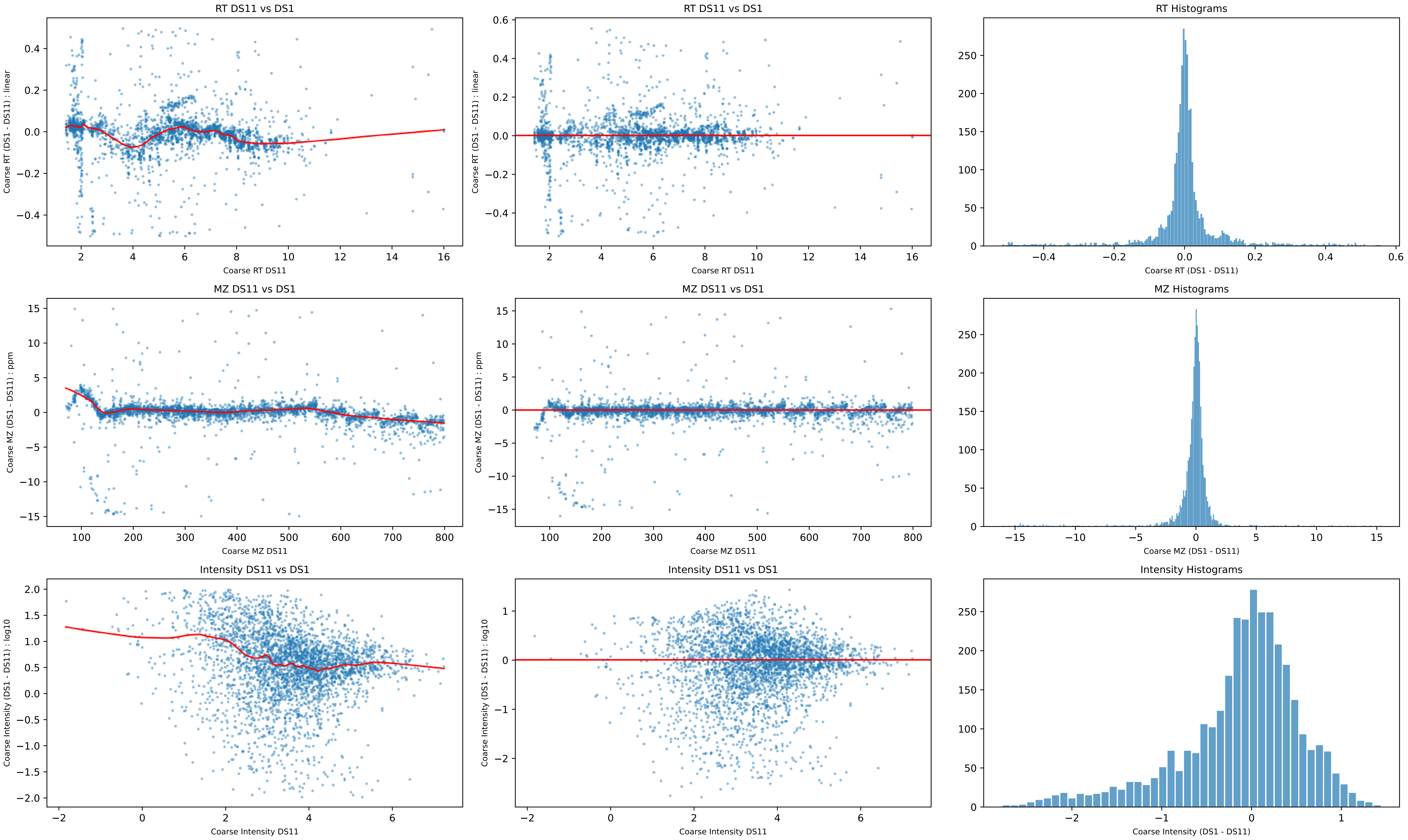
RT DS11 vs DS1
Coarse RT (DS1 - DS11) : linear
Coarse RT DS11
RT DS11 vs DS1
Coarse RT (DS1 - DS11) : linear
Coarse RT DS11
RT Histograms
Coarse RT (DS1 - DS11)
MZ DS11 vs DS1
Coarse MZ (DS1 - DS11) : ppm
Coarse MZ DS11
MZ DS11 vs DS1
Coarse MZ (DS1 - DS11) : ppm
Coarse MZ DS11
MZ Histograms
Coarse MZ (DS1 - DS11)
Intensity DS11 vs DS1
Coarse Intensity (DS1 - DS11) : log10
Coarse Intensity DS11
Intensity DS11 vs DS1
Coarse Intensity (DS1 - DS11) : log10
Coarse Intensity DS11
Intensity Histograms
Coarse Intensity (DS1 - DS11)

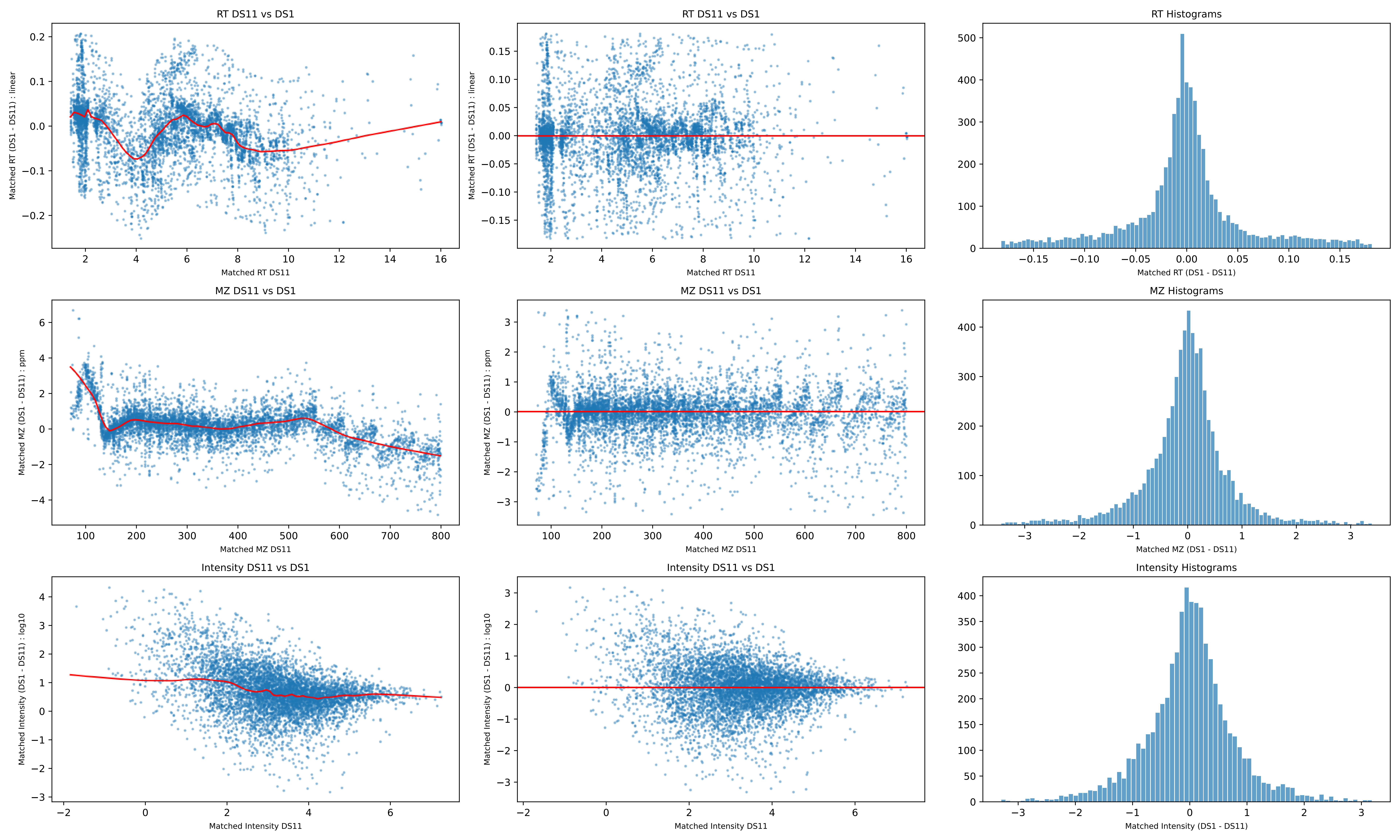

RT DS11 vs DS1
Matched RT (DS1 - DS11) : linear
Matched RT DS11
RT DS11 vs DS1
Matched RT (DS1 - DS11) : linear
Matched RT DS11
RT Histograms
Matched RT (DS1 - DS11)
MZ DS11 vs DS1
Matched MZ (DS1 - DS11) : ppm
Matched MZ DS11
MZ DS11 vs DS1
Matched MZ (DS1 - DS11) : ppm
Matched MZ DS11
MZ Histograms
Matched MZ (DS1 - DS11)
Intensity DS11 vs DS1
Matched Intensity (DS1 - DS11) : log10
Matched Intensity DS11
Intensity DS11 vs DS1
Matched Intensity (DS1 - DS11) : log10
Matched Intensity DS11
Intensity Histograms
Matched Intensity (DS1 - DS11)

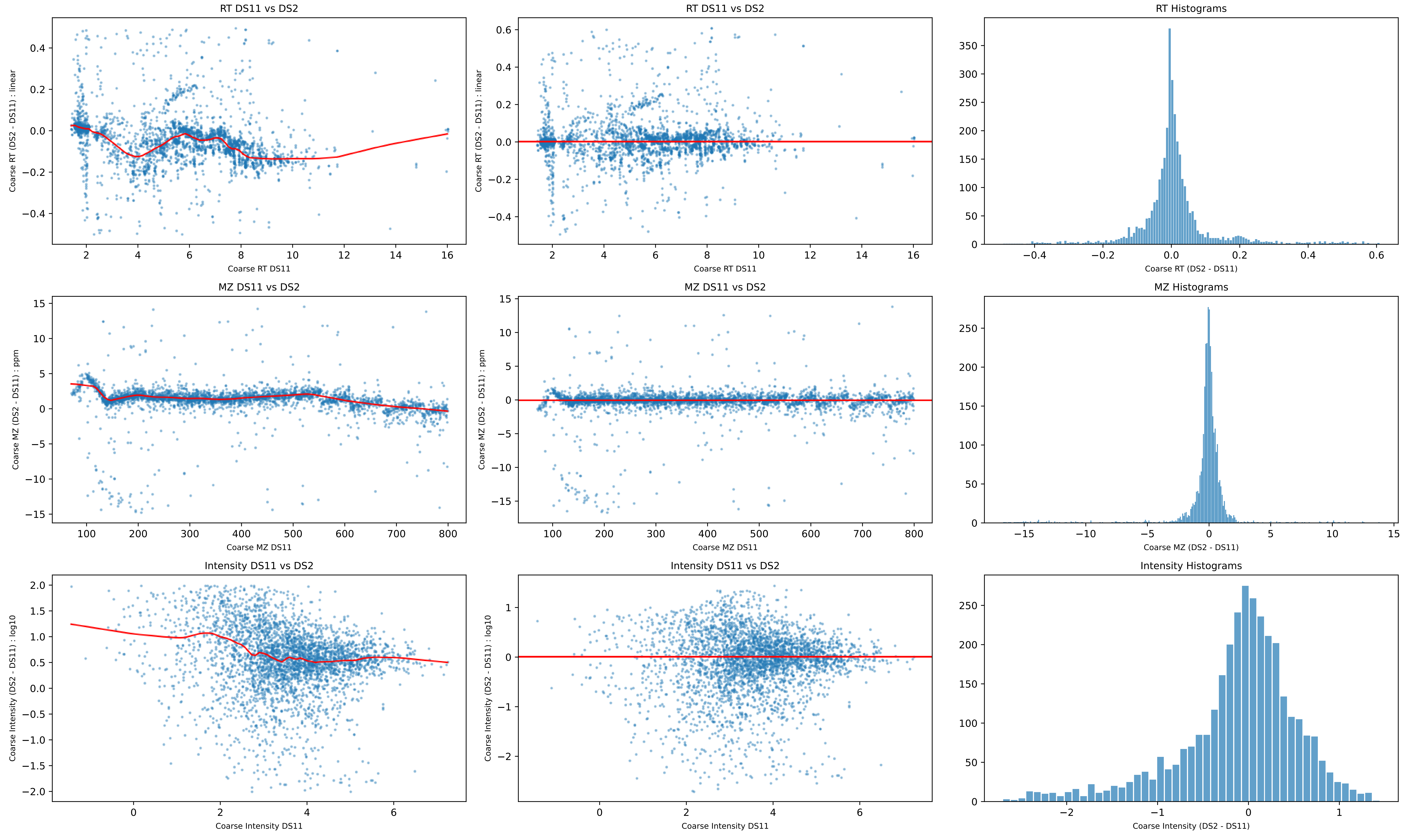
RT DS11 vs DS2
Coarse RT (DS2 - DS11) : linear
Coarse RT DS11
RT DS11 vs DS2
Coarse RT (DS2 - DS11) : linear
Coarse RT DS11
RT Histograms
Coarse RT (DS2 - DS11)
MZ DS11 vs DS2
Coarse MZ (DS2 - DS11) : ppm
Coarse MZ DS11
MZ DS11 vs DS2
Coarse MZ (DS2 - DS11) : ppm
Coarse MZ DS11
MZ Histograms
Coarse MZ (DS2 - DS11)
Intensity DS11 vs DS2
Coarse Intensity (DS2 - DS11) : log10
Coarse Intensity DS11
Intensity DS11 vs DS2
Coarse Intensity (DS2 - DS11) : log10
Coarse Intensity DS11
Intensity Histograms
Coarse Intensity (DS2 - DS11)

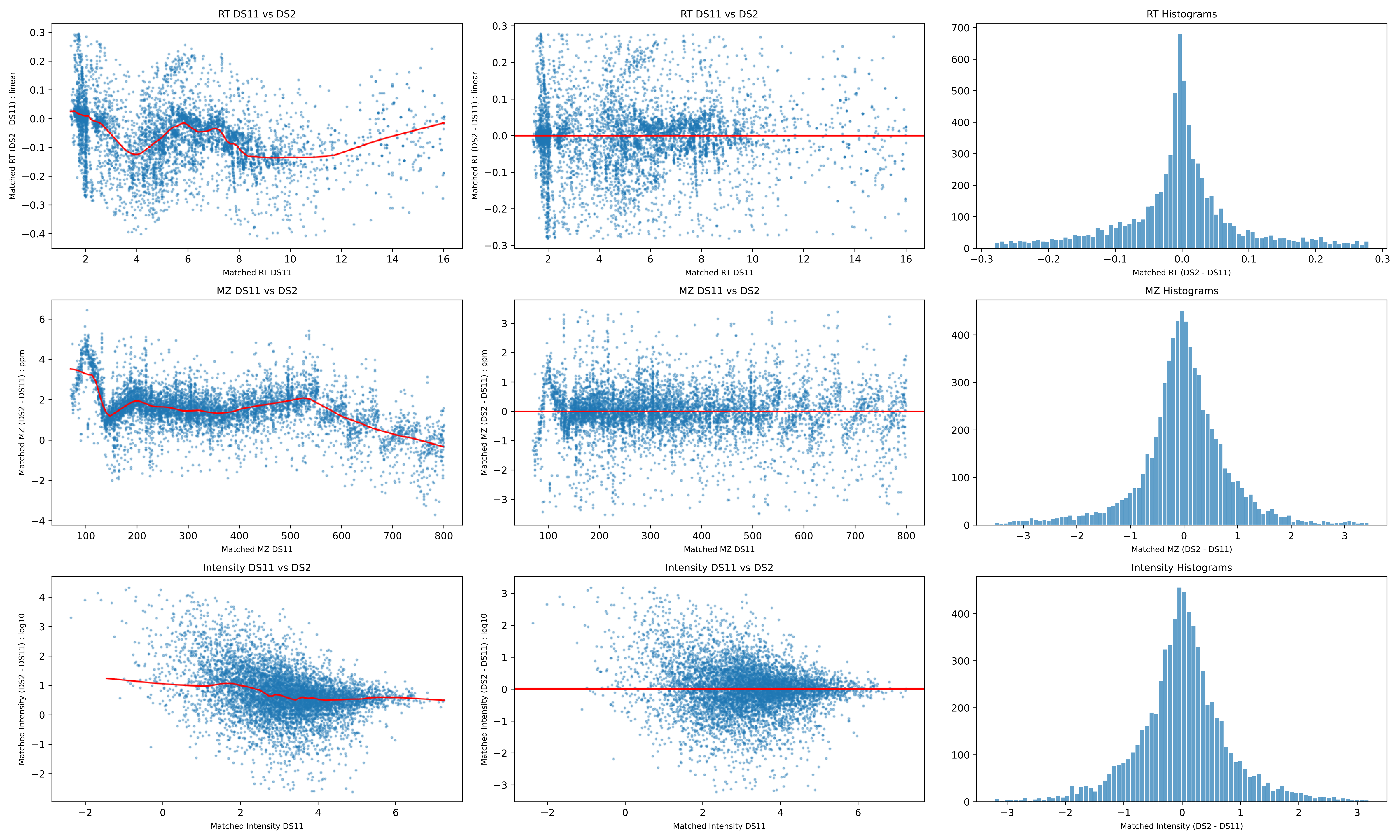
RT DS11 vs DS2
Matched RT (DS2 - DS11) : linear
Matched RT DS11
RT DS11 vs DS2
Matched RT (DS2 - DS11) : linear
Matched RT DS11
RT Histograms
Matched RT (DS2 - DS11)
MZ DS11 vs DS2
Matched MZ (DS2 - DS11) : ppm
Matched MZ DS11
MZ DS11 vs DS2
Matched MZ (DS2 - DS11) : ppm
Matched MZ DS11
MZ Histograms
Matched MZ (DS2 - DS11)
Intensity DS11 vs DS2
Matched Intensity (DS2 - DS11) : log10
Matched Intensity DS11
Intensity DS11 vs DS2
Matched Intensity (DS2 - DS11) : log10
Matched Intensity DS11
Intensity Histograms
Matched Intensity (DS2 - DS11)

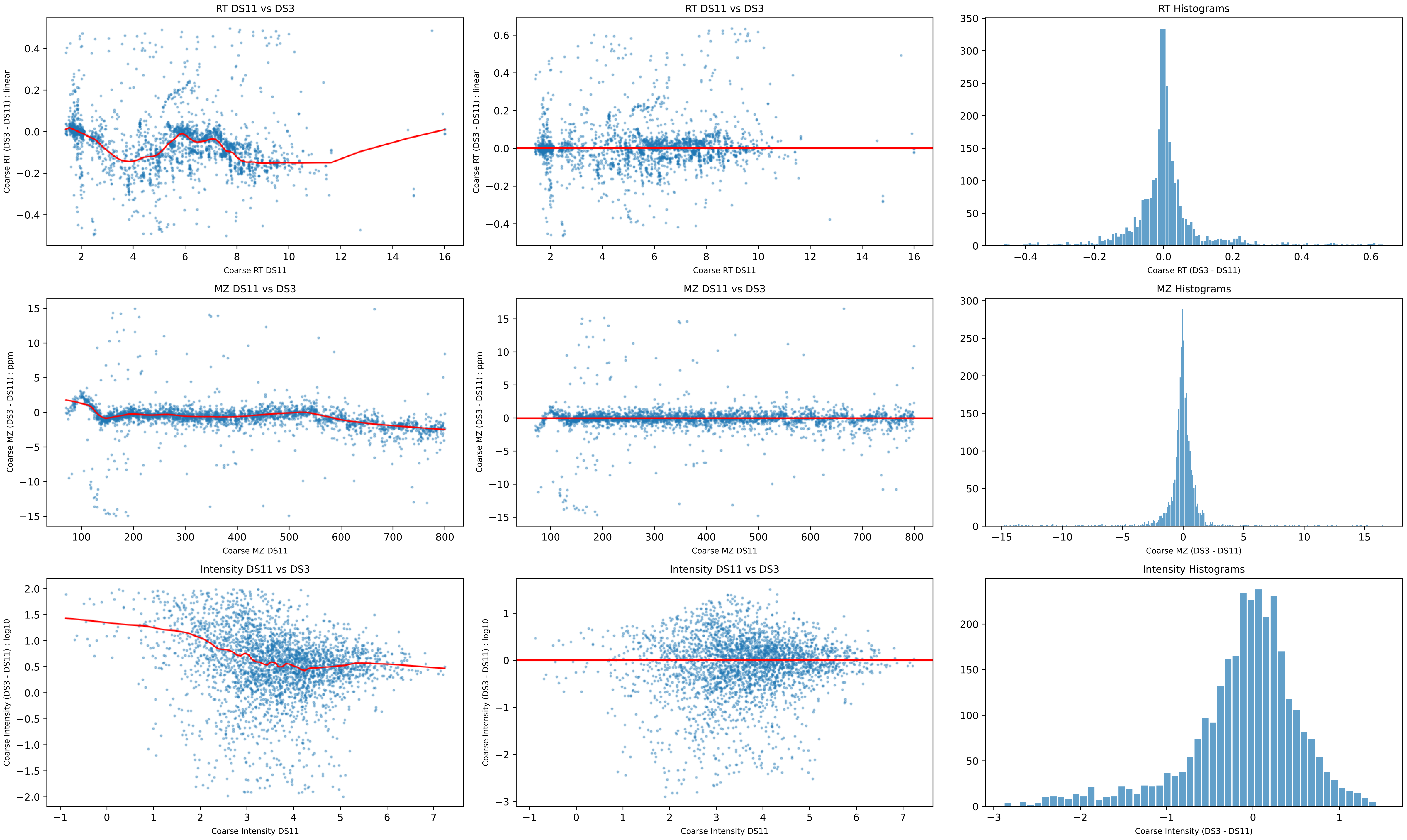
RT DS11 vs DS3
Coarse RT (DS3 - DS11) : linear
Coarse RT DS11
RT DS11 vs DS3
Coarse RT (DS3 - DS11) : linear
Coarse RT DS11
RT Histograms
Coarse RT (DS3 - DS11)
MZ DS11 vs DS3
Coarse MZ (DS3 - DS11) : ppm
Coarse MZ DS11
MZ DS11 vs DS3
Coarse MZ (DS3 - DS11) : ppm
Coarse MZ DS11
MZ Histograms
Coarse MZ (DS3 - DS11)
Intensity DS11 vs DS3
Coarse Intensity (DS3 - DS11) : log10
Coarse Intensity DS11
Intensity DS11 vs DS3
Coarse Intensity (DS3 - DS11) : log10
Coarse Intensity DS11
Intensity Histograms
Coarse Intensity (DS3 - DS11)

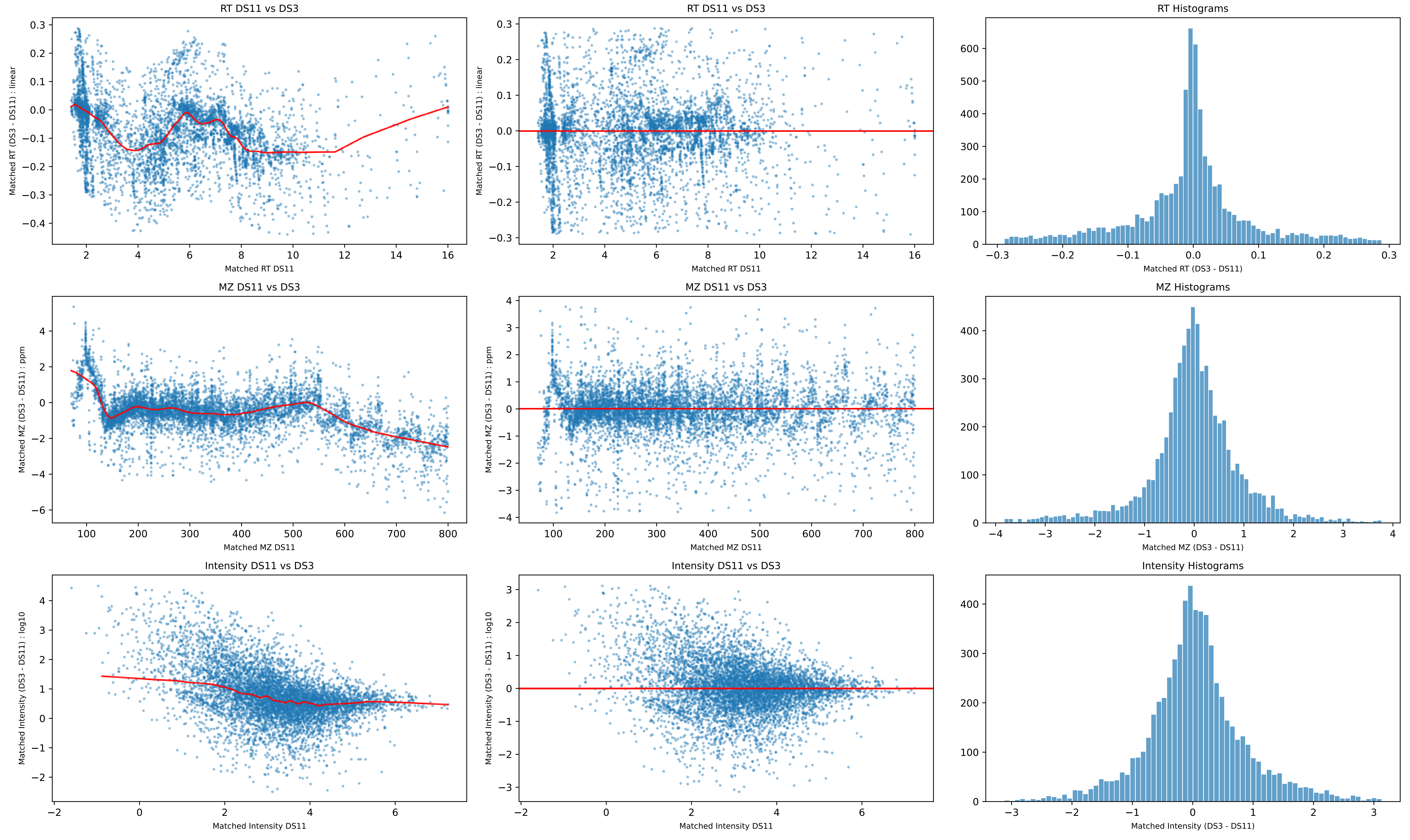
RT DS11 vs DS3
Matched RT (DS3 - DS11) : linear
Matched RT DS11
RT Histograms
Matched RT (DS3 - DS11)
MZ DS11 vs DS3
Matched MZ (DS3 - DS11) : ppm
Matched MZ DS11
MZ Histograms
Matched MZ (DS3 - DS11)
Intensity DS11 vs DS3
Matched Intensity (DS3 - DS11) : log10
Matched Intensity DS11
Intensity Histograms
Matched Intensity (DS3 - DS11)

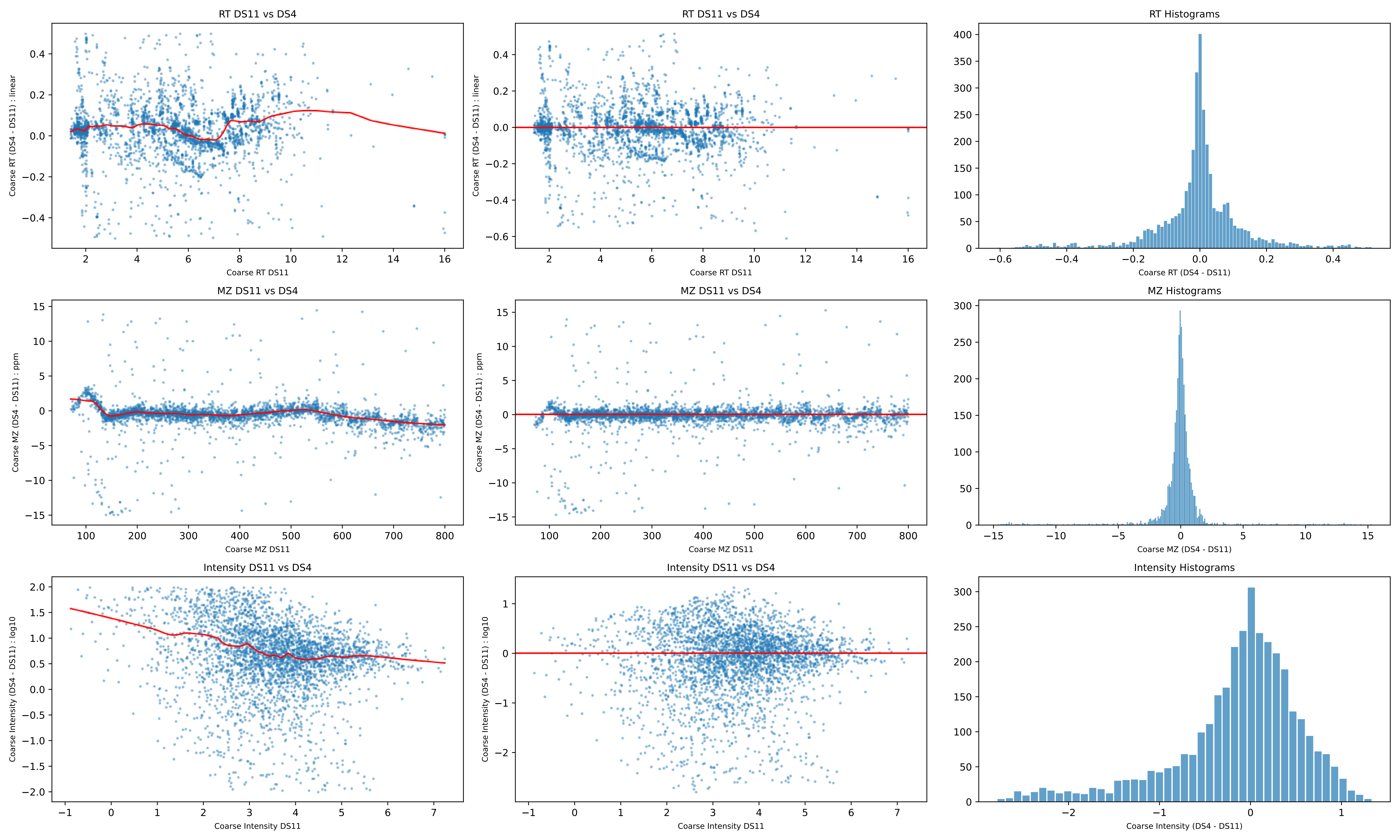

RT DS11 vs DS4
Coarse RT (DS4 - DS11) : linear
Coarse RT DS11
RT DS11 vs DS4
Coarse RT (DS4 - DS11) : linear
Coarse RT DS11
RT Histograms
Coarse RT (DS4 - DS11)
MZ DS11 vs DS4
Coarse MZ (DS4 - DS11) : ppm
Coarse MZ DS11
MZ DS11 vs DS4
Coarse MZ (DS4 - DS11) : ppm
Coarse MZ DS11
MZ Histograms
Coarse MZ (DS4 - DS11)
Intensity DS11 vs DS4
Coarse Intensity (DS4 - DS11) : log10
Coarse Intensity DS11
Intensity DS11 vs DS4
Coarse Intensity (DS4 - DS11) : log10
Coarse Intensity DS11
Intensity Histograms
Coarse Intensity (DS4 - DS11)

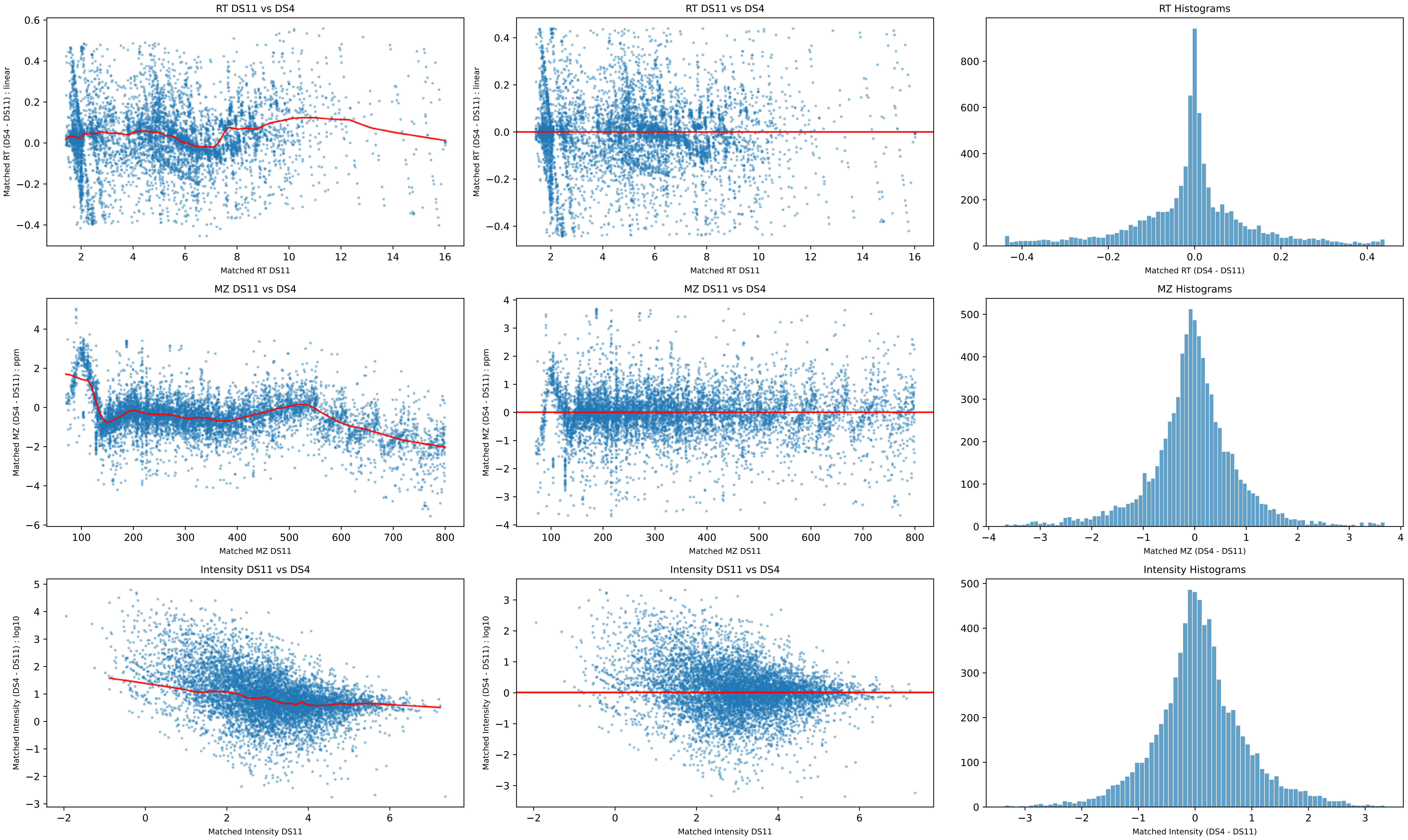

RT DS11 vs DS4
Matched RT (DS4 - DS11) : linear
Matched RT DS11
RT DS11 vs DS4
Matched RT (DS4 - DS11) : linear
Matched RT DS11
RT Histograms
Matched RT (DS4 - DS11)
MZ DS11 vs DS4
Matched MZ (DS4 - DS11) : ppm
Matched MZ DS11
MZ DS11 vs DS4
Matched MZ (DS4 - DS11) : ppm
Matched MZ DS11
MZ Histograms
Matched MZ (DS4 - DS11)
Intensity DS11 vs DS4
Matched Intensity (DS4 - DS11) : log10
Matched Intensity DS11
Intensity DS11 vs DS4
Matched Intensity (DS4 - DS11) : log10
Matched Intensity DS11
Intensity Histograms
Matched Intensity (DS4 - DS11)

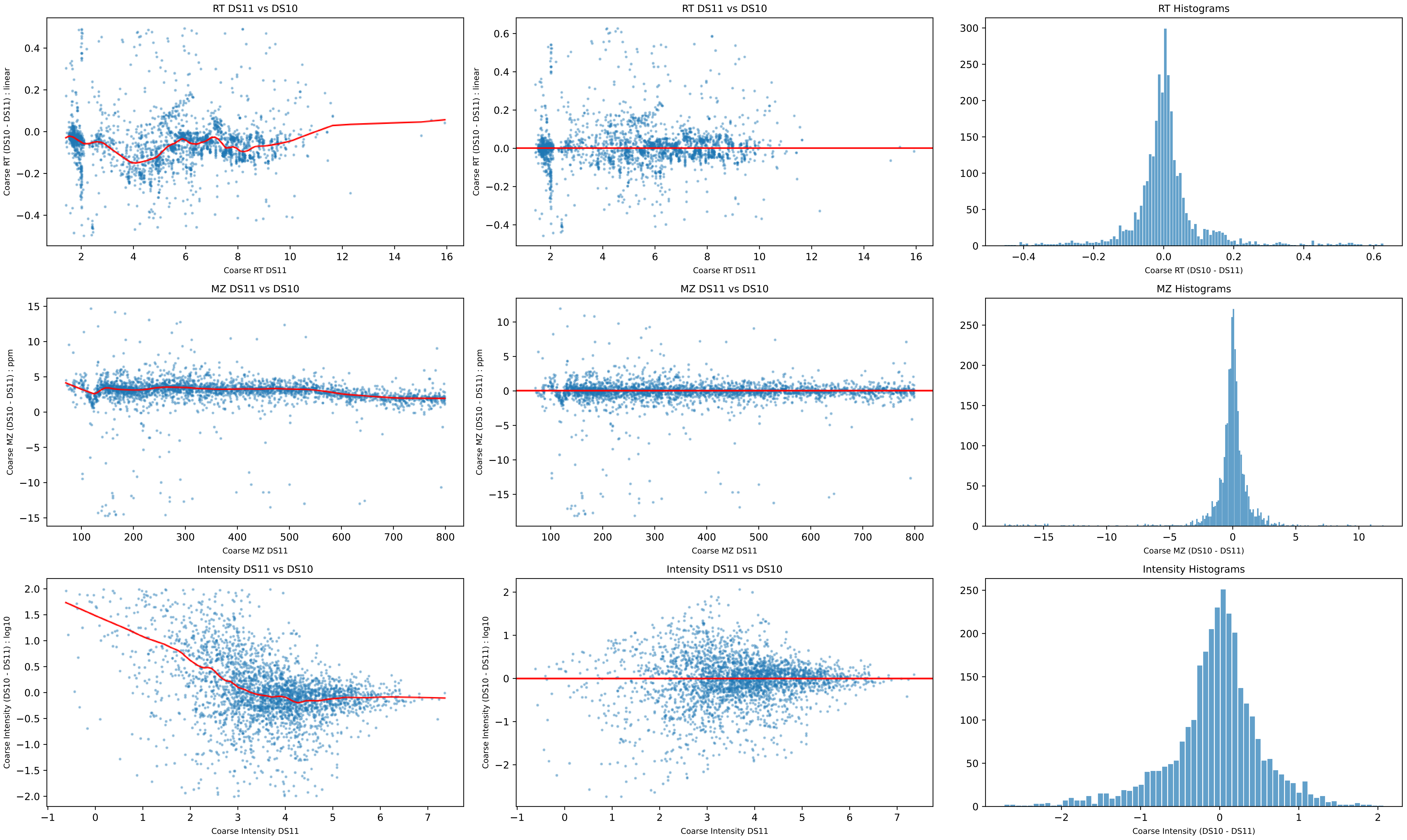
RT DS11 vs DS10
Coarse RT (DS10 - DS11) : linear
Coarse RT DS11
RT DS11 vs DS10
Coarse RT (DS10 - DS11) : linear
Coarse RT DS11
RT Histograms
Coarse RT (DS10 - DS11)
MZ DS11 vs DS10
Coarse MZ (DS10 - DS11) : ppm
Coarse MZ DS11
MZ DS11 vs DS10
Coarse MZ (DS10 - DS11) : ppm
Coarse MZ DS11
MZ Histograms
Coarse MZ (DS10 - DS11)
Intensity DS11 vs DS10
Coarse Intensity (DS10 - DS11) : log10
Coarse Intensity DS11
Intensity DS11 vs DS10
Coarse Intensity (DS10 - DS11) : log10
Coarse Intensity DS11
Intensity Histograms
Coarse Intensity (DS10 - DS11)

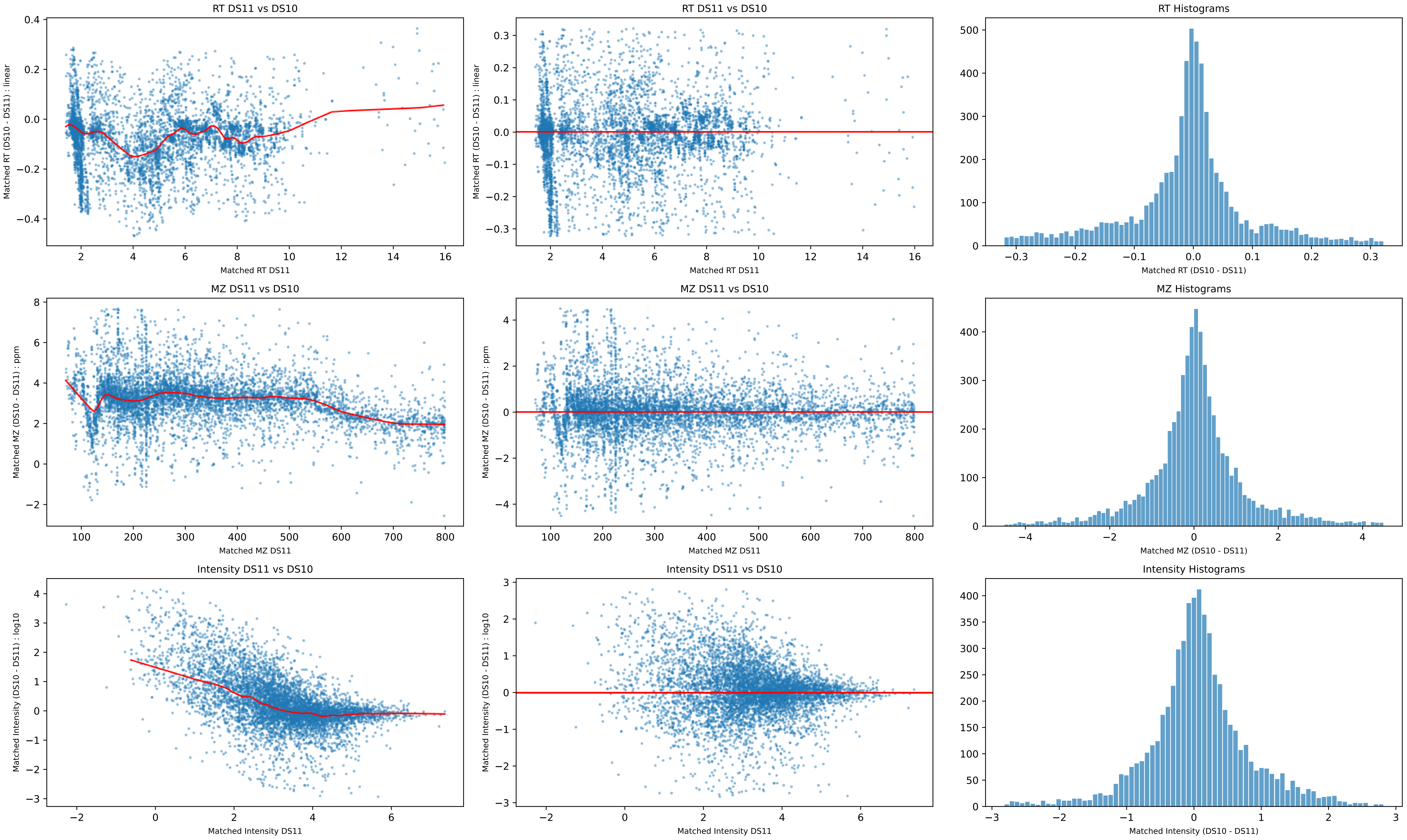
RT DS11 vs DS10
Matched RT (DS10 - DS11) : linear
Matched RT DS11
RT DS11 vs DS10
Matched RT (DS10 - DS11) : linear
Matched RT DS11
RT Histograms
Matched RT (DS10 - DS11)
MZ DS11 vs DS10
Matched MZ (DS10 - DS11) : ppm
Matched MZ DS11
MZ DS11 vs DS10
Matched MZ (DS10 - DS11) : ppm
Matched MZ DS11
MZ Histograms
Matched MZ (DS10 - DS11)
Intensity DS11 vs DS10
Matched Intensity (DS10 - DS11) : log10
Matched Intensity DS11
Intensity DS11 vs DS10
Matched Intensity (DS10 - DS11) : log10
Matched Intensity DS11
Intensity Histograms
Matched Intensity (DS10 - DS11)
