## Supplementary Reports and Code for "*Eclipse*: A Python package for alignment of two or more nontargeted LC-MS metabolomics datasets": Supplementary Report 2.pdf

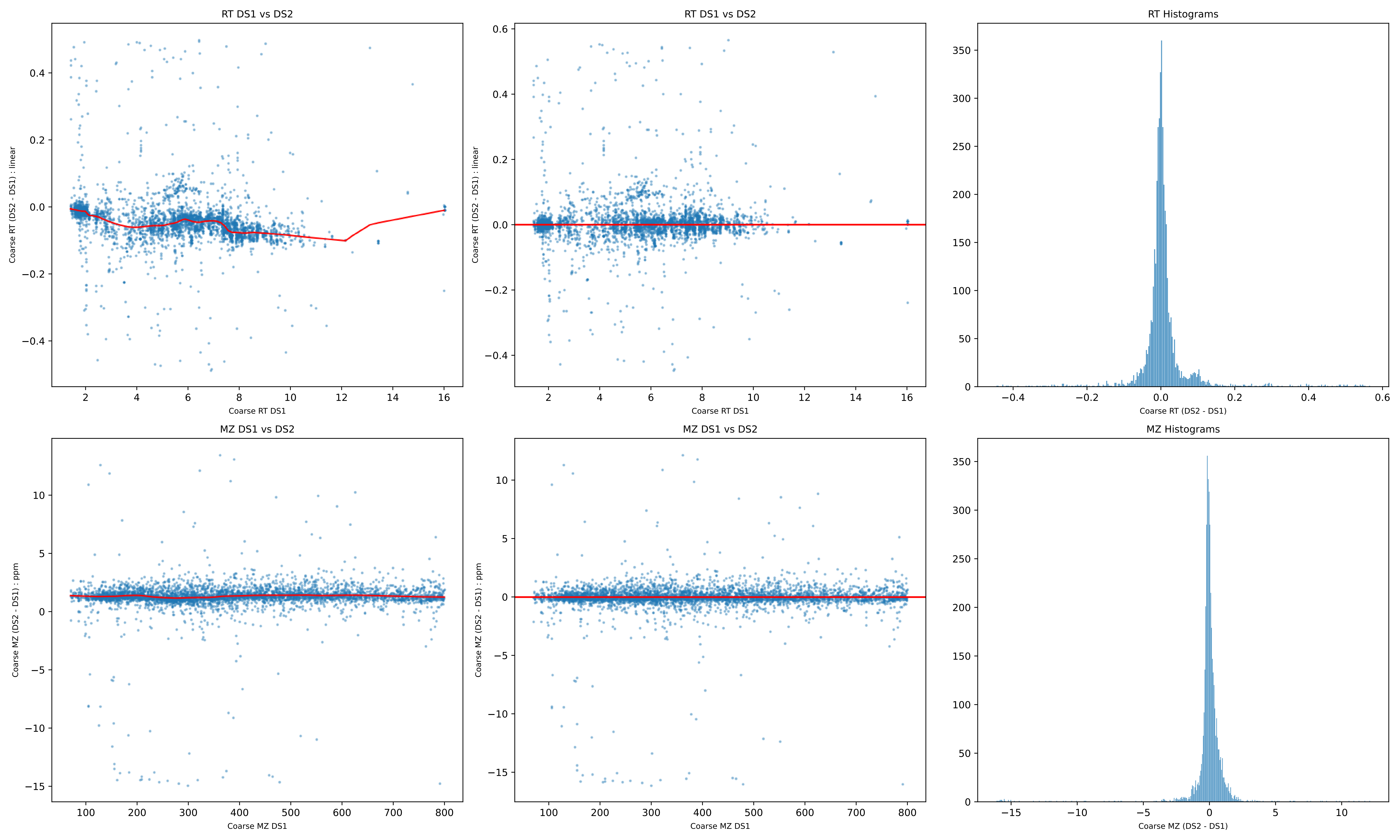
RT DS1 vs DS2
Coarse RT (DS2 - DS1) : linear
Coarse RT DS1
RT DS1 vs DS2
Coarse RT (DS2 - DS1) : linear
Coarse RT DS1
RT Histograms
Coarse RT (DS2 - DS1)
MZ DS1 vs DS2
Coarse MZ (DS2 - DS1) : ppm
Coarse MZ DS1
MZ DS1 vs DS2
Coarse MZ (DS2 - DS1) : ppm
Coarse MZ DS1
MZ Histograms
Coarse MZ (DS2 - DS1)

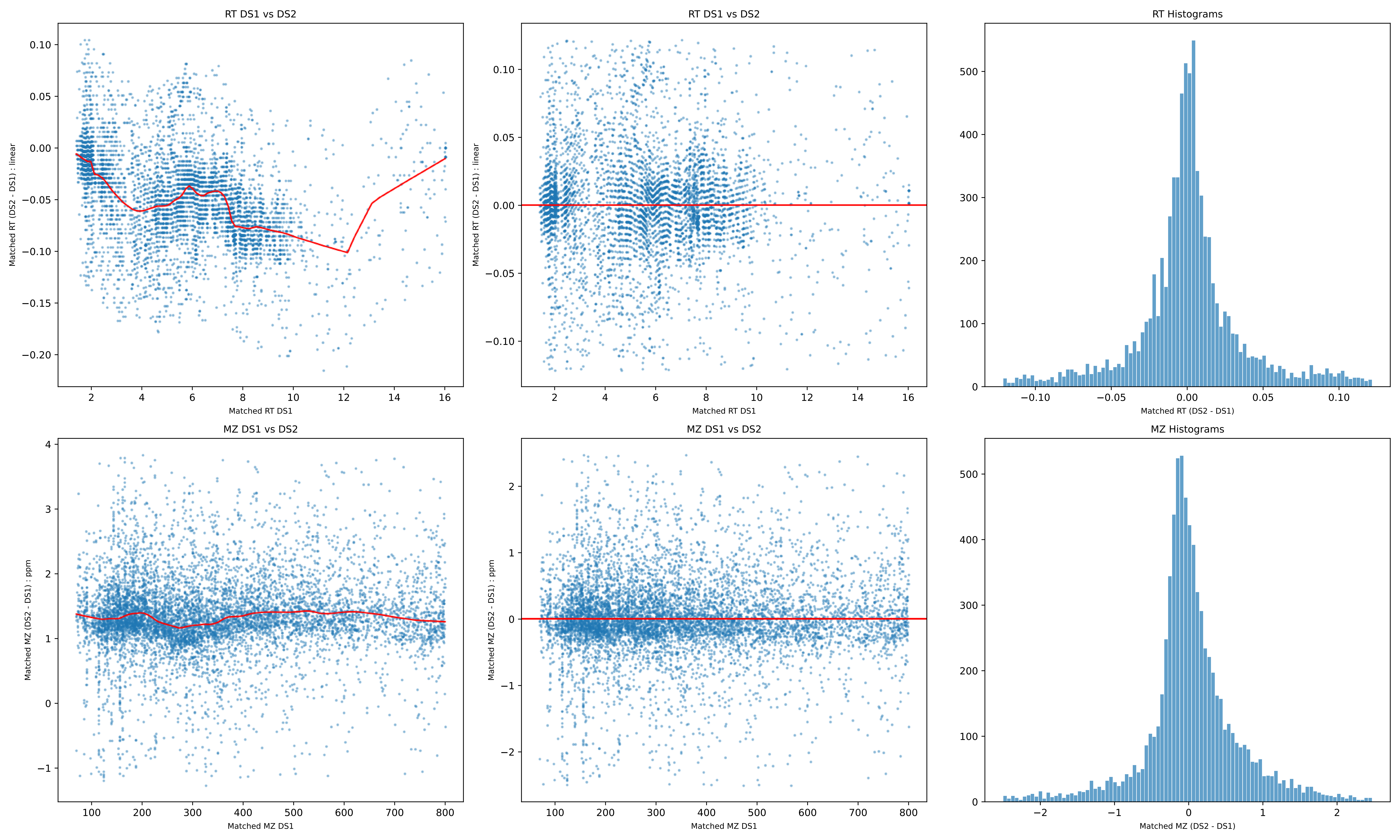
RT DS1 vs DS2
Matched RT (DS2 - DS1) : linear
Matched RT DS1
RT DS1 vs DS2
Matched RT (DS2 - DS1) : linear
Matched RT DS1
RT Histograms
Matched RT (DS2 - DS1)
MZ DS1 vs DS2
Matched MZ (DS2 - DS1) : ppm
Matched MZ DS1
MZ DS1 vs DS2
Matched MZ (DS2 - DS1) : ppm
Matched MZ DS1
MZ Histograms
Matched MZ (DS2 - DS1)

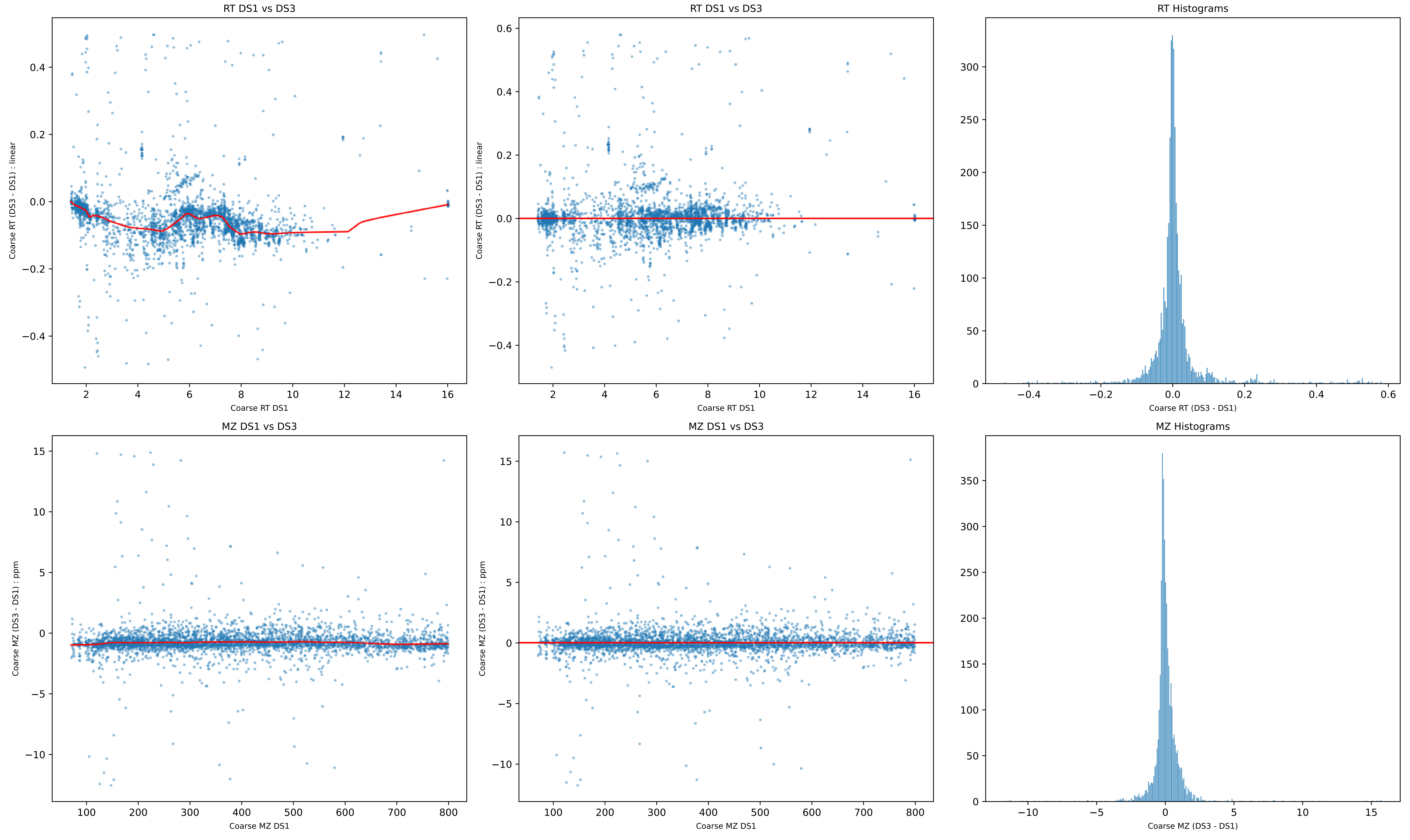

RT DS1 vs DS3
Coarse RT (DS3 - DS1) : linear
Coarse RT DS1
RT DS1 vs DS3
Coarse RT (DS3 - DS1) : linear
Coarse RT DS1
RT Histograms
Coarse RT (DS3 - DS1)
MZ DS1 vs DS3
Coarse MZ (DS3 - DS1) : ppm
Coarse MZ DS1
MZ DS1 vs DS3
Coarse MZ (DS3 - DS1) : ppm
Coarse MZ DS1
MZ Histograms
Coarse MZ (DS3 - DS1)

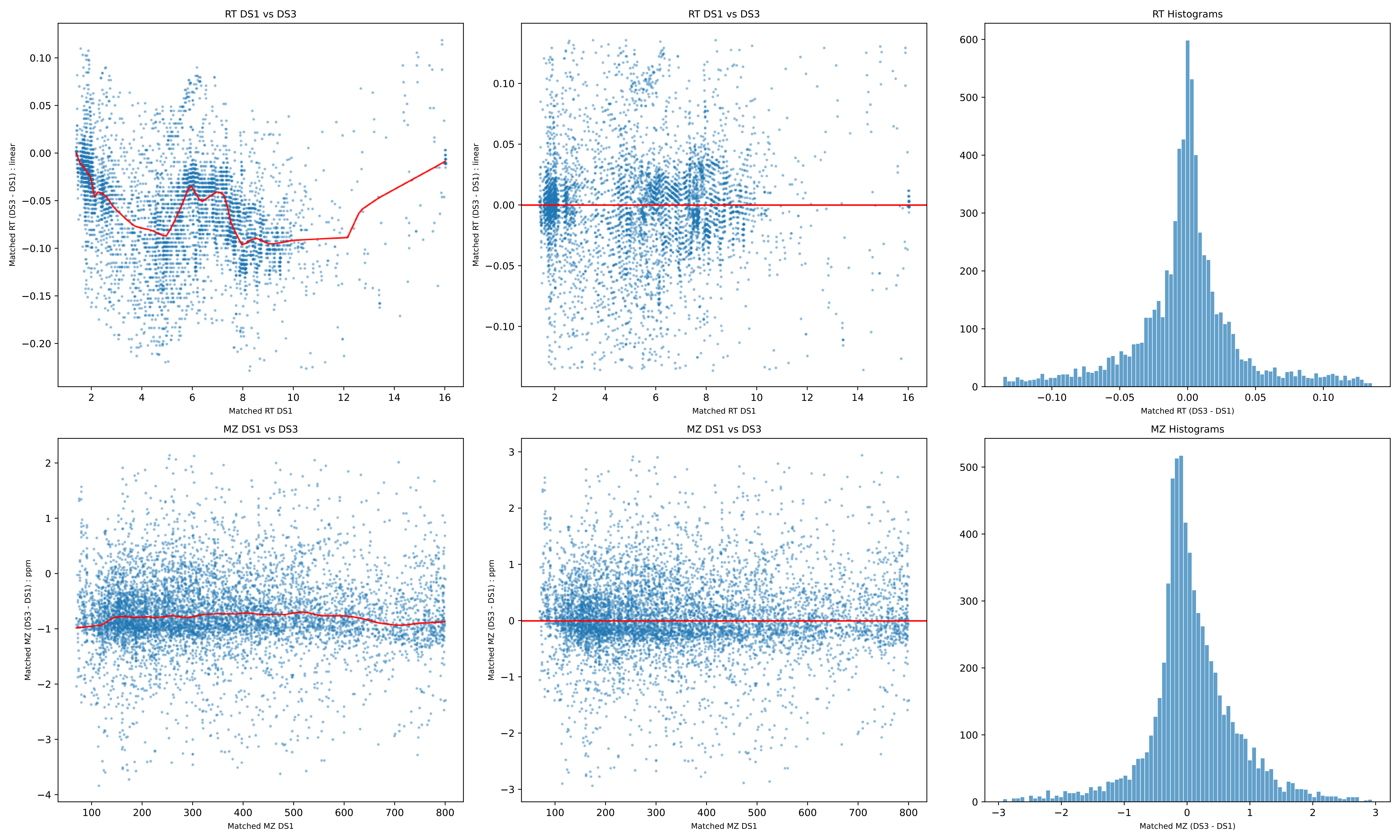
RT DS1 vs DS3
Matched RT (DS3 - DS1) : linear
Matched RT DS1
RT DS1 vs DS3
Matched RT (DS3 - DS1) : linear
Matched RT DS1
RT Histograms
Matched RT (DS3 - DS1)
MZ DS1 vs DS3
Matched MZ (DS3 - DS1) : ppm
Matched MZ DS1
MZ DS1 vs DS3
Matched MZ (DS3 - DS1) : ppm
Matched MZ DS1
MZ Histograms
Matched MZ (DS3 - DS1)

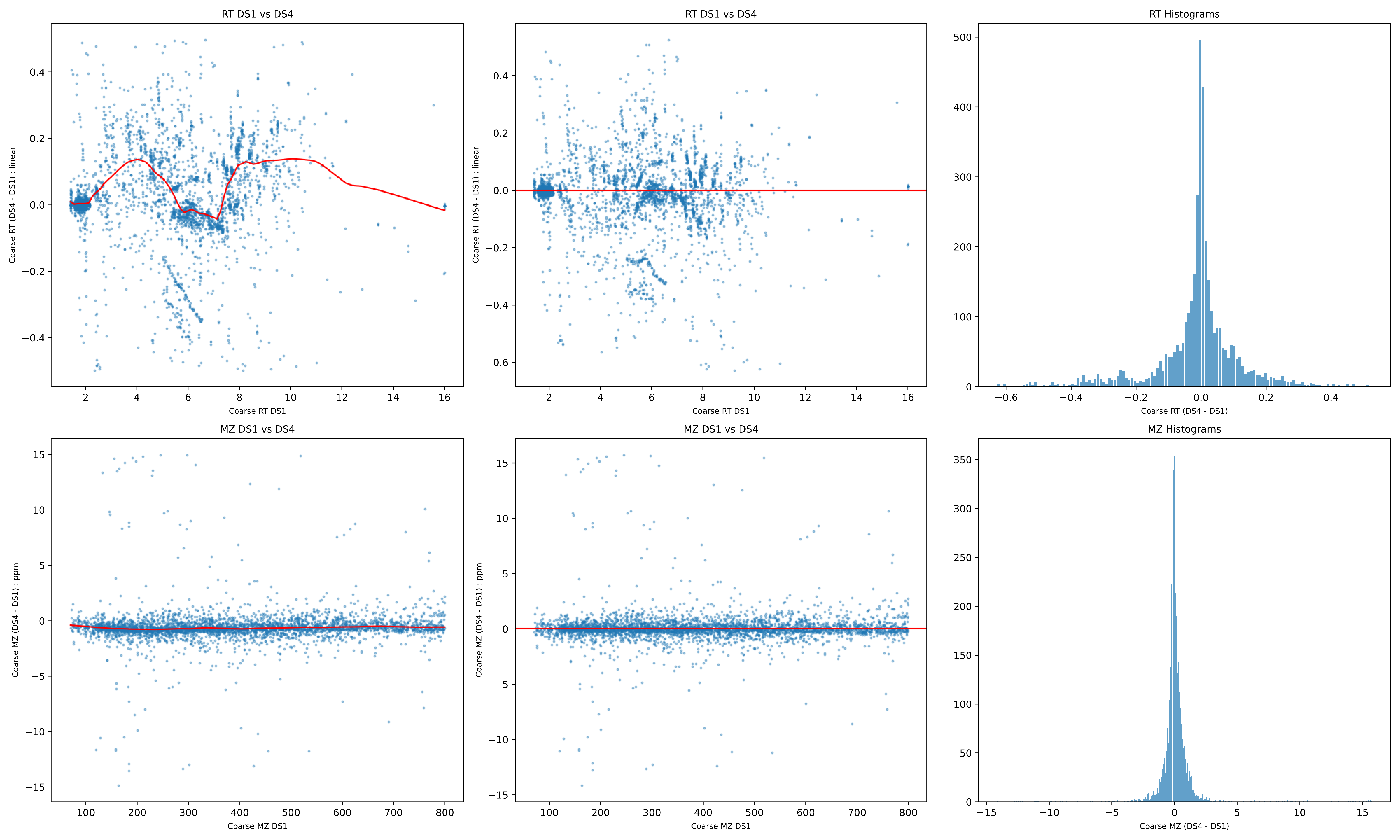
RT DS1 vs DS4
Coarse RT (DS4 - DS1) : linear
Coarse RT DS1
RT DS1 vs DS4
Coarse RT (DS4 - DS1) : linear
Coarse RT DS1
RT Histograms
Coarse RT (DS4 - DS1)
MZ DS1 vs DS4
Coarse MZ (DS4 - DS1) : ppm
Coarse MZ DS1
MZ DS1 vs DS4
Coarse MZ (DS4 - DS1) : ppm
Coarse MZ DS1
MZ Histograms
Coarse MZ (DS4 - DS1)

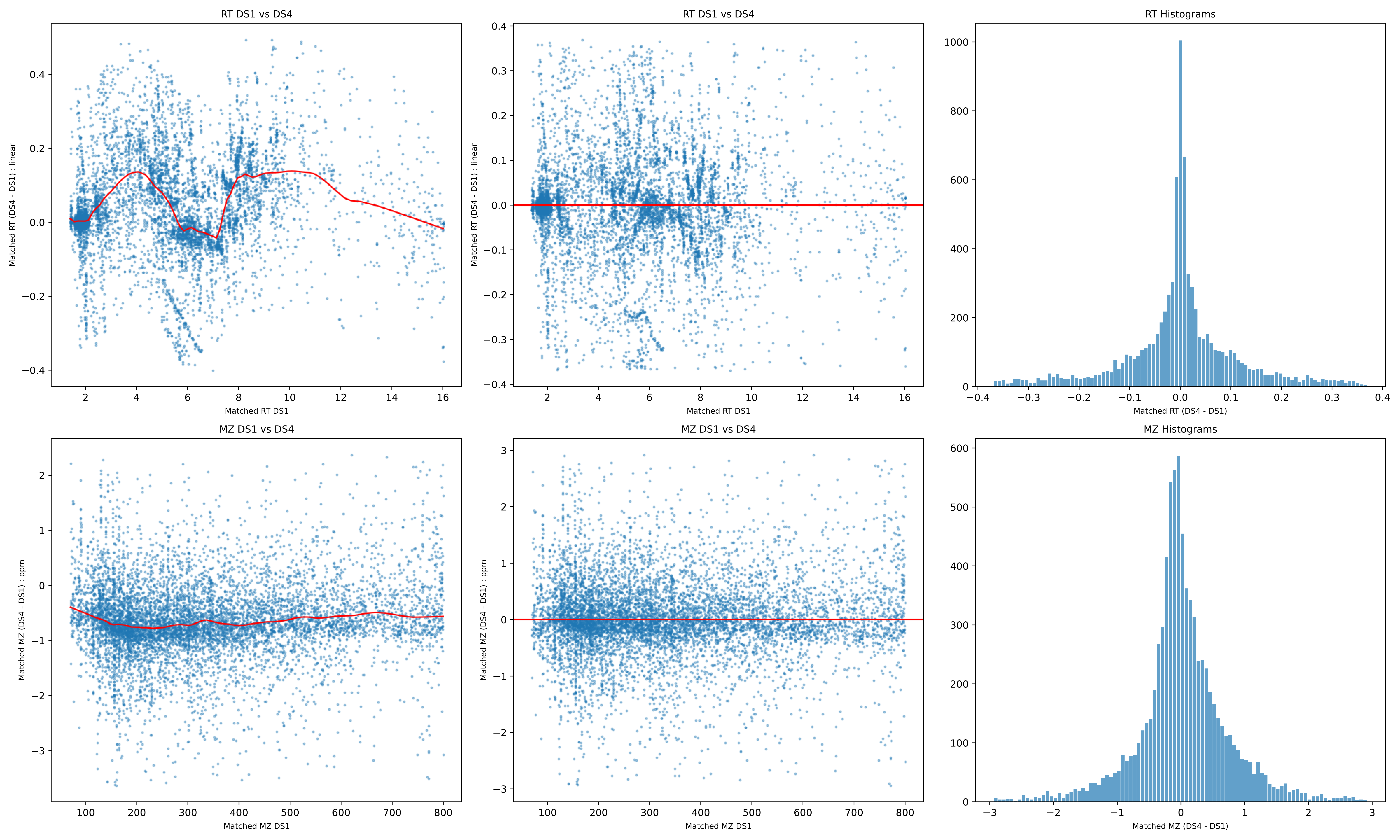
RT DS1 vs DS4
Matched RT (DS4 - DS1) : linear
Matched RT DS1
RT DS1 vs DS4
Matched RT (DS4 - DS1) : linear
Matched RT DS1
RT Histograms
Matched RT (DS4 - DS1)
MZ DS1 vs DS4
Matched MZ (DS4 - DS1) : ppm
Matched MZ DS1
MZ DS1 vs DS4
Matched MZ (DS4 - DS1) : ppm
Matched MZ DS1
MZ Histograms
Matched MZ (DS4 - DS1)

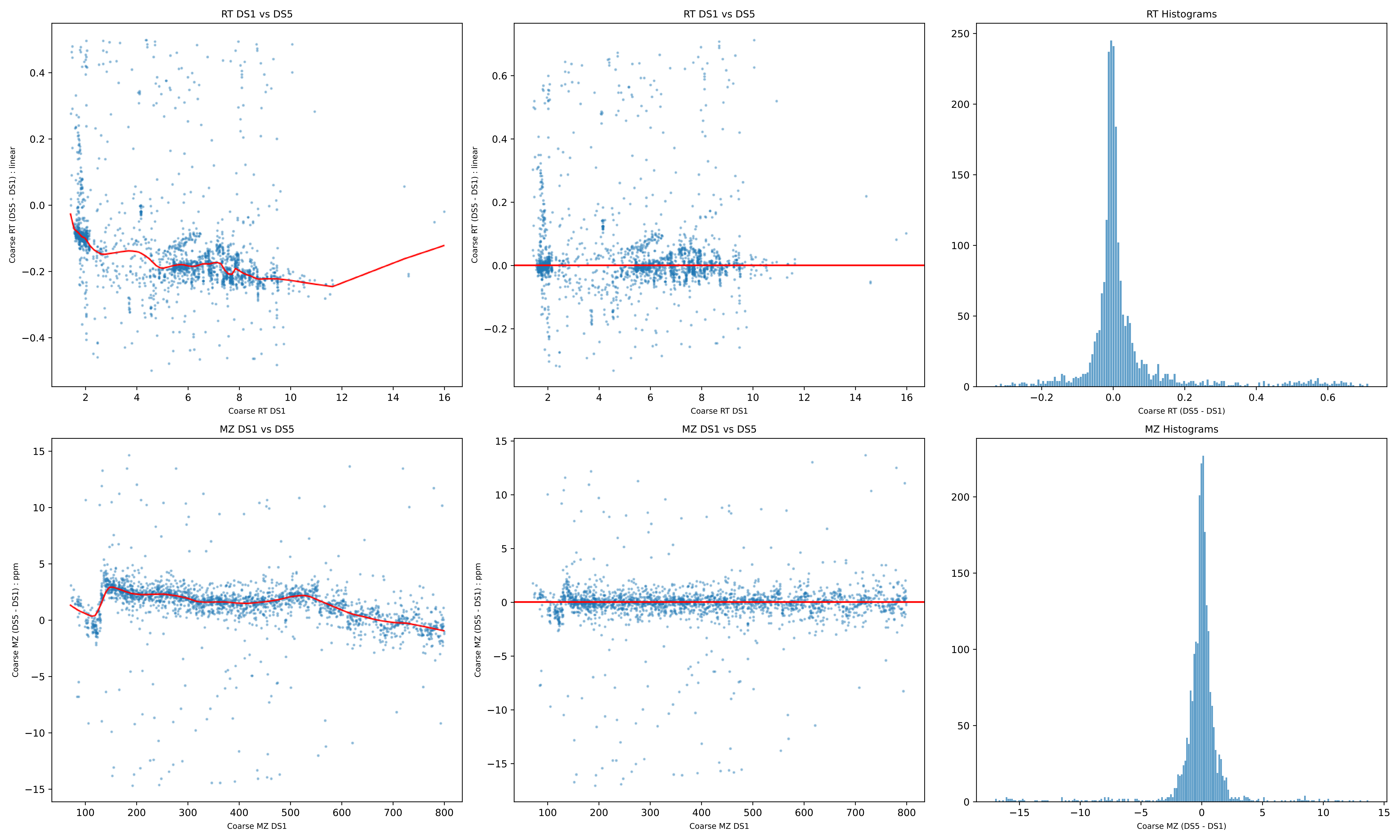
RT DS1 vs DS5
Coarse RT (DS5 - DS1) : linear
Coarse RT DS1
RT DS1 vs DS5
Coarse RT (DS5 - DS1) : linear
Coarse RT DS1
RT Histograms
Coarse RT (DS5 - DS1)
MZ DS1 vs DS5
Coarse MZ (DS5 - DS1) : ppm
Coarse MZ DS1
MZ DS1 vs DS5
Coarse MZ (DS5 - DS1) : ppm
Coarse MZ DS1
MZ Histograms
Coarse MZ (DS5 - DS1)

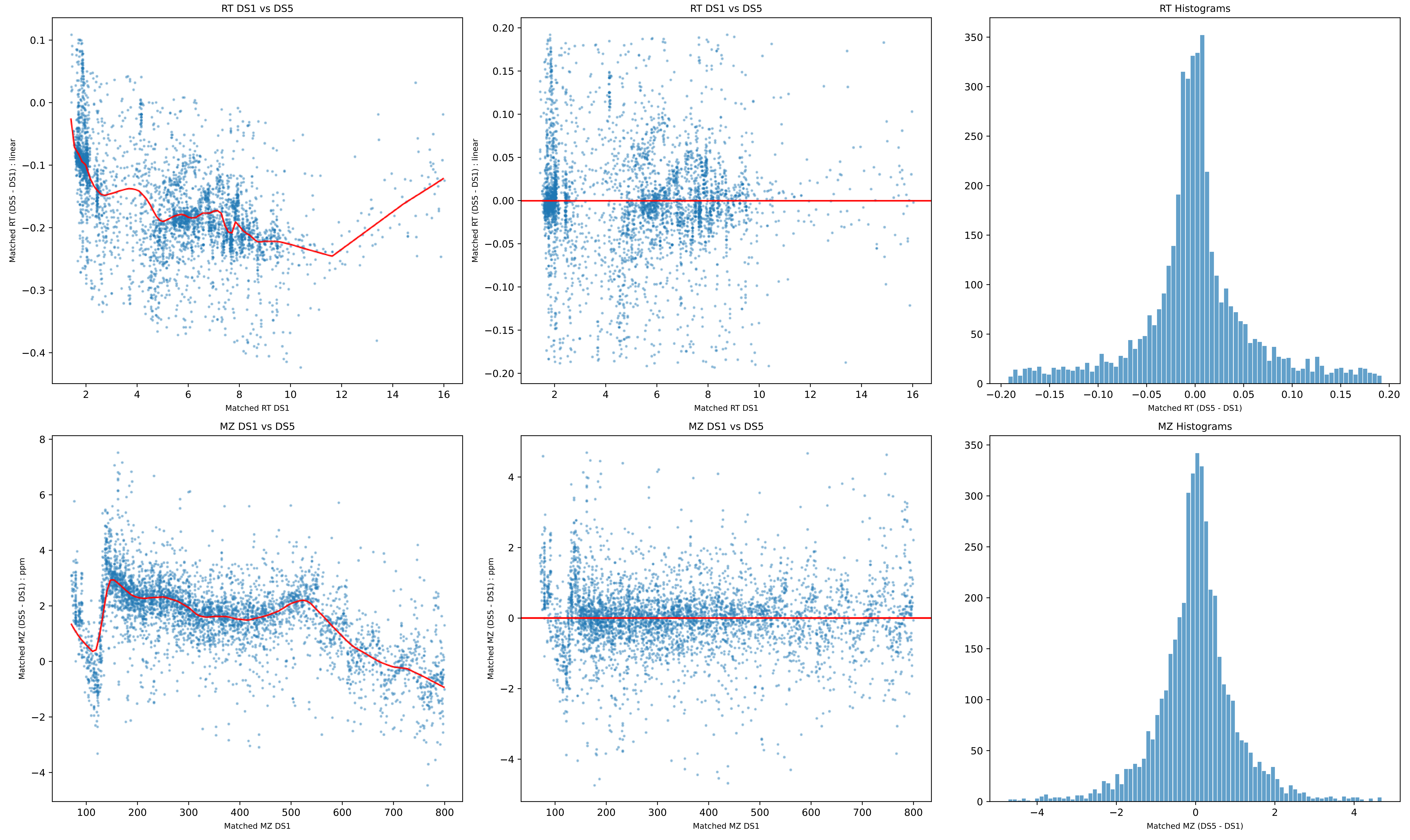
RT DS1 vs DS5
Matched RT (DS5 - DS1) : linear
Matched RT DS1
RT DS1 vs DS5
Matched RT (DS5 - DS1) : linear
Matched RT DS1
RT Histograms
Matched RT (DS5 - DS1)
MZ DS1 vs DS5
Matched MZ (DS5 - DS1) : ppm
Matched MZ DS1
MZ DS1 vs DS5
Matched MZ (DS5 - DS1) : ppm
Matched MZ DS1
MZ Histograms
Matched MZ (DS5 - DS1)

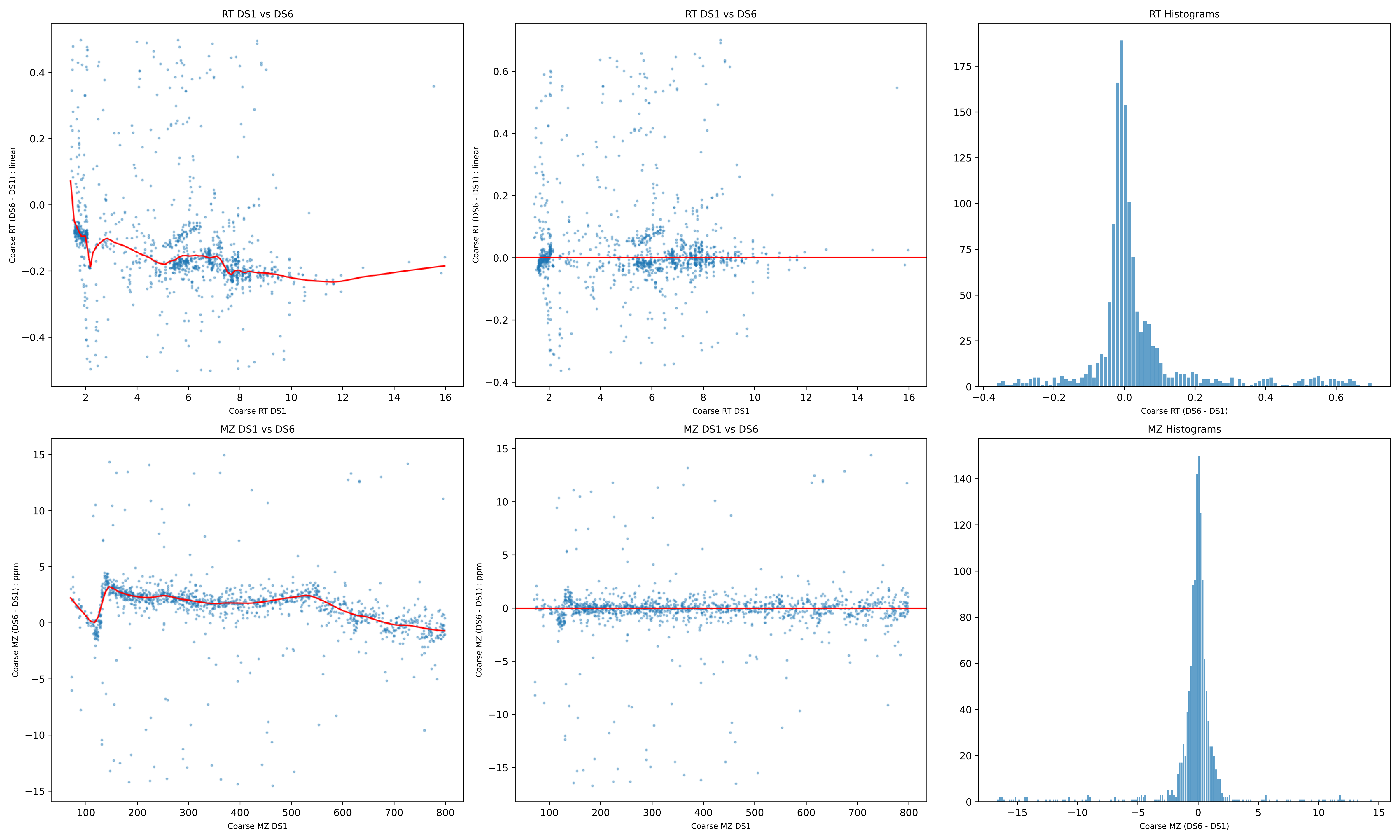

RT DS1 vs DS6
Coarse RT (DS6 - DS1) : linear
Coarse RT DS1
RT DS1 vs DS6
Coarse RT (DS6 - DS1) : linear
Coarse RT DS1
RT Histograms
Coarse RT (DS6 - DS1)
MZ DS1 vs DS6
Coarse MZ (DS6 - DS1) : ppm
Coarse MZ DS1
MZ DS1 vs DS6
Coarse MZ (DS6 - DS1) : ppm
Coarse MZ DS1
MZ Histograms
Coarse MZ (DS6 - DS1)

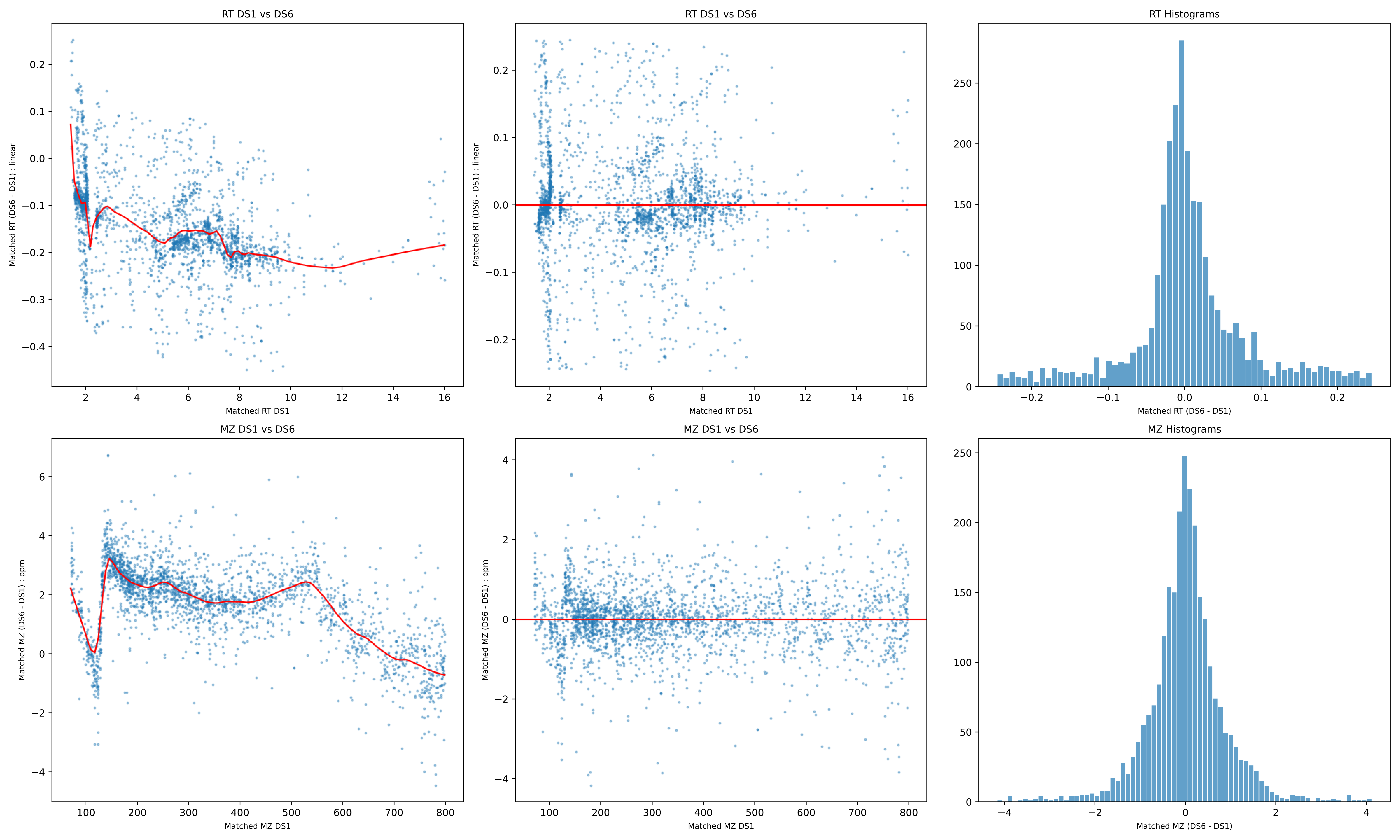
RT DS1 vs DS6
Matched RT (DS6 - DS1) : linear
Matched RT DS1
RT DS1 vs DS6
Matched RT (DS6 - DS1) : linear
Matched RT DS1
RT Histograms
Matched RT (DS6 - DS1)
MZ DS1 vs DS6
Matched MZ (DS6 - DS1) : ppm
Matched MZ DS1
MZ DS1 vs DS6
Matched MZ (DS6 - DS1) : ppm
Matched MZ DS1
MZ Histograms
Matched MZ (DS6 - DS1)

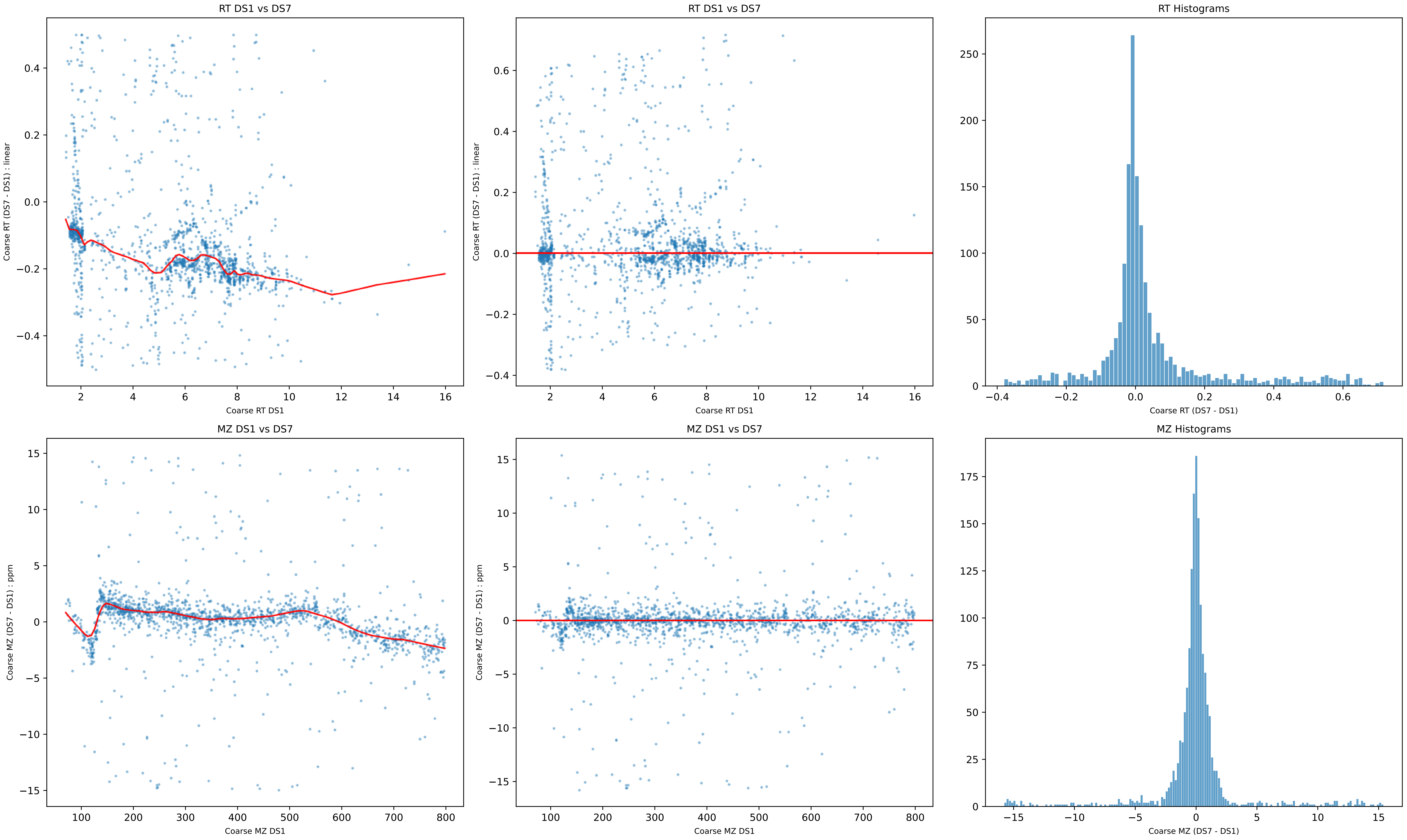

RT DS1 vs DS7
Coarse RT (DS7 - DS1) : linear
Coarse RT DS1
RT DS1 vs DS7
Coarse RT (DS7 - DS1) : linear
Coarse RT DS1
RT Histograms
Coarse RT (DS7 - DS1)
MZ DS1 vs DS7
Coarse MZ (DS7 - DS1) : ppm
Coarse MZ DS1
MZ DS1 vs DS7
Coarse MZ (DS7 - DS1) : ppm
Coarse MZ DS1
MZ Histograms
Coarse MZ (DS7 - DS1)

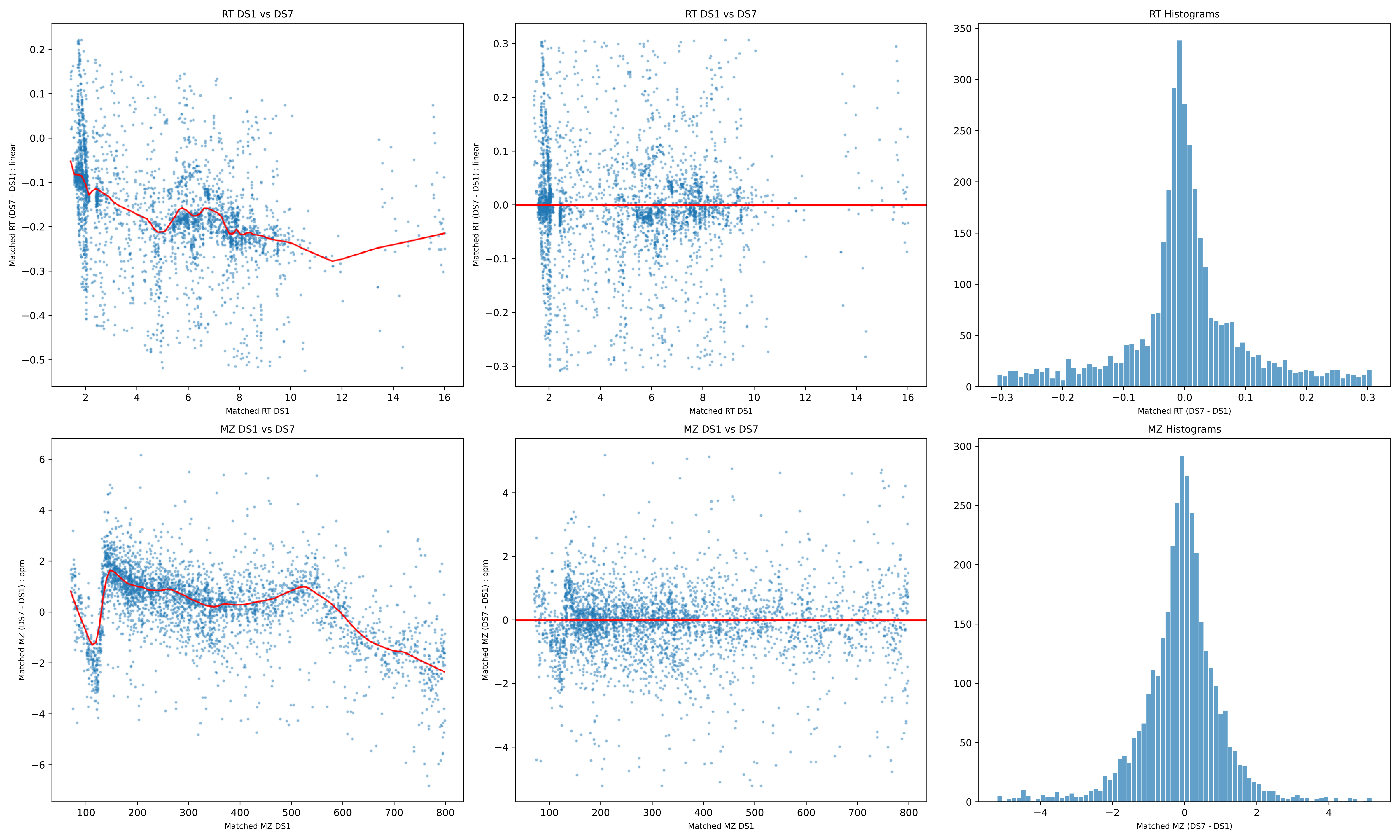
RT DS1 vs DS7
Matched RT (DS7 - DS1) : linear
Matched RT DS1
RT DS1 vs DS7
Matched RT (DS7 - DS1) : linear
Matched RT DS1
RT Histograms
Matched RT (DS7 - DS1)
MZ DS1 vs DS7
Matched MZ (DS7 - DS1) : ppm
Matched MZ DS1
MZ DS1 vs DS7
Matched MZ (DS7 - DS1) : ppm
Matched MZ DS1
MZ Histograms
Matched MZ (DS7 - DS1)

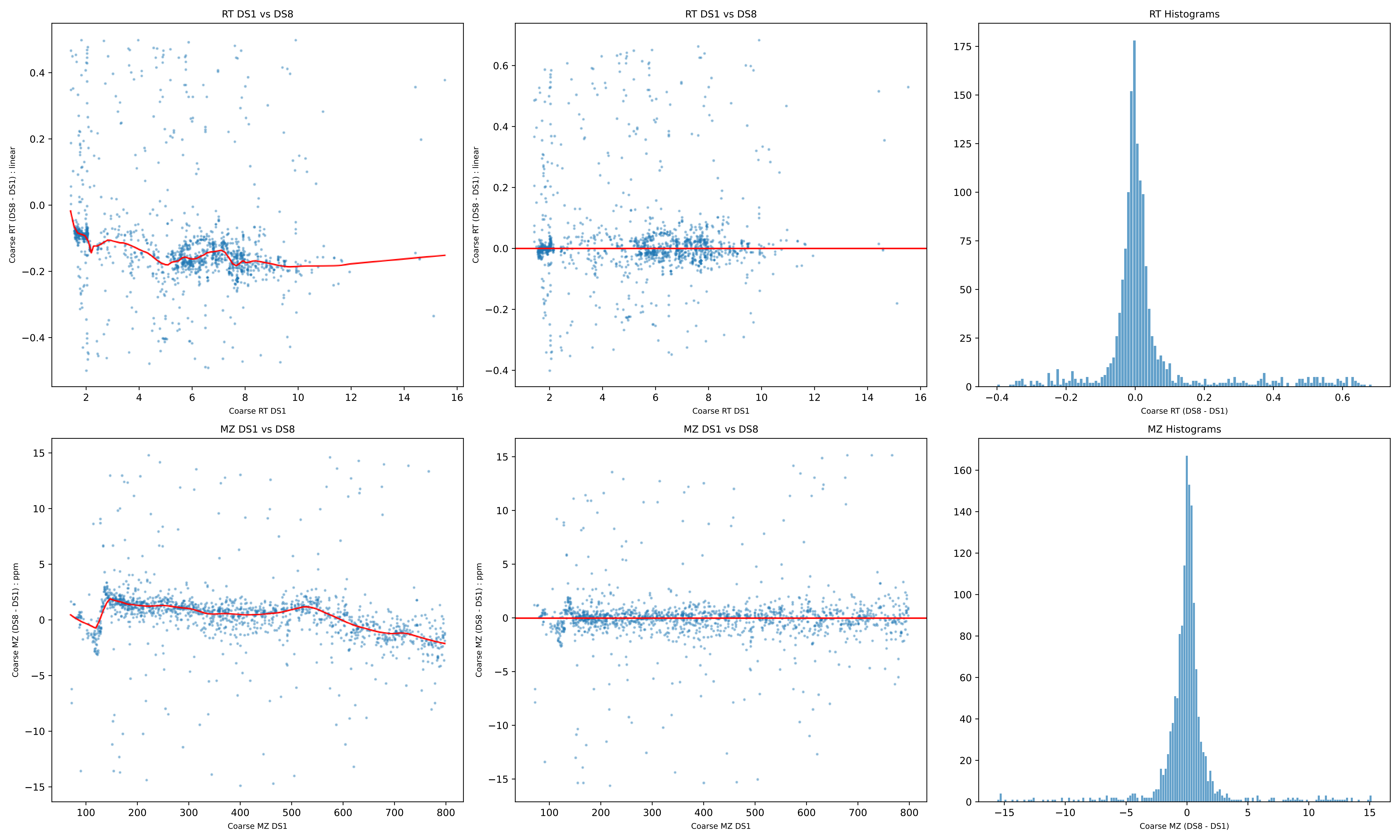

RT DS1 vs DS8
Coarse RT (DS8 - DS1) : linear
Coarse RT DS1
RT DS1 vs DS8
Coarse RT (DS8 - DS1) : linear
Coarse RT DS1
RT Histograms
Coarse RT (DS8 - DS1)
MZ DS1 vs DS8
Coarse MZ (DS8 - DS1) : ppm
Coarse MZ DS1
MZ DS1 vs DS8
Coarse MZ (DS8 - DS1) : ppm
Coarse MZ DS1
MZ Histograms
Coarse MZ (DS8 - DS1)

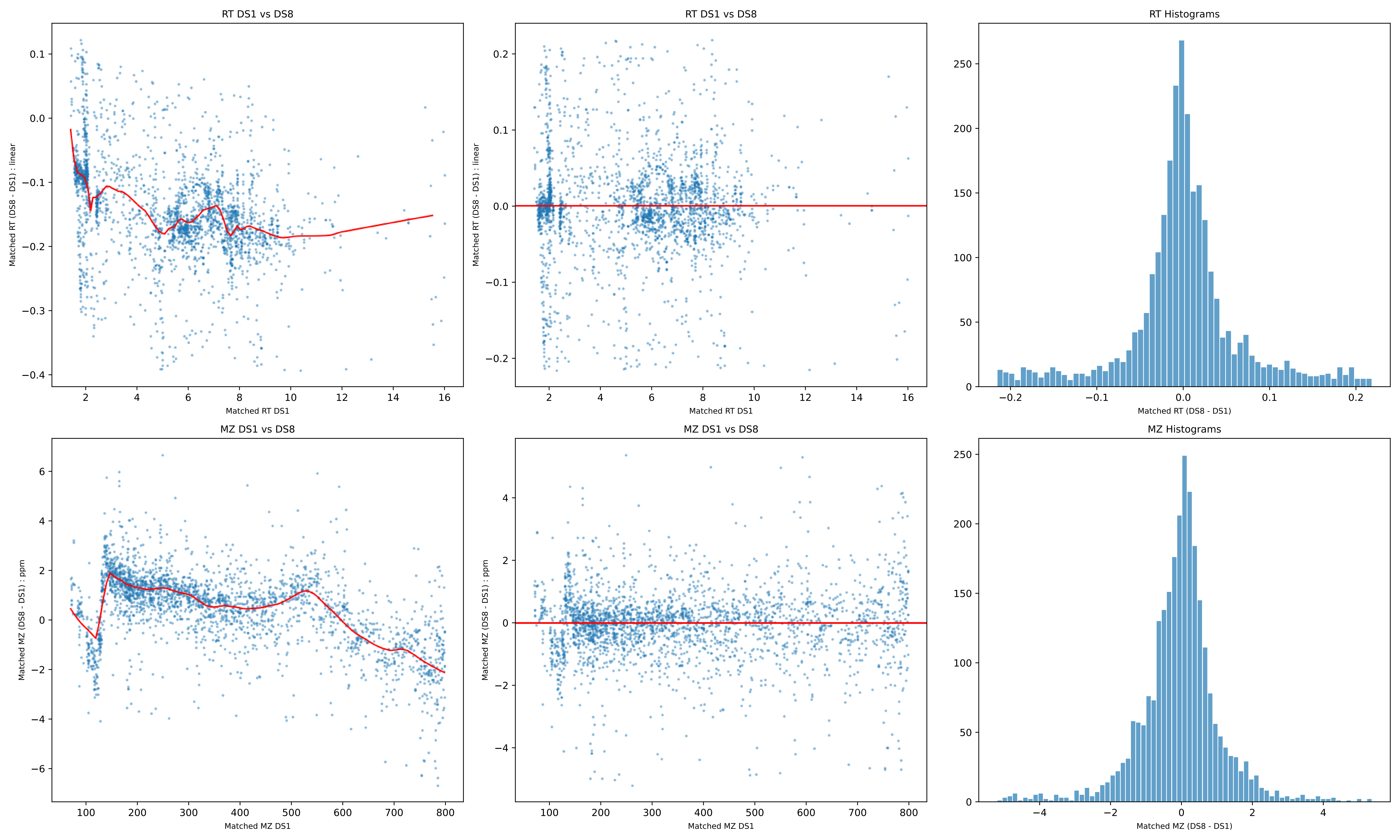
RT DS1 vs DS8
Matched RT (DS8 - DS1) : linear
Matched RT DS1
RT DS1 vs DS8
Matched RT (DS8 - DS1) : linear
Matched RT DS1
RT Histograms
Matched RT (DS8 - DS1)
MZ DS1 vs DS8
Matched MZ (DS8 - DS1) : ppm
Matched MZ DS1
MZ DS1 vs DS8
Matched MZ (DS8 - DS1) : ppm
Matched MZ DS1
MZ Histograms
Matched MZ (DS8 - DS1)

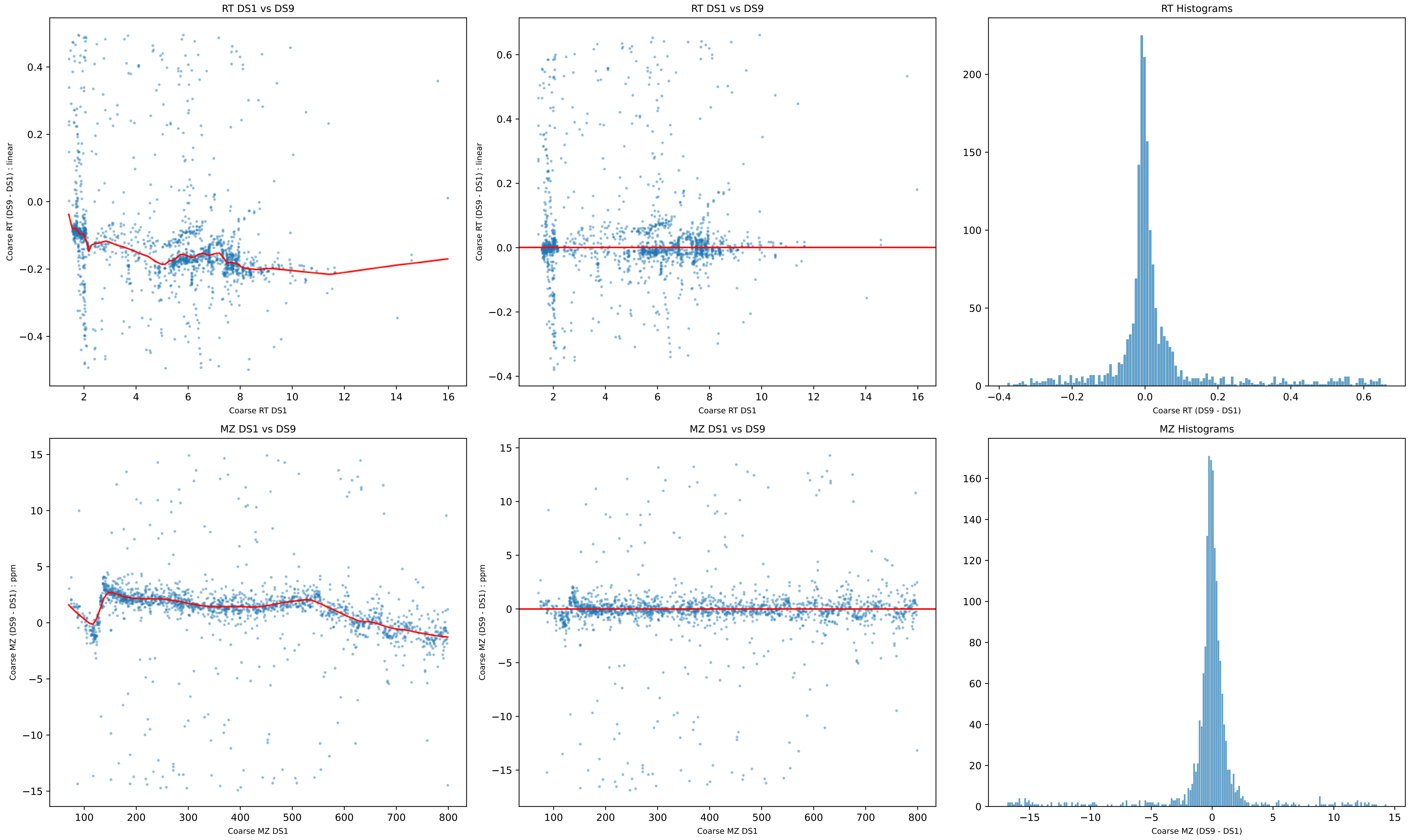
RT DS1 vs DS9
Coarse RT (DS9 - DS1) : linear
Coarse RT DS1
RT DS1 vs DS9
Coarse RT (DS9 - DS1) : linear
Coarse RT DS1
RT Histograms
Coarse RT (DS9 - DS1)
MZ DS1 vs DS9
Coarse MZ (DS9 - DS1) : ppm
Coarse MZ DS1
MZ DS1 vs DS9
Coarse MZ (DS9 - DS1) : ppm
Coarse MZ DS1
MZ Histograms
Coarse MZ (DS9 - DS1)

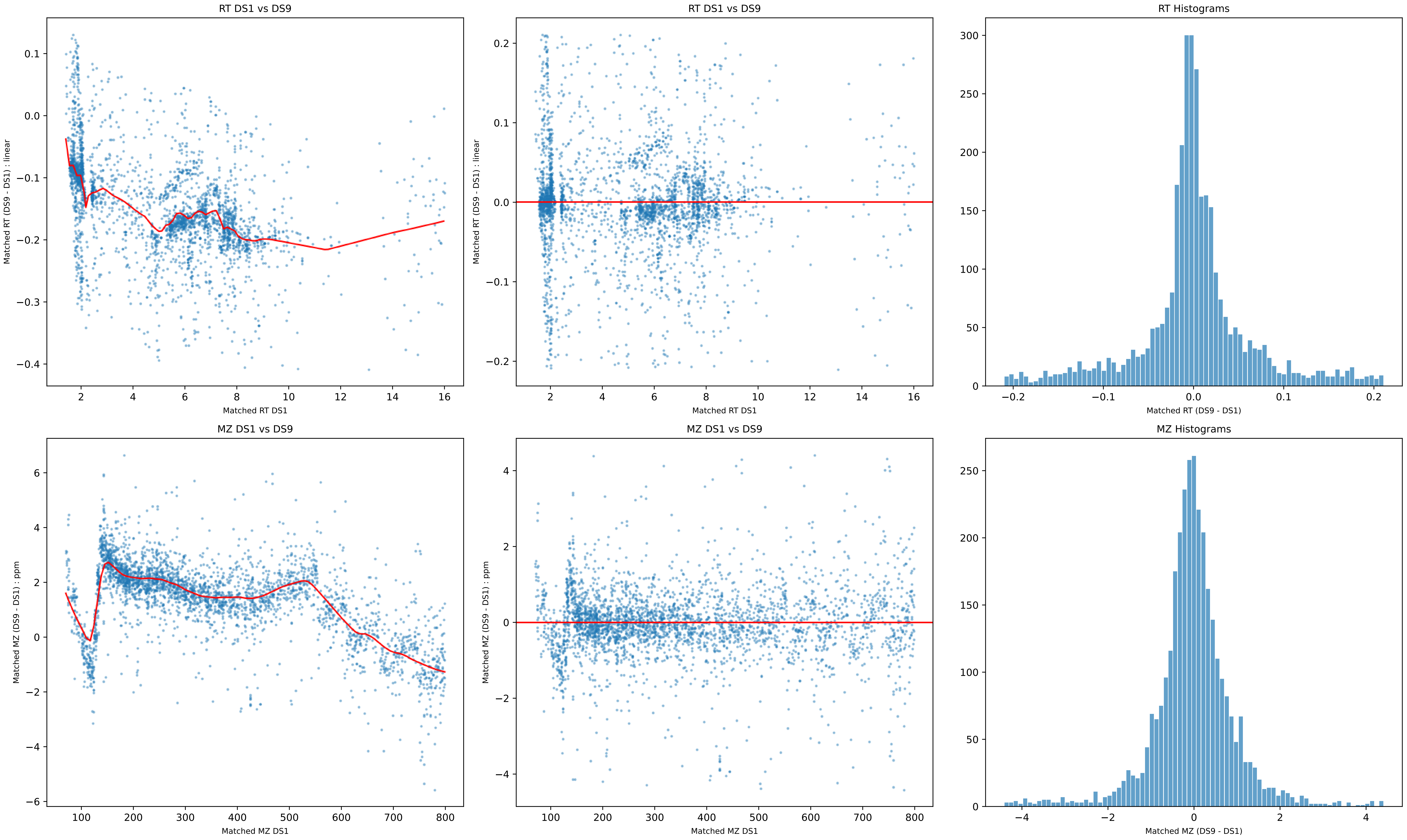
RT DS1 vs DS9
Matched RT (DS9 - DS1) : linear
Matched RT DS1
RT DS1 vs DS9
Matched RT (DS9 - DS1) : linear
Matched RT DS1
RT Histograms
Matched RT (DS9 - DS1)
MZ DS1 vs DS9
Matched MZ (DS9 - DS1) : ppm
Matched MZ DS1
MZ DS1 vs DS9
Matched MZ (DS9 - DS1) : ppm
Matched MZ DS1
MZ Histograms
Matched MZ (DS9 - DS1)

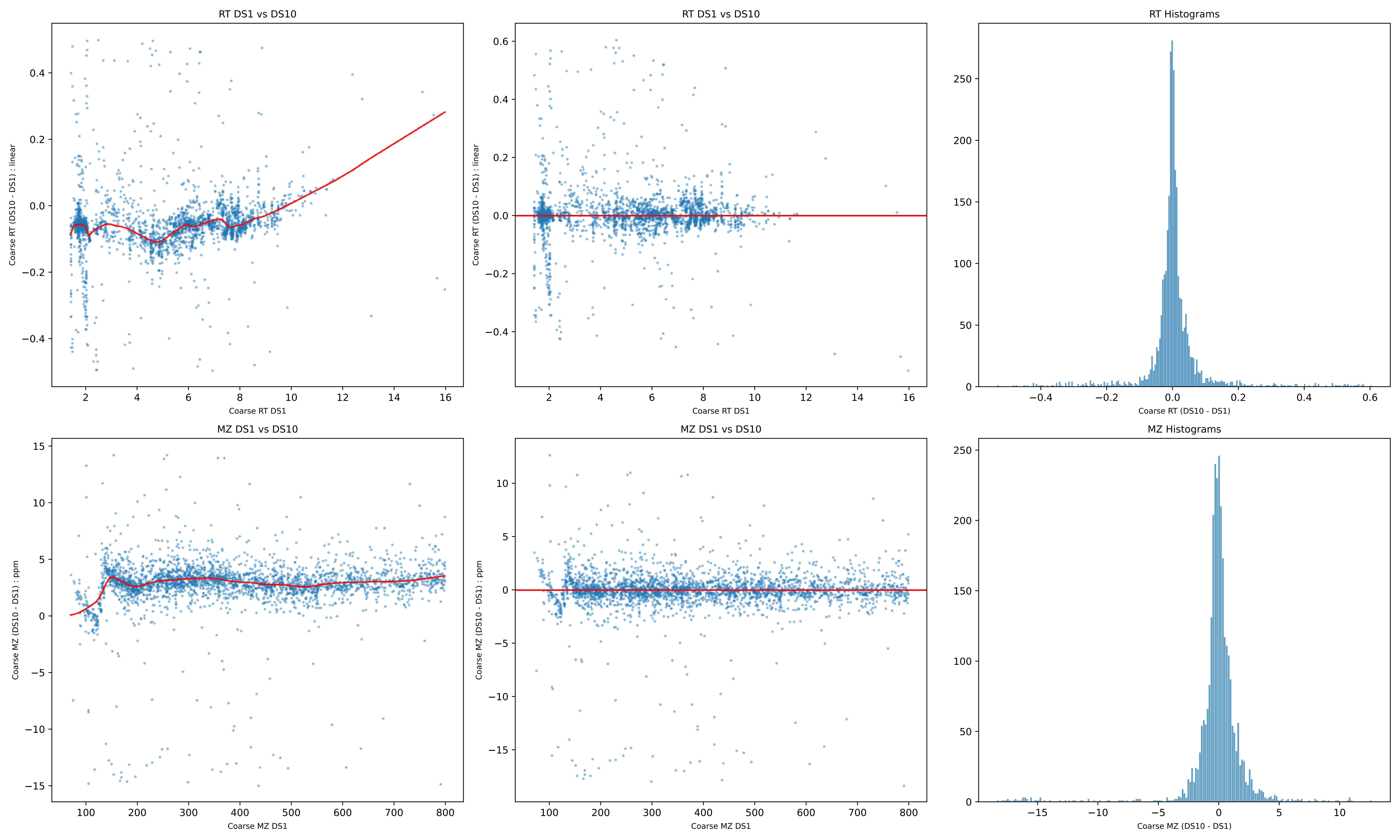
RT DS1 vs DS10
Coarse RT (DS10 - DS1) : linear
Coarse RT DS1
RT DS1 vs DS10
Coarse RT (DS10 - DS1) : linear
Coarse RT DS1
RT Histograms
Coarse RT (DS10 - DS1)
MZ DS1 vs DS10
Coarse MZ (DS10 - DS1) : ppm
Coarse MZ DS1
MZ DS1 vs DS10
Coarse MZ (DS10 - DS1) : ppm
Coarse MZ DS1
MZ Histograms
Coarse MZ (DS10 - DS1)

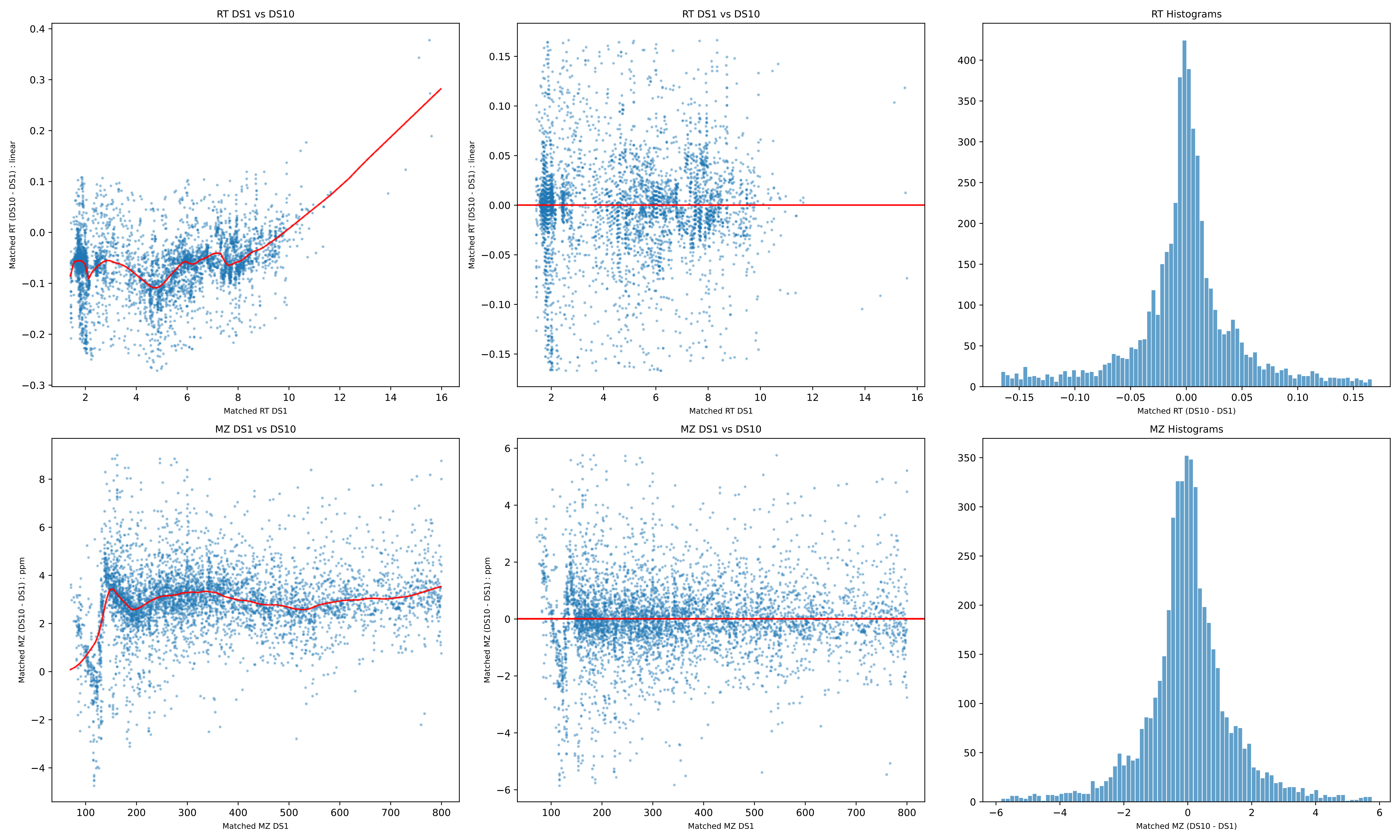
RT DS1 vs DS10
Matched RT (DS10 - DS1) : linear
Matched RT DS1
RT DS1 vs DS10
Matched RT (DS10 - DS1) : linear
Matched RT DS1
RT Histograms
Matched RT (DS10 - DS1)
MZ DS1 vs DS10
Matched MZ (DS10 - DS1) : ppm
Matched MZ DS1
MZ DS1 vs DS10
Matched MZ (DS10 - DS1) : ppm
Matched MZ DS1
MZ Histograms
Matched MZ (DS10 - DS1)

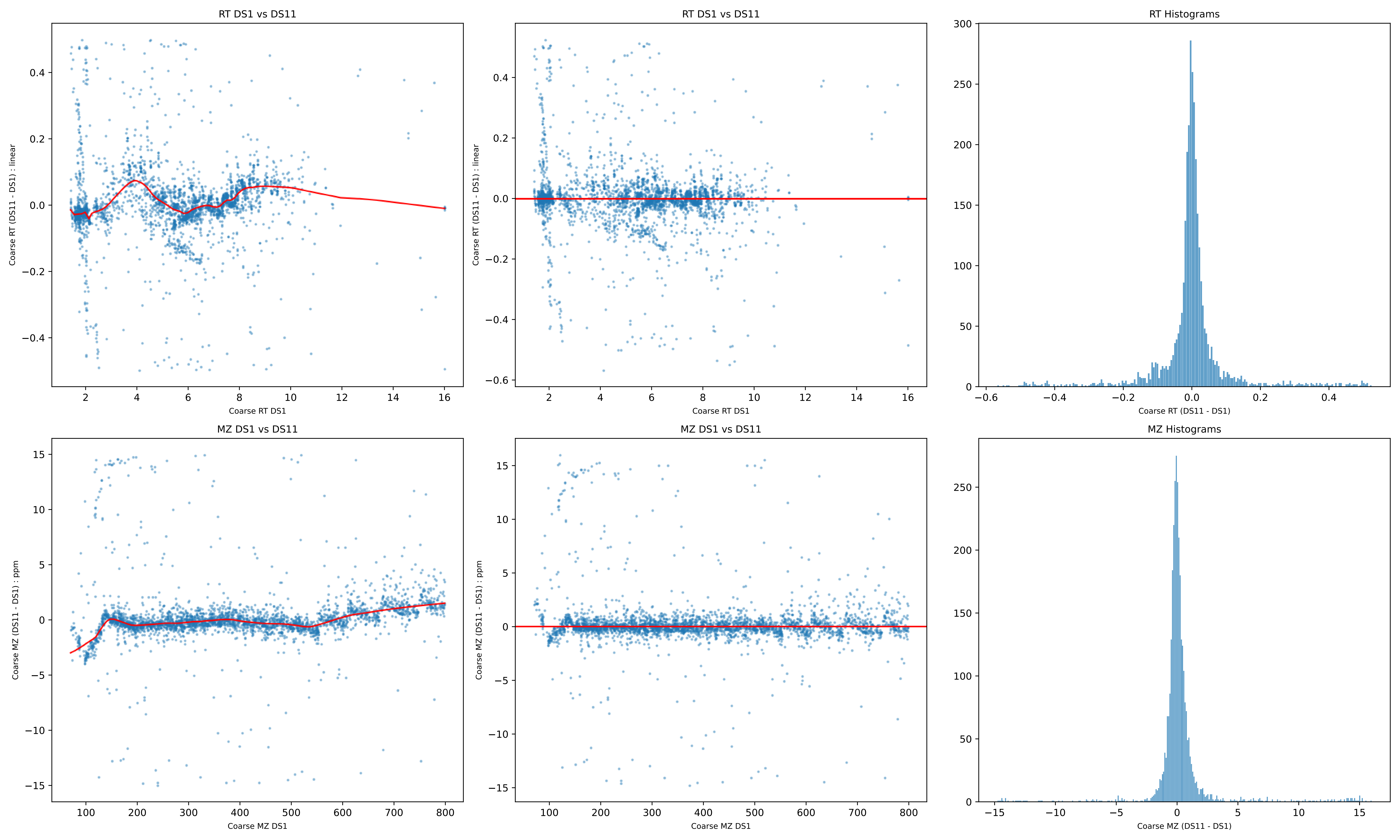
RT DS1 vs DS11
Coarse RT (DS11 - DS1) : linear
Coarse RT DS1
RT DS1 vs DS11
Coarse RT (DS11 - DS1) : linear
Coarse RT DS1
RT Histograms
Coarse RT (DS11 - DS1)
MZ DS1 vs DS11
Coarse MZ (DS11 - DS1) : ppm
Coarse MZ DS1
MZ DS1 vs DS11
Coarse MZ (DS11 - DS1) : ppm
Coarse MZ DS1
MZ Histograms
Coarse MZ (DS11 - DS1)

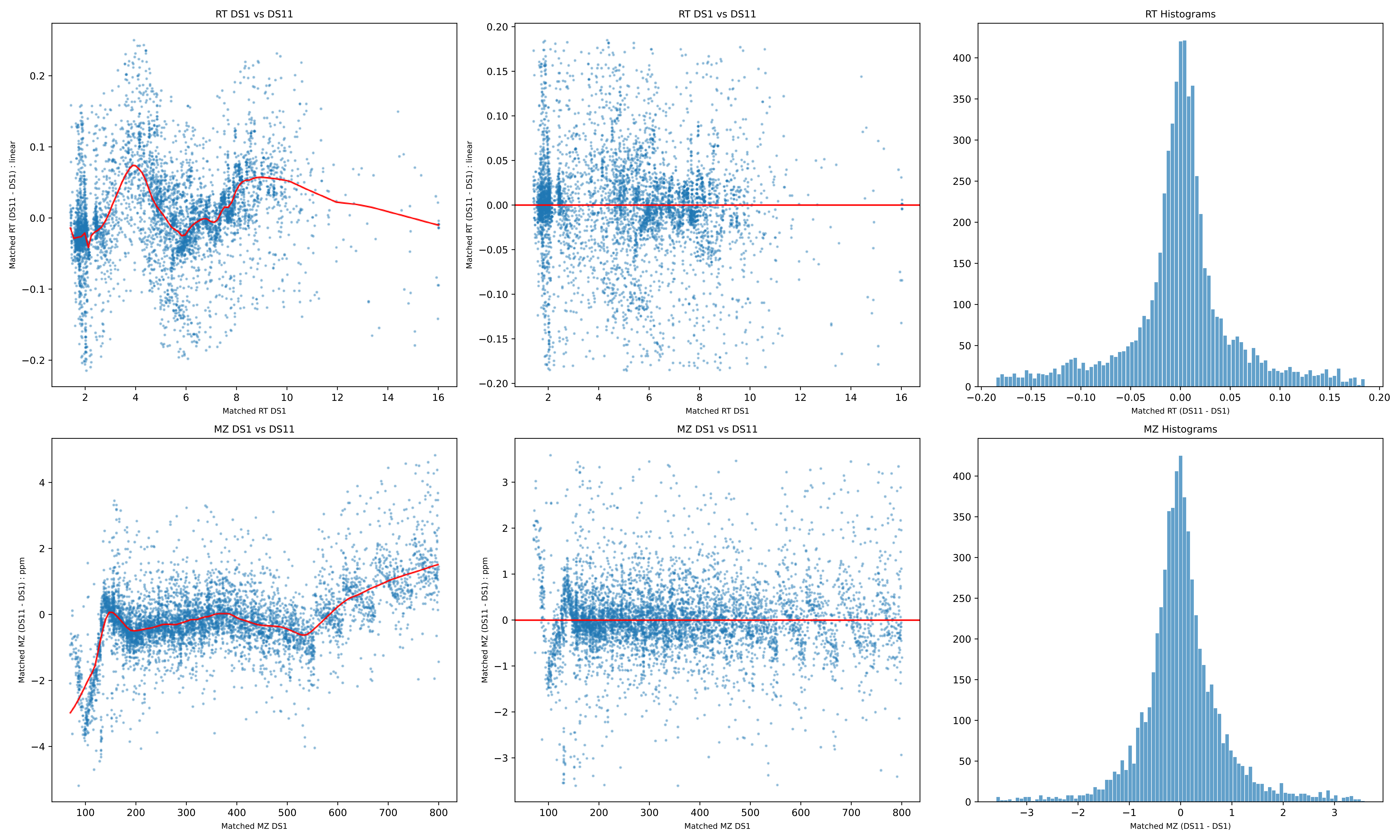
RT DS1 vs DS11
Matched RT (DS11 - DS1) : linear
Matched RT DS1
RT DS1 vs DS11
Matched RT (DS11 - DS1) : linear
Matched RT DS1
RT Histograms
Matched RT (DS11 - DS1)
MZ DS1 vs DS11
Matched MZ (DS11 - DS1) : ppm
Matched MZ DS1
MZ DS1 vs DS11
Matched MZ (DS11 - DS1) : ppm
Matched MZ DS1
MZ Histograms
Matched MZ (DS11 - DS1)

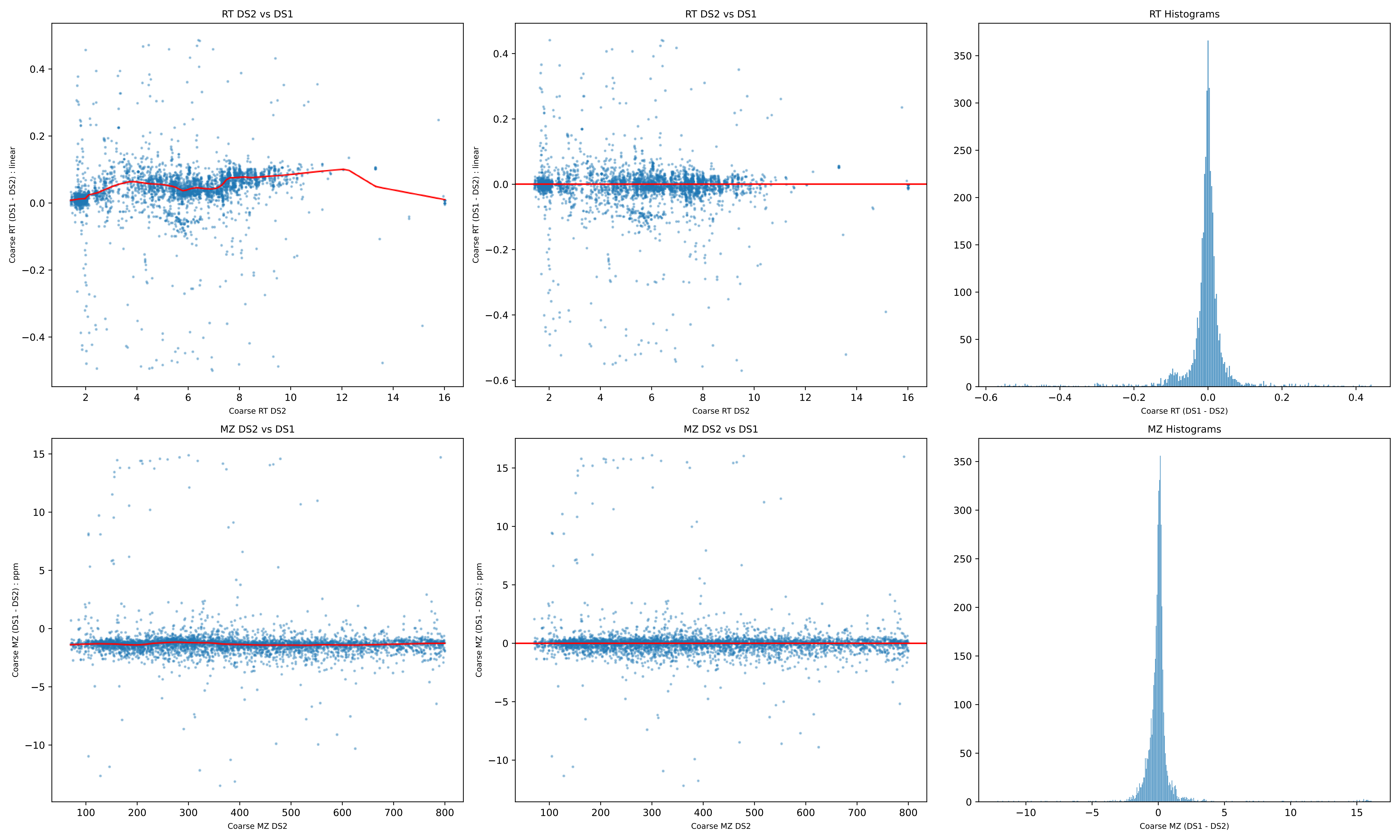
RT DS2 vs DS1
Coarse RT (DS1 - DS2) : linear
Coarse RT DS2
RT DS2 vs DS1
Coarse RT (DS1 - DS2) : linear
Coarse RT DS2
RT Histograms
Coarse RT (DS1 - DS2)
MZ DS2 vs DS1
Coarse MZ (DS1 - DS2) : ppm
Coarse MZ DS2
MZ DS2 vs DS1
Coarse MZ (DS1 - DS2) : ppm
Coarse MZ DS2
MZ Histograms
Coarse MZ (DS1 - DS2)

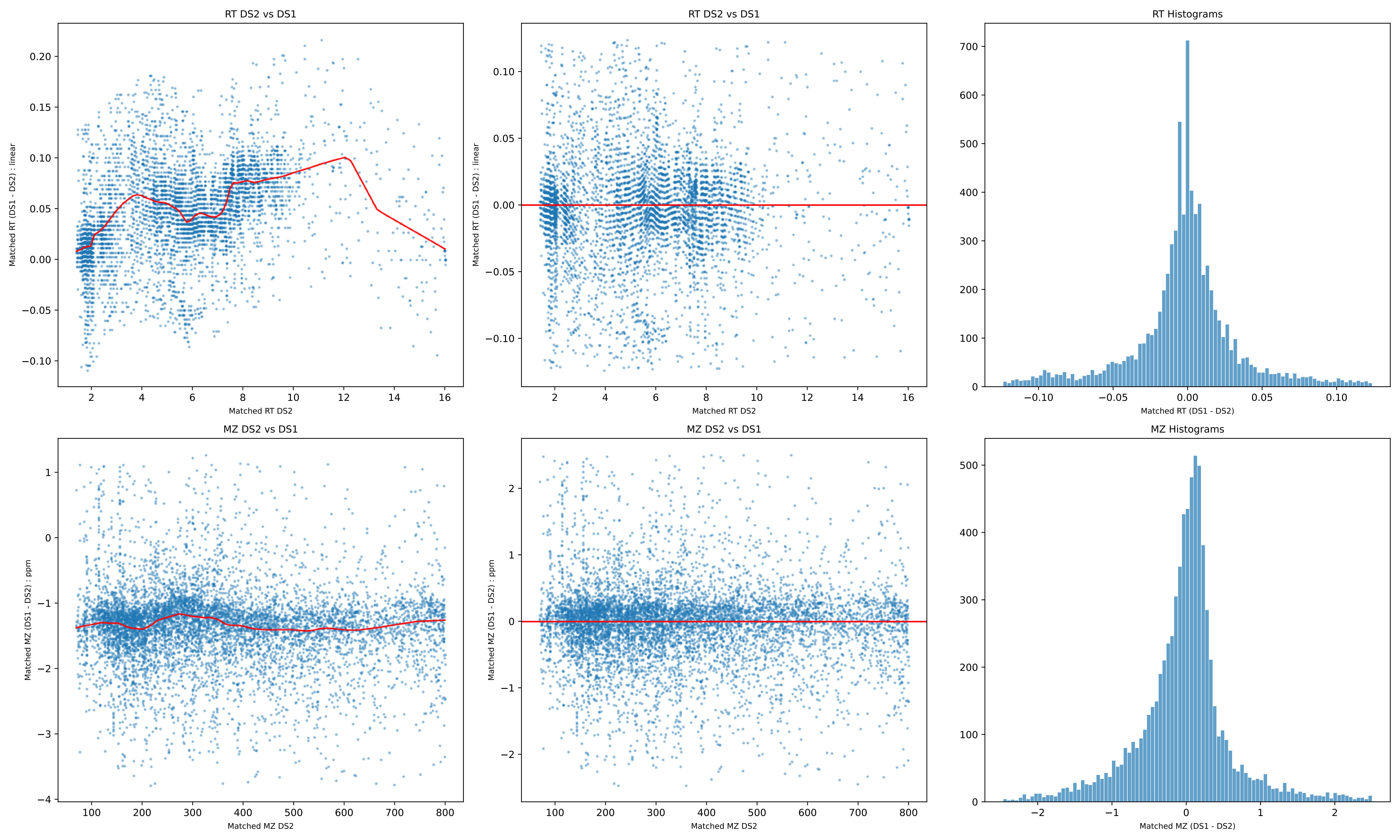

RT DS2 vs DS1
Matched RT (DS1 - DS2) : linear
Matched RT DS2
RT DS2 vs DS1
Matched RT (DS1 - DS2) : linear
Matched RT DS2
RT Histograms
Matched RT (DS1 - DS2)
MZ DS2 vs DS1
Matched MZ (DS1 - DS2) : ppm
Matched MZ DS2
MZ DS2 vs DS1
Matched MZ (DS1 - DS2) : ppm
Matched MZ DS2
MZ Histograms
Matched MZ (DS1 - DS2)

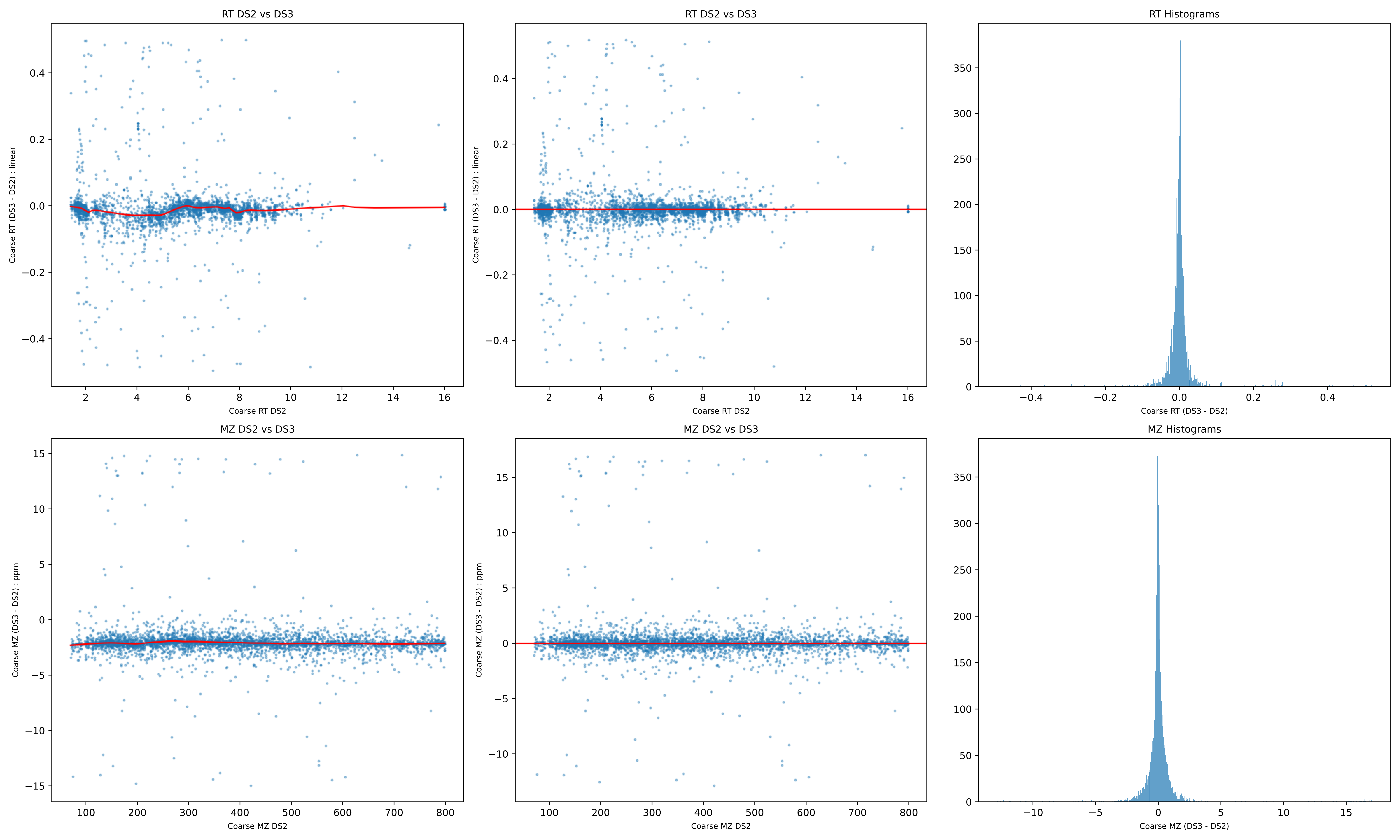
RT DS2 vs DS3
Coarse RT (DS3 - DS2) : linear
Coarse RT DS2
RT DS2 vs DS3
Coarse RT (DS3 - DS2) : linear
Coarse RT DS2
RT Histograms
Coarse RT (DS3 - DS2)
MZ DS2 vs DS3
Coarse MZ (DS3 - DS2) : ppm
Coarse MZ DS2
MZ DS2 vs DS3
Coarse MZ (DS3 - DS2) : ppm
Coarse MZ DS2
MZ Histograms
Coarse MZ (DS3 - DS2)

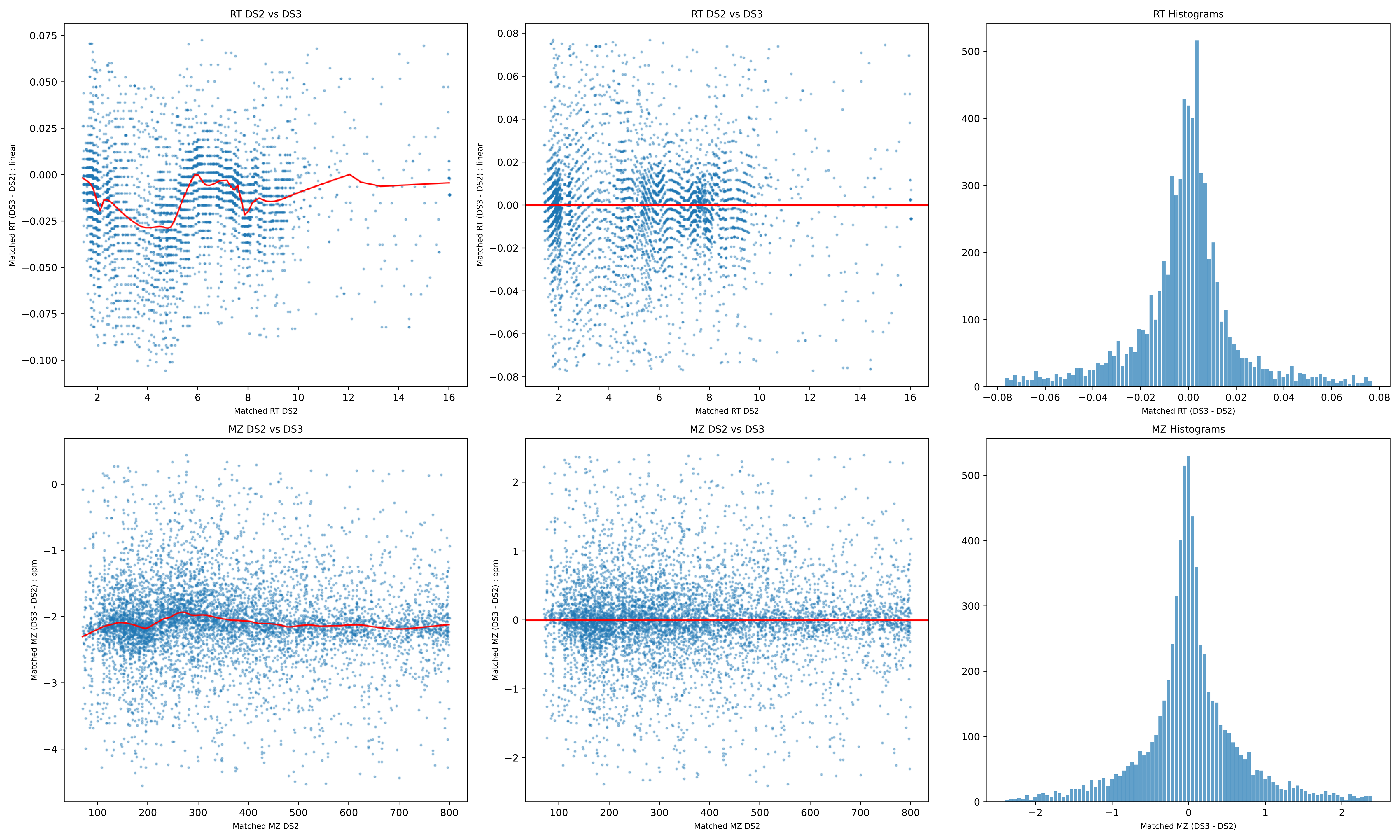

RT DS2 vs DS3
Matched RT (DS3 - DS2) : linear
Matched RT DS2
RT DS2 vs DS3
Matched RT (DS3 - DS2) : linear
Matched RT DS2
RT Histograms
Matched RT (DS3 - DS2)
MZ DS2 vs DS3
Matched MZ (DS3 - DS2) : ppm
Matched MZ DS2
MZ DS2 vs DS3
Matched MZ (DS3 - DS2) : ppm
Matched MZ DS2
MZ Histograms
Matched MZ (DS3 - DS2)

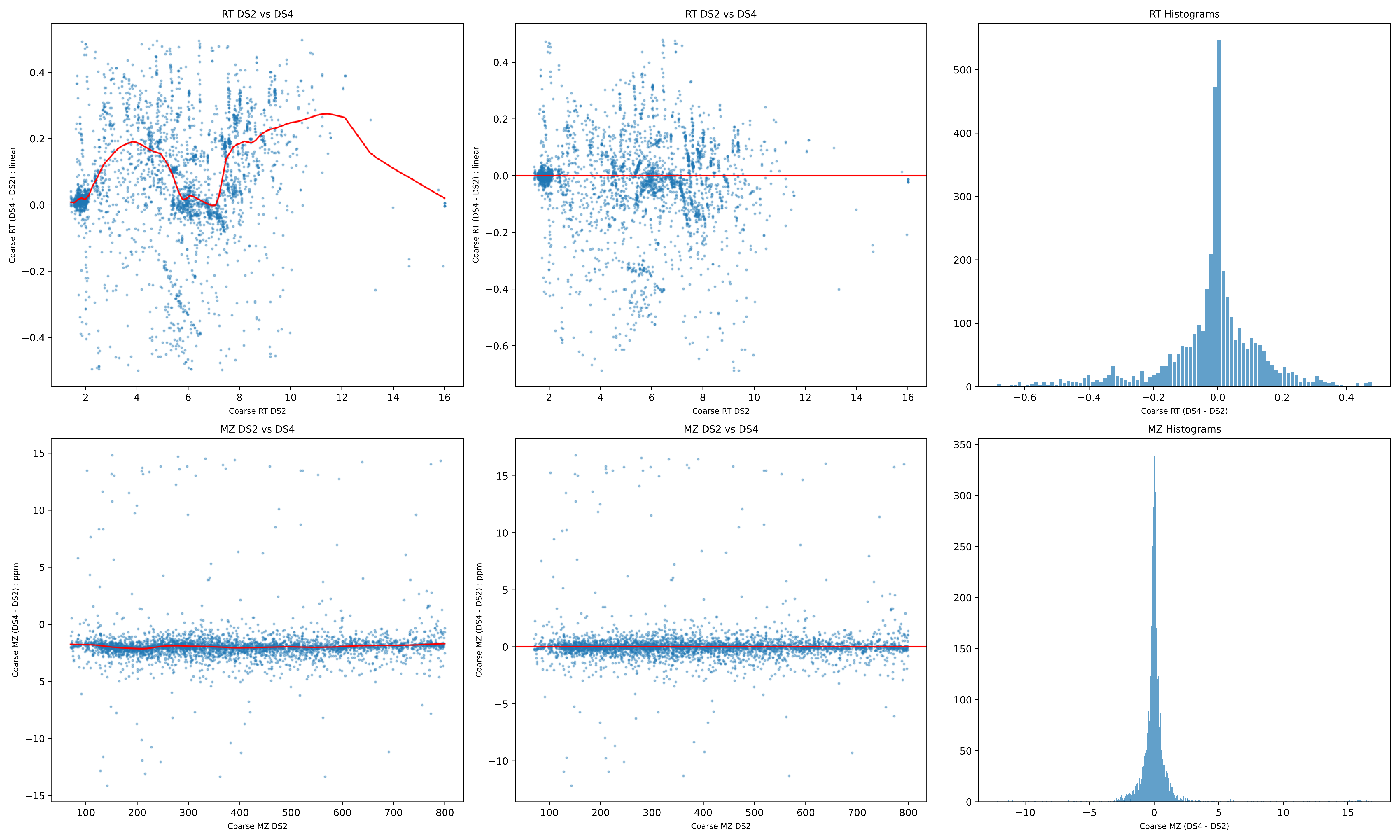
RT DS2 vs DS4
Coarse RT (DS4 - DS2) : linear
Coarse RT DS2
RT DS2 vs DS4
Coarse RT (DS4 - DS2) : linear
Coarse RT DS2
RT Histograms
Coarse RT (DS4 - DS2)
MZ DS2 vs DS4
Coarse MZ (DS4 - DS2) : ppm
Coarse MZ DS2
MZ DS2 vs DS4
Coarse MZ (DS4 - DS2) : ppm
Coarse MZ DS2
MZ Histograms
Coarse MZ (DS4 - DS2)

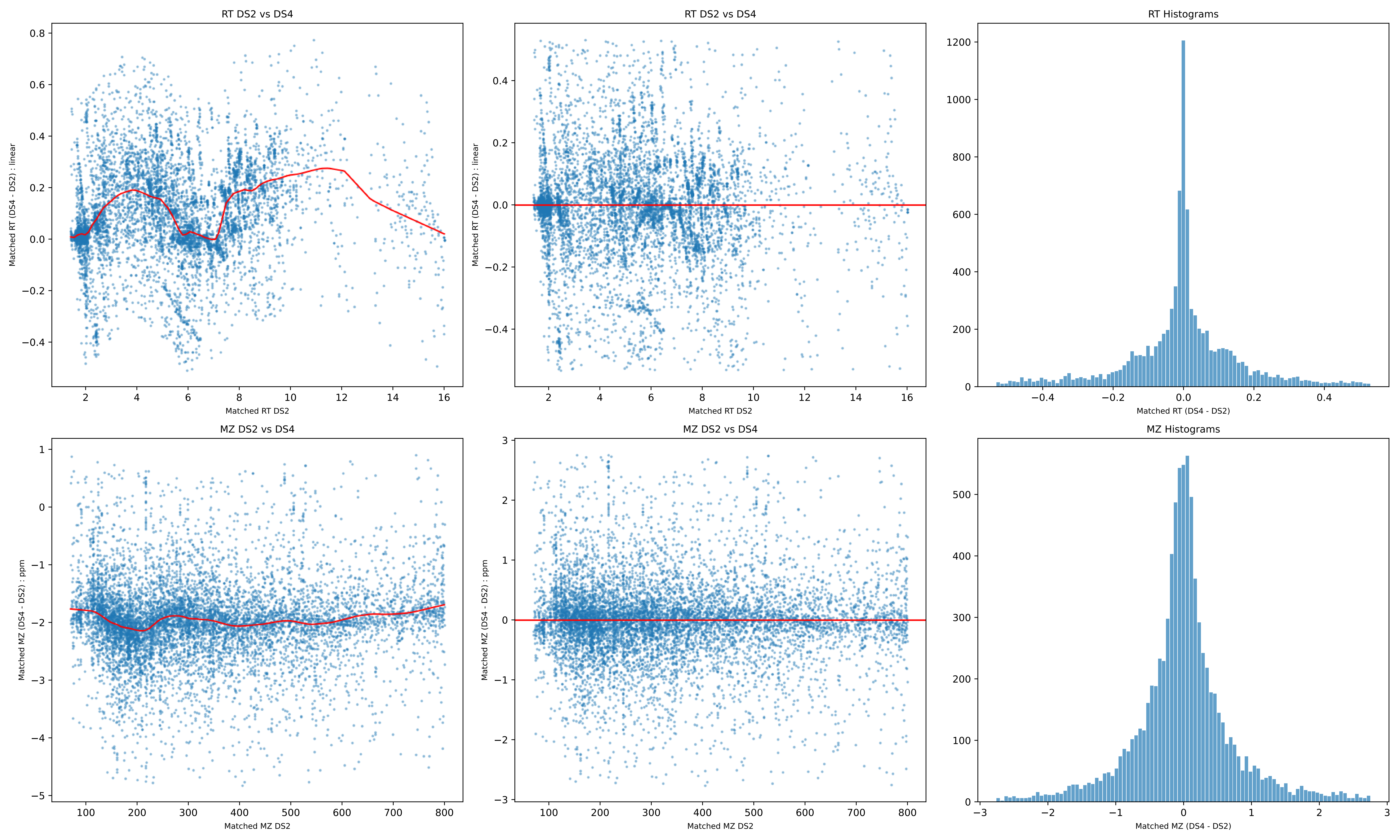
RT DS2 vs DS4
Matched RT (DS4 - DS2) : linear
Matched RT DS2
RT DS2 vs DS4
Matched RT (DS4 - DS2) : linear
Matched RT DS2
RT Histograms
Matched RT (DS4 - DS2)
MZ DS2 vs DS4
Matched MZ (DS4 - DS2) : ppm
Matched MZ DS2
MZ DS2 vs DS4
Matched MZ (DS4 - DS2) : ppm
Matched MZ DS2
MZ Histograms
Matched MZ (DS4 - DS2)

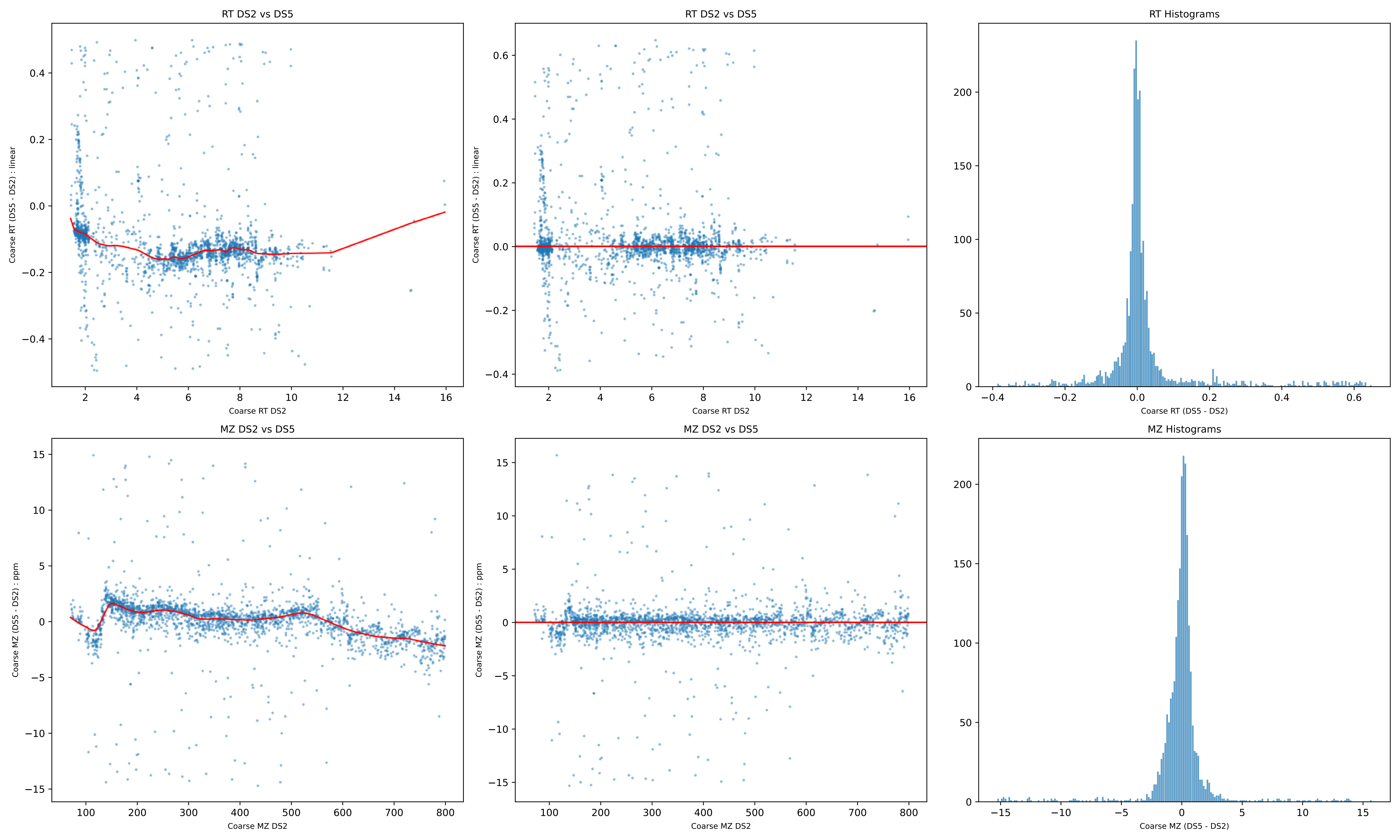

RT DS2 vs DS5
Coarse RT (DS5 - DS2) : linear
Coarse RT DS2
RT DS2 vs DS5
Coarse RT (DS5 - DS2) : linear
Coarse RT DS2
RT Histograms
Coarse RT (DS5 - DS2)
MZ DS2 vs DS5
Coarse MZ (DS5 - DS2) : ppm
Coarse MZ DS2
MZ DS2 vs DS5
Coarse MZ (DS5 - DS2) : ppm
Coarse MZ DS2
MZ Histograms
Coarse MZ (DS5 - DS2)

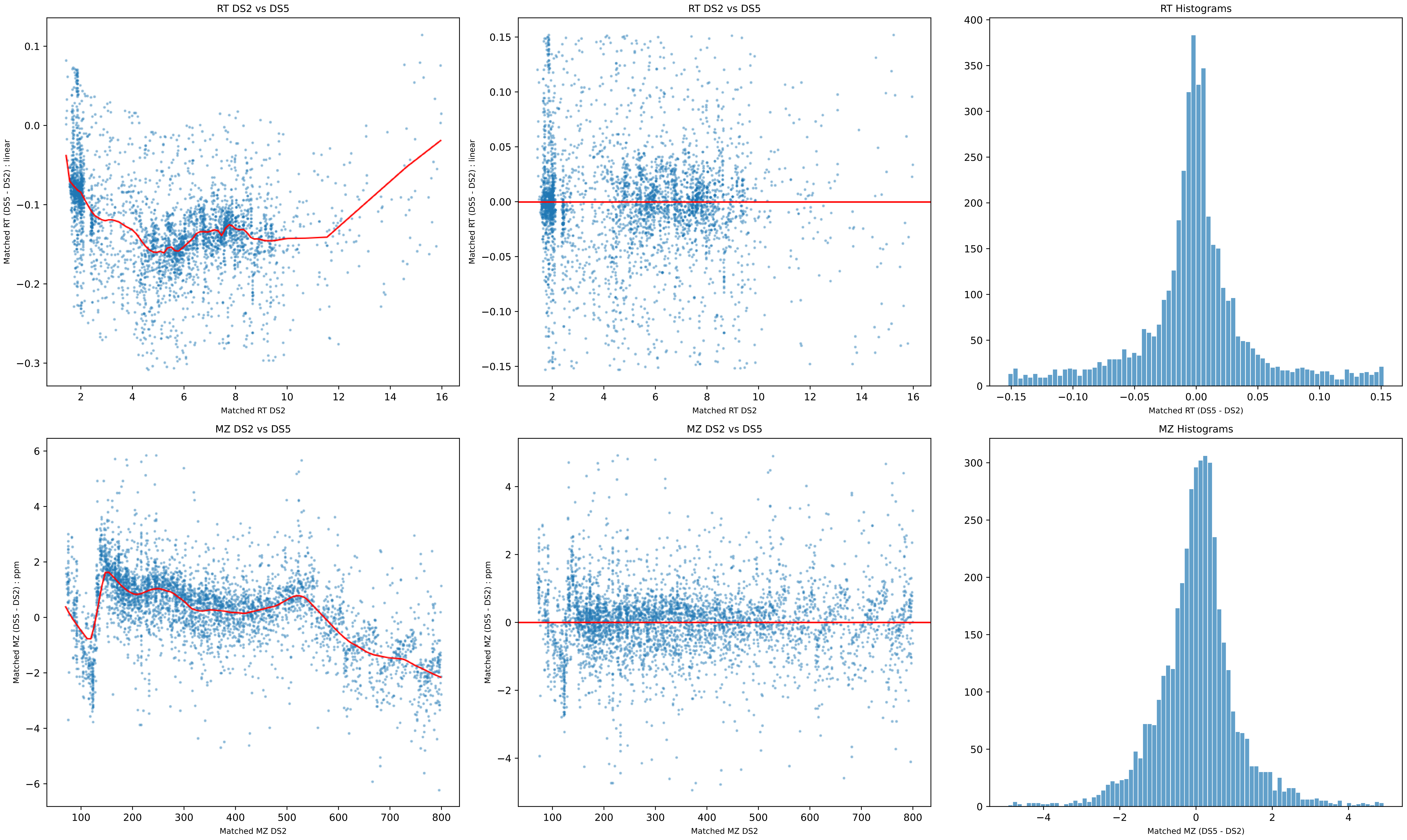
RT DS2 vs DS5
Matched RT (DS5 - DS2) : linear
Matched RT DS2
RT DS2 vs DS5
Matched RT (DS5 - DS2) : linear
Matched RT DS2
RT Histograms
Matched RT (DS5 - DS2)
MZ DS2 vs DS5
Matched MZ (DS5 - DS2) : ppm
Matched MZ DS2
MZ DS2 vs DS5
Matched MZ (DS5 - DS2) : ppm
Matched MZ DS2
MZ Histograms
Matched MZ (DS5 - DS2)

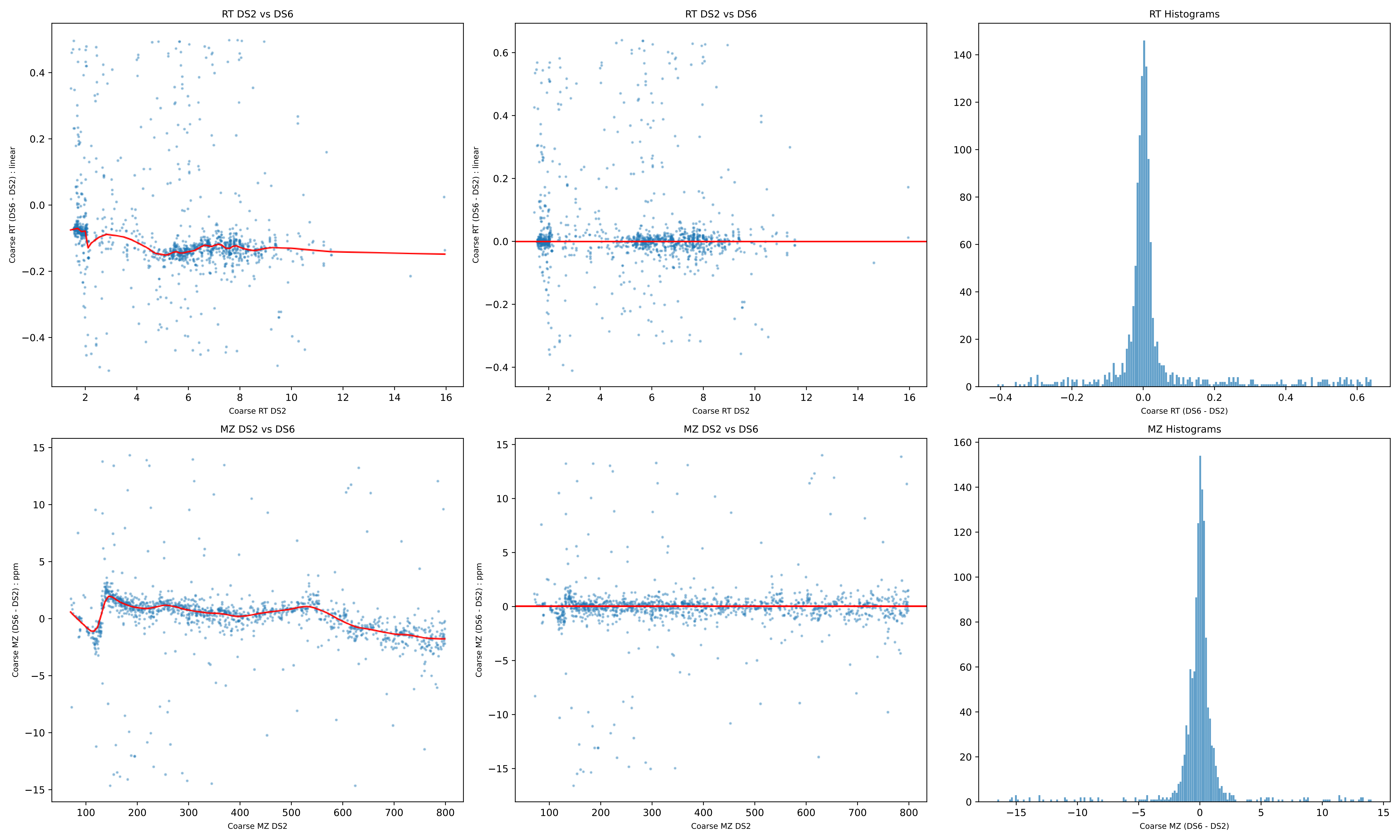
RT DS2 vs DS6
Coarse RT (DS6 - DS2) : linear
Coarse RT DS2
RT DS2 vs DS6
Coarse RT (DS6 - DS2) : linear
Coarse RT DS2
RT Histograms
Coarse RT (DS6 - DS2)
MZ DS2 vs DS6
Coarse MZ (DS6 - DS2) : ppm
Coarse MZ DS2
MZ DS2 vs DS6
Coarse MZ (DS6 - DS2) : ppm
Coarse MZ DS2
MZ Histograms
Coarse MZ (DS6 - DS2)

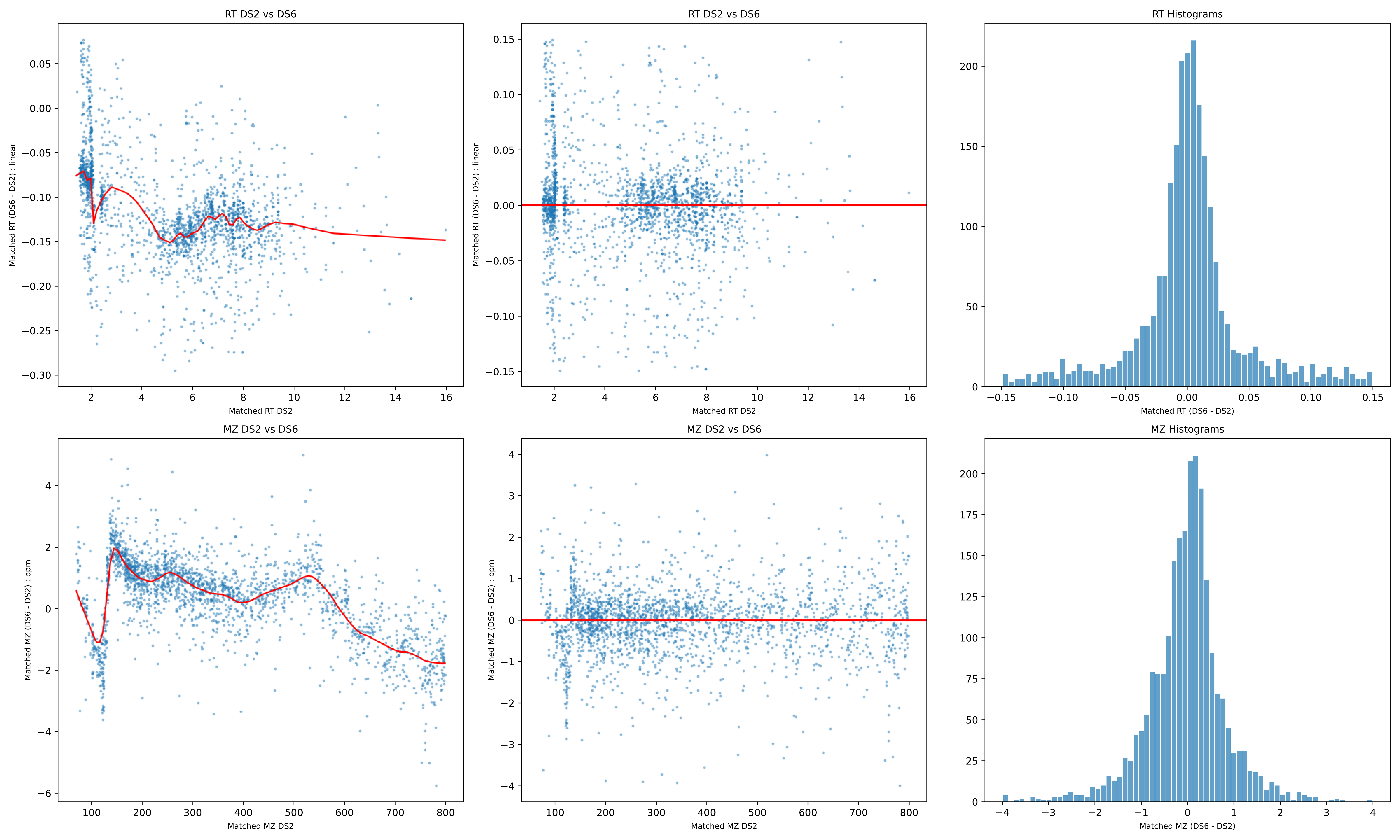
RT DS2 vs DS6
Matched RT (DS6 - DS2) : linear
Matched RT DS2
RT DS2 vs DS6
Matched RT (DS6 - DS2) : linear
Matched RT DS2
RT Histograms
Matched RT (DS6 - DS2)
MZ DS2 vs DS6
Matched MZ (DS6 - DS2) : ppm
Matched MZ DS2
MZ DS2 vs DS6
Matched MZ (DS6 - DS2) : ppm
Matched MZ DS2
MZ Histograms
Matched MZ (DS6 - DS2)

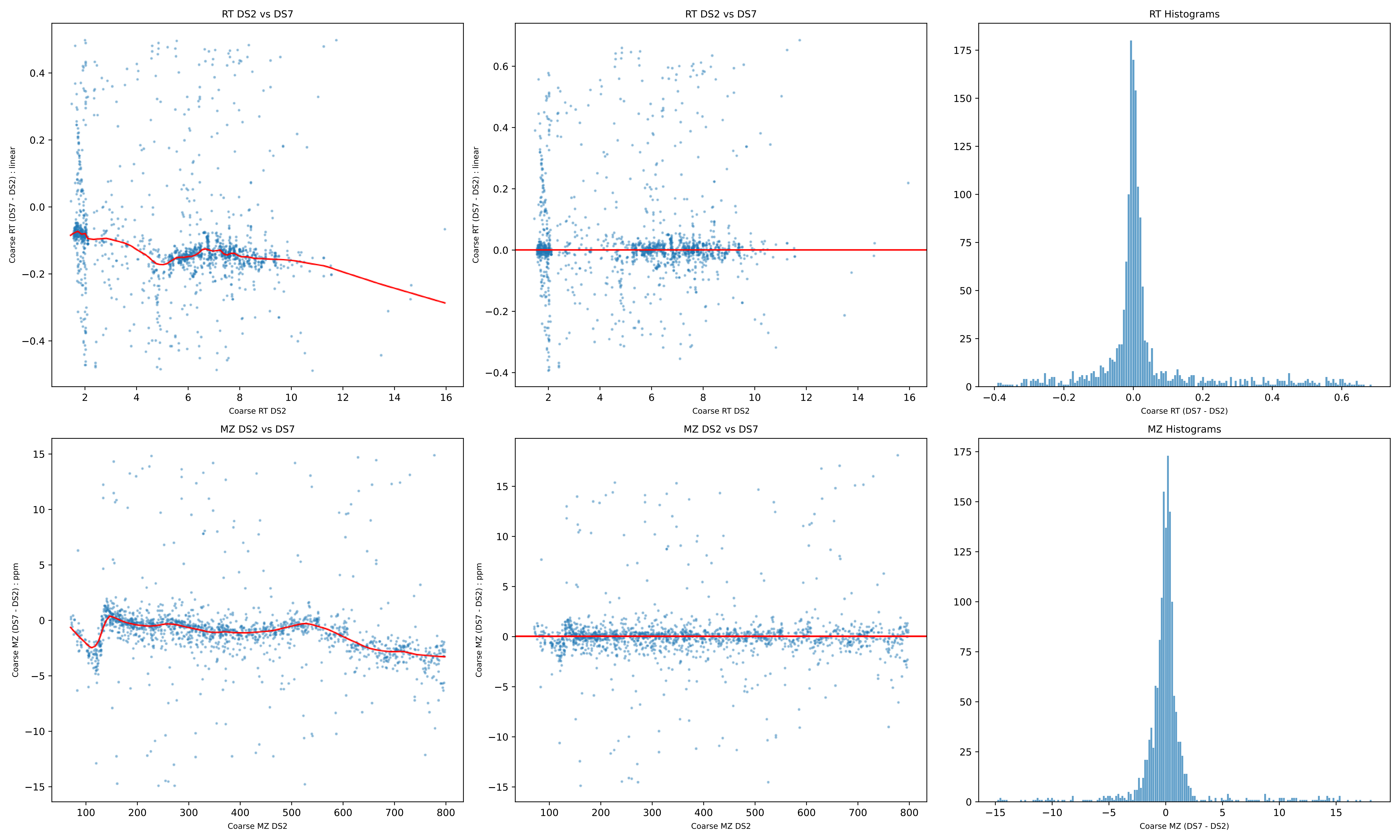

RT DS2 vs DS7
Coarse RT (DS7 - DS2) : linear
Coarse RT DS2
RT DS2 vs DS7
Coarse RT (DS7 - DS2) : linear
Coarse RT DS2
RT Histograms
Coarse RT (DS7 - DS2)
MZ DS2 vs DS7
Coarse MZ (DS7 - DS2) : ppm
Coarse MZ DS2
MZ DS2 vs DS7
Coarse MZ (DS7 - DS2) : ppm
Coarse MZ DS2
MZ Histograms
Coarse MZ (DS7 - DS2)

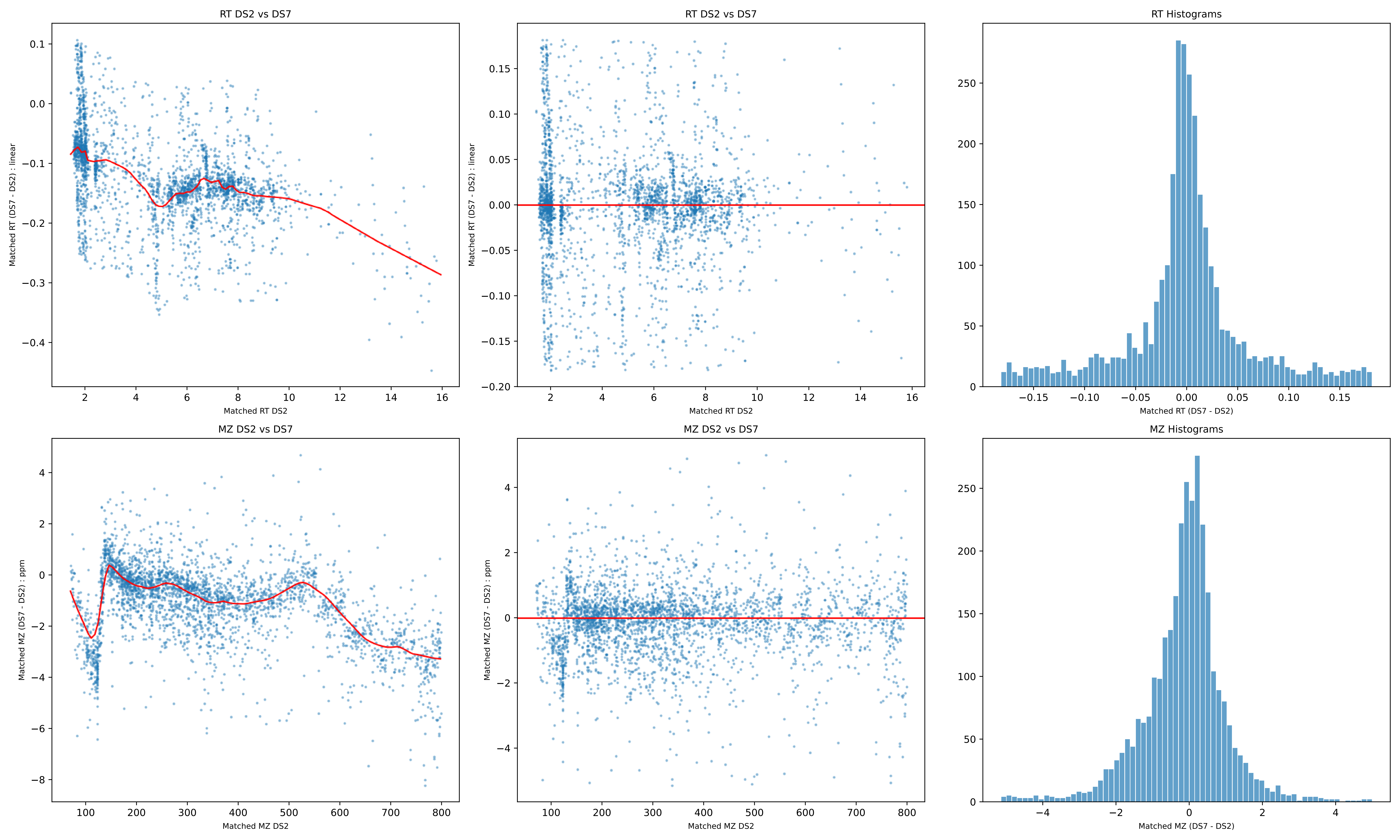
RT DS2 vs DS7
Matched RT (DS7 - DS2) : linear
Matched RT DS2
RT DS2 vs DS7
Matched RT (DS7 - DS2) : linear
Matched RT DS2
RT Histograms
Matched RT (DS7 - DS2)
MZ DS2 vs DS7
Matched MZ (DS7 - DS2) : ppm
Matched MZ DS2
MZ DS2 vs DS7
Matched MZ (DS7 - DS2) : ppm
Matched MZ DS2
MZ Histograms
Matched MZ (DS7 - DS2)

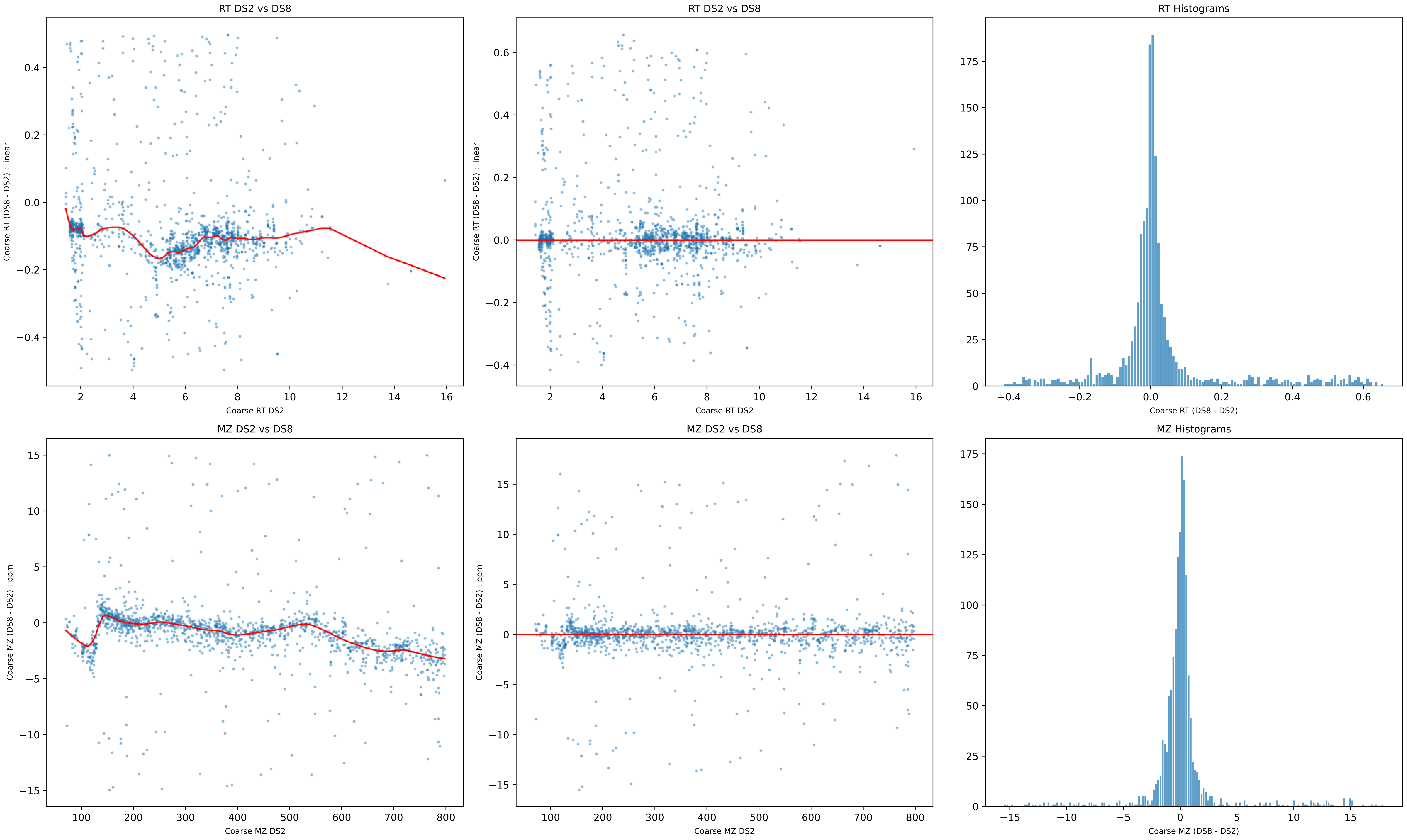

RT DS2 vs DS8
Coarse RT (DS8 - DS2) : linear
Coarse RT DS2
RT DS2 vs DS8
Coarse RT (DS8 - DS2) : linear
Coarse RT DS2
RT Histograms
Coarse RT (DS8 - DS2)
MZ DS2 vs DS8
Coarse MZ (DS8 - DS2) : ppm
Coarse MZ DS2
MZ DS2 vs DS8
Coarse MZ (DS8 - DS2) : ppm
Coarse MZ DS2
MZ Histograms
Coarse MZ (DS8 - DS2)

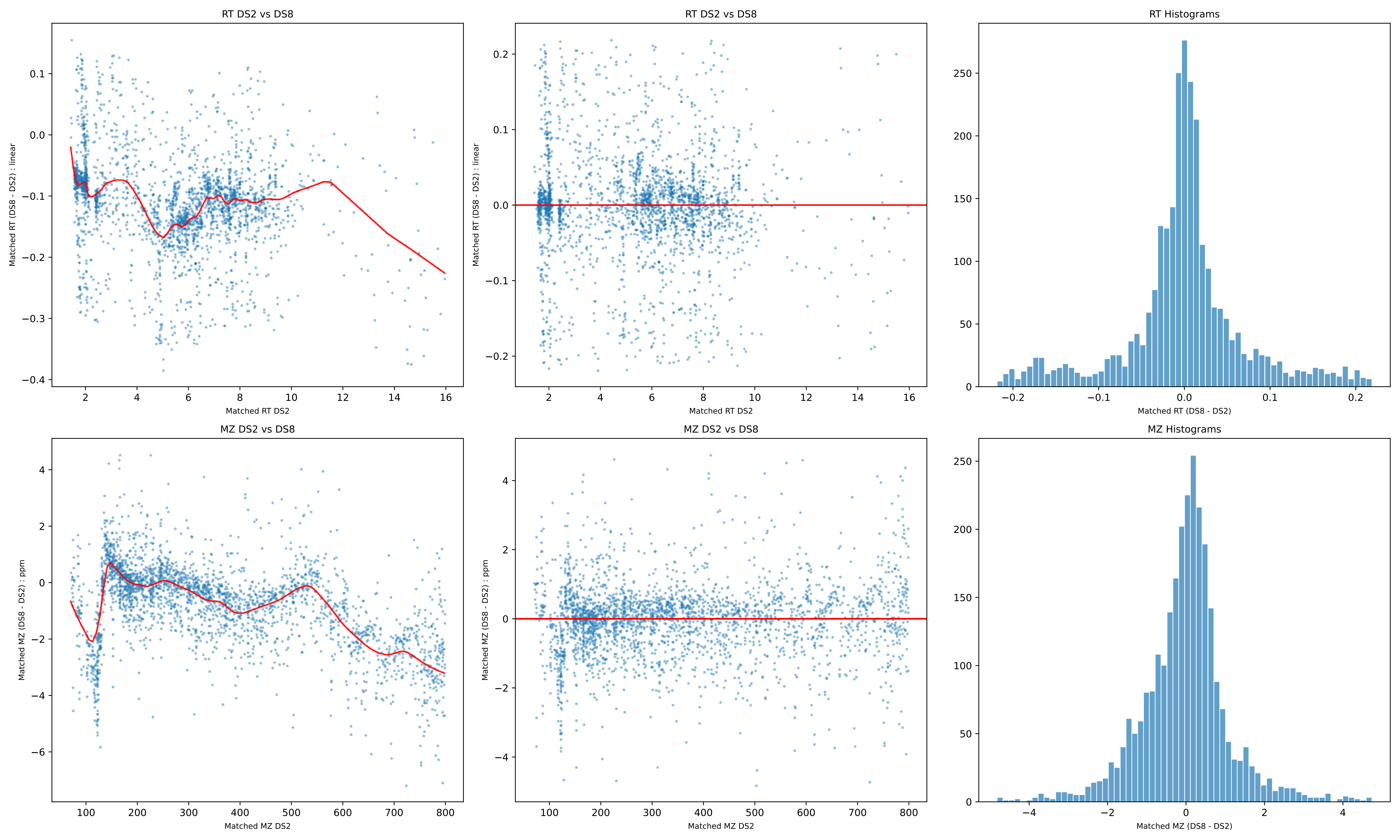
RT DS2 vs DS8
Matched RT (DS8 - DS2) : linear
Matched RT DS2
RT DS2 vs DS8
Matched RT (DS8 - DS2) : linear
Matched RT DS2
RT Histograms
Matched RT (DS8 - DS2)
MZ DS2 vs DS8
Matched MZ (DS8 - DS2) : ppm
Matched MZ DS2
MZ DS2 vs DS8
Matched MZ (DS8 - DS2) : ppm
Matched MZ DS2
MZ Histograms
Matched MZ (DS8 - DS2)

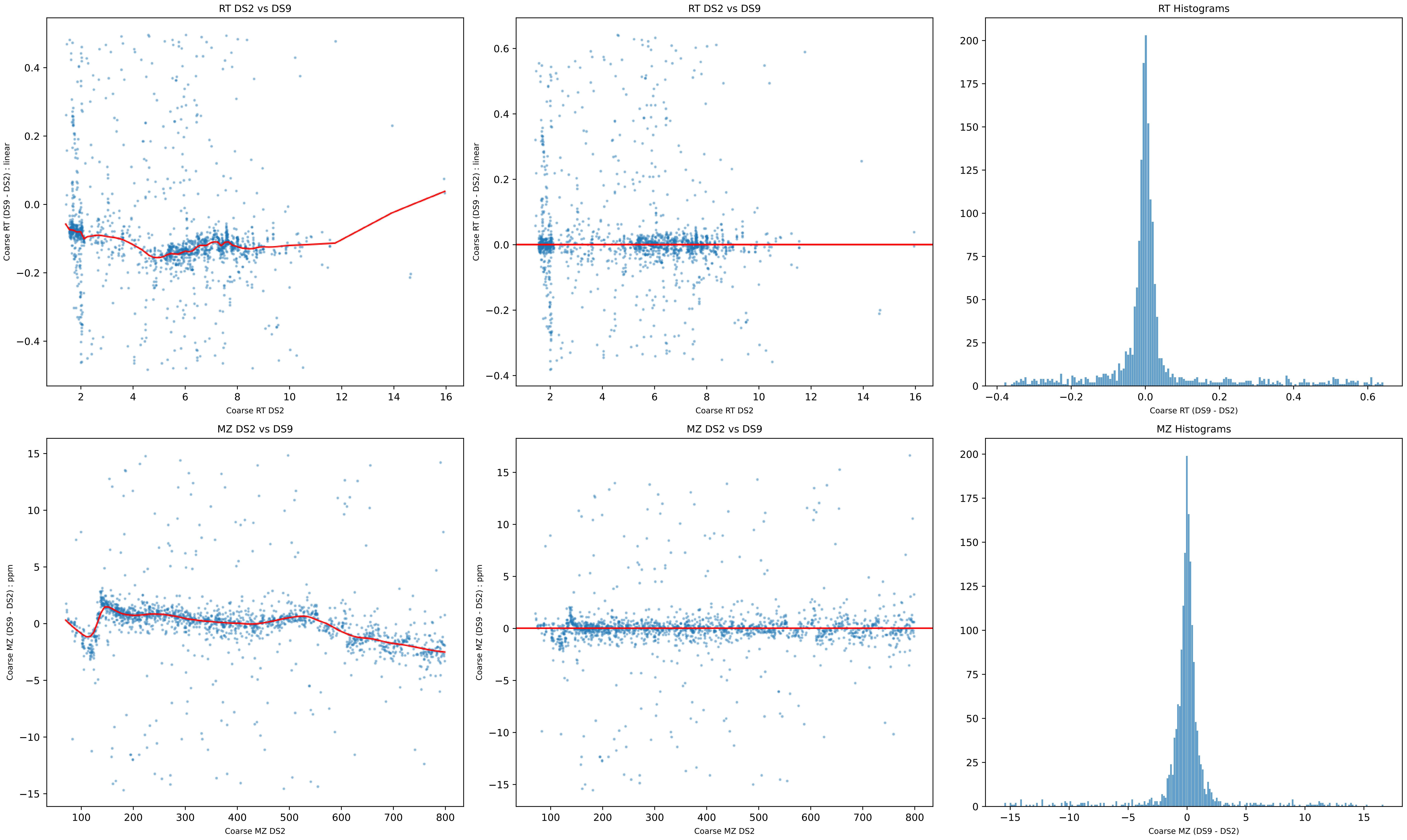

RT DS2 vs DS9
Coarse RT (DS9 - DS2) : linear
Coarse RT DS2
RT DS2 vs DS9
Coarse RT (DS9 - DS2) : linear
Coarse RT DS2
RT Histograms
Coarse RT (DS9 - DS2)
MZ DS2 vs DS9
Coarse MZ (DS9 - DS2) : ppm
Coarse MZ DS2
MZ DS2 vs DS9
Coarse MZ (DS9 - DS2) : ppm
Coarse MZ DS2
MZ Histograms
Coarse MZ (DS9 - DS2)

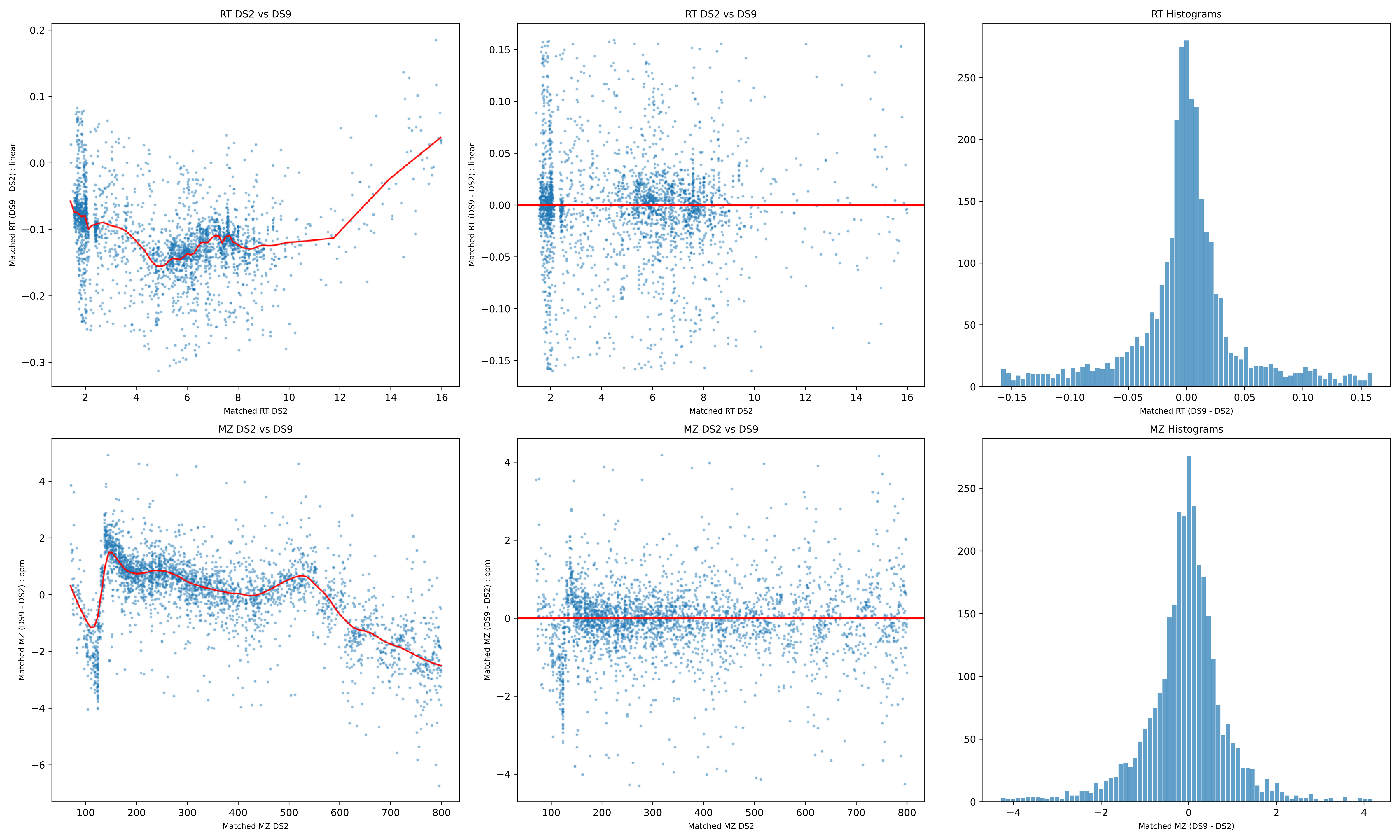
RT DS2 vs DS9
Matched RT (DS9 - DS2) : linear
Matched RT DS2
RT DS2 vs DS9
Matched RT (DS9 - DS2) : linear
Matched RT DS2
RT Histograms
Matched RT (DS9 - DS2)
MZ DS2 vs DS9
Matched MZ (DS9 - DS2) : ppm
Matched MZ DS2
MZ DS2 vs DS9
Matched MZ (DS9 - DS2) : ppm
Matched MZ DS2
MZ Histograms
Matched MZ (DS9 - DS2)

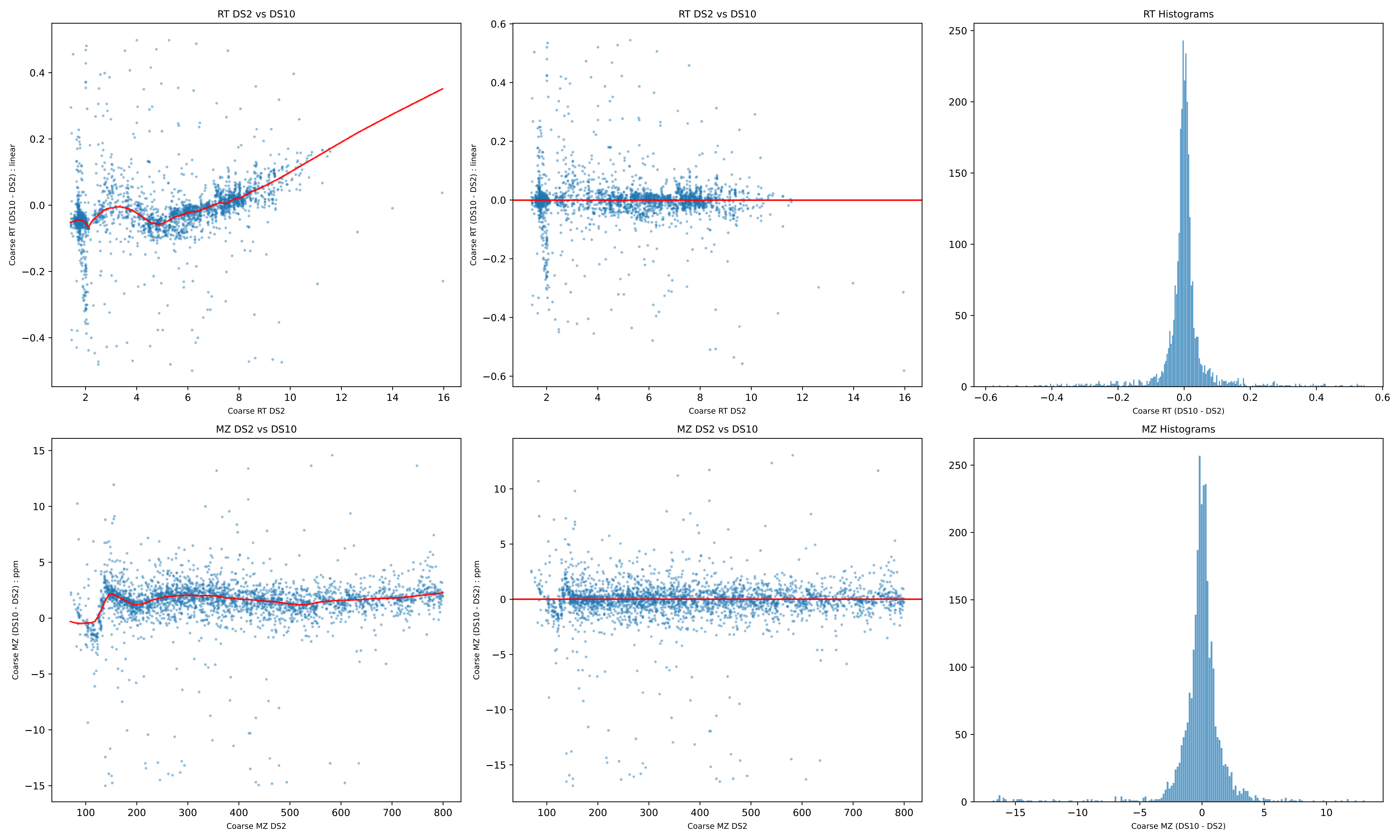
RT DS2 vs DS10
Coarse RT (DS10 - DS2) : linear
Coarse RT DS2
RT DS2 vs DS10
Coarse RT (DS10 - DS2) : linear
Coarse RT DS2
RT Histograms
Coarse RT (DS10 - DS2)
MZ DS2 vs DS10
Coarse MZ (DS10 - DS2) : ppm
Coarse MZ DS2
MZ DS2 vs DS10
Coarse MZ (DS10 - DS2) : ppm
Coarse MZ DS2
MZ Histograms
Coarse MZ (DS10 - DS2)

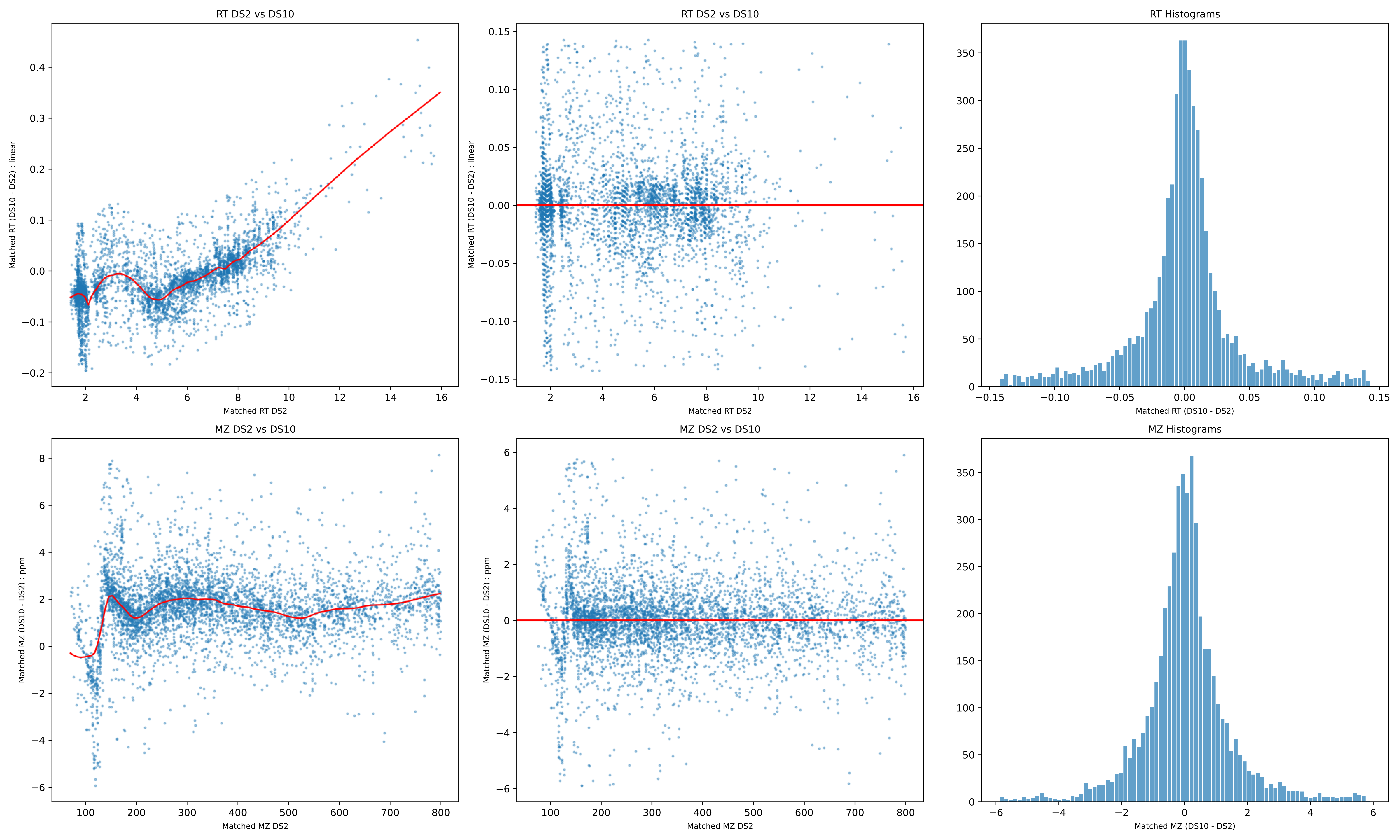
RT DS2 vs DS10
Matched RT (DS10 - DS2) : linear
Matched RT DS2
RT DS2 vs DS10
Matched RT (DS10 - DS2) : linear
Matched RT DS2
RT Histograms
Matched RT (DS10 - DS2)
MZ DS2 vs DS10
Matched MZ (DS10 - DS2) : ppm
Matched MZ DS2
MZ DS2 vs DS10
Matched MZ (DS10 - DS2) : ppm
Matched MZ DS2
MZ Histograms
Matched MZ (DS10 - DS2)

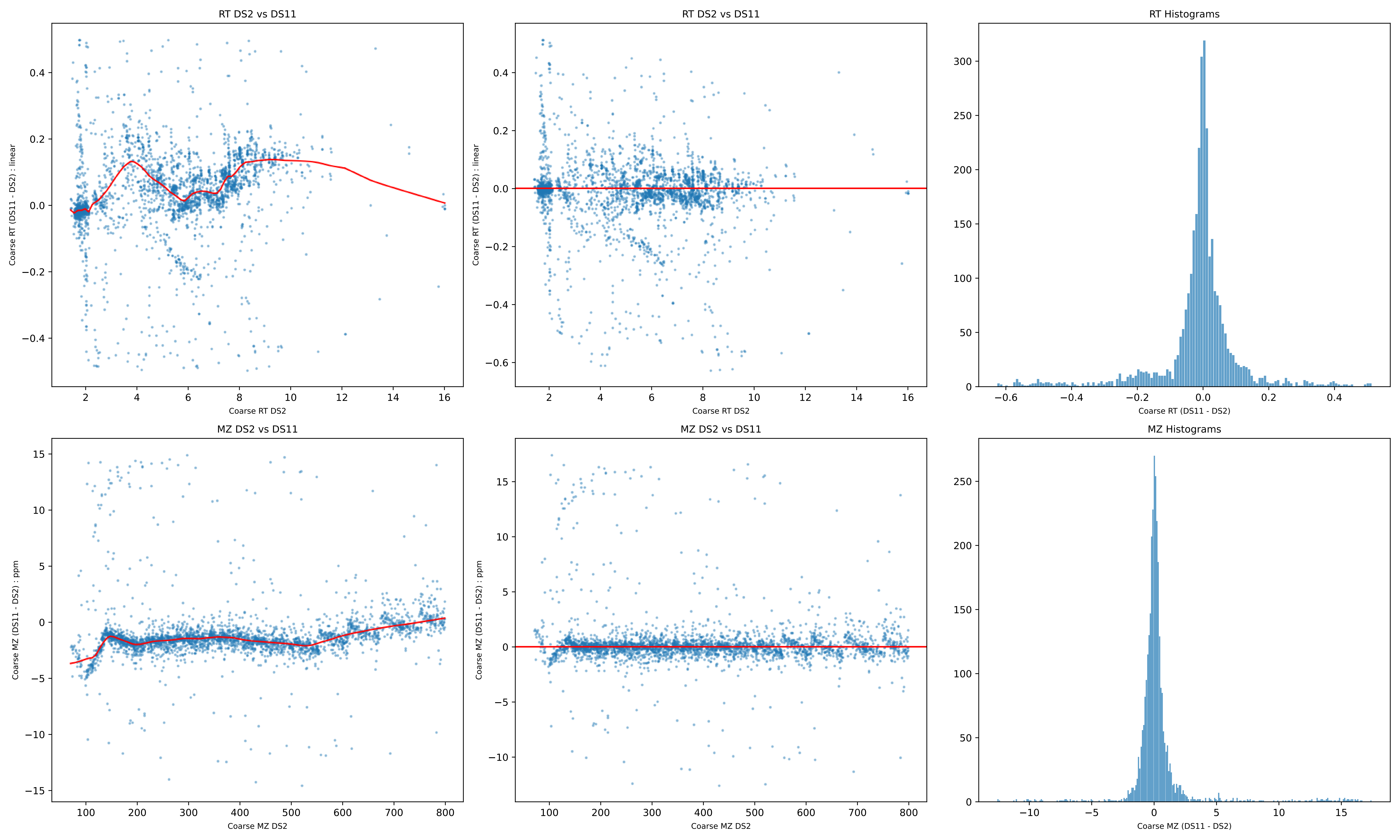
RT DS2 vs DS11
Coarse RT (DS11 - DS2) : linear
Coarse RT DS2
RT DS2 vs DS11
Coarse RT (DS11 - DS2) : linear
Coarse RT DS2
RT Histograms
Coarse RT (DS11 - DS2)
MZ DS2 vs DS11
Coarse MZ (DS11 - DS2) : ppm
Coarse MZ DS2
MZ DS2 vs DS11
Coarse MZ (DS11 - DS2) : ppm
Coarse MZ DS2
MZ Histograms
Coarse MZ (DS11 - DS2)

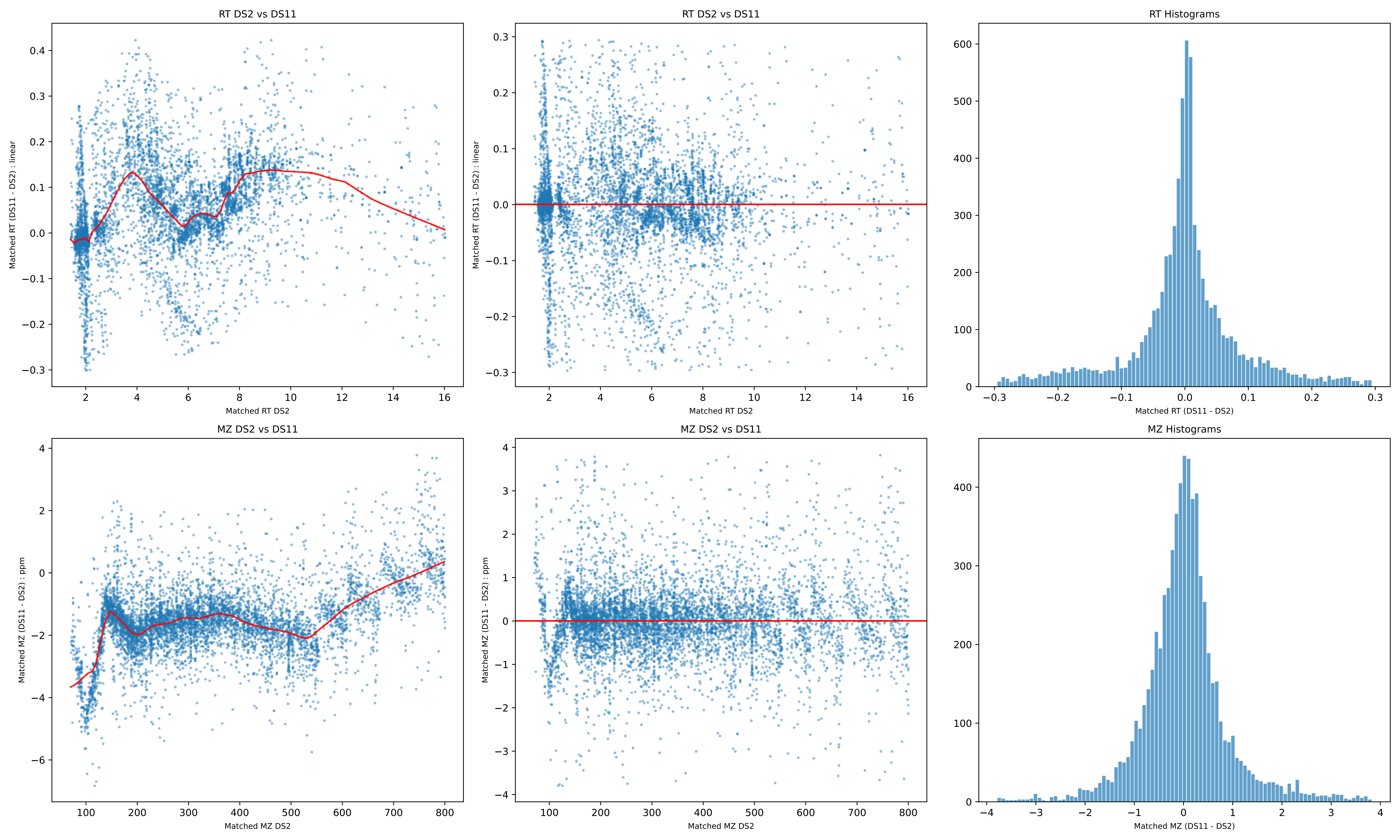
RT DS2 vs DS11
Matched RT (DS11 - DS2) : linear
Matched RT DS2
RT DS2 vs DS11
Matched RT (DS11 - DS2) : linear
Matched RT DS2
RT Histograms
Matched RT (DS11 - DS2)
MZ DS2 vs DS11
Matched MZ (DS11 - DS2) : ppm
Matched MZ DS2
MZ DS2 vs DS11
Matched MZ (DS11 - DS2) : ppm
Matched MZ DS2
MZ Histograms
Matched MZ (DS11 - DS2)

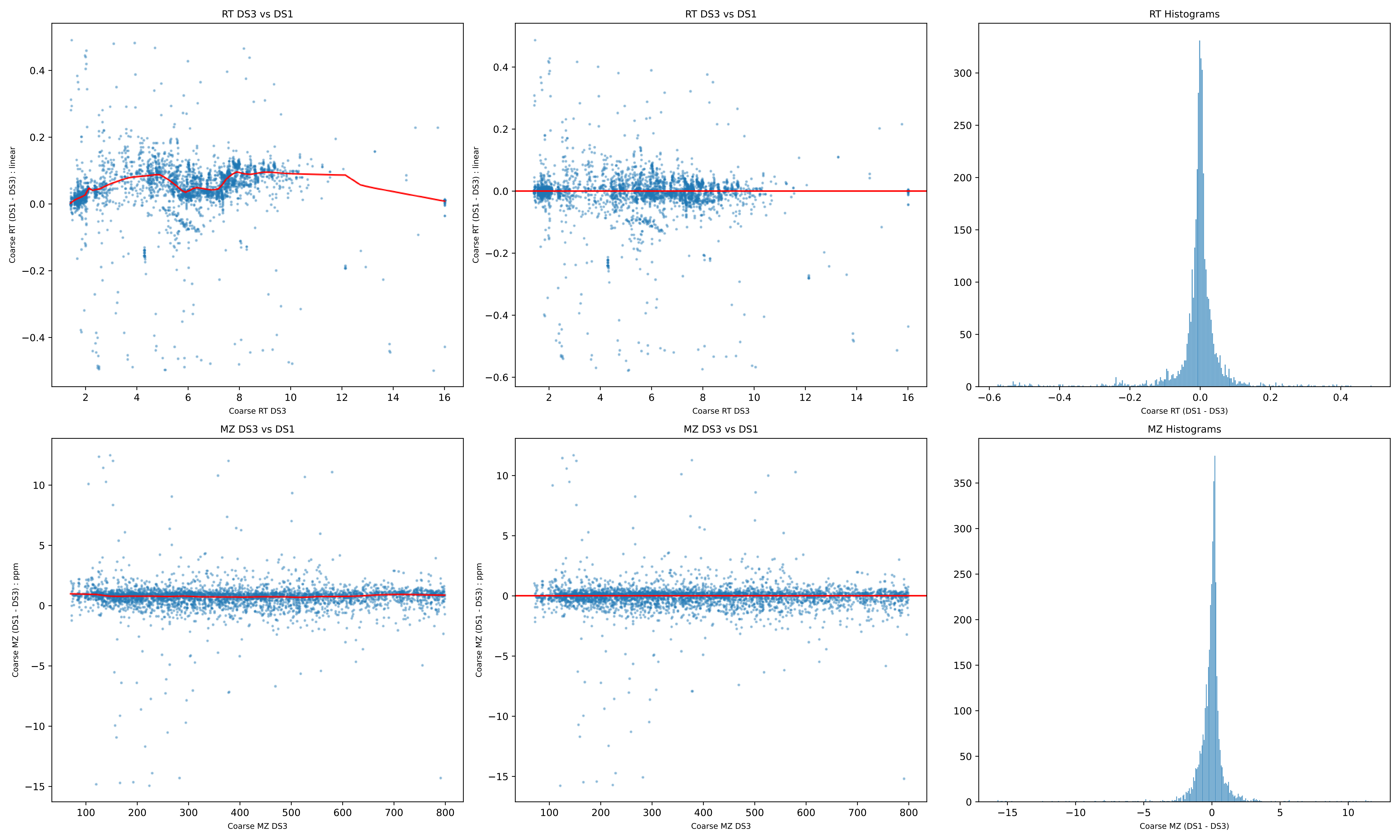

RT DS3 vs DS1
Coarse RT (DS1 - DS3) : linear
Coarse RT DS3
RT DS3 vs DS1
Coarse RT (DS1 - DS3) : linear
Coarse RT DS3
RT Histograms
Coarse RT (DS1 - DS3)
MZ DS3 vs DS1
Coarse MZ (DS1 - DS3) : ppm
Coarse MZ DS3
MZ DS3 vs DS1
Coarse MZ (DS1 - DS3) : ppm
Coarse MZ DS3
MZ Histograms
Coarse MZ (DS1 - DS3)

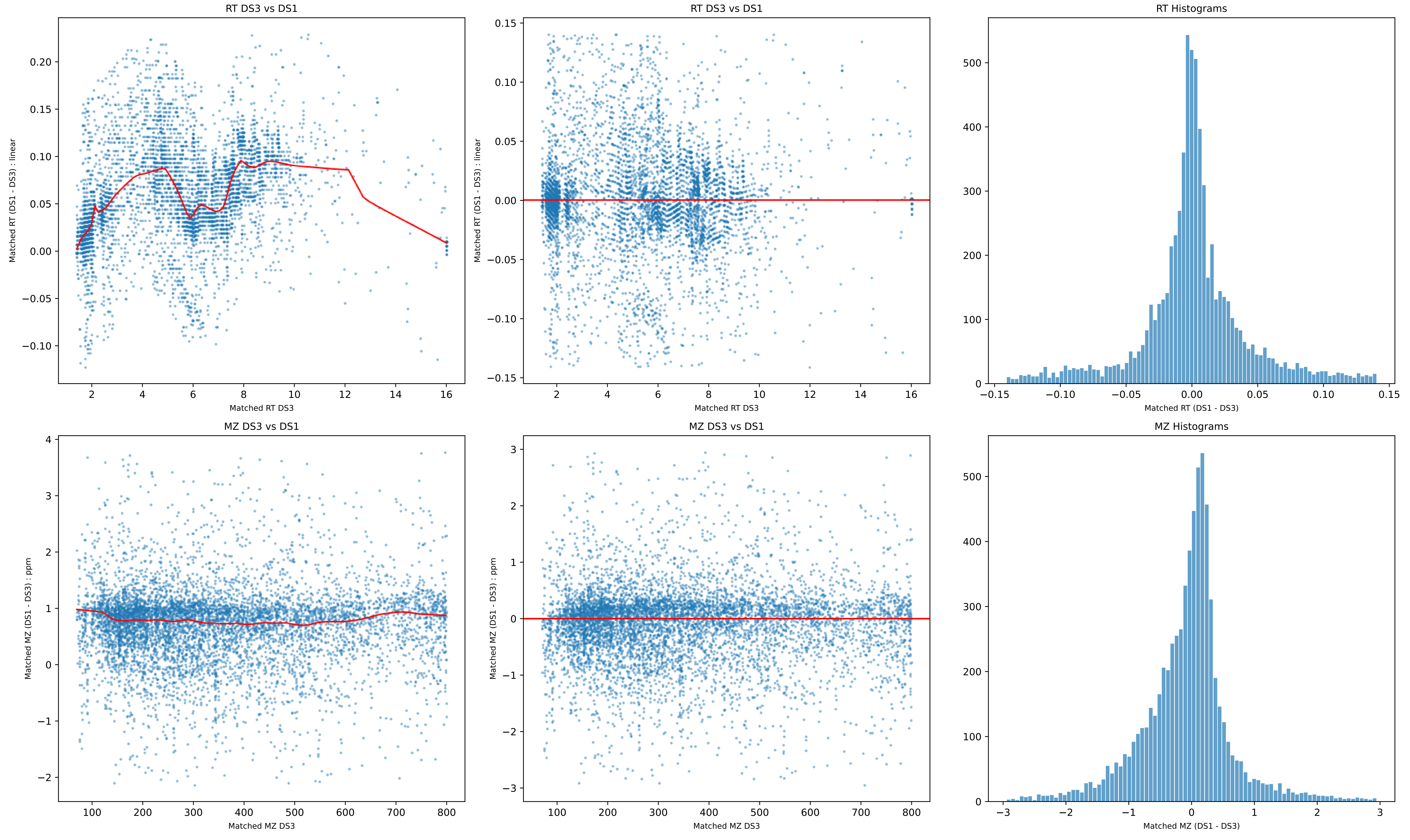

RT DS3 vs DS1
Matched RT (DS1 - DS3) : linear
Matched RT DS3
RT DS3 vs DS1
Matched RT (DS1 - DS3) : linear
Matched RT DS3
RT Histograms
Matched RT (DS1 - DS3)
MZ DS3 vs DS1
Matched MZ (DS1 - DS3) : ppm
Matched MZ DS3
MZ DS3 vs DS1
Matched MZ (DS1 - DS3) : ppm
Matched MZ DS3
MZ Histograms
Matched MZ (DS1 - DS3)

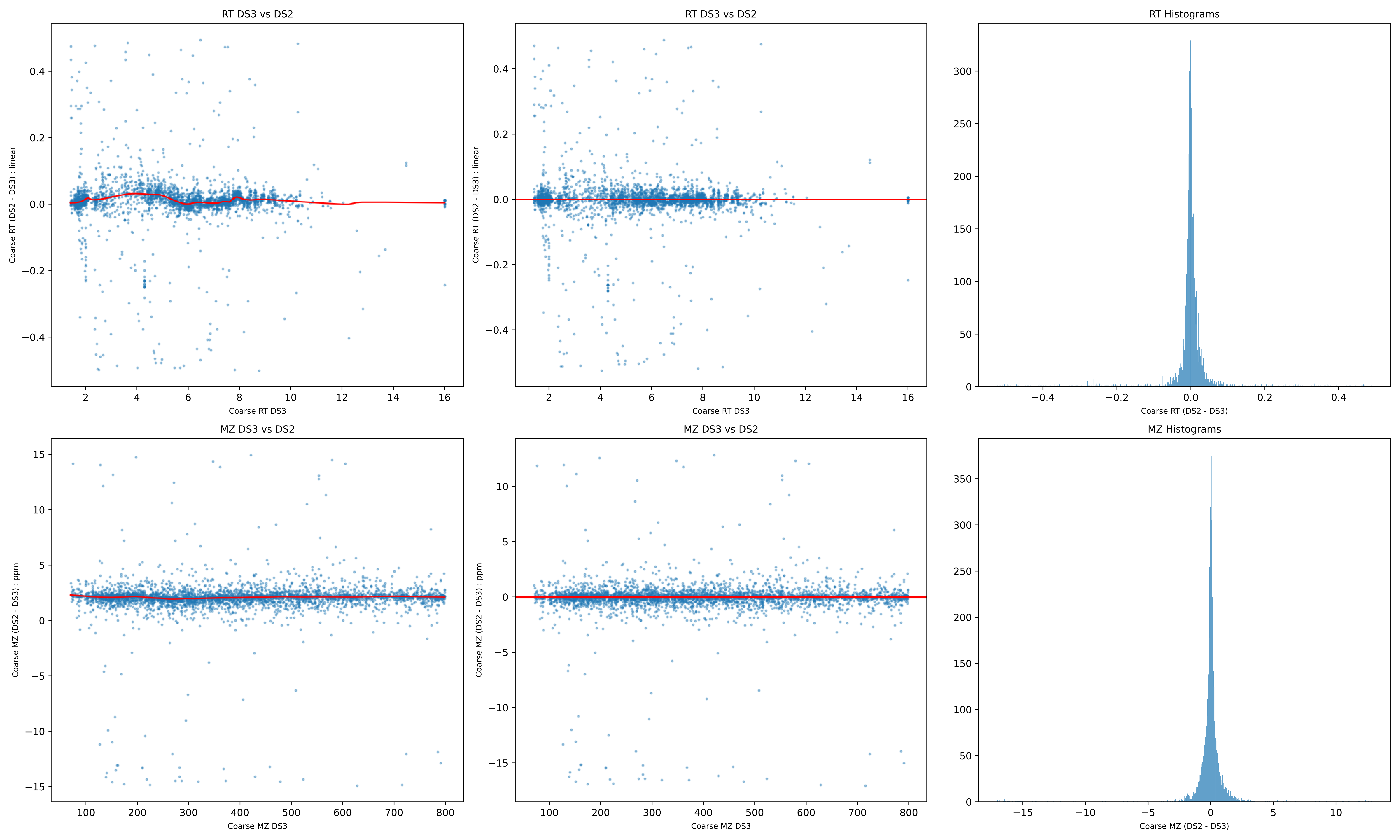
RT DS3 vs DS2
Coarse RT (DS2 - DS3) : linear
Coarse RT DS3
RT DS3 vs DS2
Coarse RT (DS2 - DS3) : linear
Coarse RT DS3
RT Histograms
Coarse RT (DS2 - DS3)
MZ DS3 vs DS2
Coarse MZ (DS2 - DS3) : ppm
Coarse MZ DS3
MZ DS3 vs DS2
Coarse MZ (DS2 - DS3) : ppm
Coarse MZ DS3
MZ Histograms
Coarse MZ (DS2 - DS3)

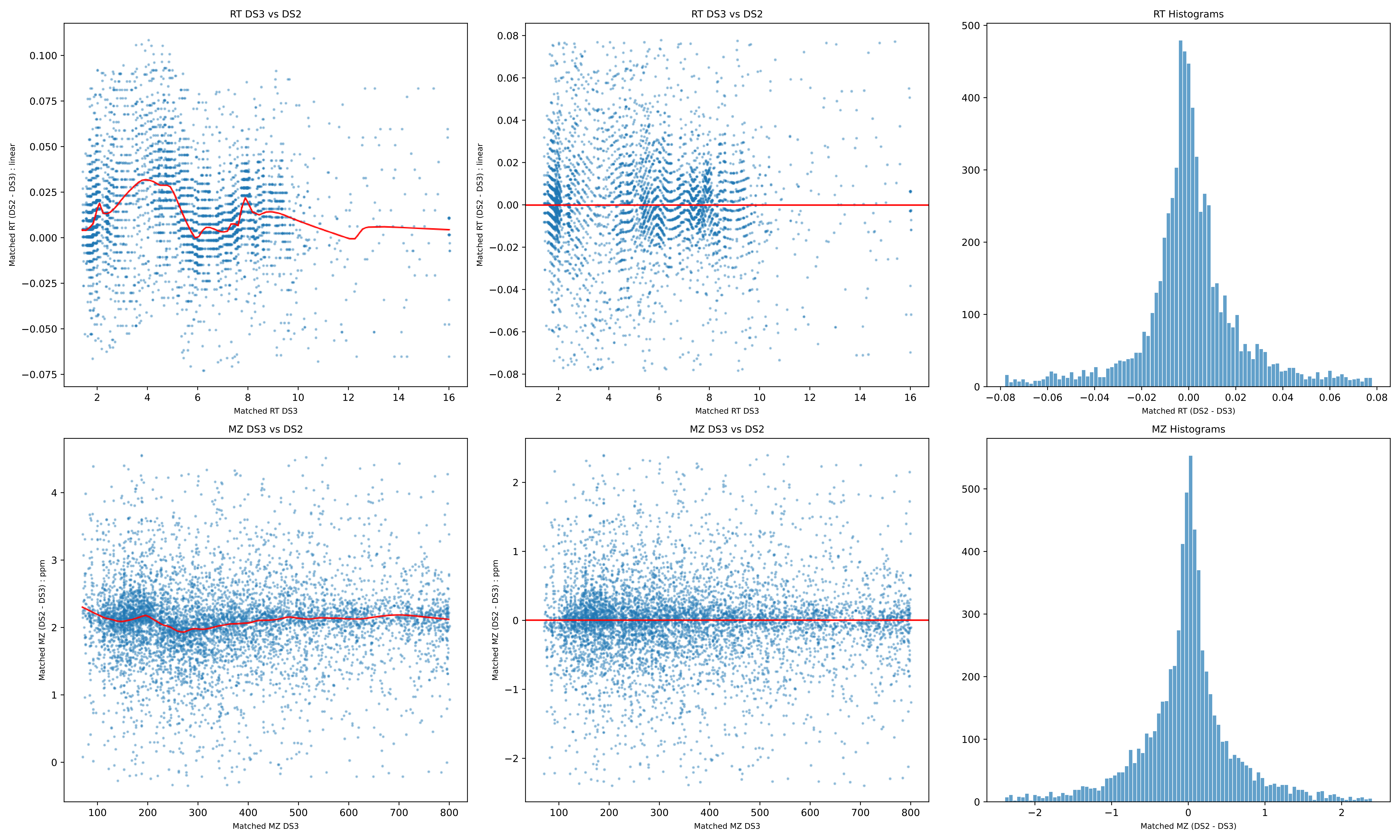
RT DS3 vs DS2
Matched RT (DS2 - DS3) : linear
Matched RT DS3
RT DS3 vs DS2
Matched RT (DS2 - DS3) : linear
Matched RT DS3
RT Histograms
Matched RT (DS2 - DS3)
MZ DS3 vs DS2
Matched MZ (DS2 - DS3) : ppm
Matched MZ DS3
MZ DS3 vs DS2
Matched MZ (DS2 - DS3) : ppm
Matched MZ DS3
MZ Histograms
Matched MZ (DS2 - DS3)

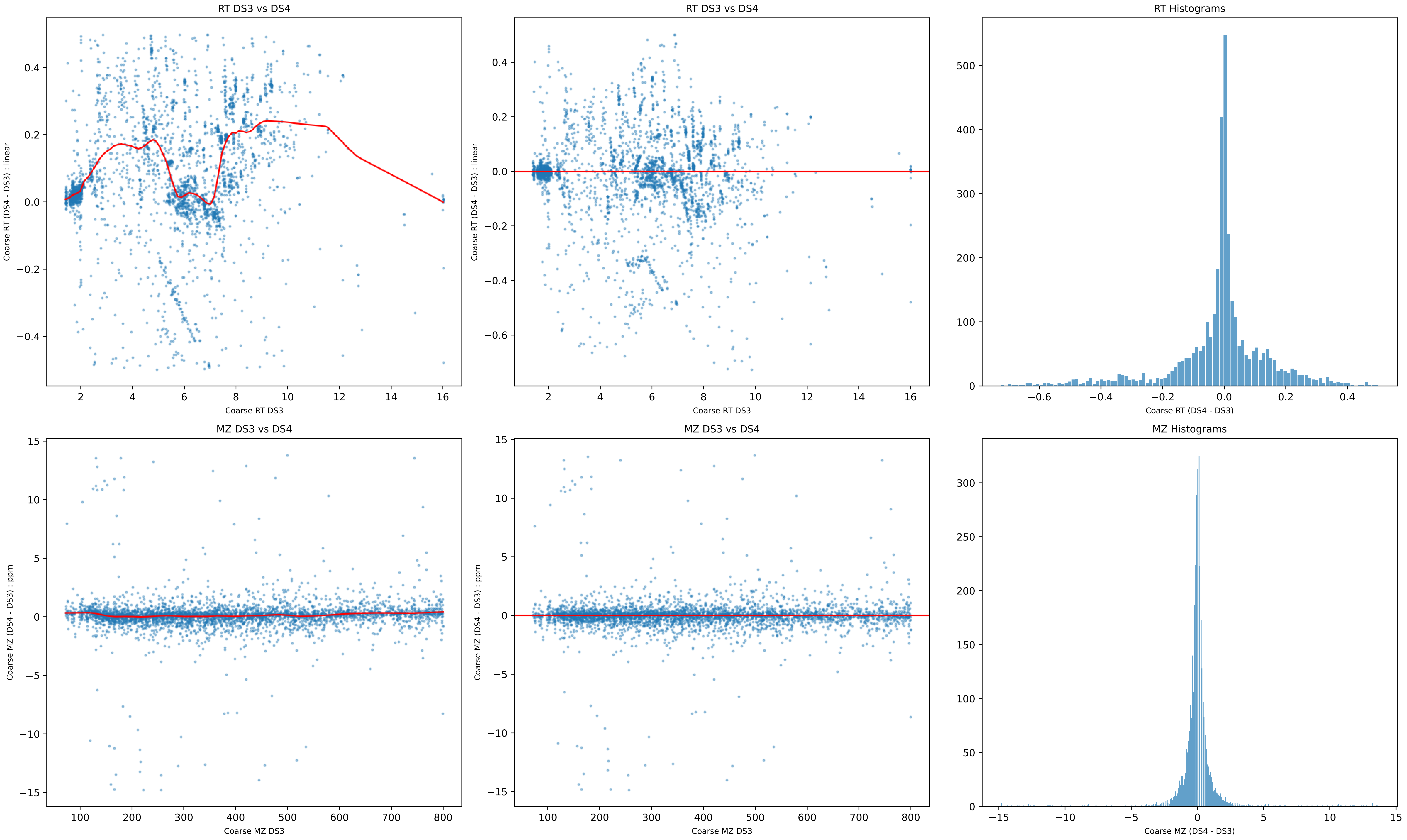
RT DS3 vs DS4
Coarse RT (DS4 - DS3) : linear
Coarse RT DS3
RT DS3 vs DS4
Coarse RT (DS4 - DS3) : linear
Coarse RT DS3
RT Histograms
Coarse RT (DS4 - DS3)
MZ DS3 vs DS4
Coarse MZ (DS4 - DS3) : ppm
Coarse MZ DS3
MZ DS3 vs DS4
Coarse MZ (DS4 - DS3) : ppm
Coarse MZ DS3
MZ Histograms
Coarse MZ (DS4 - DS3)

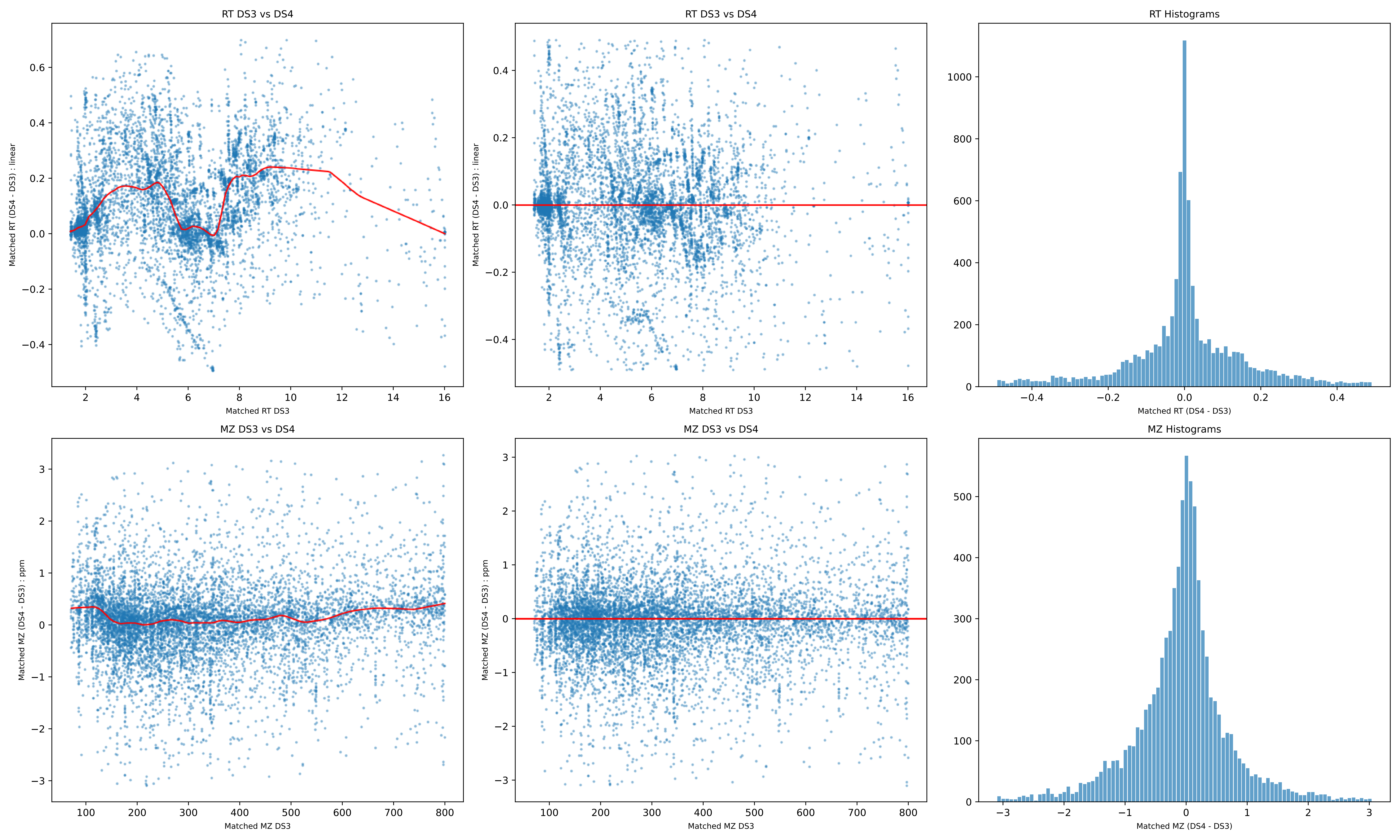
RT DS3 vs DS4
Matched RT (DS4 - DS3) : linear
Matched RT DS3
RT DS3 vs DS4
Matched RT (DS4 - DS3) : linear
Matched RT DS3
RT Histograms
Matched RT (DS4 - DS3)
MZ DS3 vs DS4
Matched MZ (DS4 - DS3) : ppm
Matched MZ DS3
MZ DS3 vs DS4
Matched MZ (DS4 - DS3) : ppm
Matched MZ DS3
MZ Histograms
Matched MZ (DS4 - DS3)

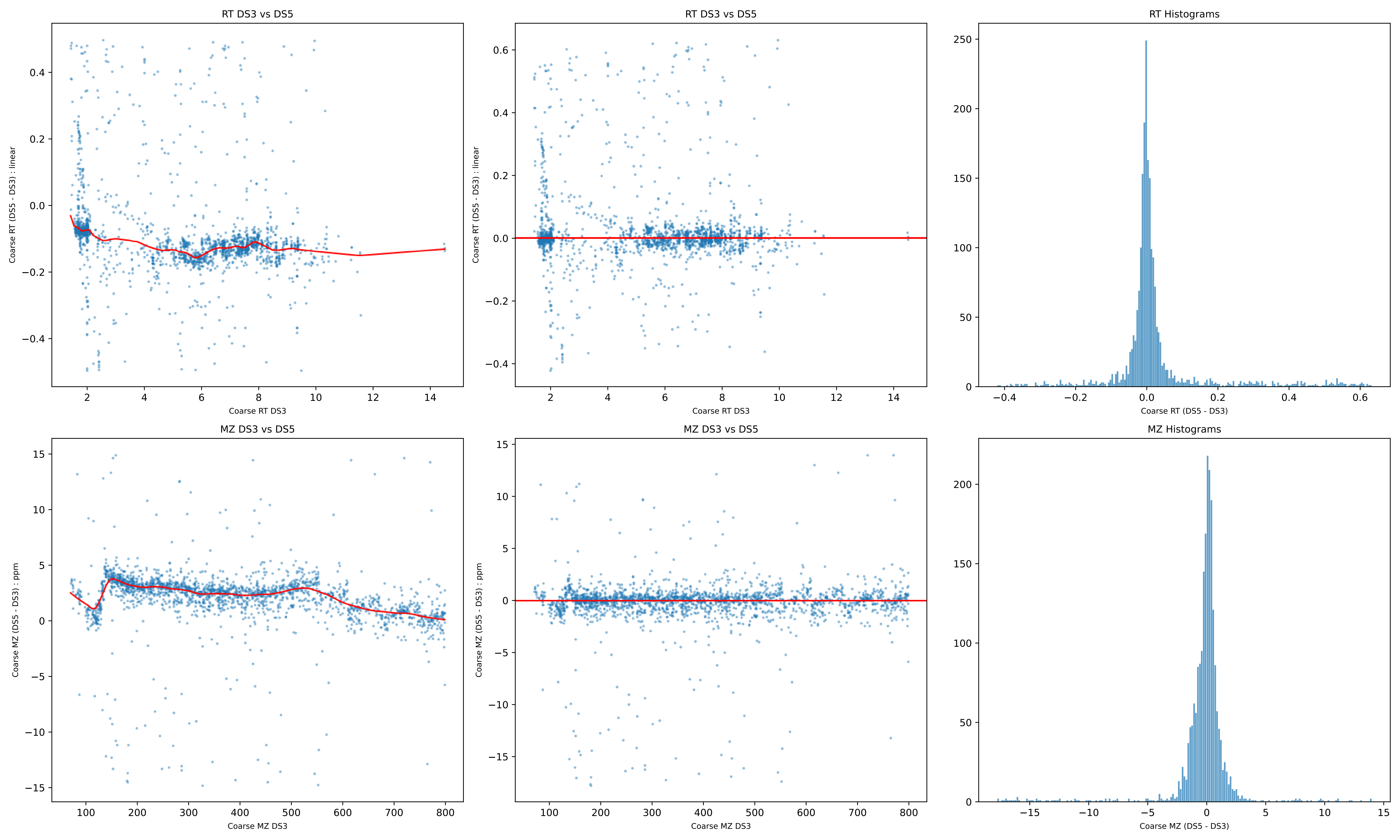
RT DS3 vs DS5
Coarse RT (DS5 - DS3) : linear
Coarse RT DS3
RT DS3 vs DS5
Coarse RT (DS5 - DS3) : linear
Coarse RT DS3
RT Histograms
Coarse RT (DS5 - DS3)
MZ DS3 vs DS5
Coarse MZ (DS5 - DS3) : ppm
Coarse MZ DS3
MZ DS3 vs DS5
Coarse MZ (DS5 - DS3) : ppm
Coarse MZ DS3
MZ Histograms
Coarse MZ (DS5 - DS3)

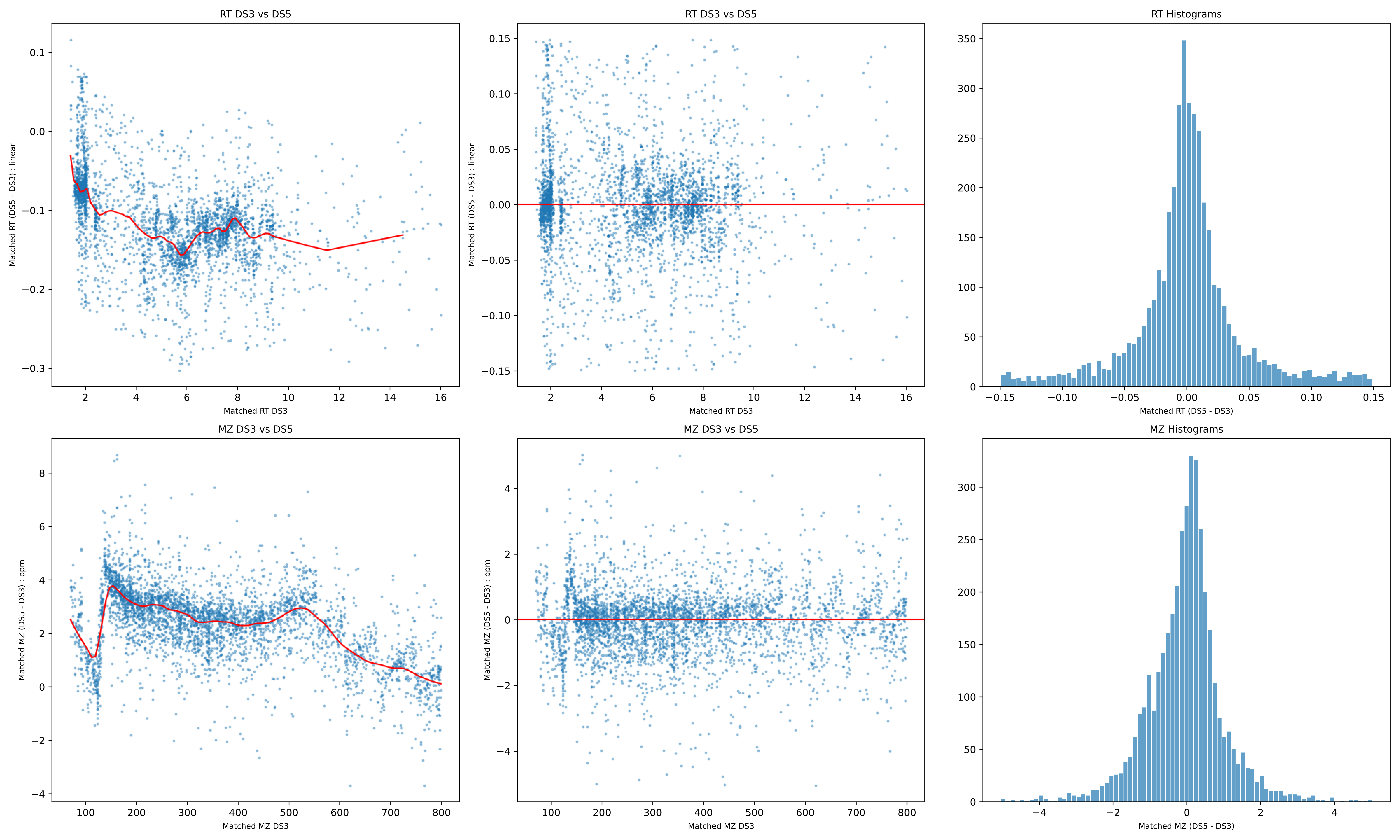
RT DS3 vs DS5
Matched RT (DS5 - DS3) : linear
Matched RT DS3
RT DS3 vs DS5
Matched RT (DS5 - DS3) : linear
Matched RT DS3
RT Histograms
Matched RT (DS5 - DS3)
MZ DS3 vs DS5
Matched MZ (DS5 - DS3) : ppm
Matched MZ DS3
MZ DS3 vs DS5
Matched MZ (DS5 - DS3) : ppm
Matched MZ DS3
MZ Histograms
Matched MZ (DS5 - DS3)

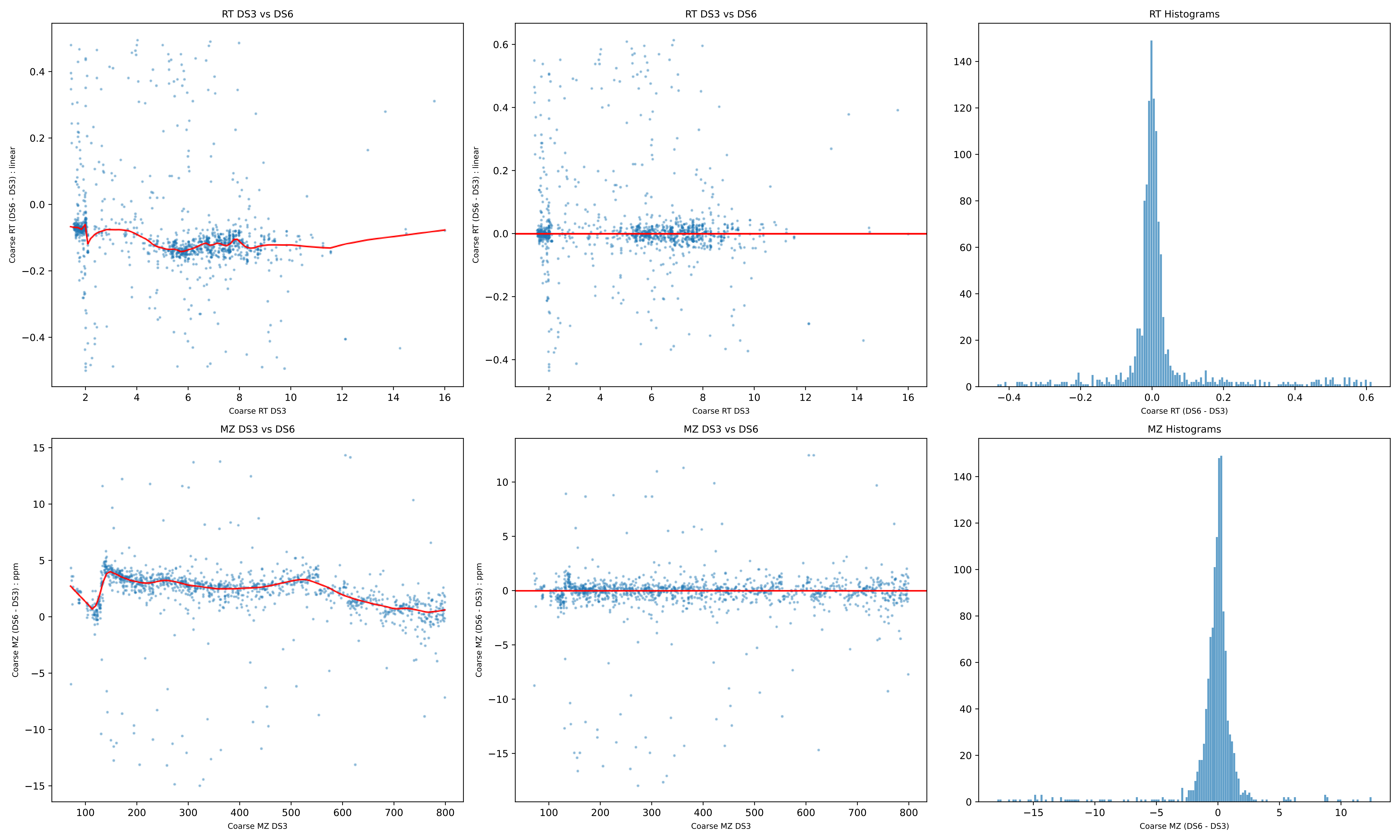

RT DS3 vs DS6
Coarse RT (DS6 - DS3) : linear
Coarse RT DS3
RT DS3 vs DS6
Coarse RT (DS6 - DS3) : linear
Coarse RT DS3
RT Histograms
Coarse RT (DS6 - DS3)
MZ DS3 vs DS6
Coarse MZ (DS6 - DS3) : ppm
Coarse MZ DS3
MZ DS3 vs DS6
Coarse MZ (DS6 - DS3) : ppm
Coarse MZ DS3
MZ Histograms
Coarse MZ (DS6 - DS3)

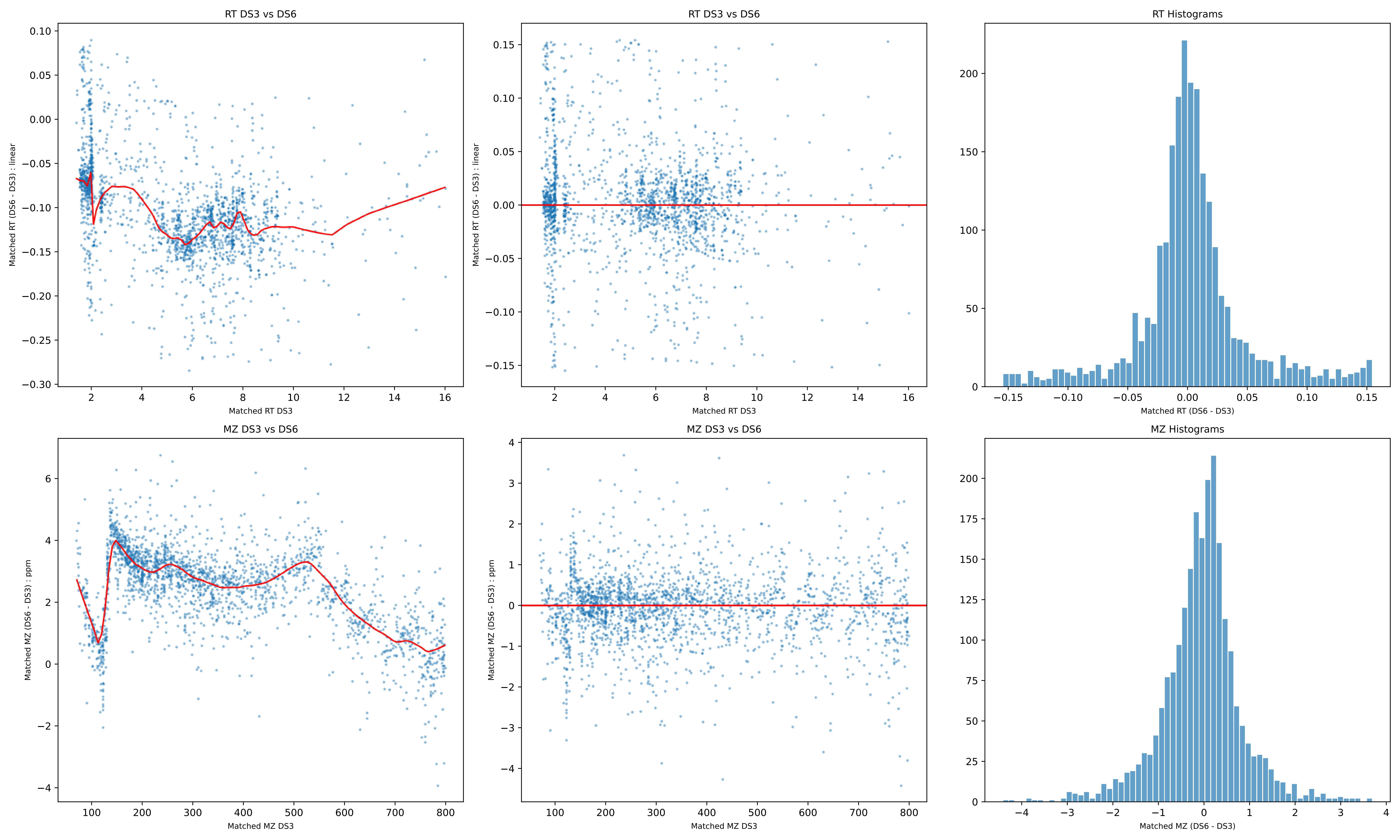
RT DS3 vs DS6
Matched RT (DS6 - DS3) : linear
Matched RT DS3
RT DS3 vs DS6
Matched RT (DS6 - DS3) : linear
Matched RT DS3
RT Histograms
Matched RT (DS6 - DS3)
MZ DS3 vs DS6
Matched MZ (DS6 - DS3) : ppm
Matched MZ DS3
MZ DS3 vs DS6
Matched MZ (DS6 - DS3) : ppm
Matched MZ DS3
MZ Histograms
Matched MZ (DS6 - DS3)

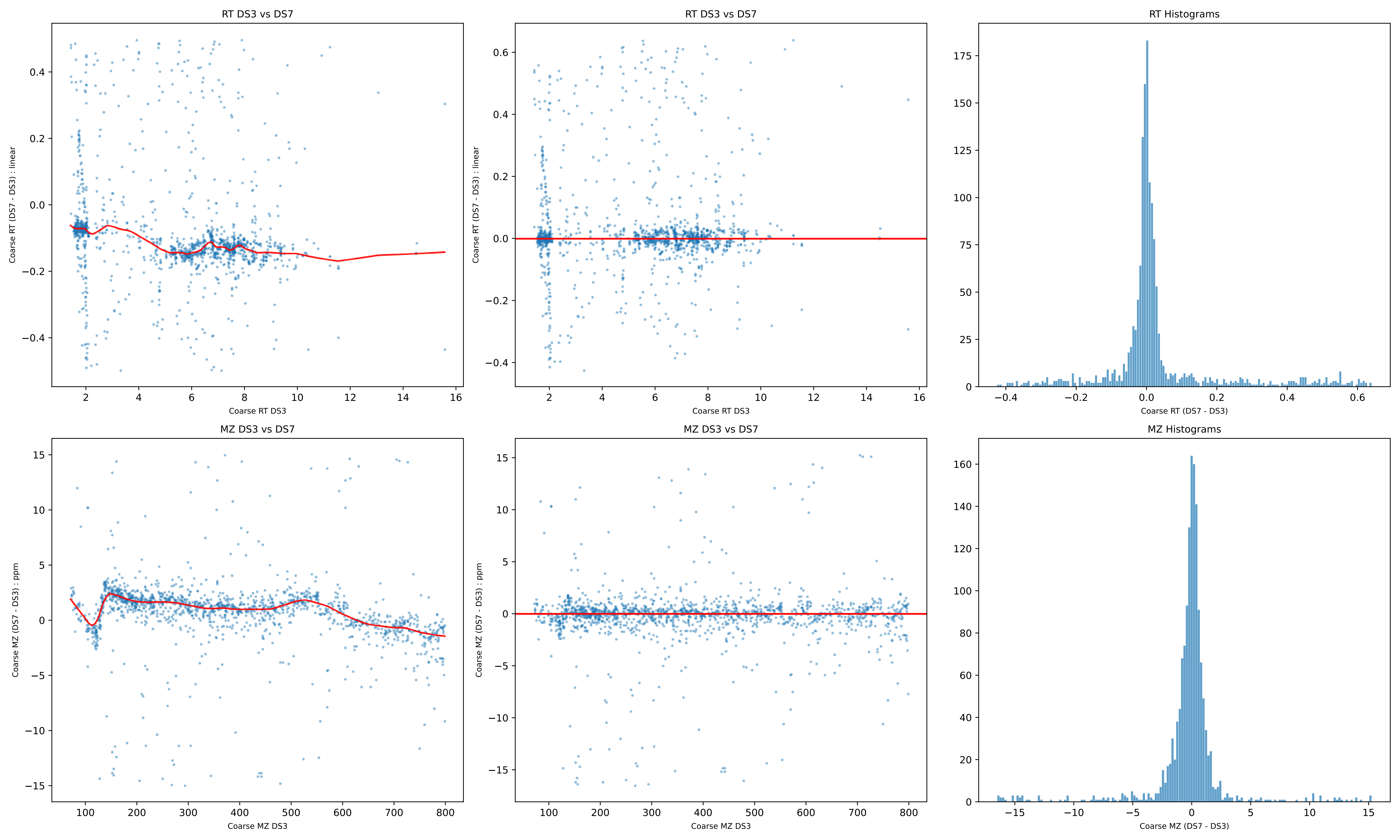
RT DS3 vs DS7
Coarse RT (DS7 - DS3) : linear
Coarse RT DS3
RT DS3 vs DS7
Coarse RT (DS7 - DS3) : linear
Coarse RT DS3
RT Histograms
Coarse RT (DS7 - DS3)
MZ DS3 vs DS7
Coarse MZ (DS7 - DS3) : ppm
Coarse MZ DS3
MZ DS3 vs DS7
Coarse MZ (DS7 - DS3) : ppm
Coarse MZ DS3
MZ Histograms
Coarse MZ (DS7 - DS3)

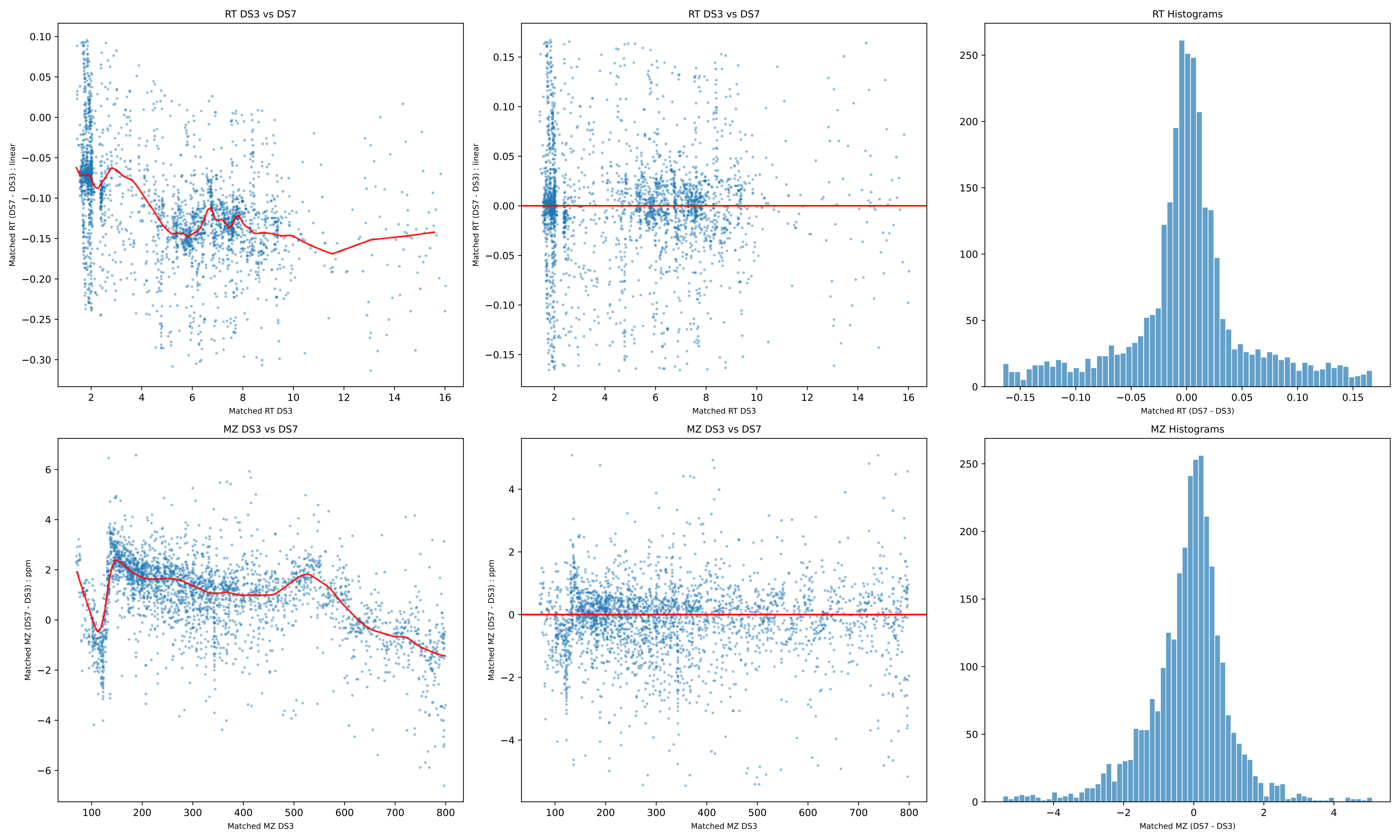

RT DS3 vs DS7
Matched RT (DS7 - DS3) : linear
Matched RT DS3
RT DS3 vs DS7
Matched RT (DS7 - DS3) : linear
Matched RT DS3
RT Histograms
Matched RT (DS7 - DS3)
MZ DS3 vs DS7
Matched MZ (DS7 - DS3) : ppm
Matched MZ DS3
MZ DS3 vs DS7
Matched MZ (DS7 - DS3) : ppm
Matched MZ DS3
MZ Histograms
Matched MZ (DS7 - DS3)

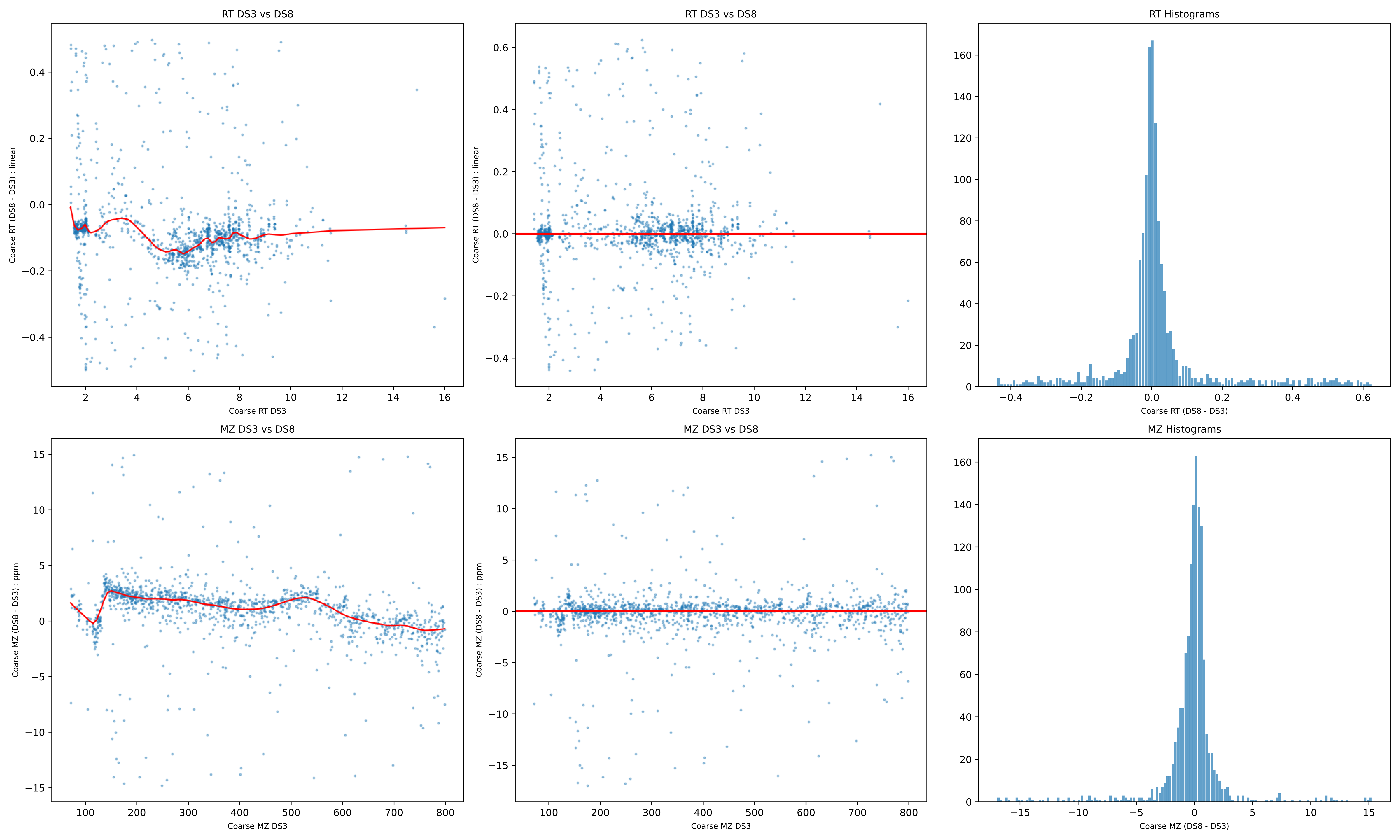

RT DS3 vs DS8
Coarse RT (DS8 - DS3) : linear
Coarse RT DS3
RT DS3 vs DS8
Coarse RT (DS8 - DS3) : linear
Coarse RT DS3
RT Histograms
Coarse RT (DS8 - DS3)
MZ DS3 vs DS8
Coarse MZ (DS8 - DS3) : ppm
Coarse MZ DS3
MZ DS3 vs DS8
Coarse MZ (DS8 - DS3) : ppm
Coarse MZ DS3
MZ Histograms
Coarse MZ (DS8 - DS3)

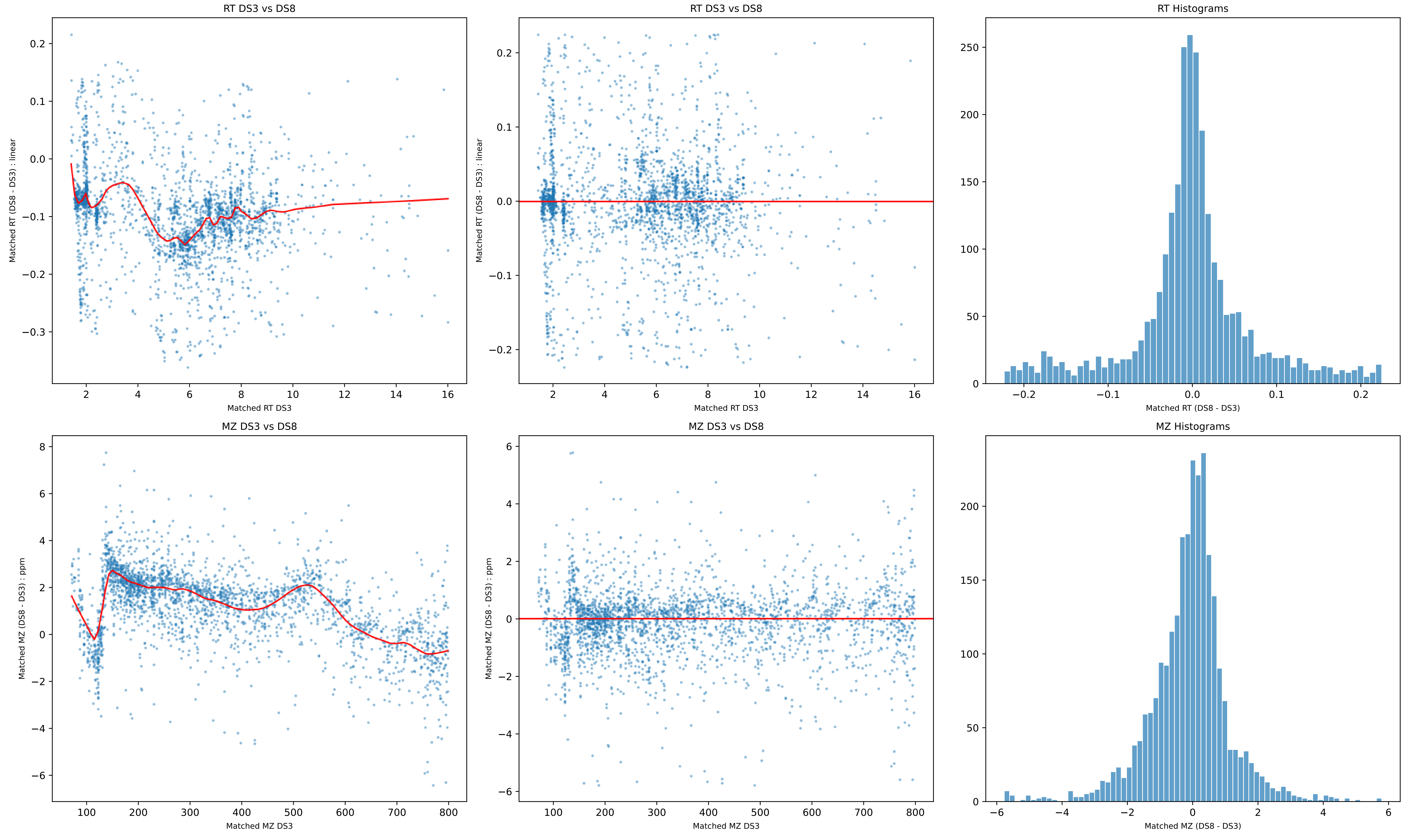
RT DS3 vs DS8
Matched RT (DS8 - DS3) : linear
Matched RT DS3
RT DS3 vs DS8
Matched RT (DS8 - DS3) : linear
Matched RT DS3
RT Histograms
Matched RT (DS8 - DS3)
MZ DS3 vs DS8
Matched MZ (DS8 - DS3) : ppm
Matched MZ DS3
MZ DS3 vs DS8
Matched MZ (DS8 - DS3) : ppm
Matched MZ DS3
MZ Histograms
Matched MZ (DS8 - DS3)

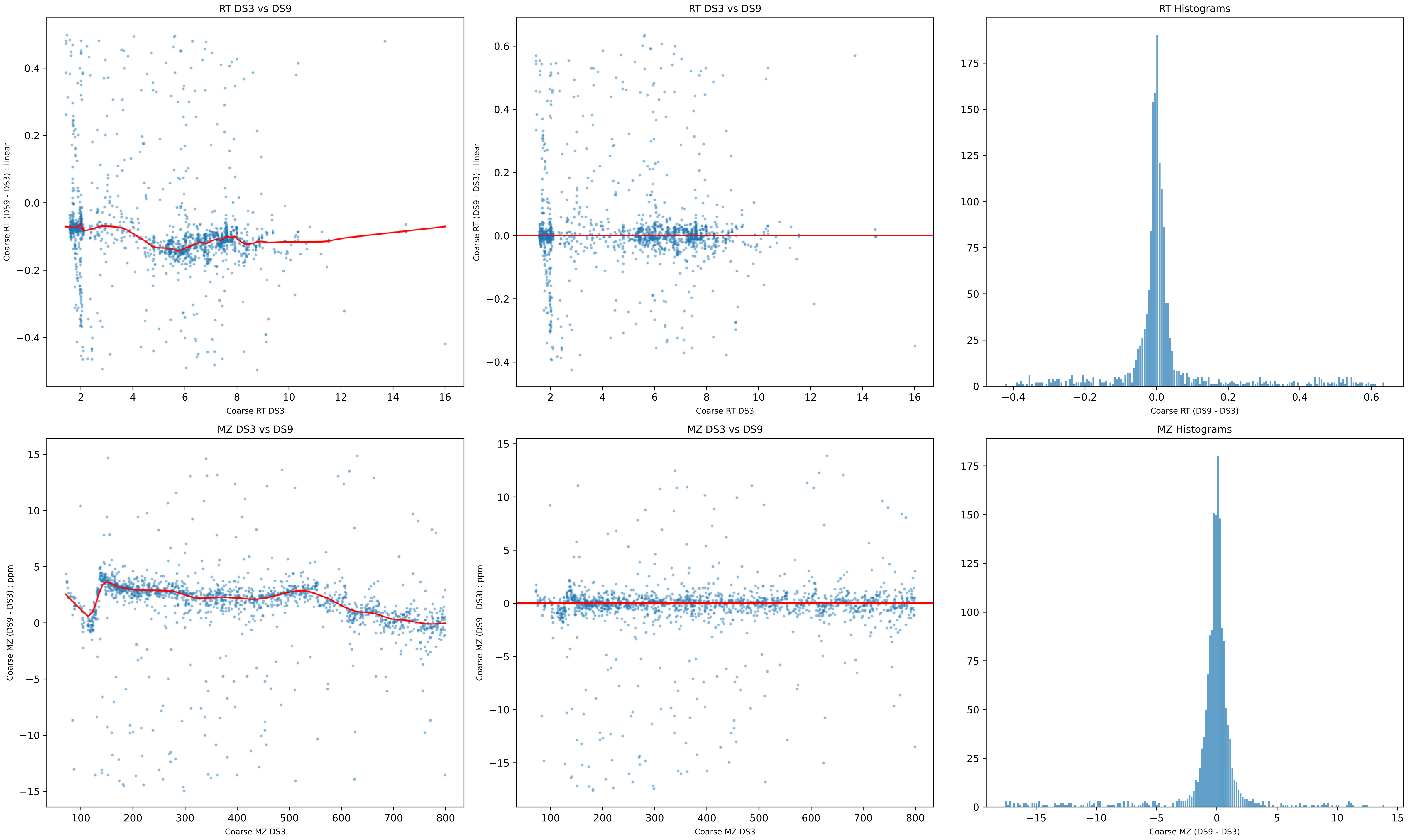
RT DS3 vs DS9
Coarse RT (DS9 - DS3) : linear
Coarse RT DS3
RT DS3 vs DS9
Coarse RT (DS9 - DS3) : linear
Coarse RT DS3
RT Histograms
Coarse RT (DS9 - DS3)
MZ DS3 vs DS9
Coarse MZ (DS9 - DS3) : ppm
Coarse MZ DS3
MZ DS3 vs DS9
Coarse MZ (DS9 - DS3) : ppm
Coarse MZ DS3
MZ Histograms
Coarse MZ (DS9 - DS3)

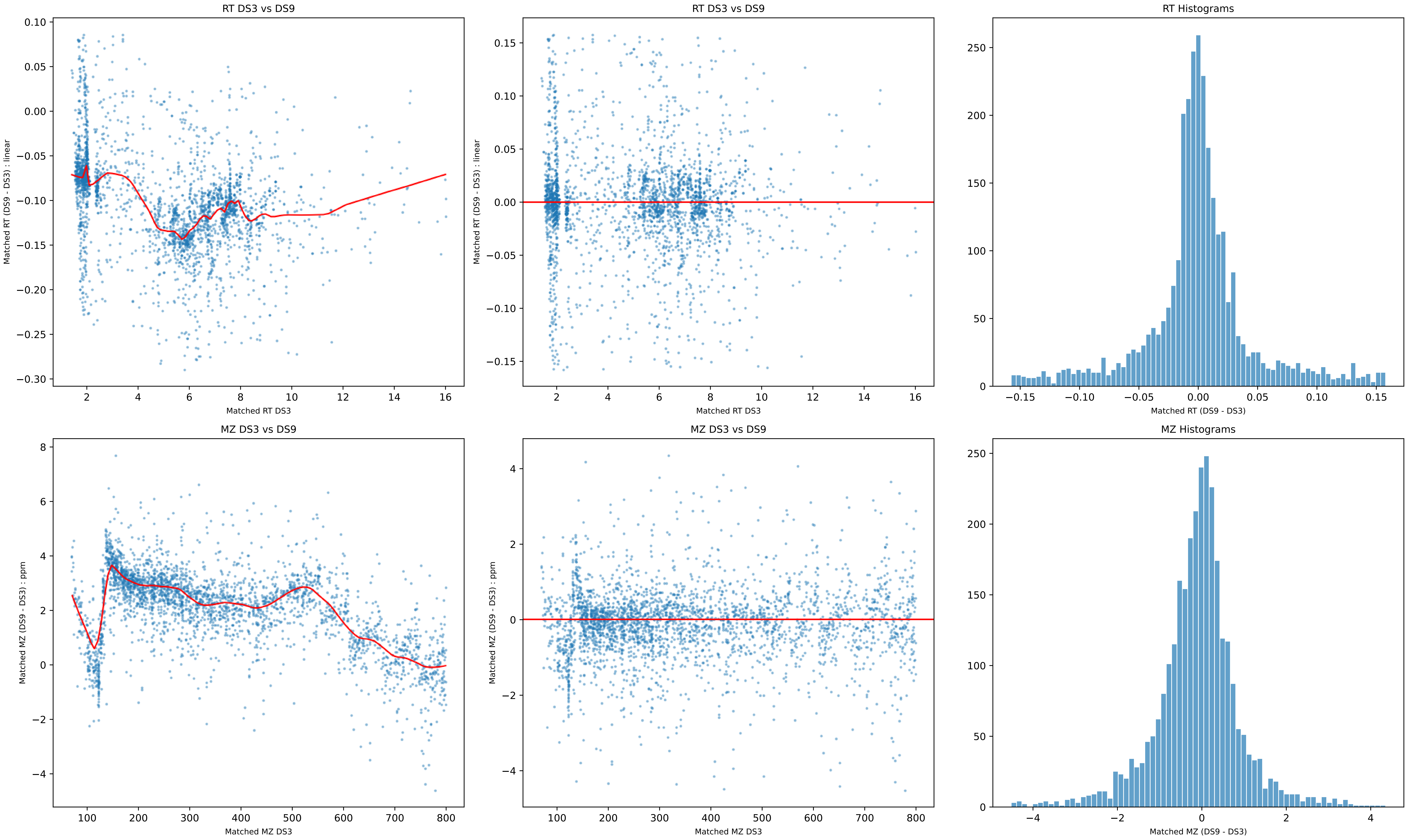
RT DS3 vs DS9
Matched RT (DS9 - DS3) : linear
Matched RT DS3
RT DS3 vs DS9
Matched RT (DS9 - DS3) : linear
Matched RT DS3
RT Histograms
Matched RT (DS9 - DS3)
MZ DS3 vs DS9
Matched MZ (DS9 - DS3) : ppm
Matched MZ DS3
MZ DS3 vs DS9
Matched MZ (DS9 - DS3) : ppm
Matched MZ DS3
MZ Histograms
Matched MZ (DS9 - DS3)

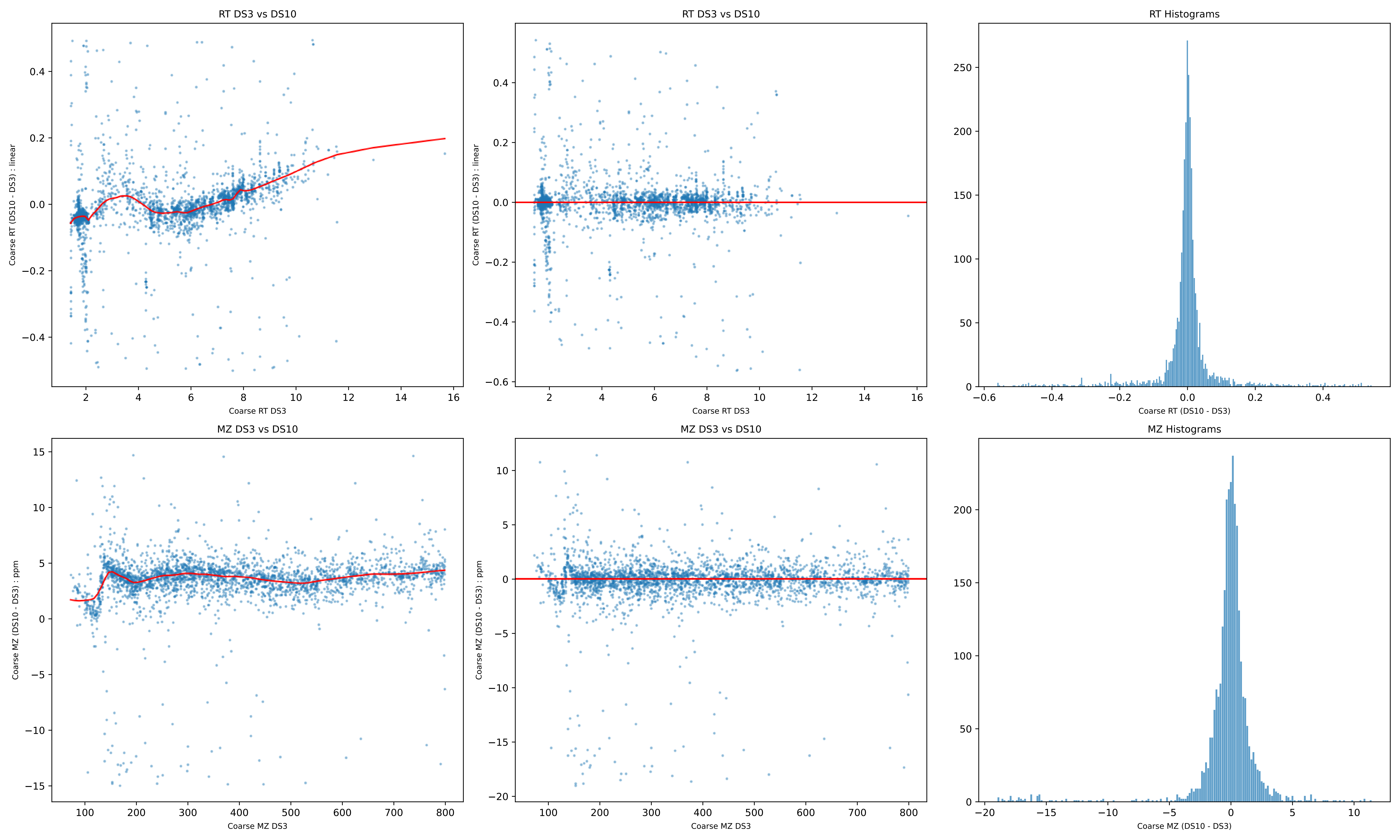
RT DS3 vs DS10
Coarse RT (DS10 - DS3) : linear
Coarse RT DS3
RT DS3 vs DS10
Coarse RT (DS10 - DS3) : linear
Coarse RT DS3
RT Histograms
Coarse RT (DS10 - DS3)
MZ DS3 vs DS10
Coarse MZ (DS10 - DS3) : ppm
Coarse MZ DS3
MZ DS3 vs DS10
Coarse MZ (DS10 - DS3) : ppm
Coarse MZ DS3
MZ Histograms
Coarse MZ (DS10 - DS3)

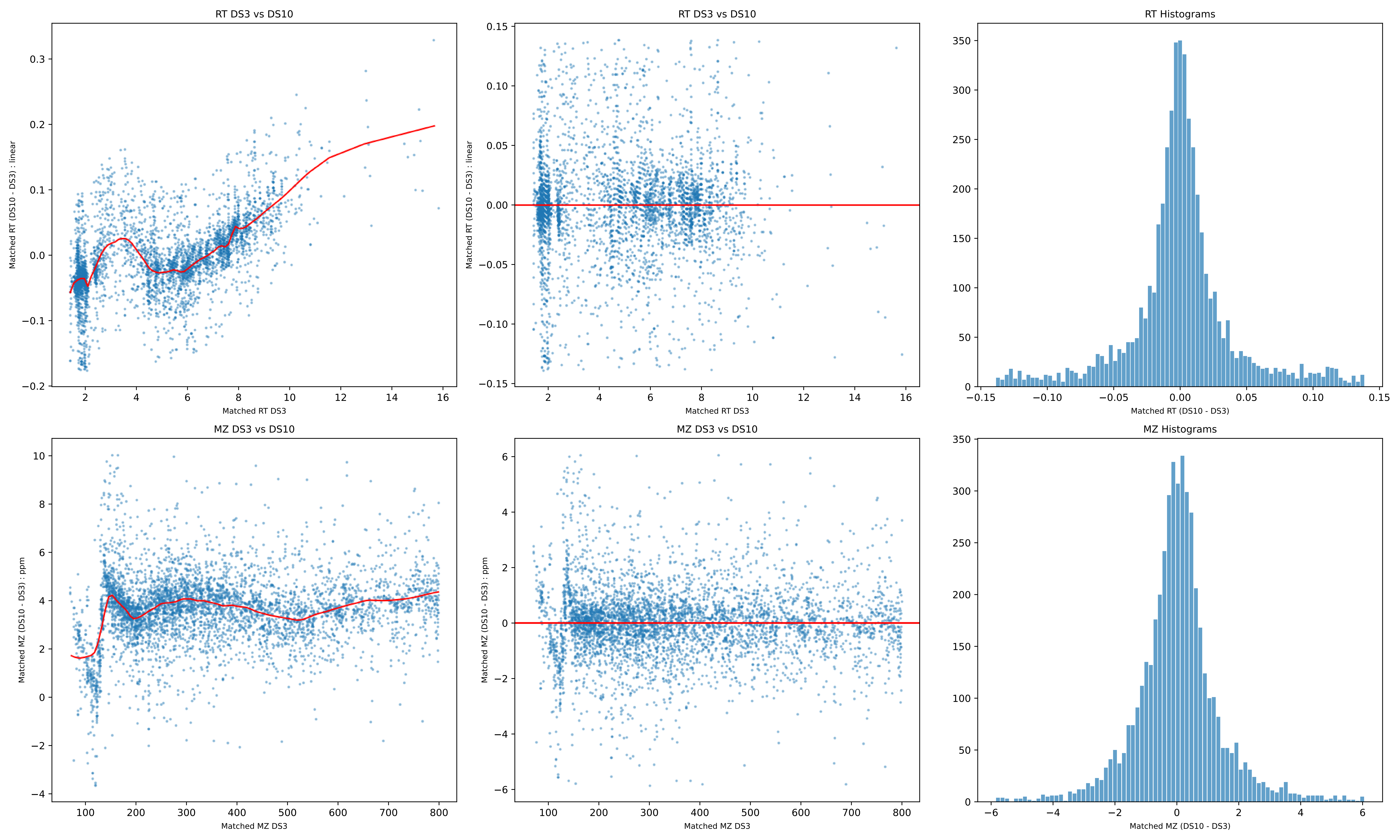

RT DS3 vs DS10
Matched RT (DS10 - DS3) : linear
Matched RT DS3
RT DS3 vs DS10
Matched RT (DS10 - DS3) : linear
Matched RT DS3
RT Histograms
Matched RT (DS10 - DS3)
MZ DS3 vs DS10
Matched MZ (DS10 - DS3) : ppm
Matched MZ DS3
MZ DS3 vs DS10
Matched MZ (DS10 - DS3) : ppm
Matched MZ DS3
MZ Histograms
Matched MZ (DS10 - DS3)

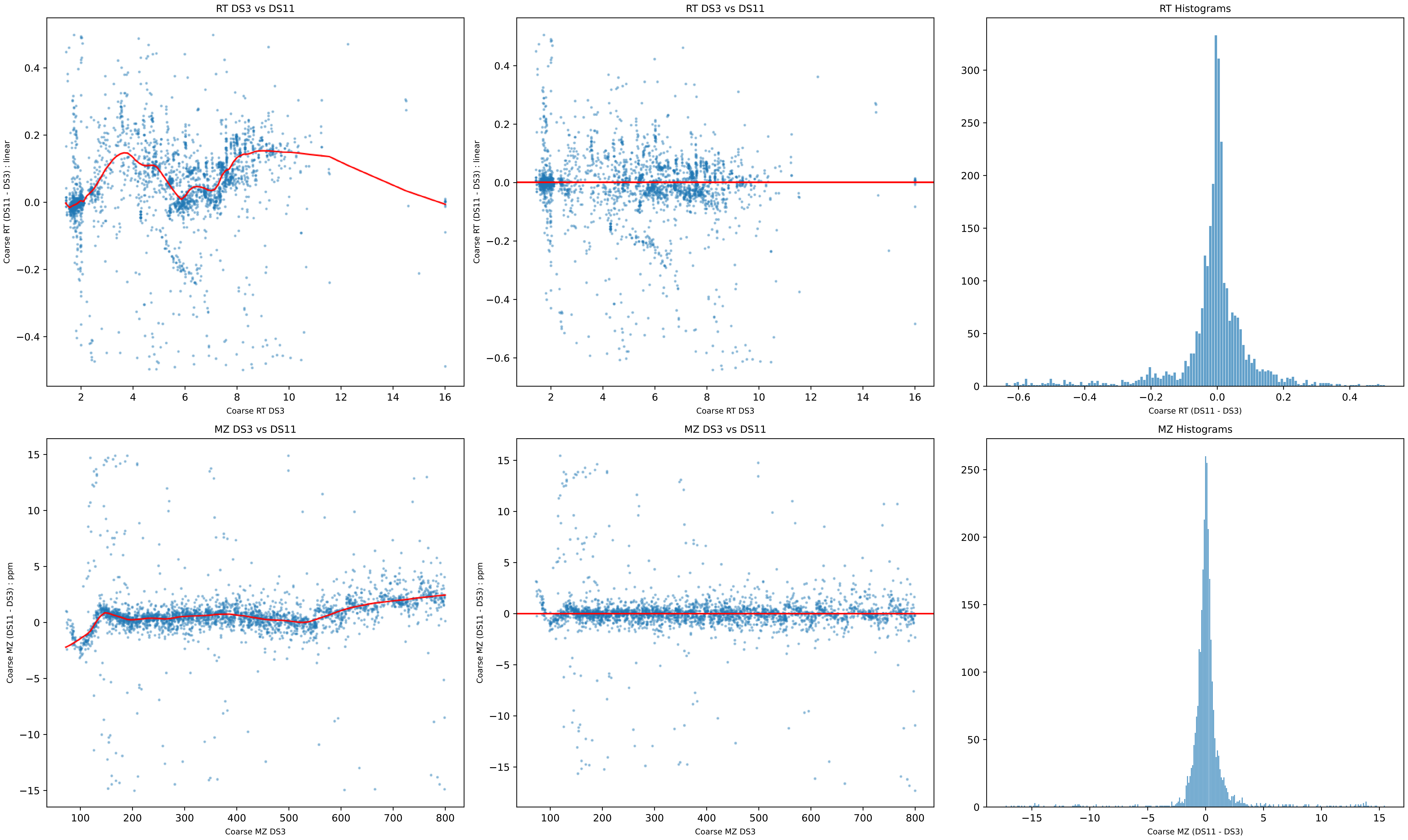
RT DS3 vs DS11
Coarse RT (DS11 - DS3) : linear
Coarse RT DS3
RT DS3 vs DS11
Coarse RT (DS11 - DS3) : linear
Coarse RT DS3
RT Histograms
Coarse RT (DS11 - DS3)
MZ DS3 vs DS11
Coarse MZ (DS11 - DS3) : ppm
Coarse MZ DS3
MZ DS3 vs DS11
Coarse MZ (DS11 - DS3) : ppm
Coarse MZ DS3
MZ Histograms
Coarse MZ (DS11 - DS3)

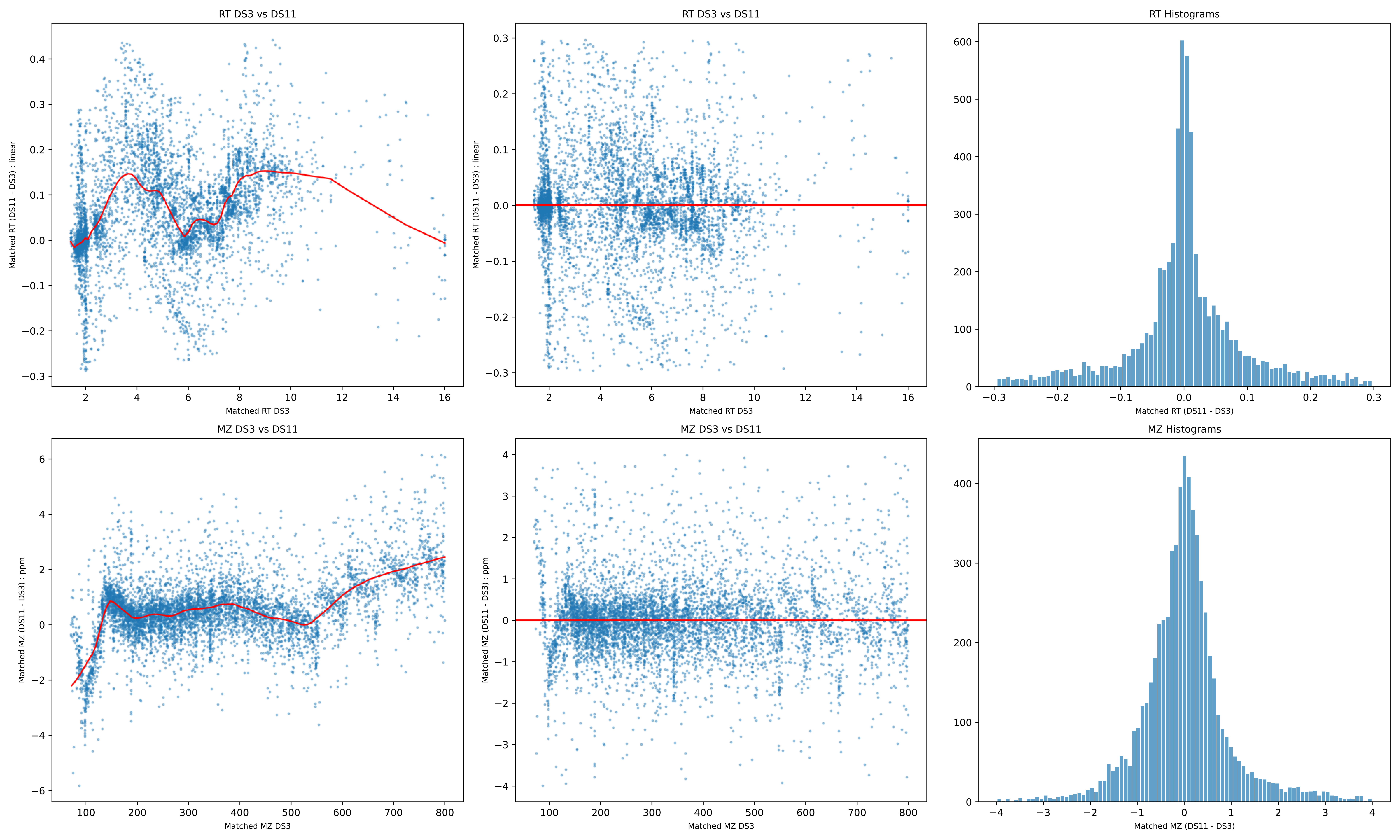

RT DS3 vs DS11
Matched RT (DS11 - DS3) : linear
Matched RT DS3
RT DS3 vs DS11
Matched RT (DS11 - DS3) : linear
Matched RT DS3
RT Histograms
Matched RT (DS11 - DS3)
MZ DS3 vs DS11
Matched MZ (DS11 - DS3) : ppm
Matched MZ DS3
MZ DS3 vs DS11
Matched MZ (DS11 - DS3) : ppm
Matched MZ DS3
MZ Histograms
Matched MZ (DS11 - DS3)

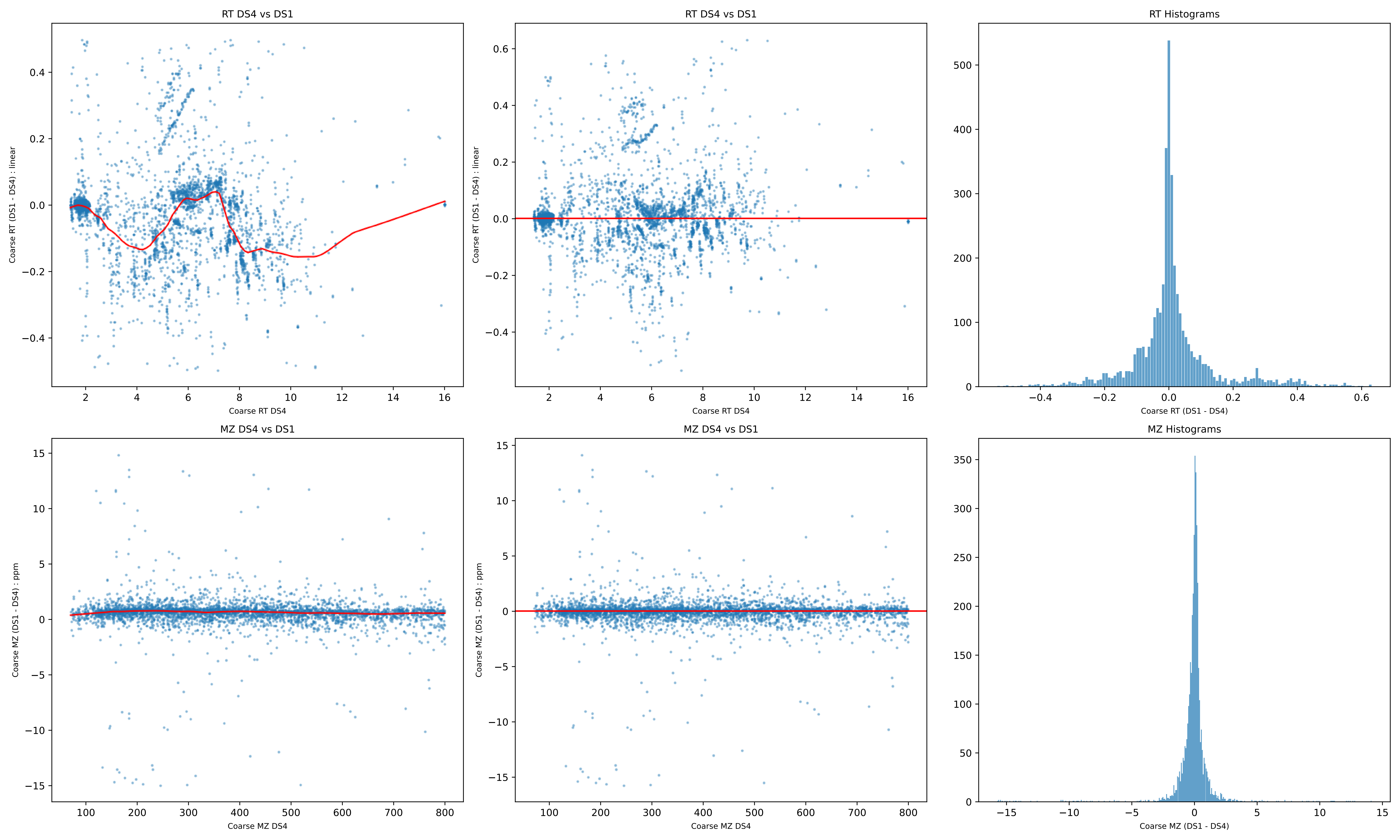

RT DS4 vs DS1
Coarse RT (DS1 - DS4) : linear
Coarse RT DS4
RT DS4 vs DS1
Coarse RT (DS1 - DS4) : linear
Coarse RT DS4
RT Histograms
Coarse RT (DS1 - DS4)
MZ DS4 vs DS1
Coarse MZ (DS1 - DS4) : ppm
Coarse MZ DS4
MZ DS4 vs DS1
Coarse MZ (DS1 - DS4) : ppm
Coarse MZ DS4
MZ Histograms
Coarse MZ (DS1 - DS4)

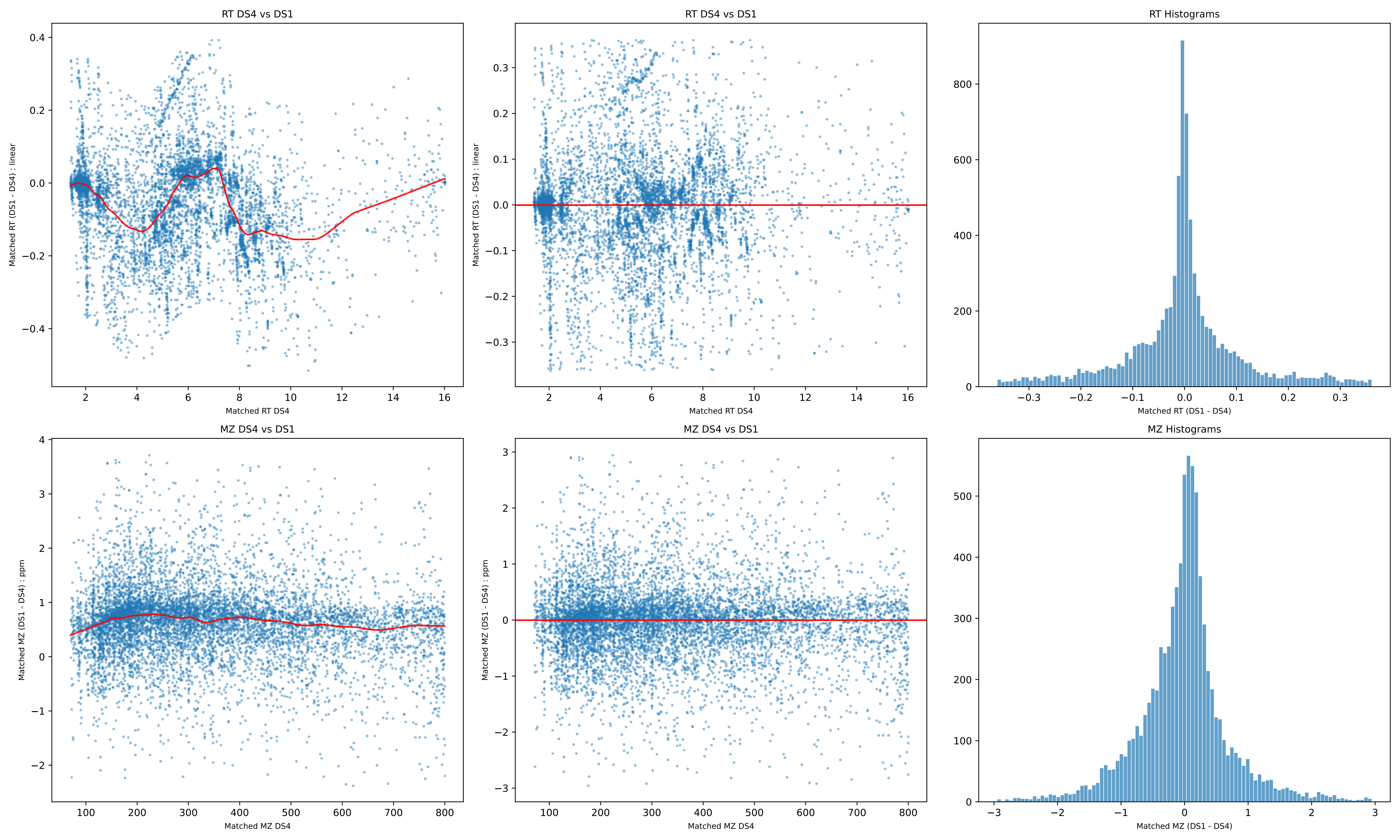

RT DS4 vs DS1
Matched RT (DS1 - DS4) : linear
Matched RT DS4
RT DS4 vs DS1
Matched RT (DS1 - DS4) : linear
Matched RT DS4
RT Histograms
Matched RT (DS1 - DS4)
MZ DS4 vs DS1
Matched MZ (DS1 - DS4) : ppm
Matched MZ DS4
MZ DS4 vs DS1
Matched MZ (DS1 - DS4) : ppm
Matched MZ DS4
MZ Histograms
Matched MZ (DS1 - DS4)

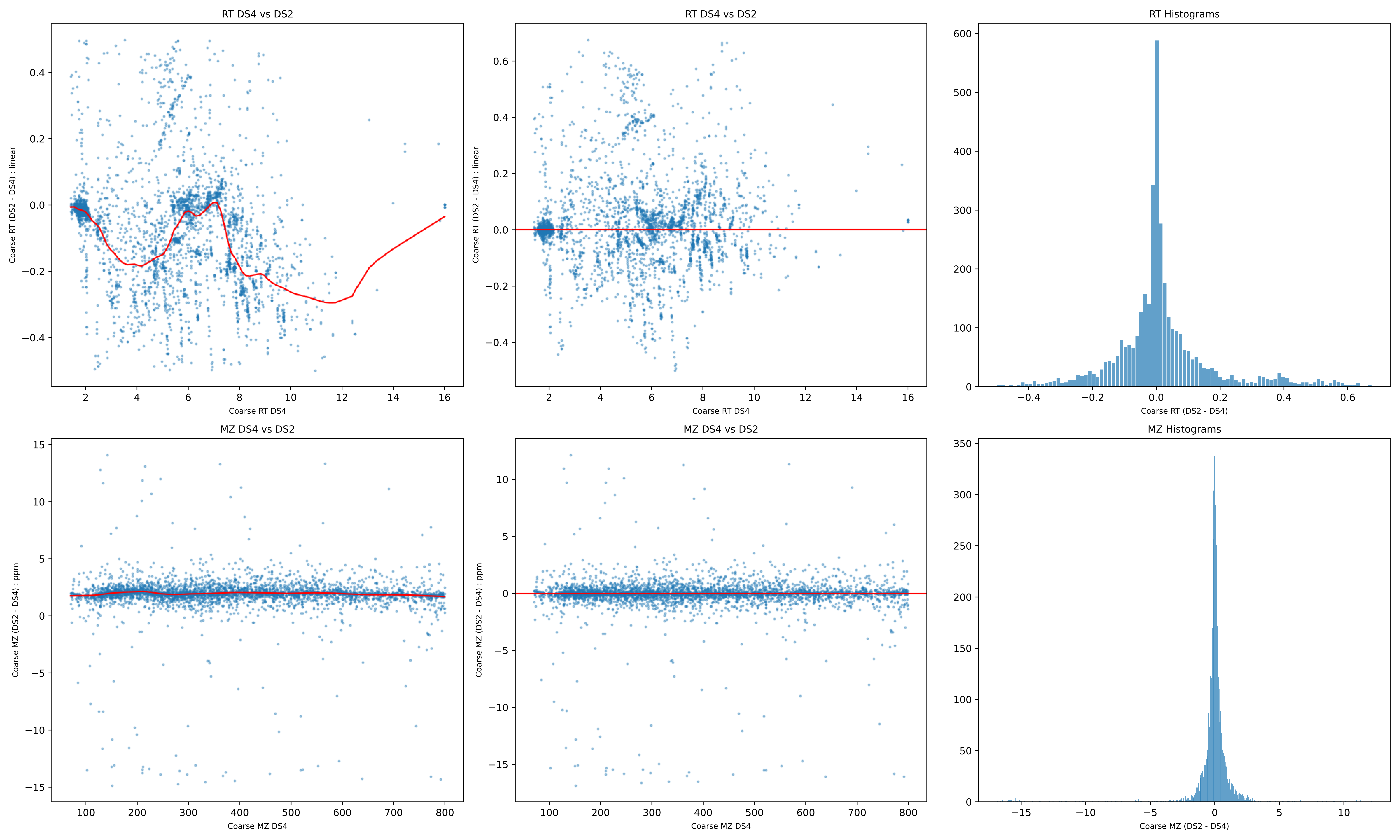
RT DS4 vs DS2
Coarse RT (DS2 - DS4) : linear
Coarse RT DS4
RT DS4 vs DS2
Coarse RT (DS2 - DS4) : linear
Coarse RT DS4
RT Histograms
Coarse RT (DS2 - DS4)
MZ DS4 vs DS2
Coarse MZ (DS2 - DS4) : ppm
Coarse MZ DS4
MZ DS4 vs DS2
Coarse MZ (DS2 - DS4) : ppm
Coarse MZ DS4
MZ Histograms
Coarse MZ (DS2 - DS4)

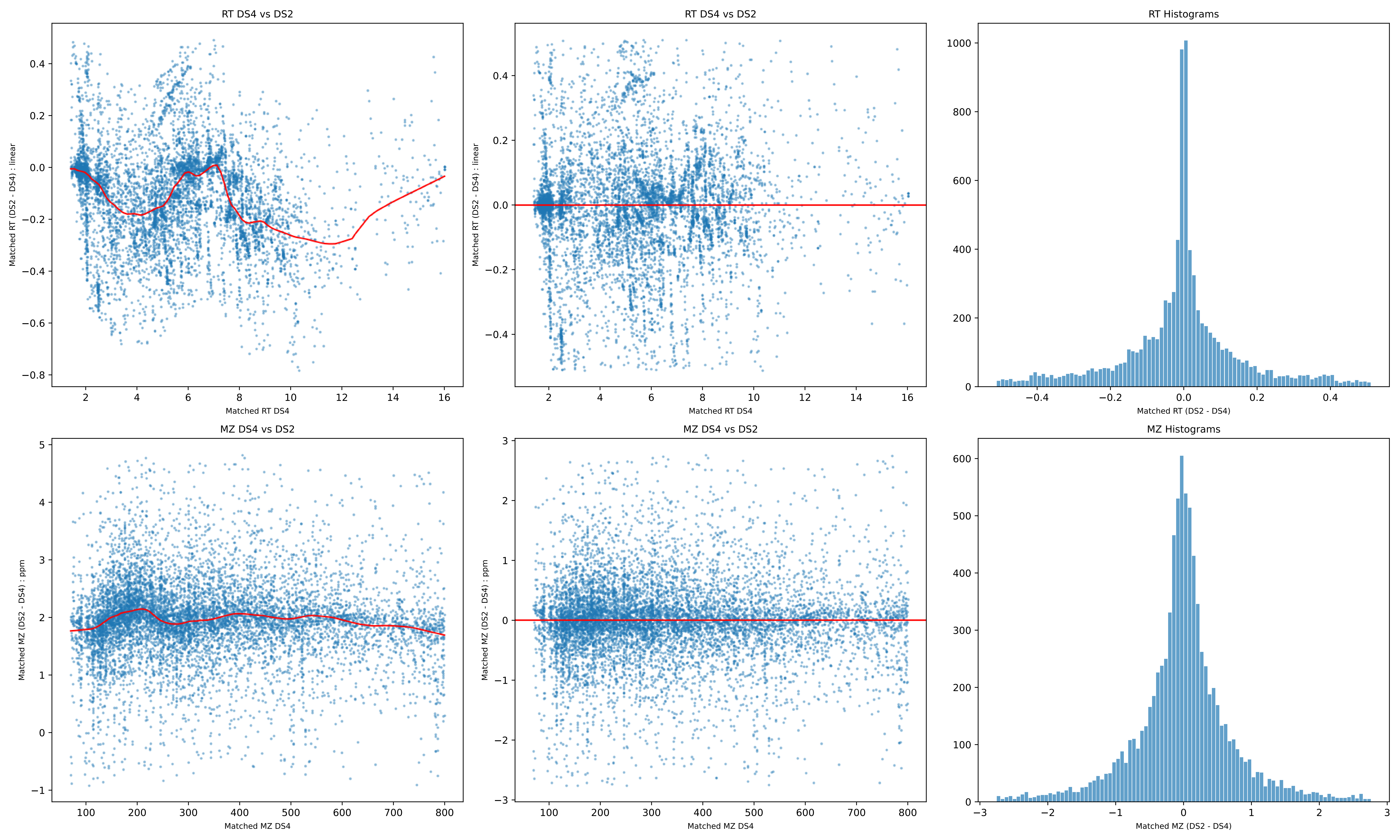
RT DS4 vs DS2
Matched RT (DS2 - DS4) : linear
Matched RT DS4
RT DS4 vs DS2
Matched RT (DS2 - DS4) : linear
Matched RT DS4
RT Histograms
Matched RT (DS2 - DS4)
MZ DS4 vs DS2
Matched MZ (DS2 - DS4) : ppm
Matched MZ DS4
MZ DS4 vs DS2
Matched MZ (DS2 - DS4) : ppm
Matched MZ DS4
MZ Histograms
Matched MZ (DS2 - DS4)

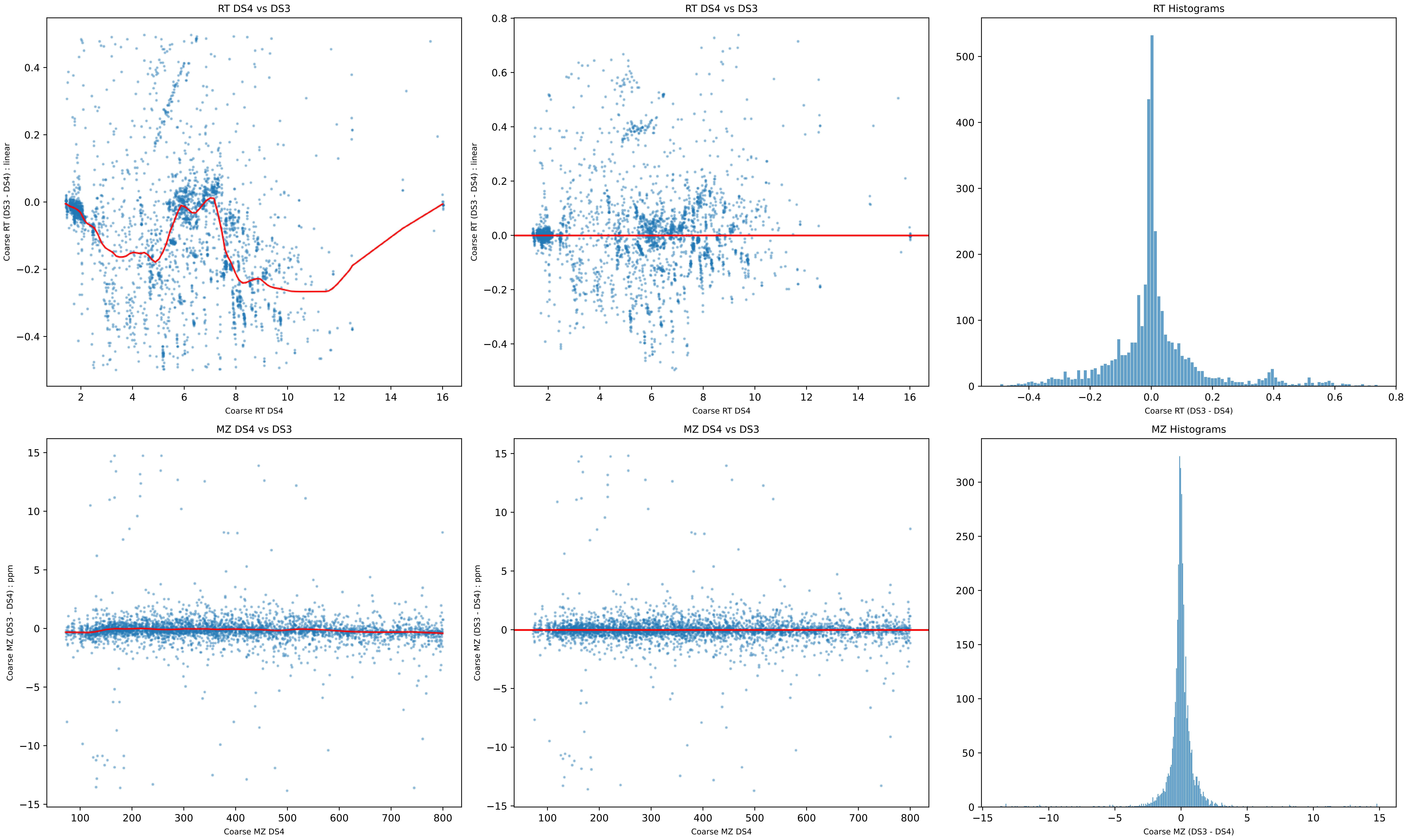
RT DS4 vs DS3
Coarse RT (DS3 - DS4) : linear
Coarse RT DS4
RT DS4 vs DS3
Coarse RT (DS3 - DS4) : linear
Coarse RT DS4
RT Histograms
Coarse RT (DS3 - DS4)
MZ DS4 vs DS3
Coarse MZ (DS3 - DS4) : ppm
Coarse MZ DS4
MZ DS4 vs DS3
Coarse MZ (DS3 - DS4) : ppm
Coarse MZ DS4
MZ Histograms
Coarse MZ (DS3 - DS4)

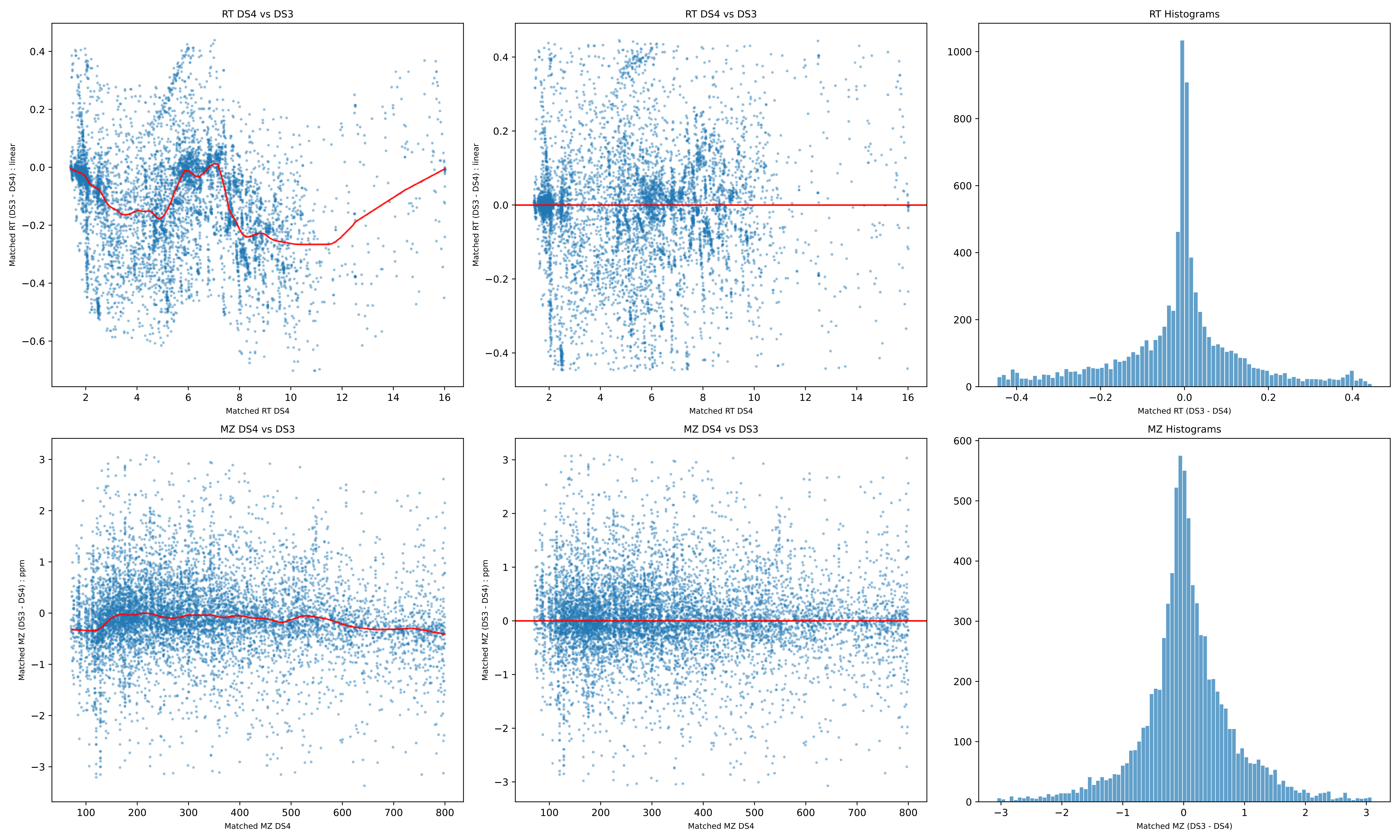
RT DS4 vs DS3
Matched RT (DS3 - DS4) : linear
Matched RT DS4
RT DS4 vs DS3
Matched RT (DS3 - DS4) : linear
Matched RT DS4
RT Histograms
Matched RT (DS3 - DS4)
MZ DS4 vs DS3
Matched MZ (DS3 - DS4) : ppm
Matched MZ DS4
MZ DS4 vs DS3
Matched MZ (DS3 - DS4) : ppm
Matched MZ DS4
MZ Histograms
Matched MZ (DS3 - DS4)

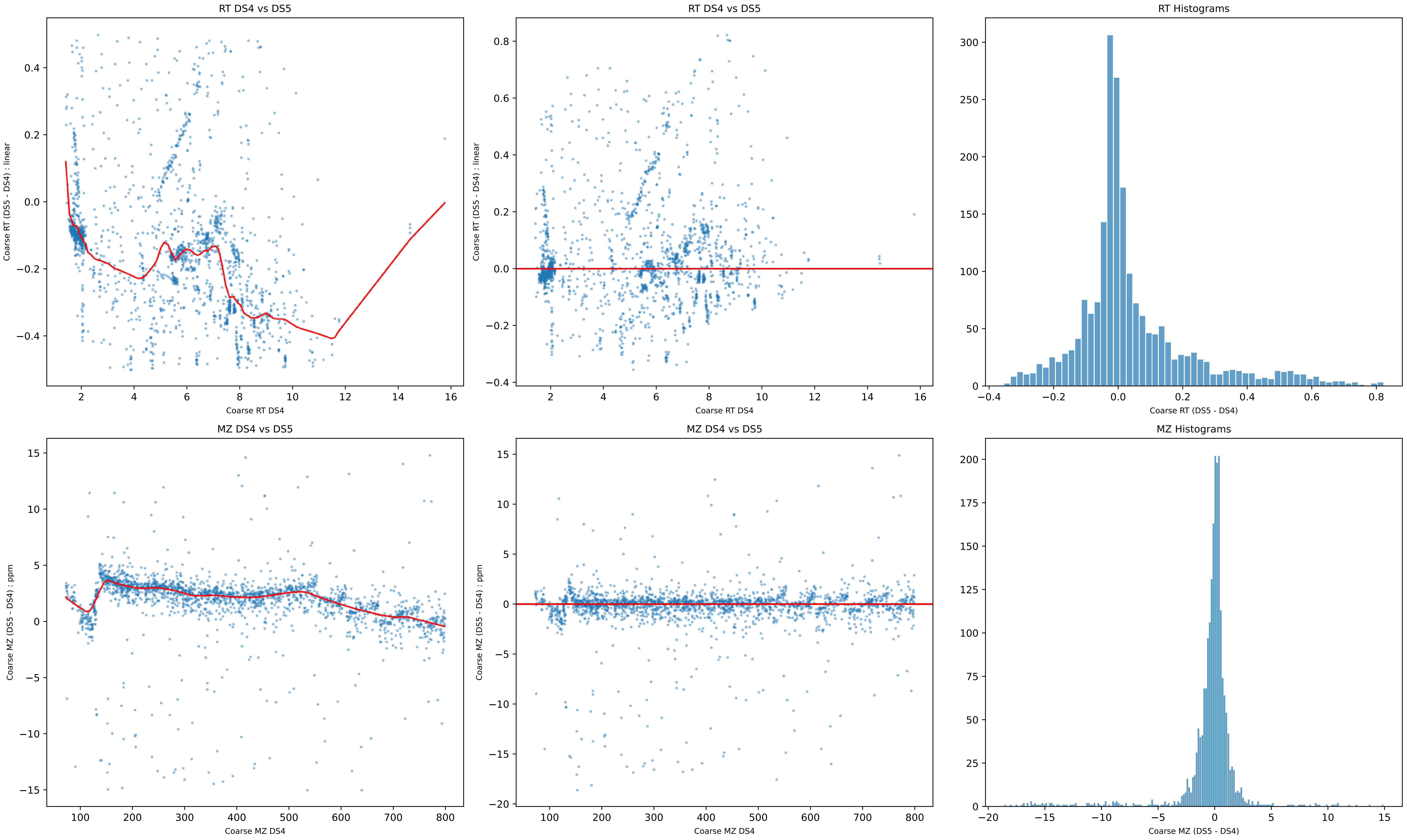
RT DS4 vs DS5
Coarse RT (DS5 - DS4) : linear
Coarse RT DS4
RT DS4 vs DS5
Coarse RT (DS5 - DS4) : linear
Coarse RT DS4
RT Histograms
Coarse RT (DS5 - DS4)
MZ DS4 vs DS5
Coarse MZ (DS5 - DS4) : ppm
Coarse MZ DS4
MZ DS4 vs DS5
Coarse MZ (DS5 - DS4) : ppm
Coarse MZ DS4
MZ Histograms
Coarse MZ (DS5 - DS4)

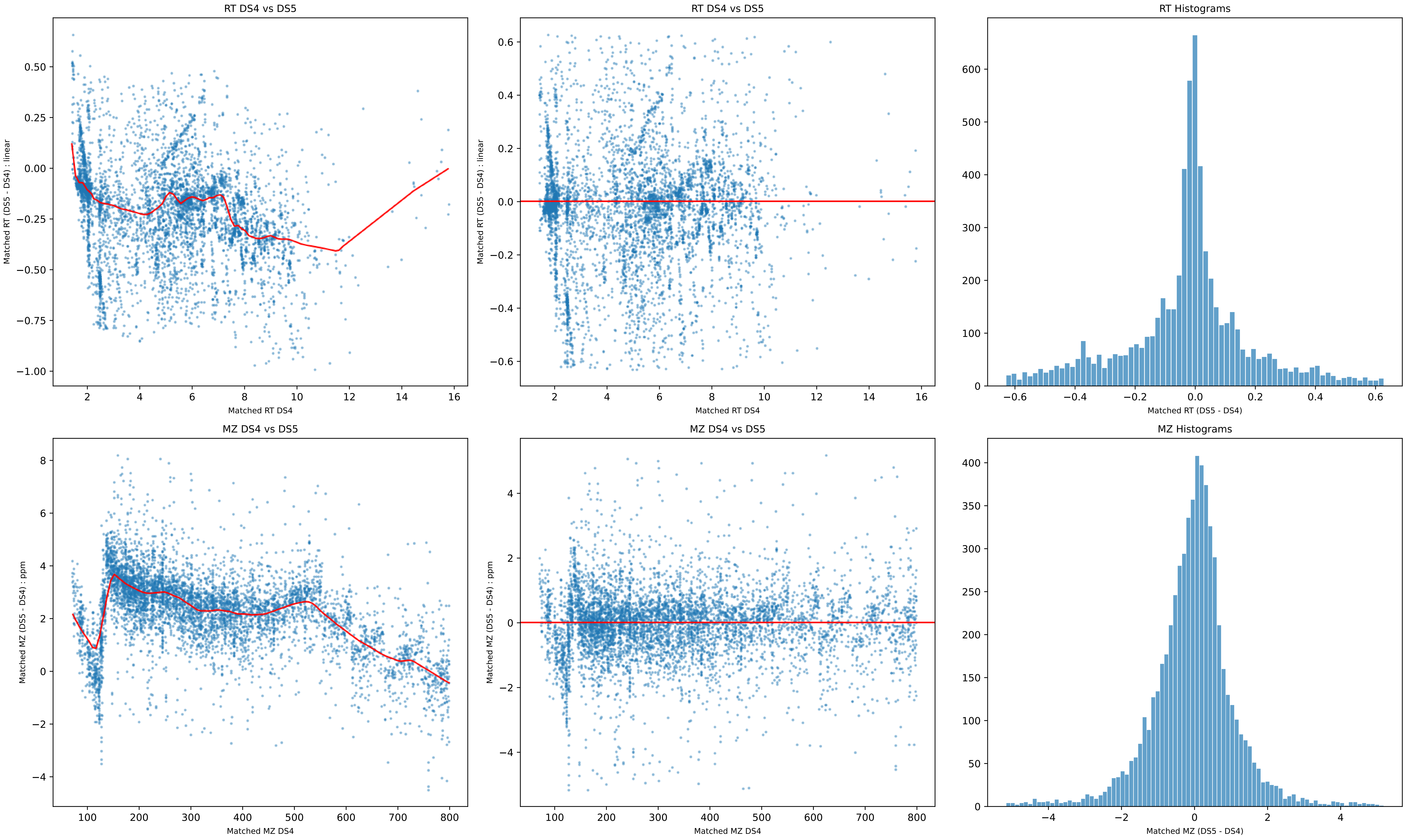
RT DS4 vs DS5
Matched RT (DS5 - DS4) : linear
Matched RT DS4
RT DS4 vs DS5
Matched RT (DS5 - DS4) : linear
Matched RT DS4
RT Histograms
Matched RT (DS5 - DS4)
MZ DS4 vs DS5
Matched MZ (DS5 - DS4) : ppm
Matched MZ DS4
MZ DS4 vs DS5
Matched MZ (DS5 - DS4) : ppm
Matched MZ DS4
MZ Histograms
Matched MZ (DS5 - DS4)

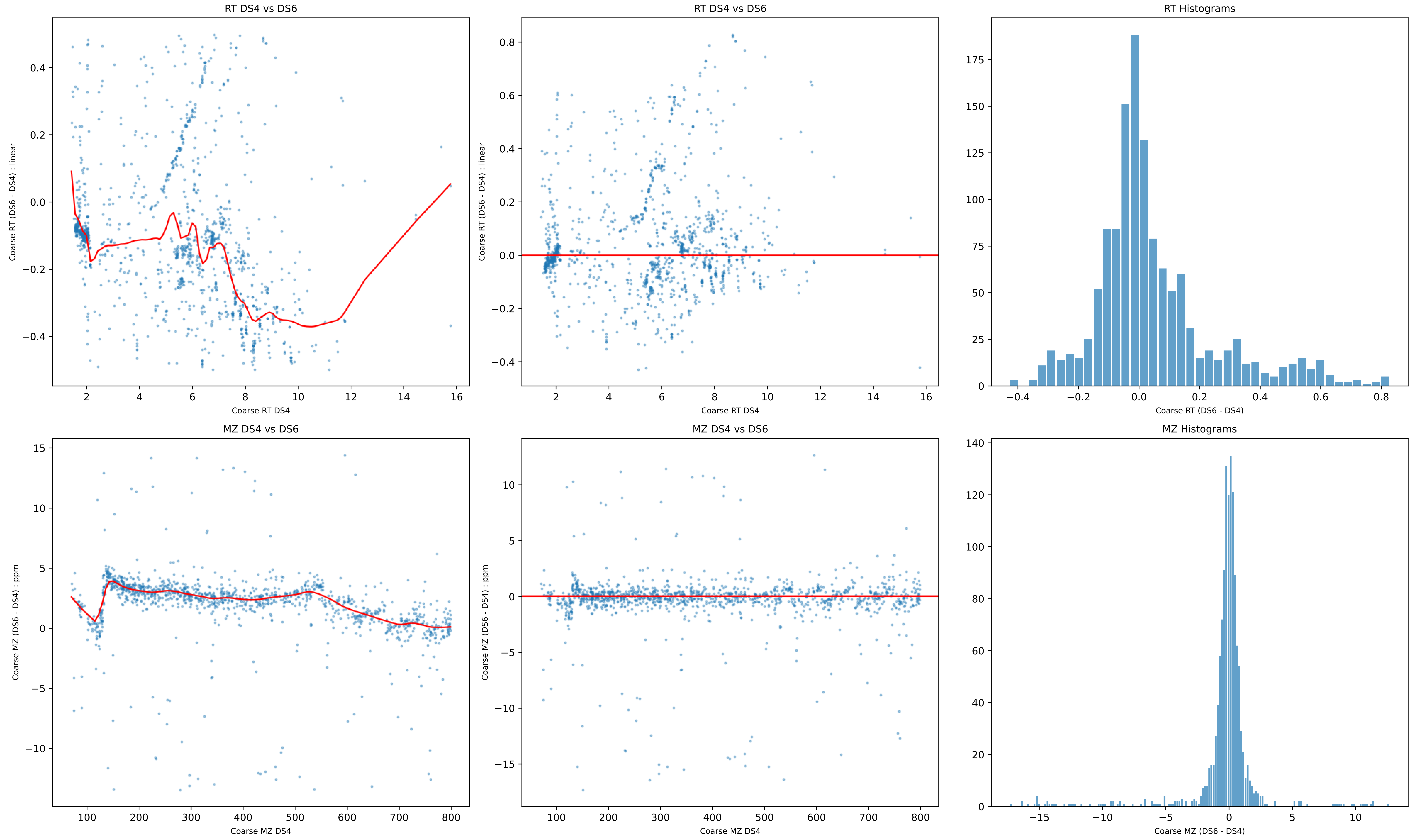
RT DS4 vs DS6
Coarse RT (DS6 - DS4) : linear
Coarse RT DS4
RT DS4 vs DS6
Coarse RT (DS6 - DS4) : linear
Coarse RT DS4
RT Histograms
Coarse RT (DS6 - DS4)
MZ DS4 vs DS6
Coarse MZ (DS6 - DS4) : ppm
Coarse MZ DS4
MZ DS4 vs DS6
Coarse MZ (DS6 - DS4) : ppm
Coarse MZ DS4
MZ Histograms
Coarse MZ (DS6 - DS4)

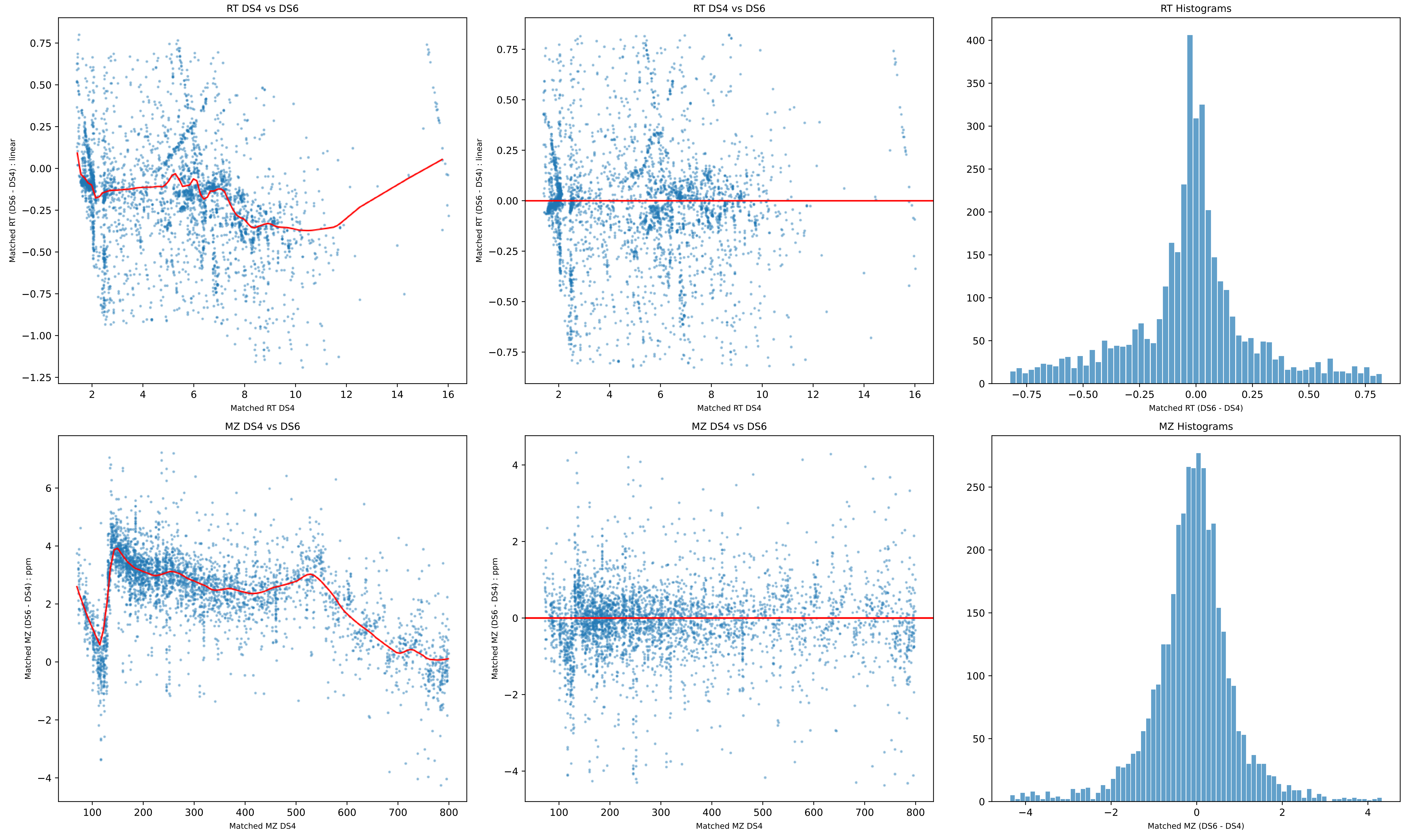
RT DS4 vs DS6
Matched RT (DS6 - DS4) : linear
Matched RT DS4
RT DS4 vs DS6
Matched RT (DS6 - DS4) : linear
Matched RT DS4
RT Histograms
Matched RT (DS6 - DS4)
MZ DS4 vs DS6
Matched MZ (DS6 - DS4) : ppm
Matched MZ DS4
MZ DS4 vs DS6
Matched MZ (DS6 - DS4) : ppm
Matched MZ DS4
MZ Histograms
Matched MZ (DS6 - DS4)

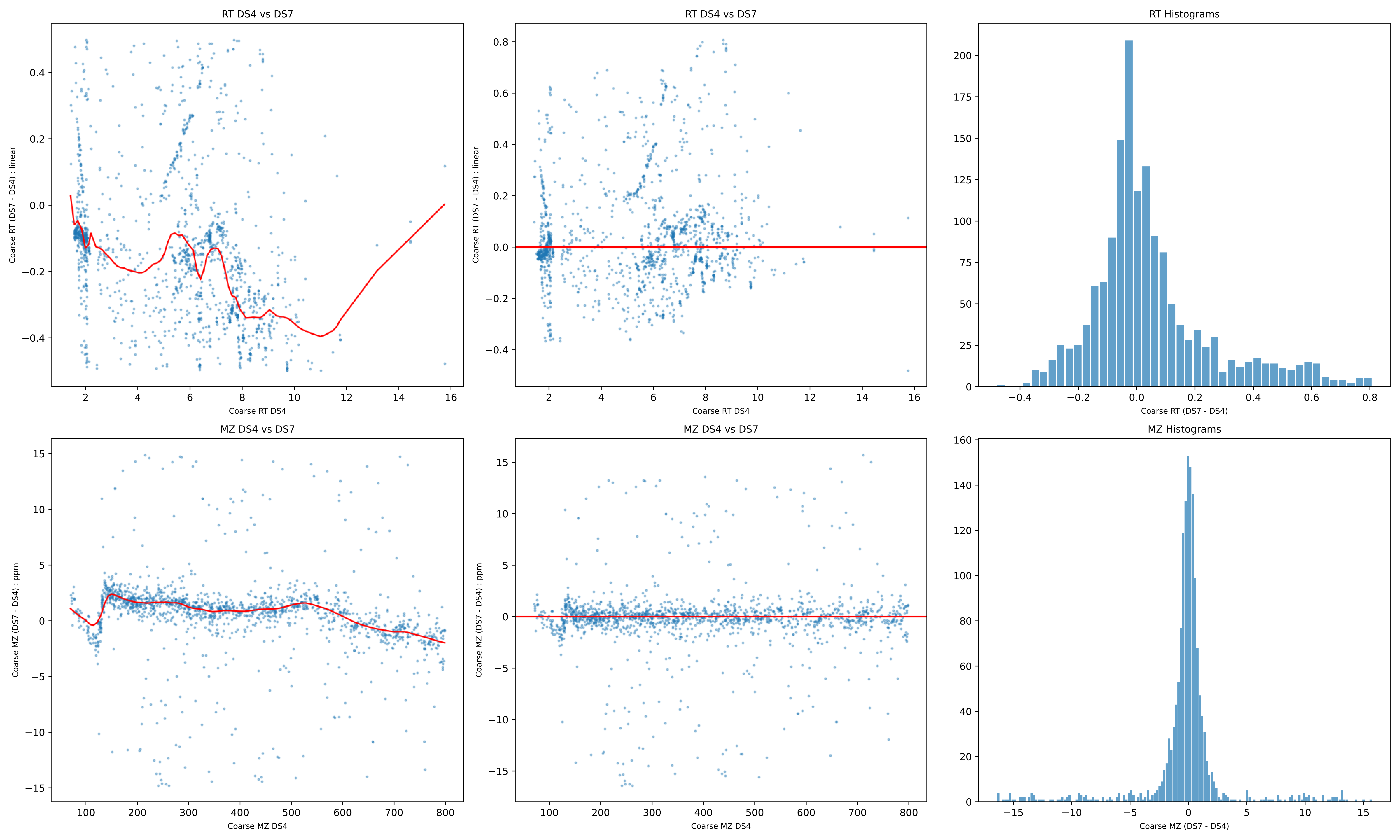

RT DS4 vs DS7
Coarse RT (DS7 - DS4) : linear
Coarse RT DS4
RT DS4 vs DS7
Coarse RT (DS7 - DS4) : linear
Coarse RT DS4
RT Histograms
Coarse RT (DS7 - DS4)
MZ DS4 vs DS7
Coarse MZ (DS7 - DS4) : ppm
Coarse MZ DS4
MZ DS4 vs DS7
Coarse MZ (DS7 - DS4) : ppm
Coarse MZ DS4
MZ Histograms
Coarse MZ (DS7 - DS4)

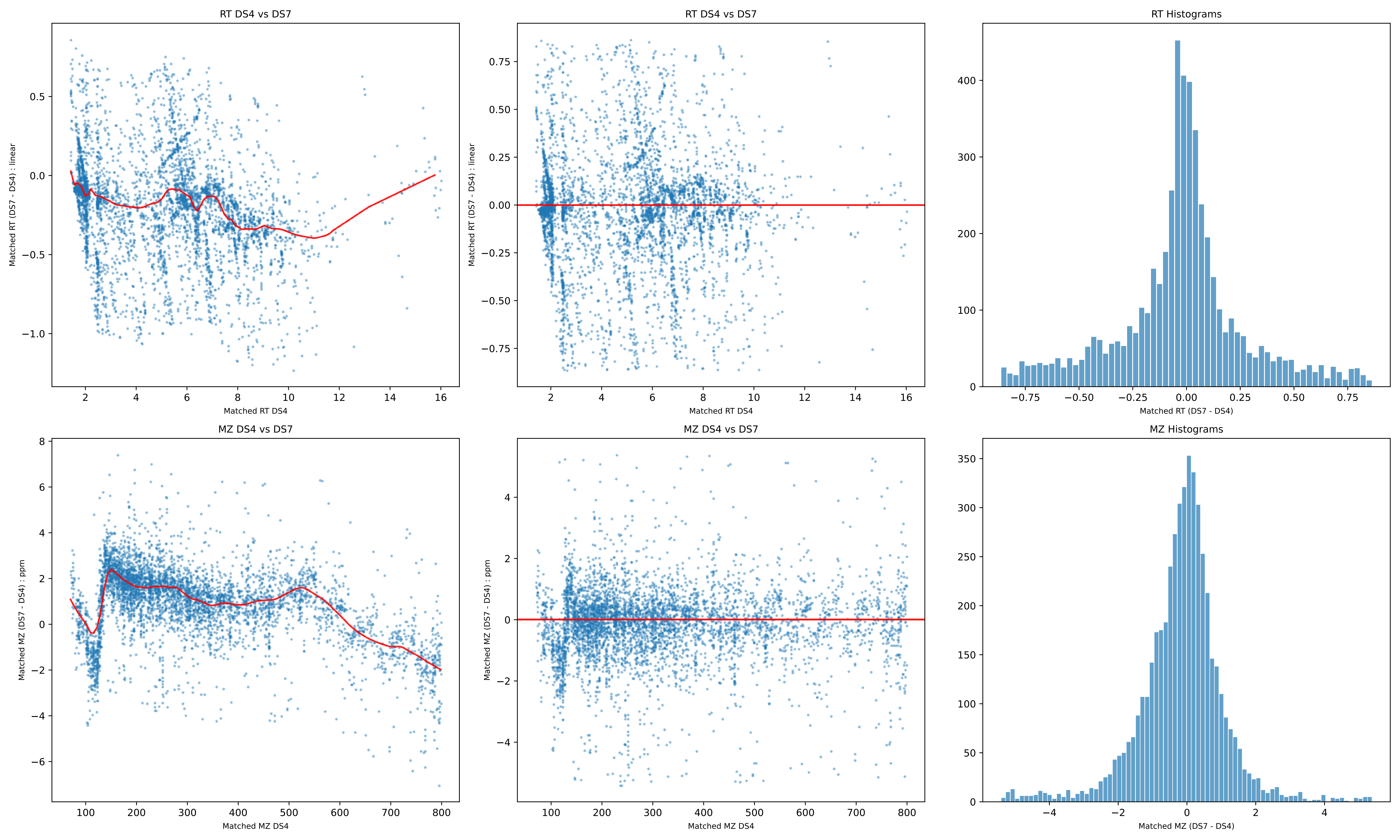
RT DS4 vs DS7
Matched RT (DS7 - DS4) : linear
Matched RT DS4
RT DS4 vs DS7
Matched RT (DS7 - DS4) : linear
Matched RT DS4
RT Histograms
Matched RT (DS7 - DS4)
MZ DS4 vs DS7
Matched MZ (DS7 - DS4) : ppm
Matched MZ DS4
MZ DS4 vs DS7
Matched MZ (DS7 - DS4) : ppm
Matched MZ DS4
MZ Histograms
Matched MZ (DS7 - DS4)

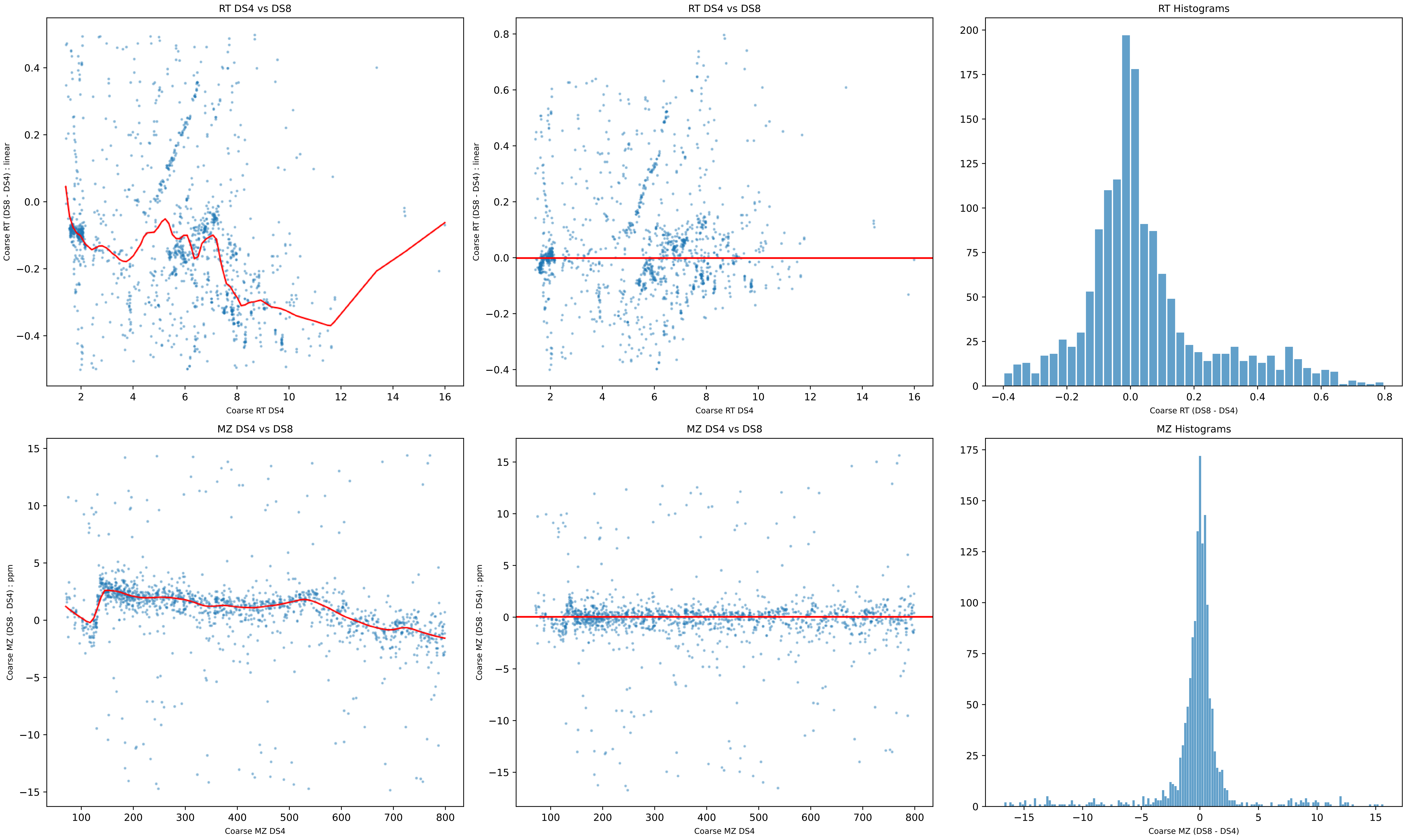

RT DS4 vs DS8
Coarse RT (DS8 - DS4) : linear
Coarse RT DS4
RT DS4 vs DS8
Coarse RT (DS8 - DS4) : linear
Coarse RT DS4
RT Histograms
Coarse RT (DS8 - DS4)
MZ DS4 vs DS8
Coarse MZ (DS8 - DS4) : ppm
Coarse MZ DS4
MZ DS4 vs DS8
Coarse MZ (DS8 - DS4) : ppm
Coarse MZ DS4
MZ Histograms
Coarse MZ (DS8 - DS4)

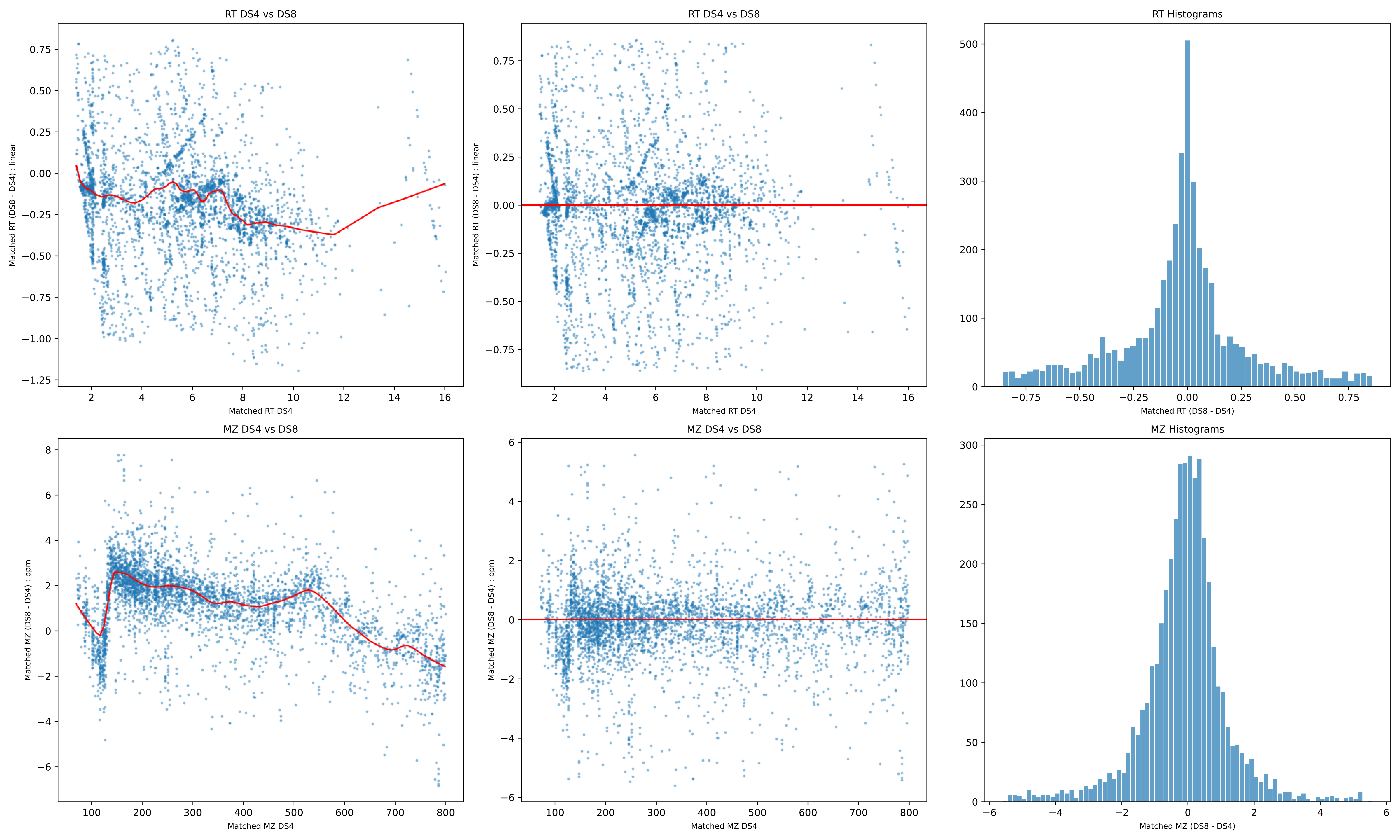

RT DS4 vs DS8
Matched RT (DS8 - DS4) : linear
Matched RT DS4
RT DS4 vs DS8
Matched RT (DS8 - DS4) : linear
Matched RT DS4
RT Histograms
Matched RT (DS8 - DS4)
MZ DS4 vs DS8
Matched MZ (DS8 - DS4) : ppm
Matched MZ DS4
MZ DS4 vs DS8
Matched MZ (DS8 - DS4) : ppm
Matched MZ DS4
MZ Histograms
Matched MZ (DS8 - DS4)

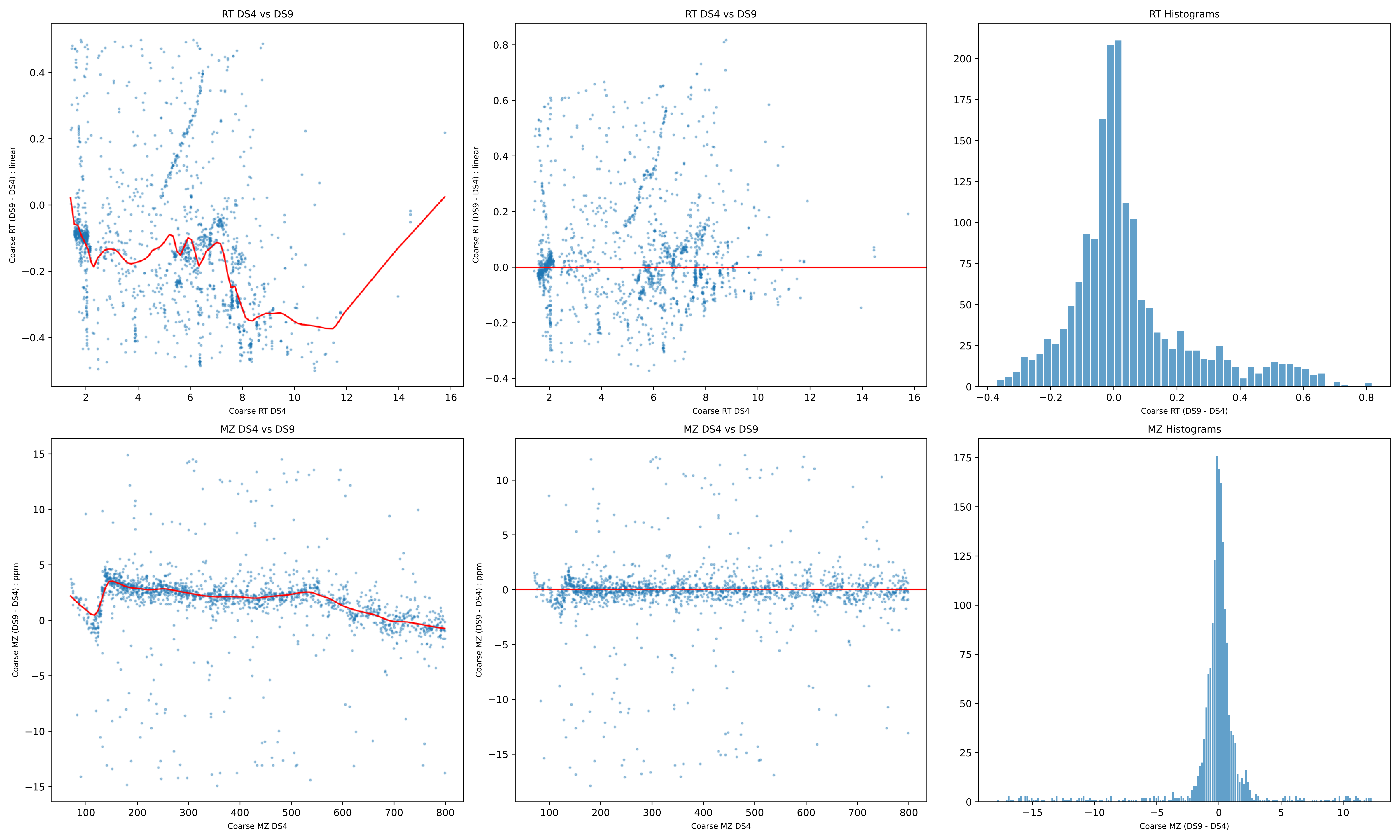
RT DS4 vs DS9
Coarse RT (DS9 - DS4) : linear
Coarse RT DS4
RT DS4 vs DS9
Coarse RT (DS9 - DS4) : linear
Coarse RT DS4
RT Histograms
Coarse RT (DS9 - DS4)
MZ DS4 vs DS9
Coarse MZ (DS9 - DS4) : ppm
Coarse MZ DS4
MZ DS4 vs DS9
Coarse MZ (DS9 - DS4) : ppm
Coarse MZ DS4
MZ Histograms
Coarse MZ (DS9 - DS4)

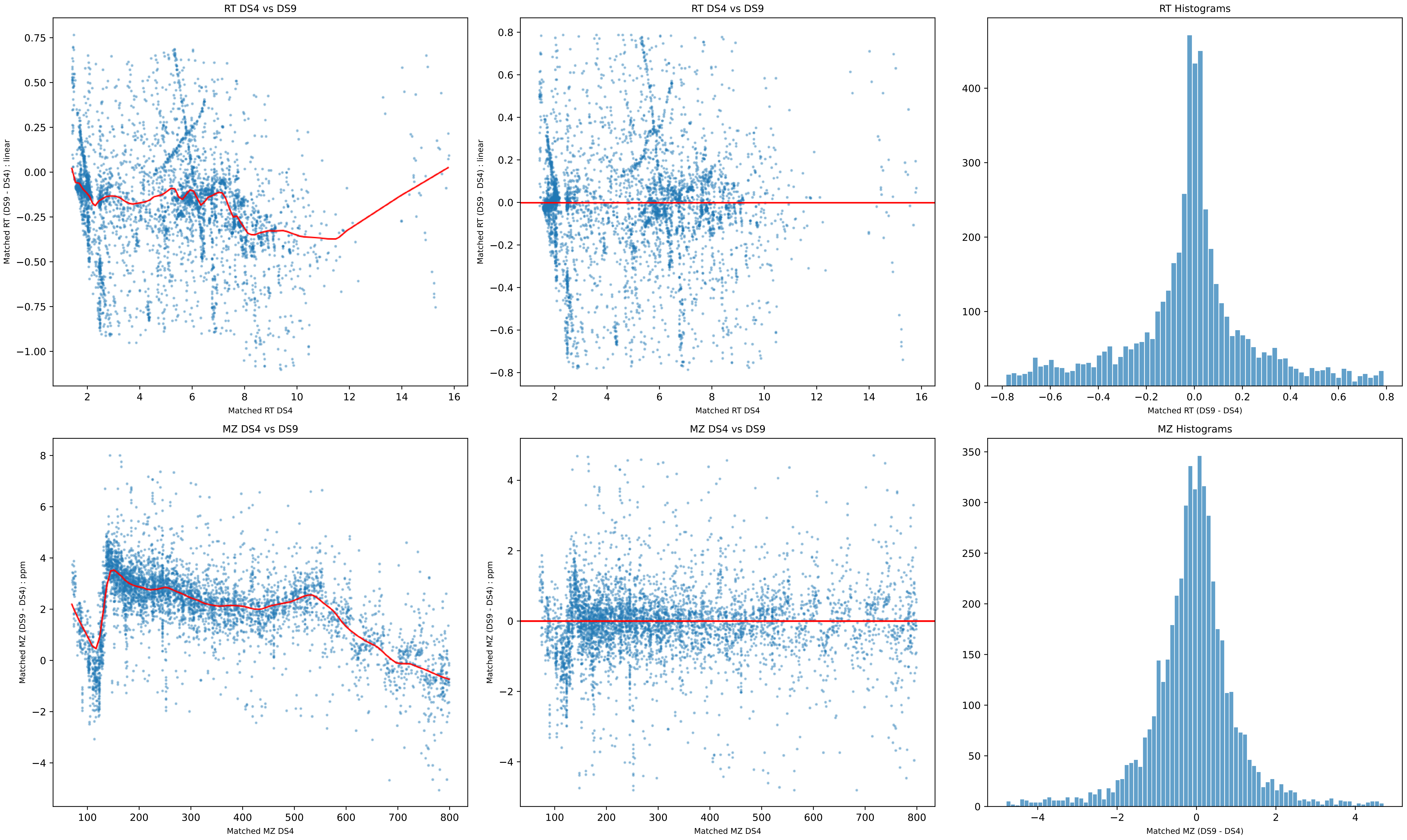
RT DS4 vs DS9
Matched RT (DS9 - DS4) : linear
Matched RT DS4
RT DS4 vs DS9
Matched RT (DS9 - DS4) : linear
Matched RT DS4
RT Histograms
Matched RT (DS9 - DS4)
MZ DS4 vs DS9
Matched MZ (DS9 - DS4) : ppm
Matched MZ DS4
MZ DS4 vs DS9
Matched MZ (DS9 - DS4) : ppm
Matched MZ DS4
MZ Histograms
Matched MZ (DS9 - DS4)

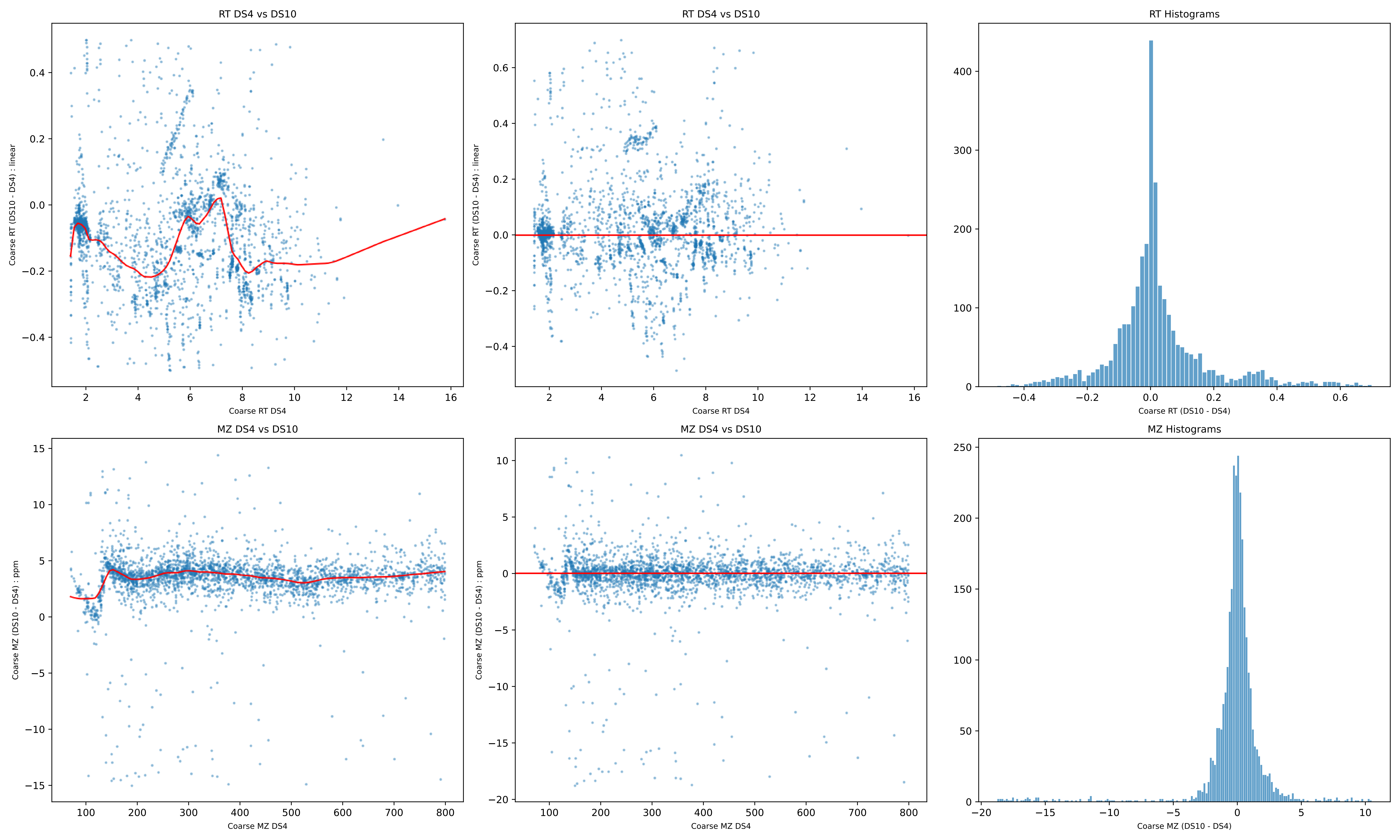

RT DS4 vs DS10
Coarse RT (DS10 - DS4) : linear
Coarse RT DS4
RT DS4 vs DS10
Coarse RT (DS10 - DS4) : linear
Coarse RT DS4
RT Histograms
Coarse RT (DS10 - DS4)
MZ DS4 vs DS10
Coarse MZ (DS10 - DS4) : ppm
Coarse MZ DS4
MZ DS4 vs DS10
Coarse MZ (DS10 - DS4) : ppm
Coarse MZ DS4
MZ Histograms
Coarse MZ (DS10 - DS4)

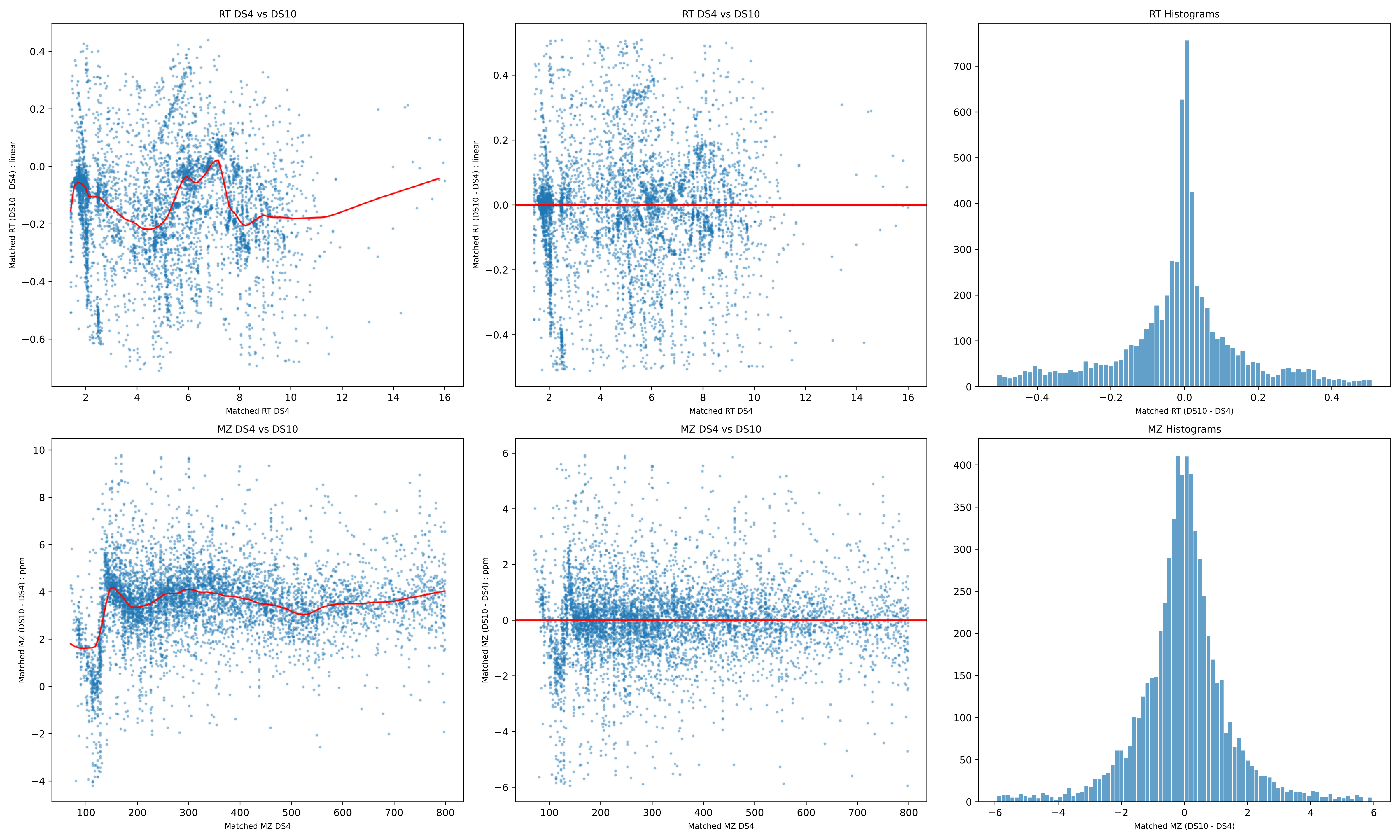
RT DS4 vs DS10
Matched RT (DS10 - DS4) : linear
Matched RT DS4
RT DS4 vs DS10
Matched RT (DS10 - DS4) : linear
Matched RT DS4
RT Histograms
Matched RT (DS10 - DS4)
MZ DS4 vs DS10
Matched MZ (DS10 - DS4) : ppm
Matched MZ DS4
MZ DS4 vs DS10
Matched MZ (DS10 - DS4) : ppm
Matched MZ DS4
MZ Histograms
Matched MZ (DS10 - DS4)

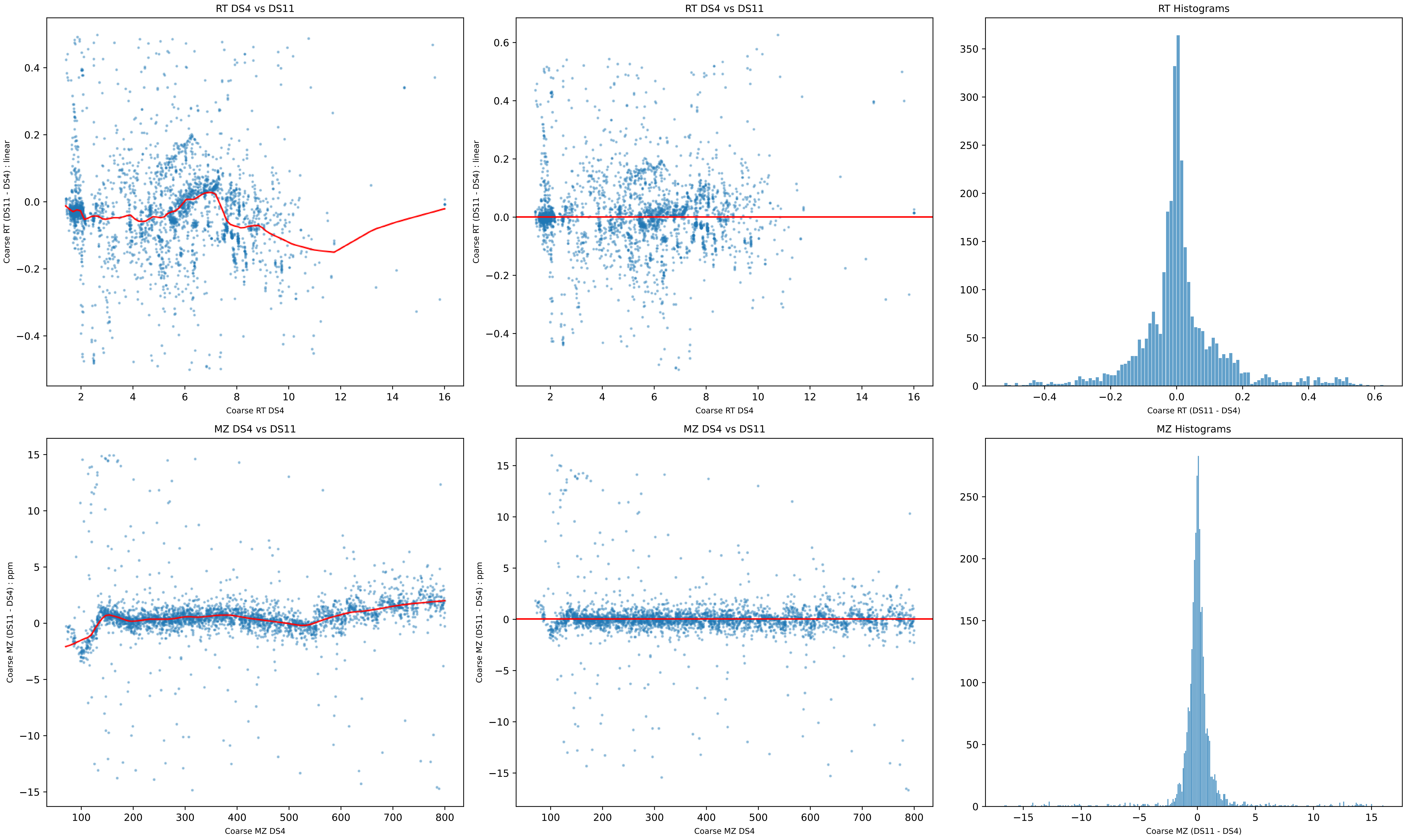
RT DS4 vs DS11
Coarse RT (DS11 - DS4) : linear
Coarse RT DS4
RT DS4 vs DS11
Coarse RT (DS11 - DS4) : linear
Coarse RT DS4
RT Histograms
Coarse RT (DS11 - DS4)
MZ DS4 vs DS11
Coarse MZ (DS11 - DS4) : ppm
Coarse MZ DS4
MZ DS4 vs DS11
Coarse MZ (DS11 - DS4) : ppm
Coarse MZ DS4
MZ Histograms
Coarse MZ (DS11 - DS4)

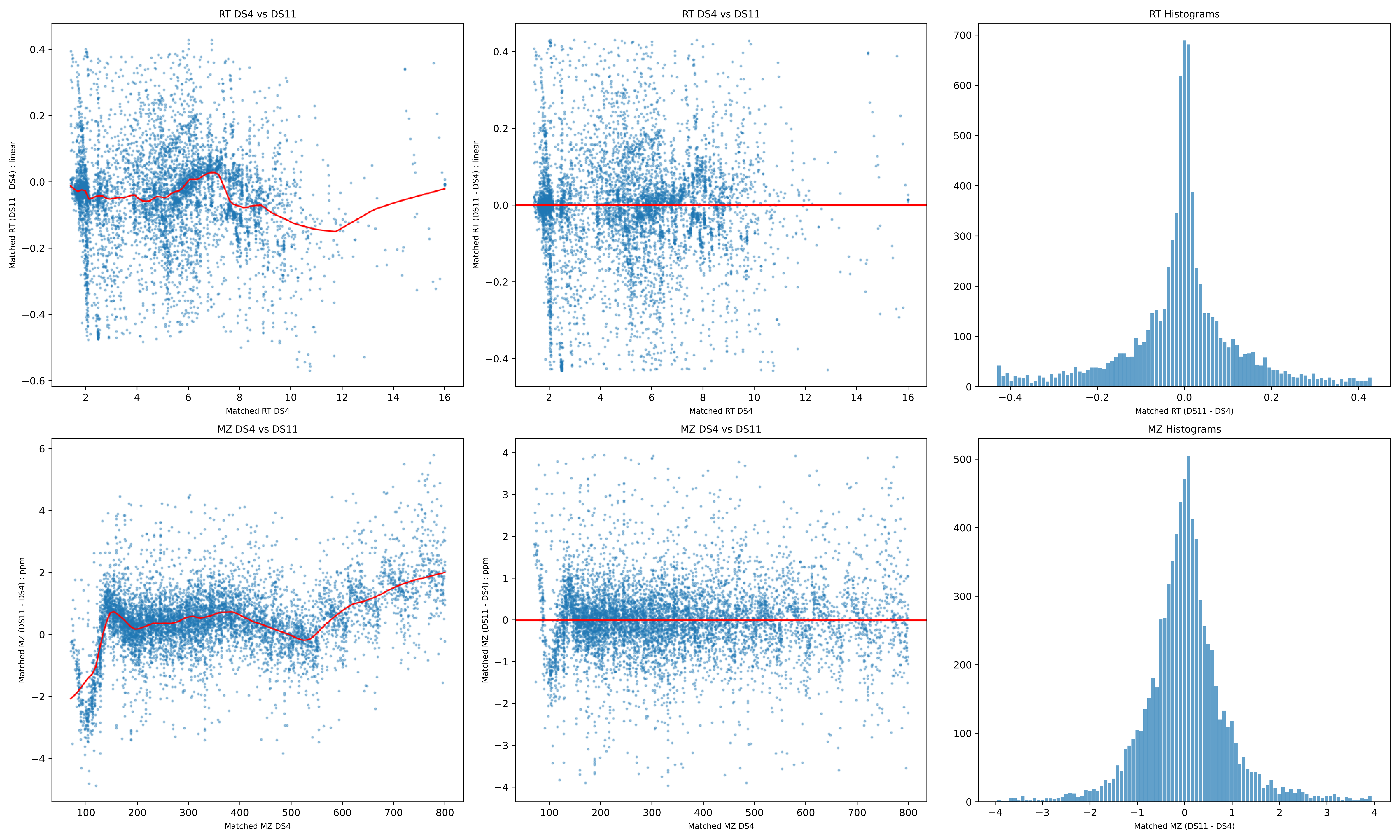
RT DS4 vs DS11
Matched RT (DS11 - DS4) : linear
Matched RT DS4
RT DS4 vs DS11
Matched RT (DS11 - DS4) : linear
Matched RT DS4
RT Histograms
Matched RT (DS11 - DS4)
MZ DS4 vs DS11
Matched MZ (DS11 - DS4) : ppm
Matched MZ DS4
MZ DS4 vs DS11
Matched MZ (DS11 - DS4) : ppm
Matched MZ DS4
MZ Histograms
Matched MZ (DS11 - DS4)

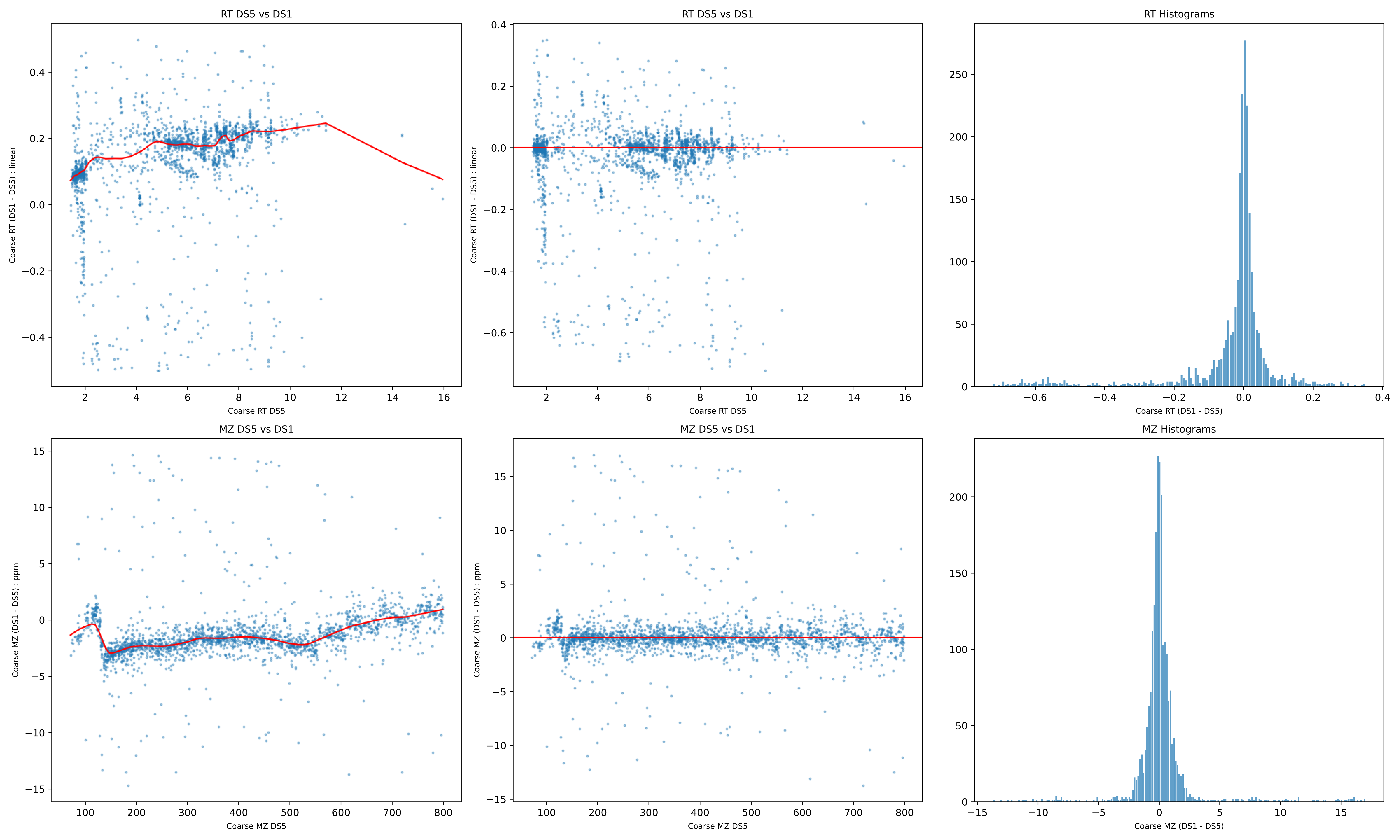
RT DS5 vs DS1
Coarse RT (DS1 - DS5) : linear
Coarse RT DS5
RT DS5 vs DS1
Coarse RT (DS1 - DS5) : linear
Coarse RT DS5
RT Histograms
Coarse RT (DS1 - DS5)
MZ DS5 vs DS1
Coarse MZ (DS1 - DS5) : ppm
Coarse MZ DS5
MZ DS5 vs DS1
Coarse MZ (DS1 - DS5) : ppm
Coarse MZ DS5
MZ Histograms
Coarse MZ (DS1 - DS5)

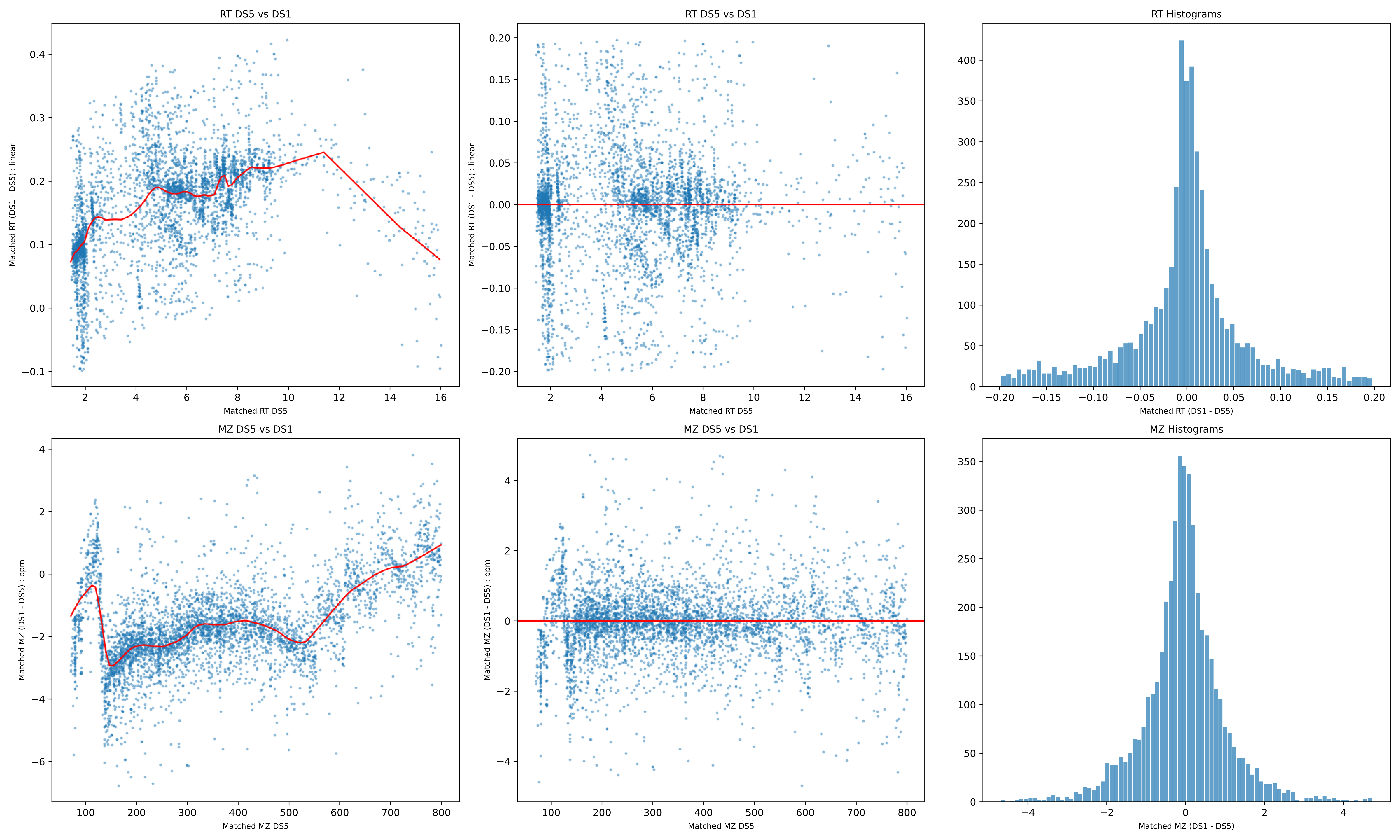

RT DS5 vs DS1
Matched RT (DS1 - DS5) : linear
Matched RT DS5
RT DS5 vs DS1
Matched RT (DS1 - DS5) : linear
Matched RT DS5
RT Histograms
Matched RT (DS1 - DS5)
MZ DS5 vs DS1
Matched MZ (DS1 - DS5) : ppm
Matched MZ DS5
MZ DS5 vs DS1
Matched MZ (DS1 - DS5) : ppm
Matched MZ DS5
MZ Histograms
Matched MZ (DS1 - DS5)

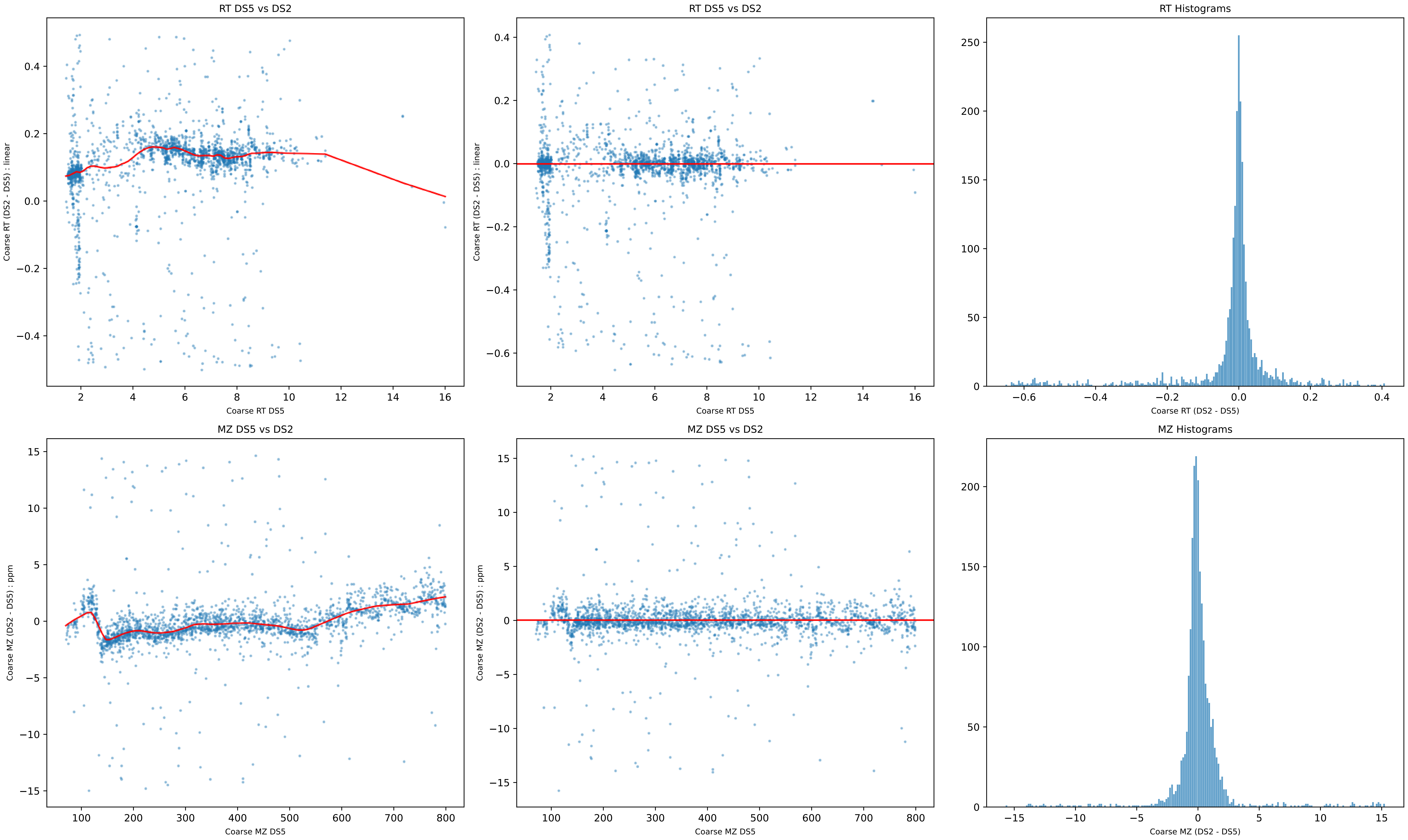

RT DS5 vs DS2
Coarse RT (DS2 - DS5) : linear
Coarse RT DS5
RT DS5 vs DS2
Coarse RT (DS2 - DS5) : linear
Coarse RT DS5
RT Histograms
Coarse RT (DS2 - DS5)
MZ DS5 vs DS2
Coarse MZ (DS2 - DS5) : ppm
Coarse MZ DS5
MZ DS5 vs DS2
Coarse MZ (DS2 - DS5) : ppm
Coarse MZ DS5
MZ Histograms
Coarse MZ (DS2 - DS5)

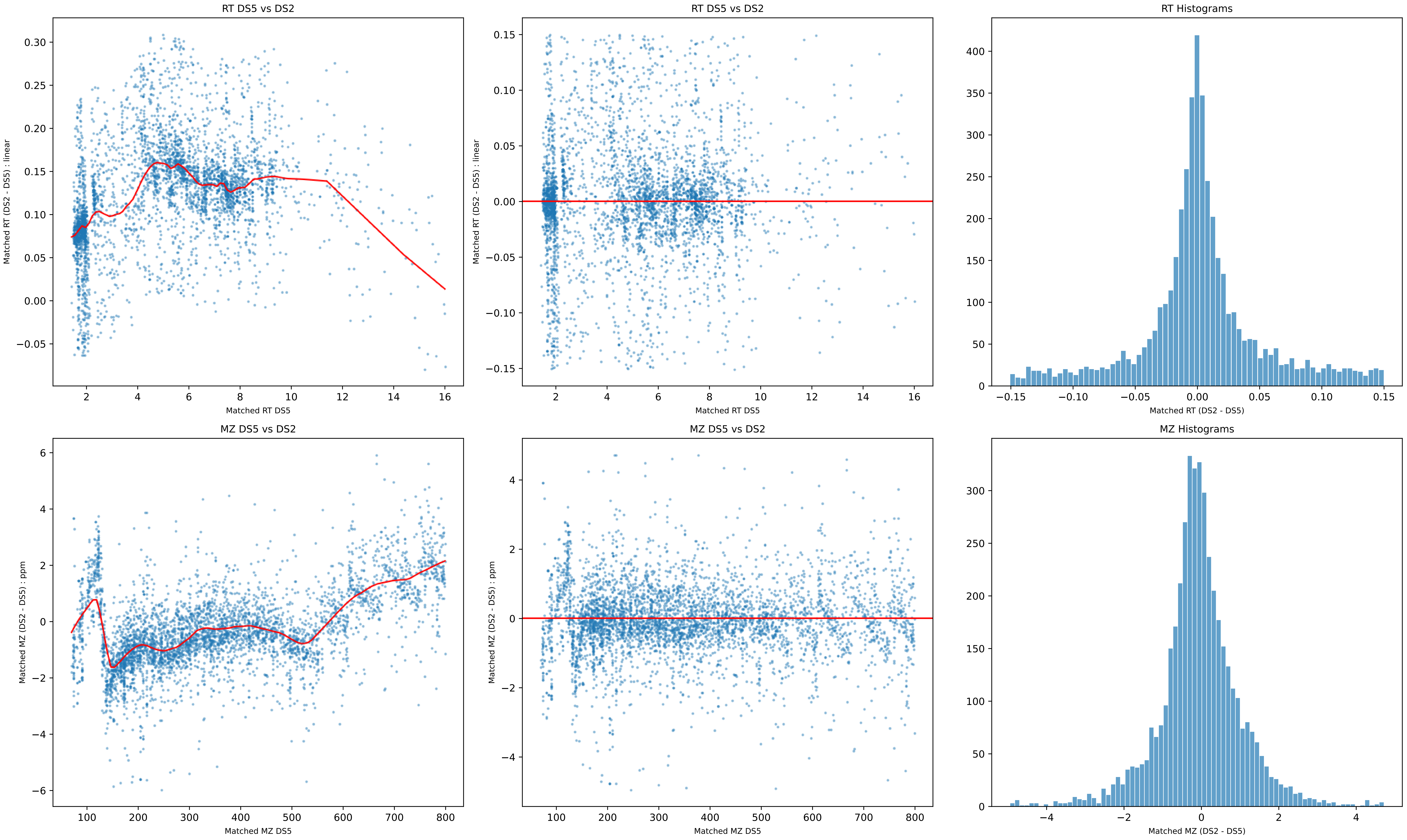
RT DS5 vs DS2
Matched RT (DS2 - DS5) : linear
Matched RT DS5
RT DS5 vs DS2
Matched RT (DS2 - DS5) : linear
Matched RT DS5
RT Histograms
Matched RT (DS2 - DS5)
MZ DS5 vs DS2
Matched MZ (DS2 - DS5) : ppm
Matched MZ DS5
MZ DS5 vs DS2
Matched MZ (DS2 - DS5) : ppm
Matched MZ DS5
MZ Histograms
Matched MZ (DS2 - DS5)

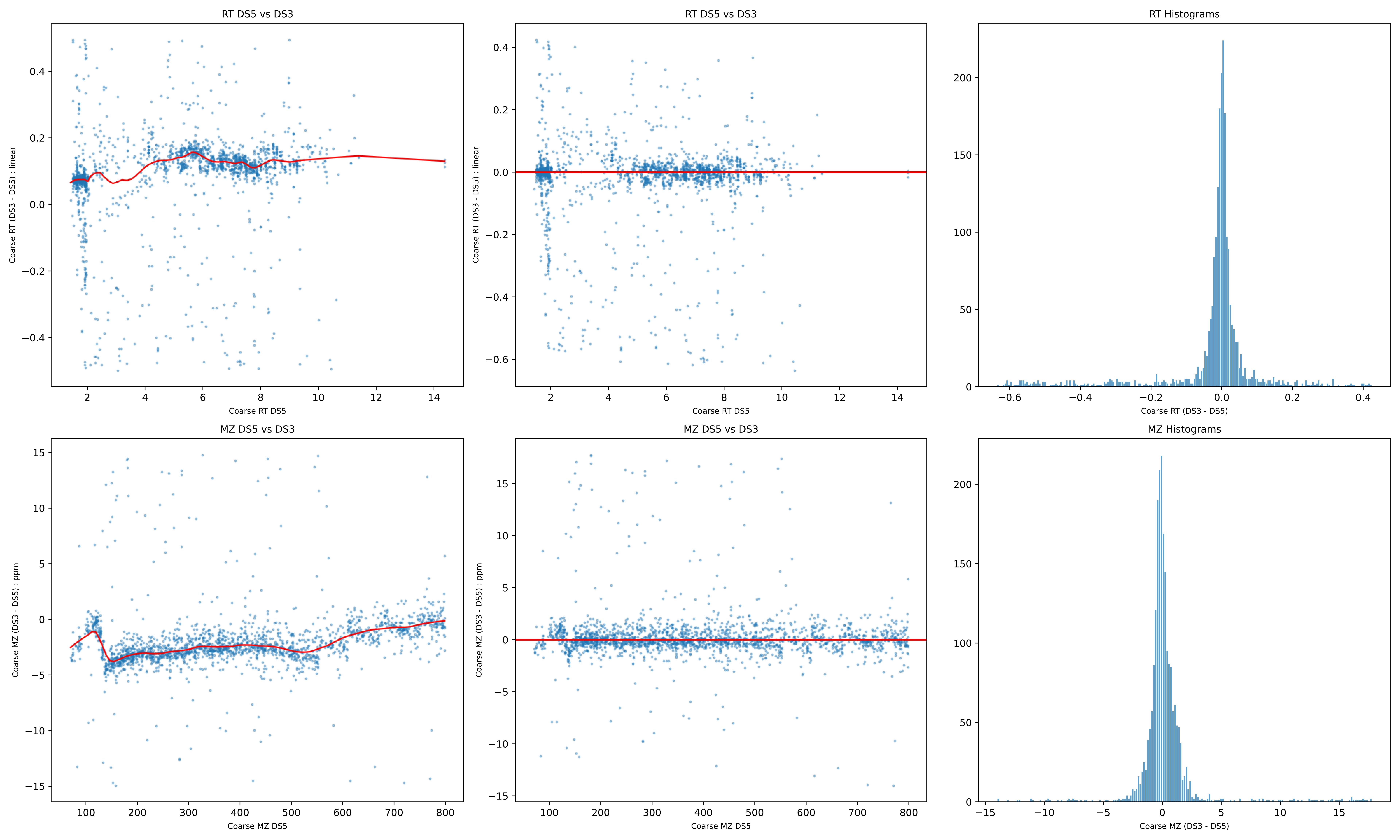

RT DS5 vs DS3
Coarse RT (DS3 - DS5) : linear
Coarse RT DS5
RT DS5 vs DS3
Coarse RT (DS3 - DS5) : linear
Coarse RT DS5
RT Histograms
Coarse RT (DS3 - DS5)
MZ DS5 vs DS3
Coarse MZ (DS3 - DS5) : ppm
Coarse MZ DS5
MZ DS5 vs DS3
Coarse MZ (DS3 - DS5) : ppm
Coarse MZ DS5
MZ Histograms
Coarse MZ (DS3 - DS5)

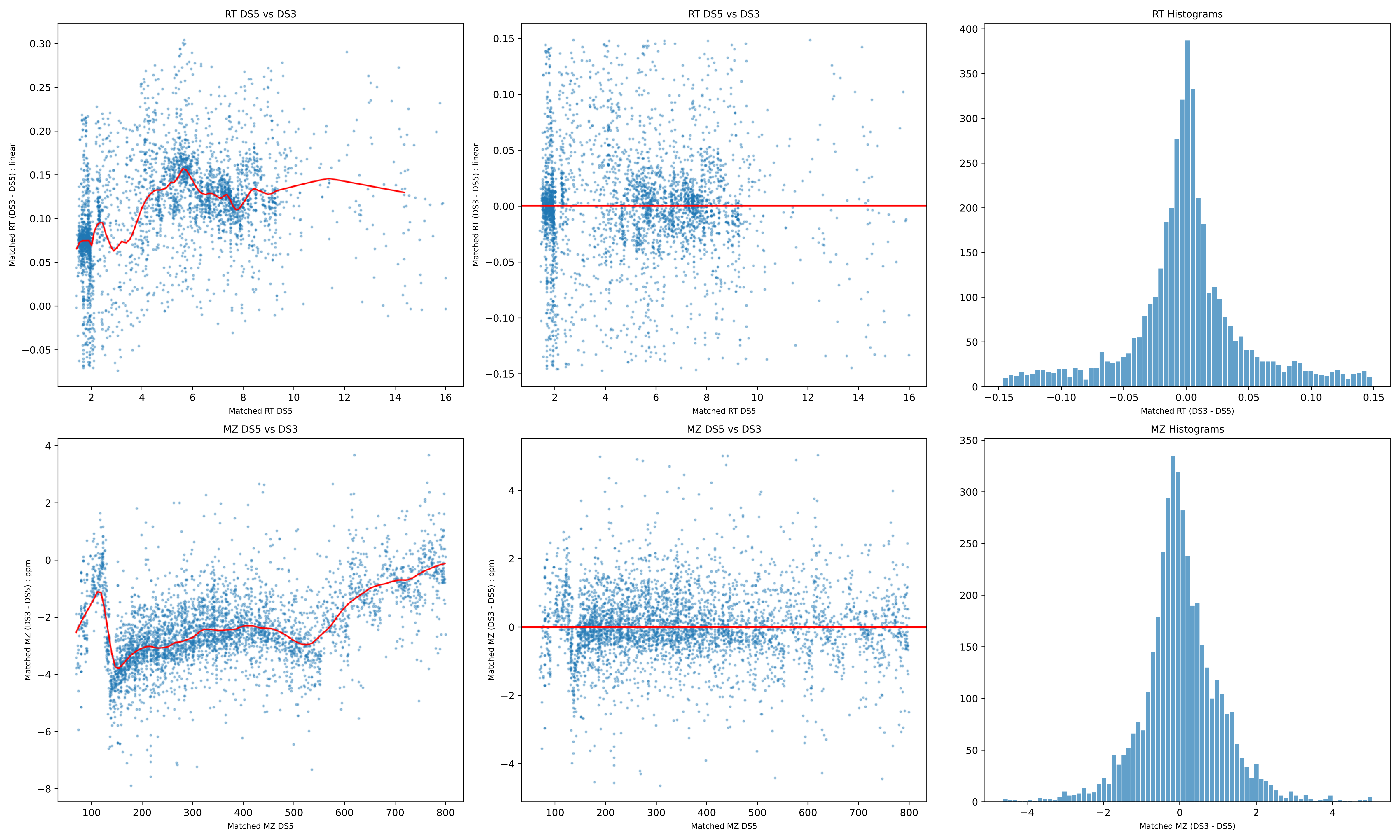
RT DS5 vs DS3
Matched RT (DS3 - DS5) : linear
Matched RT DS5
RT DS5 vs DS3
Matched RT (DS3 - DS5) : linear
Matched RT DS5
RT Histograms
Matched RT (DS3 - DS5)
MZ DS5 vs DS3
Matched MZ (DS3 - DS5) : ppm
Matched MZ DS5
MZ DS5 vs DS3
Matched MZ (DS3 - DS5) : ppm
Matched MZ DS5
MZ Histograms
Matched MZ (DS3 - DS5)

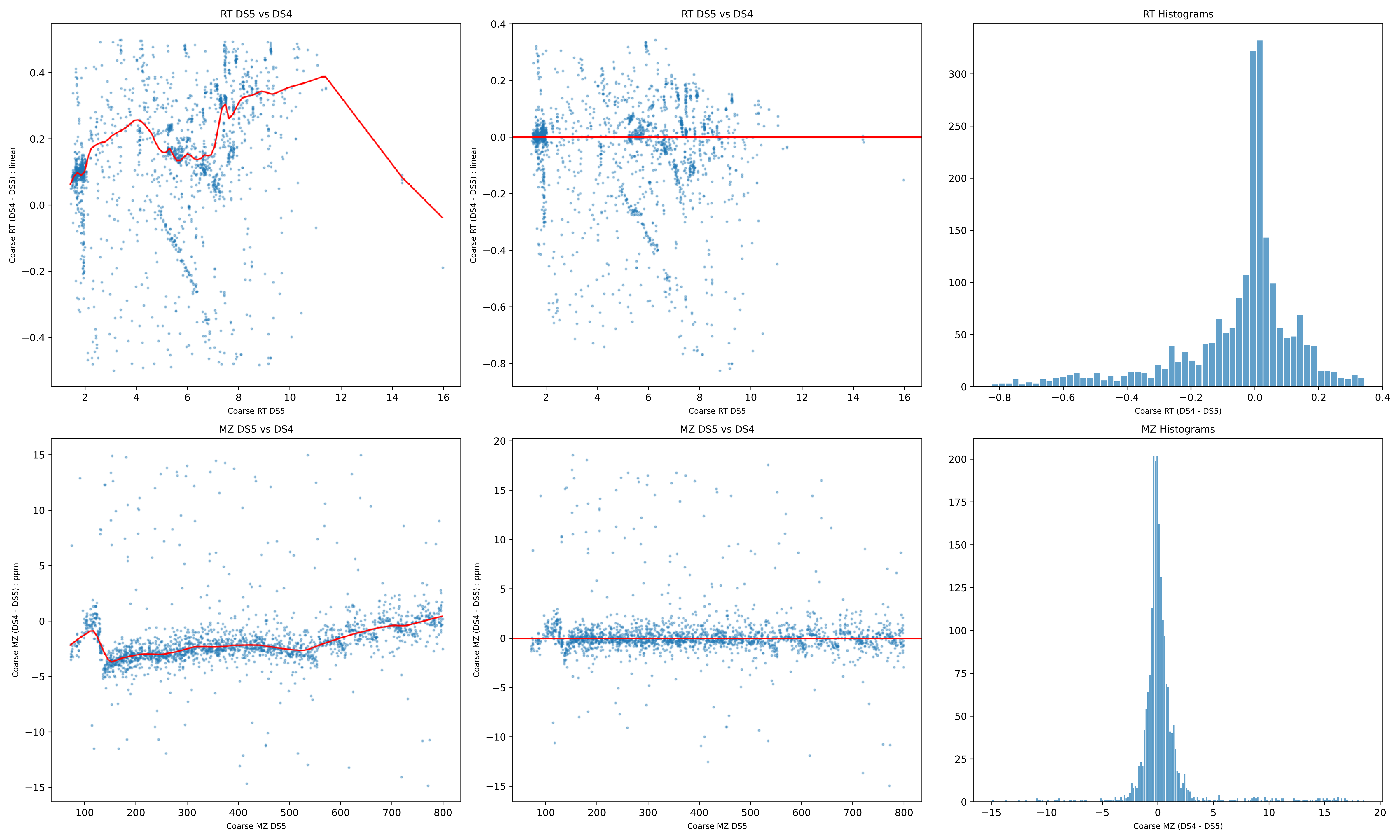

RT DS5 vs DS4
Coarse RT (DS4 - DS5) : linear
Coarse RT DS5
RT DS5 vs DS4
Coarse RT (DS4 - DS5) : linear
Coarse RT DS5
RT Histograms
Coarse RT (DS4 - DS5)
MZ DS5 vs DS4
Coarse MZ (DS4 - DS5) : ppm
Coarse MZ DS5
MZ DS5 vs DS4
Coarse MZ (DS4 - DS5) : ppm
Coarse MZ DS5
MZ Histograms
Coarse MZ (DS4 - DS5)

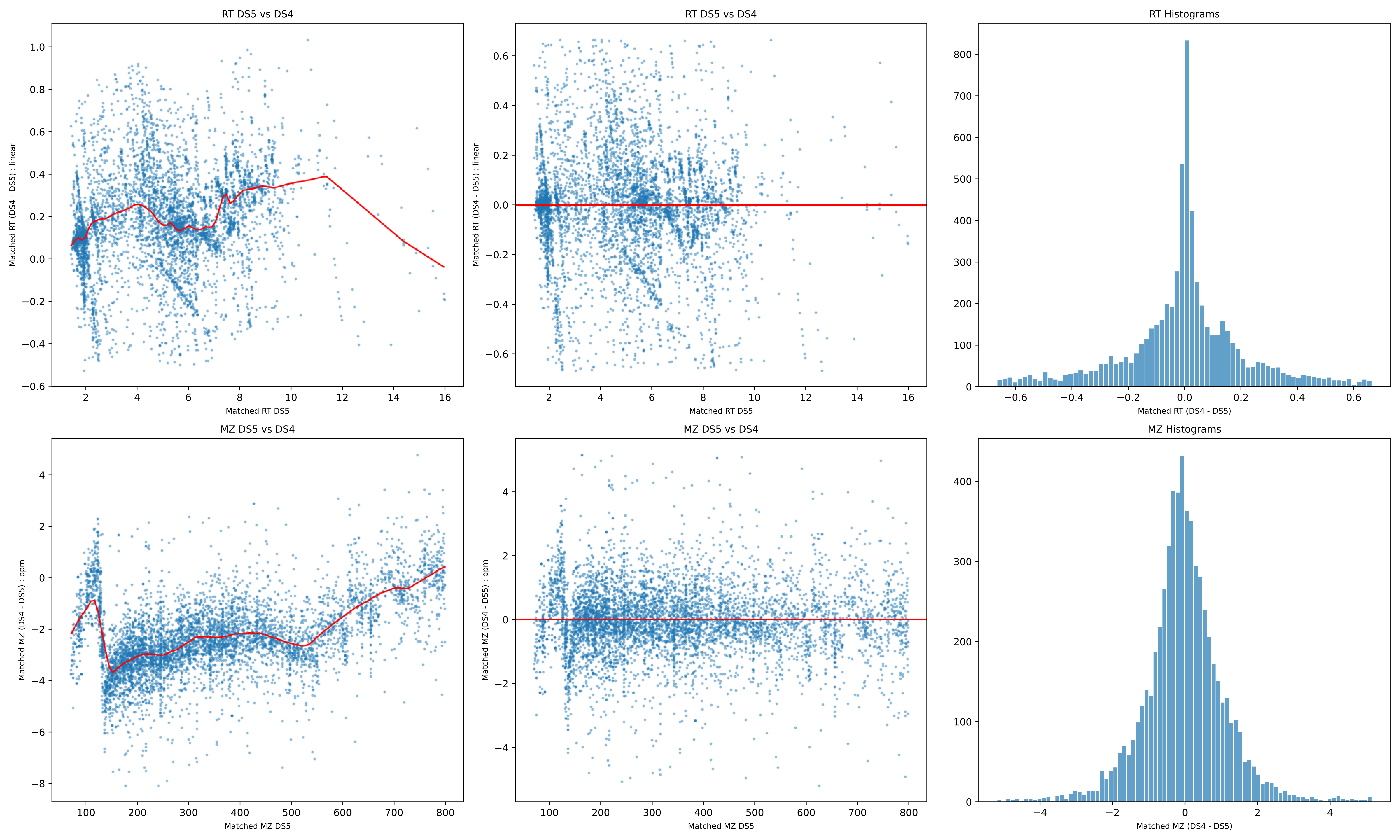

RT DS5 vs DS4
Matched RT (DS4 - DS5) : linear
Matched RT DS5
RT DS5 vs DS4
Matched RT (DS4 - DS5) : linear
Matched RT DS5
RT Histograms
Matched RT (DS4 - DS5)
MZ DS5 vs DS4
Matched MZ (DS4 - DS5) : ppm
Matched MZ DS5
MZ DS5 vs DS4
Matched MZ (DS4 - DS5) : ppm
Matched MZ DS5
MZ Histograms
Matched MZ (DS4 - DS5)

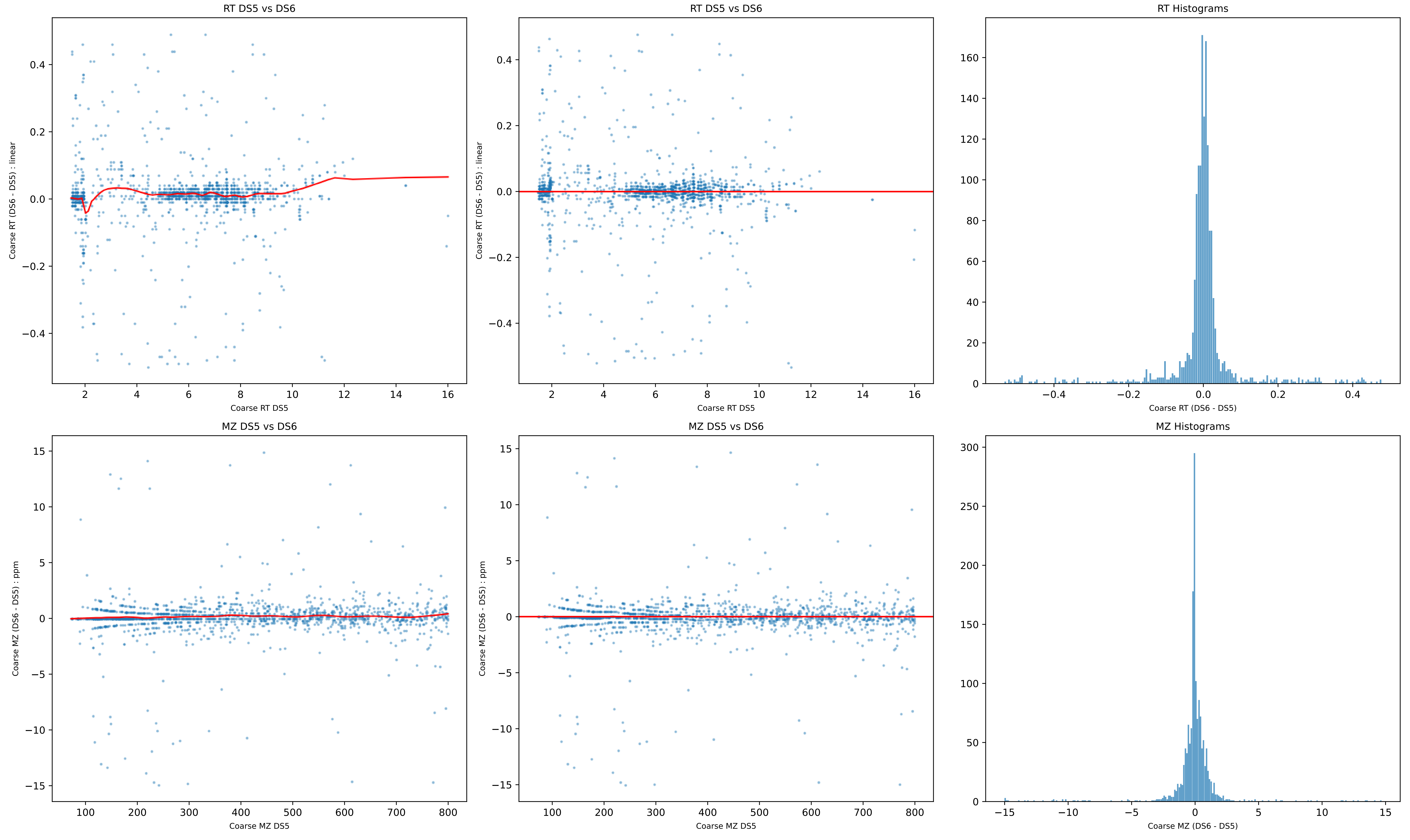

RT DS5 vs DS6
Coarse RT (DS6 - DS5) : linear
Coarse RT DS5
RT DS5 vs DS6
Coarse RT (DS6 - DS5) : linear
Coarse RT DS5
RT Histograms
Coarse RT (DS6 - DS5)
MZ DS5 vs DS6
Coarse MZ (DS6 - DS5) : ppm
Coarse MZ DS5
MZ DS5 vs DS6
Coarse MZ (DS6 - DS5) : ppm
Coarse MZ DS5
MZ Histograms
Coarse MZ (DS6 - DS5)

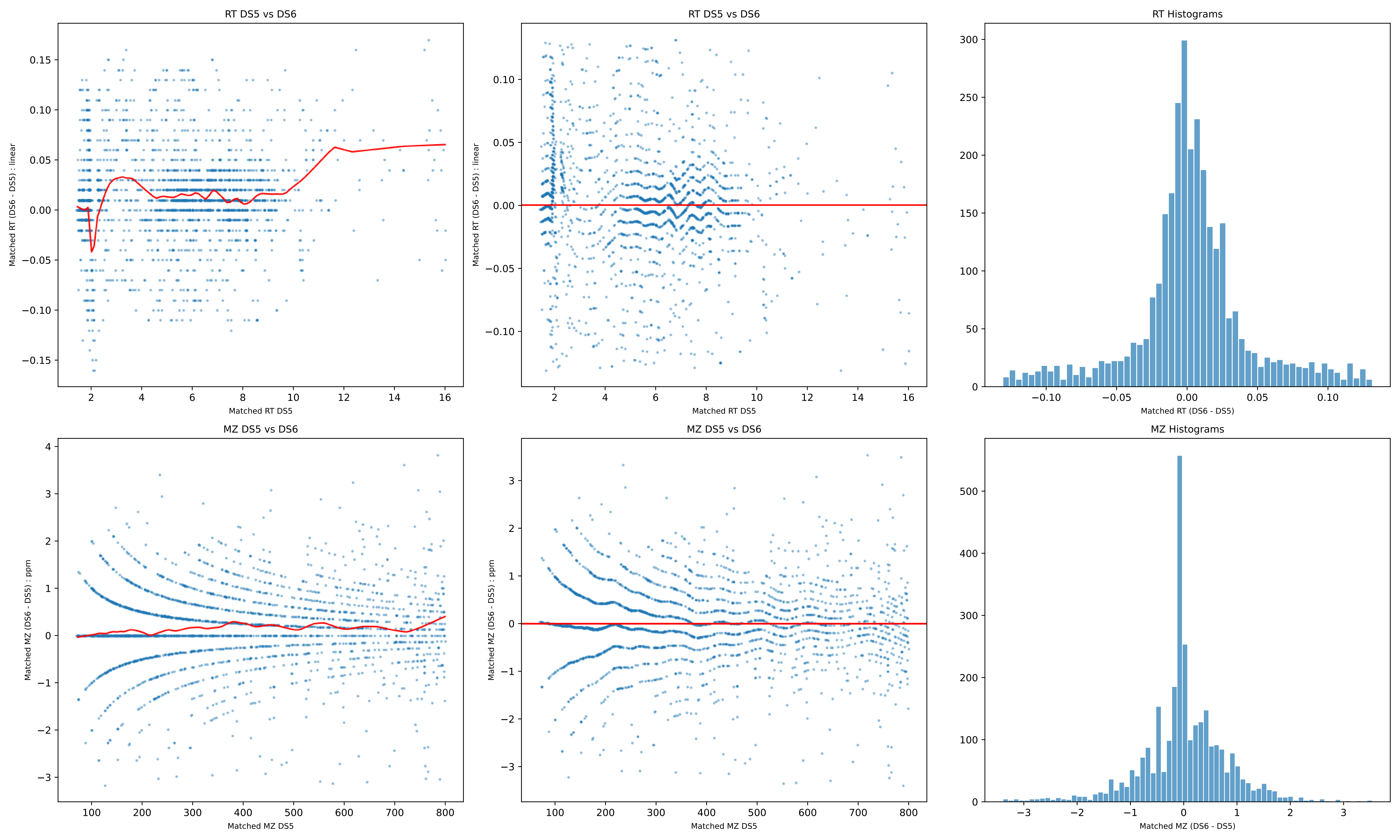
RT DS5 vs DS6
Matched RT (DS6 - DS5) : linear
Matched RT DS5
RT DS5 vs DS6
Matched RT (DS6 - DS5) : linear
Matched RT DS5
RT Histograms
Matched RT (DS6 - DS5)
MZ DS5 vs DS6
Matched MZ (DS6 - DS5) : ppm
Matched MZ DS5
MZ DS5 vs DS6
Matched MZ (DS6 - DS5) : ppm
Matched MZ DS5
MZ Histograms
Matched MZ (DS6 - DS5)

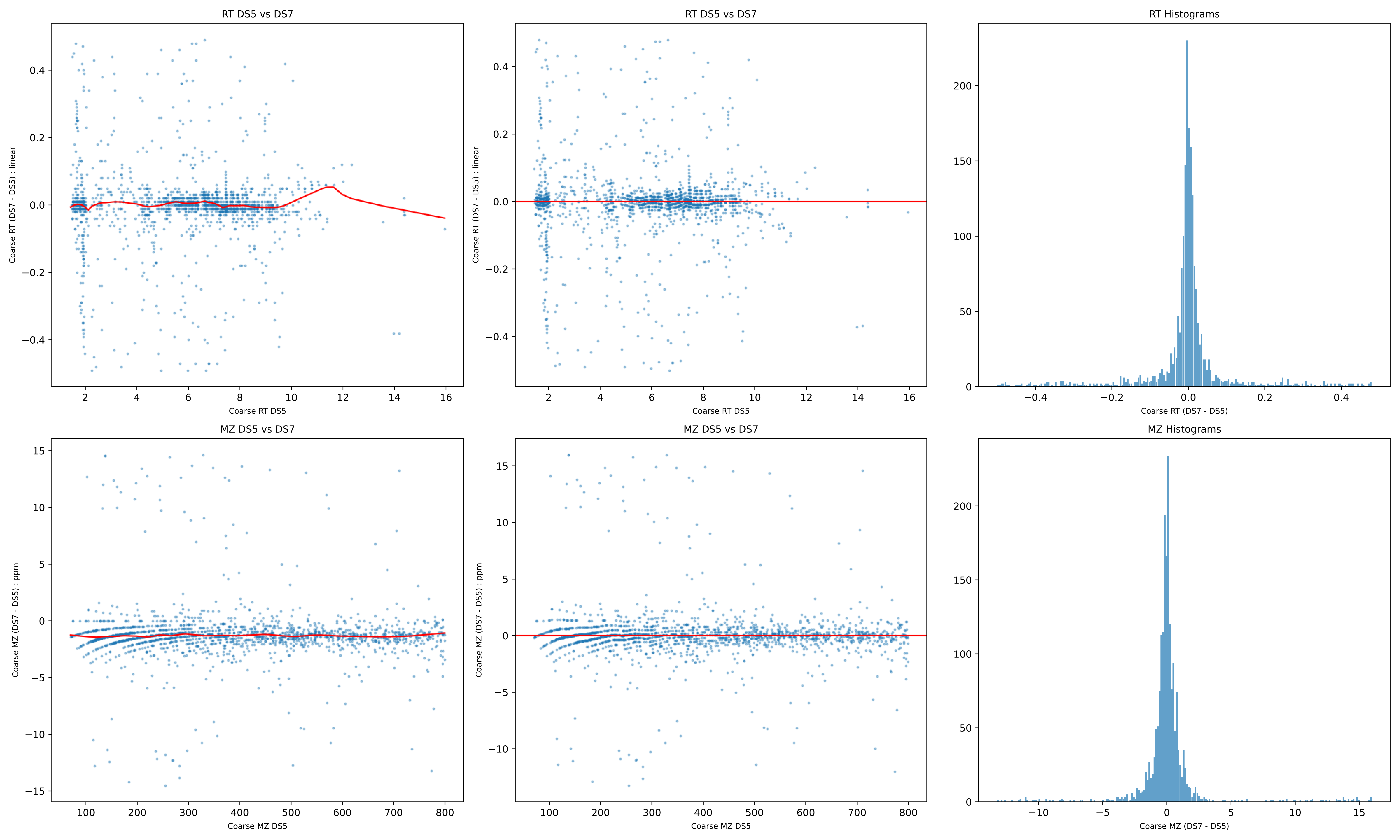
RT DS5 vs DS7
Coarse RT (DS7 - DS5) : linear
Coarse RT DS5
RT DS5 vs DS7
Coarse RT (DS7 - DS5) : linear
Coarse RT DS5
RT Histograms
Coarse RT (DS7 - DS5)
MZ DS5 vs DS7
Coarse MZ (DS7 - DS5) : ppm
Coarse MZ DS5
MZ DS5 vs DS7
Coarse MZ (DS7 - DS5) : ppm
Coarse MZ DS5
MZ Histograms
Coarse MZ (DS7 - DS5)

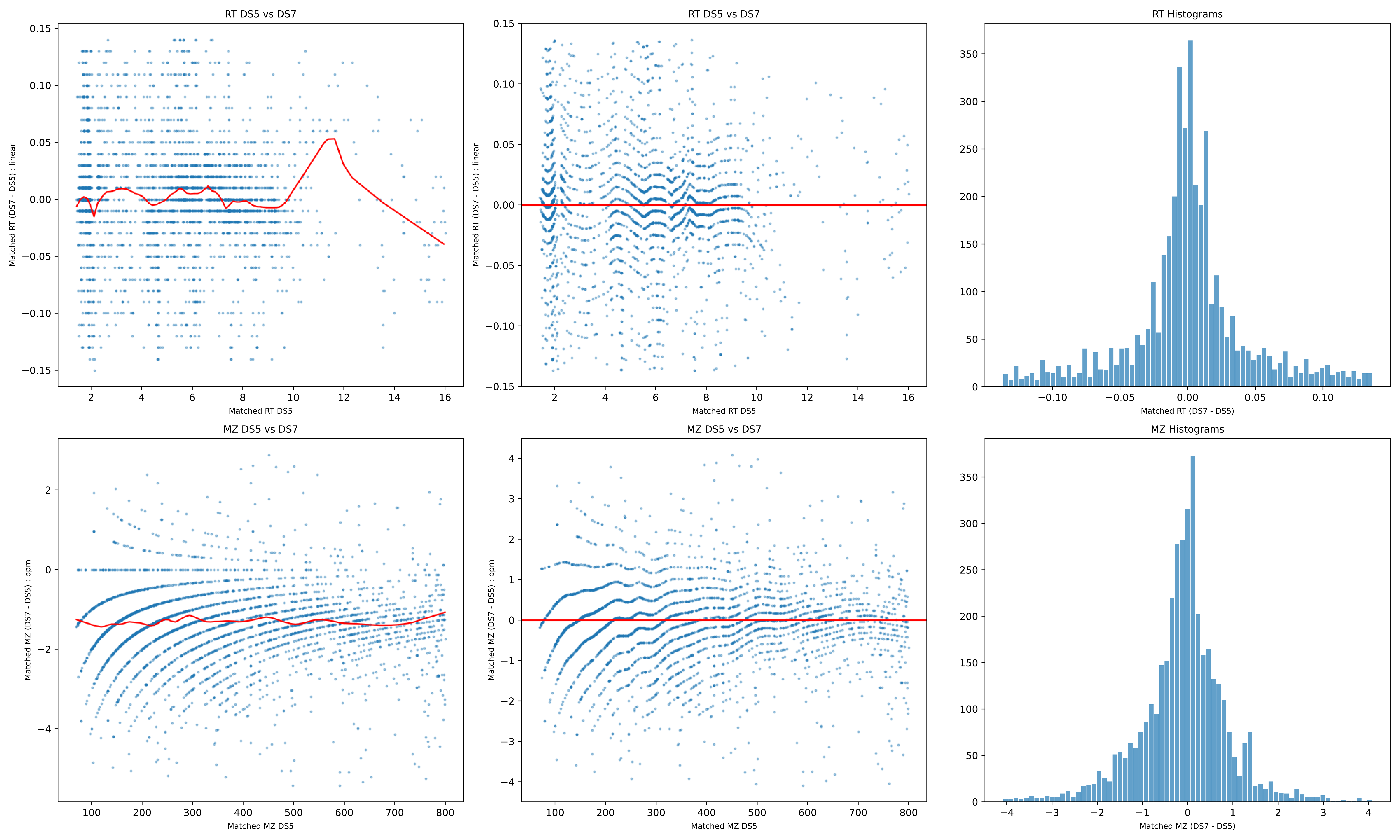
RT DS5 vs DS7
Matched RT (DS7 - DS5) : linear
Matched RT DS5
RT DS5 vs DS7
Matched RT (DS7 - DS5) : linear
Matched RT DS5
RT Histograms
Matched RT (DS7 - DS5)
MZ DS5 vs DS7
Matched MZ (DS7 - DS5) : ppm
Matched MZ DS5
MZ DS5 vs DS7
Matched MZ (DS7 - DS5) : ppm
Matched MZ DS5
MZ Histograms
Matched MZ (DS7 - DS5)

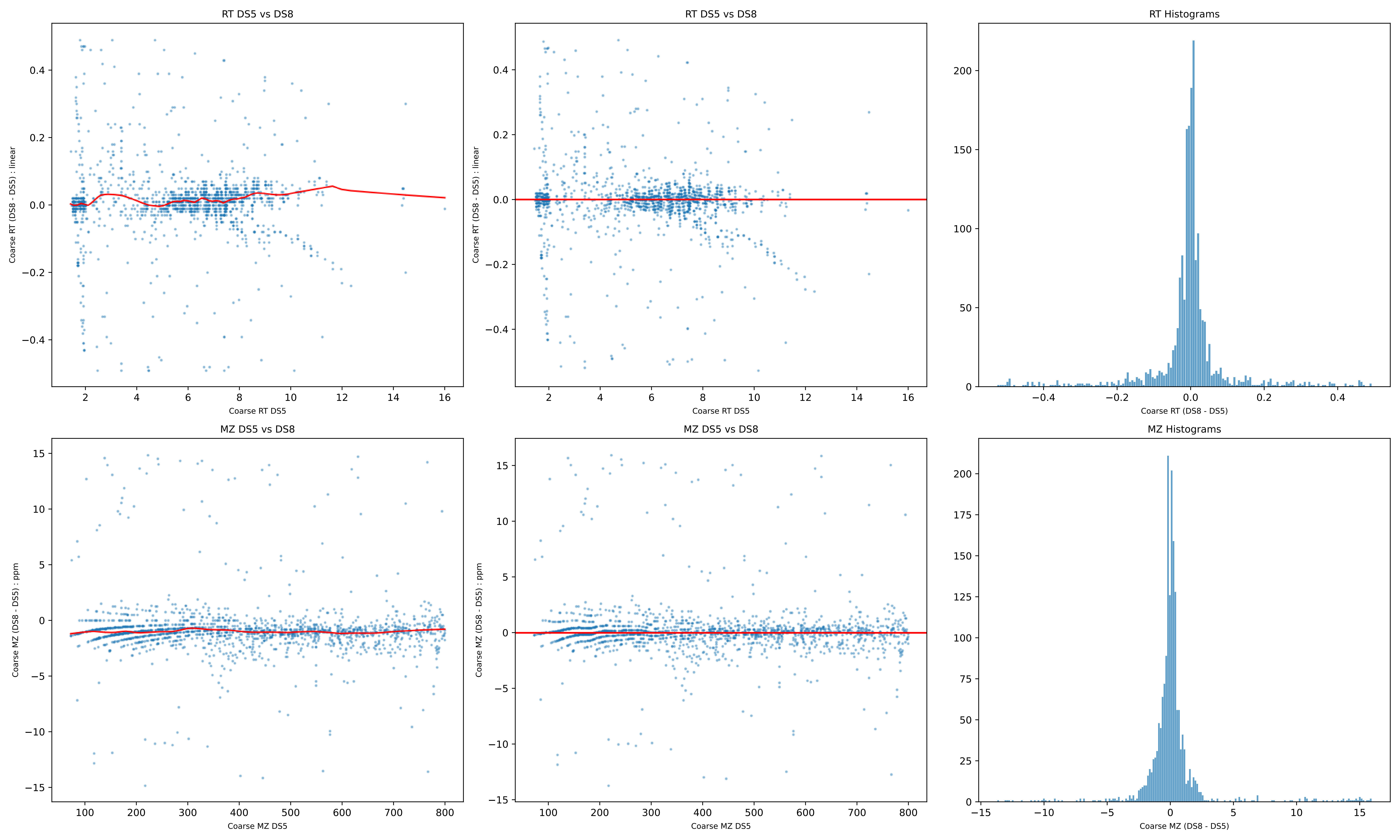
RT DS5 vs DS8
Coarse RT (DS8 - DS5) : linear
Coarse RT DS5
RT DS5 vs DS8
Coarse RT (DS8 - DS5) : linear
Coarse RT DS5
RT Histograms
Coarse RT (DS8 - DS5)
MZ DS5 vs DS8
Coarse MZ (DS8 - DS5) : ppm
Coarse MZ DS5
MZ DS5 vs DS8
Coarse MZ (DS8 - DS5) : ppm
Coarse MZ DS5
MZ Histograms
Coarse MZ (DS8 - DS5)

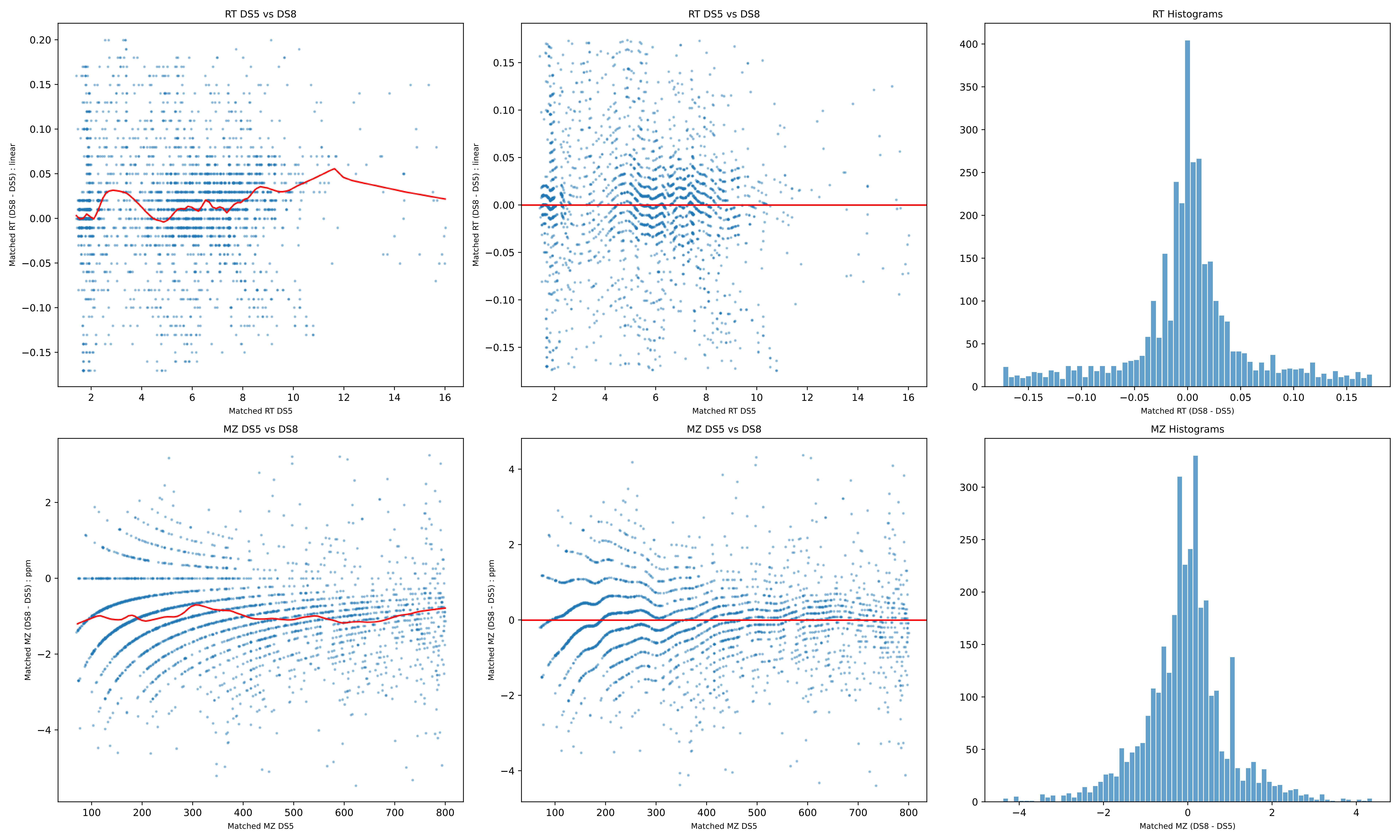
RT DS5 vs DS8
Matched RT (DS8 - DS5) : linear
Matched RT DS5
RT DS5 vs DS8
Matched RT (DS8 - DS5) : linear
Matched RT DS5
RT Histograms
Matched RT (DS8 - DS5)
MZ DS5 vs DS8
Matched MZ (DS8 - DS5) : ppm
Matched MZ DS5
MZ DS5 vs DS8
Matched MZ (DS8 - DS5) : ppm
Matched MZ DS5
MZ Histograms
Matched MZ (DS8 - DS5)

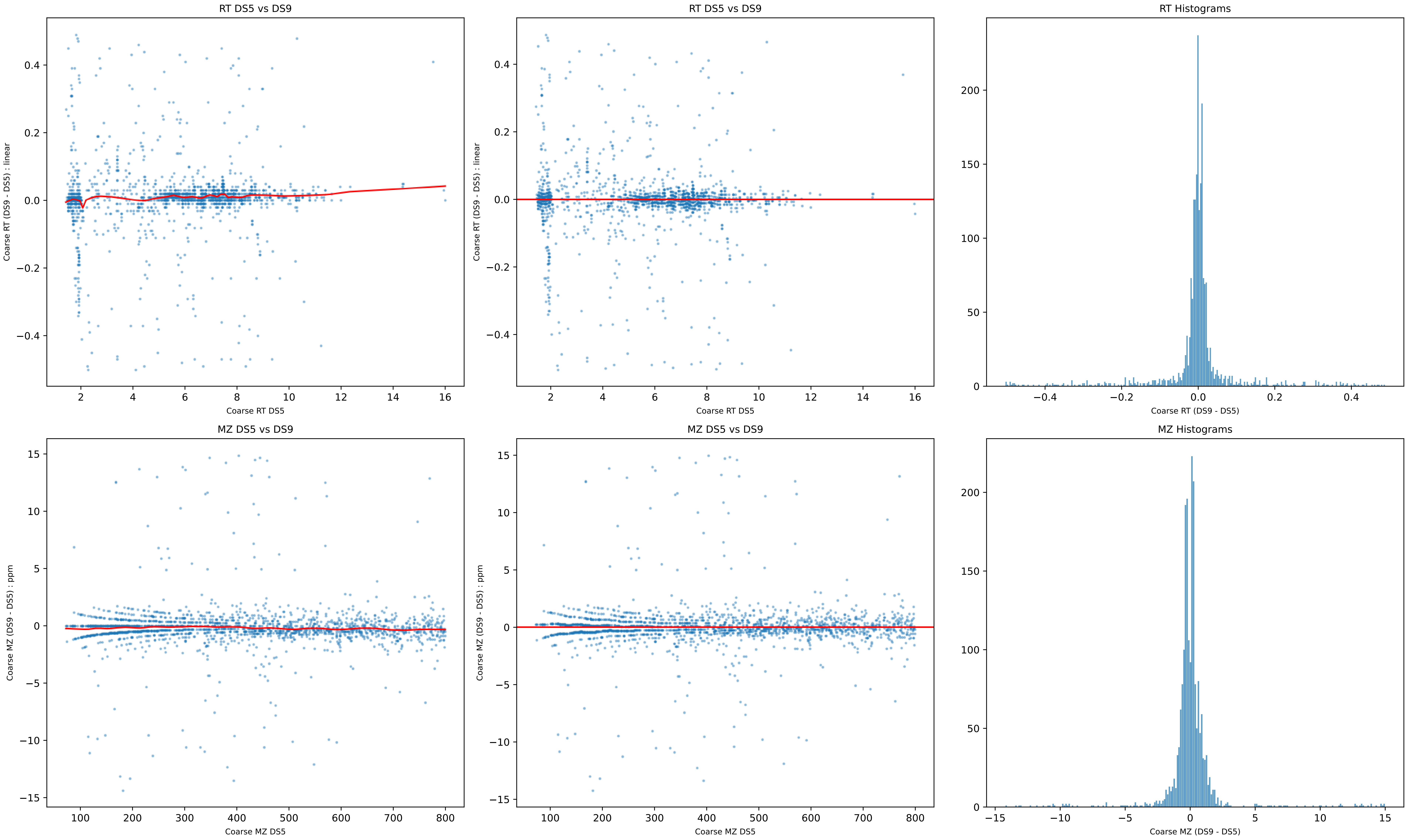
RT DS5 vs DS9
Coarse RT (DS9 - DS5) : linear
Coarse RT DS5
RT DS5 vs DS9
Coarse RT (DS9 - DS5) : linear
Coarse RT DS5
RT Histograms
Coarse RT (DS9 - DS5)
MZ DS5 vs DS9
Coarse MZ (DS9 - DS5) : ppm
Coarse MZ DS5
MZ DS5 vs DS9
Coarse MZ (DS9 - DS5) : ppm
Coarse MZ DS5
MZ Histograms
Coarse MZ (DS9 - DS5)

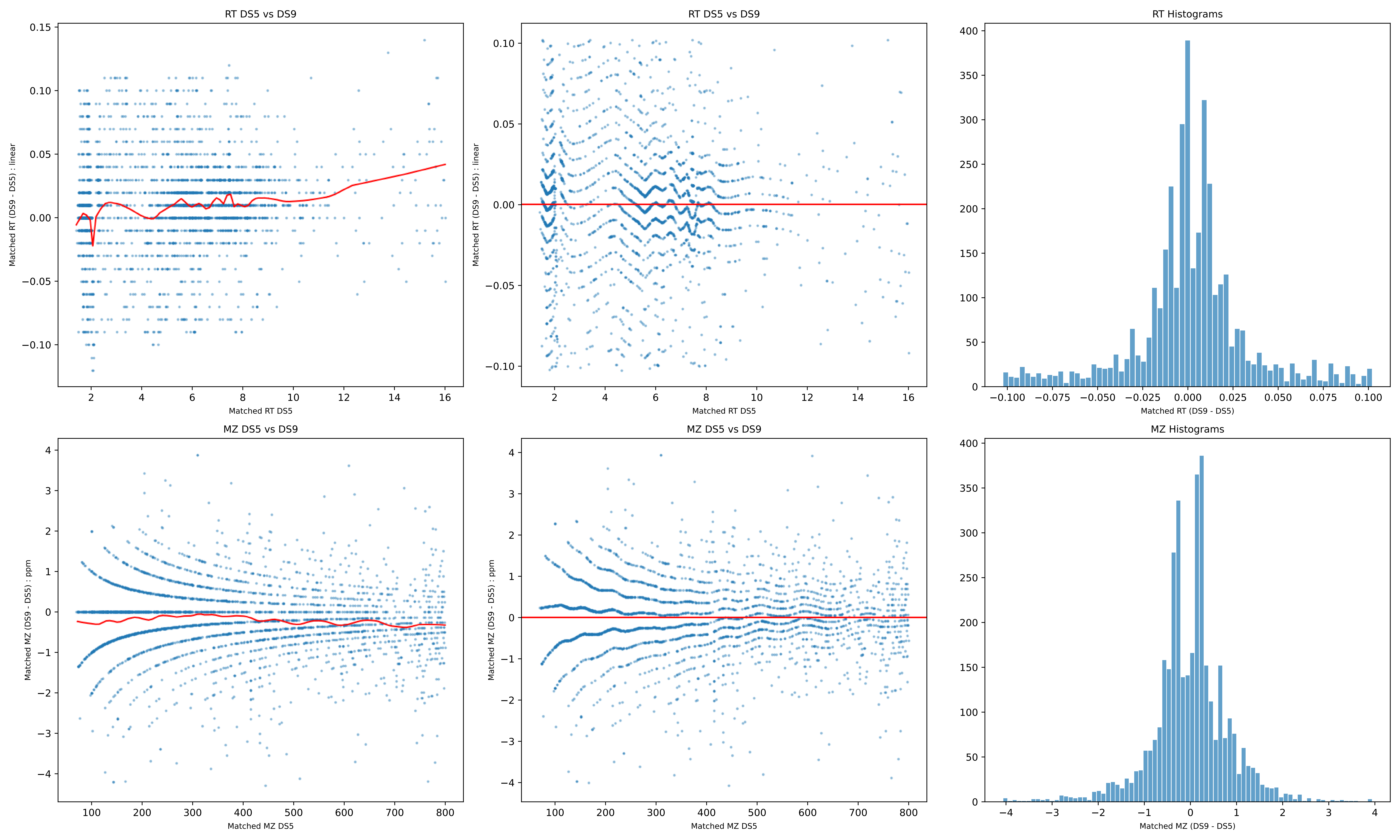
RT DS5 vs DS9
Matched RT (DS9 - DS5) : linear
Matched RT DS5
RT DS5 vs DS9
Matched RT (DS9 - DS5) : linear
Matched RT DS5
RT Histograms
Matched RT (DS9 - DS5)
MZ DS5 vs DS9
Matched MZ (DS9 - DS5) : ppm
Matched MZ DS5
MZ DS5 vs DS9
Matched MZ (DS9 - DS5) : ppm
Matched MZ DS5
MZ Histograms
Matched MZ (DS9 - DS5)

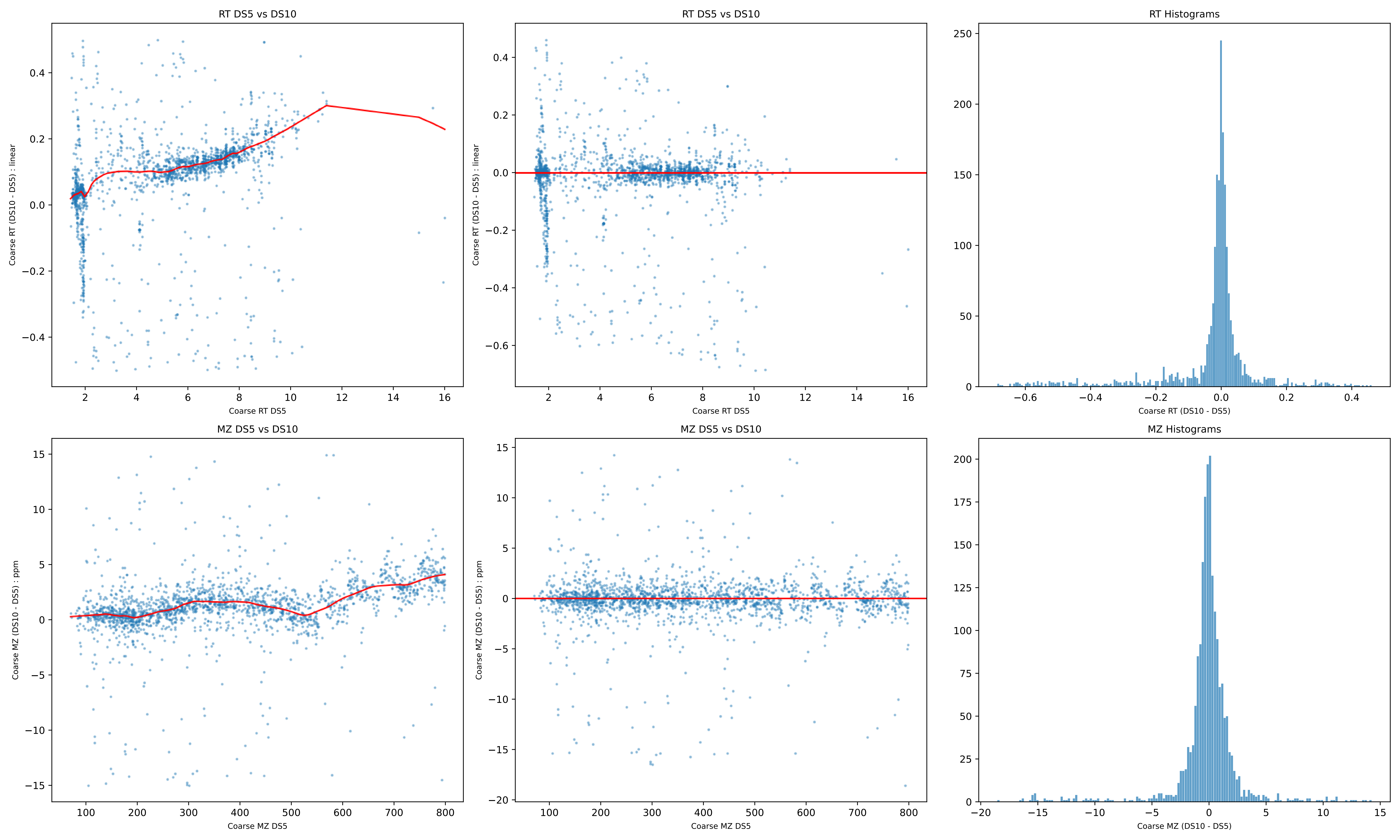

RT DS5 vs DS10
Coarse RT (DS10 - DS5) : linear
Coarse RT DS5
RT DS5 vs DS10
Coarse RT (DS10 - DS5) : linear
Coarse RT DS5
RT Histograms
Coarse RT (DS10 - DS5)
MZ DS5 vs DS10
Coarse MZ (DS10 - DS5) : ppm
Coarse MZ DS5
MZ DS5 vs DS10
Coarse MZ (DS10 - DS5) : ppm
Coarse MZ DS5
MZ Histograms
Coarse MZ (DS10 - DS5)

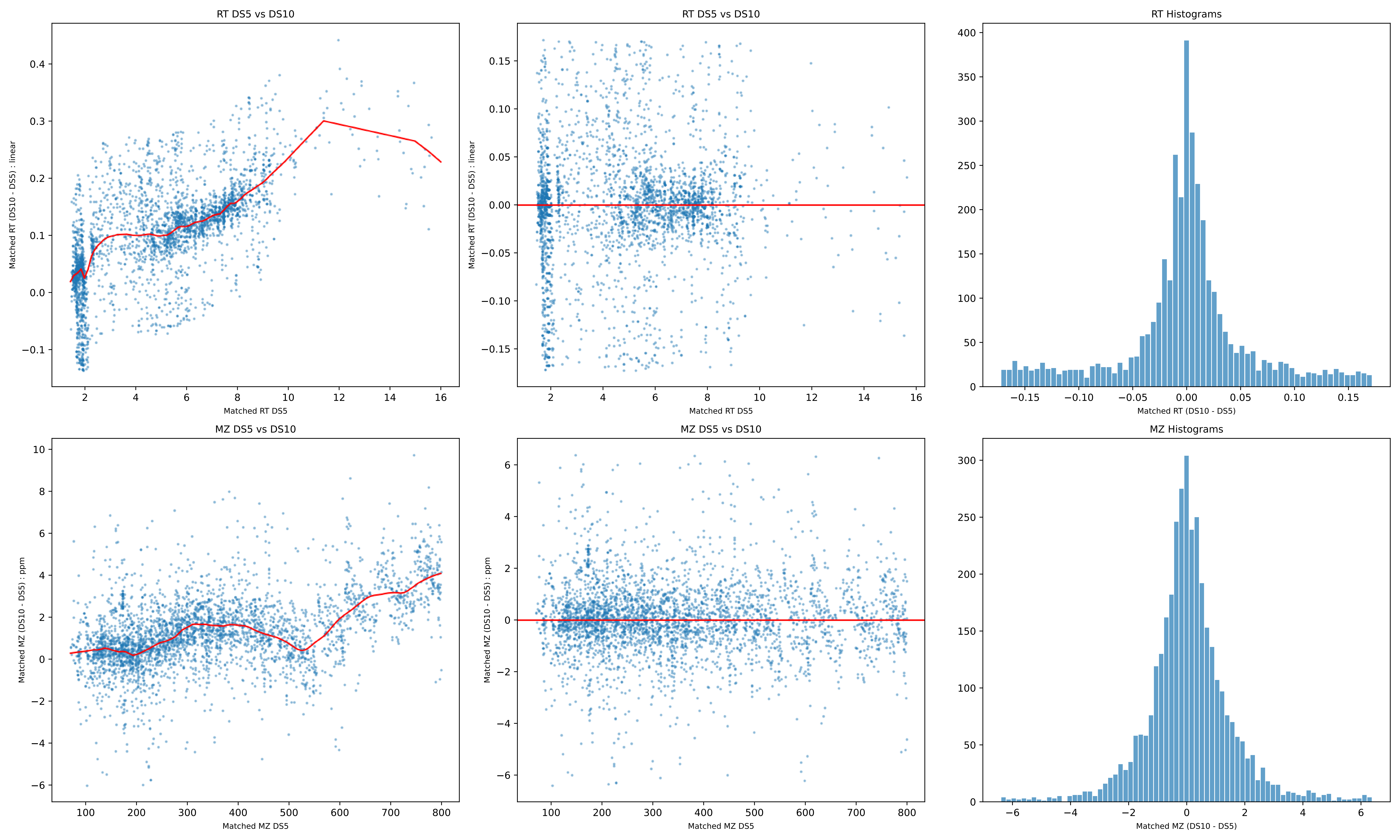

RT DS5 vs DS10
Matched RT (DS10 - DS5) : linear
Matched RT DS5
RT DS5 vs DS10
Matched RT (DS10 - DS5) : linear
Matched RT DS5
RT Histograms
Matched RT (DS10 - DS5)
MZ DS5 vs DS10
Matched MZ (DS10 - DS5) : ppm
Matched MZ DS5
MZ DS5 vs DS10
Matched MZ (DS10 - DS5) : ppm
Matched MZ DS5
MZ Histograms
Matched MZ (DS10 - DS5)

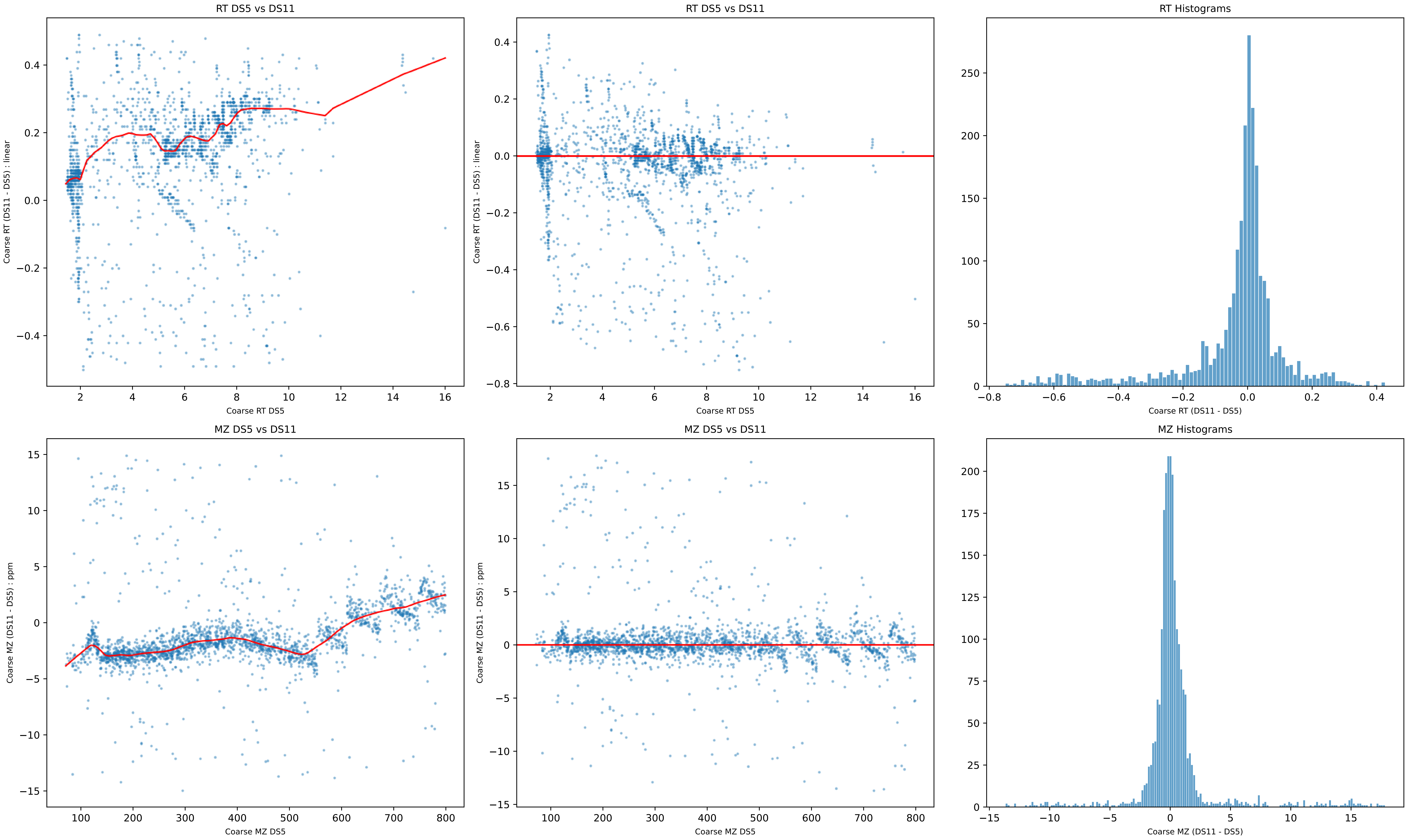
RT DS5 vs DS11
Coarse RT (DS11 - DS5) : linear
Coarse RT DS5
RT DS5 vs DS11
Coarse RT (DS11 - DS5) : linear
Coarse RT DS5
RT Histograms
Coarse RT (DS11 - DS5)
MZ DS5 vs DS11
Coarse MZ (DS11 - DS5) : ppm
Coarse MZ DS5
MZ DS5 vs DS11
Coarse MZ (DS11 - DS5) : ppm
Coarse MZ DS5
MZ Histograms
Coarse MZ (DS11 - DS5)

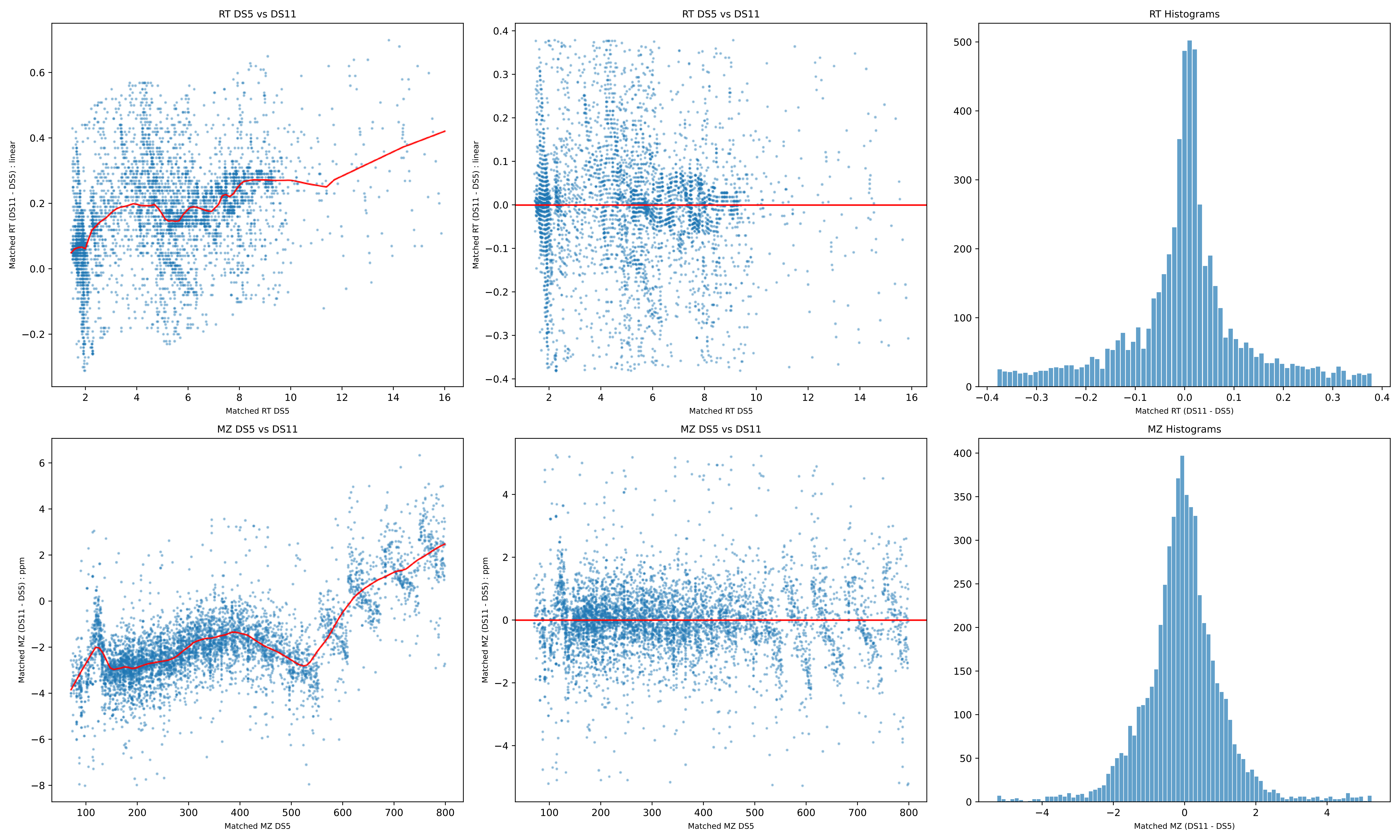

RT DS5 vs DS11
Matched RT (DS11 - DS5) : linear
Matched RT DS5
RT DS5 vs DS11
Matched RT (DS11 - DS5) : linear
Matched RT DS5
RT Histograms
Matched RT (DS11 - DS5)
MZ DS5 vs DS11
Matched MZ (DS11 - DS5) : ppm
Matched MZ DS5
MZ DS5 vs DS11
Matched MZ (DS11 - DS5) : ppm
Matched MZ DS5
MZ Histograms
Matched MZ (DS11 - DS5)

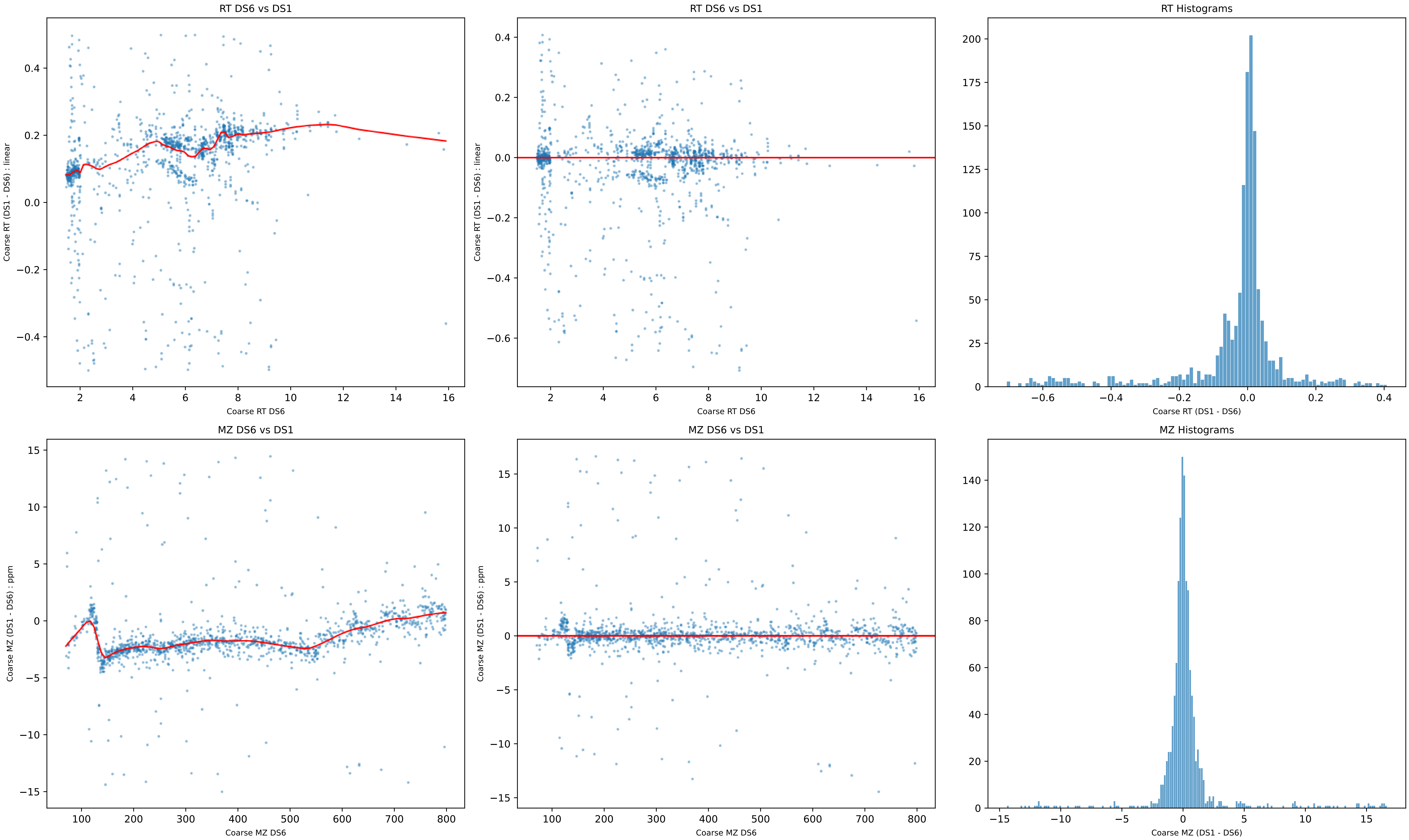
RT DS6 vs DS1
Coarse RT (DS1 - DS6) : linear
Coarse RT DS6
RT DS6 vs DS1
Coarse RT (DS1 - DS6) : linear
Coarse RT DS6
RT Histograms
Coarse RT (DS1 - DS6)
MZ DS6 vs DS1
Coarse MZ (DS1 - DS6) : ppm
Coarse MZ DS6
MZ DS6 vs DS1
Coarse MZ (DS1 - DS6) : ppm
Coarse MZ DS6
MZ Histograms
Coarse MZ (DS1 - DS6)

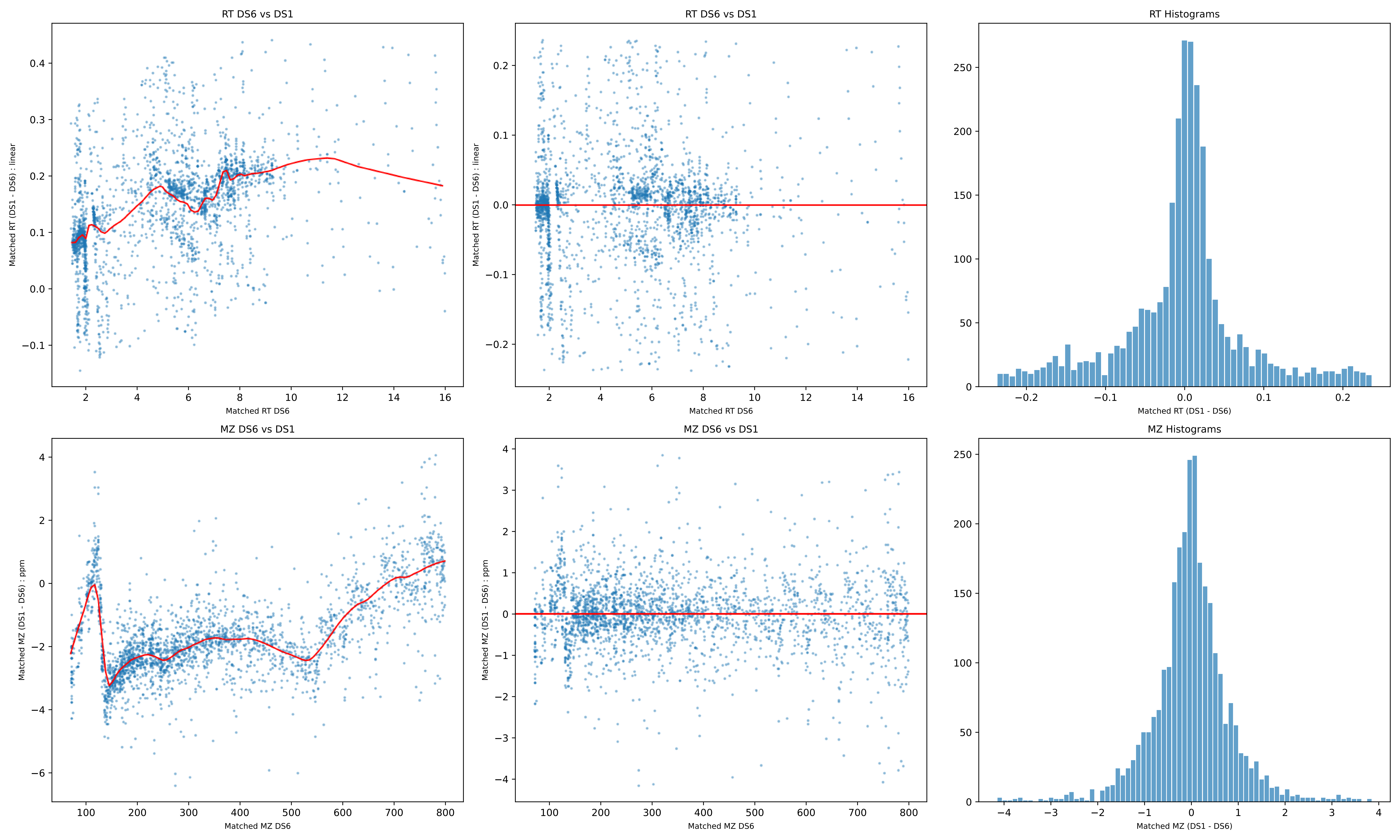

RT DS6 vs DS1
Matched RT (DS1 - DS6) : linear
Matched RT DS6
RT DS6 vs DS1
Matched RT (DS1 - DS6) : linear
Matched RT DS6
RT Histograms
Matched RT (DS1 - DS6)
MZ DS6 vs DS1
Matched MZ (DS1 - DS6) : ppm
Matched MZ DS6
MZ DS6 vs DS1
Matched MZ (DS1 - DS6) : ppm
Matched MZ DS6
MZ Histograms
Matched MZ (DS1 - DS6)

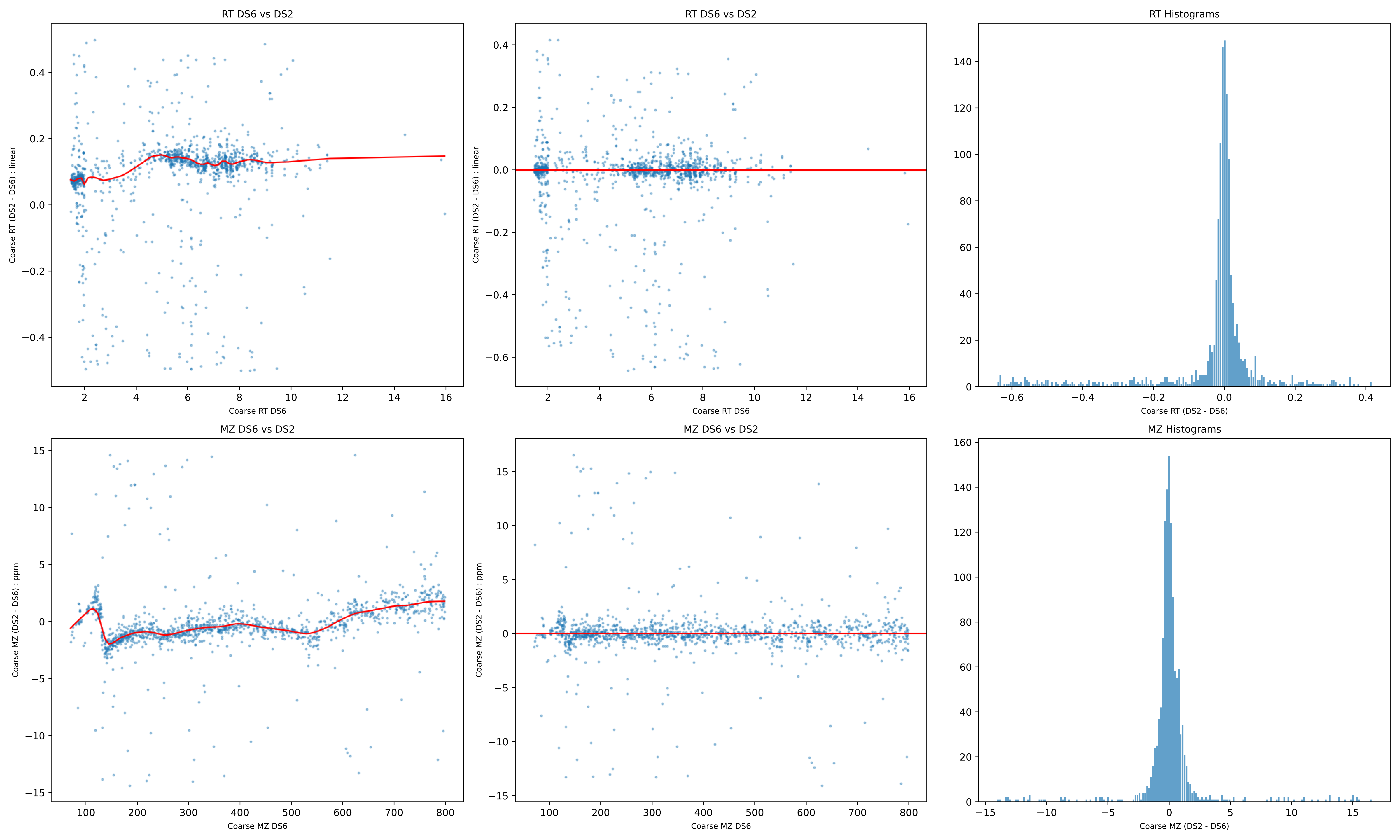
RT DS6 vs DS2
Coarse RT (DS2 - DS6) : linear
Coarse RT DS6
RT DS6 vs DS2
Coarse RT (DS2 - DS6) : linear
Coarse RT DS6
RT Histograms
Coarse RT (DS2 - DS6)
MZ DS6 vs DS2
Coarse MZ (DS2 - DS6) : ppm
Coarse MZ DS6
MZ DS6 vs DS2
Coarse MZ (DS2 - DS6) : ppm
Coarse MZ DS6
MZ Histograms
Coarse MZ (DS2 - DS6)

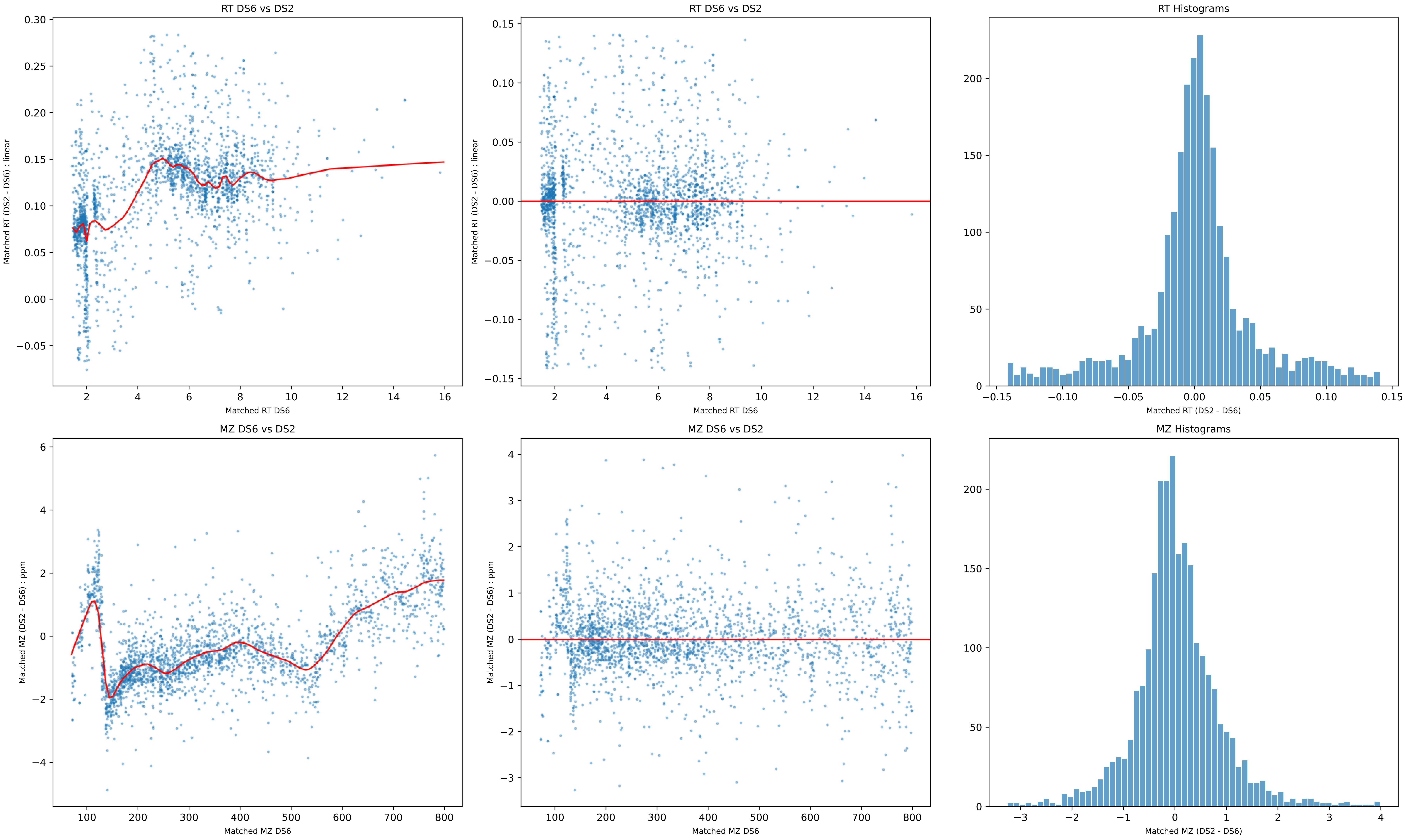
RT DS6 vs DS2
Matched RT (DS2 - DS6) : linear
Matched RT DS6
RT DS6 vs DS2
Matched RT (DS2 - DS6) : linear
Matched RT DS6
RT Histograms
Matched RT (DS2 - DS6)
MZ DS6 vs DS2
Matched MZ (DS2 - DS6) : ppm
Matched MZ DS6
MZ DS6 vs DS2
Matched MZ (DS2 - DS6) : ppm
Matched MZ DS6
MZ Histograms
Matched MZ (DS2 - DS6)

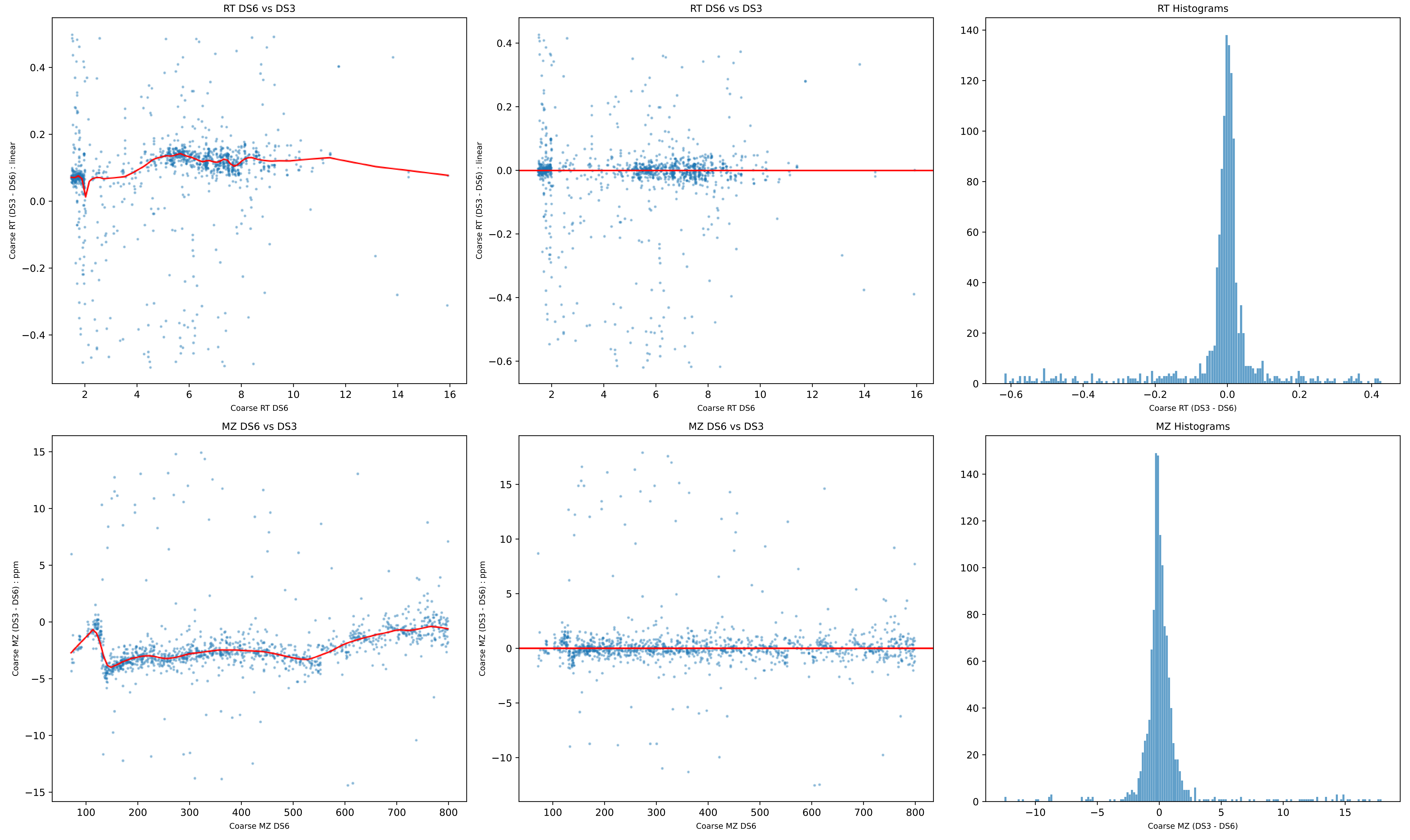

RT DS6 vs DS3
Coarse RT (DS3 - DS6) : linear
Coarse RT DS6
RT DS6 vs DS3
Coarse RT (DS3 - DS6) : linear
Coarse RT DS6
RT Histograms
Coarse RT (DS3 - DS6)
MZ DS6 vs DS3
Coarse MZ (DS3 - DS6) : ppm
Coarse MZ DS6
MZ DS6 vs DS3
Coarse MZ (DS3 - DS6) : ppm
Coarse MZ DS6
MZ Histograms
Coarse MZ (DS3 - DS6)

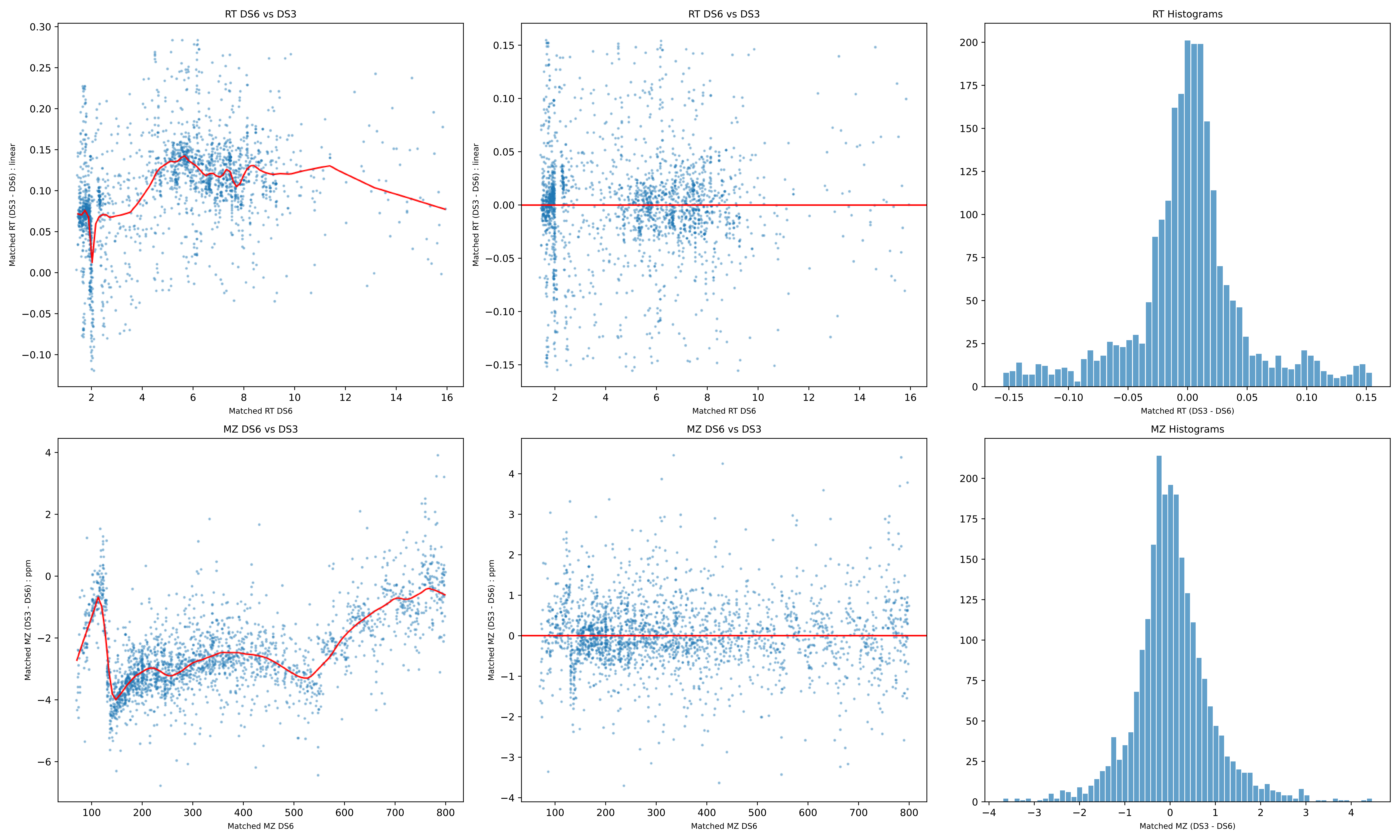

RT DS6 vs DS3
Matched RT (DS3 - DS6) : linear
Matched RT DS6
RT DS6 vs DS3
Matched RT (DS3 - DS6) : linear
Matched RT DS6
RT Histograms
Matched RT (DS3 - DS6)
MZ DS6 vs DS3
Matched MZ (DS3 - DS6) : ppm
Matched MZ DS6
MZ DS6 vs DS3
Matched MZ (DS3 - DS6) : ppm
Matched MZ DS6
MZ Histograms
Matched MZ (DS3 - DS6)

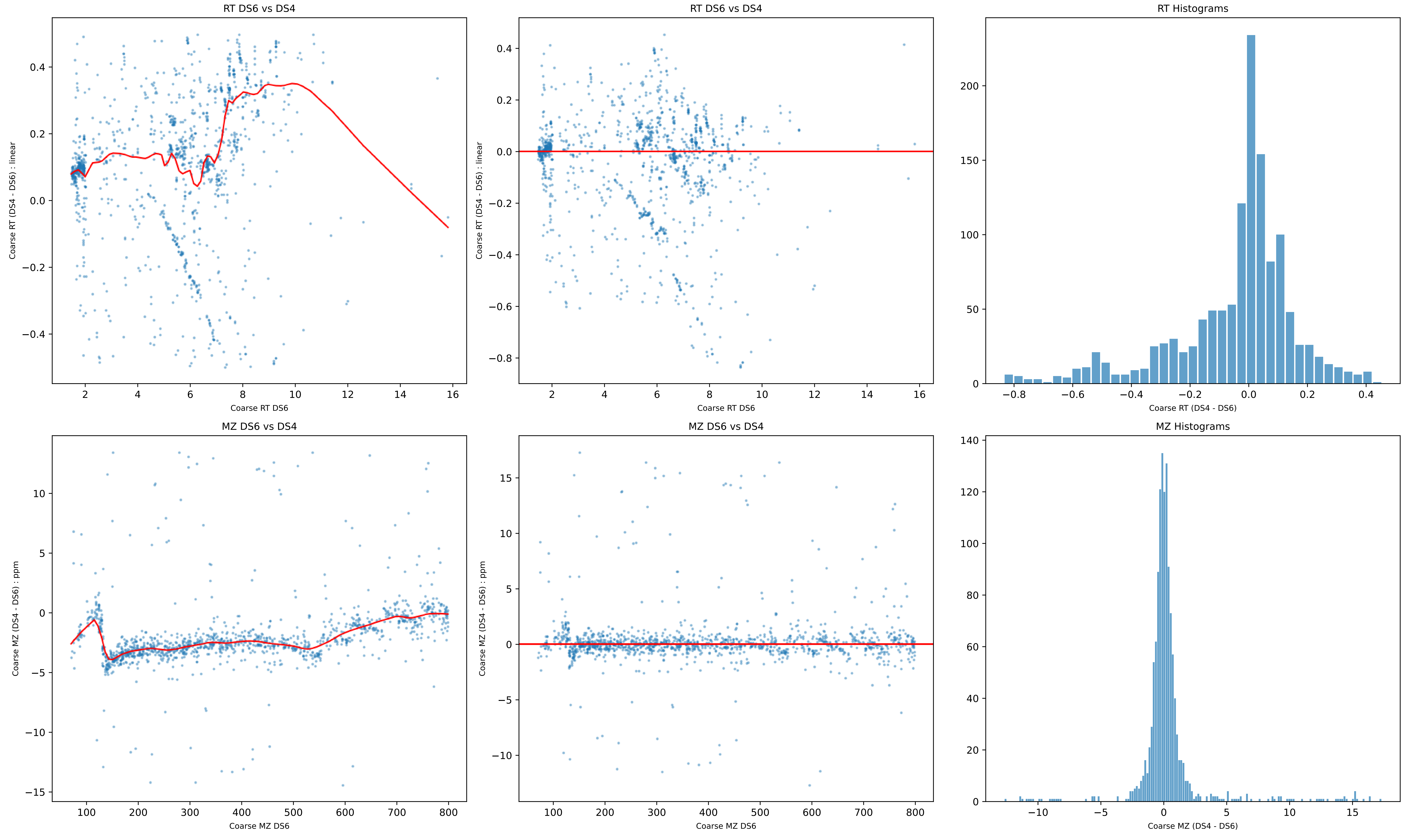
RT DS6 vs DS4
Coarse RT (DS4 - DS6) : linear
Coarse RT DS6
RT DS6 vs DS4
Coarse RT (DS4 - DS6) : linear
Coarse RT DS6
RT Histograms
Coarse RT (DS4 - DS6)
MZ DS6 vs DS4
Coarse MZ (DS4 - DS6) : ppm
Coarse MZ DS6
MZ DS6 vs DS4
Coarse MZ (DS4 - DS6) : ppm
Coarse MZ DS6
MZ Histograms
Coarse MZ (DS4 - DS6)

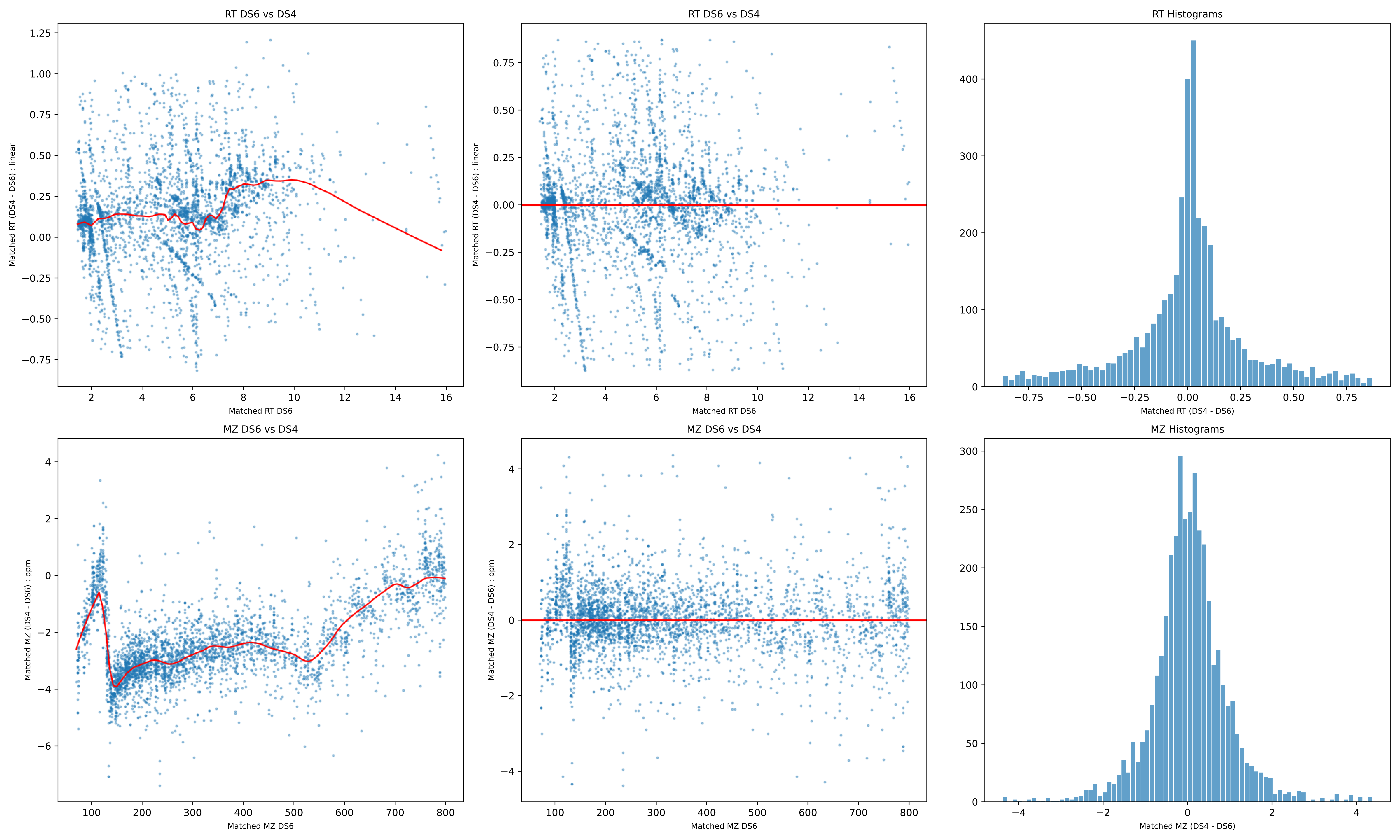

RT DS6 vs DS4
Matched RT (DS4 - DS6) : linear
Matched RT DS6
RT DS6 vs DS4
Matched RT (DS4 - DS6) : linear
Matched RT DS6
RT Histograms
Matched RT (DS4 - DS6)
MZ DS6 vs DS4
Matched MZ (DS4 - DS6) : ppm
Matched MZ DS6
MZ DS6 vs DS4
Matched MZ (DS4 - DS6) : ppm
Matched MZ DS6
MZ Histograms
Matched MZ (DS4 - DS6)

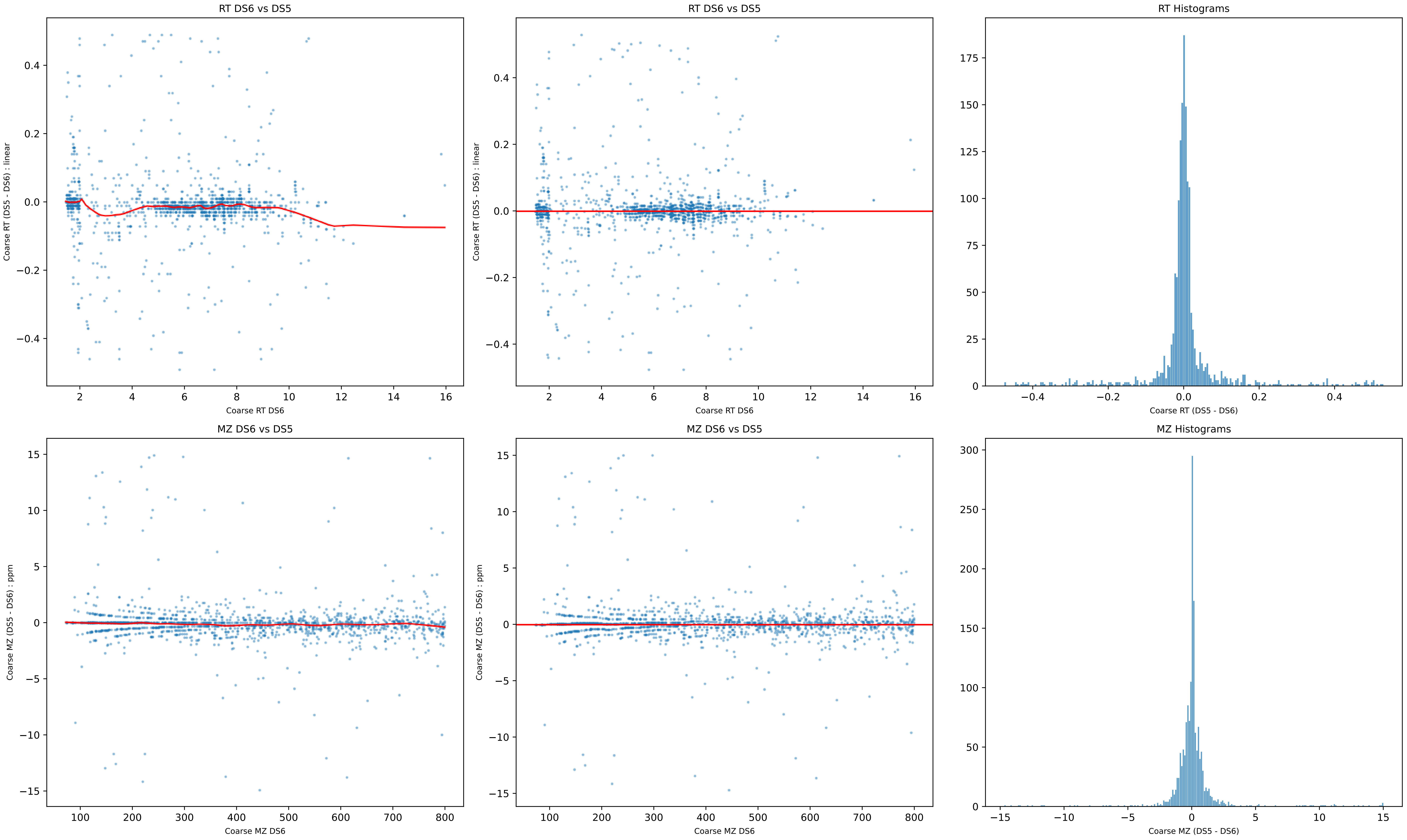

RT DS6 vs DS5
Coarse RT (DS5 - DS6) : linear
Coarse RT DS6
RT DS6 vs DS5
Coarse RT (DS5 - DS6) : linear
Coarse RT DS6
RT Histograms
Coarse RT (DS5 - DS6)
MZ DS6 vs DS5
Coarse MZ (DS5 - DS6) : ppm
Coarse MZ DS6
MZ DS6 vs DS5
Coarse MZ (DS5 - DS6) : ppm
Coarse MZ DS6
MZ Histograms
Coarse MZ (DS5 - DS6)

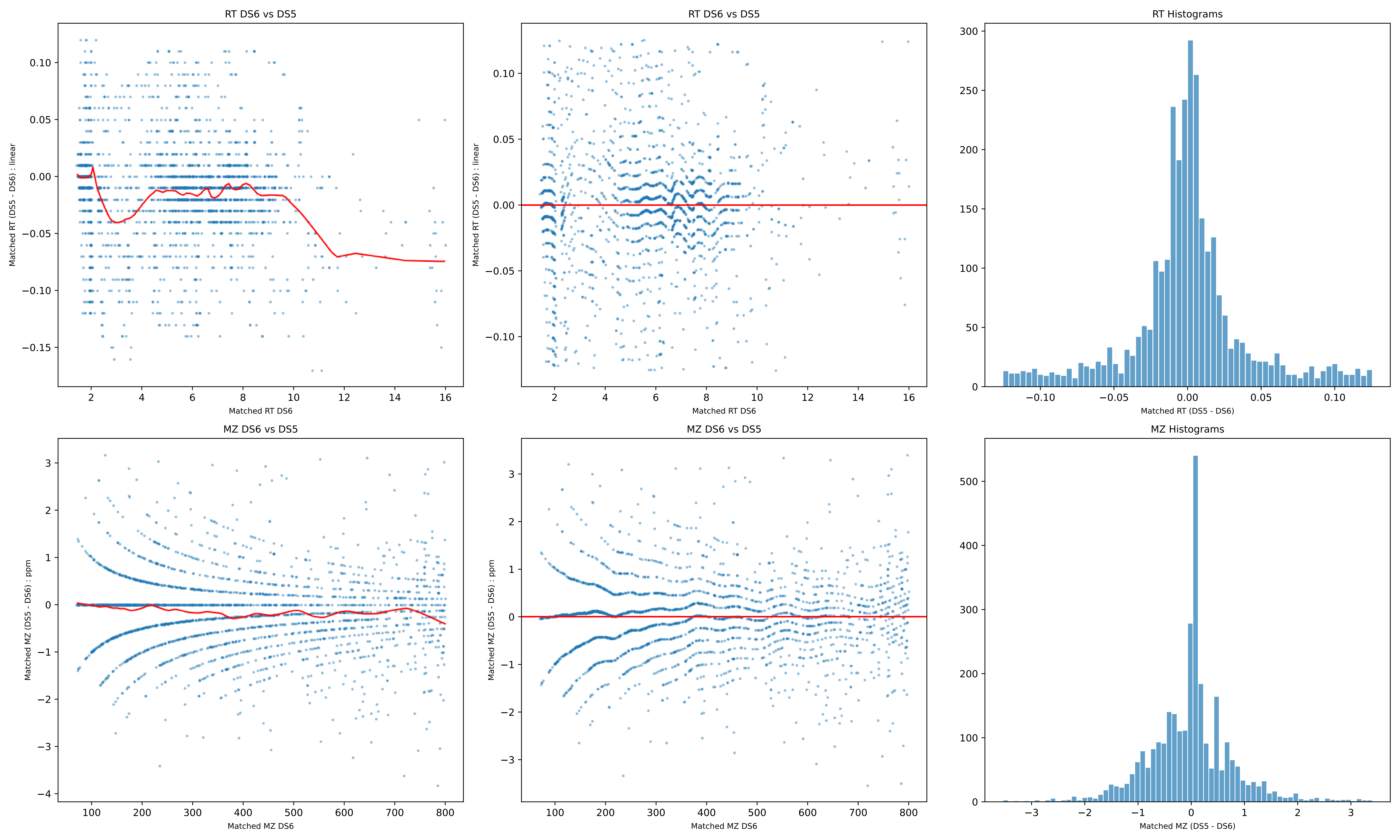

RT DS6 vs DS5
Matched RT (DS5 - DS6) : linear
Matched RT DS6
RT DS6 vs DS5
Matched RT (DS5 - DS6) : linear
Matched RT DS6
RT Histograms
Matched RT (DS5 - DS6)
MZ DS6 vs DS5
Matched MZ (DS5 - DS6) : ppm
Matched MZ DS6
MZ DS6 vs DS5
Matched MZ (DS5 - DS6) : ppm
Matched MZ DS6
MZ Histograms
Matched MZ (DS5 - DS6)

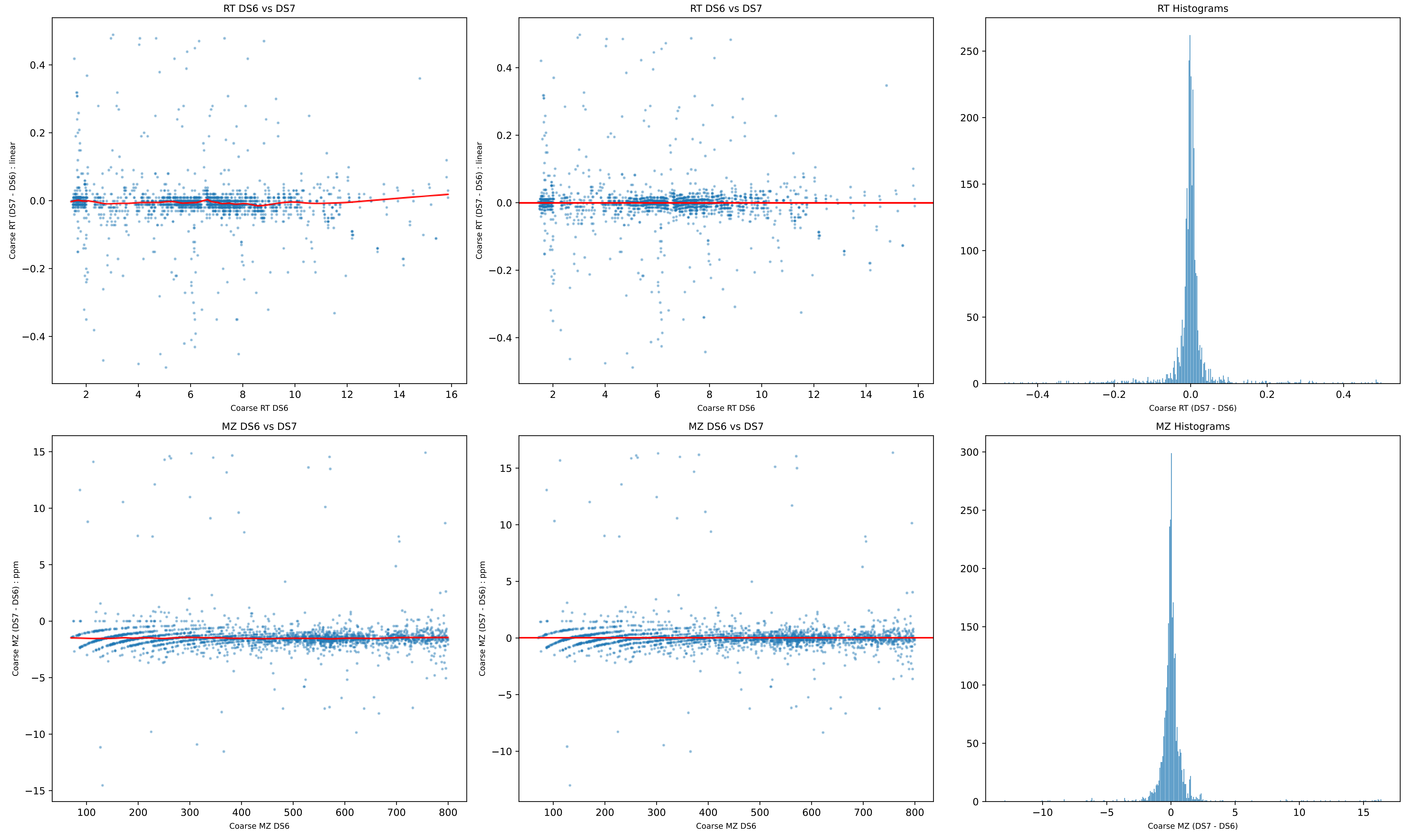
RT DS6 vs DS7
Coarse RT (DS7 - DS6) : linear
Coarse RT DS6
RT DS6 vs DS7
Coarse RT (DS7 - DS6) : linear
Coarse RT DS6
RT Histograms
Coarse RT (DS7 - DS6)
MZ DS6 vs DS7
Coarse MZ (DS7 - DS6) : ppm
Coarse MZ DS6
MZ DS6 vs DS7
Coarse MZ (DS7 - DS6) : ppm
Coarse MZ DS6
MZ Histograms
Coarse MZ (DS7 - DS6)

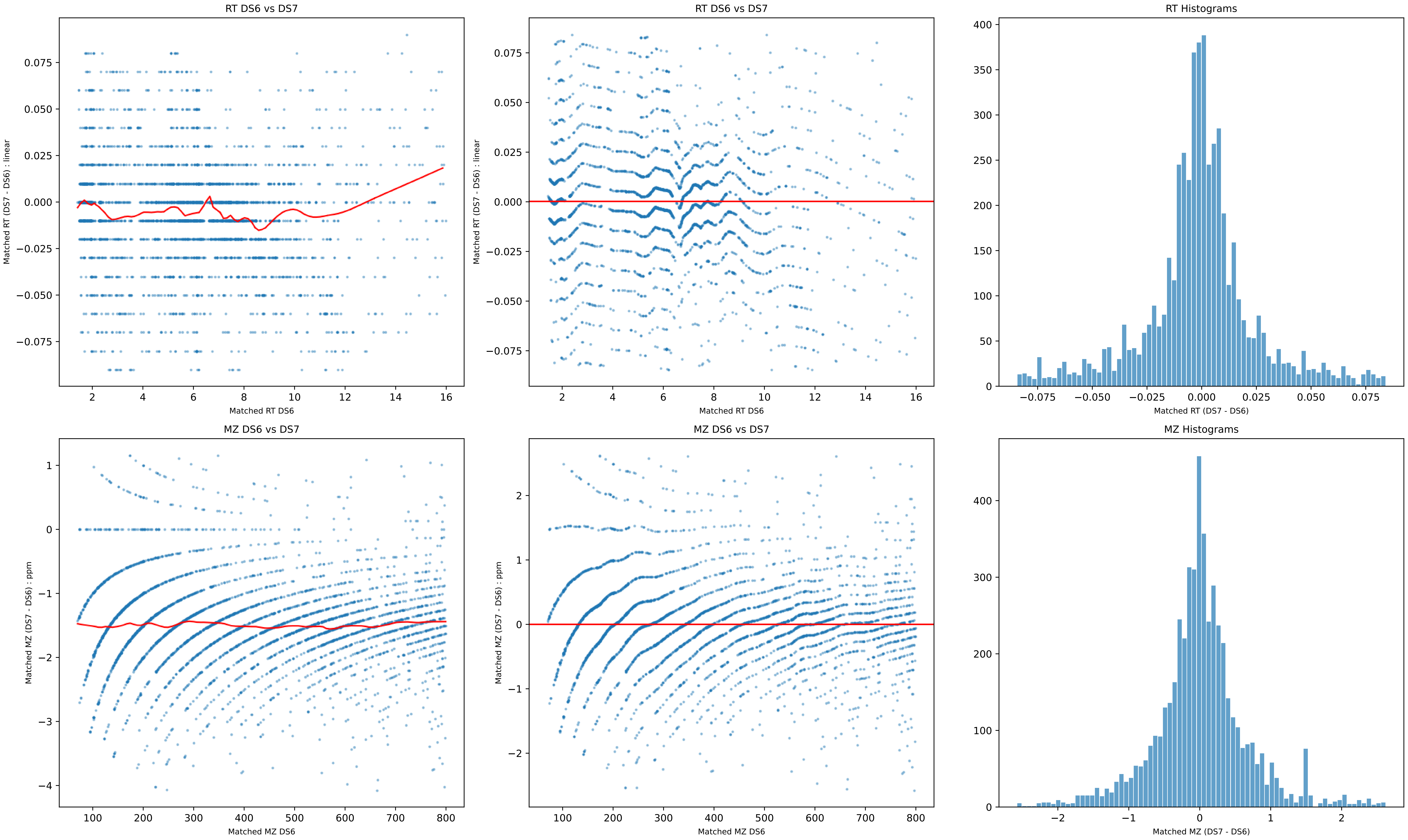
RT DS6 vs DS7
Matched RT (DS7 - DS6) : linear
Matched RT DS6
RT DS6 vs DS7
Matched RT (DS7 - DS6) : linear
Matched RT DS6
RT Histograms
Matched RT (DS7 - DS6)
MZ DS6 vs DS7
Matched MZ (DS7 - DS6) : ppm
Matched MZ DS6
MZ DS6 vs DS7
Matched MZ (DS7 - DS6) : ppm
Matched MZ DS6
MZ Histograms
Matched MZ (DS7 - DS6)

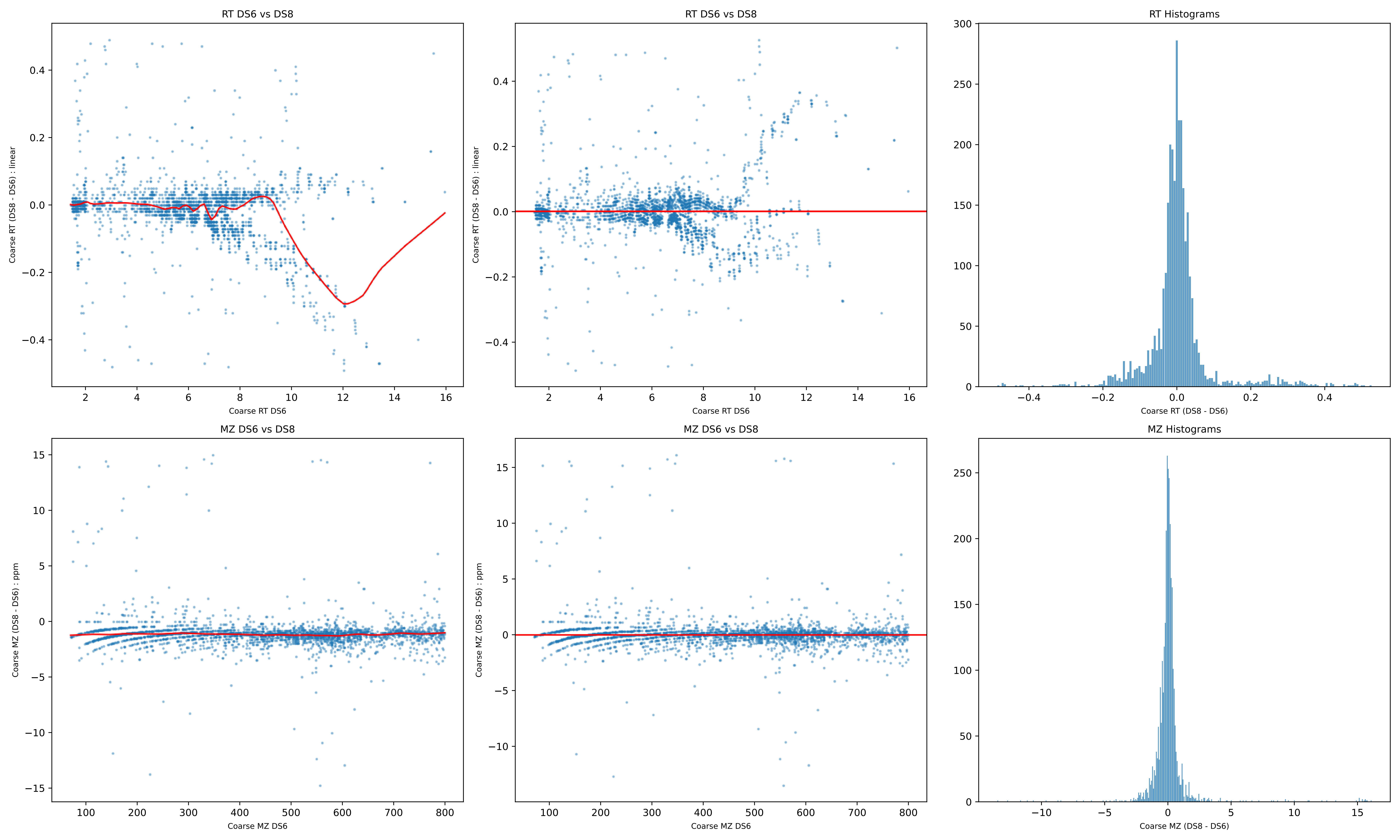
RT DS6 vs DS8
Coarse RT (DS8 - DS6) : linear
Coarse RT DS6
RT DS6 vs DS8
Coarse RT (DS8 - DS6) : linear
Coarse RT DS6
RT Histograms
Coarse RT (DS8 - DS6)
MZ DS6 vs DS8
Coarse MZ (DS8 - DS6) : ppm
Coarse MZ DS6
MZ DS6 vs DS8
Coarse MZ (DS8 - DS6) : ppm
Coarse MZ DS6
MZ Histograms
Coarse MZ (DS8 - DS6)

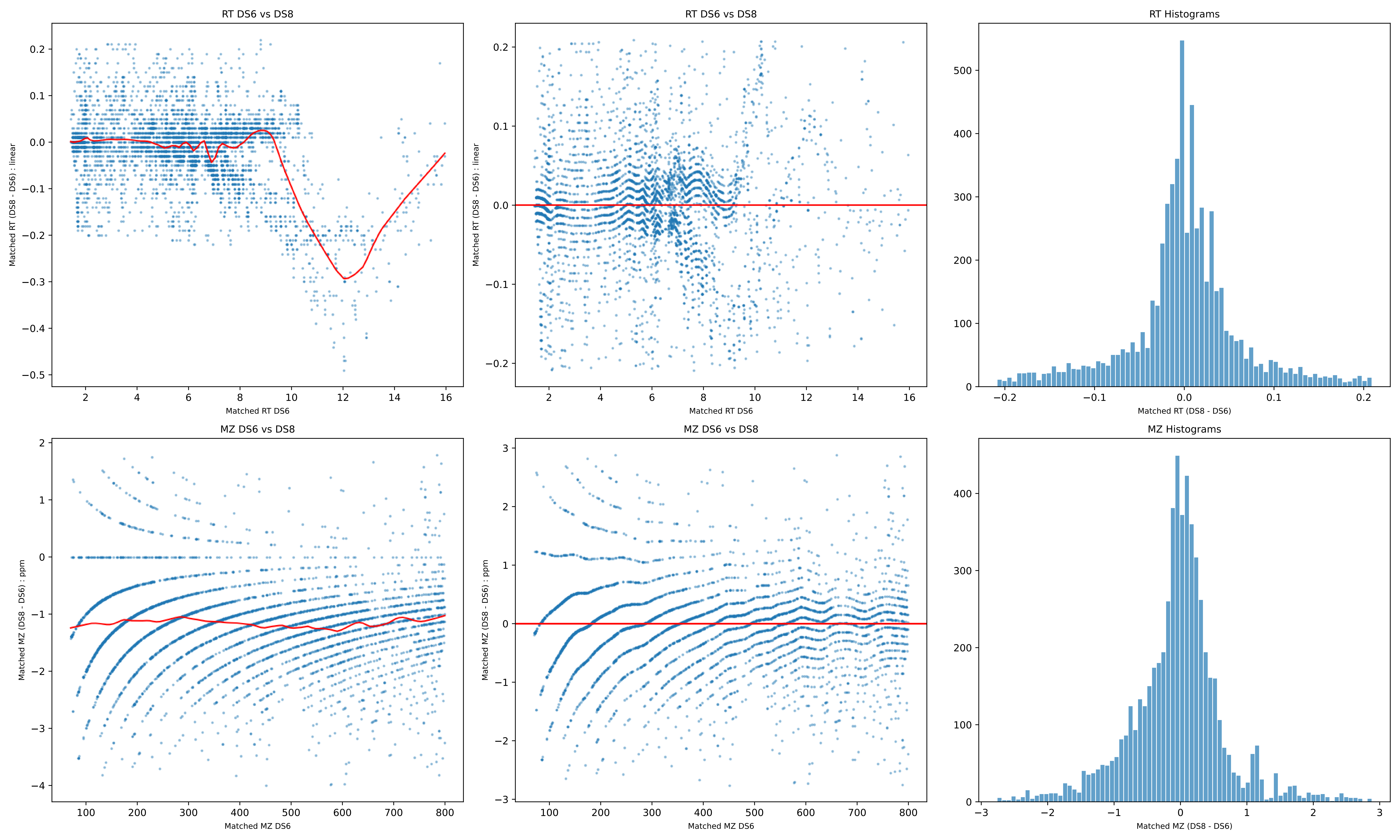
RT DS6 vs DS8
Matched RT (DS8 - DS6) : linear
Matched RT DS6
RT DS6 vs DS8
Matched RT (DS8 - DS6) : linear
Matched RT DS6
RT Histograms
Matched RT (DS8 - DS6)
MZ DS6 vs DS8
Matched MZ (DS8 - DS6) : ppm
Matched MZ DS6
MZ DS6 vs DS8
Matched MZ (DS8 - DS6) : ppm
Matched MZ DS6
MZ Histograms
Matched MZ (DS8 - DS6)

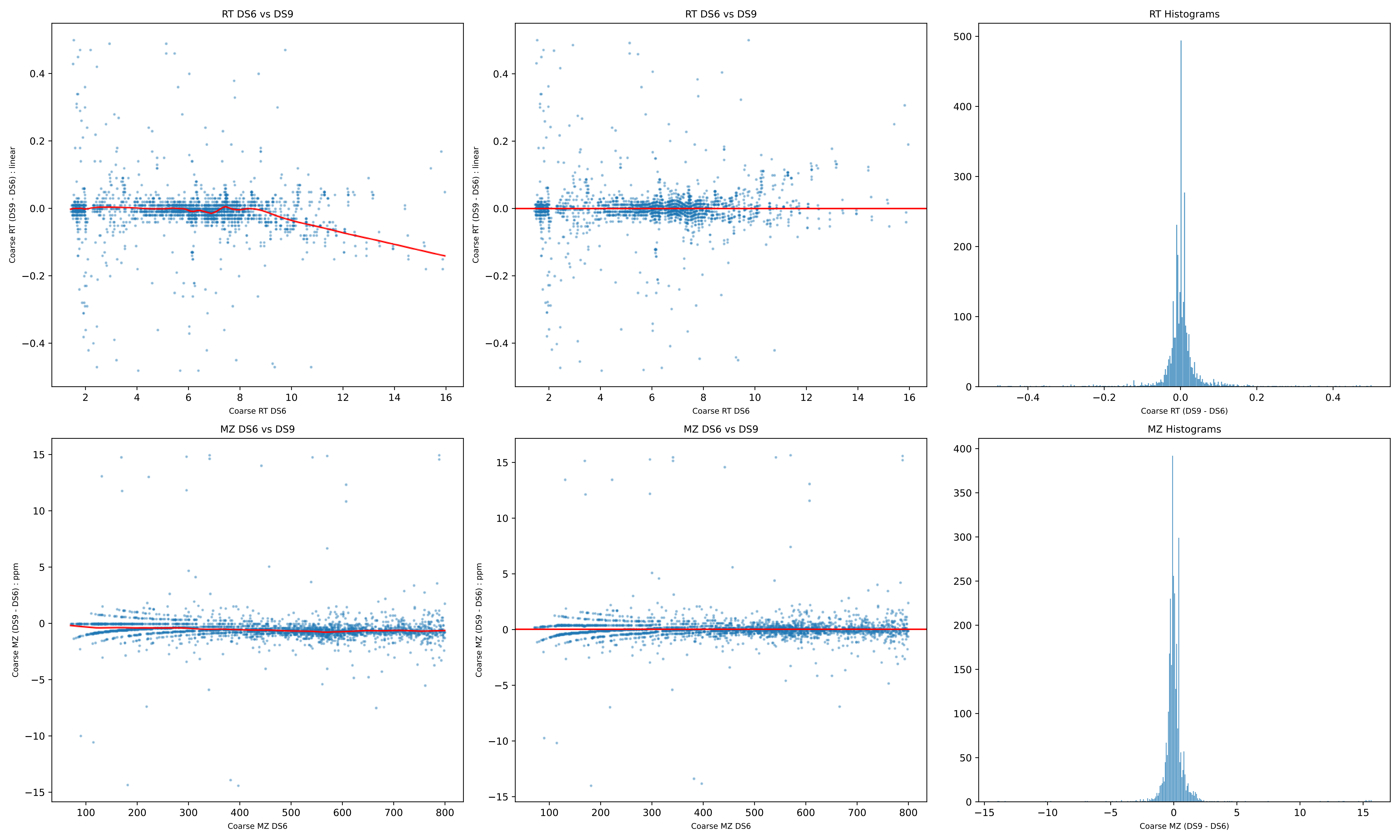
RT DS6 vs DS9
Coarse RT (DS9 - DS6) : linear
Coarse RT DS6
RT DS6 vs DS9
Coarse RT (DS9 - DS6) : linear
Coarse RT DS6
RT Histograms
Coarse RT (DS9 - DS6)
MZ DS6 vs DS9
Coarse MZ (DS9 - DS6) : ppm
Coarse MZ DS6
MZ DS6 vs DS9
Coarse MZ (DS9 - DS6) : ppm
Coarse MZ DS6
MZ Histograms
Coarse MZ (DS9 - DS6)

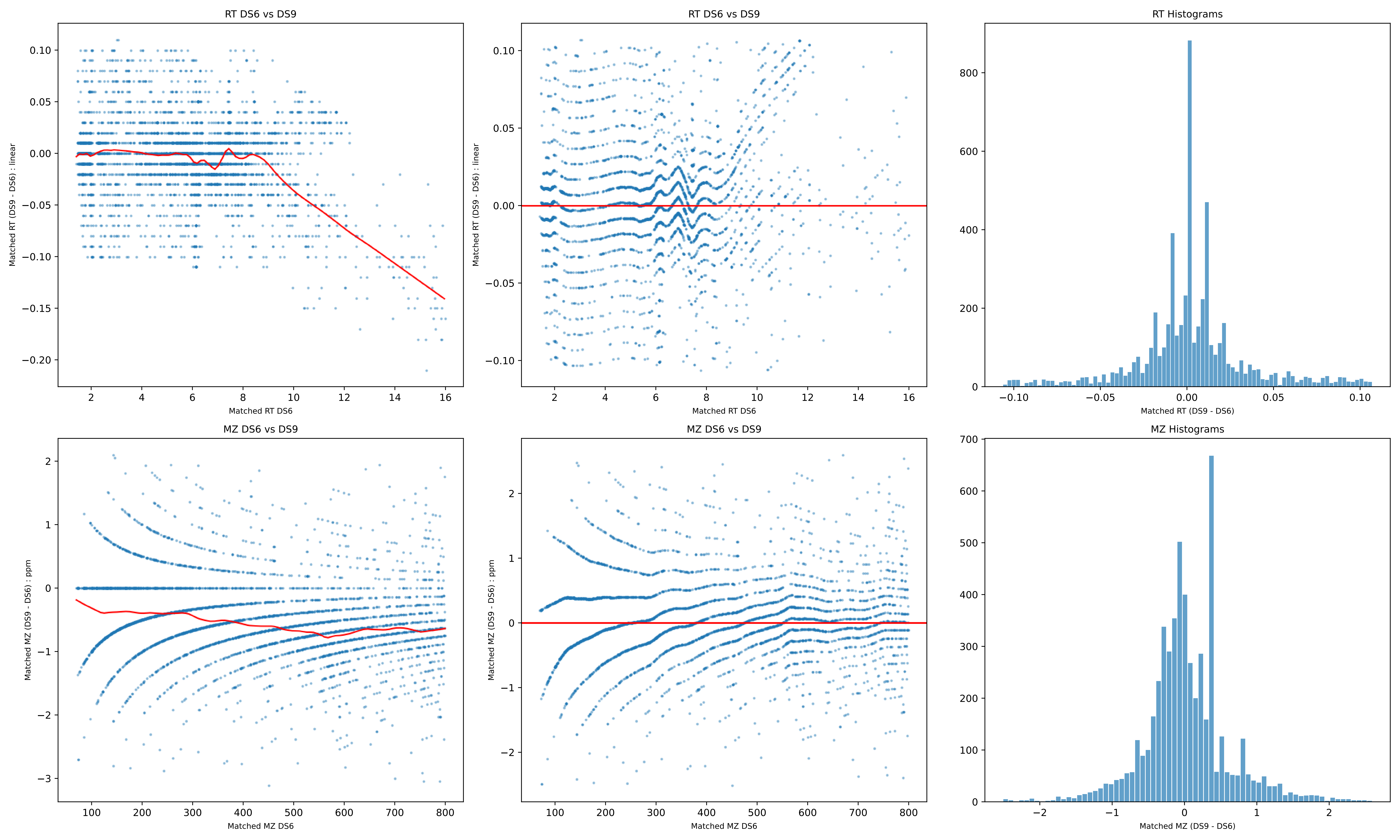
RT DS6 vs DS9
Matched RT (DS9 - DS6) : linear
Matched RT DS6
RT DS6 vs DS9
Matched RT (DS9 - DS6) : linear
Matched RT DS6
RT Histograms
Matched RT (DS9 - DS6)
MZ DS6 vs DS9
Matched MZ (DS9 - DS6) : ppm
Matched MZ DS6
MZ DS6 vs DS9
Matched MZ (DS9 - DS6) : ppm
Matched MZ DS6
MZ Histograms
Matched MZ (DS9 - DS6)

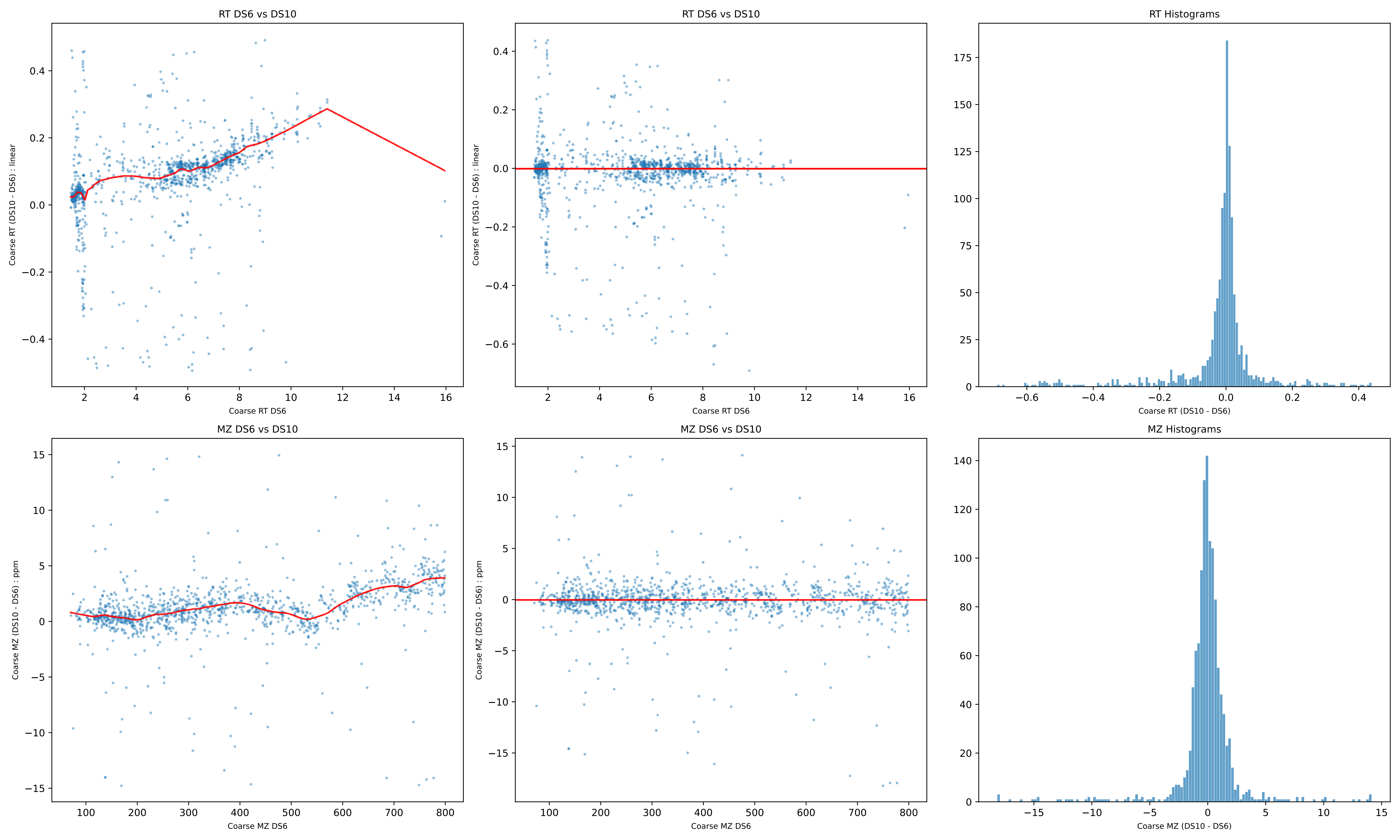
RT DS6 vs DS10
Coarse RT (DS10 - DS6) : linear
Coarse RT DS6
RT DS6 vs DS10
Coarse RT (DS10 - DS6) : linear
Coarse RT DS6
RT Histograms
Coarse RT (DS10 - DS6)
MZ DS6 vs DS10
Coarse MZ (DS10 - DS6) : ppm
Coarse MZ DS6
MZ DS6 vs DS10
Coarse MZ (DS10 - DS6) : ppm
Coarse MZ DS6
MZ Histograms
Coarse MZ (DS10 - DS6)

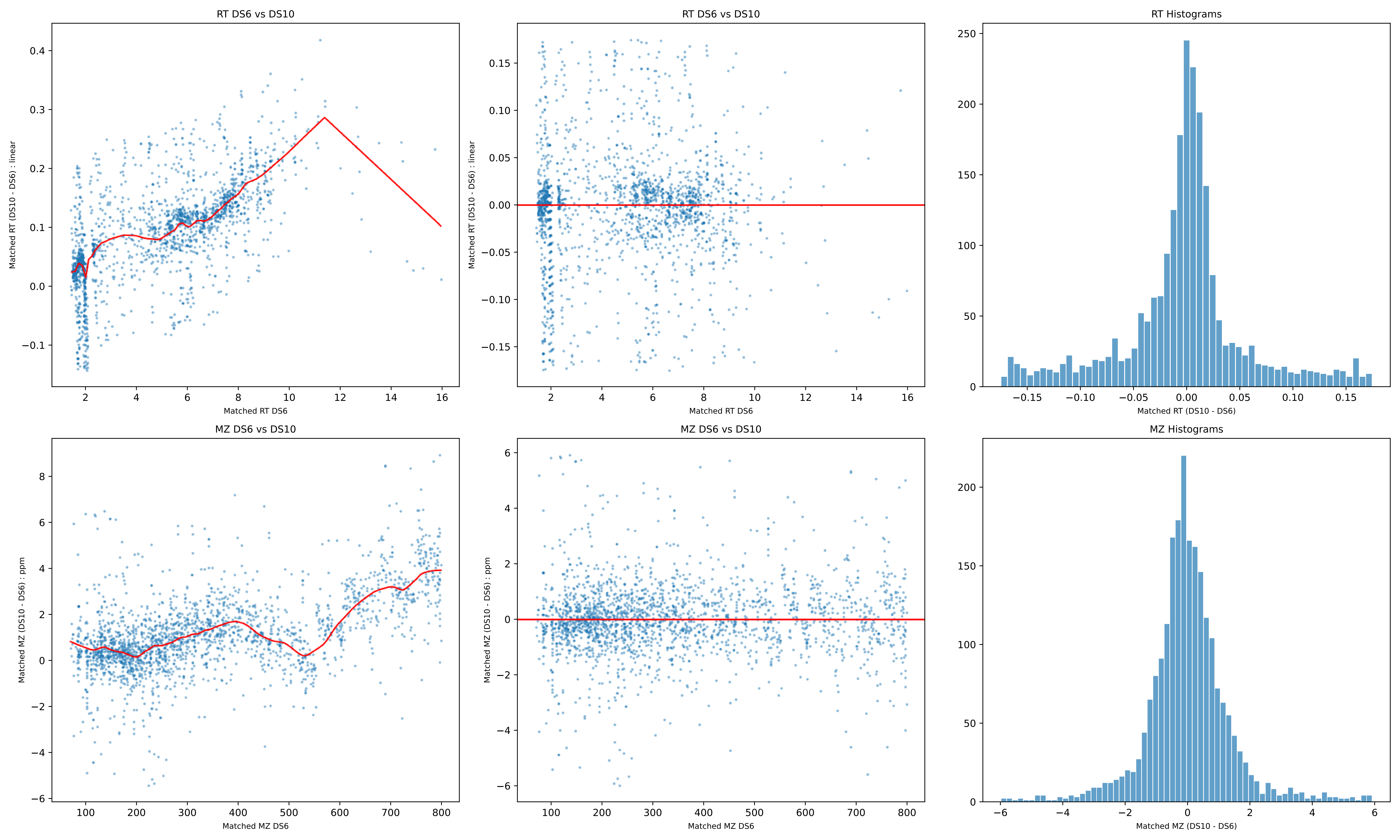

RT DS6 vs DS10
Matched RT (DS10 - DS6) : linear
Matched RT DS6
RT DS6 vs DS10
Matched RT (DS10 - DS6) : linear
Matched RT DS6
RT Histograms
Matched RT (DS10 - DS6)
MZ DS6 vs DS10
Matched MZ (DS10 - DS6) : ppm
Matched MZ DS6
MZ DS6 vs DS10
Matched MZ (DS10 - DS6) : ppm
Matched MZ DS6
MZ Histograms
Matched MZ (DS10 - DS6)

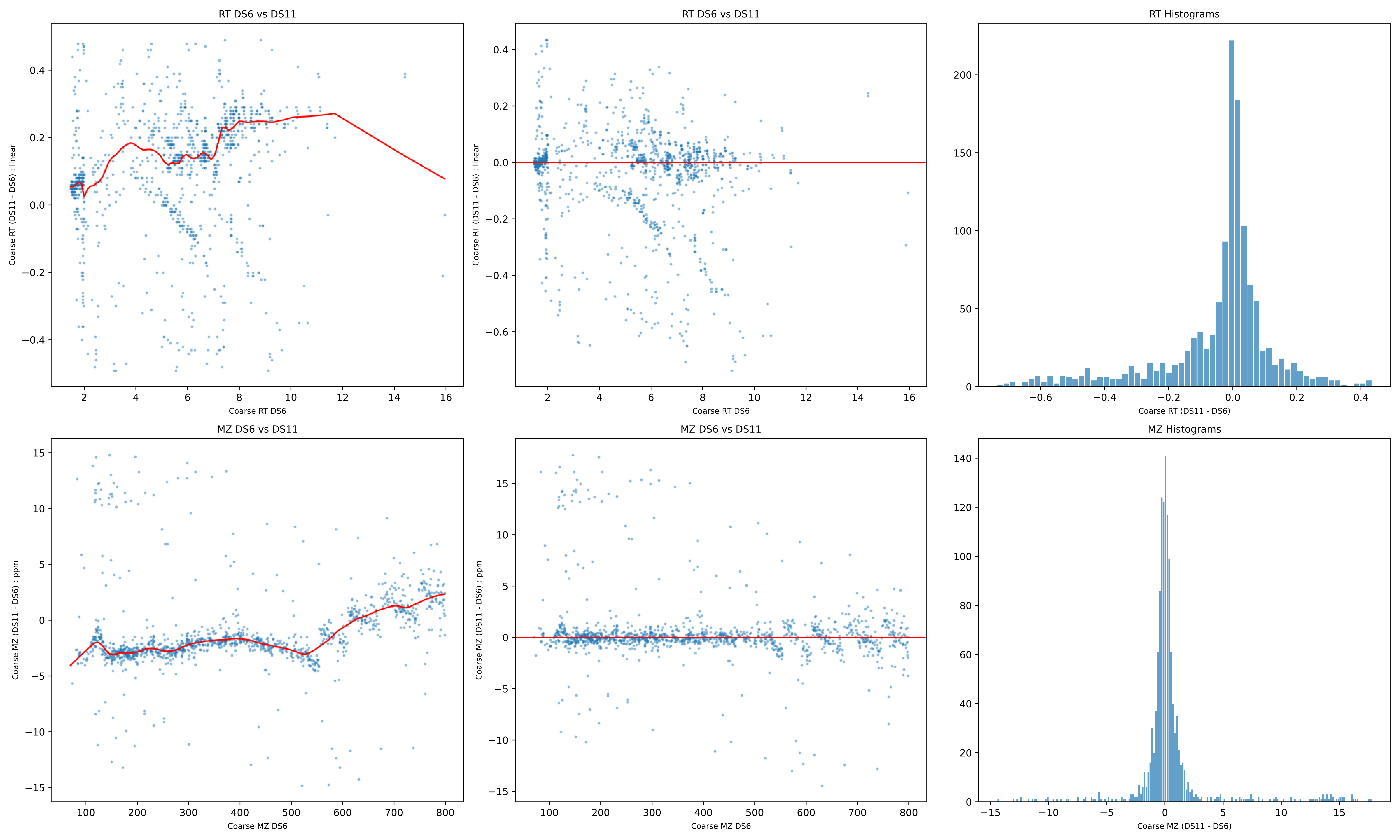

RT DS6 vs DS11
Coarse RT (DS11 - DS6) : linear
Coarse RT DS6
RT DS6 vs DS11
Coarse RT (DS11 - DS6) : linear
Coarse RT DS6
RT Histograms
Coarse RT (DS11 - DS6)
MZ DS6 vs DS11
Coarse MZ (DS11 - DS6) : ppm
Coarse MZ DS6
MZ DS6 vs DS11
Coarse MZ (DS11 - DS6) : ppm
Coarse MZ DS6
MZ Histograms
Coarse MZ (DS11 - DS6)

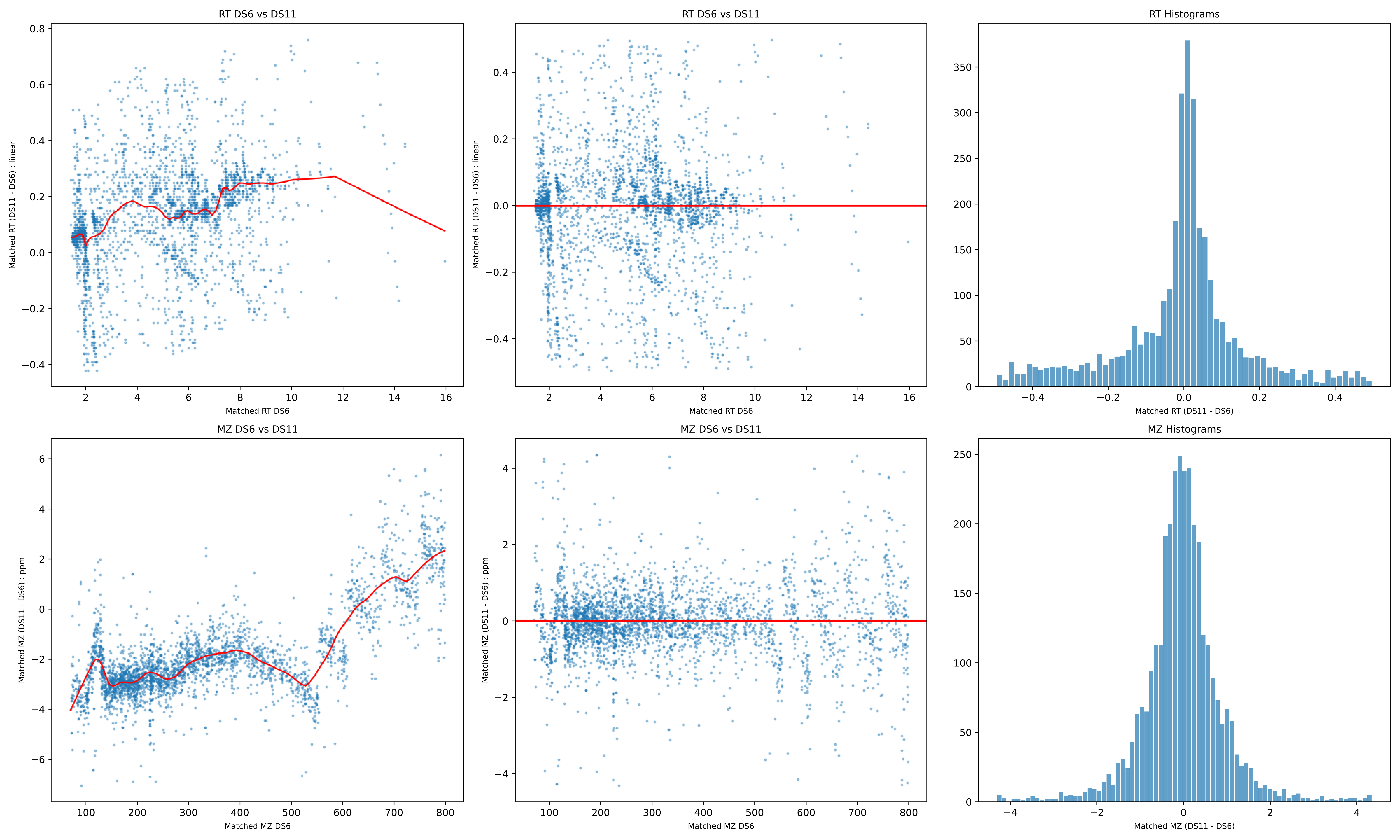

RT DS6 vs DS11
Matched RT (DS11 - DS6) : linear
Matched RT DS6
RT DS6 vs DS11
Matched RT (DS11 - DS6) : linear
Matched RT DS6
RT Histograms
Matched RT (DS11 - DS6)
MZ DS6 vs DS11
Matched MZ (DS11 - DS6) : ppm
Matched MZ DS6
MZ DS6 vs DS11
Matched MZ (DS11 - DS6) : ppm
Matched MZ DS6
MZ Histograms
Matched MZ (DS11 - DS6)

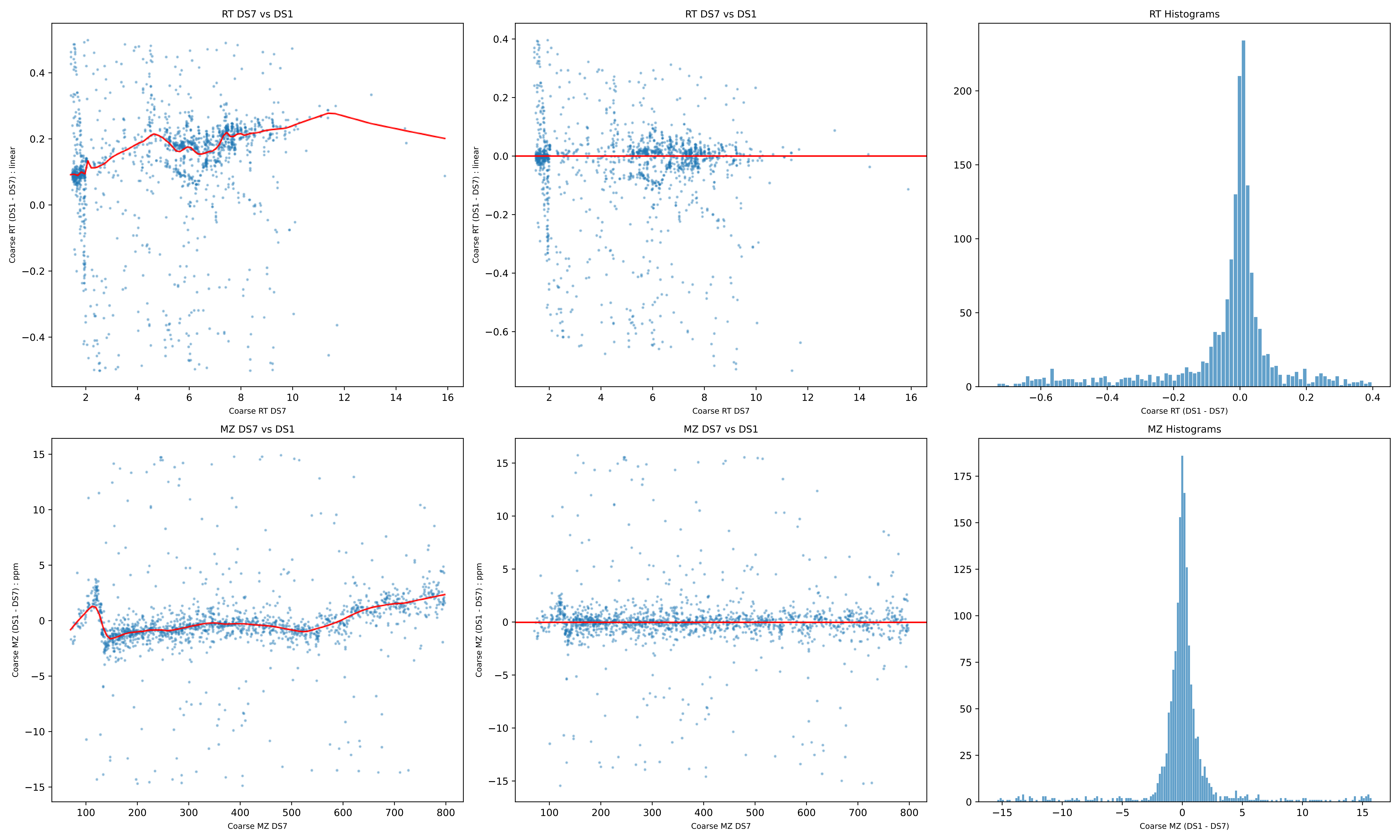
RT DS7 vs DS1
Coarse RT (DS1 - DS7) : linear
Coarse RT DS7
RT DS7 vs DS1
Coarse RT (DS1 - DS7) : linear
Coarse RT DS7
RT Histograms
Coarse RT (DS1 - DS7)
MZ DS7 vs DS1
Coarse MZ (DS1 - DS7) : ppm
Coarse MZ DS7
MZ DS7 vs DS1
Coarse MZ (DS1 - DS7) : ppm
Coarse MZ DS7
MZ Histograms
Coarse MZ (DS1 - DS7)

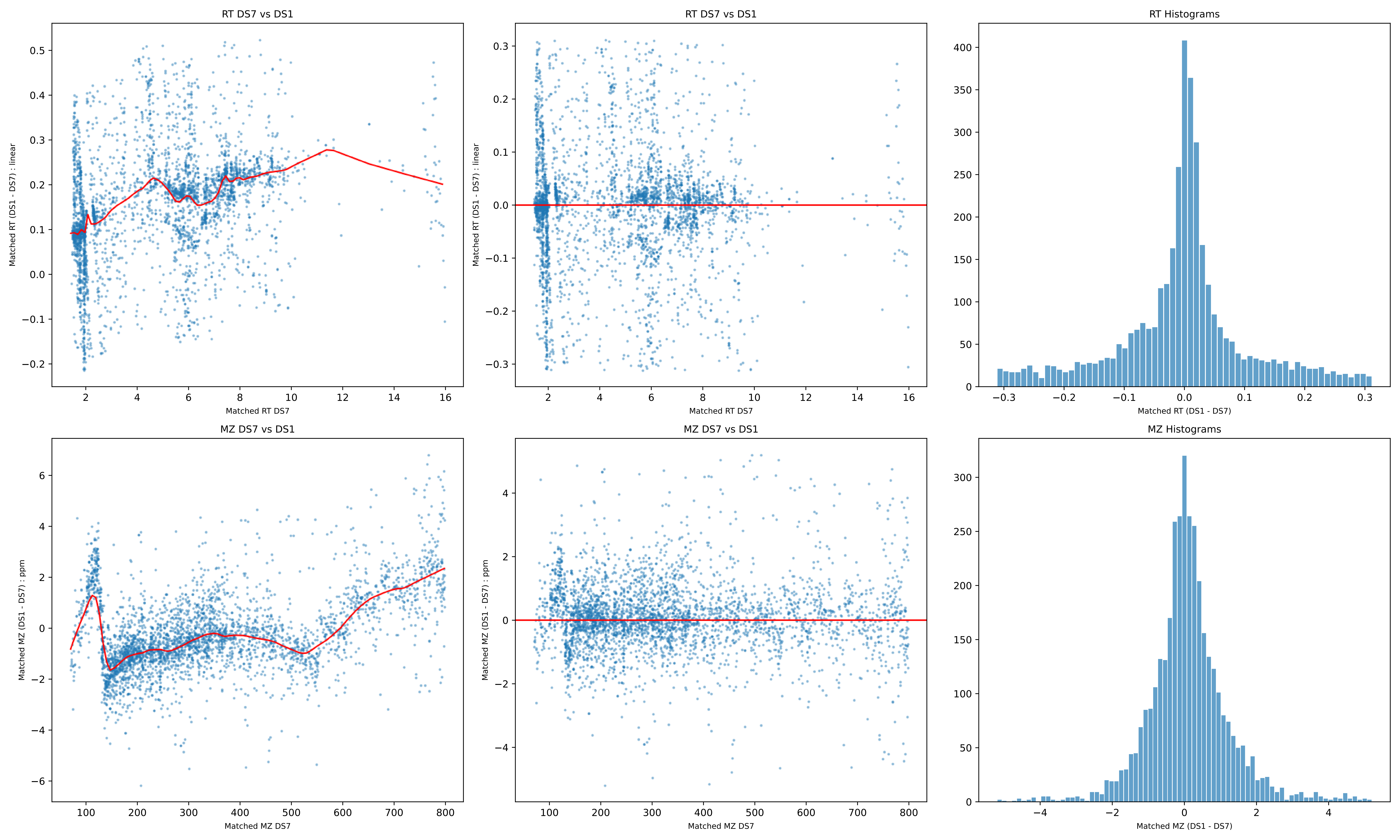
RT DS7 vs DS1
Matched RT (DS1 - DS7) : linear
Matched RT DS7
RT DS7 vs DS1
Matched RT (DS1 - DS7) : linear
Matched RT DS7
RT Histograms
Matched RT (DS1 - DS7)
MZ DS7 vs DS1
Matched MZ (DS1 - DS7) : ppm
Matched MZ DS7
MZ DS7 vs DS1
Matched MZ (DS1 - DS7) : ppm
Matched MZ DS7
MZ Histograms
Matched MZ (DS1 - DS7)

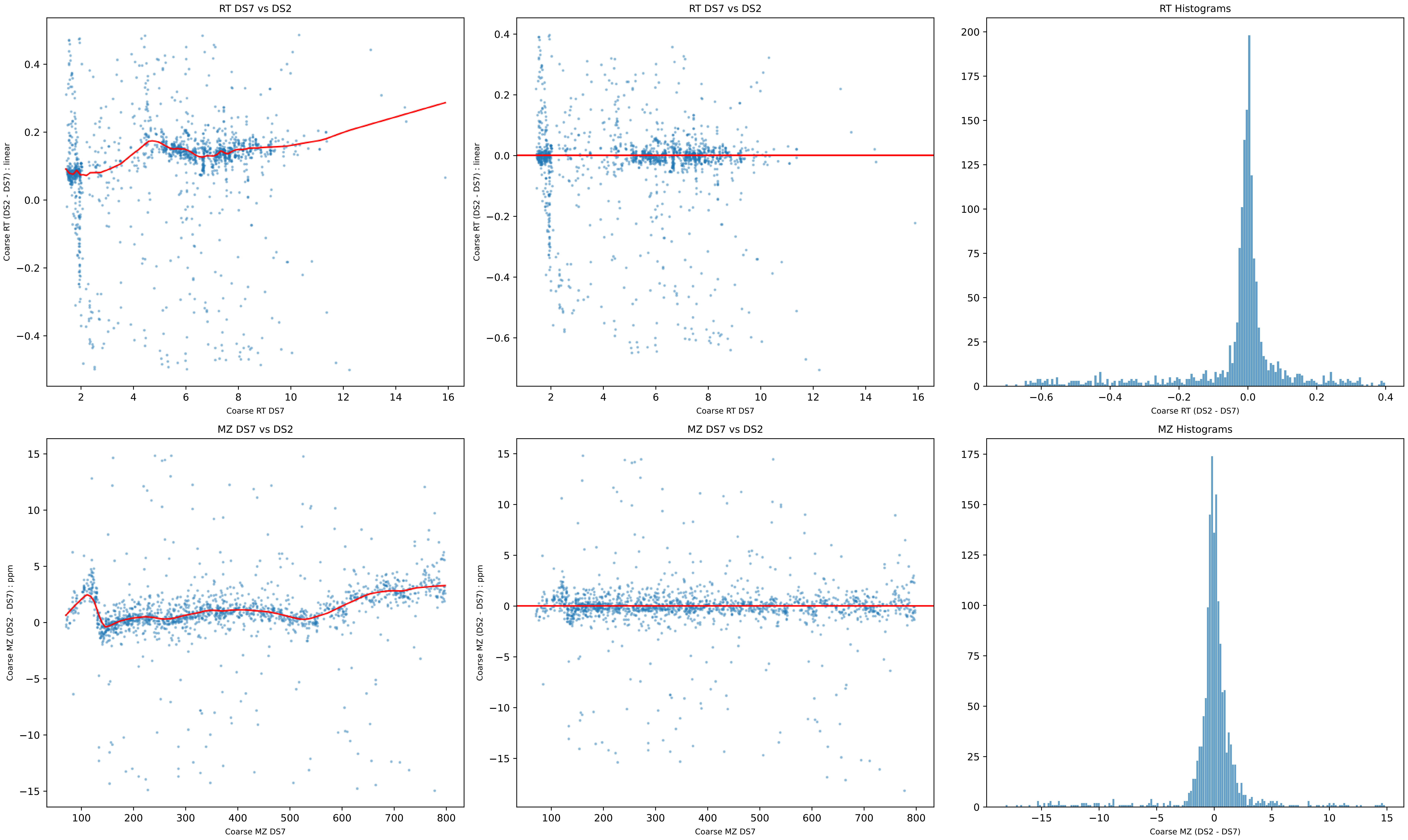
RT DS7 vs DS2
Coarse RT (DS2 - DS7) : linear
Coarse RT DS7
RT DS7 vs DS2
Coarse RT (DS2 - DS7) : linear
Coarse RT DS7
RT Histograms
Coarse RT (DS2 - DS7)
MZ DS7 vs DS2
Coarse MZ (DS2 - DS7) : ppm
Coarse MZ DS7
MZ DS7 vs DS2
Coarse MZ (DS2 - DS7) : ppm
Coarse MZ DS7
MZ Histograms
Coarse MZ (DS2 - DS7)

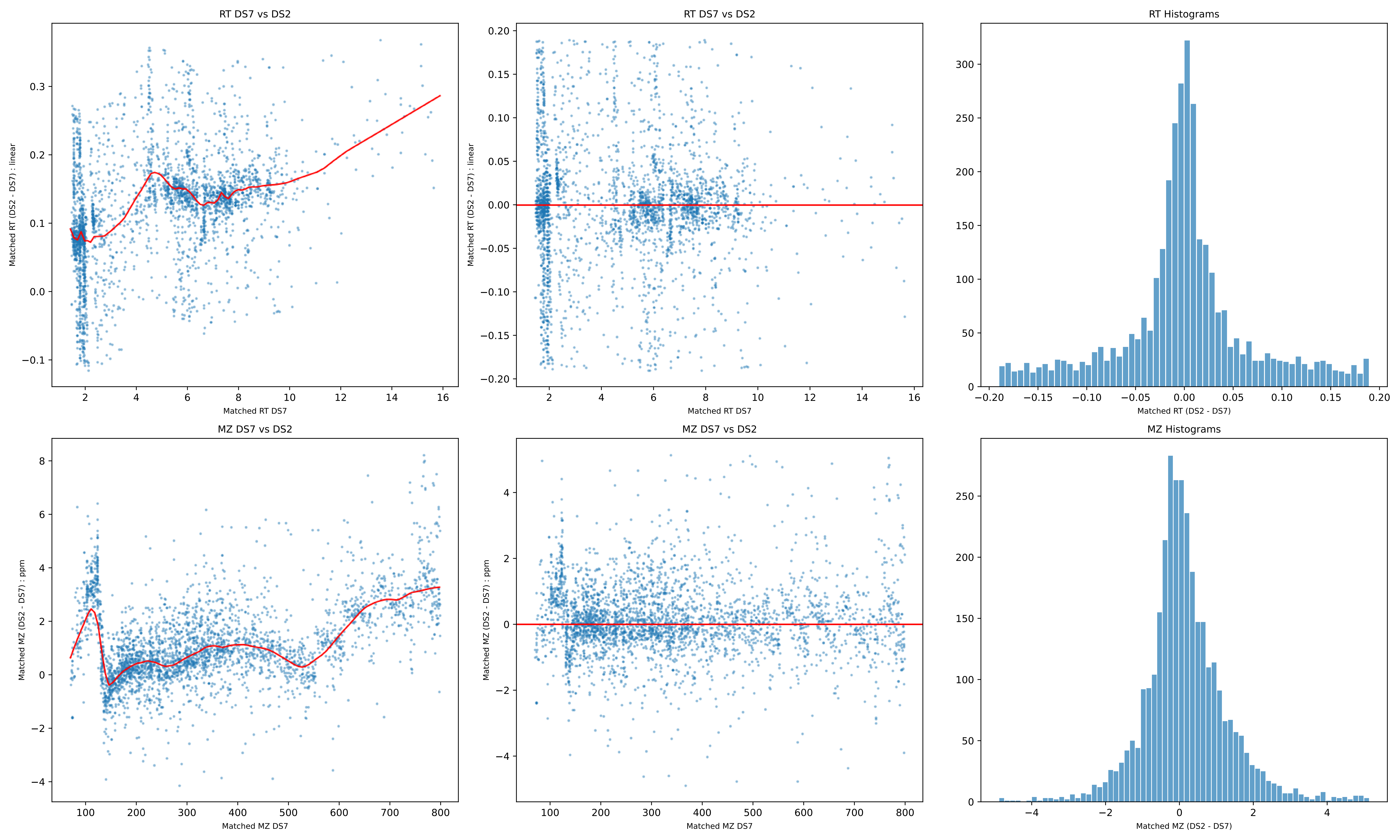
RT DS7 vs DS2
Matched RT (DS2 - DS7) : linear
Matched RT DS7
RT DS7 vs DS2
Matched RT (DS2 - DS7) : linear
Matched RT DS7
RT Histograms
Matched RT (DS2 - DS7)
MZ DS7 vs DS2
Matched MZ (DS2 - DS7) : ppm
Matched MZ DS7
MZ DS7 vs DS2
Matched MZ (DS2 - DS7) : ppm
Matched MZ DS7
MZ Histograms
Matched MZ (DS2 - DS7)

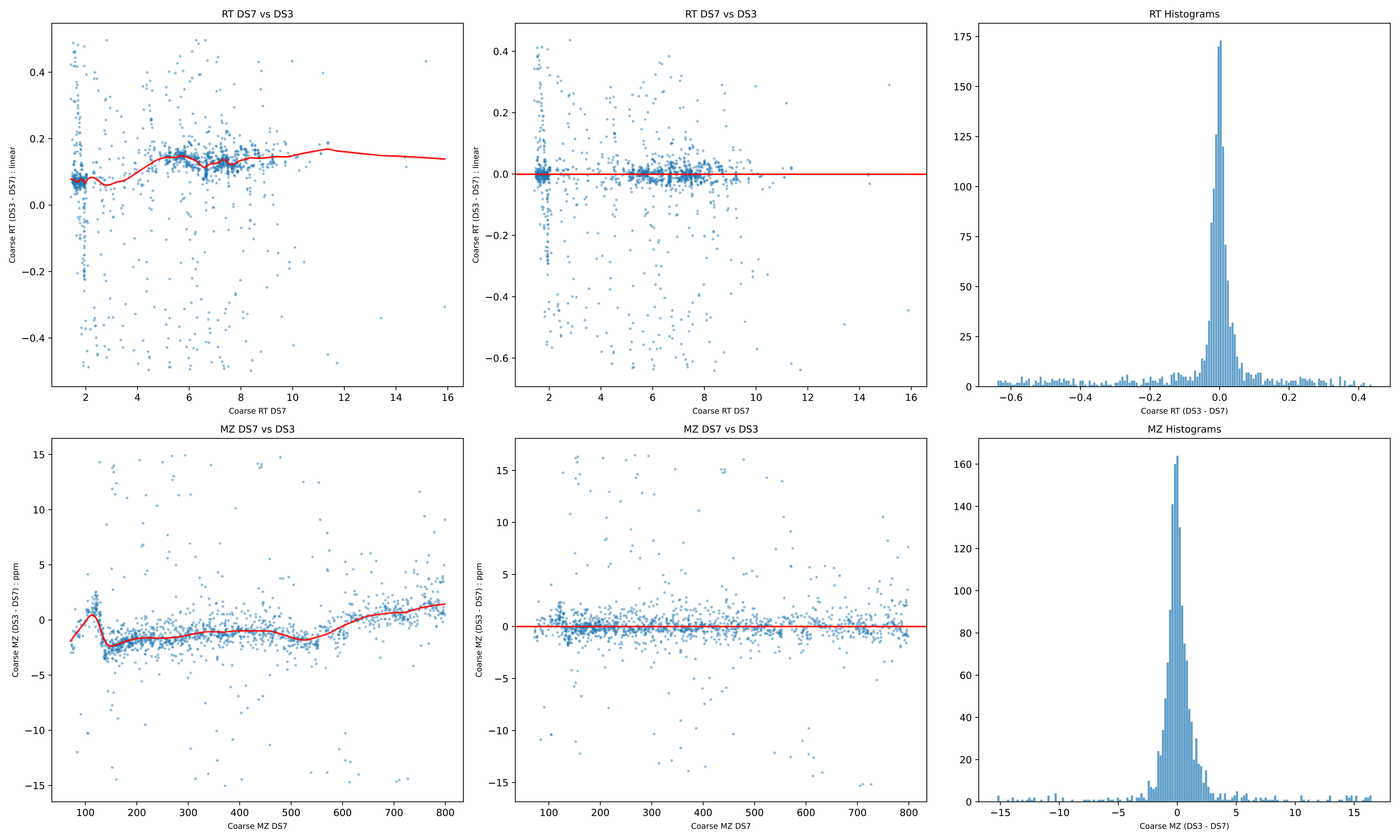
RT DS7 vs DS3
Coarse RT (DS3 - DS7) : linear
Coarse RT DS7
RT DS7 vs DS3
Coarse RT (DS3 - DS7) : linear
Coarse RT DS7
RT Histograms
Coarse RT (DS3 - DS7)
MZ DS7 vs DS3
Coarse MZ (DS3 - DS7) : ppm
Coarse MZ DS7
MZ DS7 vs DS3
Coarse MZ (DS3 - DS7) : ppm
Coarse MZ DS7
MZ Histograms
Coarse MZ (DS3 - DS7)

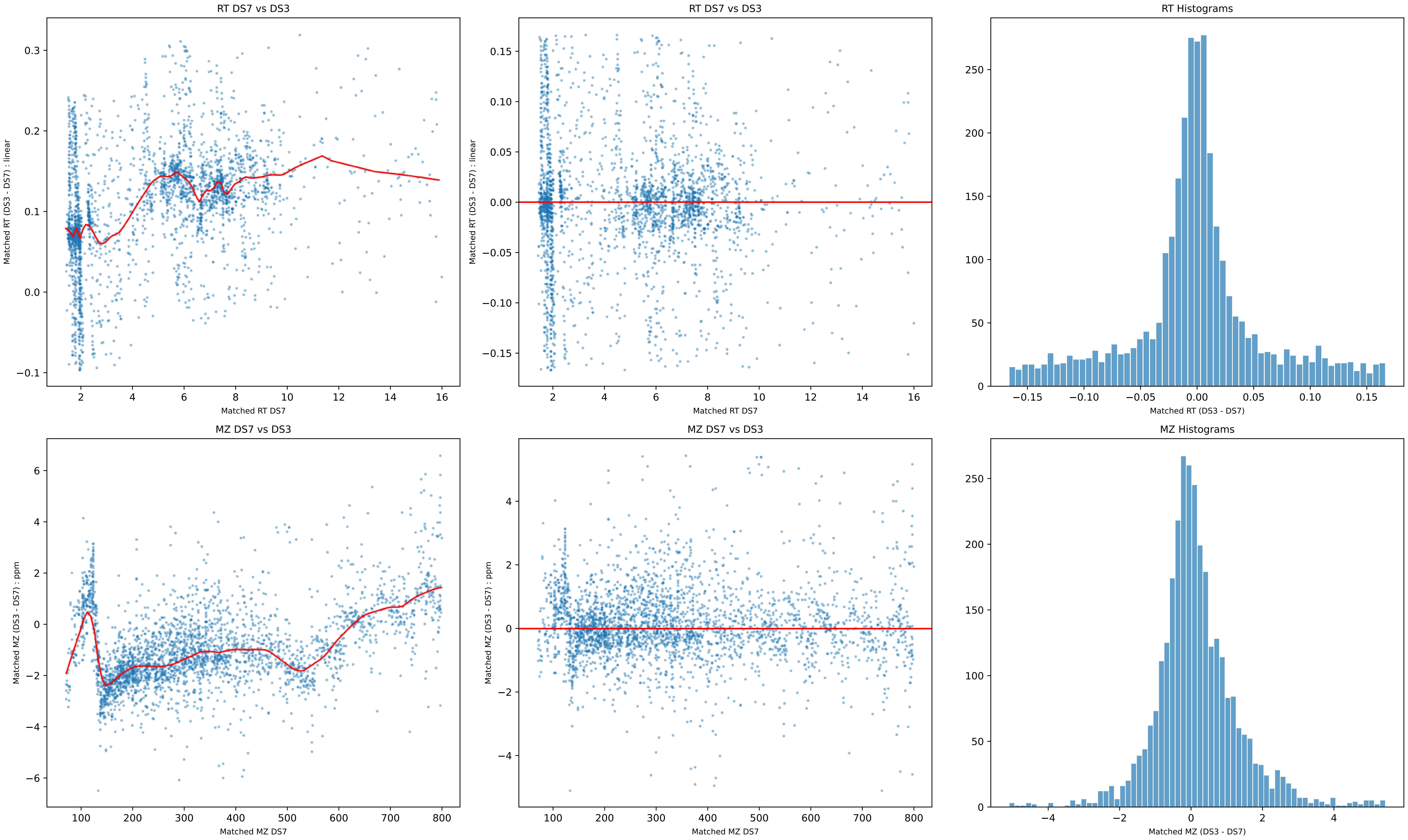

RT DS7 vs DS3
Matched RT (DS3 - DS7) : linear
Matched RT DS7
RT DS7 vs DS3
Matched RT (DS3 - DS7) : linear
Matched RT DS7
RT Histograms
Matched RT (DS3 - DS7)
MZ DS7 vs DS3
Matched MZ (DS3 - DS7) : ppm
Matched MZ DS7
MZ DS7 vs DS3
Matched MZ (DS3 - DS7) : ppm
Matched MZ DS7
MZ Histograms
Matched MZ (DS3 - DS7)

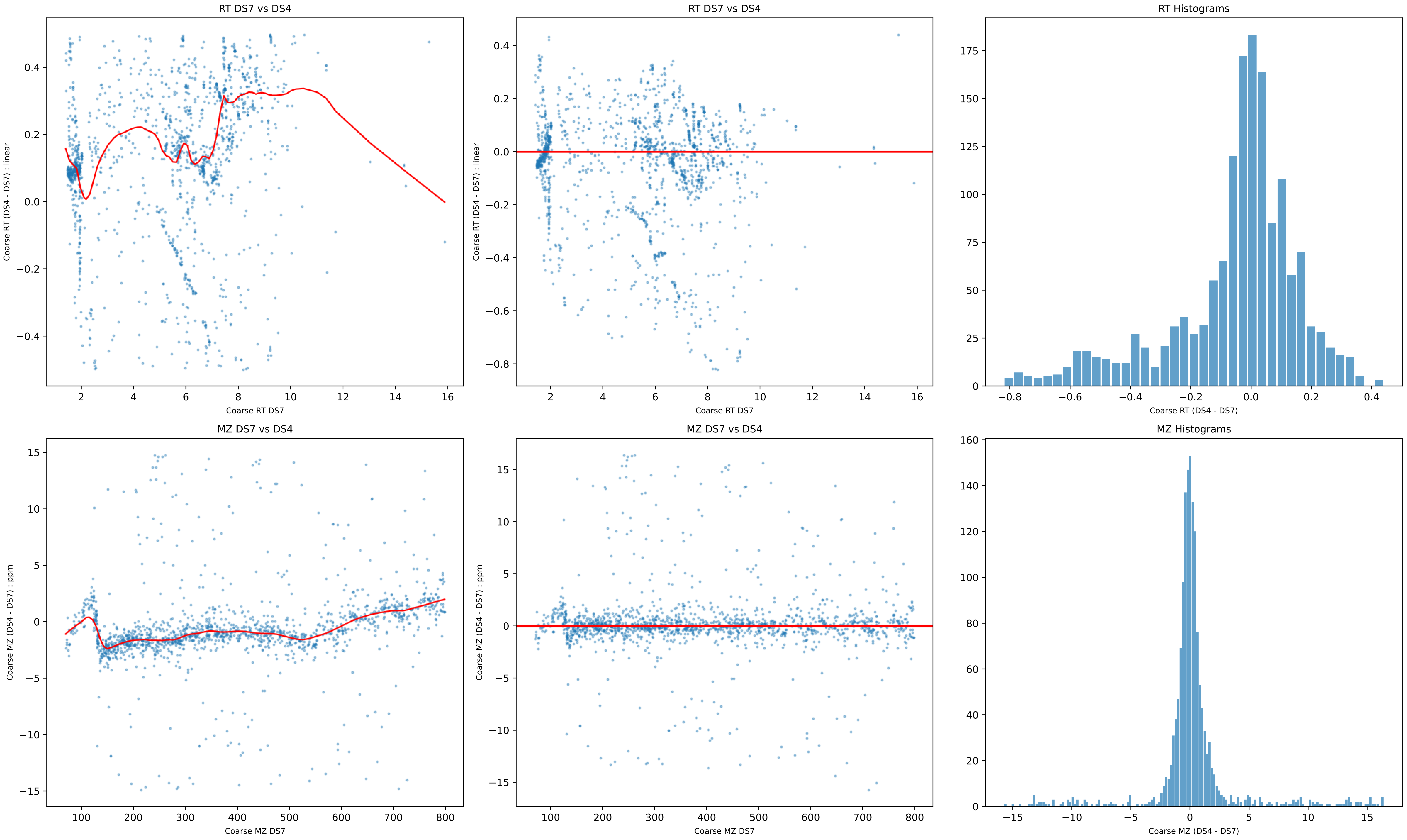
RT DS7 vs DS4
Coarse RT (DS4 - DS7) : linear
Coarse RT DS7
RT DS7 vs DS4
Coarse RT (DS4 - DS7) : linear
Coarse RT DS7
RT Histograms
Coarse RT (DS4 - DS7)
MZ DS7 vs DS4
Coarse MZ (DS4 - DS7) : ppm
Coarse MZ DS7
MZ DS7 vs DS4
Coarse MZ (DS4 - DS7) : ppm
Coarse MZ DS7
MZ Histograms
Coarse MZ (DS4 - DS7)

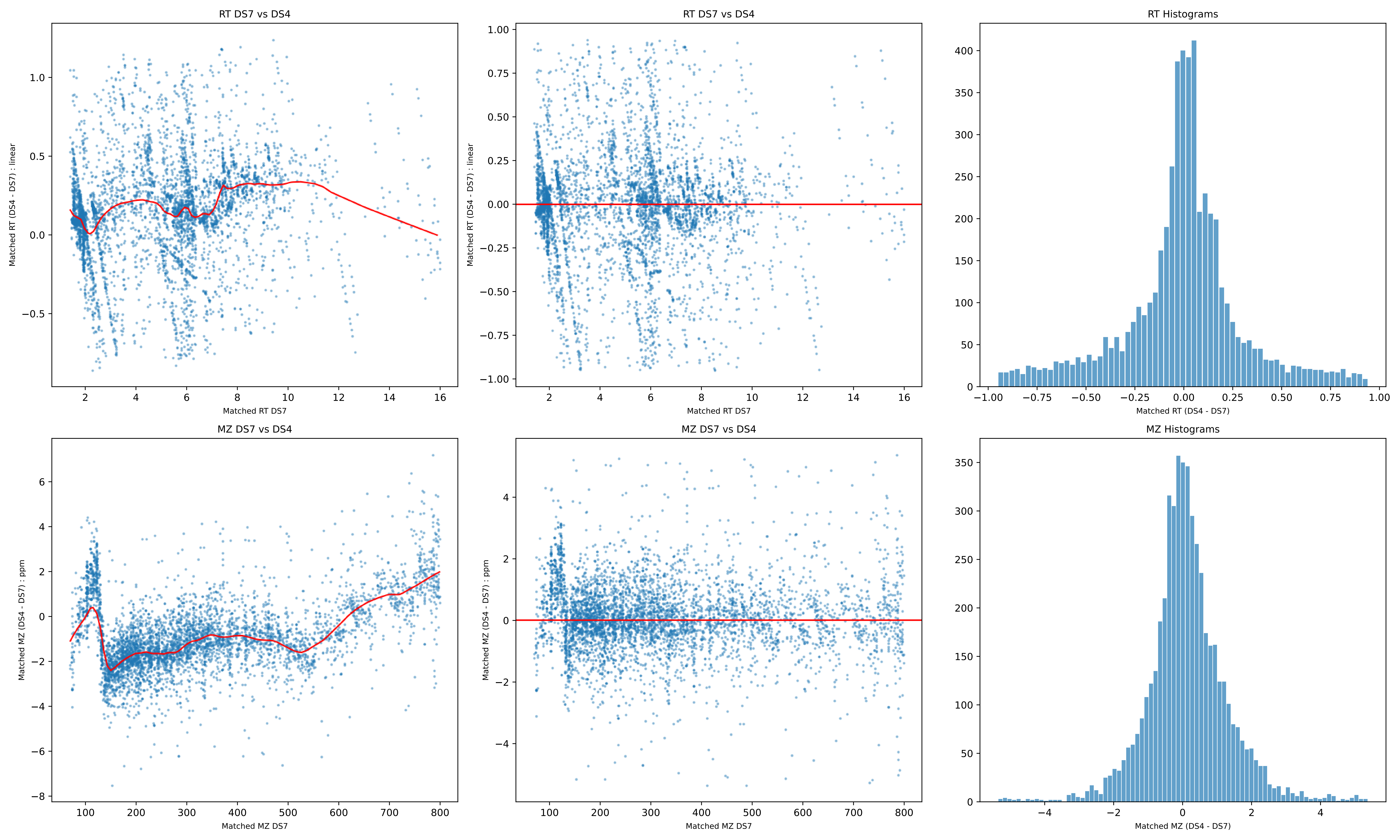

RT DS7 vs DS4
Matched RT (DS4 - DS7) : linear
Matched RT DS7
RT DS7 vs DS4
Matched RT (DS4 - DS7) : linear
Matched RT DS7
RT Histograms
Matched RT (DS4 - DS7)
MZ DS7 vs DS4
Matched MZ (DS4 - DS7) : ppm
Matched MZ DS7
MZ DS7 vs DS4
Matched MZ (DS4 - DS7) : ppm
Matched MZ DS7
MZ Histograms
Matched MZ (DS4 - DS7)

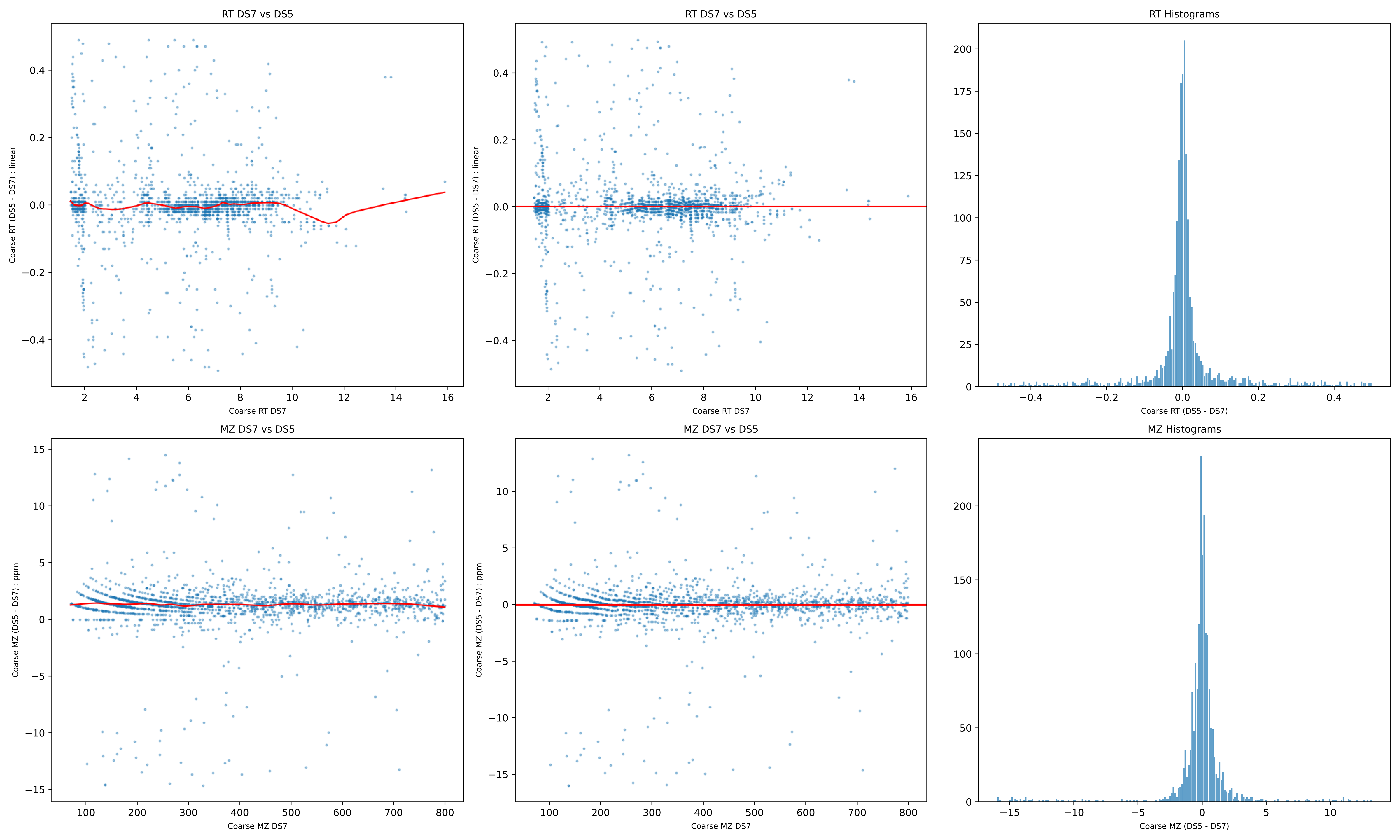

RT DS7 vs DS5
Coarse RT (DS5 - DS7) : linear
Coarse RT DS7
RT DS7 vs DS5
Coarse RT (DS5 - DS7) : linear
Coarse RT DS7
RT Histograms
Coarse RT (DS5 - DS7)
MZ DS7 vs DS5
Coarse MZ (DS5 - DS7) : ppm
Coarse MZ DS7
MZ DS7 vs DS5
Coarse MZ (DS5 - DS7) : ppm
Coarse MZ DS7
MZ Histograms
Coarse MZ (DS5 - DS7)

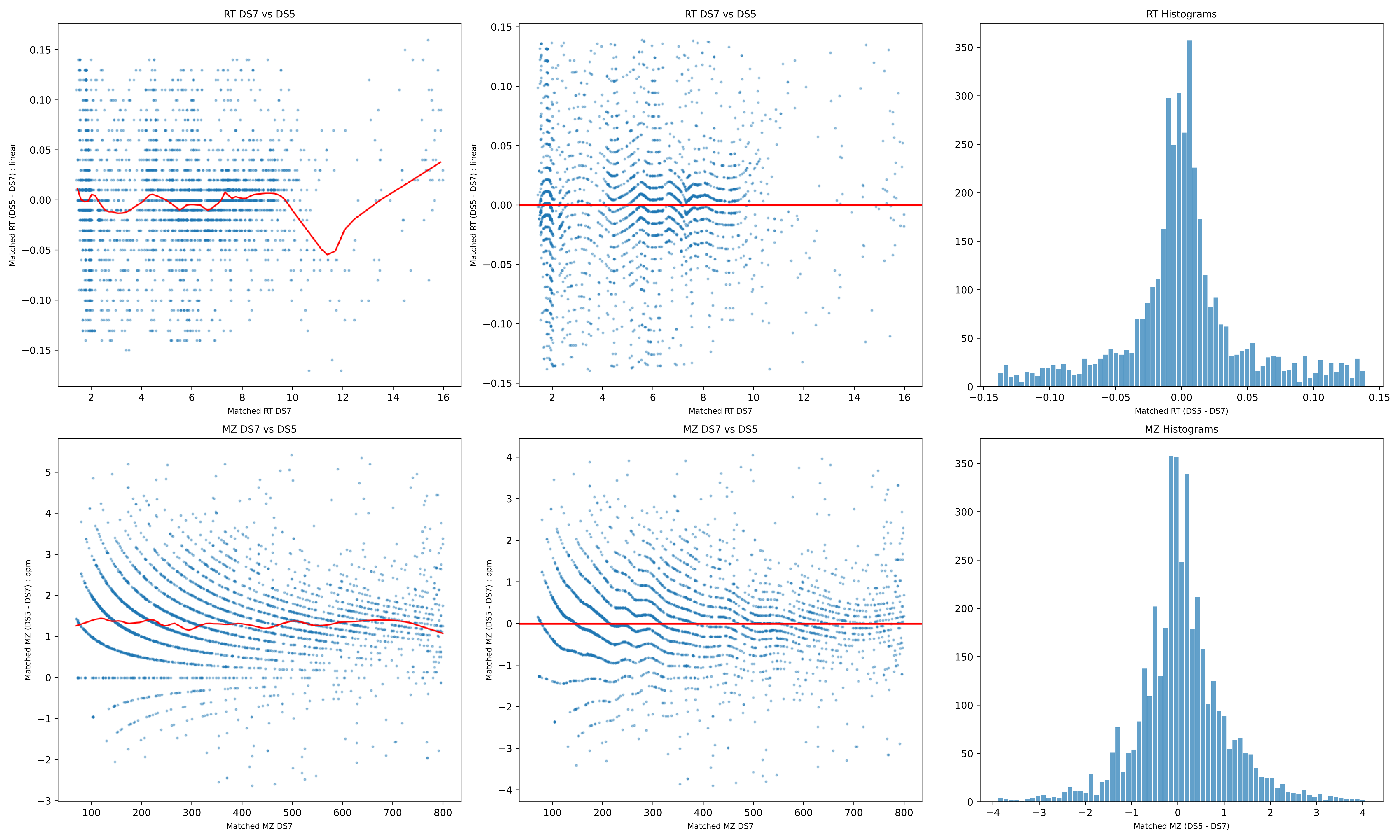
RT DS7 vs DS5
Matched RT (DS5 - DS7) : linear
Matched RT DS7
RT DS7 vs DS5
Matched RT (DS5 - DS7) : linear
Matched RT DS7
RT Histograms
Matched RT (DS5 - DS7)
MZ DS7 vs DS5
Matched MZ (DS5 - DS7) : ppm
Matched MZ DS7
MZ DS7 vs DS5
Matched MZ (DS5 - DS7) : ppm
Matched MZ DS7
MZ Histograms
Matched MZ (DS5 - DS7)

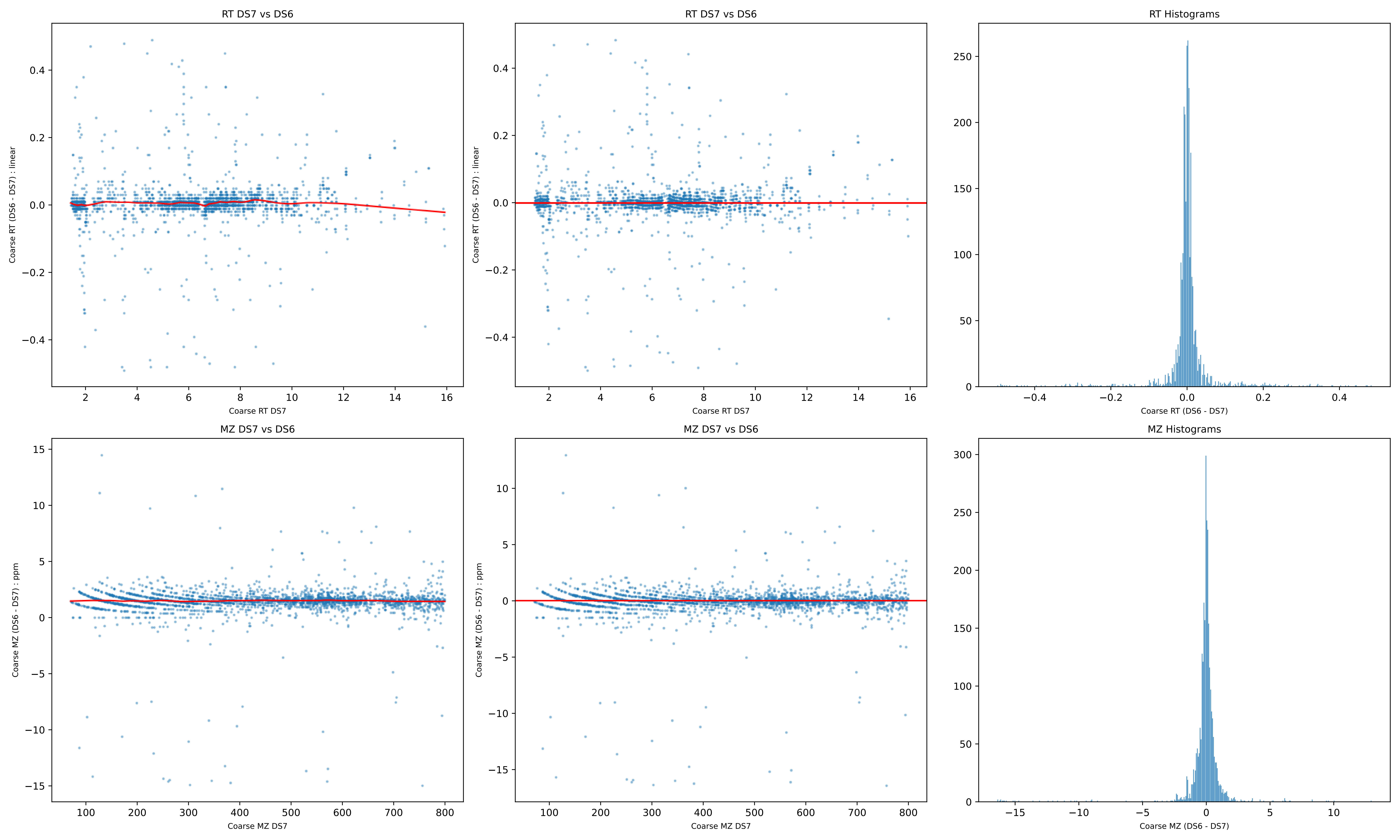
RT DS7 vs DS6
Coarse RT (DS6 - DS7) : linear
Coarse RT DS7
RT DS7 vs DS6
Coarse RT (DS6 - DS7) : linear
Coarse RT DS7
RT Histograms
Coarse RT (DS6 - DS7)
MZ DS7 vs DS6
Coarse MZ (DS6 - DS7) : ppm
Coarse MZ DS7
MZ DS7 vs DS6
Coarse MZ (DS6 - DS7) : ppm
Coarse MZ DS7
MZ Histograms
Coarse MZ (DS6 - DS7)

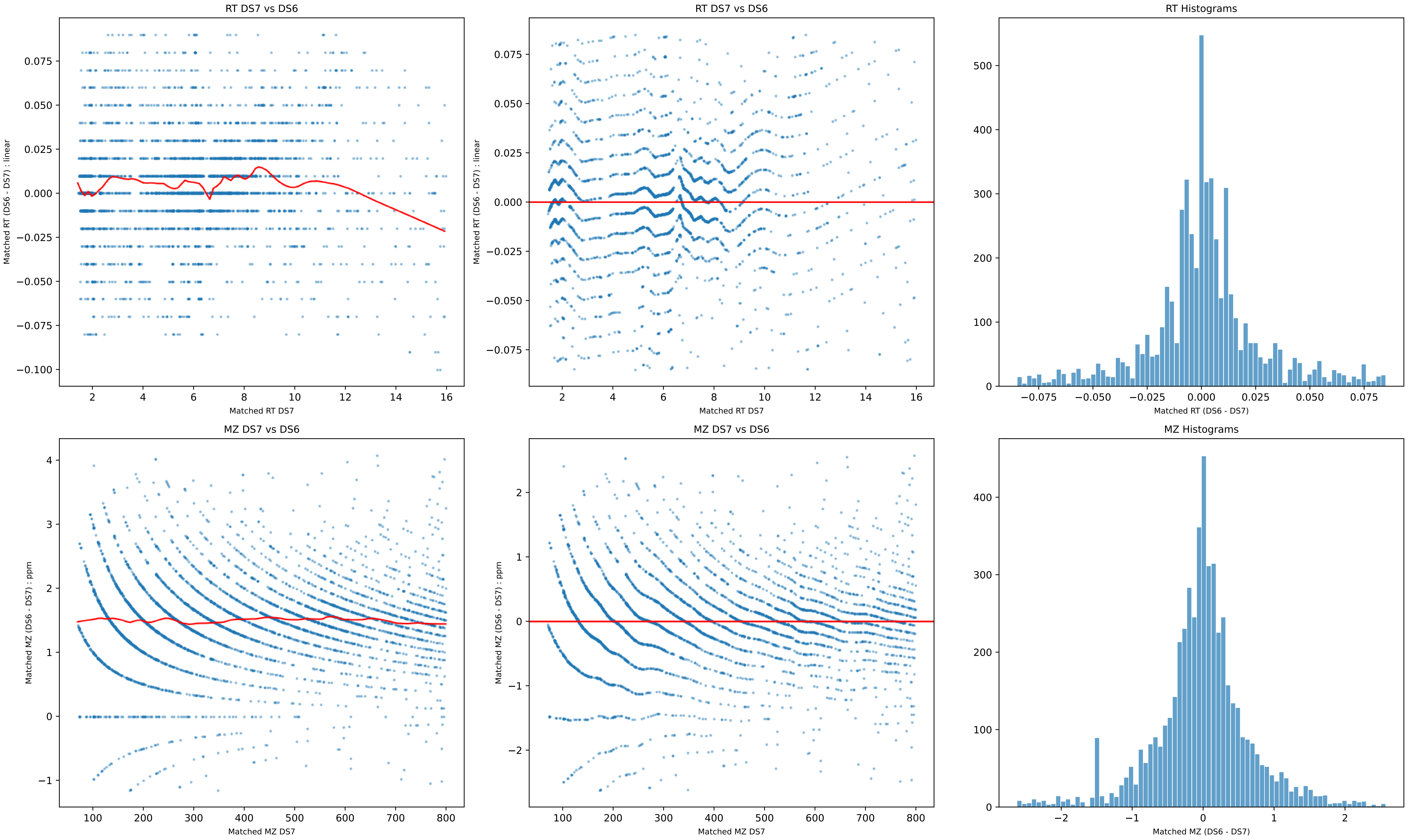
RT DS7 vs DS6
Matched RT (DS6 - DS7) : linear
Matched RT DS7
RT DS7 vs DS6
Matched RT (DS6 - DS7) : linear
Matched RT DS7
RT Histograms
Matched RT (DS6 - DS7)
MZ DS7 vs DS6
Matched MZ (DS6 - DS7) : ppm
Matched MZ DS7
MZ DS7 vs DS6
Matched MZ (DS6 - DS7) : ppm
Matched MZ DS7
MZ Histograms
Matched MZ (DS6 - DS7)

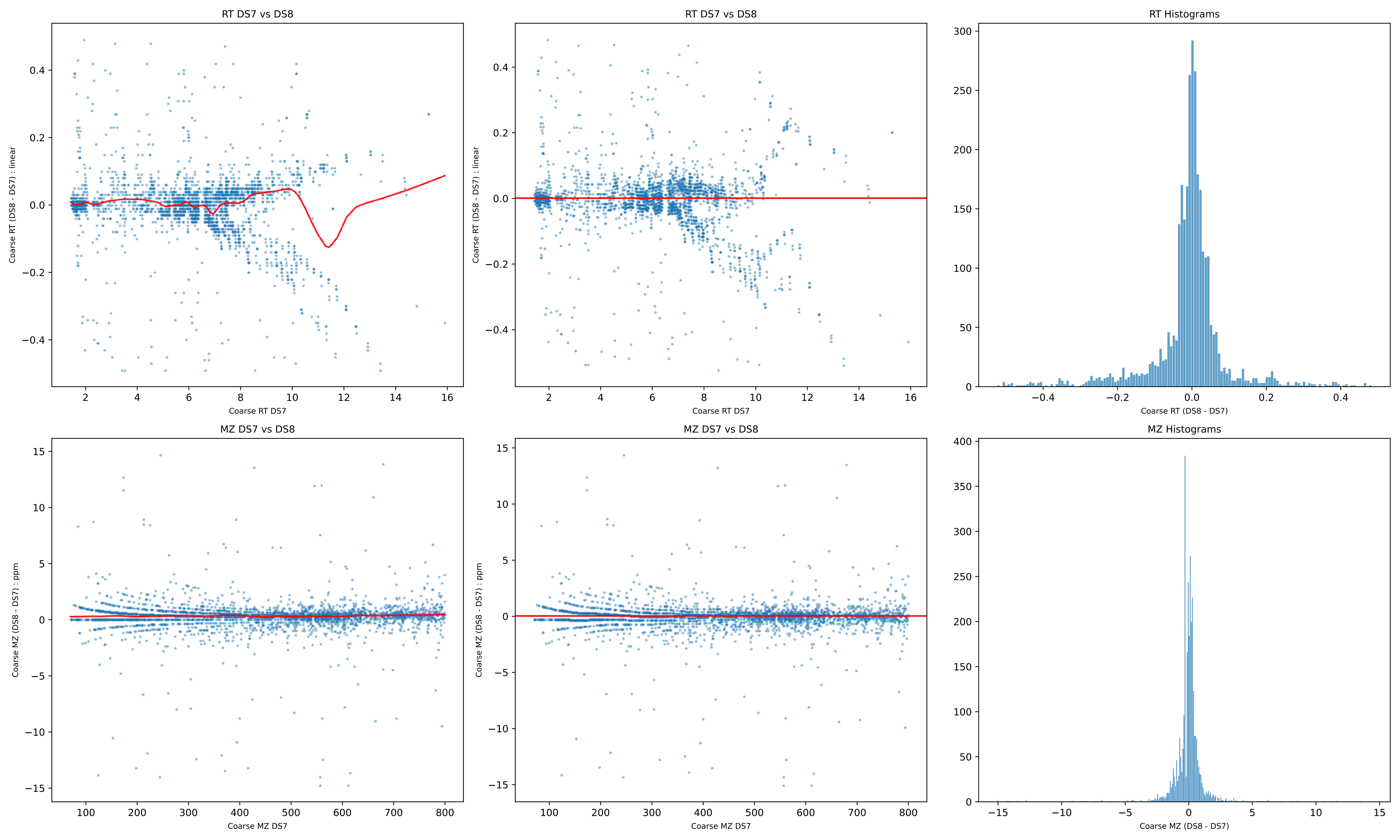
RT DS7 vs DS8
Coarse RT (DS8 - DS7) : linear
Coarse RT DS7
RT DS7 vs DS8
Coarse RT (DS8 - DS7) : linear
Coarse RT DS7
RT Histograms
Coarse RT (DS8 - DS7)
MZ DS7 vs DS8
Coarse MZ (DS8 - DS7) : ppm
Coarse MZ DS7
MZ DS7 vs DS8
Coarse MZ (DS8 - DS7) : ppm
Coarse MZ DS7
MZ Histograms
Coarse MZ (DS8 - DS7)

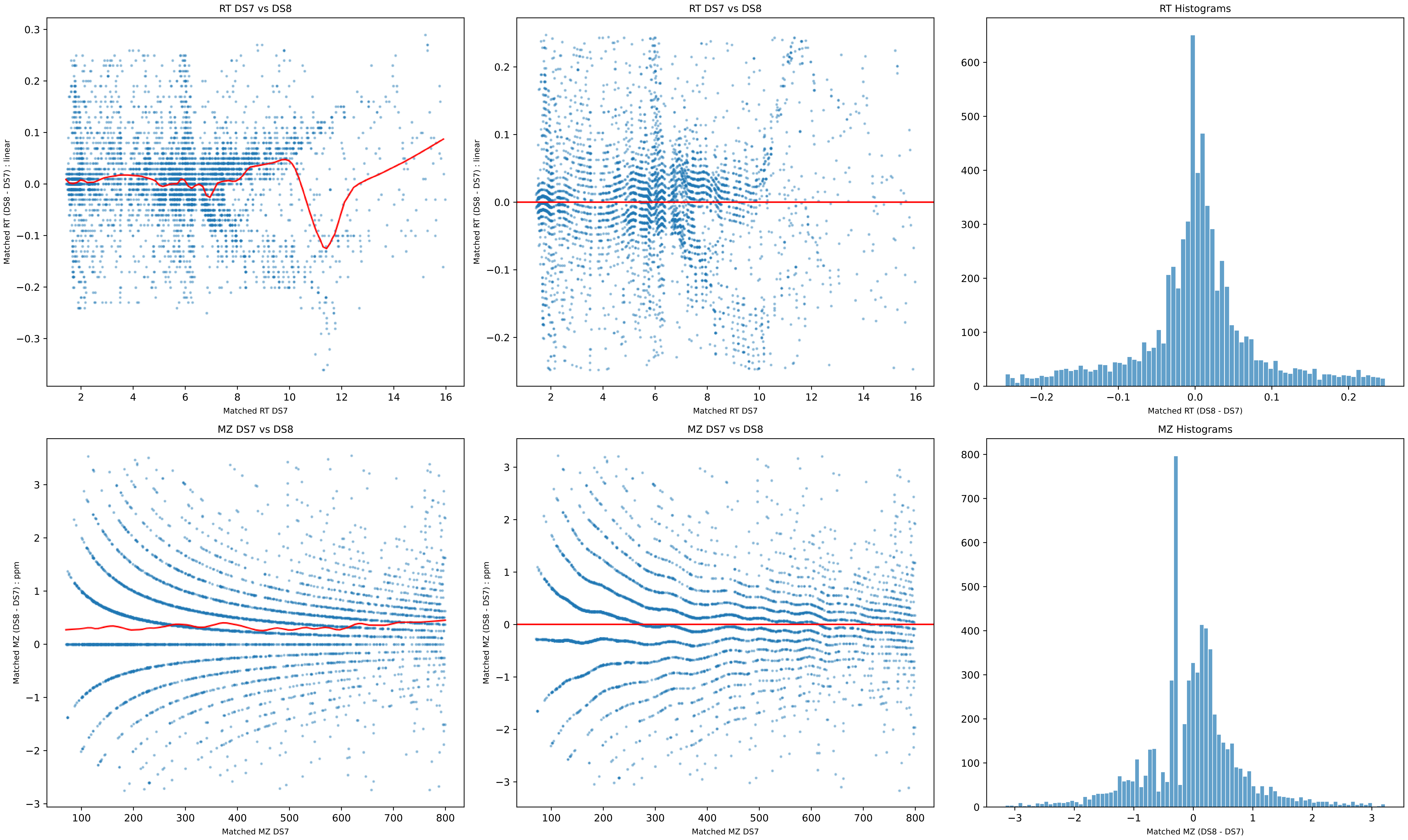
RT DS7 vs DS8
Matched RT (DS8 - DS7) : linear
Matched RT DS7
RT DS7 vs DS8
Matched RT (DS8 - DS7) : linear
Matched RT DS7
RT Histograms
Matched RT (DS8 - DS7)
MZ DS7 vs DS8
Matched MZ (DS8 - DS7) : ppm
Matched MZ DS7
MZ DS7 vs DS8
Matched MZ (DS8 - DS7) : ppm
Matched MZ DS7
MZ Histograms
Matched MZ (DS8 - DS7)

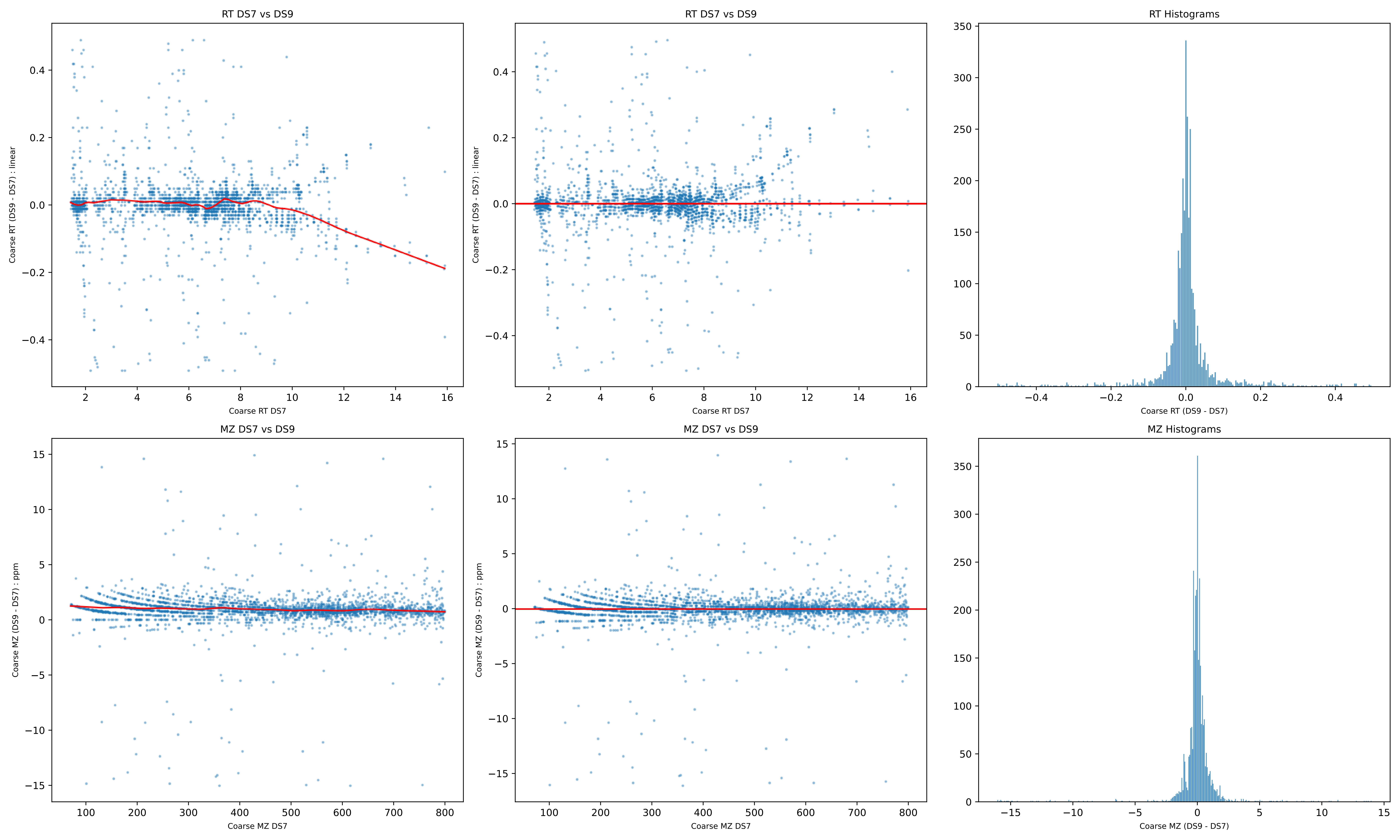
RT DS7 vs DS9
Coarse RT (DS9 - DS7) : linear
Coarse RT DS7
RT DS7 vs DS9
Coarse RT (DS9 - DS7) : linear
Coarse RT DS7
RT Histograms
Coarse RT (DS9 - DS7)
MZ DS7 vs DS9
Coarse MZ (DS9 - DS7) : ppm
Coarse MZ DS7
MZ DS7 vs DS9
Coarse MZ (DS9 - DS7) : ppm
Coarse MZ DS7
MZ Histograms
Coarse MZ (DS9 - DS7)

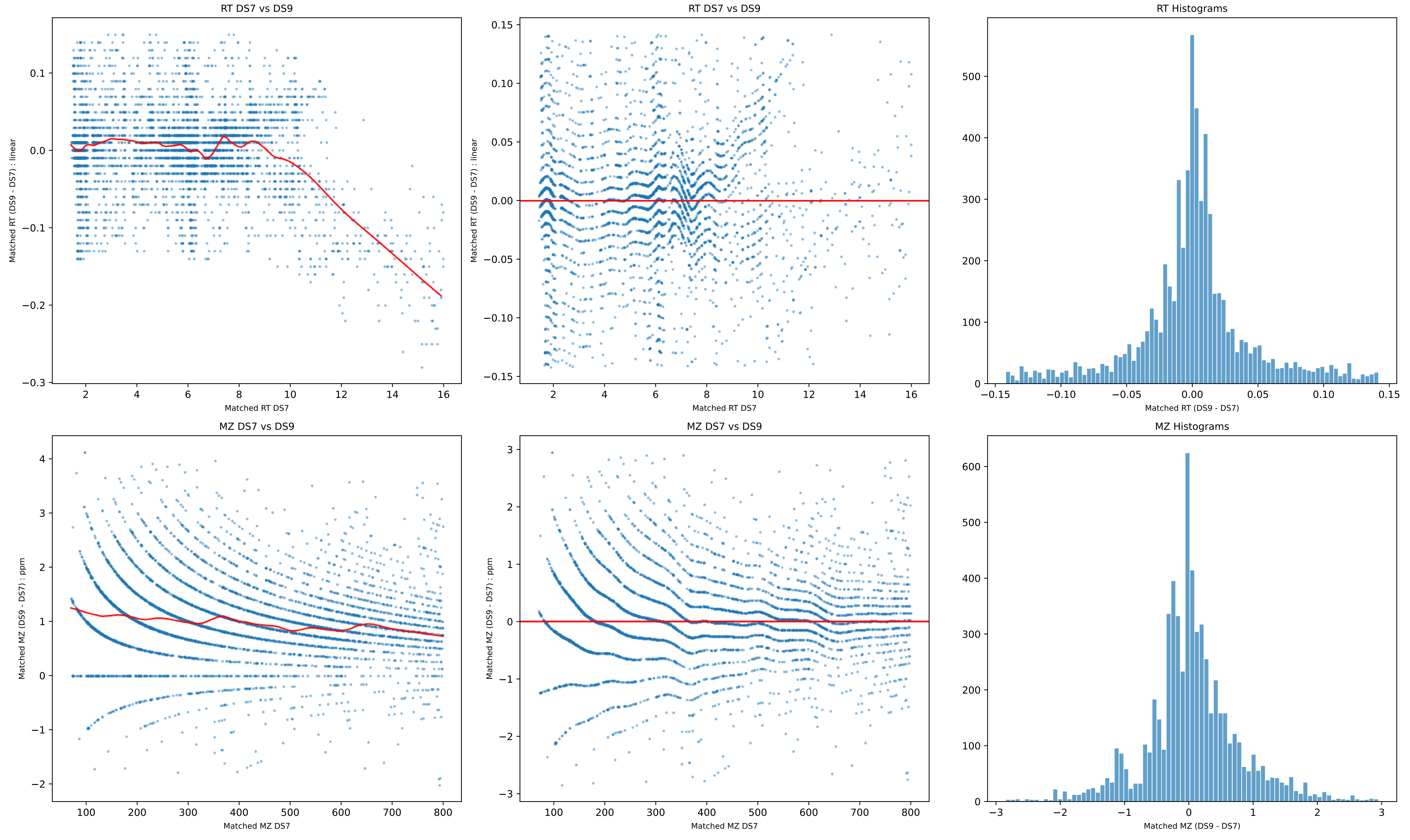
RT DS7 vs DS9
Matched RT (DS9 - DS7) : linear
Matched RT DS7
RT DS7 vs DS9
Matched RT (DS9 - DS7) : linear
Matched RT DS7
RT Histograms
Matched RT (DS9 - DS7)
MZ DS7 vs DS9
Matched MZ (DS9 - DS7) : ppm
Matched MZ DS7
MZ DS7 vs DS9
Matched MZ (DS9 - DS7) : ppm
Matched MZ DS7
MZ Histograms
Matched MZ (DS9 - DS7)

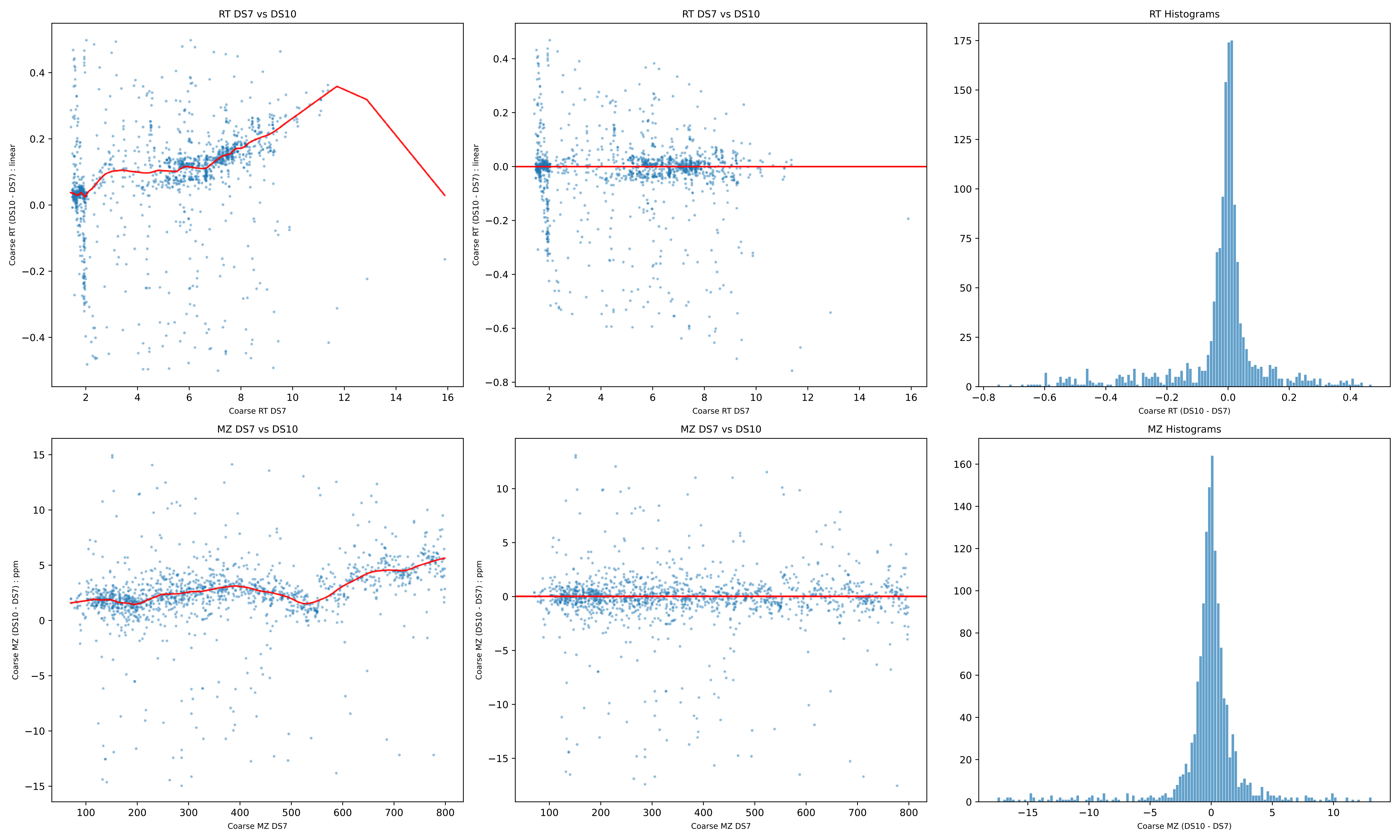

RT DS7 vs DS10
RT DS7 vs DS10
RT Histograms
Coarse RT (DS10 - DS7) : linear
Coarse RT DS7
Coarse RT (DS10 - DS7)
MZ DS7 vs DS10
MZ DS7 vs DS10
MZ Histograms
Coarse MZ (DS10 - DS7) : ppm
Coarse MZ DS7
Coarse MZ (DS10 - DS7)

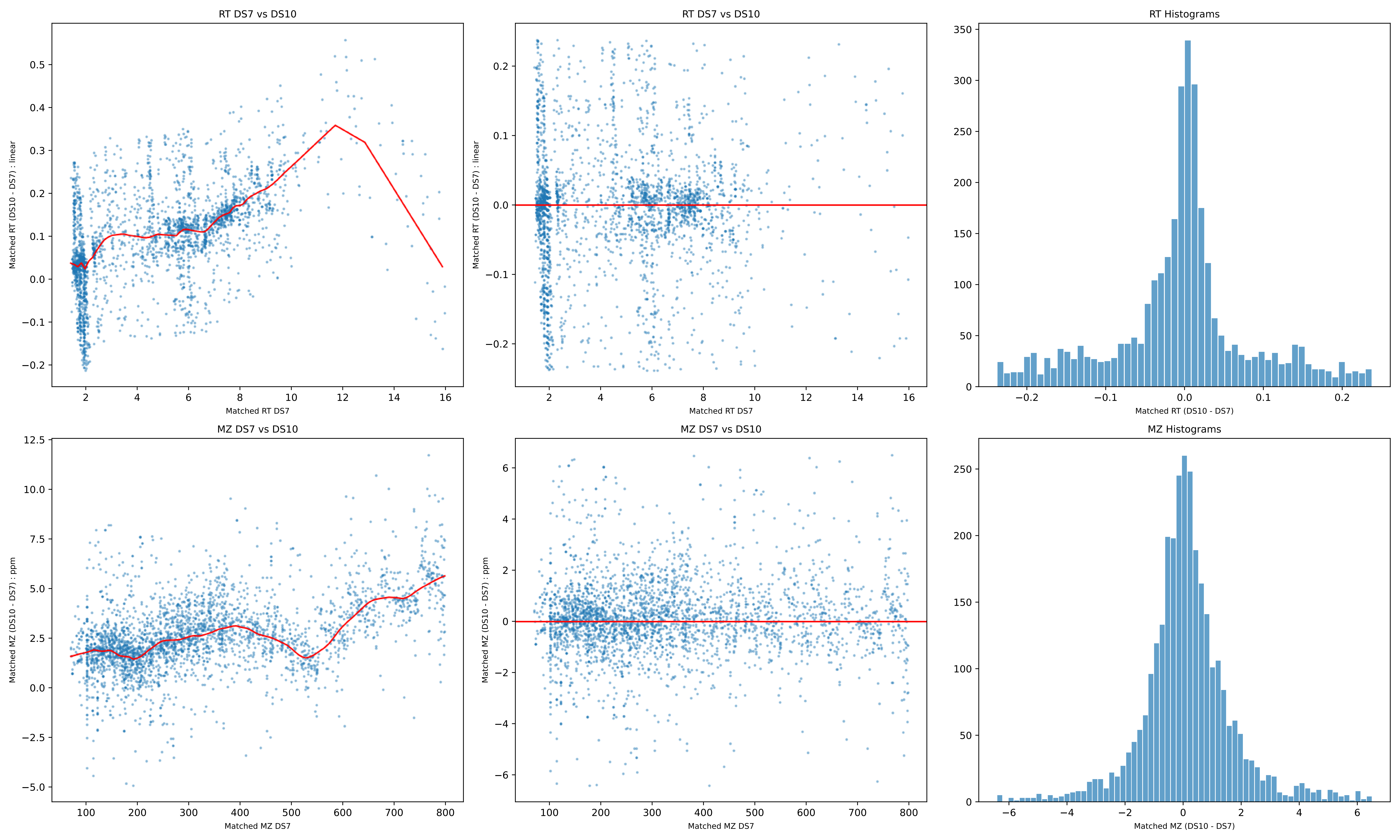
RT DS7 vs DS10
Matched RT (DS10 - DS7) : linear
Matched RT DS7
RT DS7 vs DS10
Matched RT (DS10 - DS7) : linear
Matched RT DS7
RT Histograms
Matched RT (DS10 - DS7)
MZ DS7 vs DS10
Matched MZ (DS10 - DS7) : ppm
Matched MZ DS7
MZ DS7 vs DS10
Matched MZ (DS10 - DS7) : ppm
Matched MZ DS7
MZ Histograms
Matched MZ (DS10 - DS7)

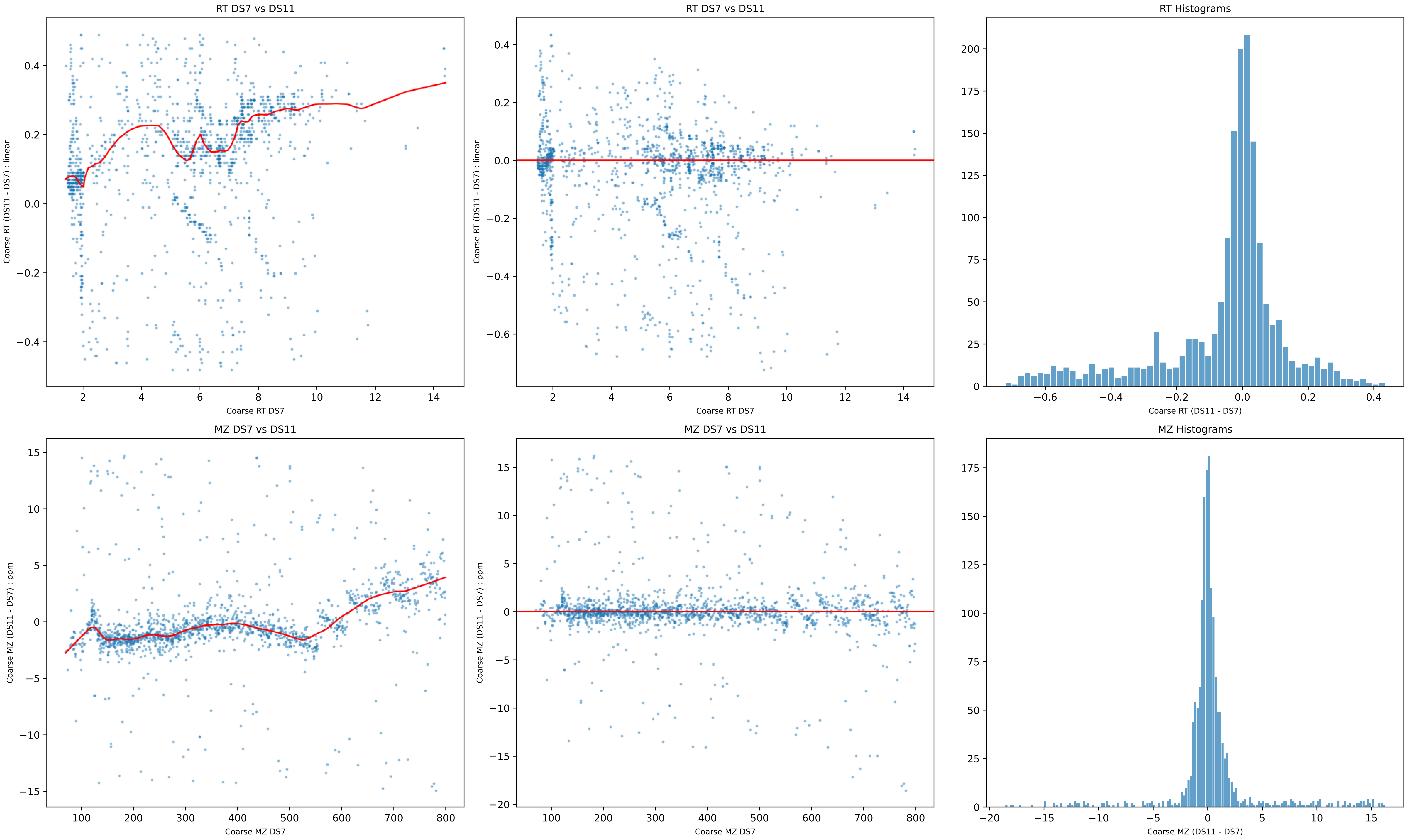
RT DS7 vs DS11
Coarse RT (DS11 - DS7) : linear
Coarse RT DS7
RT DS7 vs DS11
Coarse RT (DS11 - DS7) : linear
Coarse RT DS7
RT Histograms
Coarse RT (DS11 - DS7)
MZ DS7 vs DS11
Coarse MZ (DS11 - DS7) : ppm
Coarse MZ DS7
MZ DS7 vs DS11
Coarse MZ (DS11 - DS7) : ppm
Coarse MZ DS7
MZ Histograms
Coarse MZ (DS11 - DS7)

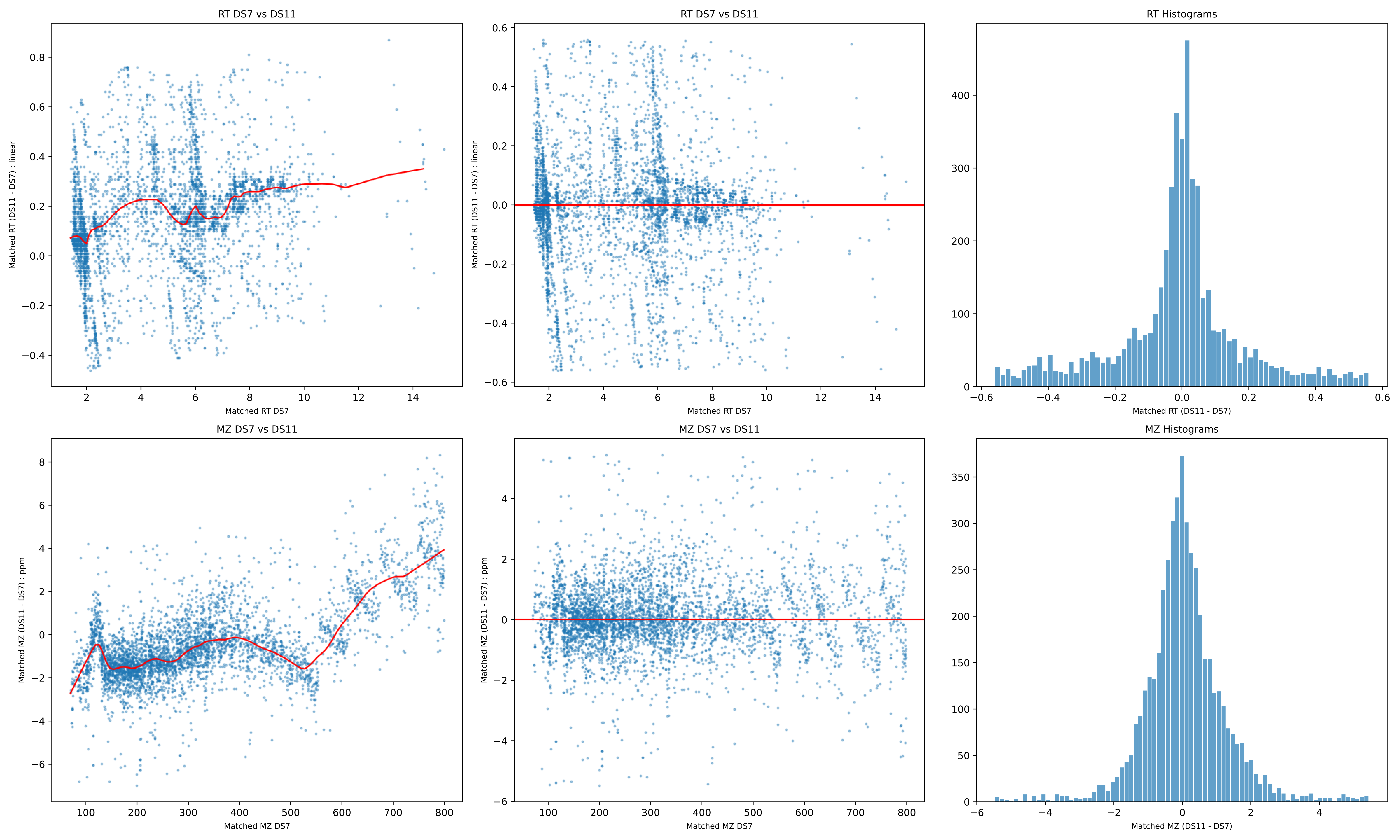
RT DS7 vs DS11
Matched RT (DS11 - DS7) : linear
Matched RT DS7
RT DS7 vs DS11
Matched RT (DS11 - DS7) : linear
Matched RT DS7
RT Histograms
Matched RT (DS11 - DS7)
MZ DS7 vs DS11
Matched MZ (DS11 - DS7) : ppm
Matched MZ DS7
MZ DS7 vs DS11
Matched MZ (DS11 - DS7) : ppm
Matched MZ DS7
MZ Histograms
Matched MZ (DS11 - DS7)

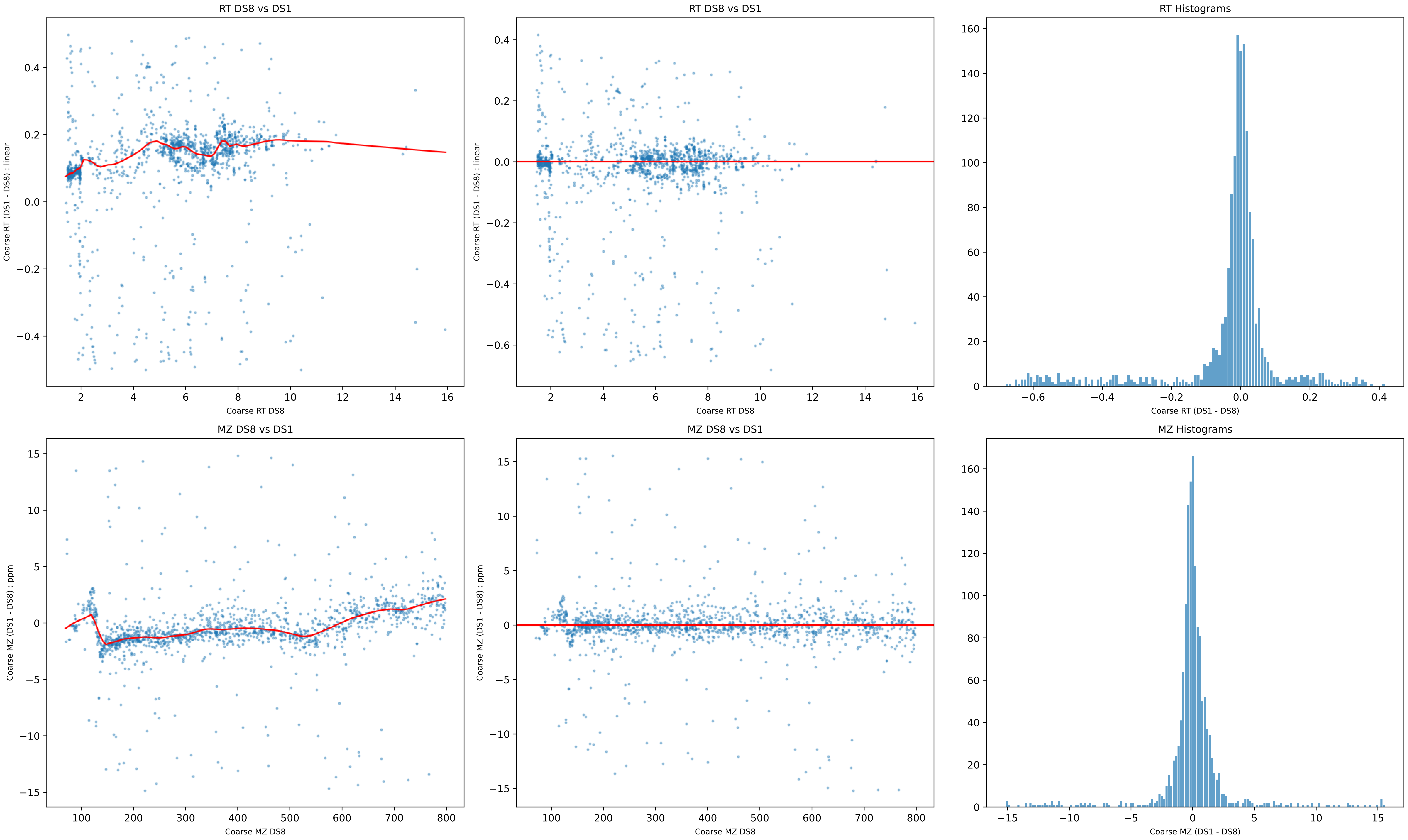

RT DS8 vs DS1
Coarse RT (DS1 - DS8) : linear
Coarse RT DS8
RT DS8 vs DS1
Coarse RT (DS1 - DS8) : linear
Coarse RT DS8
RT Histograms
Coarse RT (DS1 - DS8)
MZ DS8 vs DS1
Coarse MZ (DS1 - DS8) : ppm
Coarse MZ DS8
MZ DS8 vs DS1
Coarse MZ (DS1 - DS8) : ppm
Coarse MZ DS8
MZ Histograms
Coarse MZ (DS1 - DS8)

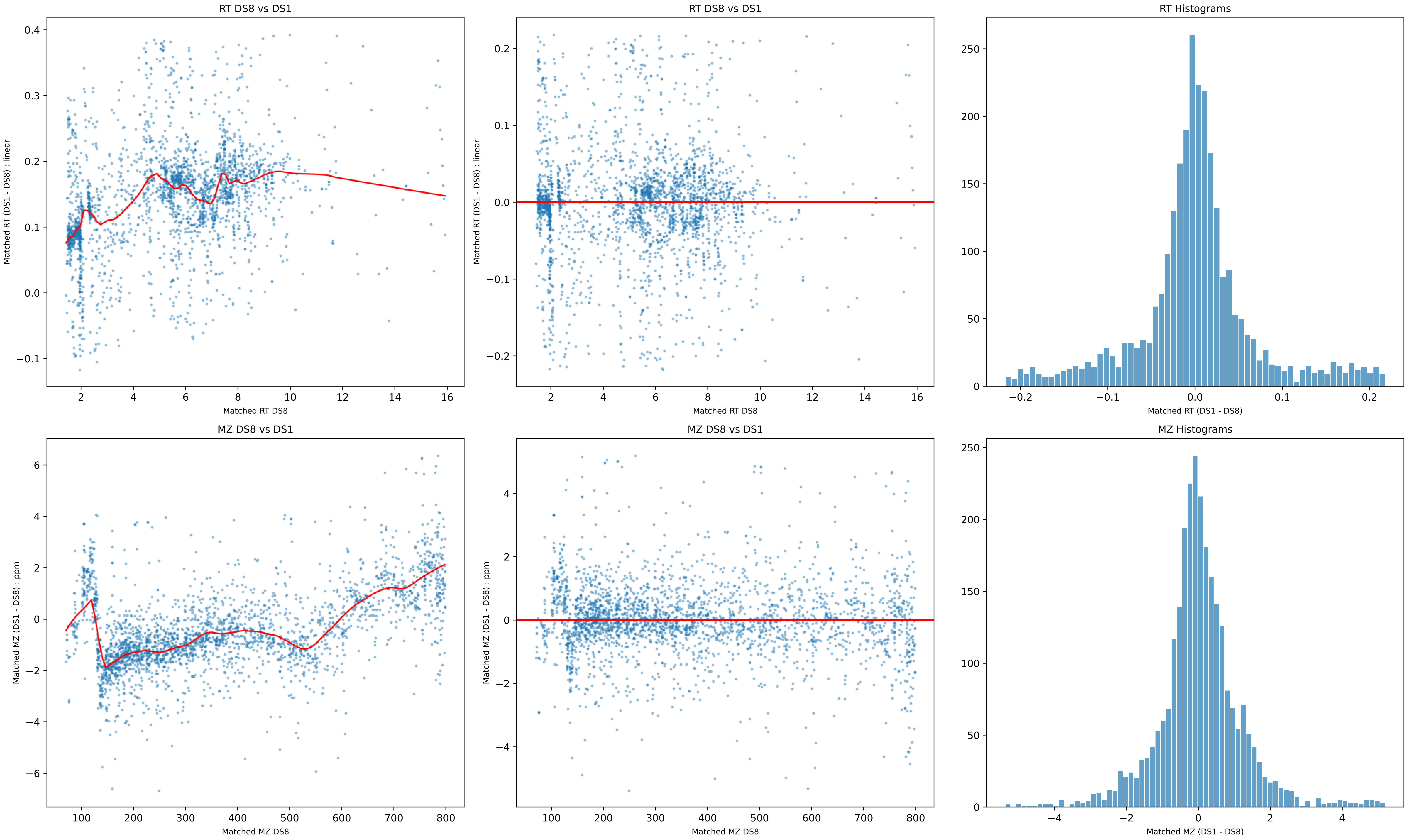

RT DS8 vs DS1
Matched RT (DS1 - DS8) : linear
Matched RT DS8
RT DS8 vs DS1
Matched RT (DS1 - DS8) : linear
Matched RT DS8
RT Histograms
Matched RT (DS1 - DS8)
MZ DS8 vs DS1
Matched MZ (DS1 - DS8) : ppm
Matched MZ DS8
MZ DS8 vs DS1
Matched MZ (DS1 - DS8) : ppm
Matched MZ DS8
MZ Histograms
Matched MZ (DS1 - DS8)

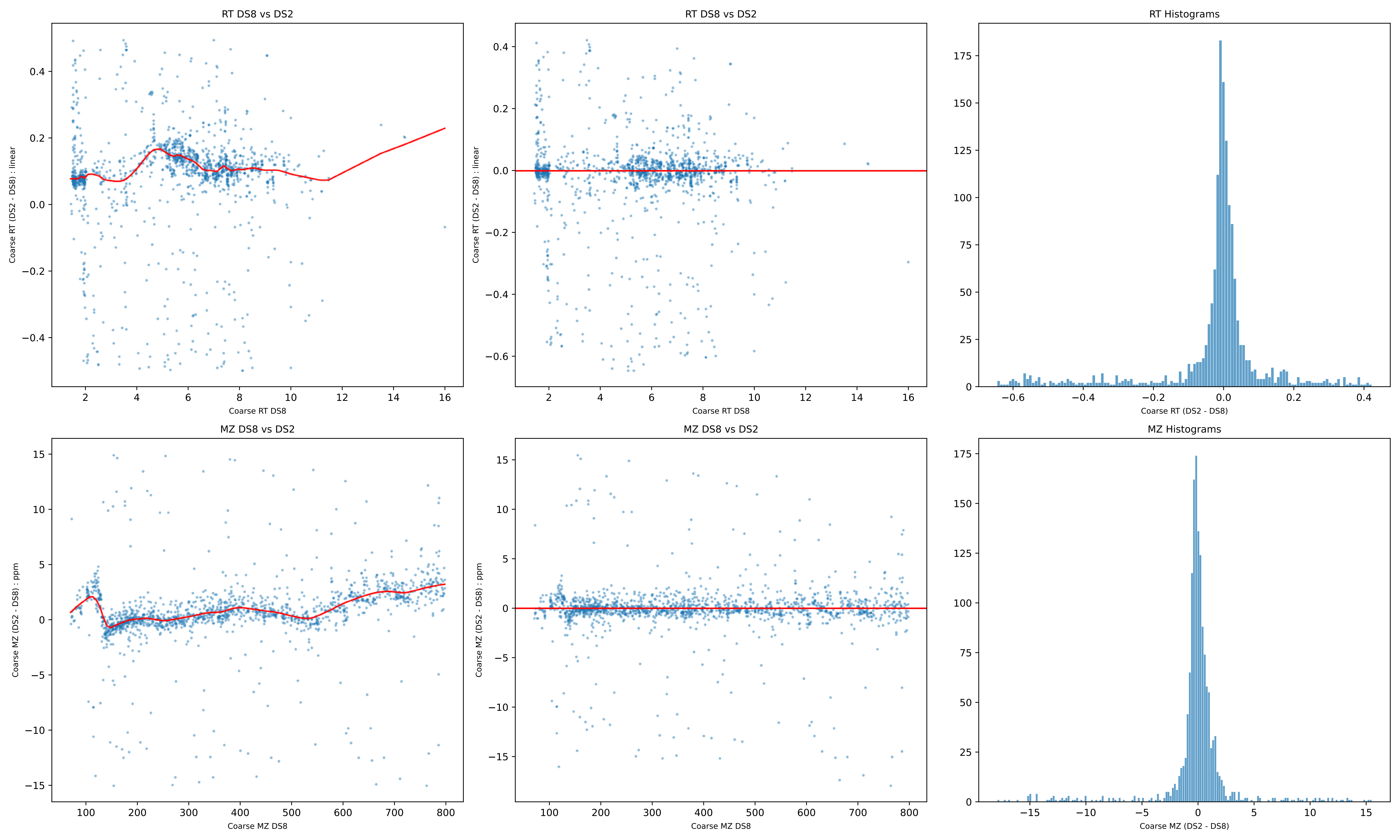
RT DS8 vs DS2
Coarse RT (DS2 - DS8) : linear
Coarse RT DS8
RT DS8 vs DS2
Coarse RT (DS2 - DS8) : linear
Coarse RT DS8
RT Histograms
Coarse RT (DS2 - DS8)
MZ DS8 vs DS2
Coarse MZ (DS2 - DS8) : ppm
Coarse MZ DS8
MZ DS8 vs DS2
Coarse MZ (DS2 - DS8) : ppm
Coarse MZ DS8
MZ Histograms
Coarse MZ (DS2 - DS8)

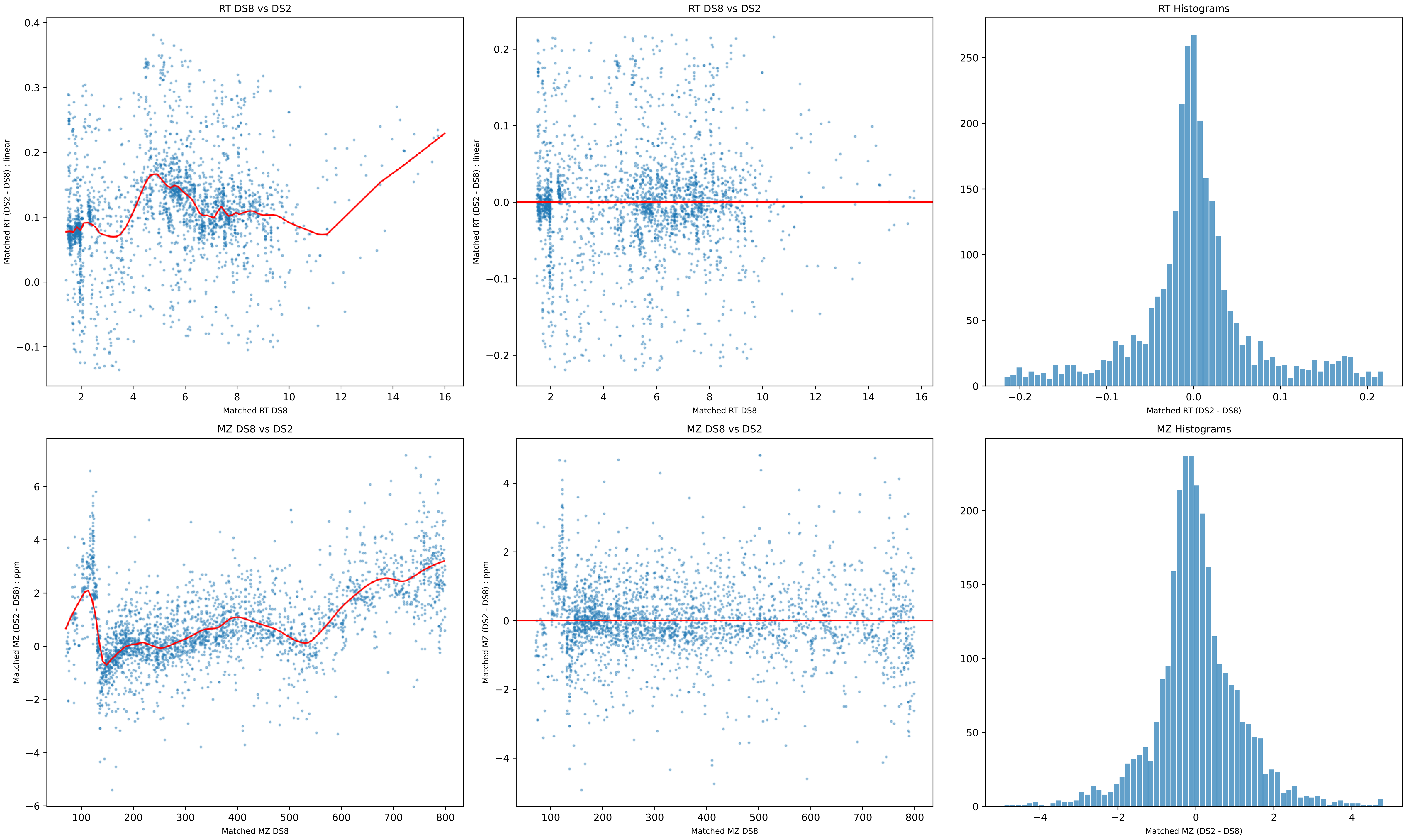
RT DS8 vs DS2
Matched RT (DS2 - DS8) : linear
Matched RT DS8
RT DS8 vs DS2
Matched RT (DS2 - DS8) : linear
Matched RT DS8
RT Histograms
Matched RT (DS2 - DS8)
MZ DS8 vs DS2
Matched MZ (DS2 - DS8) : ppm
Matched MZ DS8
MZ DS8 vs DS2
Matched MZ (DS2 - DS8) : ppm
Matched MZ DS8
MZ Histograms
Matched MZ (DS2 - DS8)

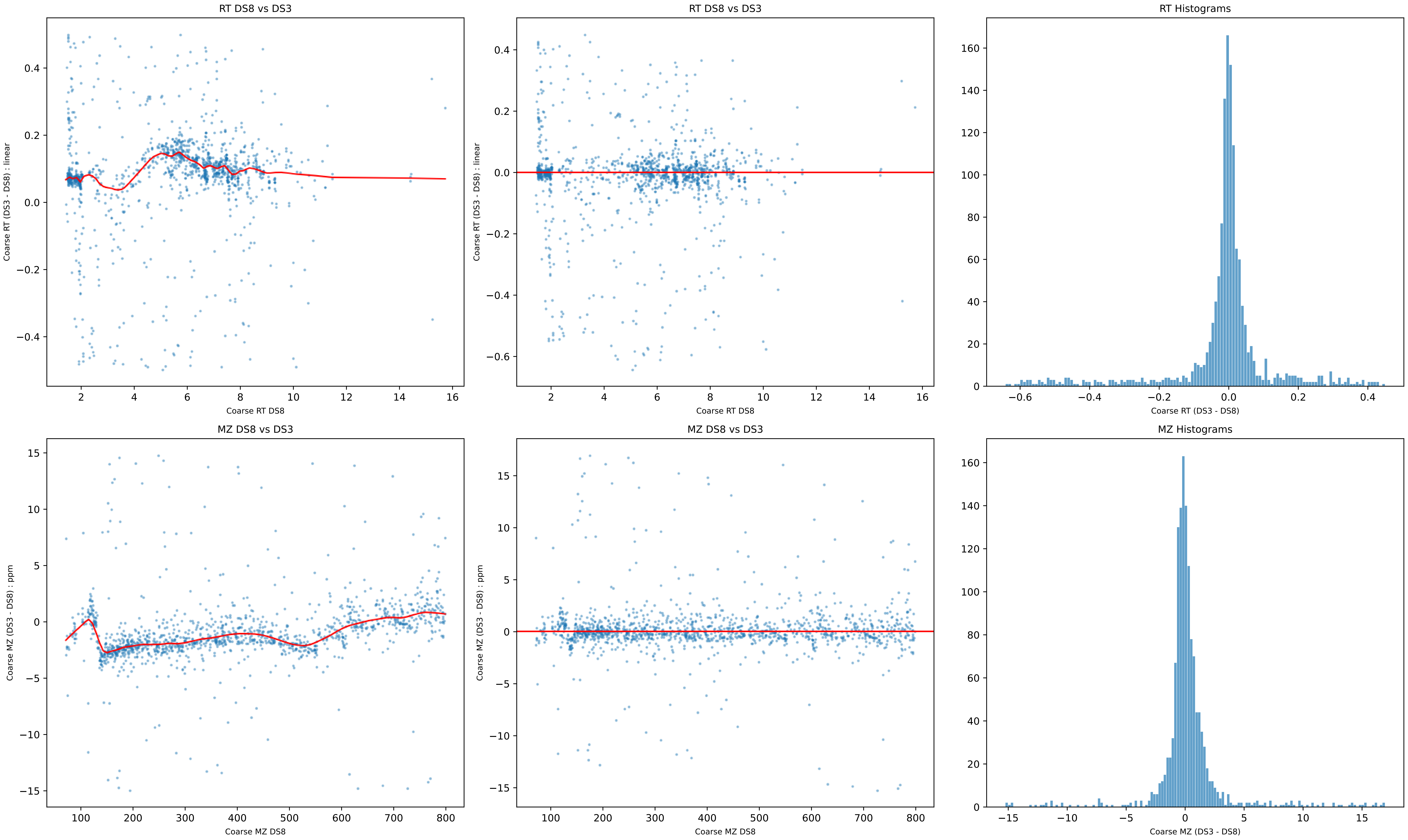
RT DS8 vs DS3
Coarse RT (DS3 - DS8) : linear
Coarse RT DS8
RT DS8 vs DS3
Coarse RT (DS3 - DS8) : linear
Coarse RT DS8
RT Histograms
Coarse RT (DS3 - DS8)
MZ DS8 vs DS3
Coarse MZ (DS3 - DS8) : ppm
Coarse MZ DS8
MZ DS8 vs DS3
Coarse MZ (DS3 - DS8) : ppm
Coarse MZ DS8
MZ Histograms
Coarse MZ (DS3 - DS8)

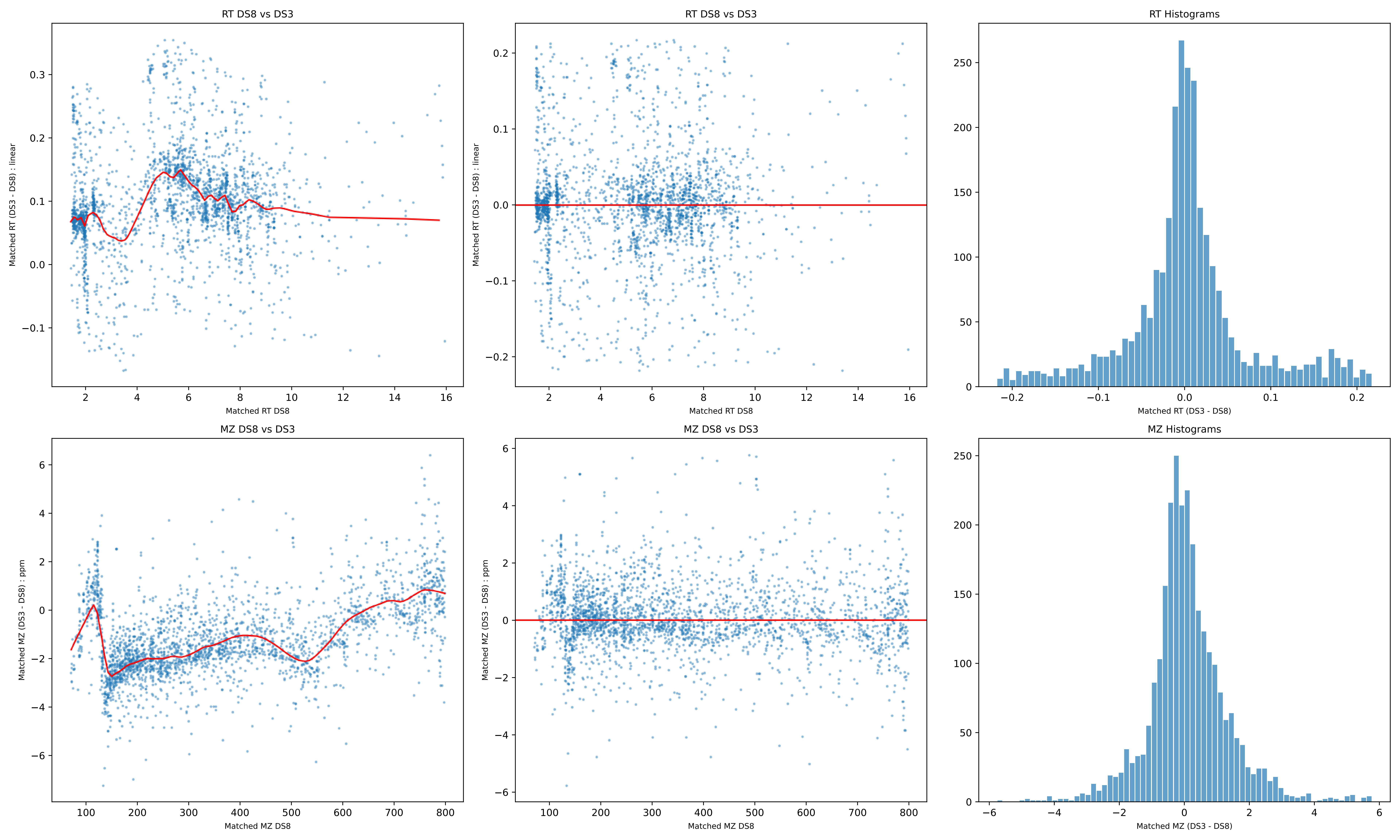
RT DS8 vs DS3
Matched RT (DS3 - DS8) : linear
Matched RT DS8
RT DS8 vs DS3
Matched RT (DS3 - DS8) : linear
Matched RT DS8
RT Histograms
Matched RT (DS3 - DS8)
MZ DS8 vs DS3
Matched MZ (DS3 - DS8) : ppm
Matched MZ DS8
MZ DS8 vs DS3
Matched MZ (DS3 - DS8) : ppm
Matched MZ DS8
MZ Histograms
Matched MZ (DS3 - DS8)

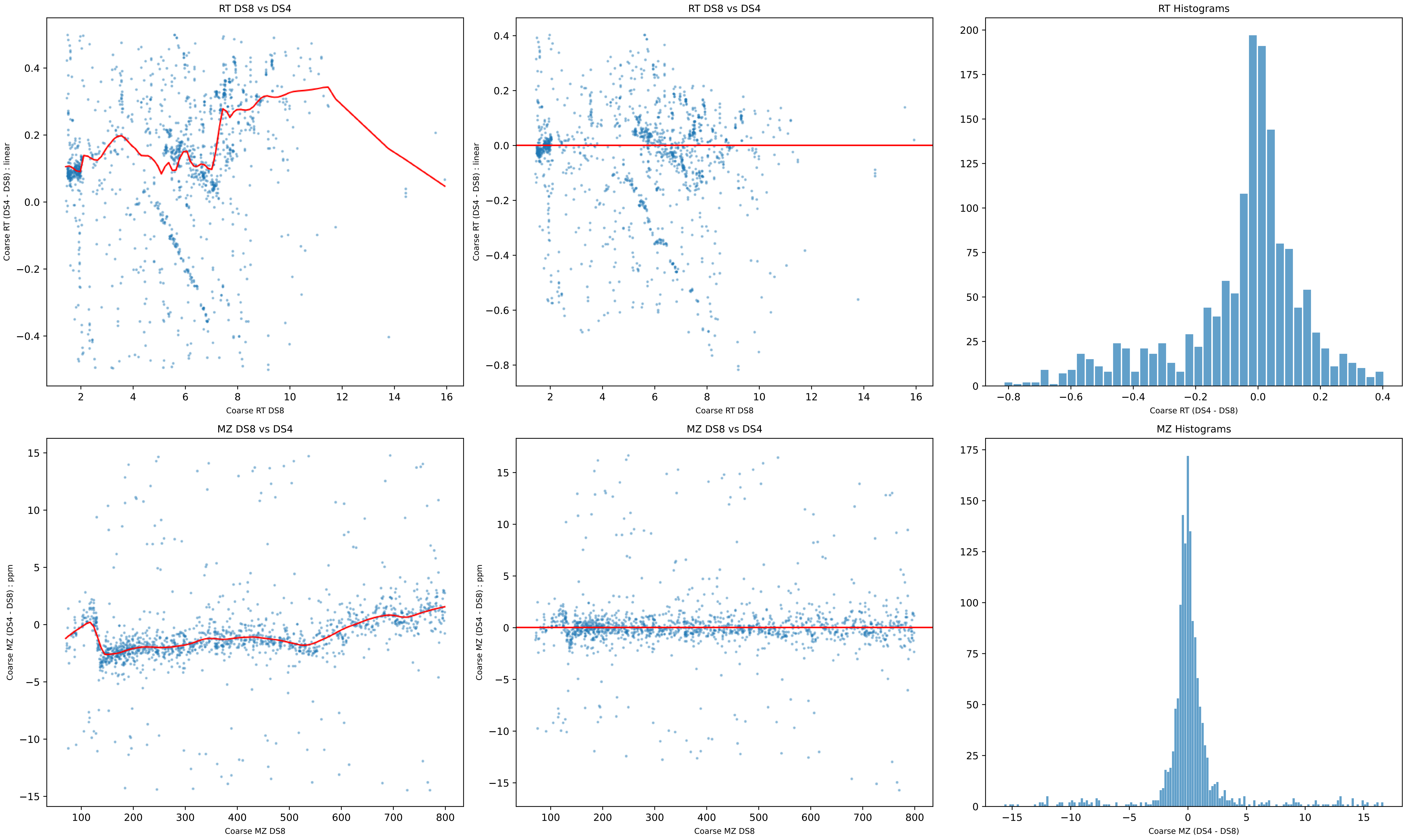
RT DS8 vs DS4
Coarse RT (DS4 - DS8) : linear
Coarse RT DS8
RT DS8 vs DS4
Coarse RT (DS4 - DS8) : linear
Coarse RT DS8
RT Histograms
Coarse RT (DS4 - DS8)
MZ DS8 vs DS4
Coarse MZ (DS4 - DS8) : ppm
Coarse MZ DS8
MZ DS8 vs DS4
Coarse MZ (DS4 - DS8) : ppm
Coarse MZ DS8
MZ Histograms
Coarse MZ (DS4 - DS8)

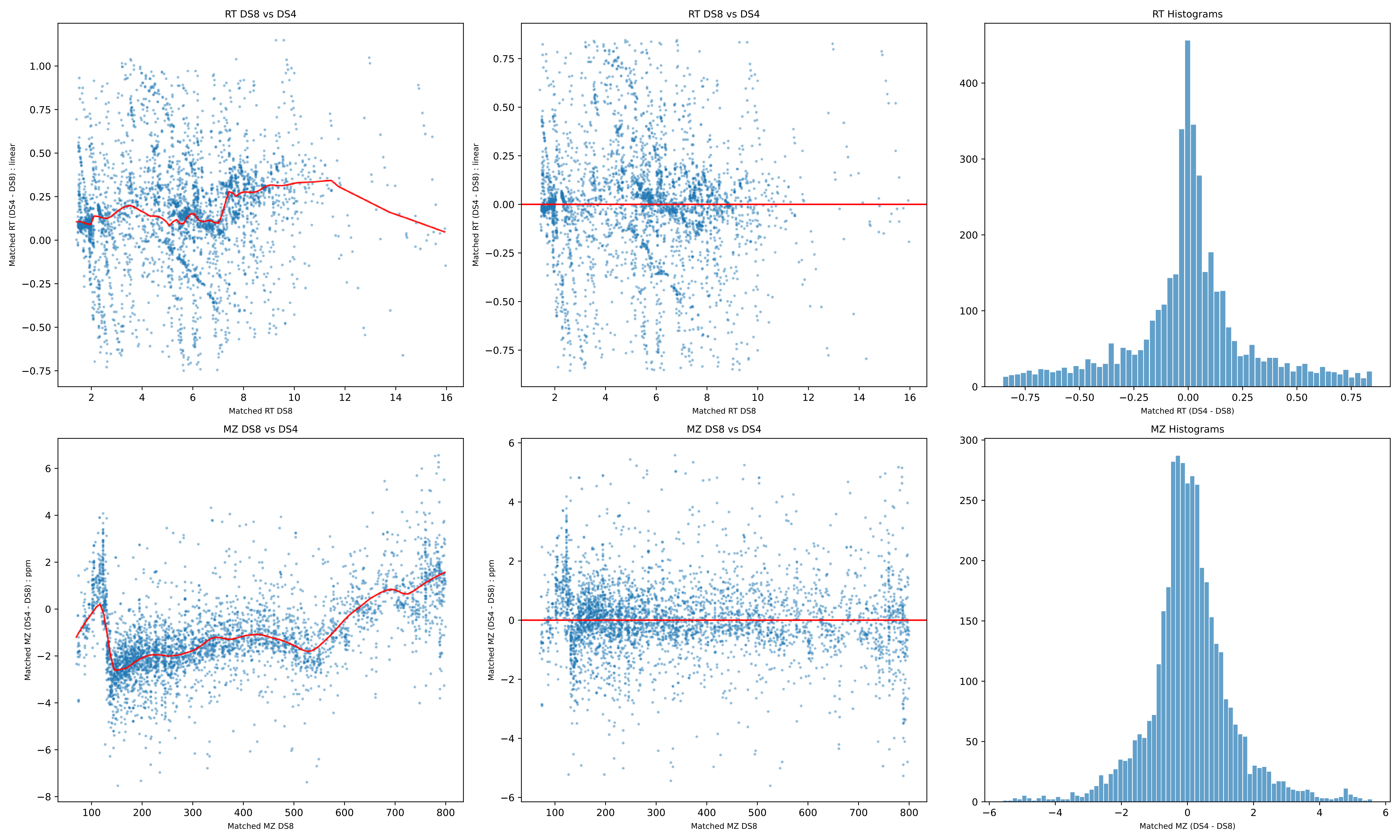

RT DS8 vs DS4
Matched RT (DS4 - DS8) : linear
Matched RT DS8
RT DS8 vs DS4
Matched RT (DS4 - DS8) : linear
Matched RT DS8
RT Histograms
Matched RT (DS4 - DS8)
MZ DS8 vs DS4
Matched MZ (DS4 - DS8) : ppm
Matched MZ DS8
MZ DS8 vs DS4
Matched MZ (DS4 - DS8) : ppm
Matched MZ DS8
MZ Histograms
Matched MZ (DS4 - DS8)

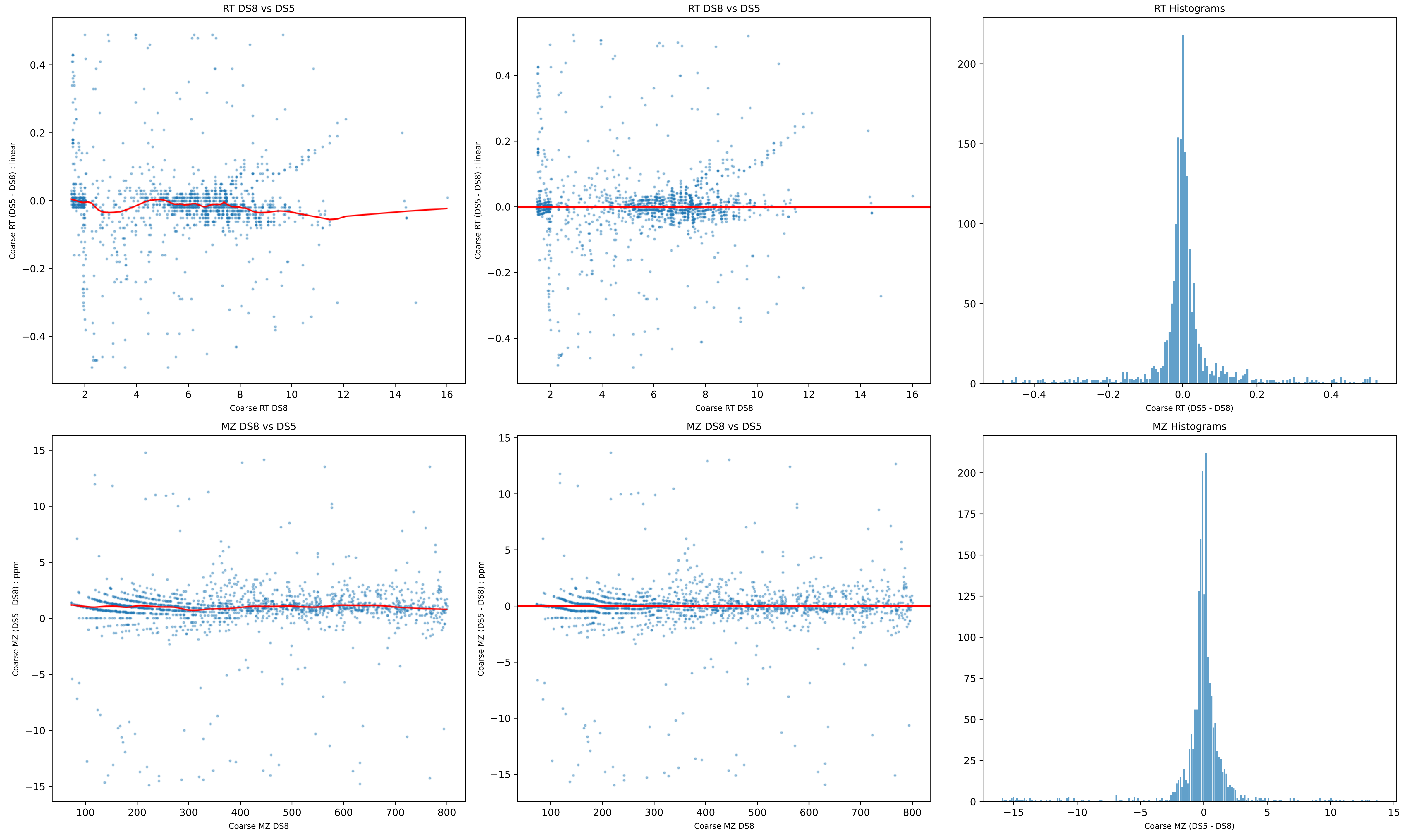
RT DS8 vs DS5
Coarse RT (DS5 - DS8) : linear
Coarse RT DS8
RT DS8 vs DS5
Coarse RT (DS5 - DS8) : linear
Coarse RT DS8
RT Histograms
Coarse RT (DS5 - DS8)
MZ DS8 vs DS5
Coarse MZ (DS5 - DS8) : ppm
Coarse MZ DS8
MZ DS8 vs DS5
Coarse MZ (DS5 - DS8) : ppm
Coarse MZ DS8
MZ Histograms
Coarse MZ (DS5 - DS8)

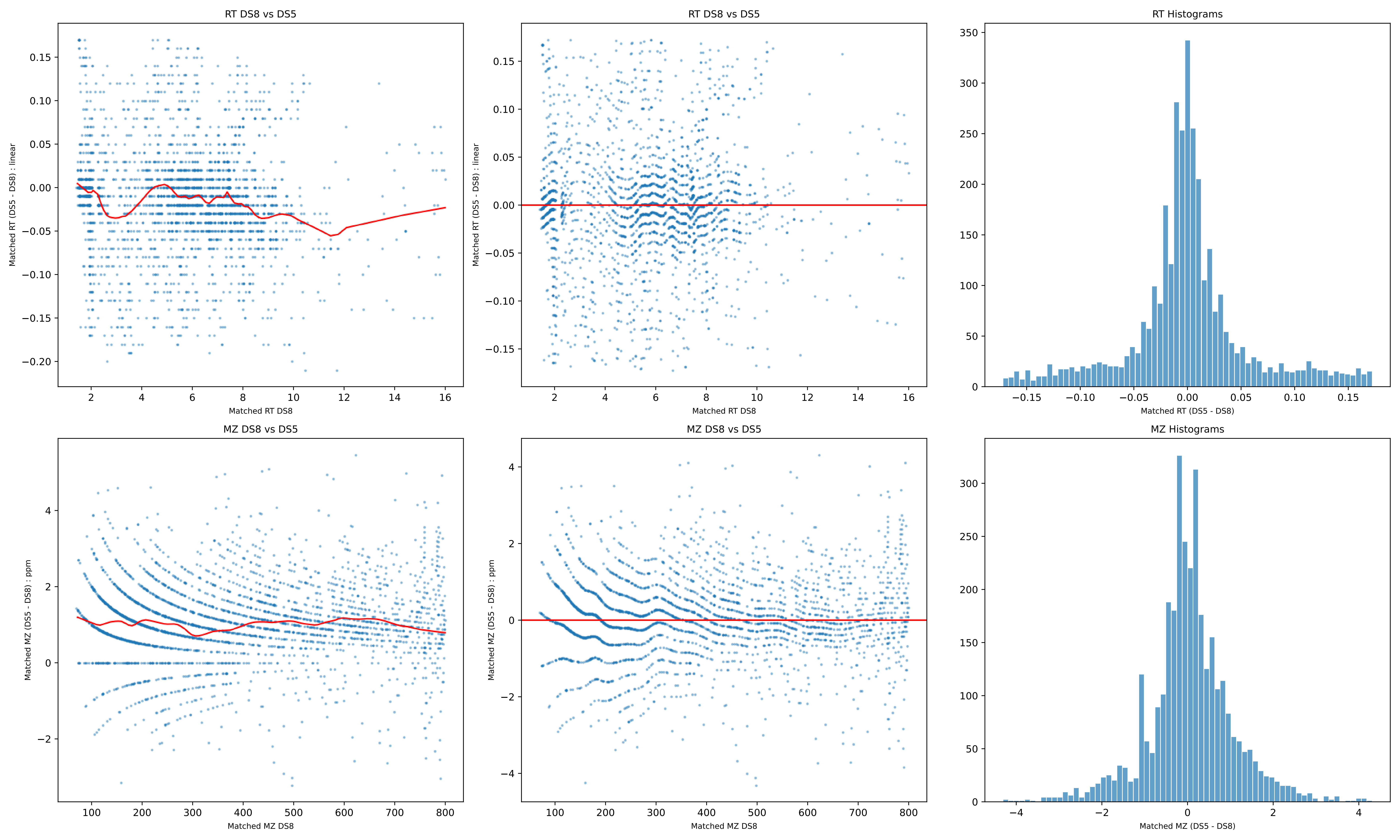

RT DS8 vs DS5
Matched RT (DS5 - DS8) : linear
Matched RT DS8
RT DS8 vs DS5
Matched RT (DS5 - DS8) : linear
Matched RT DS8
RT Histograms
Matched RT (DS5 - DS8)
MZ DS8 vs DS5
Matched MZ (DS5 - DS8) : ppm
Matched MZ DS8
MZ DS8 vs DS5
Matched MZ (DS5 - DS8) : ppm
Matched MZ DS8
MZ Histograms
Matched MZ (DS5 - DS8)

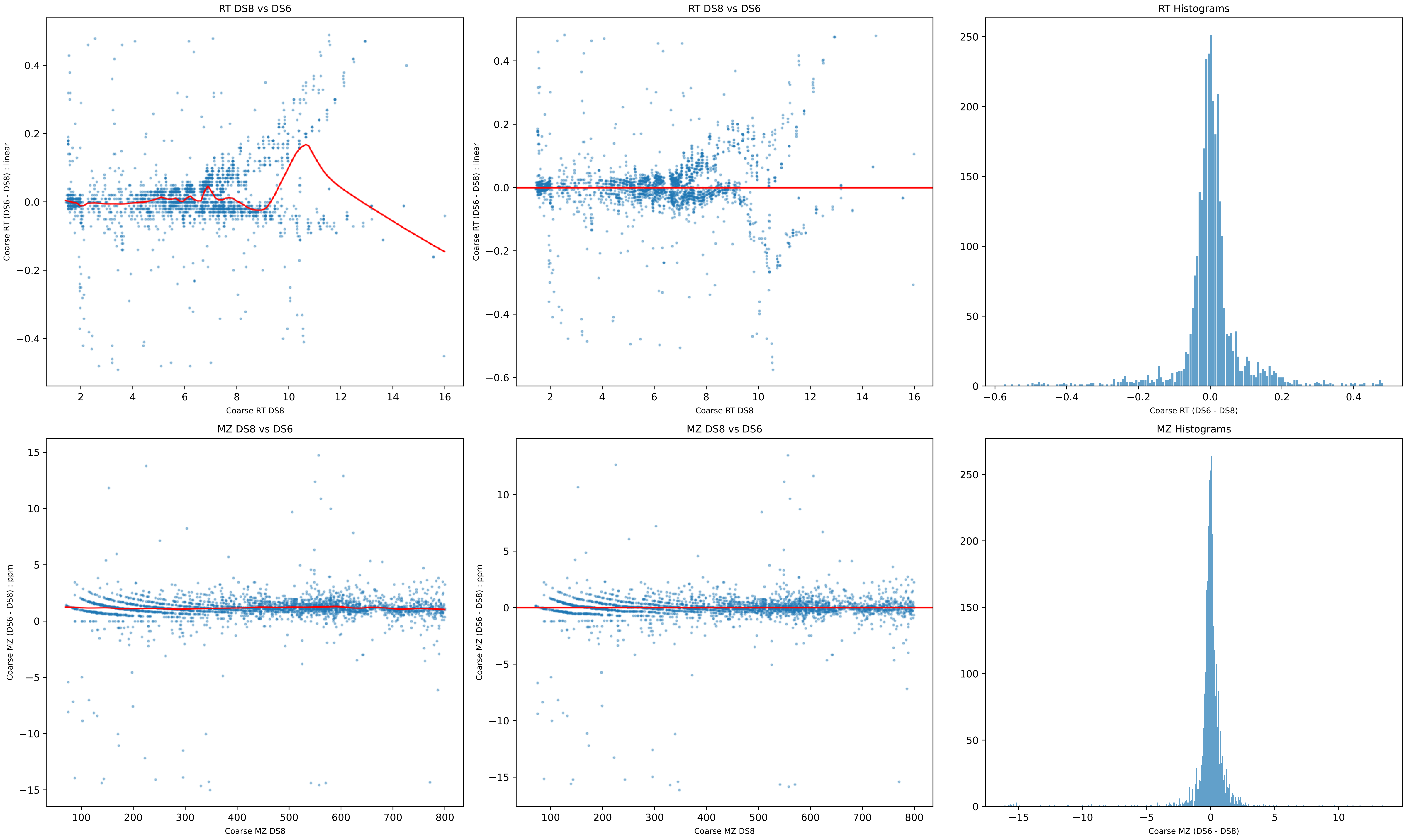

RT DS8 vs DS6
Coarse RT (DS6 - DS8) : linear
Coarse RT DS8
RT DS8 vs DS6
Coarse RT (DS6 - DS8) : linear
Coarse RT DS8
RT Histograms
Coarse RT (DS6 - DS8)
MZ DS8 vs DS6
Coarse MZ (DS6 - DS8) : ppm
Coarse MZ DS8
MZ DS8 vs DS6
Coarse MZ (DS6 - DS8) : ppm
Coarse MZ DS8
MZ Histograms
Coarse MZ (DS6 - DS8)

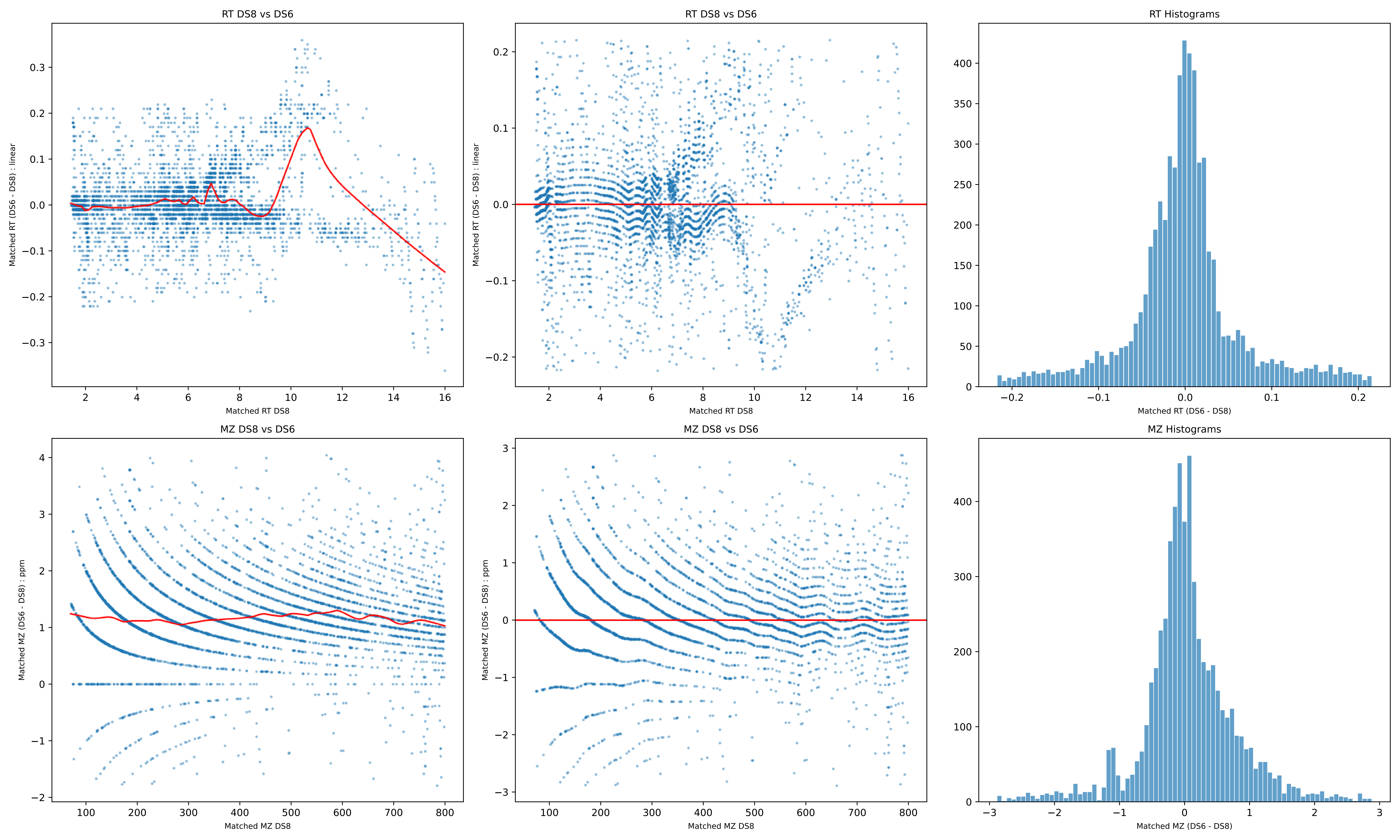
RT DS8 vs DS6
Matched RT (DS6 - DS8) : linear
Matched RT DS8
RT DS8 vs DS6
Matched RT (DS6 - DS8) : linear
Matched RT DS8
RT Histograms
Matched RT (DS6 - DS8)
MZ DS8 vs DS6
Matched MZ (DS6 - DS8) : ppm
Matched MZ DS8
MZ DS8 vs DS6
Matched MZ (DS6 - DS8) : ppm
Matched MZ DS8
MZ Histograms
Matched MZ (DS6 - DS8)

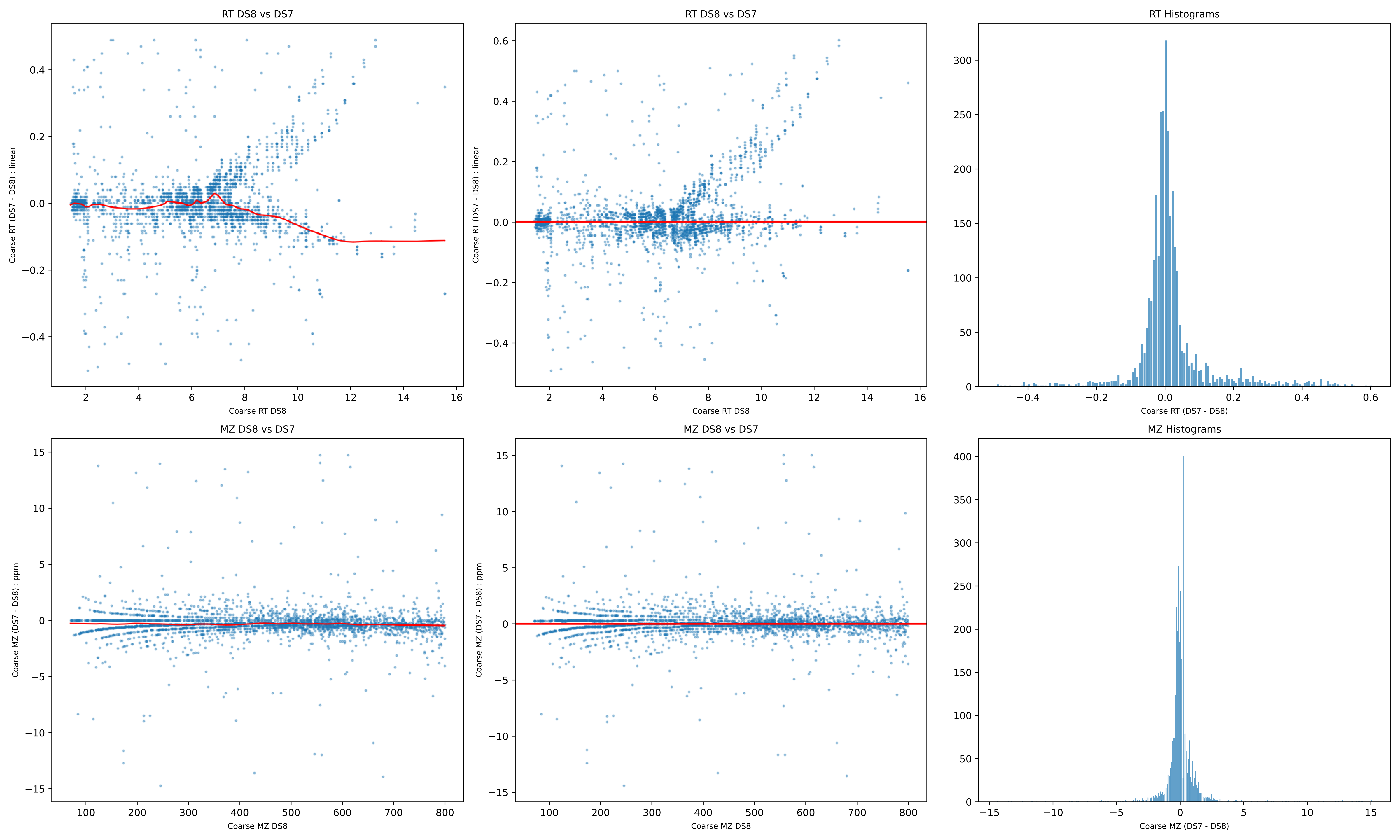
RT DS8 vs DS7
Coarse RT (DS7 - DS8) : linear
Coarse RT DS8
RT Histograms
Coarse RT (DS7 - DS8)
MZ DS8 vs DS7
Coarse MZ (DS7 - DS8) : ppm
Coarse MZ DS8
MZ Histograms
Coarse MZ (DS7 - DS8)

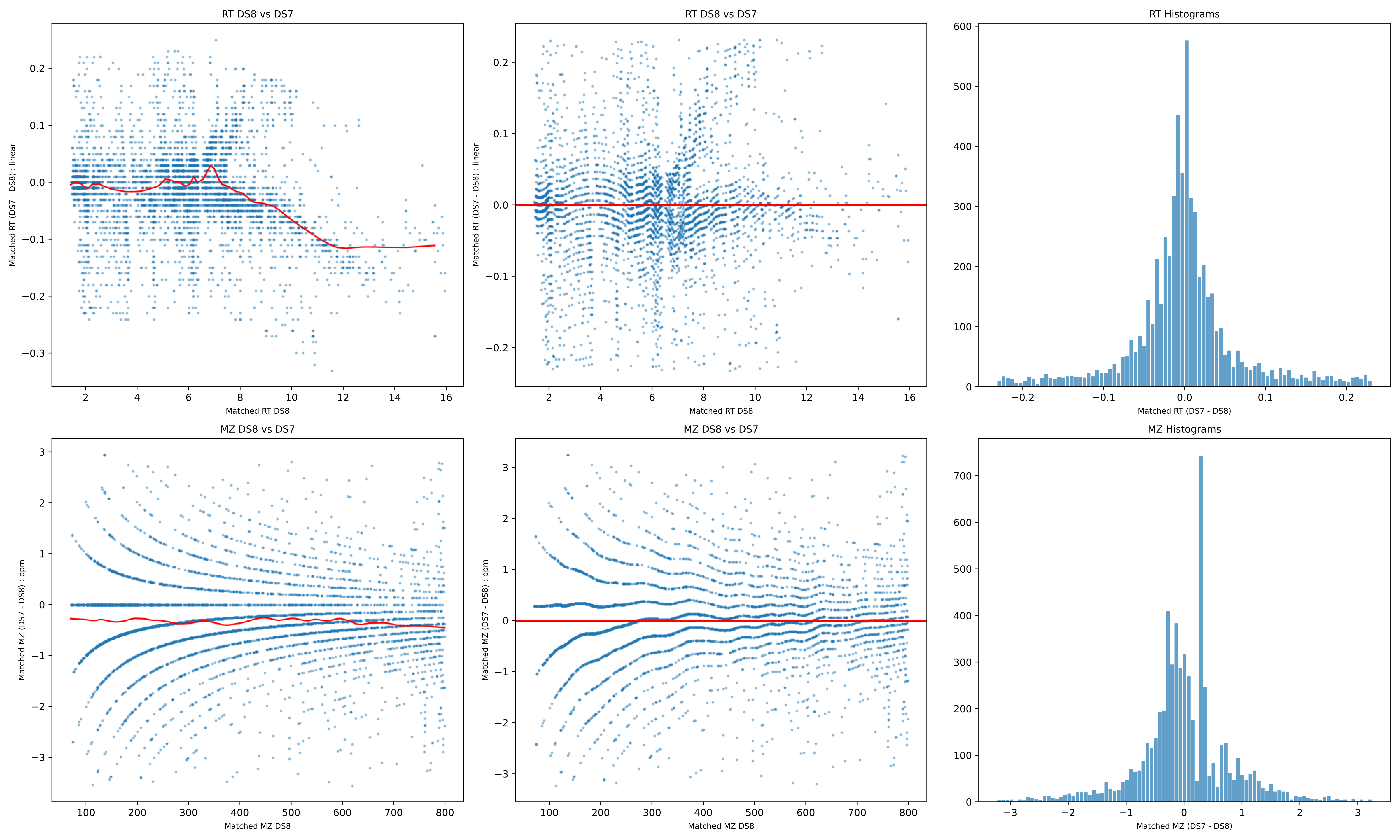
RT DS8 vs DS7
Matched RT (DS7 - DS8) : linear
Matched RT DS8
RT DS8 vs DS7
Matched RT (DS7 - DS8) : linear
Matched RT DS8
RT Histograms
Matched RT (DS7 - DS8)
MZ DS8 vs DS7
Matched MZ (DS7 - DS8) : ppm
Matched MZ DS8
MZ DS8 vs DS7
Matched MZ (DS7 - DS8) : ppm
Matched MZ DS8
MZ Histograms
Matched MZ (DS7 - DS8)

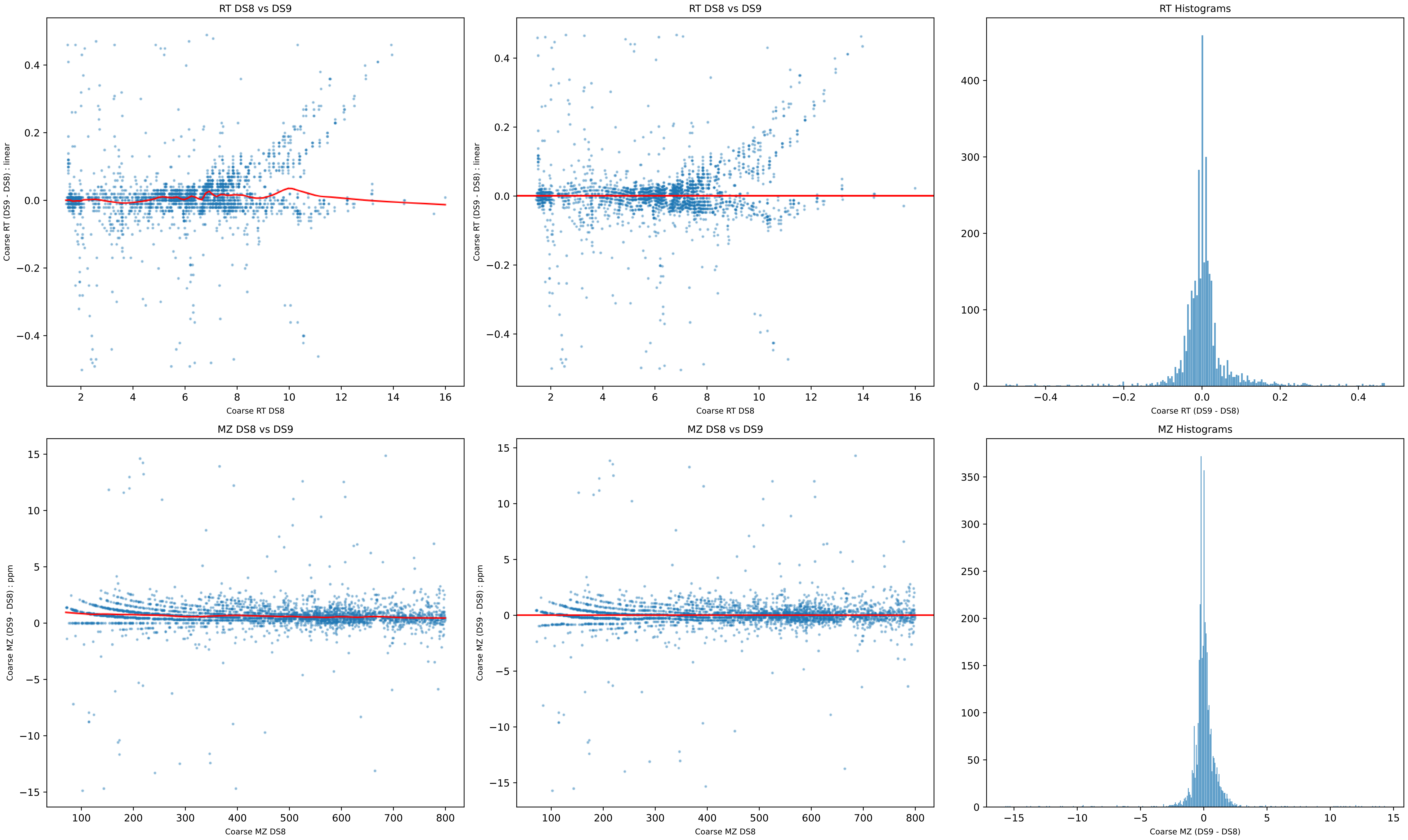
RT DS8 vs DS9
Coarse RT (DS9 - DS8) : linear
Coarse RT DS8
RT DS8 vs DS9
Coarse RT (DS9 - DS8) : linear
Coarse RT DS8
RT Histograms
Coarse RT (DS9 - DS8)
MZ DS8 vs DS9
Coarse MZ (DS9 - DS8) : ppm
Coarse MZ DS8
MZ DS8 vs DS9
Coarse MZ (DS9 - DS8) : ppm
Coarse MZ DS8
MZ Histograms
Coarse MZ (DS9 - DS8)

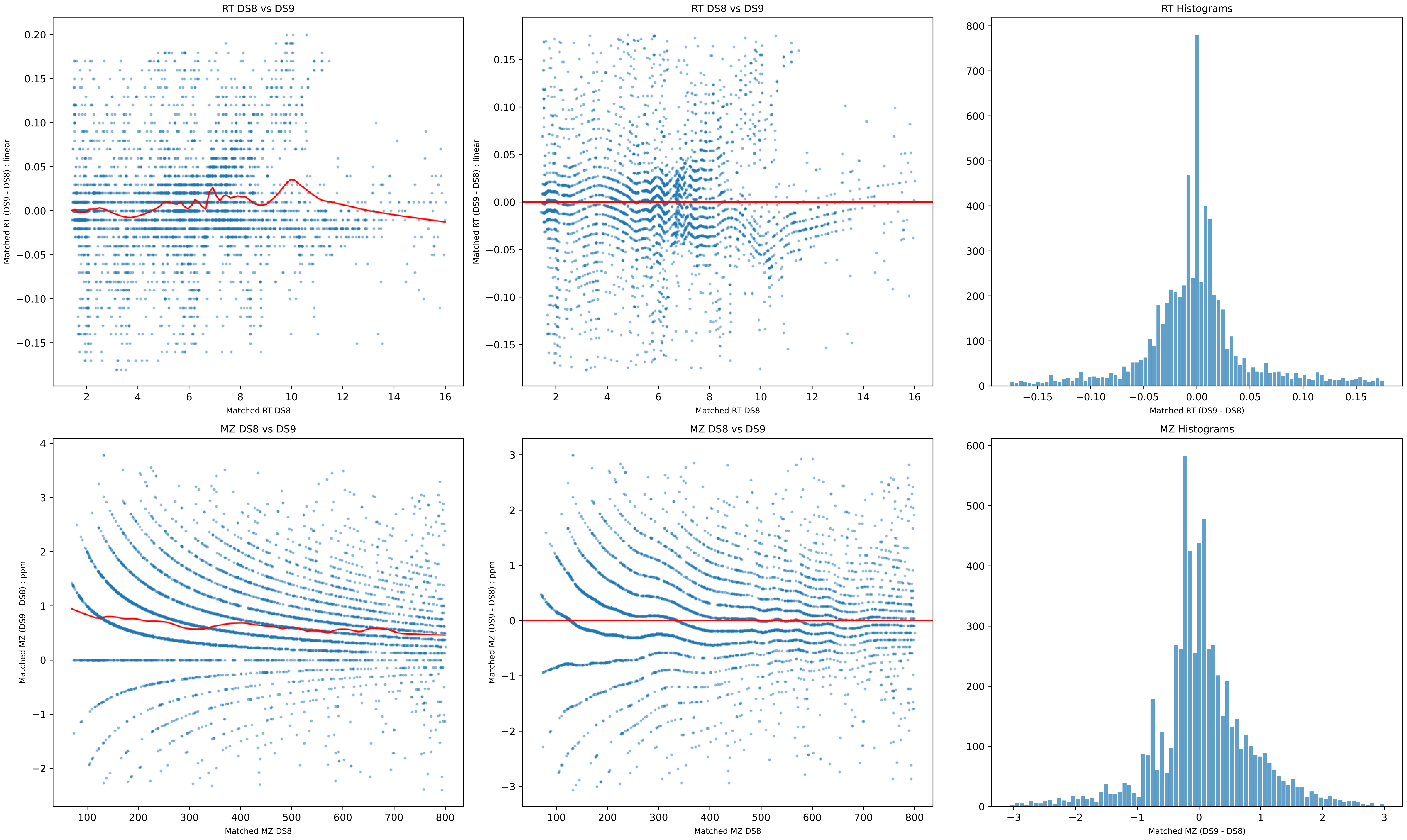
RT DS8 vs DS9
Matched RT (DS9 - DS8) : linear
Matched RT DS8
RT DS8 vs DS9
Matched RT (DS9 - DS8) : linear
Matched RT DS8
RT Histograms
Matched RT (DS9 - DS8)
MZ DS8 vs DS9
Matched MZ (DS9 - DS8) : ppm
Matched MZ DS8
MZ DS8 vs DS9
Matched MZ (DS9 - DS8) : ppm
Matched MZ DS8
MZ Histograms
Matched MZ (DS9 - DS8)

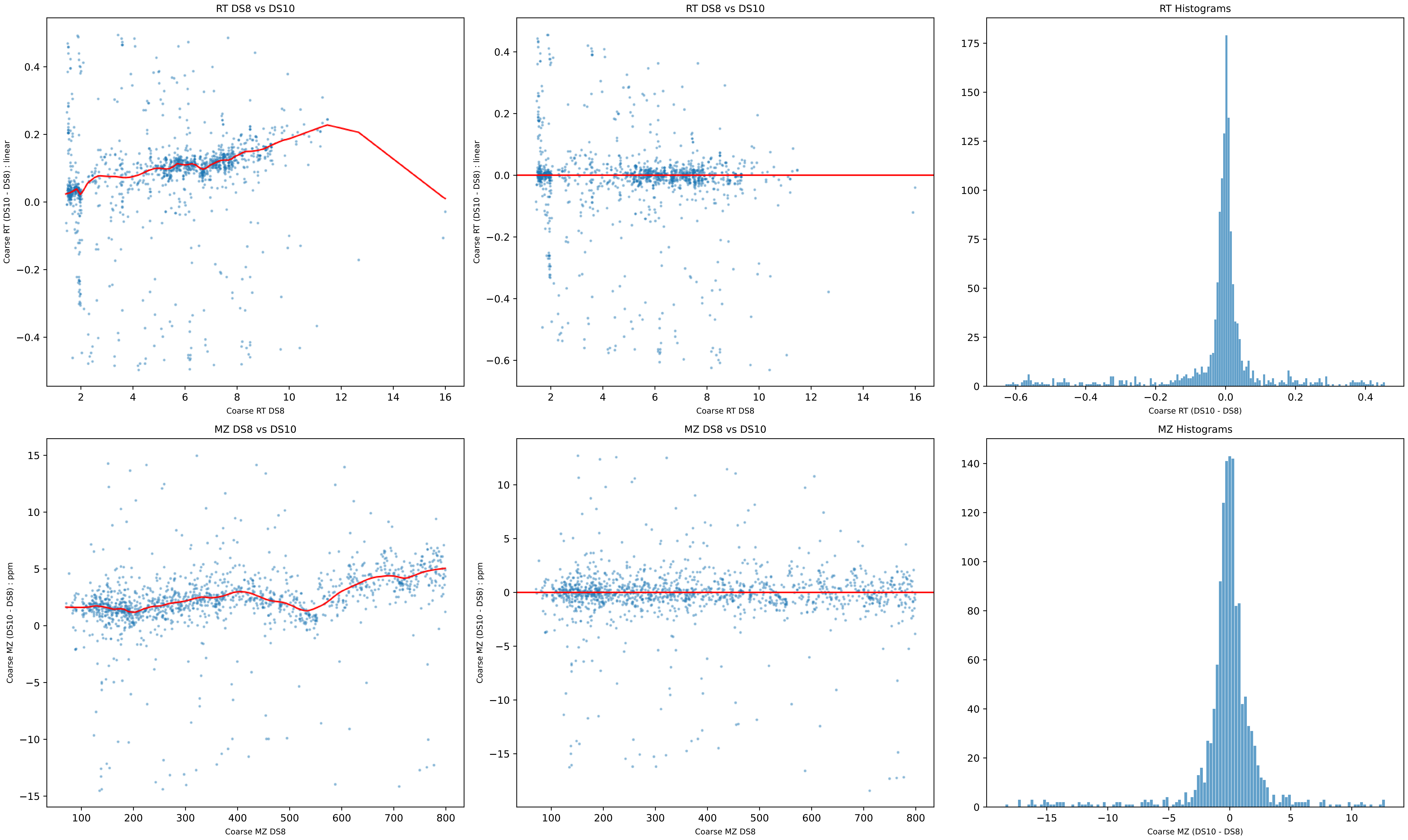
RT DS8 vs DS10
Coarse RT (DS10 - DS8) : linear
Coarse RT DS8
RT DS8 vs DS10
Coarse RT (DS10 - DS8) : linear
Coarse RT DS8
RT Histograms
Coarse RT (DS10 - DS8)
MZ DS8 vs DS10
Coarse MZ (DS10 - DS8) : ppm
Coarse MZ DS8
MZ DS8 vs DS10
Coarse MZ (DS10 - DS8) : ppm
Coarse MZ DS8
MZ Histograms
Coarse MZ (DS10 - DS8)

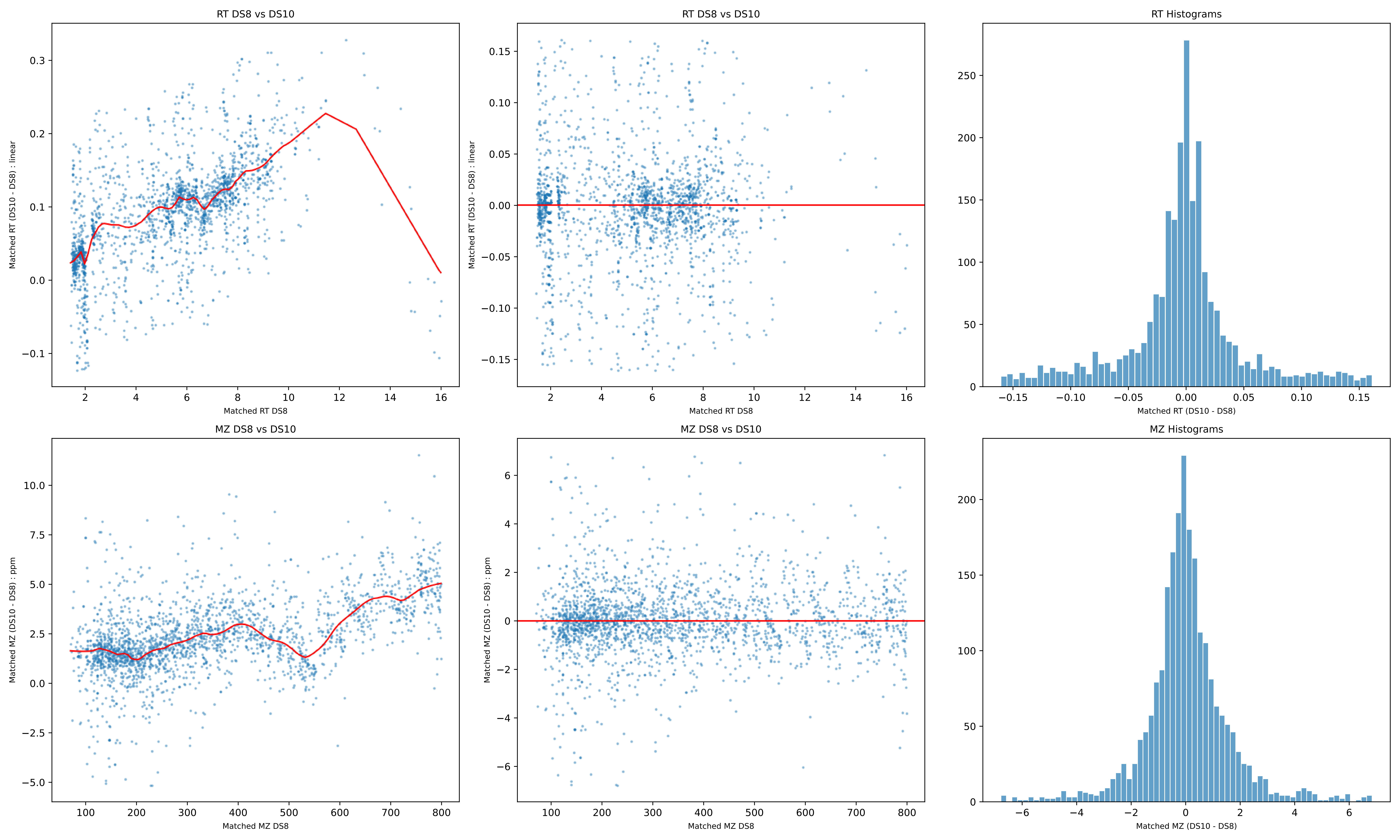
RT DS8 vs DS10
Matched RT (DS10 - DS8) : linear
Matched RT DS8
RT DS8 vs DS10
Matched RT (DS10 - DS8) : linear
Matched RT DS8
RT Histograms
Matched RT (DS10 - DS8)
MZ DS8 vs DS10
Matched MZ (DS10 - DS8) : ppm
Matched MZ DS8
MZ DS8 vs DS10
Matched MZ (DS10 - DS8) : ppm
Matched MZ DS8
MZ Histograms
Matched MZ (DS10 - DS8)

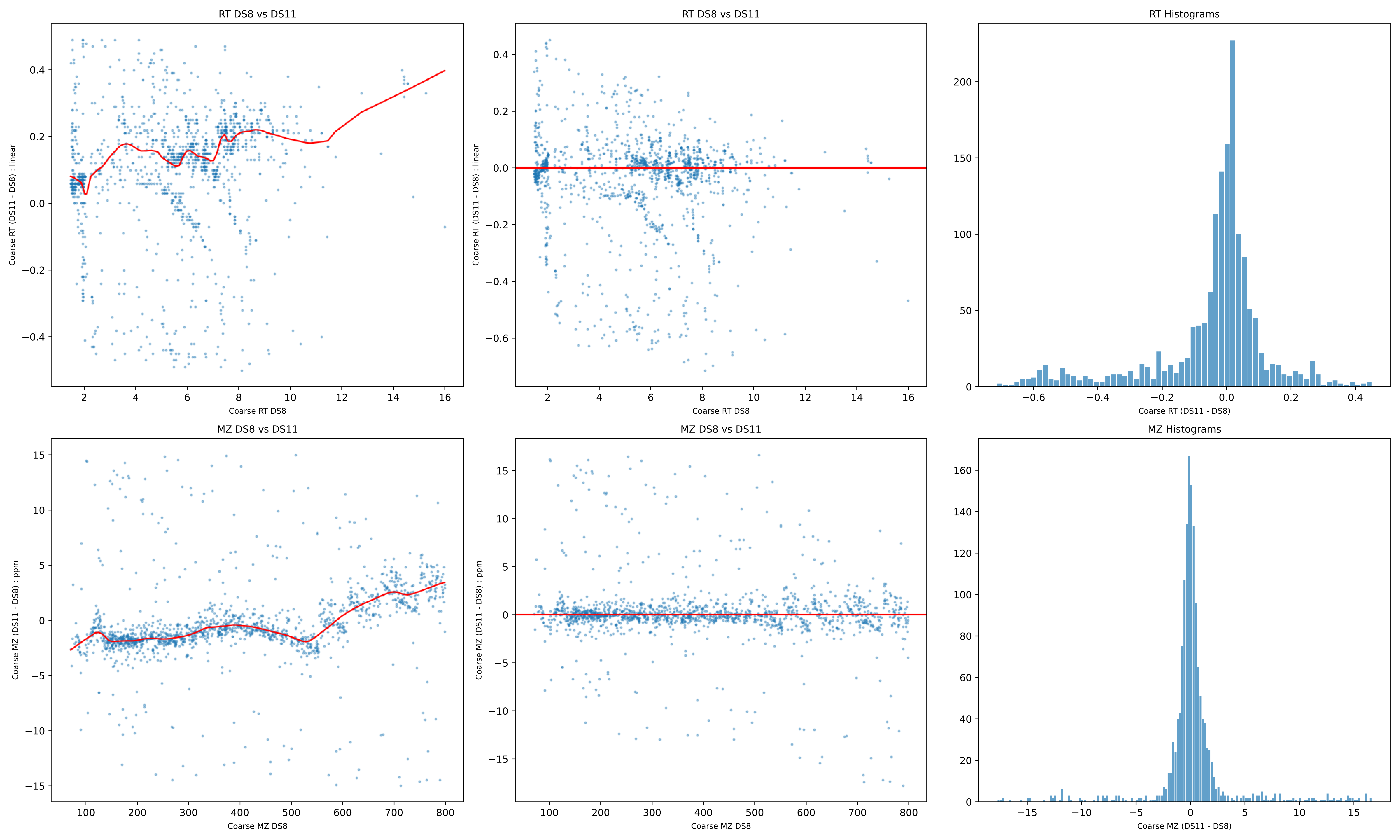
RT DS8 vs DS11
Coarse RT (DS11 - DS8) : linear
Coarse RT DS8
RT DS8 vs DS11
Coarse RT (DS11 - DS8) : linear
Coarse RT DS8
RT Histograms
Coarse RT (DS11 - DS8)
MZ DS8 vs DS11
Coarse MZ (DS11 - DS8) : ppm
Coarse MZ DS8
MZ DS8 vs DS11
Coarse MZ (DS11 - DS8) : ppm
Coarse MZ DS8
MZ Histograms
Coarse MZ (DS11 - DS8)

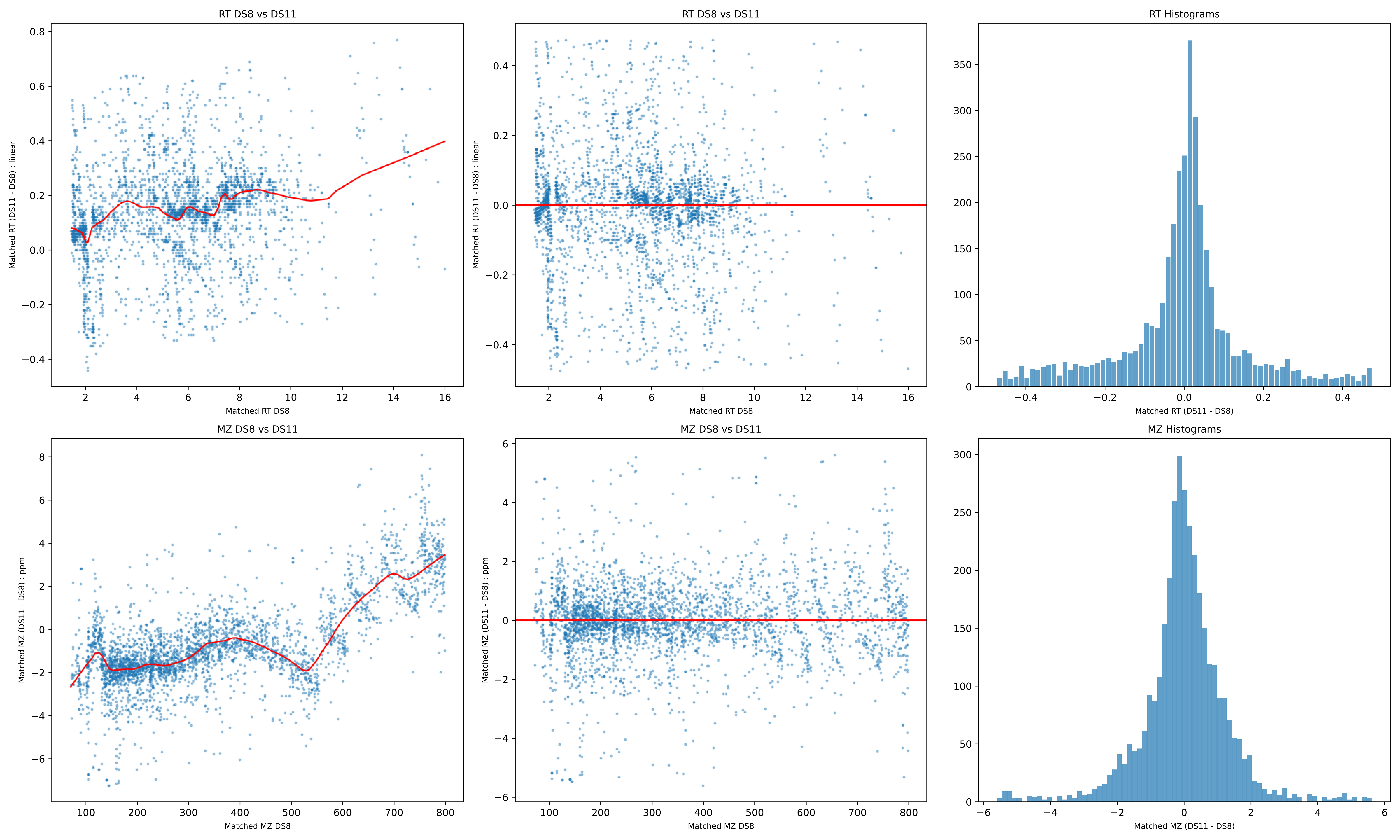

RT DS8 vs DS11
Matched RT (DS11 - DS8) : linear
Matched RT DS8
RT DS8 vs DS11
Matched RT (DS11 - DS8) : linear
Matched RT DS8
RT Histograms
Matched RT (DS11 - DS8)
MZ DS8 vs DS11
Matched MZ (DS11 - DS8) : ppm
Matched MZ DS8
MZ DS8 vs DS11
Matched MZ (DS11 - DS8) : ppm
Matched MZ DS8
MZ Histograms
Matched MZ (DS11 - DS8)

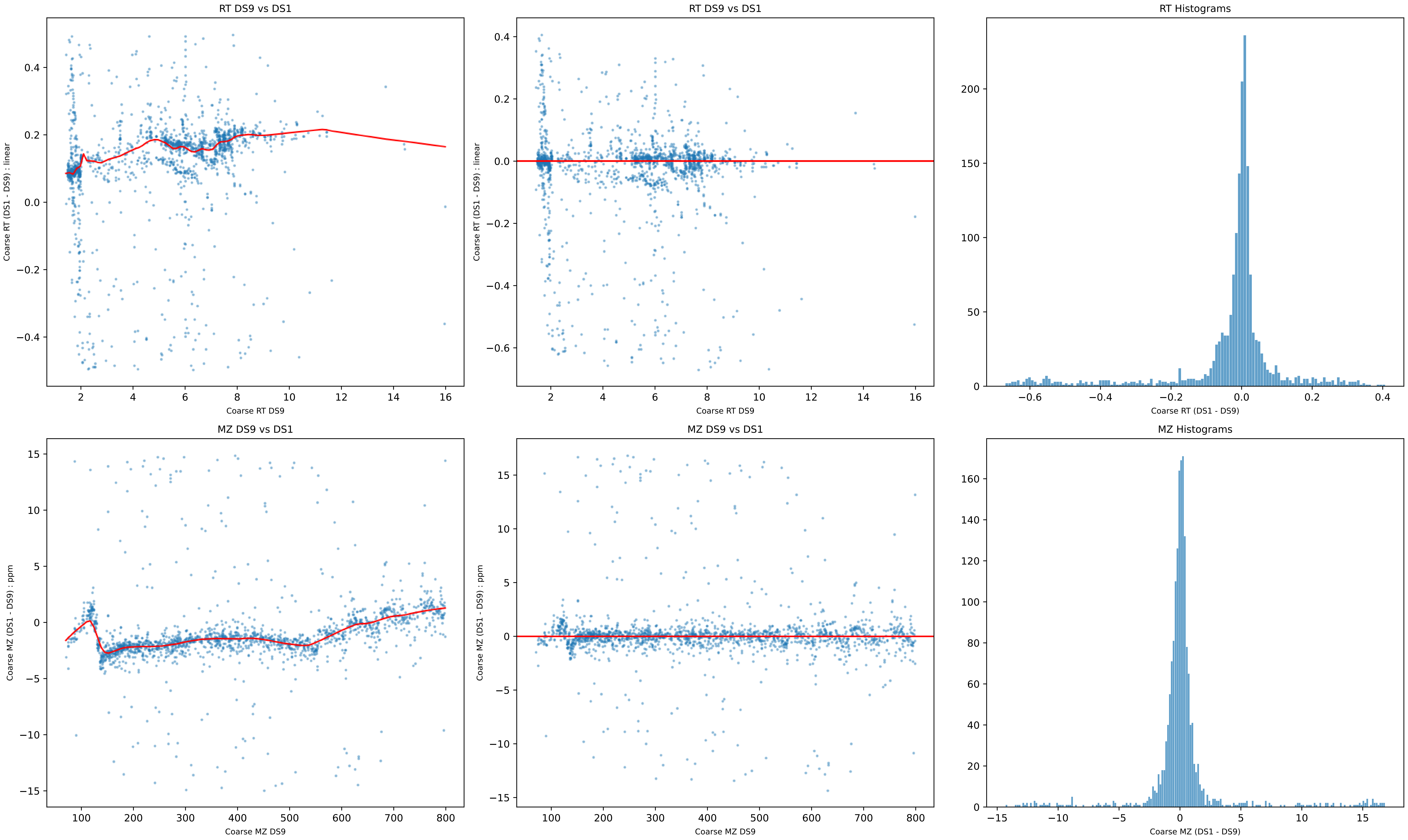
RT DS9 vs DS1
Coarse RT (DS1 - DS9) : linear
Coarse RT DS9
RT DS9 vs DS1
Coarse RT (DS1 - DS9) : linear
Coarse RT DS9
RT Histograms
Coarse RT (DS1 - DS9)
MZ DS9 vs DS1
Coarse MZ (DS1 - DS9) : ppm
Coarse MZ DS9
MZ DS9 vs DS1
Coarse MZ (DS1 - DS9) : ppm
Coarse MZ DS9
MZ Histograms
Coarse MZ (DS1 - DS9)

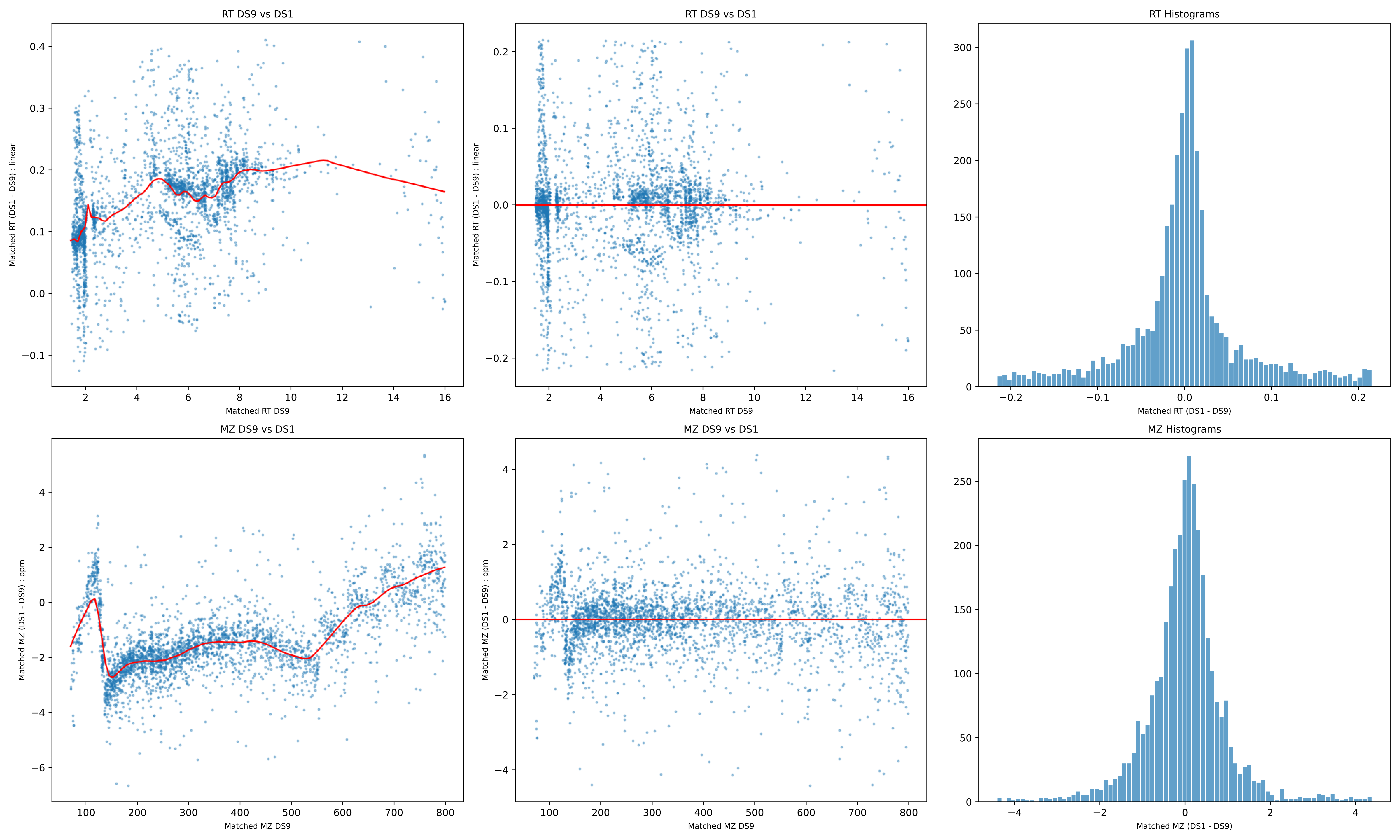

RT DS9 vs DS1
Matched RT (DS1 - DS9) : linear
Matched RT DS9
RT DS9 vs DS1
Matched RT (DS1 - DS9) : linear
Matched RT DS9
RT Histograms
Matched RT (DS1 - DS9)
MZ DS9 vs DS1
Matched MZ (DS1 - DS9) : ppm
Matched MZ DS9
MZ DS9 vs DS1
Matched MZ (DS1 - DS9) : ppm
Matched MZ DS9
MZ Histograms
Matched MZ (DS1 - DS9)

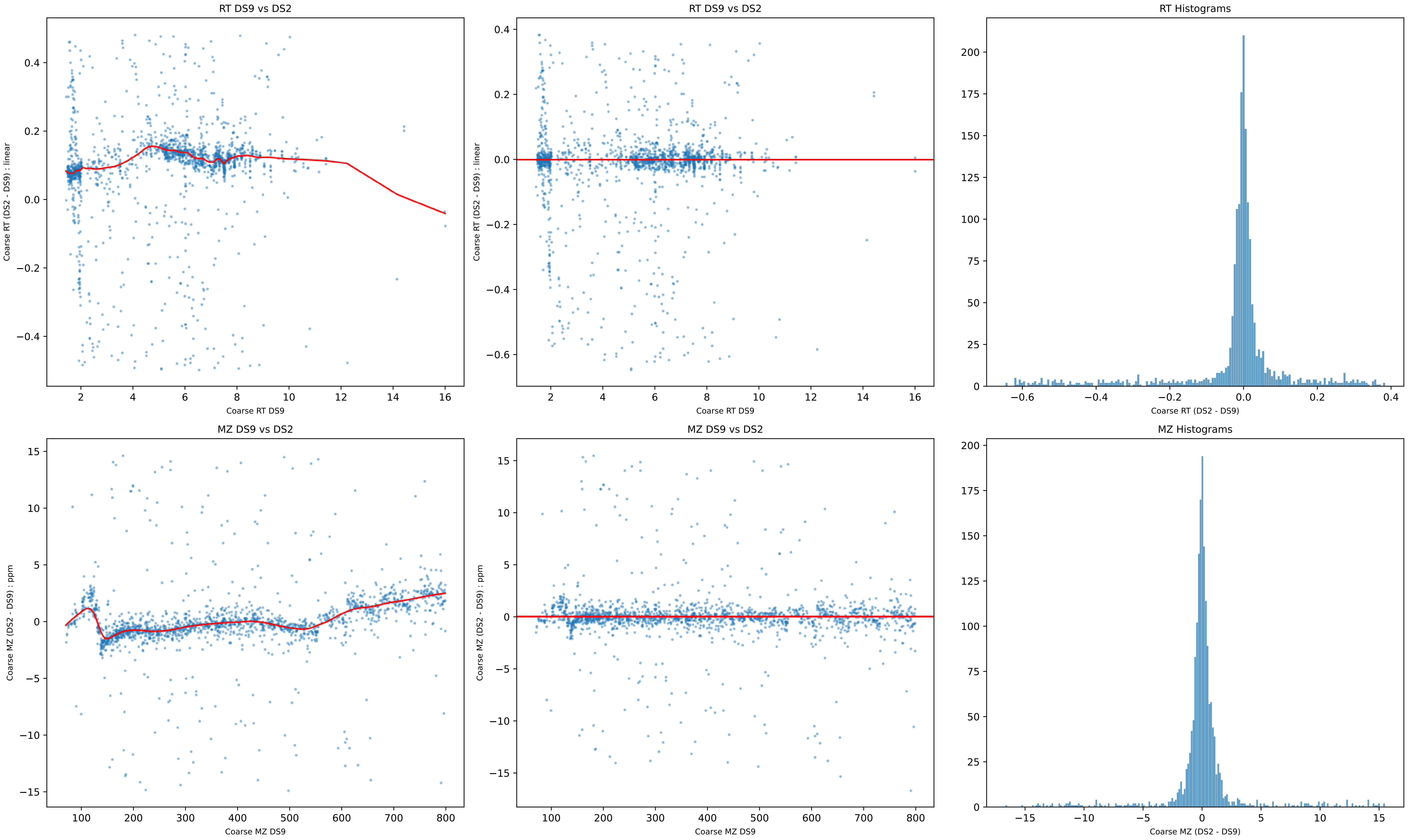
RT DS9 vs DS2
Coarse RT (DS2 - DS9) : linear
Coarse RT DS9
RT DS9 vs DS2
Coarse RT (DS2 - DS9) : linear
Coarse RT DS9
RT Histograms
Coarse RT (DS2 - DS9)
MZ DS9 vs DS2
Coarse MZ (DS2 - DS9) : ppm
Coarse MZ DS9
MZ DS9 vs DS2
Coarse MZ (DS2 - DS9) : ppm
Coarse MZ DS9
MZ Histograms
Coarse MZ (DS2 - DS9)

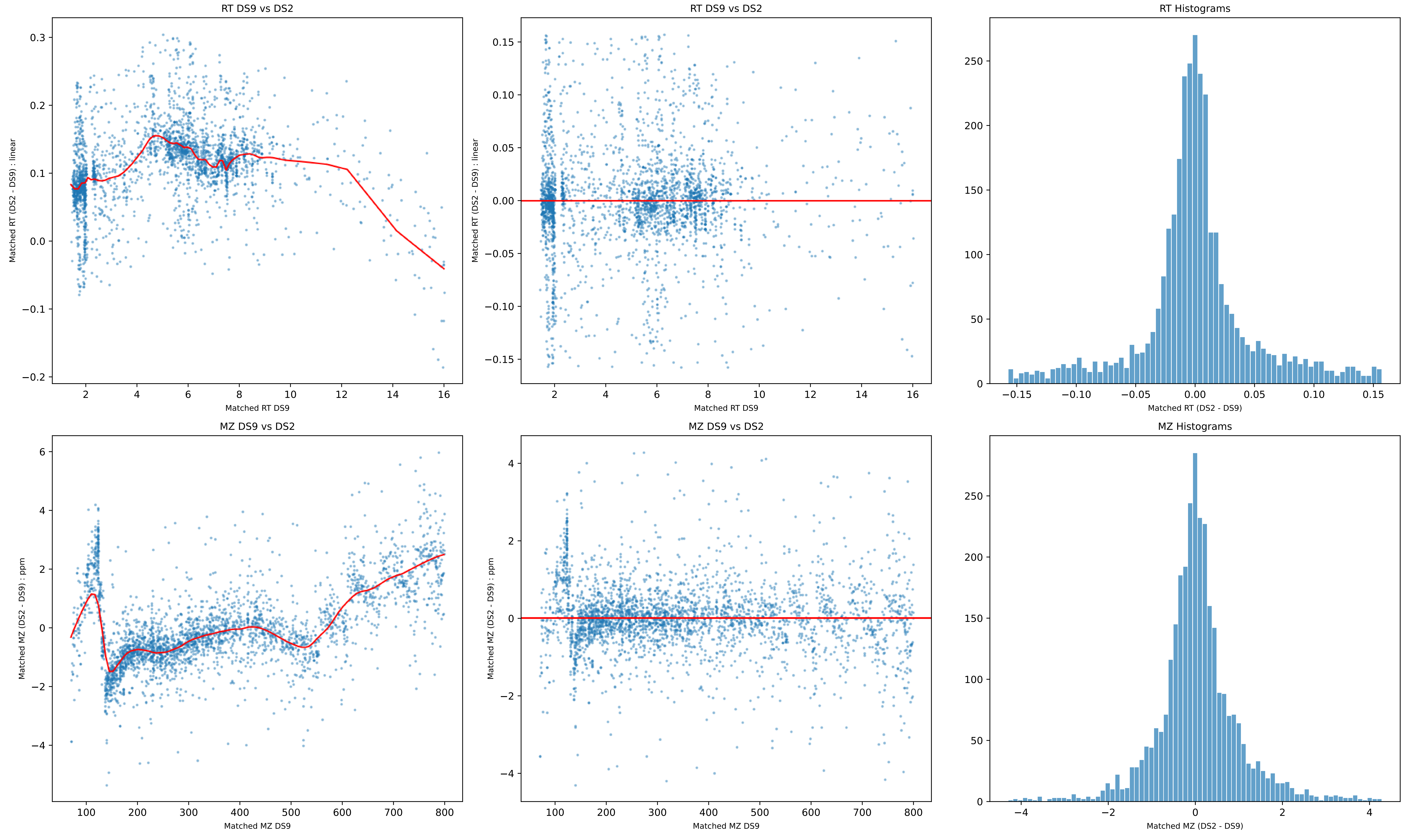

RT DS9 vs DS2
Matched RT (DS2 - DS9) : linear
Matched RT DS9
RT DS9 vs DS2
Matched RT (DS2 - DS9) : linear
Matched RT DS9
RT Histograms
Matched RT (DS2 - DS9)
MZ DS9 vs DS2
Matched MZ (DS2 - DS9) : ppm
Matched MZ DS9
MZ DS9 vs DS2
Matched MZ (DS2 - DS9) : ppm
Matched MZ DS9
MZ Histograms
Matched MZ (DS2 - DS9)

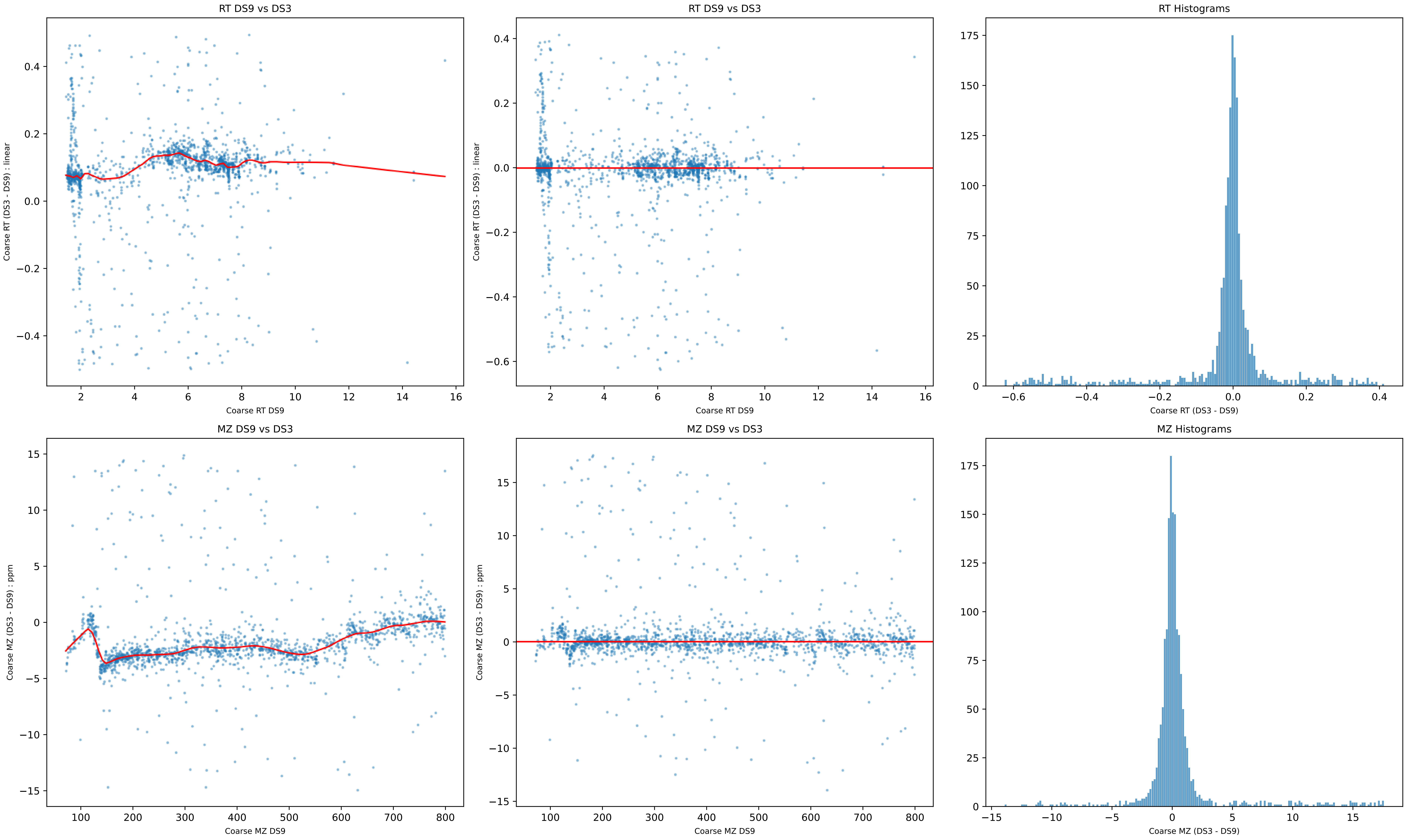
RT DS9 vs DS3
Coarse RT (DS3 - DS9) : linear
Coarse RT DS9
RT DS9 vs DS3
Coarse RT (DS3 - DS9) : linear
Coarse RT DS9
RT Histograms
Coarse RT (DS3 - DS9)
MZ DS9 vs DS3
Coarse MZ (DS3 - DS9) : ppm
Coarse MZ DS9
MZ DS9 vs DS3
Coarse MZ (DS3 - DS9) : ppm
Coarse MZ DS9
MZ Histograms
Coarse MZ (DS3 - DS9)

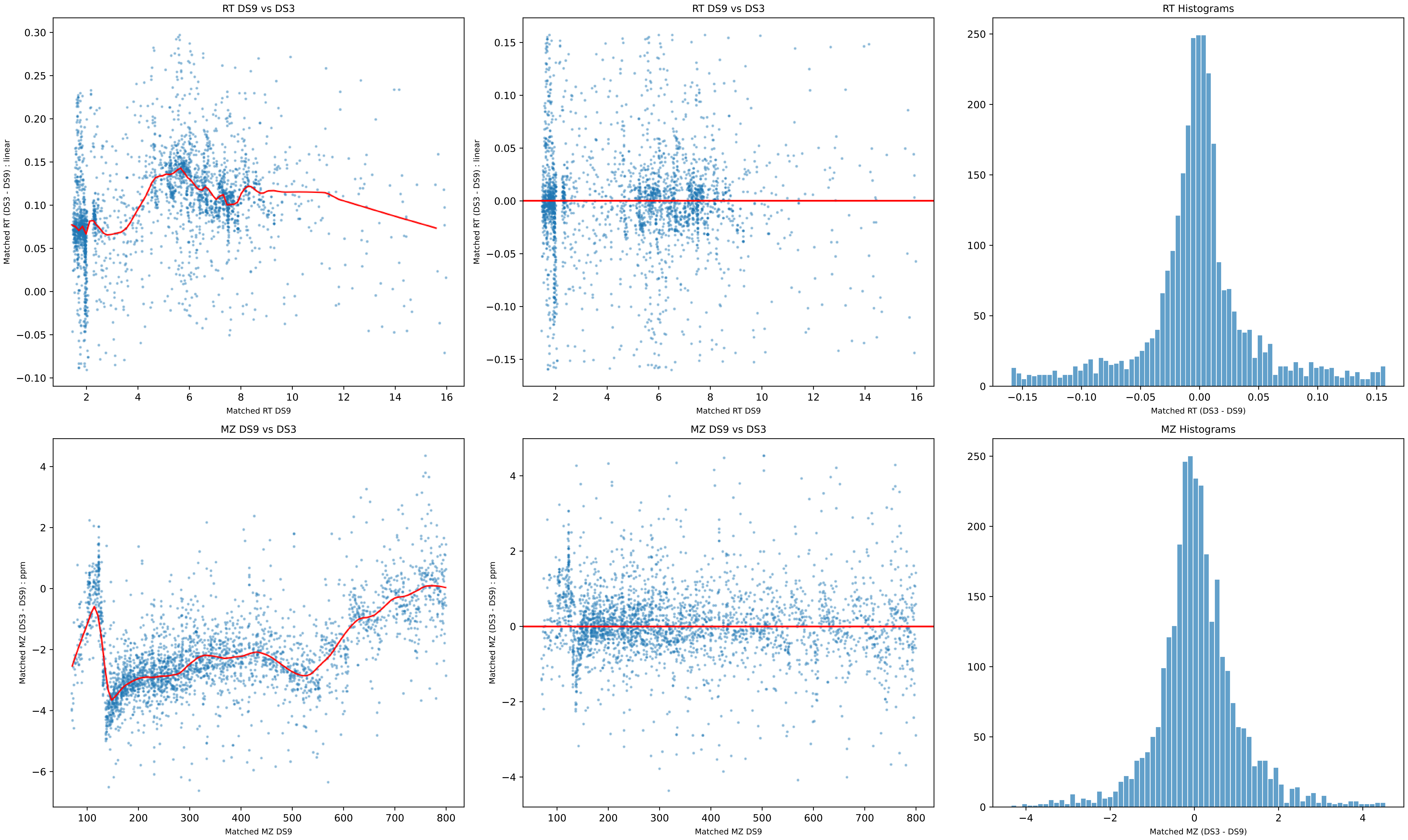
RT DS9 vs DS3
Matched RT (DS3 - DS9) : linear
Matched RT DS9
RT DS9 vs DS3
Matched RT (DS3 - DS9) : linear
Matched RT DS9
RT Histograms
Matched RT (DS3 - DS9)
MZ DS9 vs DS3
Matched MZ (DS3 - DS9) : ppm
Matched MZ DS9
MZ DS9 vs DS3
Matched MZ (DS3 - DS9) : ppm
Matched MZ DS9
MZ Histograms
Matched MZ (DS3 - DS9)

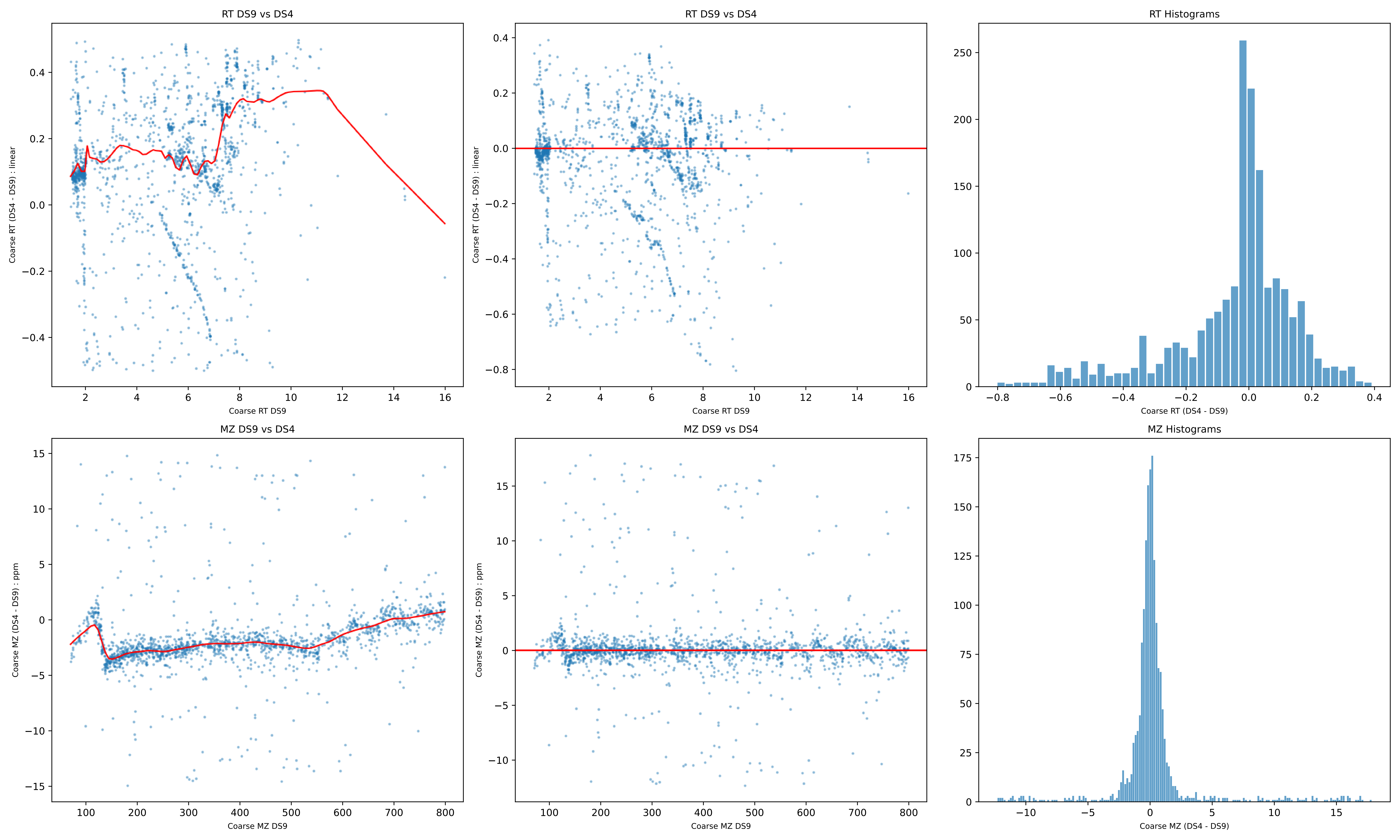
RT DS9 vs DS4
Coarse RT (DS4 - DS9) : linear
Coarse RT DS9
RT DS9 vs DS4
Coarse RT (DS4 - DS9) : linear
Coarse RT DS9
RT Histograms
Coarse RT (DS4 - DS9)
MZ DS9 vs DS4
Coarse MZ (DS4 - DS9) : ppm
Coarse MZ DS9
MZ DS9 vs DS4
Coarse MZ (DS4 - DS9) : ppm
Coarse MZ DS9
MZ Histograms
Coarse MZ (DS4 - DS9)

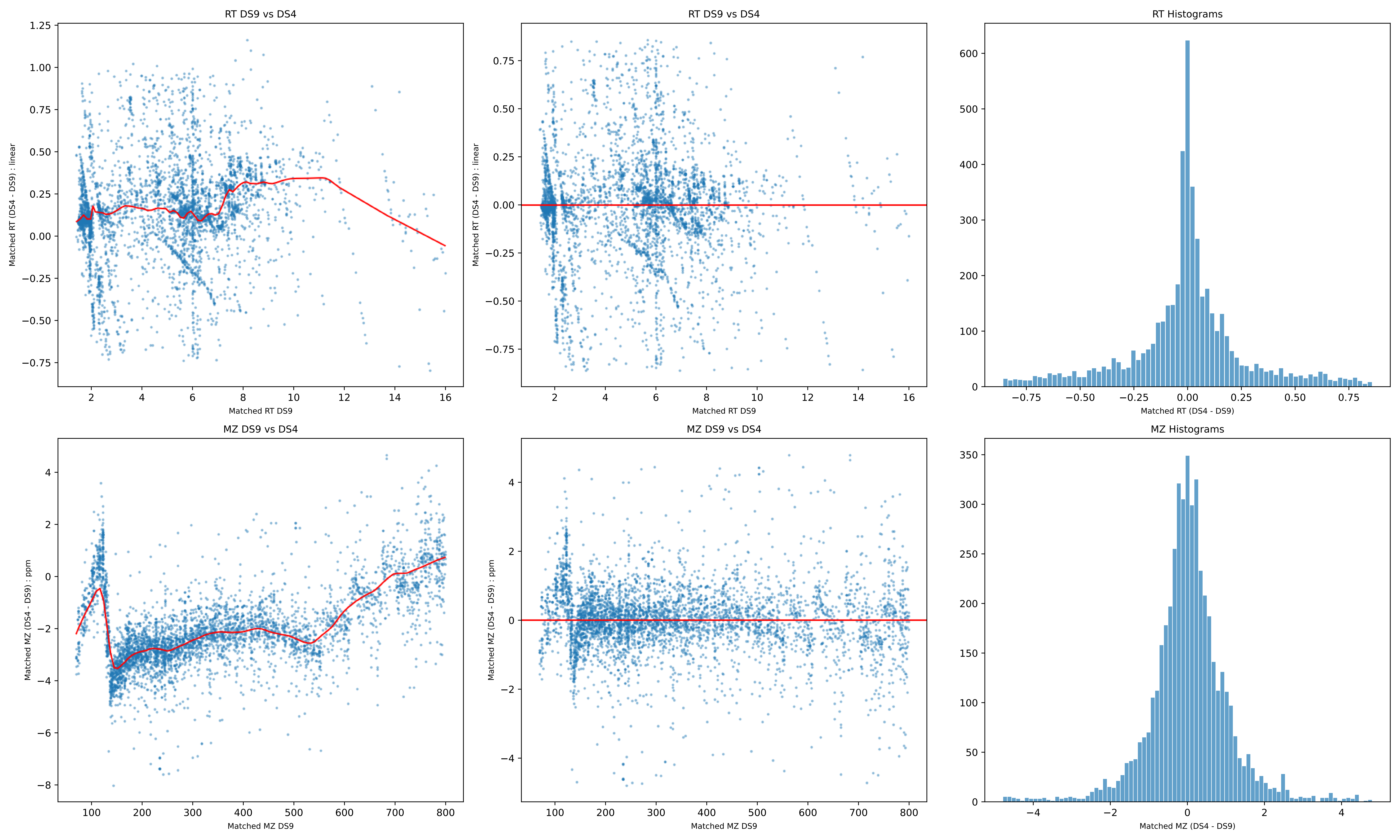
RT DS9 vs DS4
Matched RT (DS4 - DS9) : linear
Matched RT DS9
RT DS9 vs DS4
Matched RT (DS4 - DS9) : linear
Matched RT DS9
RT Histograms
Matched RT (DS4 - DS9)
MZ DS9 vs DS4
Matched MZ (DS4 - DS9) : ppm
Matched MZ DS9
MZ DS9 vs DS4
Matched MZ (DS4 - DS9) : ppm
Matched MZ DS9
MZ Histograms
Matched MZ (DS4 - DS9)

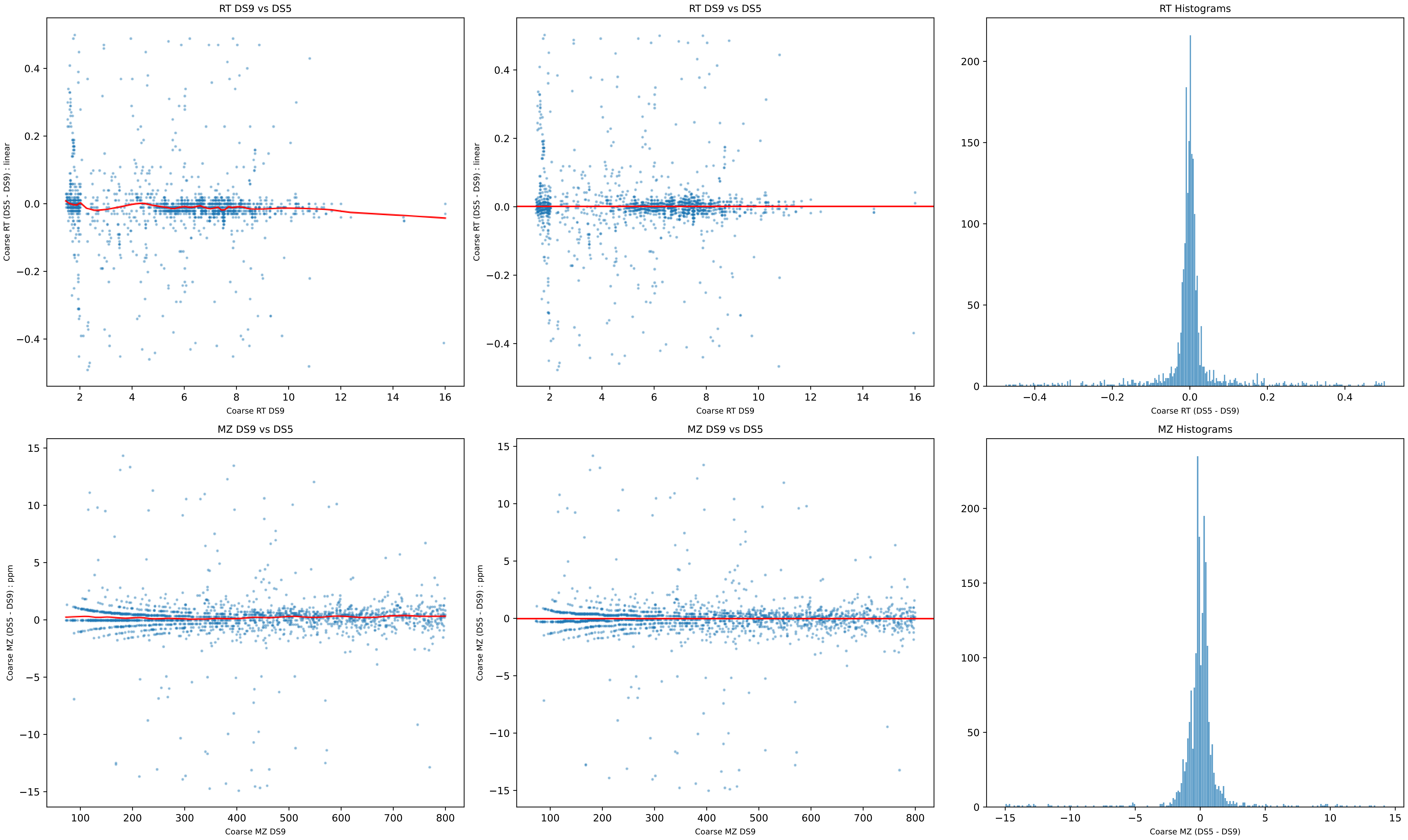
RT DS9 vs DS5
Coarse RT (DS5 - DS9) : linear
Coarse RT DS9
RT DS9 vs DS5
Coarse RT (DS5 - DS9) : linear
Coarse RT DS9
RT Histograms
Coarse RT (DS5 - DS9)
MZ DS9 vs DS5
Coarse MZ (DS5 - DS9) : ppm
Coarse MZ DS9
MZ DS9 vs DS5
Coarse MZ (DS5 - DS9) : ppm
Coarse MZ DS9
MZ Histograms
Coarse MZ (DS5 - DS9)

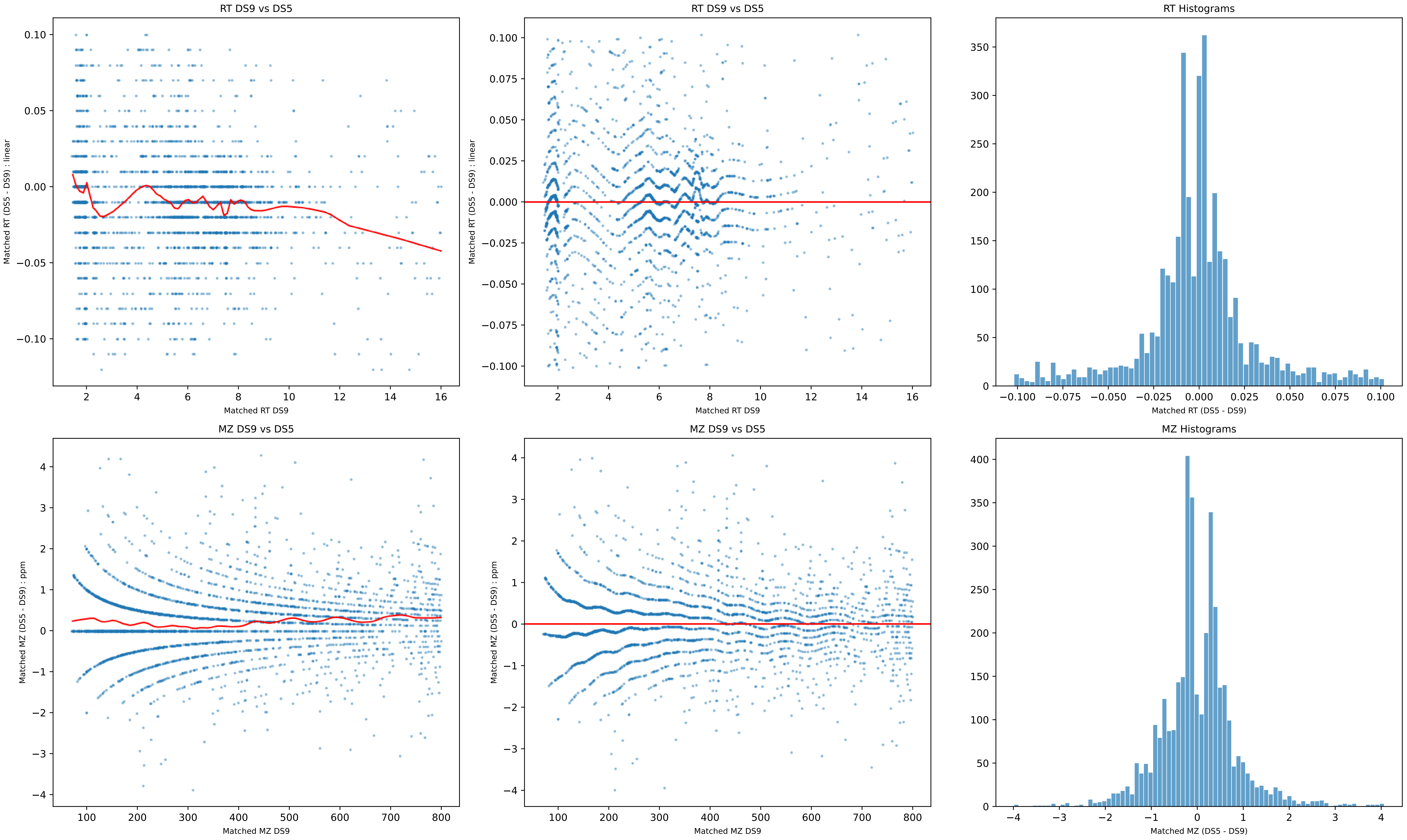
RT DS9 vs DS5
Matched RT (DS5 - DS9) : linear
Matched RT DS9
RT DS9 vs DS5
Matched RT (DS5 - DS9) : linear
Matched RT DS9
RT Histograms
Matched RT (DS5 - DS9)
MZ DS9 vs DS5
Matched MZ (DS5 - DS9) : ppm
Matched MZ DS9
MZ DS9 vs DS5
Matched MZ (DS5 - DS9) : ppm
Matched MZ DS9
MZ Histograms
Matched MZ (DS5 - DS9)

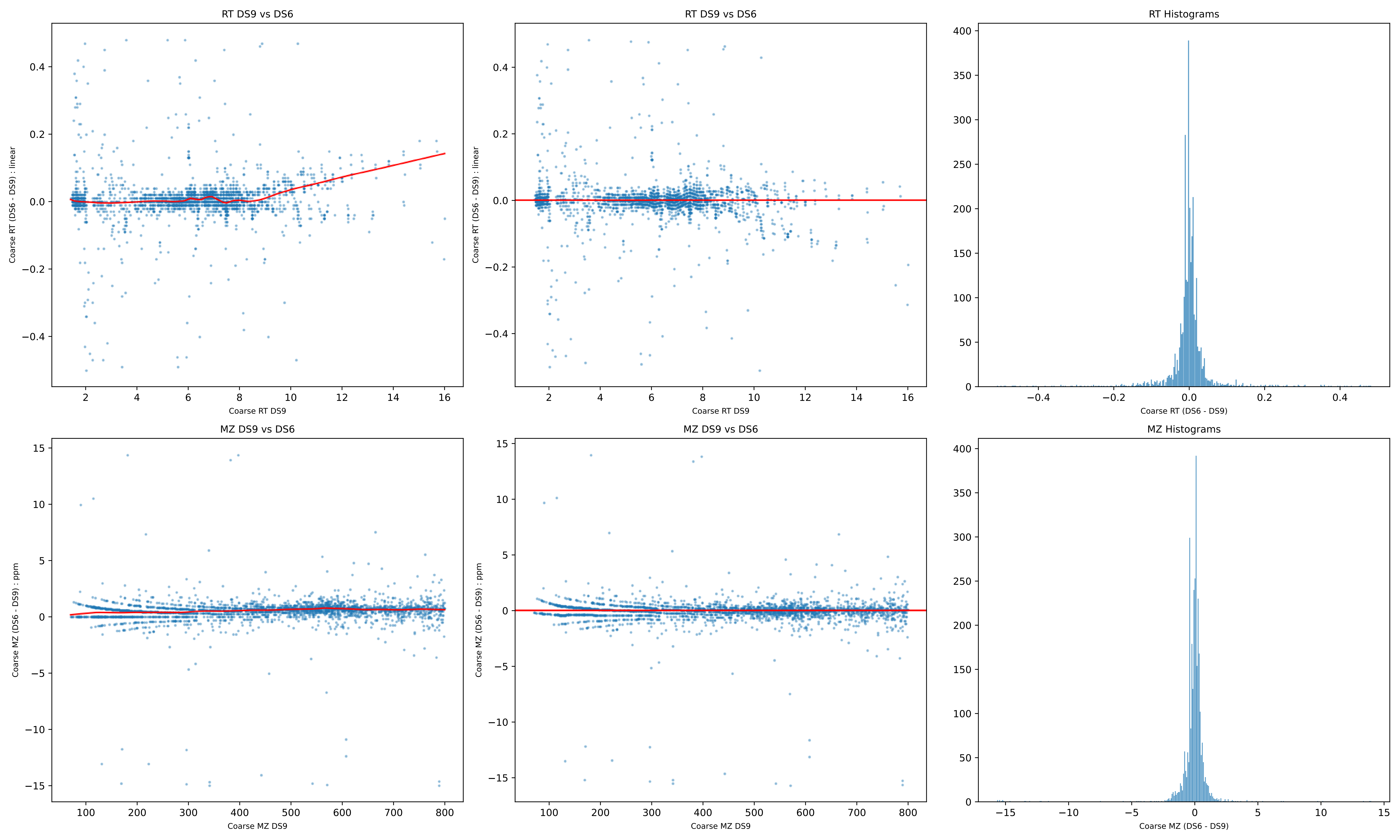

RT DS9 vs DS6
Coarse RT (DS6 - DS9) : linear
Coarse RT DS9
RT DS9 vs DS6
Coarse RT (DS6 - DS9) : linear
Coarse RT DS9
RT Histograms
Coarse RT (DS6 - DS9)
MZ DS9 vs DS6
Coarse MZ (DS6 - DS9) : ppm
Coarse MZ DS9
MZ DS9 vs DS6
Coarse MZ (DS6 - DS9) : ppm
Coarse MZ DS9
MZ Histograms
Coarse MZ (DS6 - DS9)

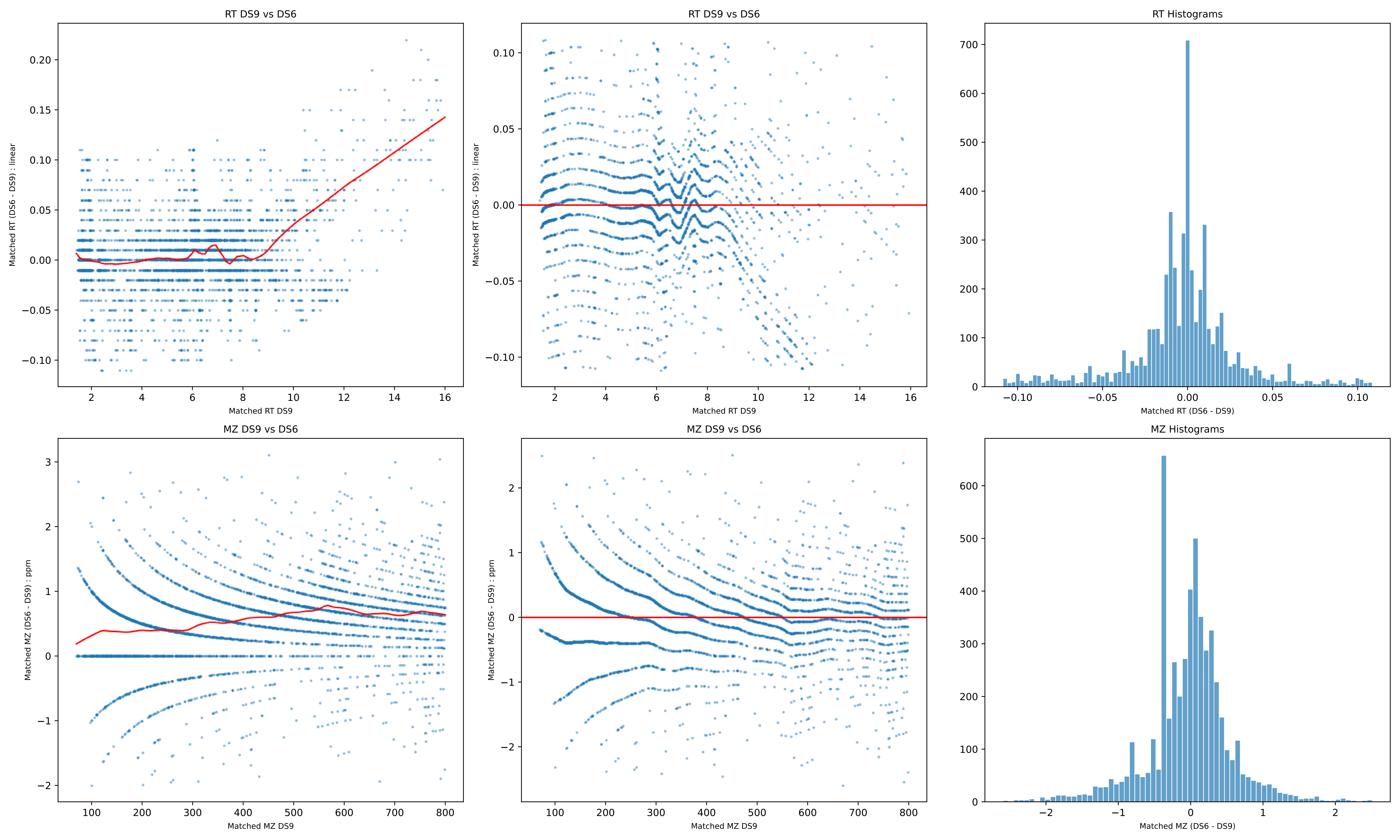
RT DS9 vs DS6
Matched RT (DS6 - DS9) : linear
Matched RT DS9
RT DS9 vs DS6
Matched RT (DS6 - DS9) : linear
Matched RT DS9
RT Histograms
Matched RT (DS6 - DS9)
MZ DS9 vs DS6
Matched MZ (DS6 - DS9) : ppm
Matched MZ DS9
MZ DS9 vs DS6
Matched MZ (DS6 - DS9) : ppm
Matched MZ DS9
MZ Histograms
Matched MZ (DS6 - DS9)

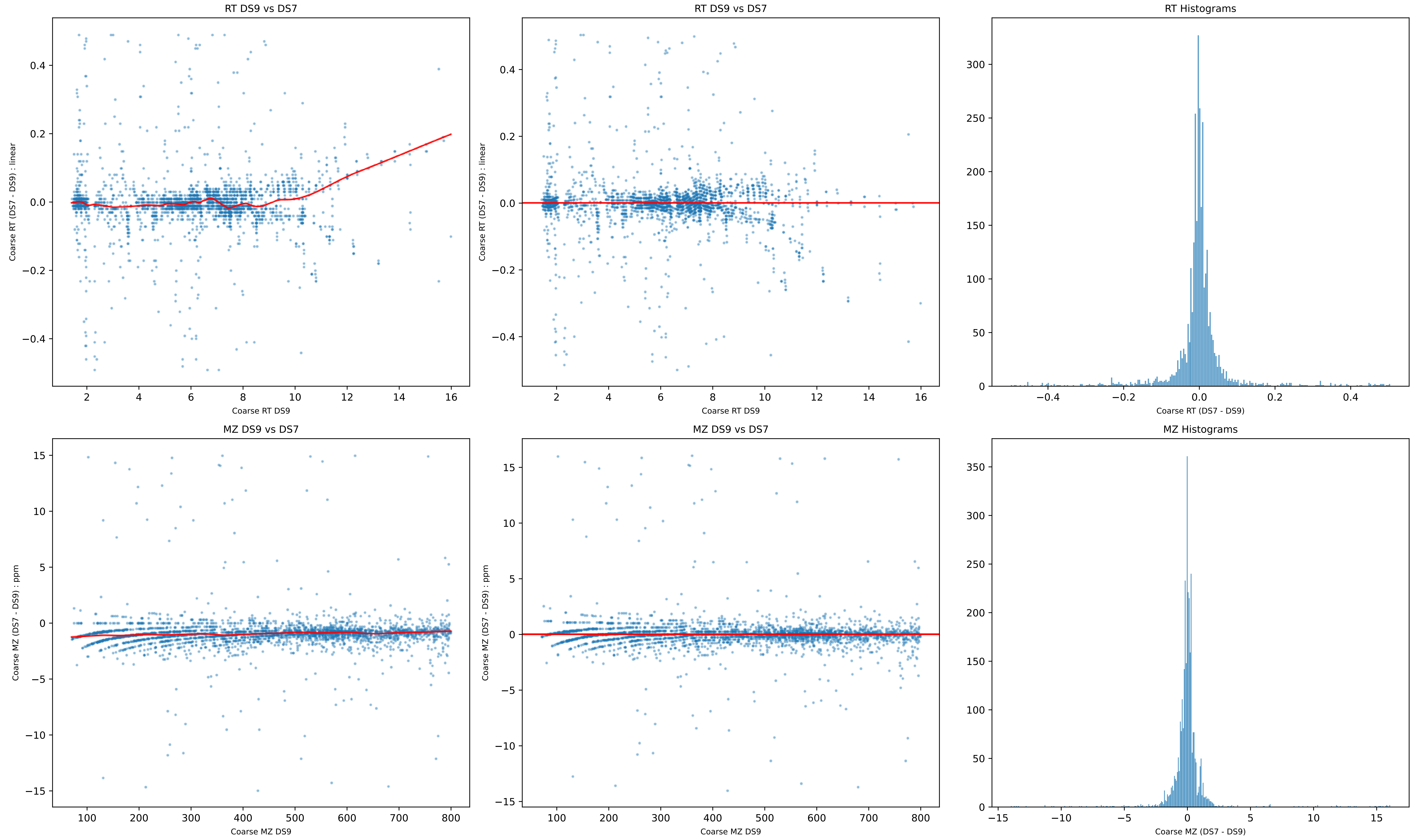
RT DS9 vs DS7
Coarse RT (DS7 - DS9) : linear
Coarse RT DS9
RT DS9 vs DS7
Coarse RT (DS7 - DS9) : linear
Coarse RT DS9
RT Histograms
Coarse RT (DS7 - DS9)
MZ DS9 vs DS7
Coarse MZ (DS7 - DS9) : ppm
Coarse MZ DS9
MZ DS9 vs DS7
Coarse MZ (DS7 - DS9) : ppm
Coarse MZ DS9
MZ Histograms
Coarse MZ (DS7 - DS9)

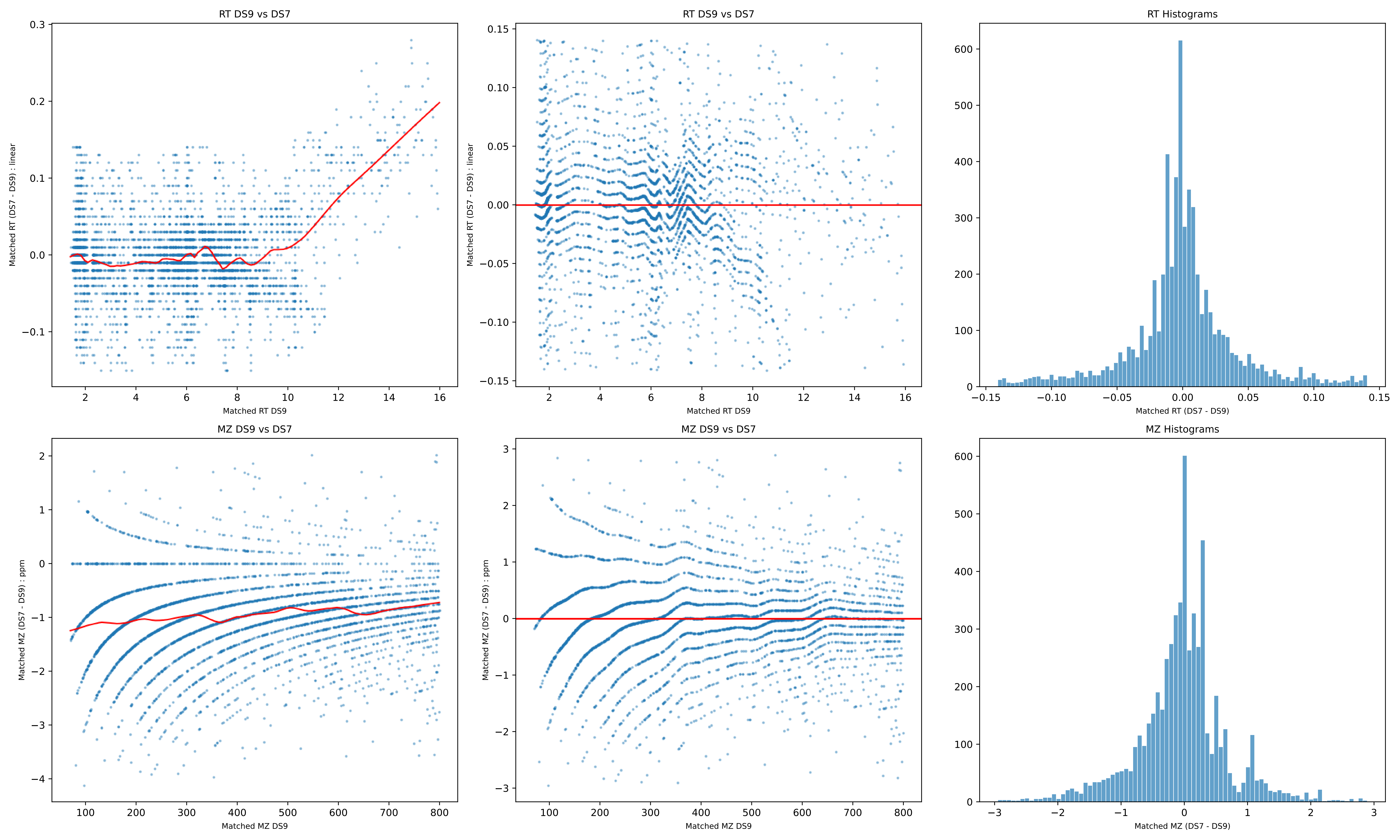
RT DS9 vs DS7
Matched RT (DS7 - DS9) : linear
Matched RT DS9
RT DS9 vs DS7
Matched RT (DS7 - DS9) : linear
Matched RT DS9
RT Histograms
Matched RT (DS7 - DS9)
MZ DS9 vs DS7
Matched MZ (DS7 - DS9) : ppm
Matched MZ DS9
MZ DS9 vs DS7
Matched MZ (DS7 - DS9) : ppm
Matched MZ DS9
MZ Histograms
Matched MZ (DS7 - DS9)

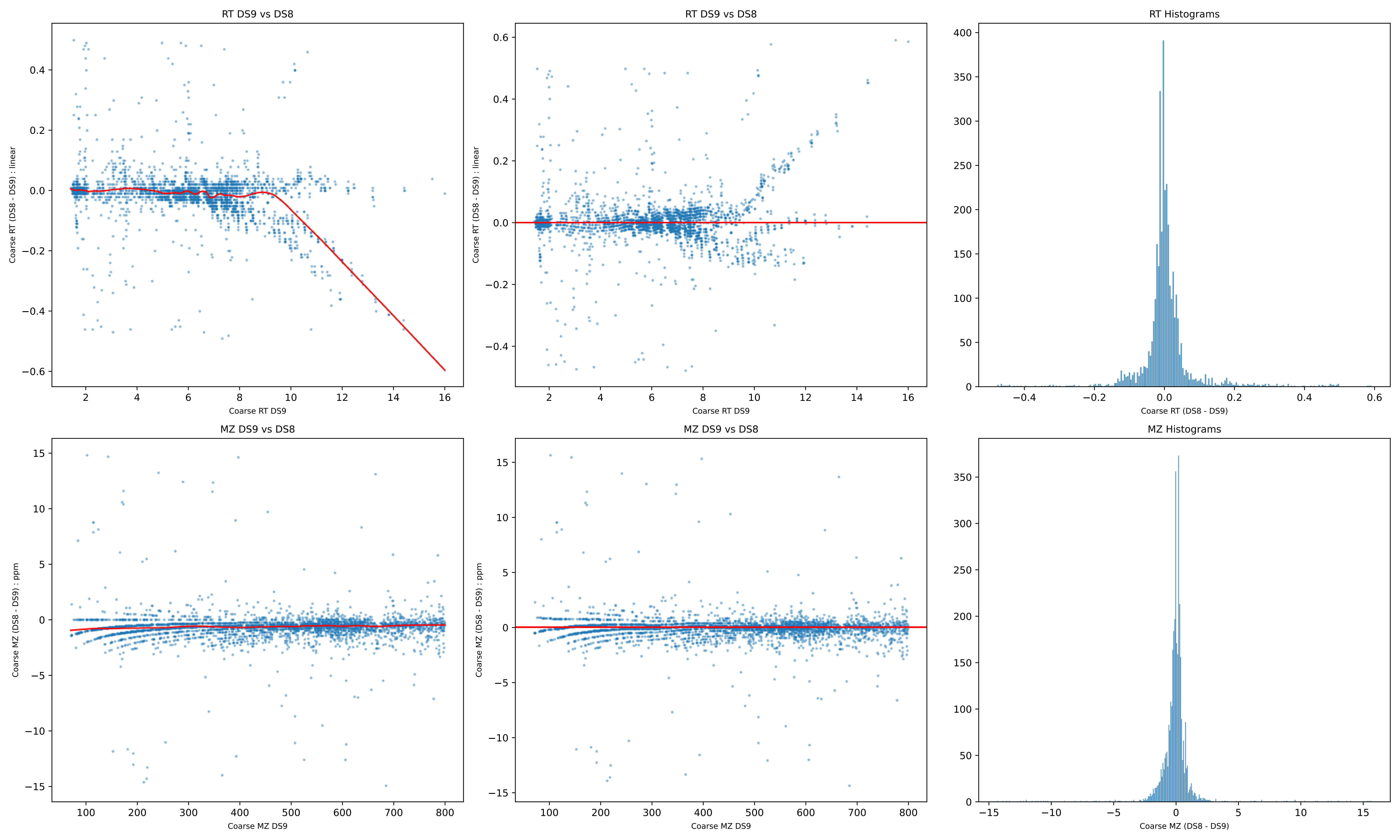
RT DS9 vs DS8
Coarse RT (DS8 - DS9) : linear
Coarse RT DS9
RT DS9 vs DS8
Coarse RT (DS8 - DS9) : linear
Coarse RT DS9
RT Histograms
Coarse RT (DS8 - DS9)
MZ DS9 vs DS8
Coarse MZ (DS8 - DS9) : ppm
Coarse MZ DS9
MZ DS9 vs DS8
Coarse MZ (DS8 - DS9) : ppm
Coarse MZ DS9
MZ Histograms
Coarse MZ (DS8 - DS9)

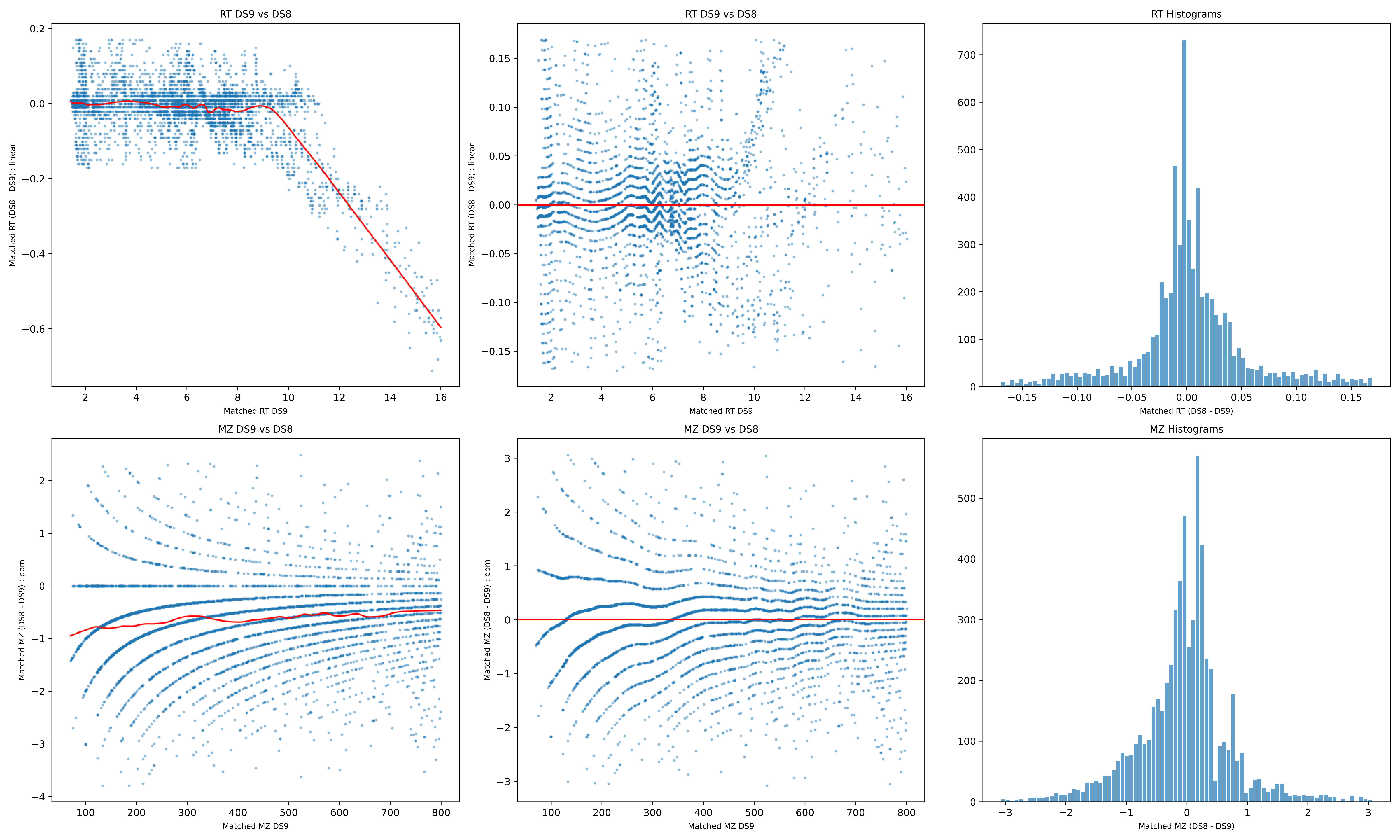

RT DS9 vs DS8
Matched RT (DS8 - DS9) : linear
Matched RT DS9
RT DS9 vs DS8
Matched RT (DS8 - DS9) : linear
Matched RT DS9
RT Histograms
Matched RT (DS8 - DS9)
MZ DS9 vs DS8
Matched MZ (DS8 - DS9) : ppm
Matched MZ DS9
MZ DS9 vs DS8
Matched MZ (DS8 - DS9) : ppm
Matched MZ DS9
MZ Histograms
Matched MZ (DS8 - DS9)

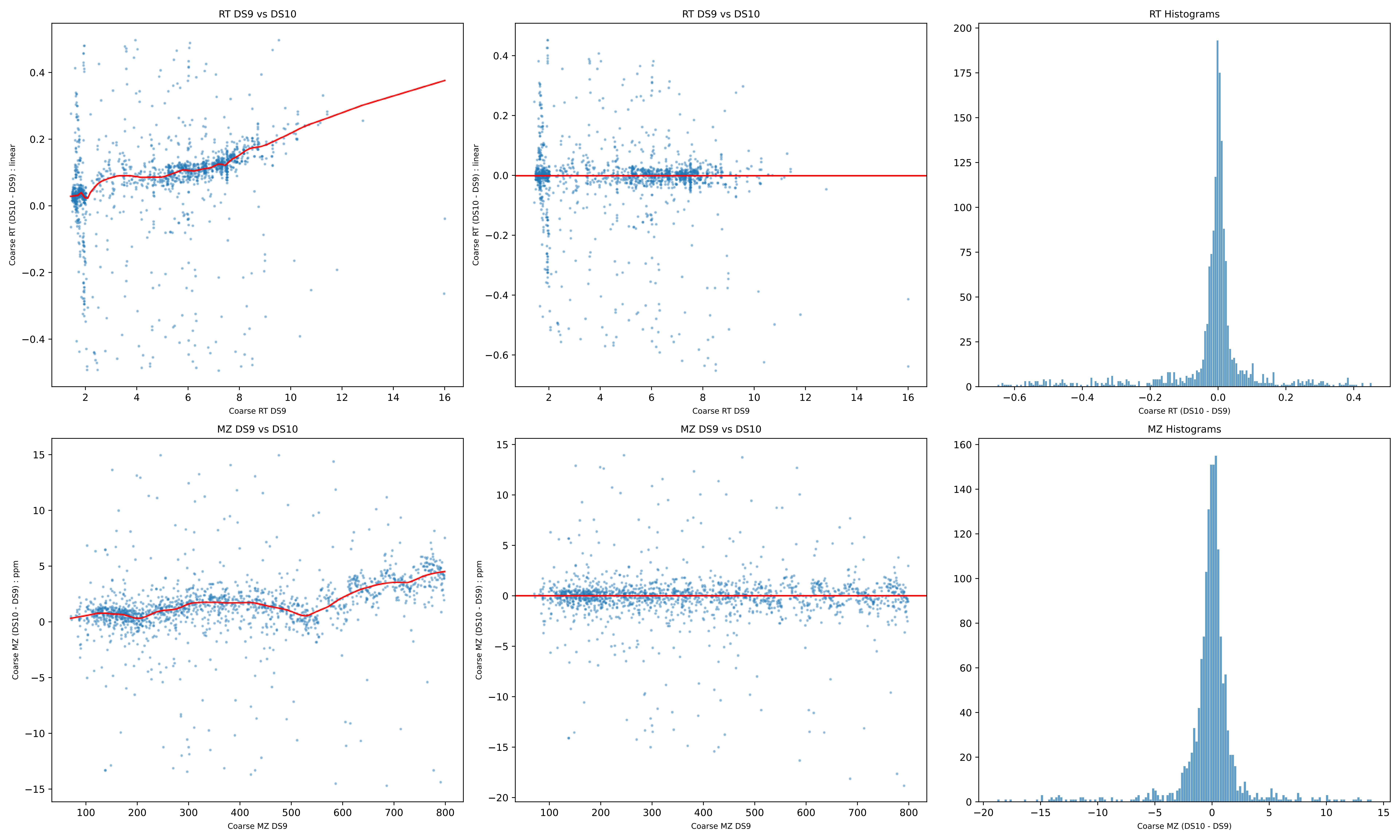

RT DS9 vs DS10
Coarse RT (DS10 - DS9) : linear
Coarse RT DS9
RT DS9 vs DS10
Coarse RT (DS10 - DS9) : linear
Coarse RT DS9
RT Histograms
Coarse RT (DS10 - DS9)
MZ DS9 vs DS10
Coarse MZ (DS10 - DS9) : ppm
Coarse MZ DS9
MZ DS9 vs DS10
Coarse MZ (DS10 - DS9) : ppm
Coarse MZ DS9
MZ Histograms
Coarse MZ (DS10 - DS9)

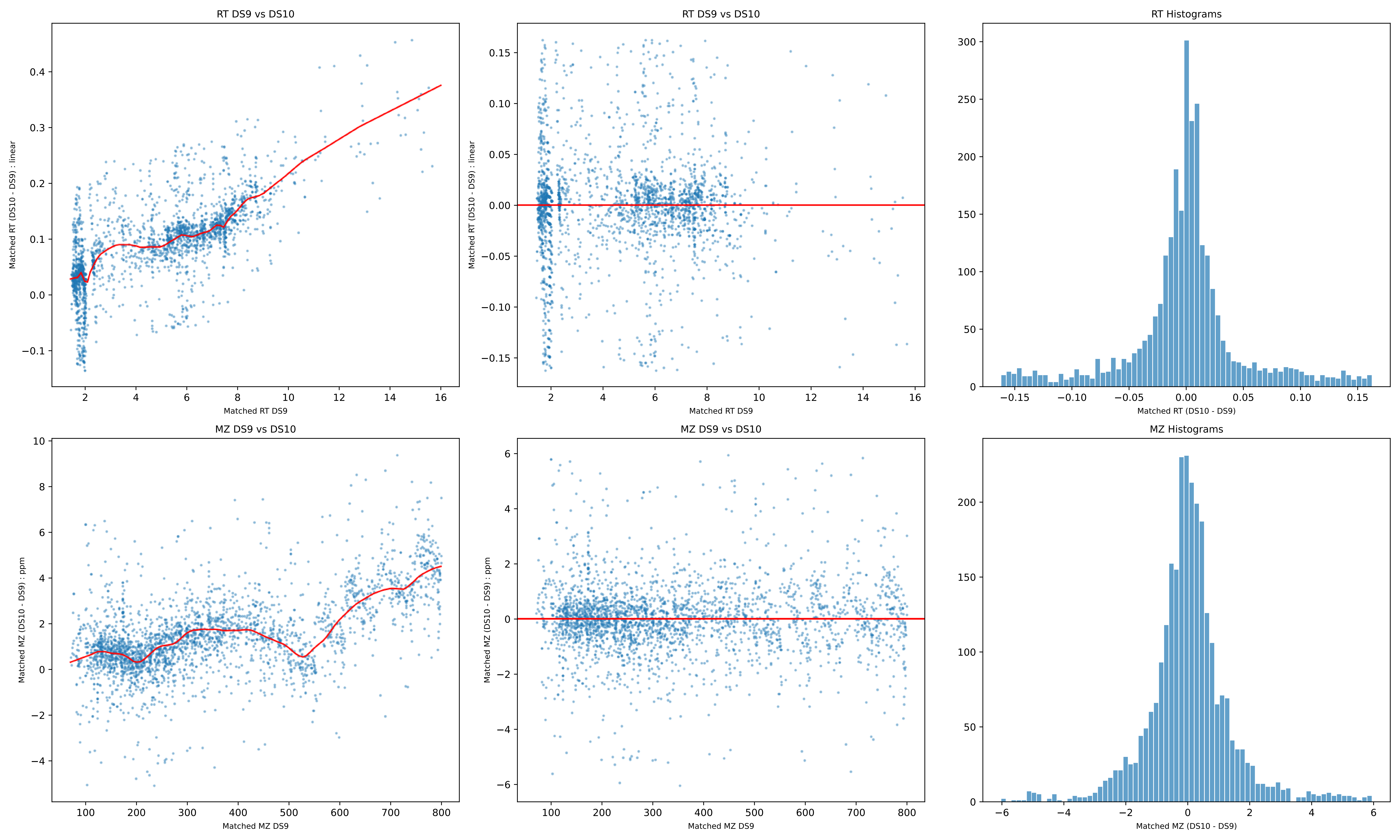
RT DS9 vs DS10
Matched RT (DS10 - DS9) : linear
Matched RT DS9
RT DS9 vs DS10
Matched RT (DS10 - DS9) : linear
Matched RT DS9
RT Histograms
Matched RT (DS10 - DS9)
MZ DS9 vs DS10
Matched MZ (DS10 - DS9) : ppm
Matched MZ DS9
MZ DS9 vs DS10
Matched MZ (DS10 - DS9) : ppm
Matched MZ DS9
MZ Histograms
Matched MZ (DS10 - DS9)

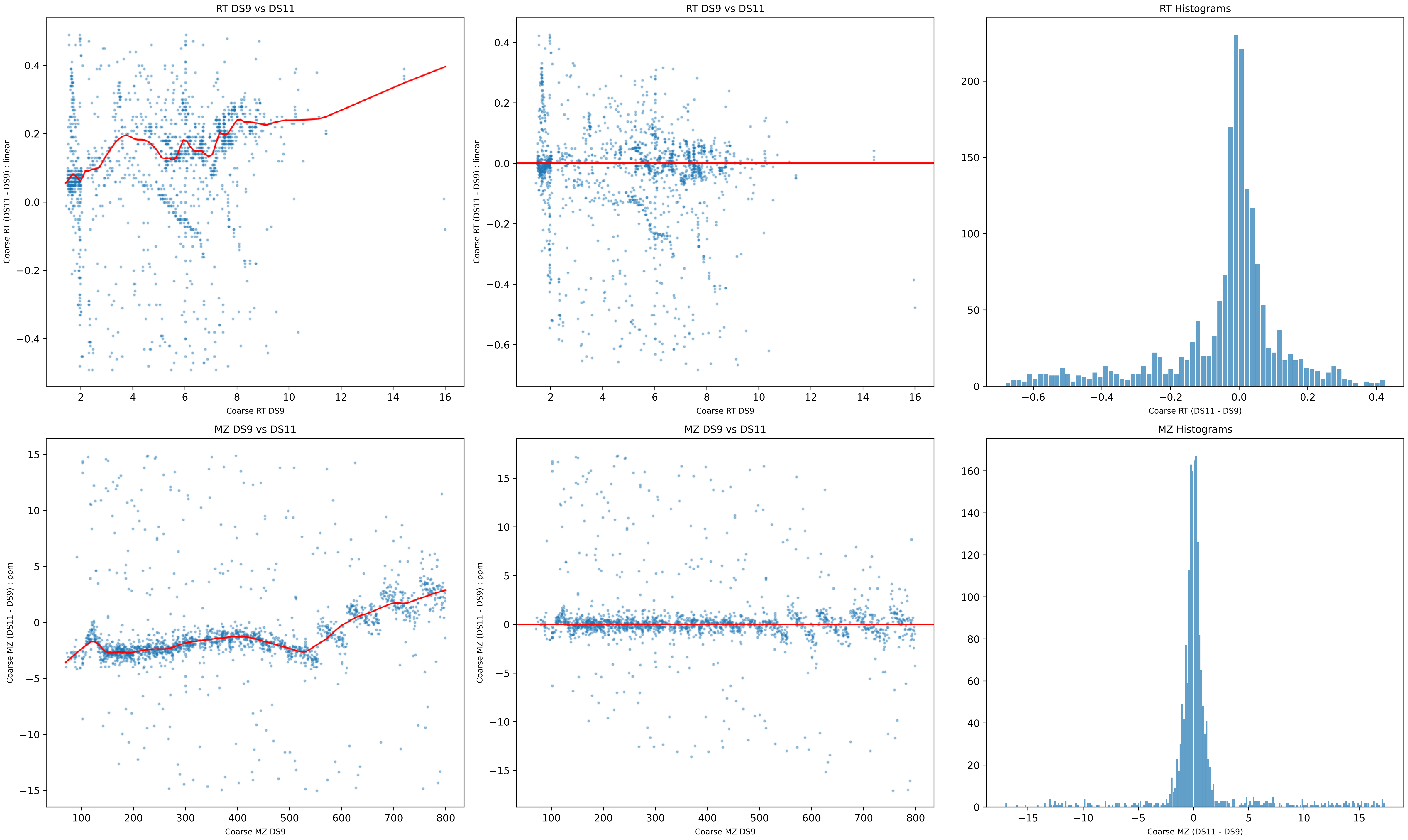

RT DS9 vs DS11
Coarse RT (DS11 - DS9) : linear
Coarse RT DS9
RT DS9 vs DS11
Coarse RT (DS11 - DS9) : linear
Coarse RT DS9
RT Histograms
Coarse RT (DS11 - DS9)
MZ DS9 vs DS11
Coarse MZ (DS11 - DS9) : ppm
Coarse MZ DS9
MZ DS9 vs DS11
Coarse MZ (DS11 - DS9) : ppm
Coarse MZ DS9
MZ Histograms
Coarse MZ (DS11 - DS9)

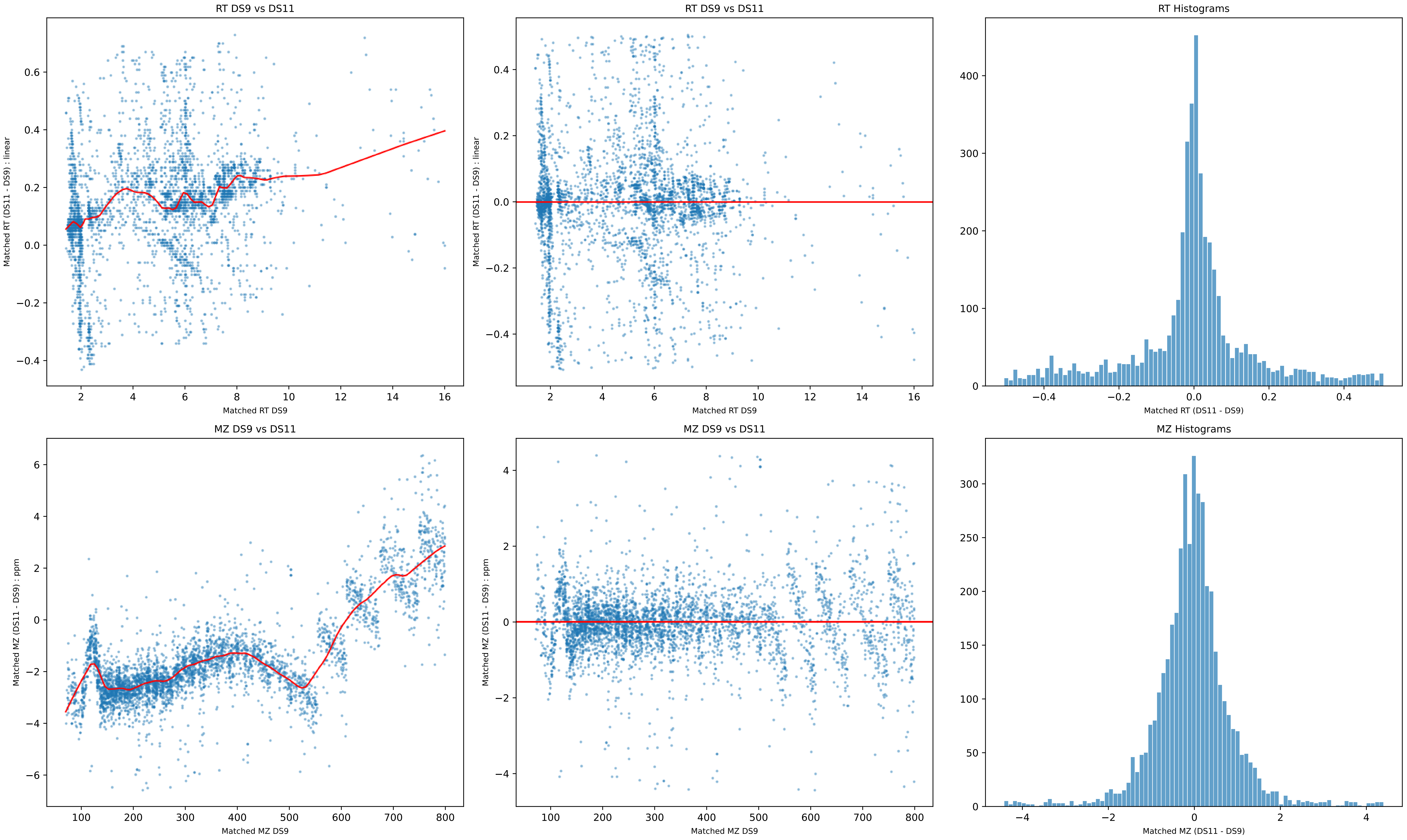

RT DS9 vs DS11
Matched RT (DS11 - DS9) : linear
Matched RT DS9
RT DS9 vs DS11
Matched RT (DS11 - DS9) : linear
Matched RT DS9
RT Histograms
Matched RT (DS11 - DS9)
MZ DS9 vs DS11
Matched MZ (DS11 - DS9) : ppm
Matched MZ DS9
MZ DS9 vs DS11
Matched MZ (DS11 - DS9) : ppm
Matched MZ DS9
MZ Histograms
Matched MZ (DS11 - DS9)

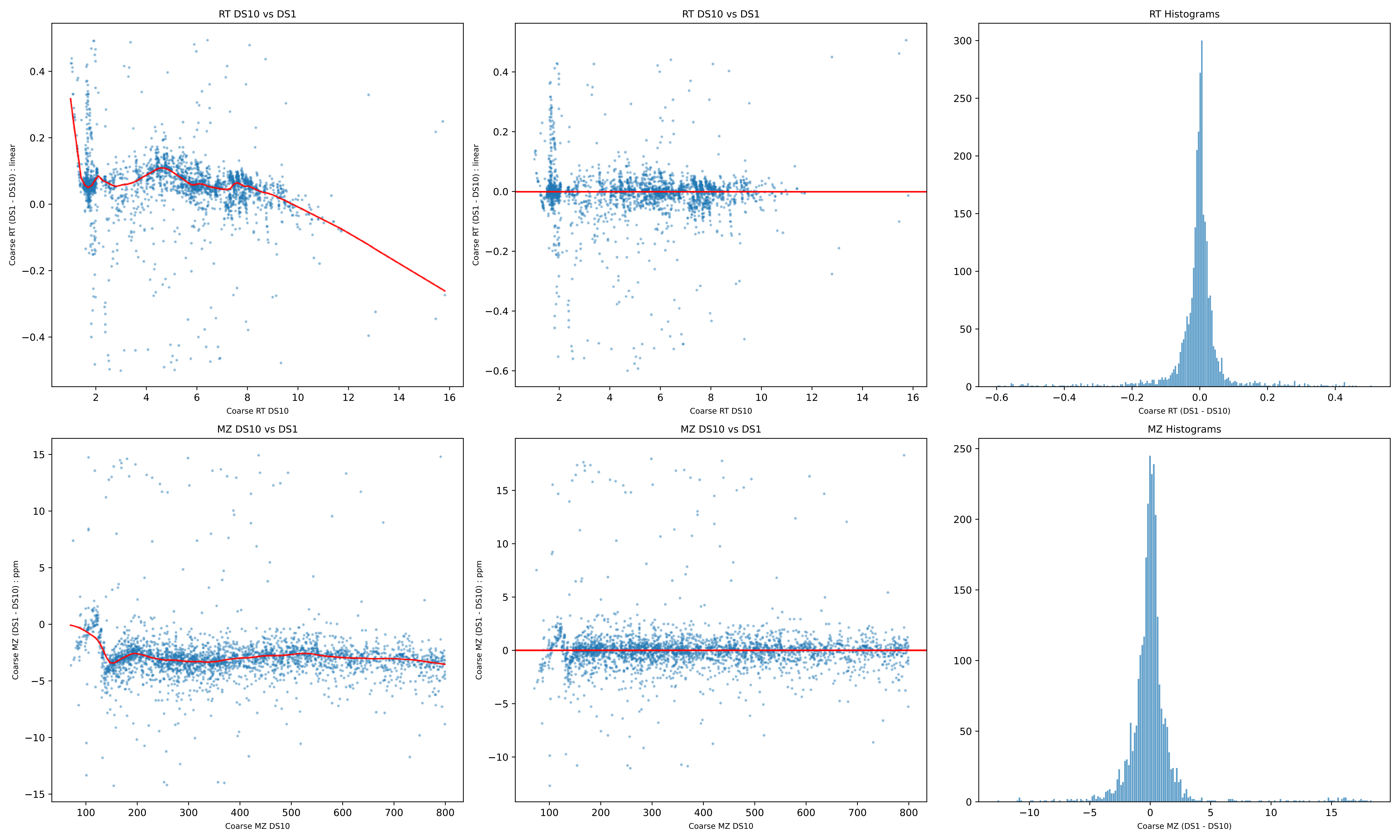
RT DS10 vs DS1
Coarse RT (DS1 - DS10) : linear
Coarse RT DS10
RT DS10 vs DS1
Coarse RT (DS1 - DS10) : linear
Coarse RT DS10
RT Histograms
Coarse RT (DS1 - DS10)
MZ DS10 vs DS1
Coarse MZ (DS1 - DS10) : ppm
Coarse MZ DS10
MZ DS10 vs DS1
Coarse MZ (DS1 - DS10) : ppm
Coarse MZ DS10
MZ Histograms
Coarse MZ (DS1 - DS10)

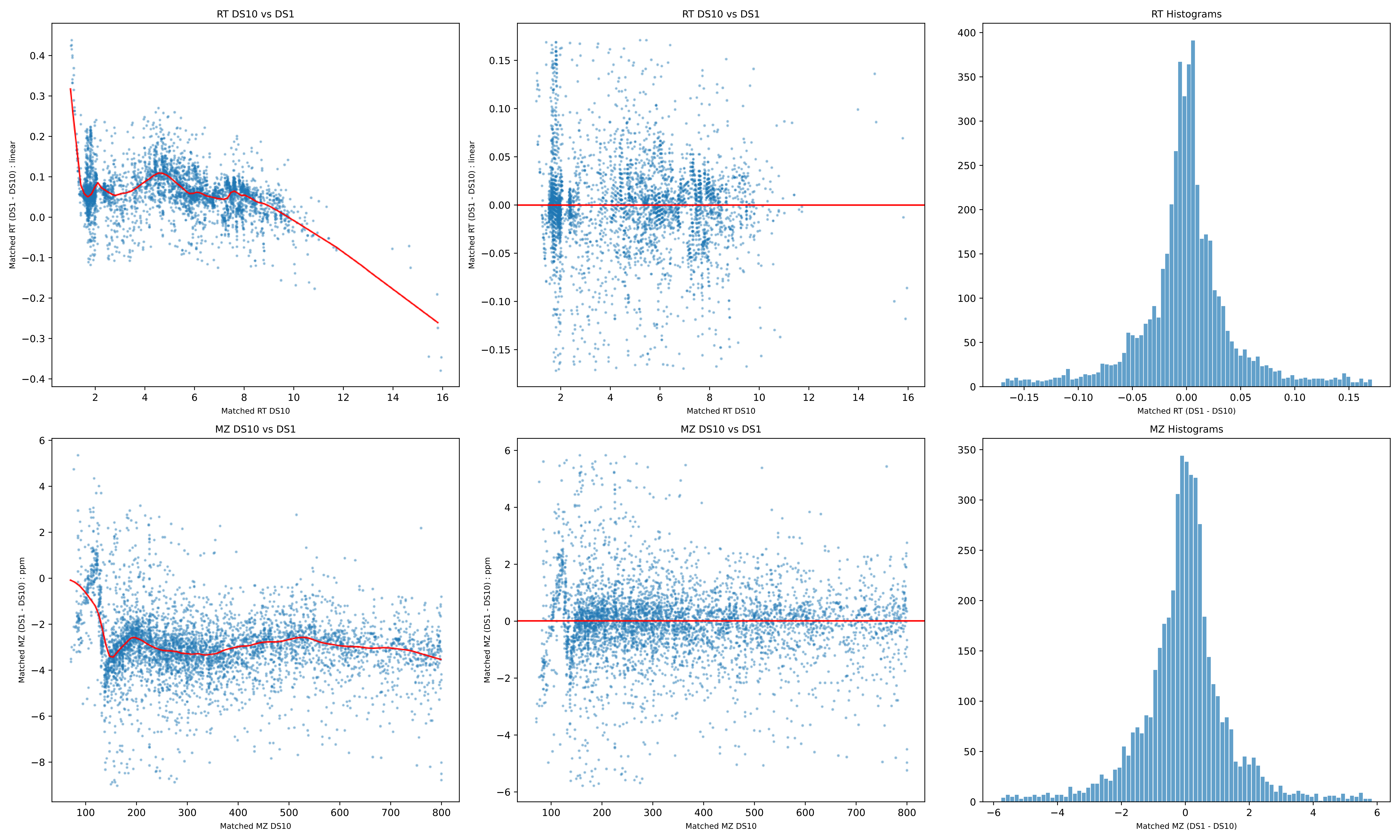
RT DS10 vs DS1
Matched RT (DS1 - DS10) : linear
Matched RT DS10
RT DS10 vs DS1
Matched RT (DS1 - DS10) : linear
Matched RT DS10
RT Histograms
Matched RT (DS1 - DS10)
MZ DS10 vs DS1
Matched MZ (DS1 - DS10) : ppm
Matched MZ DS10
MZ DS10 vs DS1
Matched MZ (DS1 - DS10) : ppm
Matched MZ DS10
MZ Histograms
Matched MZ (DS1 - DS10)

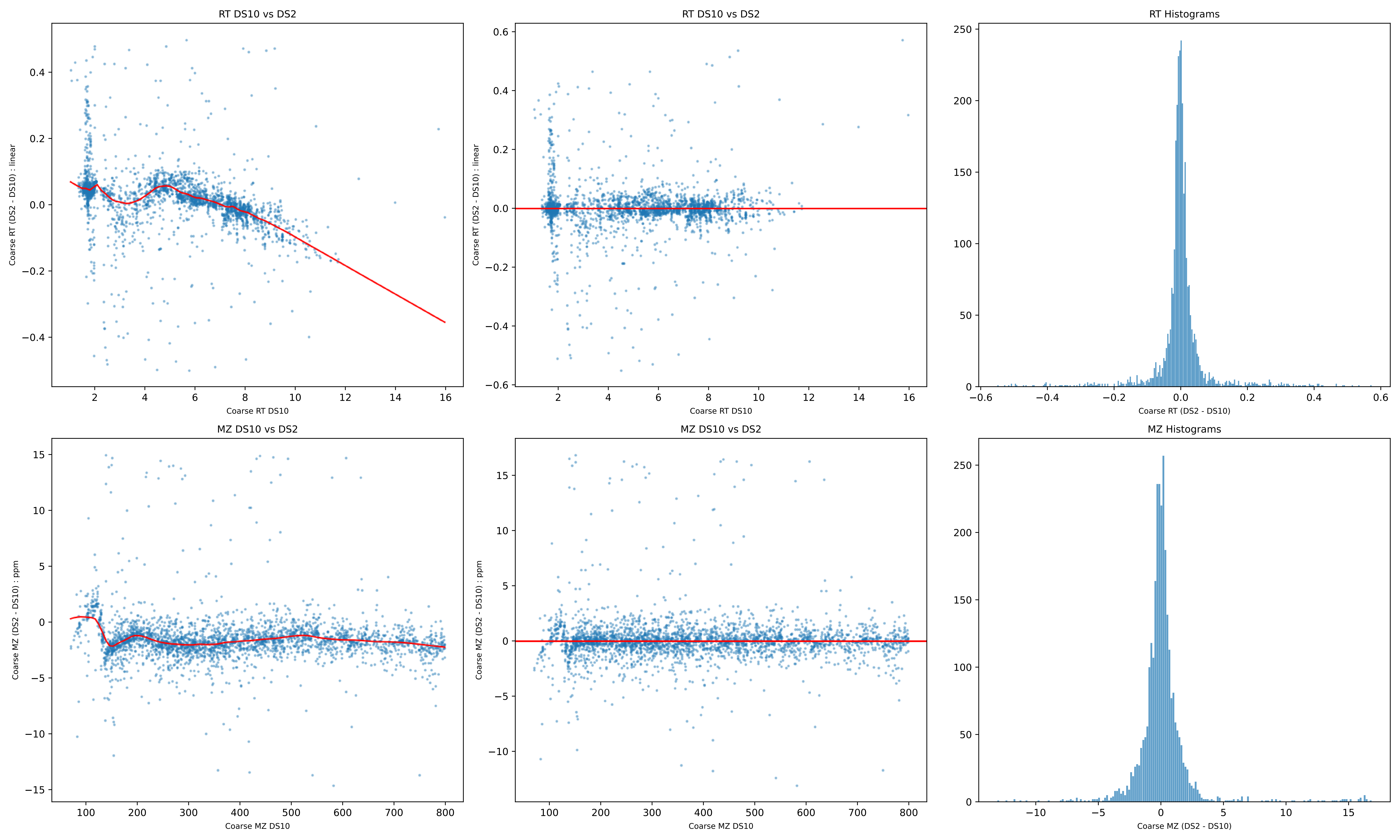
RT DS10 vs DS2
Coarse RT (DS2 - DS10) : linear
Coarse RT DS10
RT DS10 vs DS2
Coarse RT (DS2 - DS10) : linear
Coarse RT DS10
RT Histograms
Coarse RT (DS2 - DS10)
MZ DS10 vs DS2
Coarse MZ (DS2 - DS10) : ppm
Coarse MZ DS10
MZ DS10 vs DS2
Coarse MZ (DS2 - DS10) : ppm
Coarse MZ DS10
MZ Histograms
Coarse MZ (DS2 - DS10)

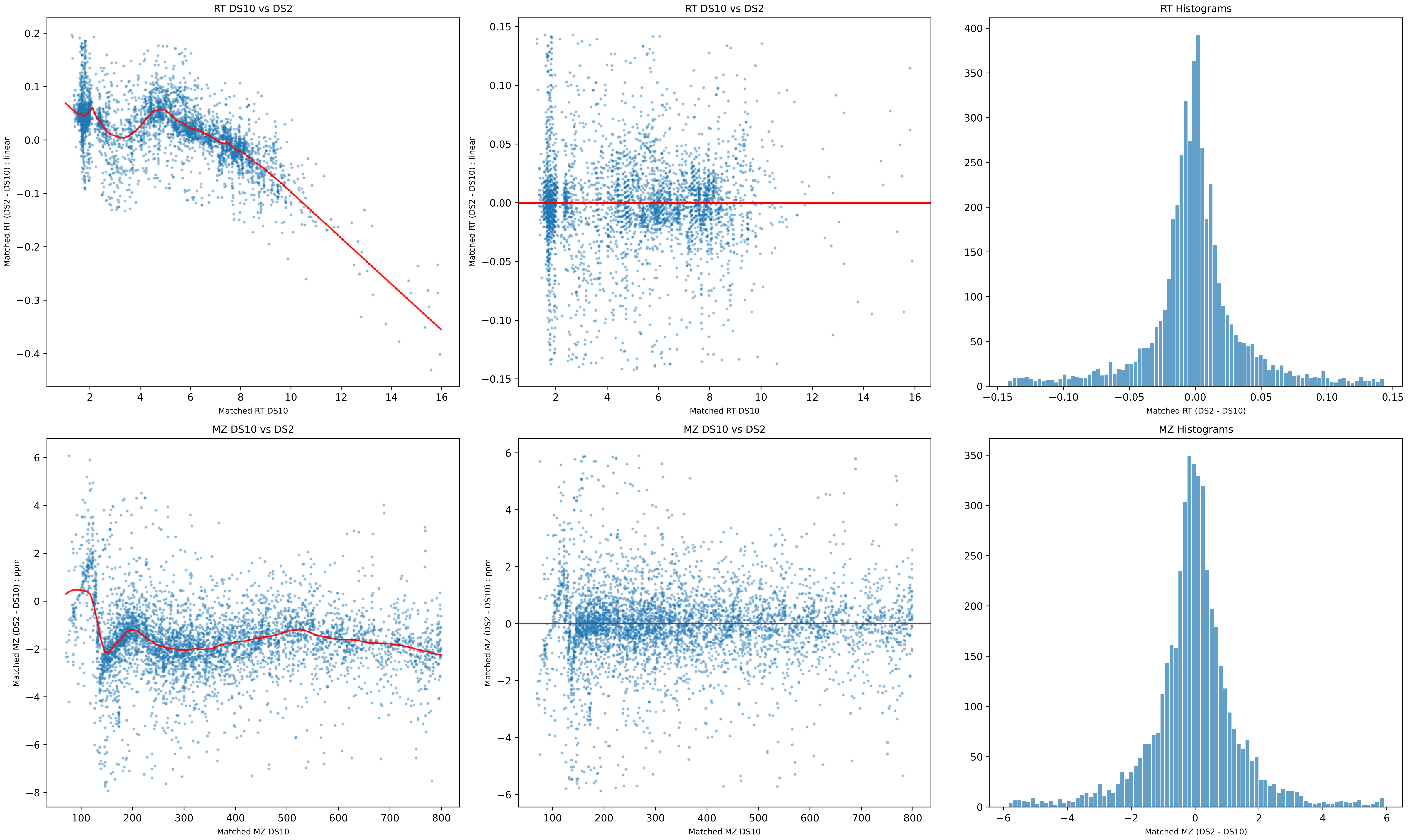
RT DS10 vs DS2
Matched RT (DS2 - DS10) : linear
Matched RT DS10
RT DS10 vs DS2
Matched RT (DS2 - DS10) : linear
Matched RT DS10
RT Histograms
Matched RT (DS2 - DS10)
MZ DS10 vs DS2
Matched MZ (DS2 - DS10) : ppm
Matched MZ DS10
MZ DS10 vs DS2
Matched MZ (DS2 - DS10) : ppm
Matched MZ DS10
MZ Histograms
Matched MZ (DS2 - DS10)

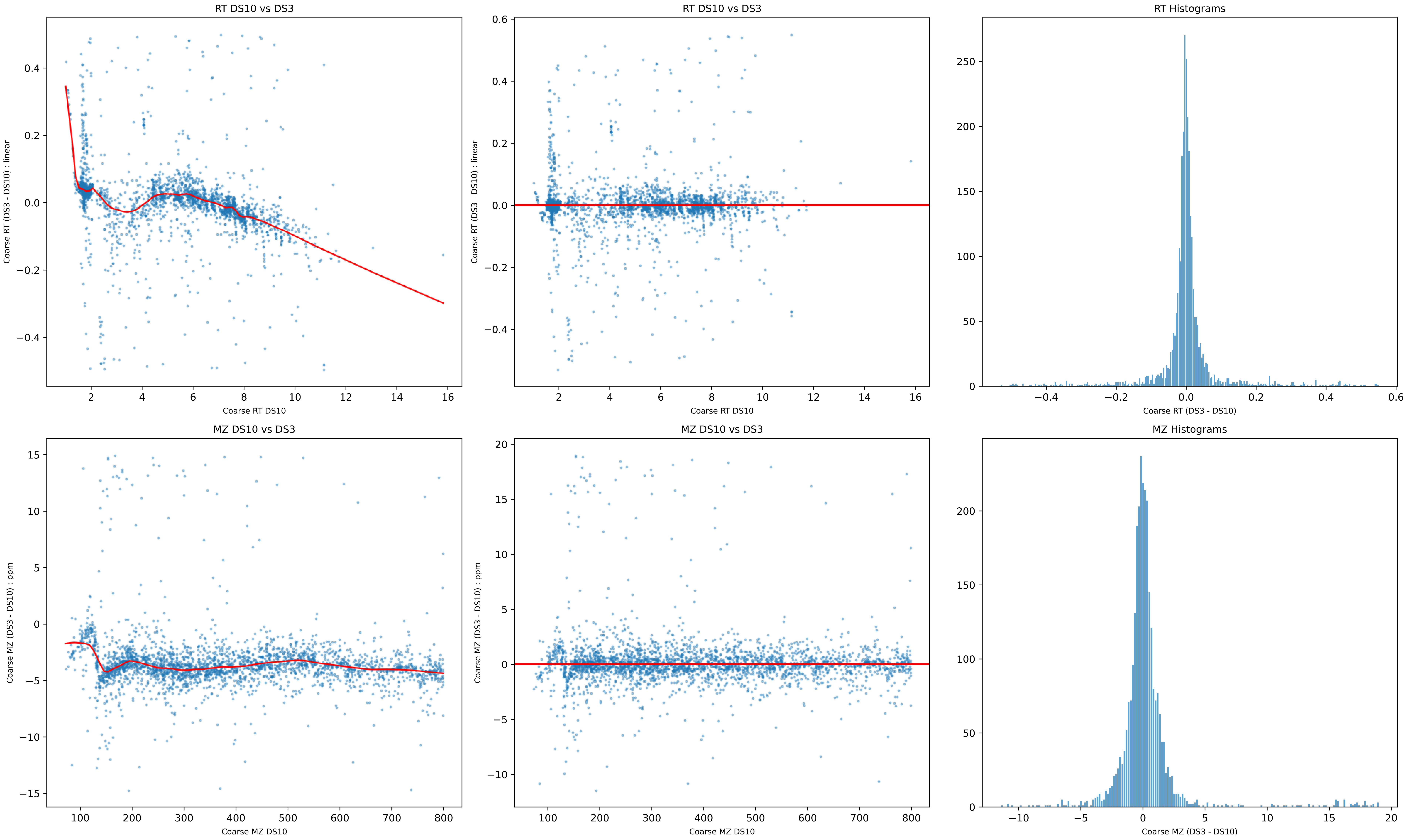

RT DS10 vs DS3
Coarse RT (DS3 - DS10) : linear
Coarse RT DS10
RT DS10 vs DS3
Coarse RT (DS3 - DS10) : linear
Coarse RT DS10
RT Histograms
Coarse RT (DS3 - DS10)
MZ DS10 vs DS3
Coarse MZ (DS3 - DS10) : ppm
Coarse MZ DS10
MZ DS10 vs DS3
Coarse MZ (DS3 - DS10) : ppm
Coarse MZ DS10
MZ Histograms
Coarse MZ (DS3 - DS10)

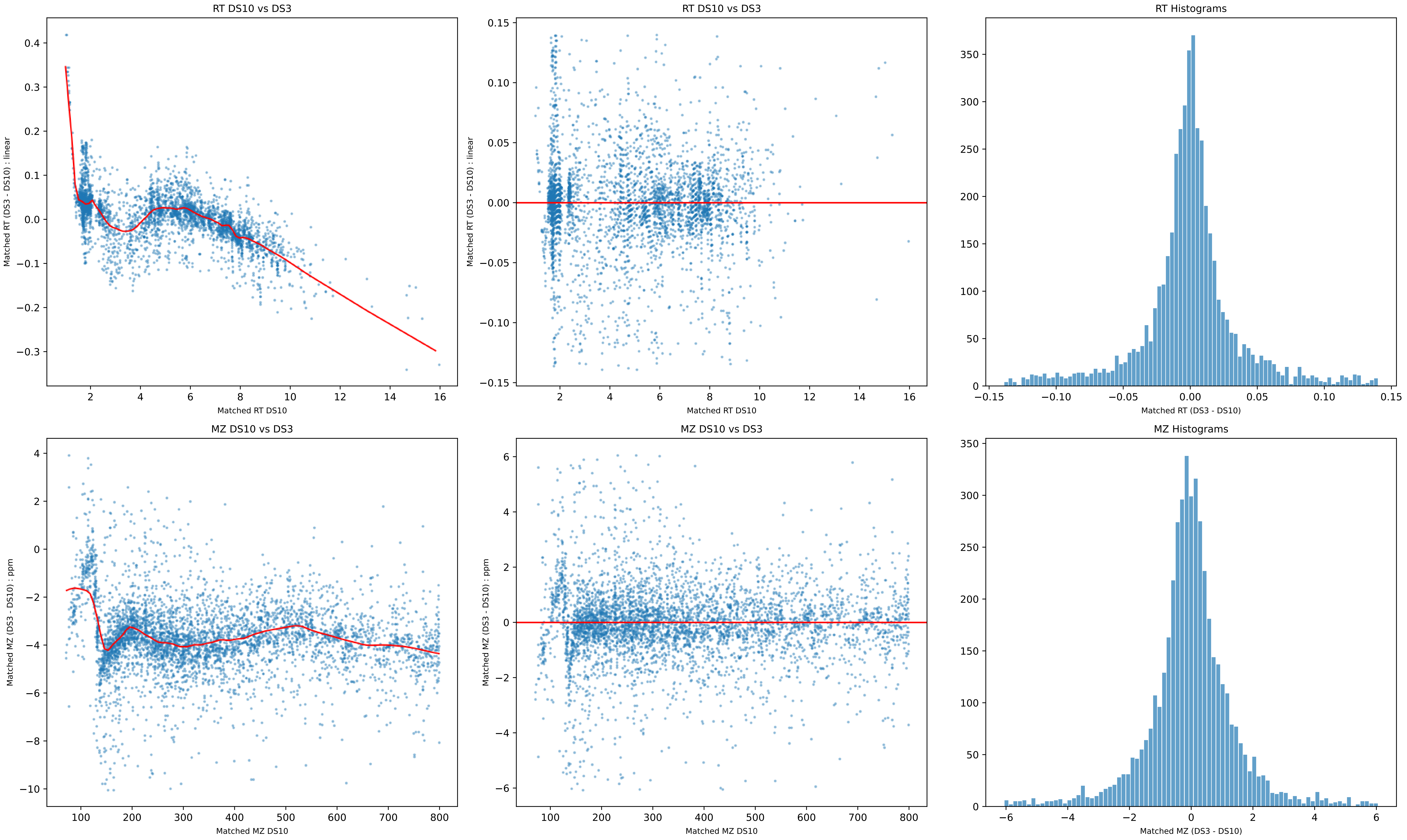

RT DS10 vs DS3
Matched RT (DS3 - DS10) : linear
Matched RT DS10
RT DS10 vs DS3
Matched RT (DS3 - DS10) : linear
Matched RT DS10
RT Histograms
Matched RT (DS3 - DS10)
MZ DS10 vs DS3
Matched MZ (DS3 - DS10) : ppm
Matched MZ DS10
MZ DS10 vs DS3
Matched MZ (DS3 - DS10) : ppm
Matched MZ DS10
MZ Histograms
Matched MZ (DS3 - DS10)

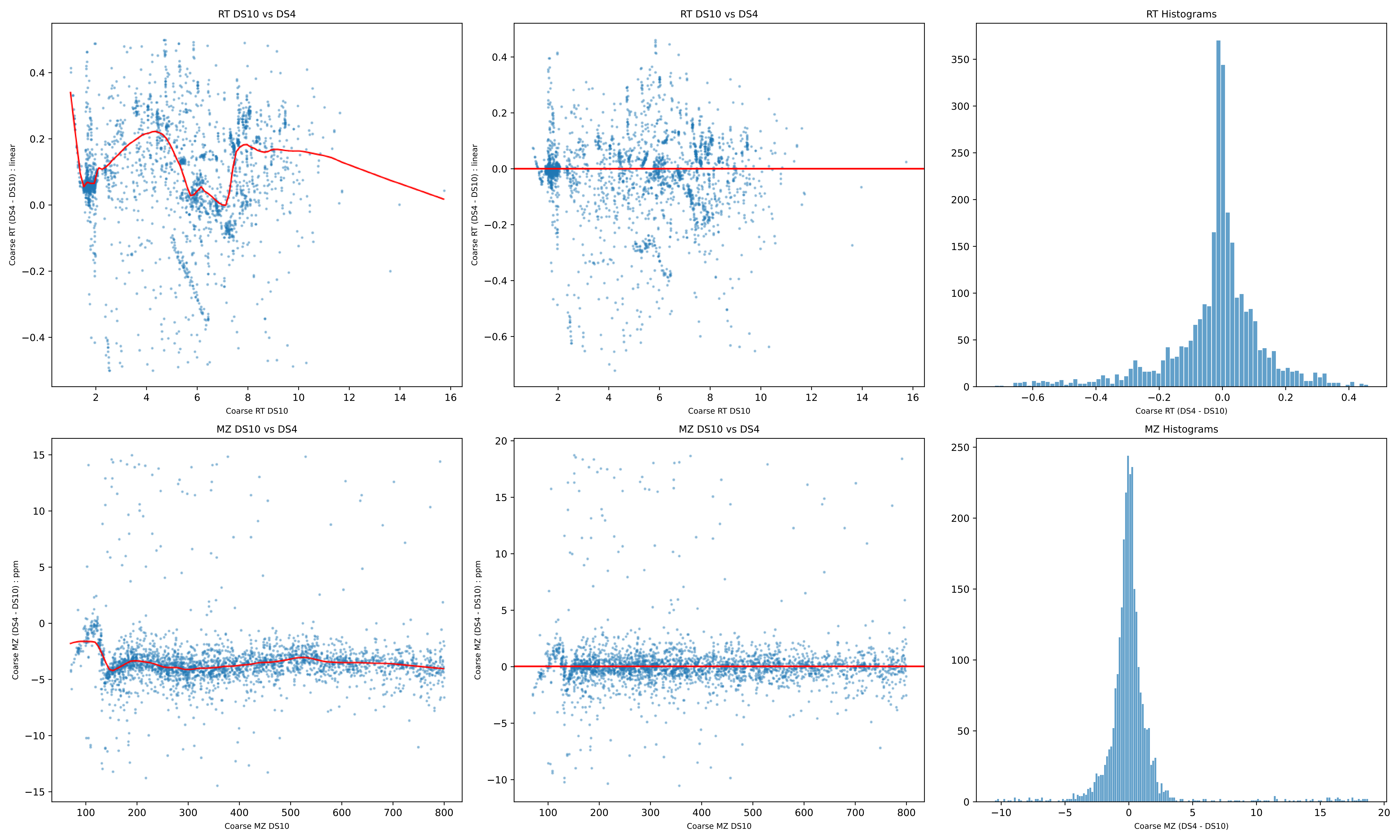
RT DS10 vs DS4
Coarse RT DS10
Coarse RT (DS4 - DS10) : linear
RT DS10 vs DS4
Coarse RT DS10
Coarse RT (DS4 - DS10) : linear
RT Histograms
Coarse RT (DS4 - DS10)
MZ DS10 vs DS4
Coarse MZ DS10
Coarse MZ (DS4 - DS10) : ppm
MZ DS10 vs DS4
Coarse MZ DS10
Coarse MZ (DS4 - DS10) : ppm
MZ Histograms
Coarse MZ (DS4 - DS10)

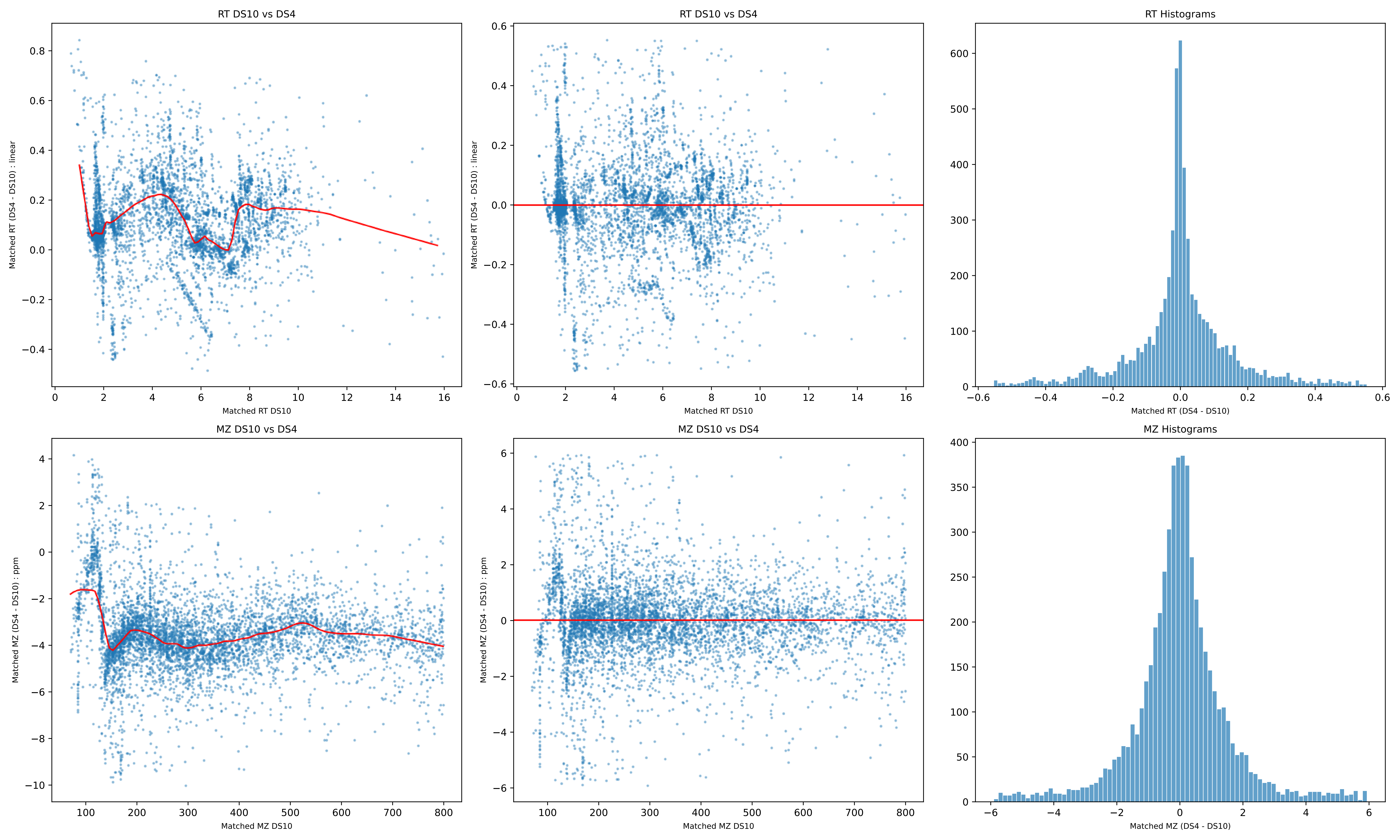

RT DS10 vs DS4
Matched RT (DS4 - DS10) : linear
Matched RT DS10
RT DS10 vs DS4
Matched RT (DS4 - DS10) : linear
Matched RT DS10
RT Histograms
Matched RT (DS4 - DS10)
MZ DS10 vs DS4
Matched MZ (DS4 - DS10) : ppm
Matched MZ DS10
MZ DS10 vs DS4
Matched MZ (DS4 - DS10) : ppm
Matched MZ DS10
MZ Histograms
Matched MZ (DS4 - DS10)

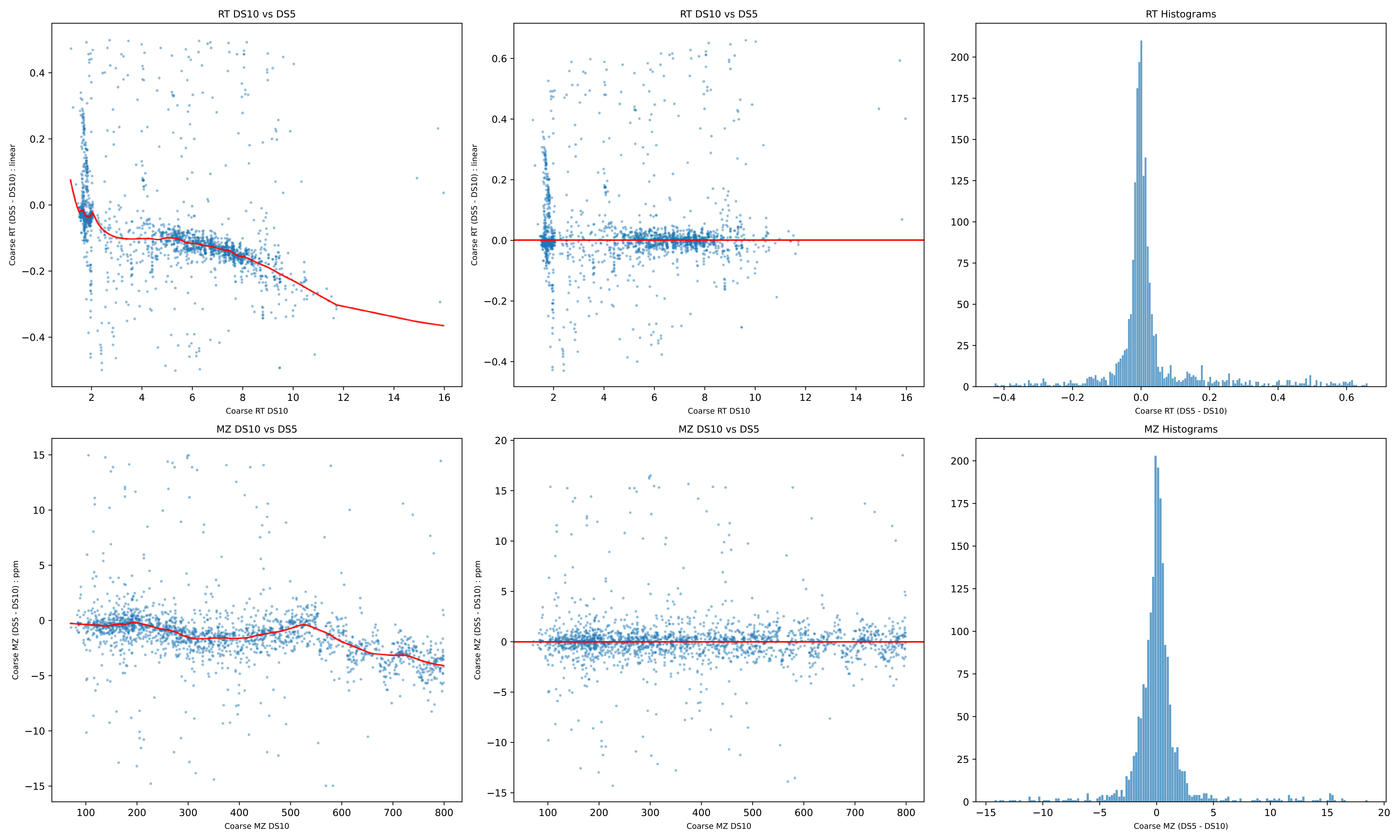

RT DS10 vs DS5
Coarse RT (DS5 - DS10) : linear
Coarse RT DS10
RT DS10 vs DS5
Coarse RT (DS5 - DS10) : linear
Coarse RT DS10
RT Histograms
Coarse RT (DS5 - DS10)
MZ DS10 vs DS5
Coarse MZ (DS5 - DS10) : ppm
Coarse MZ DS10
MZ DS10 vs DS5
Coarse MZ (DS5 - DS10) : ppm
Coarse MZ DS10
MZ Histograms
Coarse MZ (DS5 - DS10)

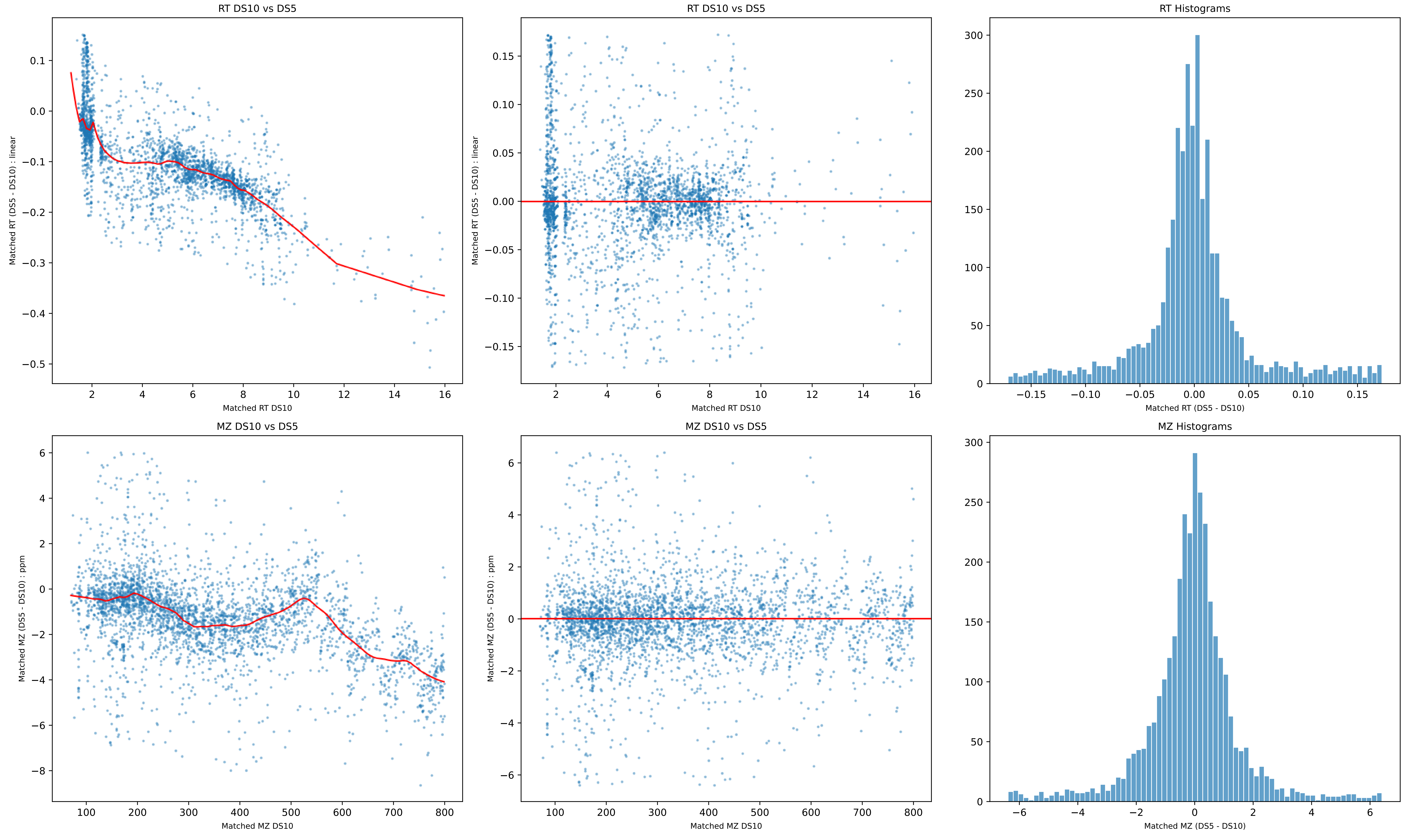

RT DS10 vs DS5
Matched RT (DS5 - DS10) : linear
Matched RT DS10
RT DS10 vs DS5
Matched RT (DS5 - DS10) : linear
Matched RT DS10
RT Histograms
Matched RT (DS5 - DS10)
MZ DS10 vs DS5
Matched MZ (DS5 - DS10) : ppm
Matched MZ DS10
MZ DS10 vs DS5
Matched MZ (DS5 - DS10) : ppm
Matched MZ DS10
MZ Histograms
Matched MZ (DS5 - DS10)

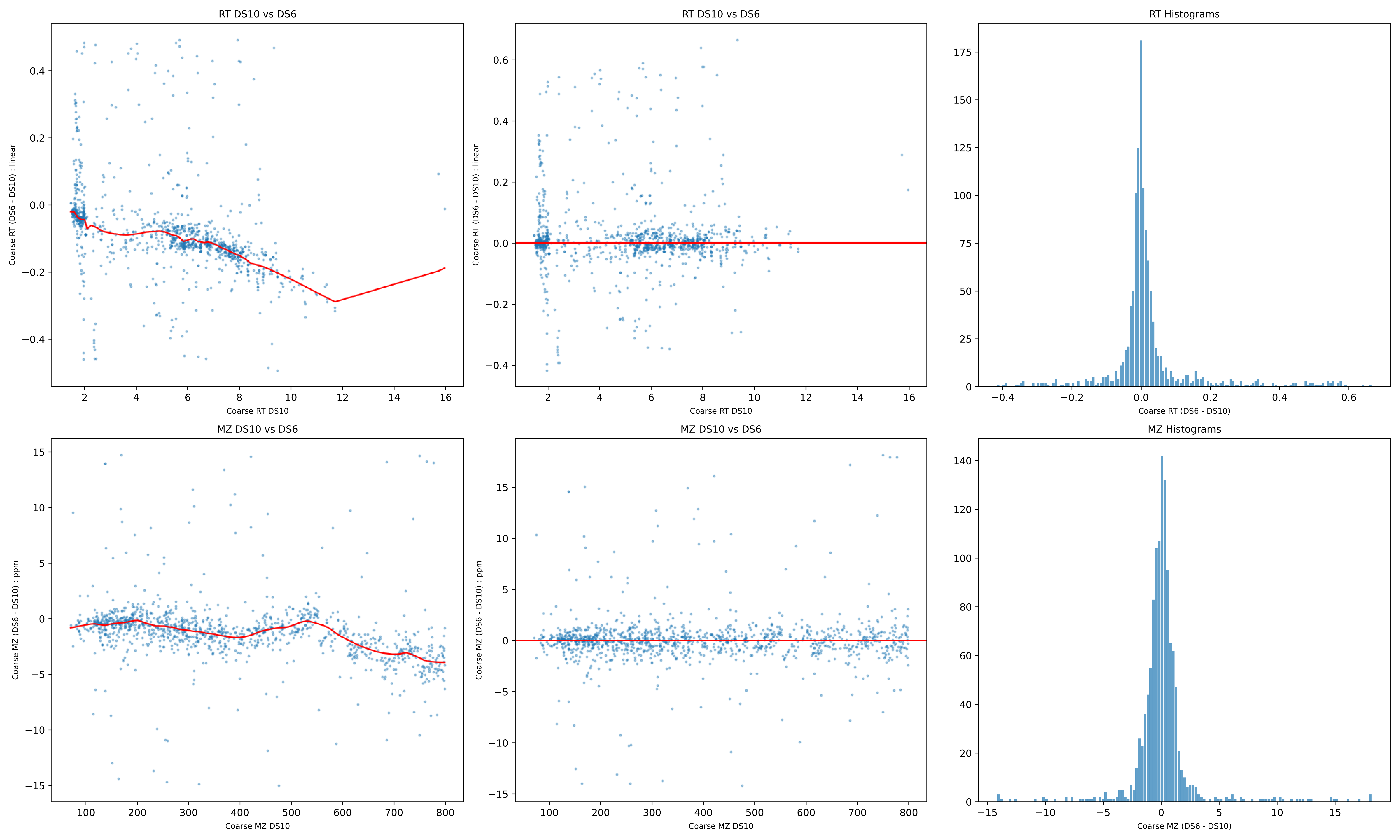

RT DS10 vs DS6
Coarse RT (DS6 - DS10) : linear
Coarse RT DS10
RT DS10 vs DS6
Coarse RT (DS6 - DS10) : linear
Coarse RT DS10
RT Histograms
Coarse RT (DS6 - DS10)
MZ DS10 vs DS6
Coarse MZ (DS6 - DS10) : ppm
Coarse MZ DS10
MZ DS10 vs DS6
Coarse MZ (DS6 - DS10) : ppm
Coarse MZ DS10
MZ Histograms
Coarse MZ (DS6 - DS10)

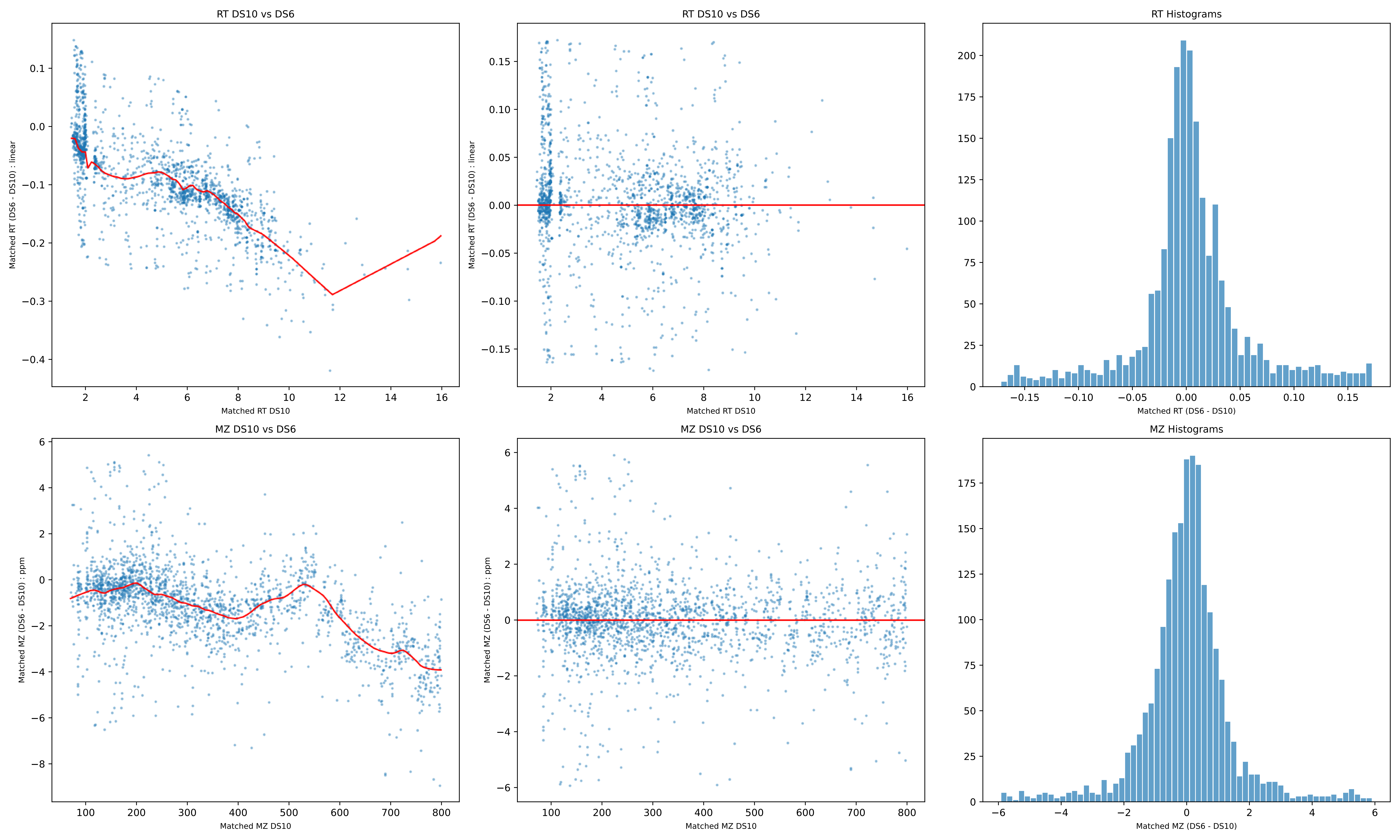
RT DS10 vs DS6
Matched RT (DS6 - DS10) : linear
Matched RT DS10
RT DS10 vs DS6
Matched RT (DS6 - DS10) : linear
Matched RT DS10
RT Histograms
Matched RT (DS6 - DS10)
MZ DS10 vs DS6
Matched MZ (DS6 - DS10) : ppm
Matched MZ DS10
MZ DS10 vs DS6
Matched MZ (DS6 - DS10) : ppm
Matched MZ DS10
MZ Histograms
Matched MZ (DS6 - DS10)

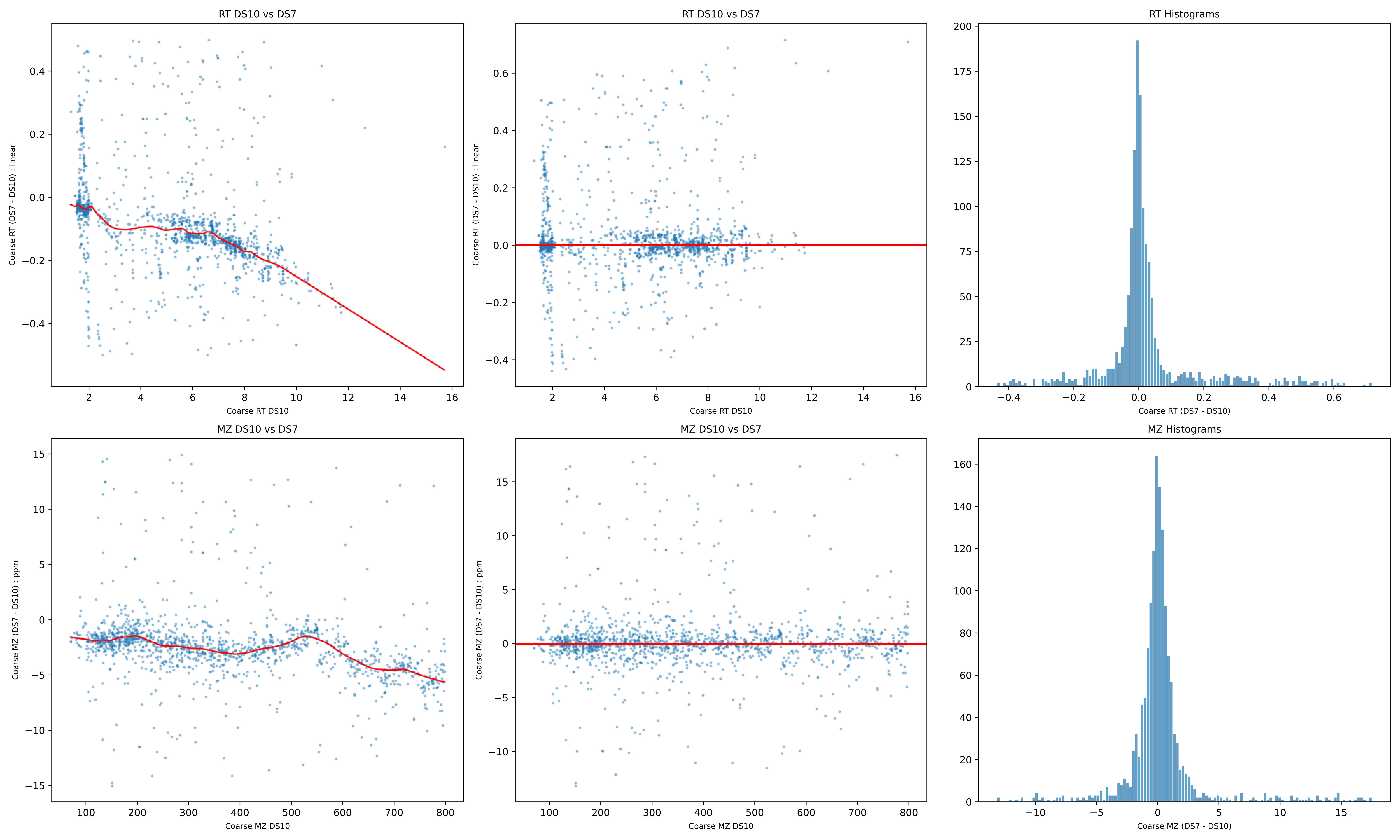
RT DS10 vs DS7
Coarse RT (DS7 - DS10) : linear
Coarse RT DS10
RT DS10 vs DS7
Coarse RT (DS7 - DS10) : linear
Coarse RT DS10
RT Histograms
Coarse RT (DS7 - DS10)
MZ DS10 vs DS7
Coarse MZ (DS7 - DS10) : ppm
Coarse MZ DS10
MZ DS10 vs DS7
Coarse MZ (DS7 - DS10) : ppm
Coarse MZ DS10
MZ Histograms
Coarse MZ (DS7 - DS10)

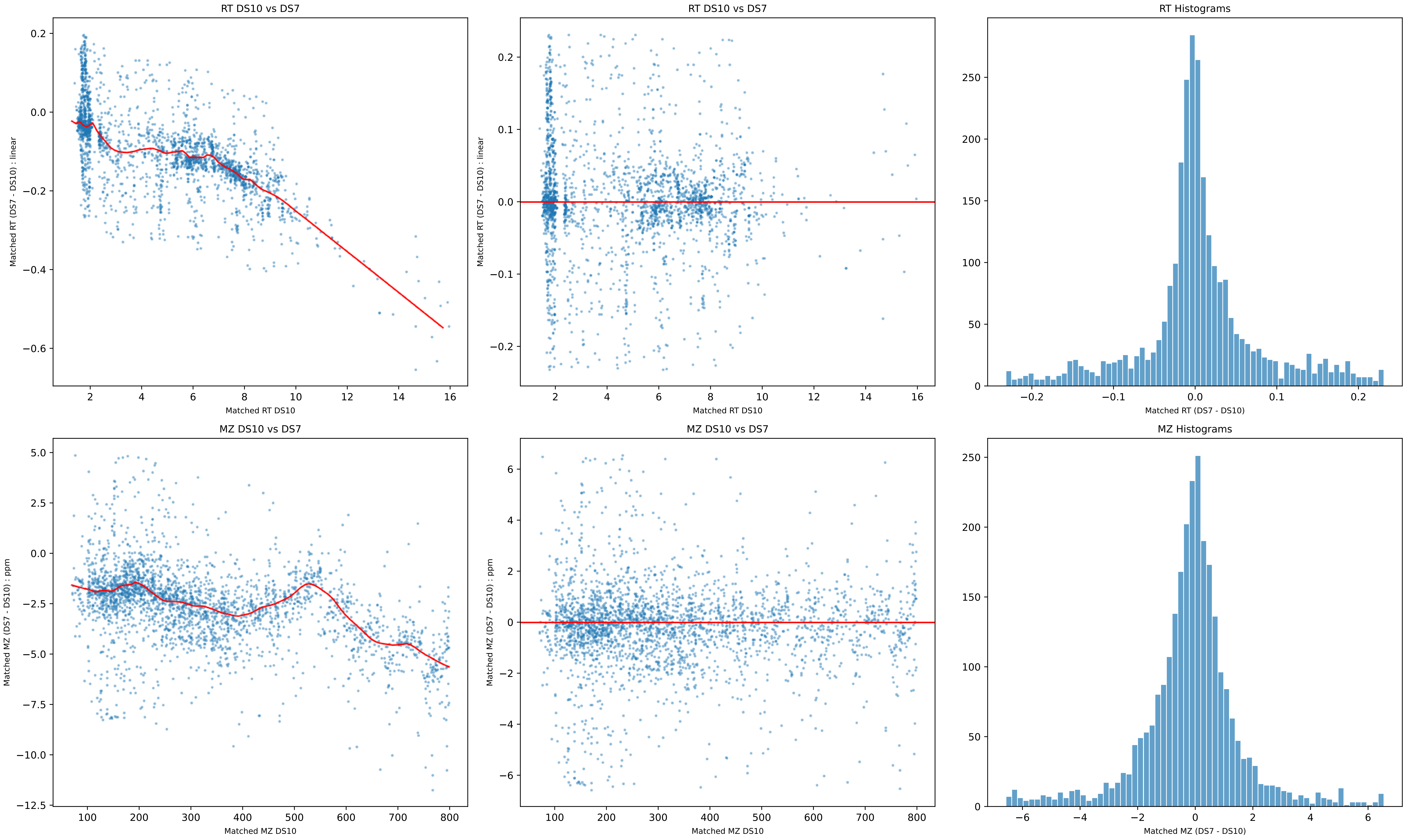
RT DS10 vs DS7
Matched RT (DS7 - DS10) : linear
Matched RT DS10
RT DS10 vs DS7
Matched RT (DS7 - DS10) : linear
Matched RT DS10
RT Histograms
Matched RT (DS7 - DS10)
MZ DS10 vs DS7
Matched MZ (DS7 - DS10) : ppm
Matched MZ DS10
MZ DS10 vs DS7
Matched MZ (DS7 - DS10) : ppm
Matched MZ DS10
MZ Histograms
Matched MZ (DS7 - DS10)

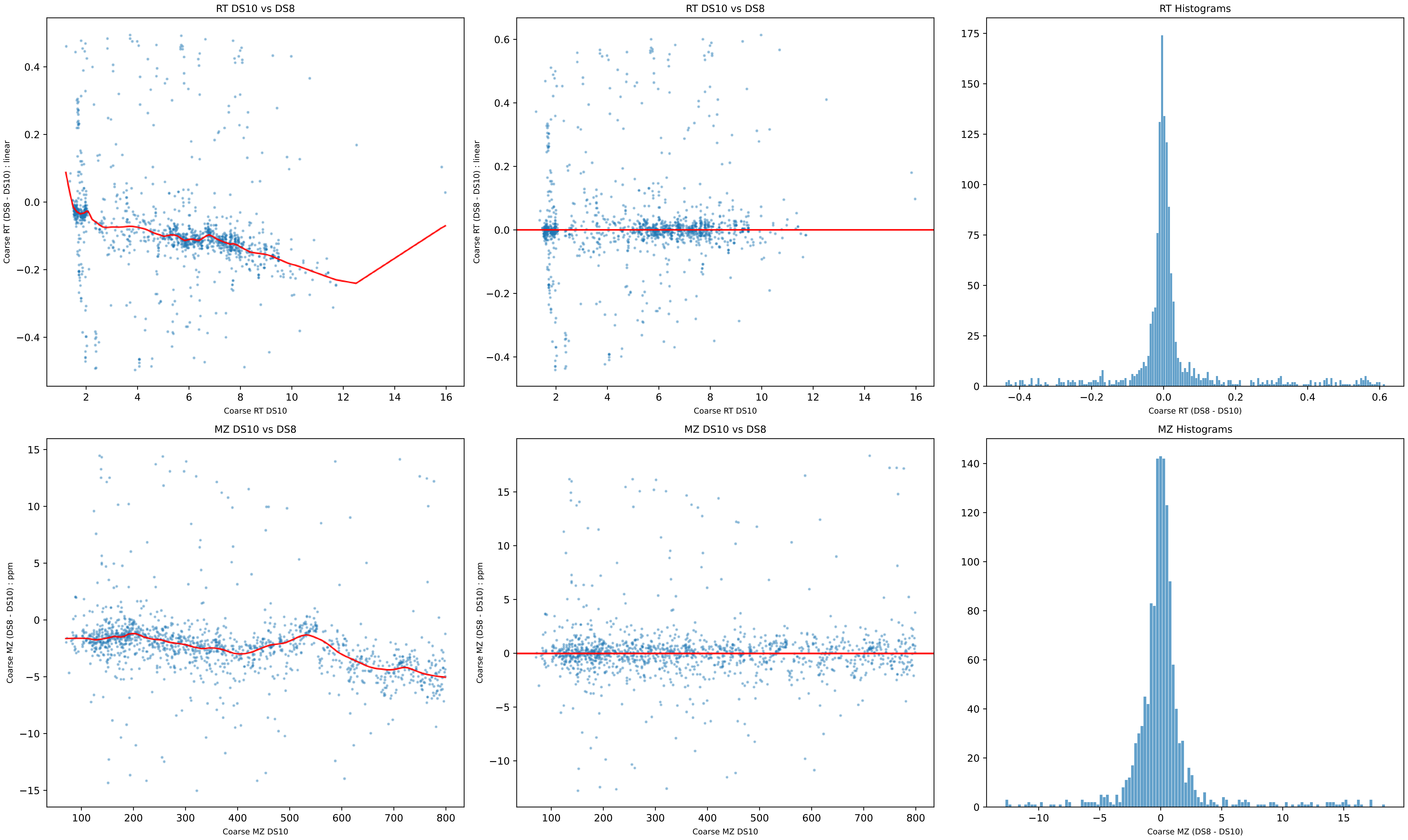

RT DS10 vs DS8
Coarse RT (DS8 - DS10) : linear
Coarse RT DS10
RT DS10 vs DS8
Coarse RT (DS8 - DS10) : linear
Coarse RT DS10
RT Histograms
Coarse RT (DS8 - DS10)
MZ DS10 vs DS8
Coarse MZ (DS8 - DS10) : ppm
Coarse MZ DS10
MZ DS10 vs DS8
Coarse MZ (DS8 - DS10) : ppm
Coarse MZ DS10
MZ Histograms
Coarse MZ (DS8 - DS10)

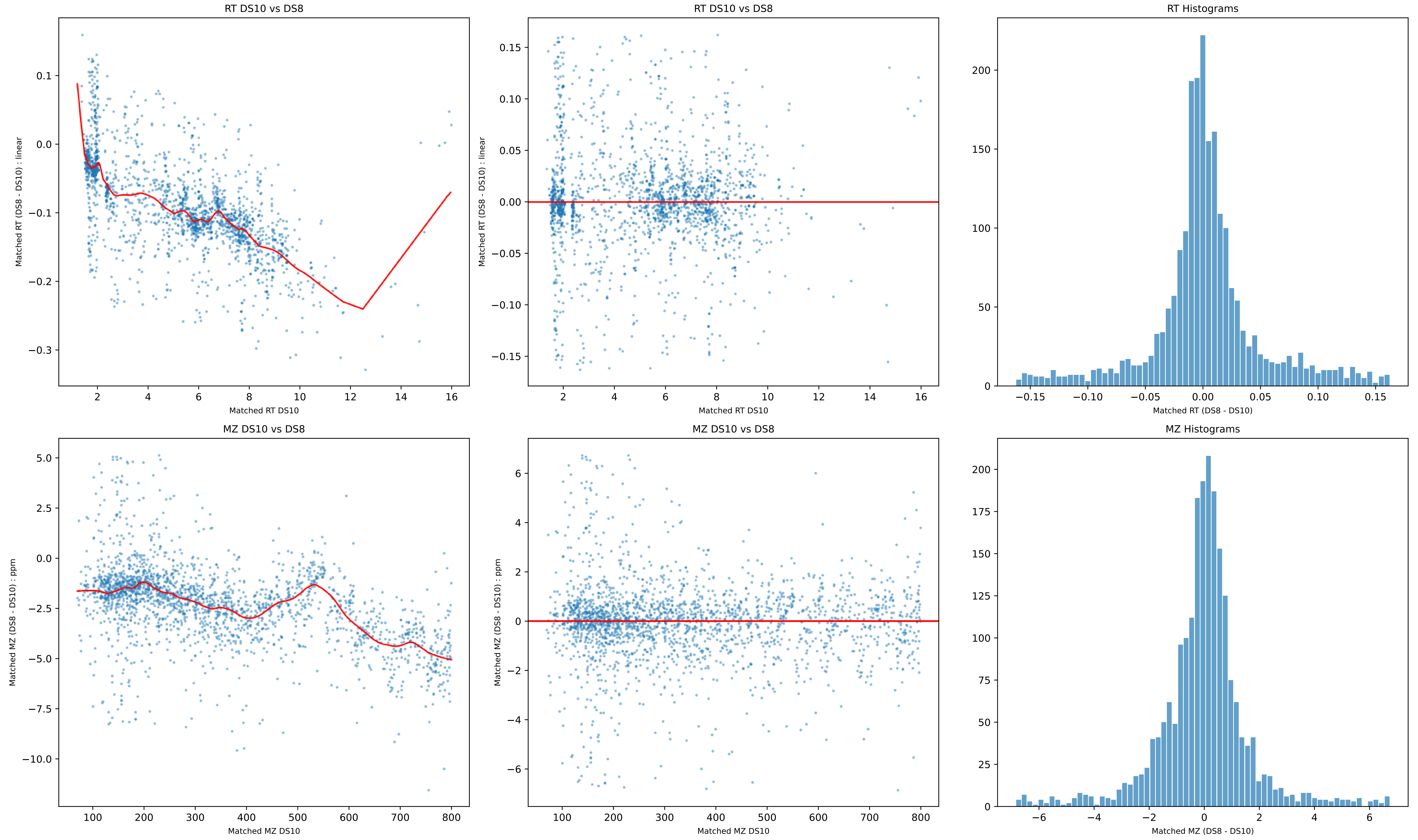

RT DS10 vs DS8
Matched RT (DS8 - DS10) : linear
Matched RT DS10
RT DS10 vs DS8
Matched RT (DS8 - DS10) : linear
Matched RT DS10
RT Histograms
Matched RT (DS8 - DS10)
MZ DS10 vs DS8
Matched MZ (DS8 - DS10) : ppm
Matched MZ DS10
MZ DS10 vs DS8
Matched MZ (DS8 - DS10) : ppm
Matched MZ DS10
MZ Histograms
Matched MZ (DS8 - DS10)

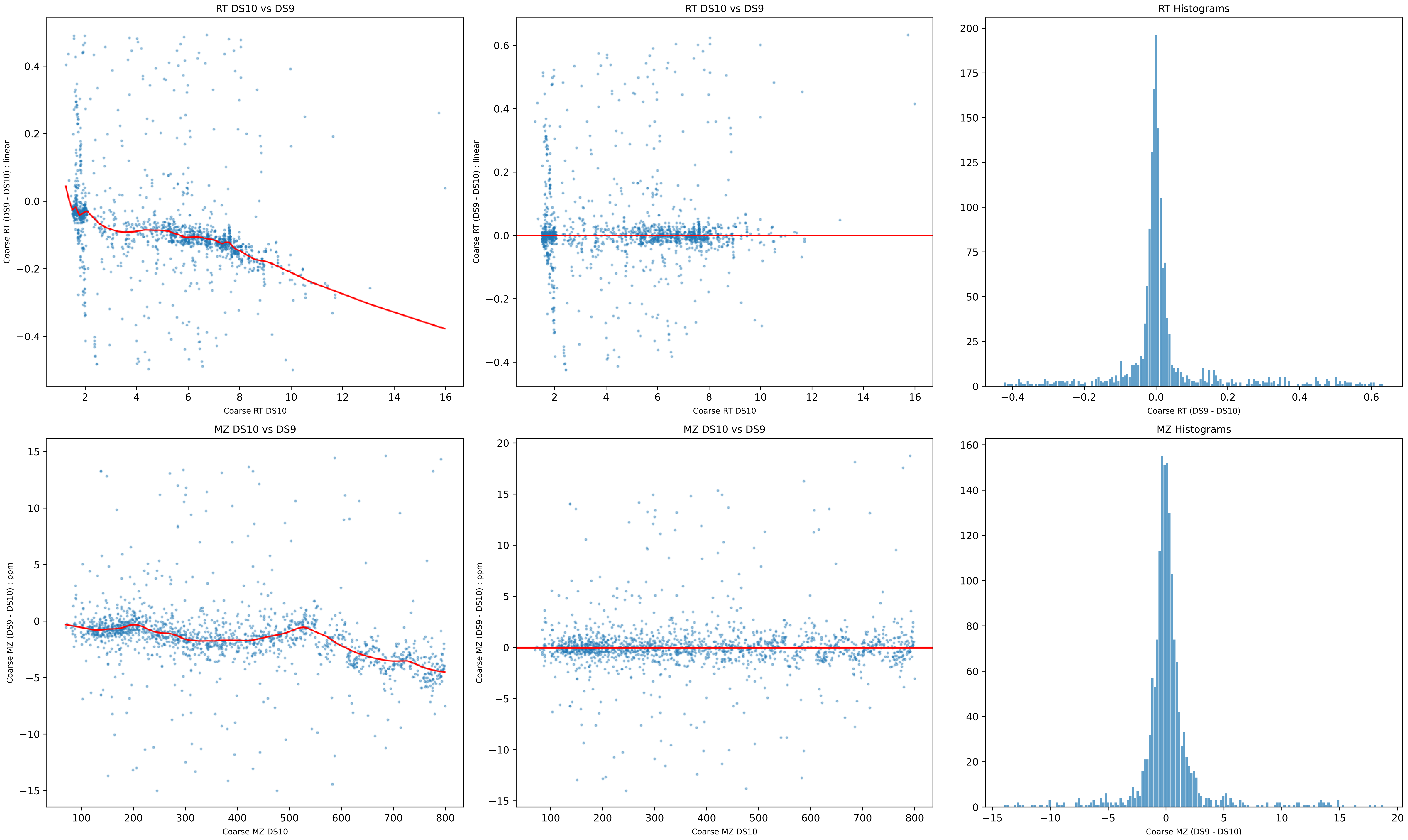
RT DS10 vs DS9
Coarse RT (DS9 - DS10) : linear
Coarse RT DS10
RT DS10 vs DS9
Coarse RT (DS9 - DS10) : linear
Coarse RT DS10
RT Histograms
Coarse RT (DS9 - DS10)
MZ DS10 vs DS9
Coarse MZ (DS9 - DS10) : ppm
Coarse MZ DS10
MZ DS10 vs DS9
Coarse MZ (DS9 - DS10) : ppm
Coarse MZ DS10
MZ Histograms
Coarse MZ (DS9 - DS10)

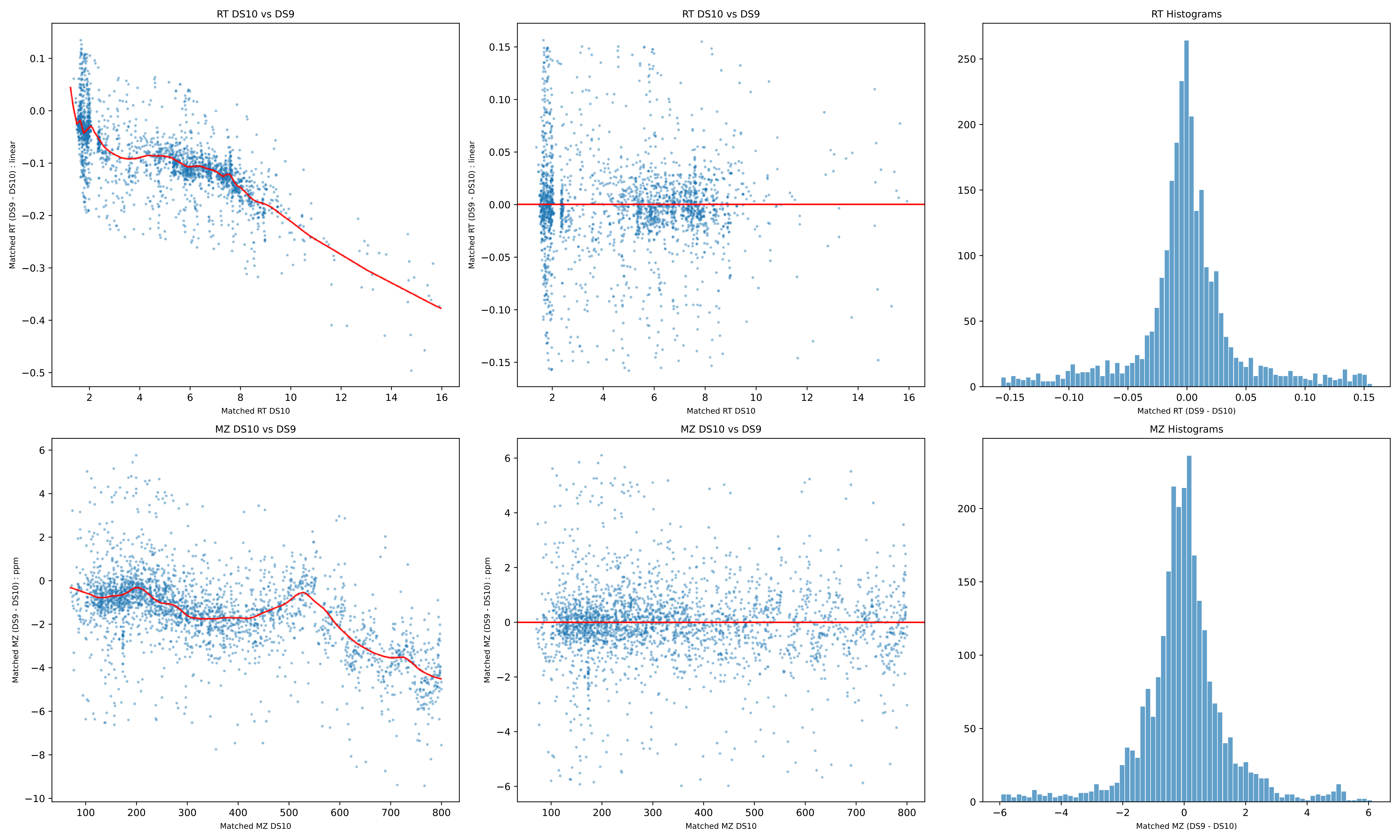
RT DS10 vs DS9
Matched RT (DS9 - DS10) : linear
Matched RT DS10
RT DS10 vs DS9
Matched RT (DS9 - DS10) : linear
Matched RT DS10
RT Histograms
Matched RT (DS9 - DS10)
MZ DS10 vs DS9
Matched MZ (DS9 - DS10) : ppm
Matched MZ DS10
MZ DS10 vs DS9
Matched MZ (DS9 - DS10) : ppm
Matched MZ DS10
MZ Histograms
Matched MZ (DS9 - DS10)

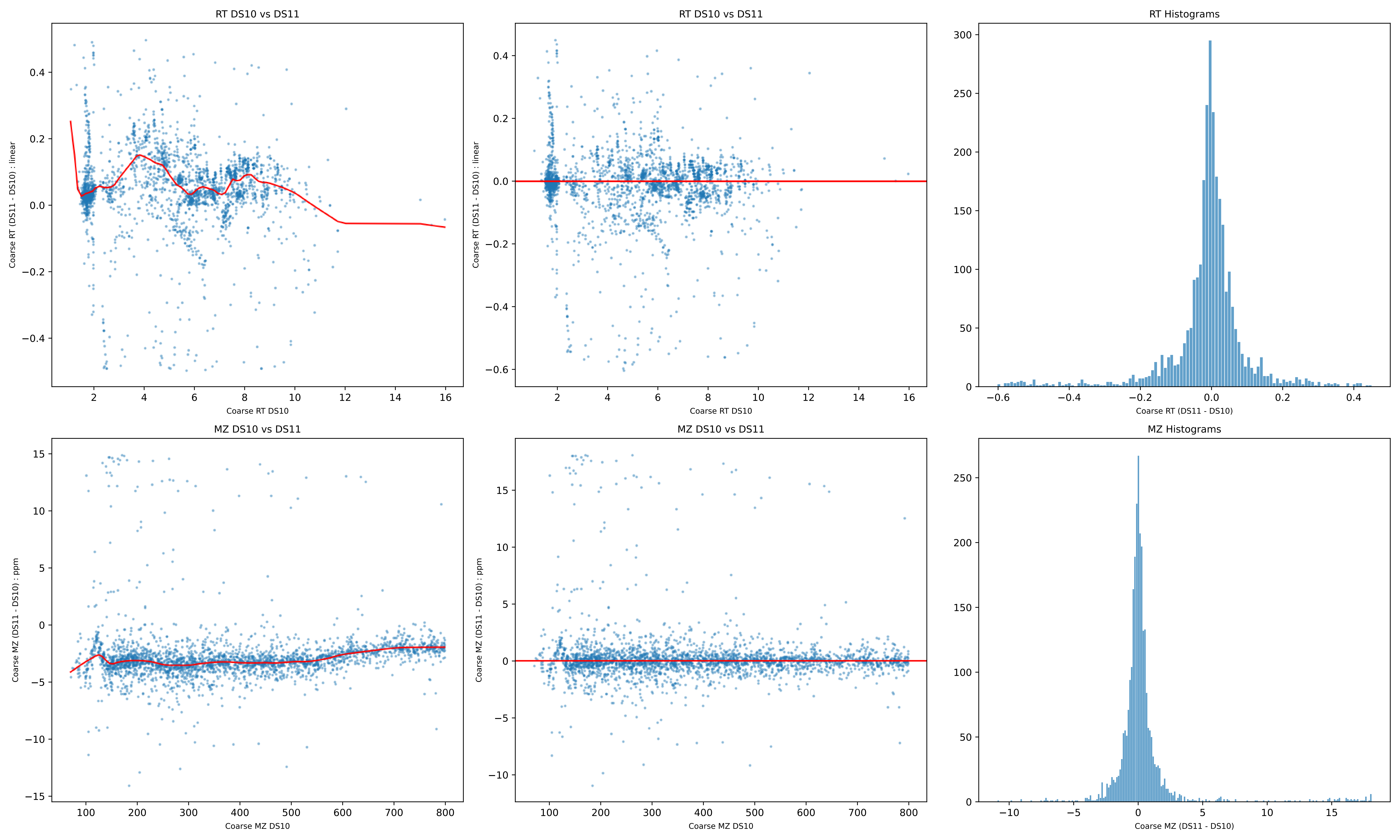
RT DS10 vs DS11
Coarse RT (DS11 - DS10) : linear
Coarse RT DS10
RT DS10 vs DS11
Coarse RT (DS11 - DS10) : linear
Coarse RT DS10
RT Histograms
Coarse RT (DS11 - DS10)
MZ DS10 vs DS11
Coarse MZ (DS11 - DS10) : ppm
Coarse MZ DS10
MZ DS10 vs DS11
Coarse MZ (DS11 - DS10) : ppm
Coarse MZ DS10
MZ Histograms
Coarse MZ (DS11 - DS10)

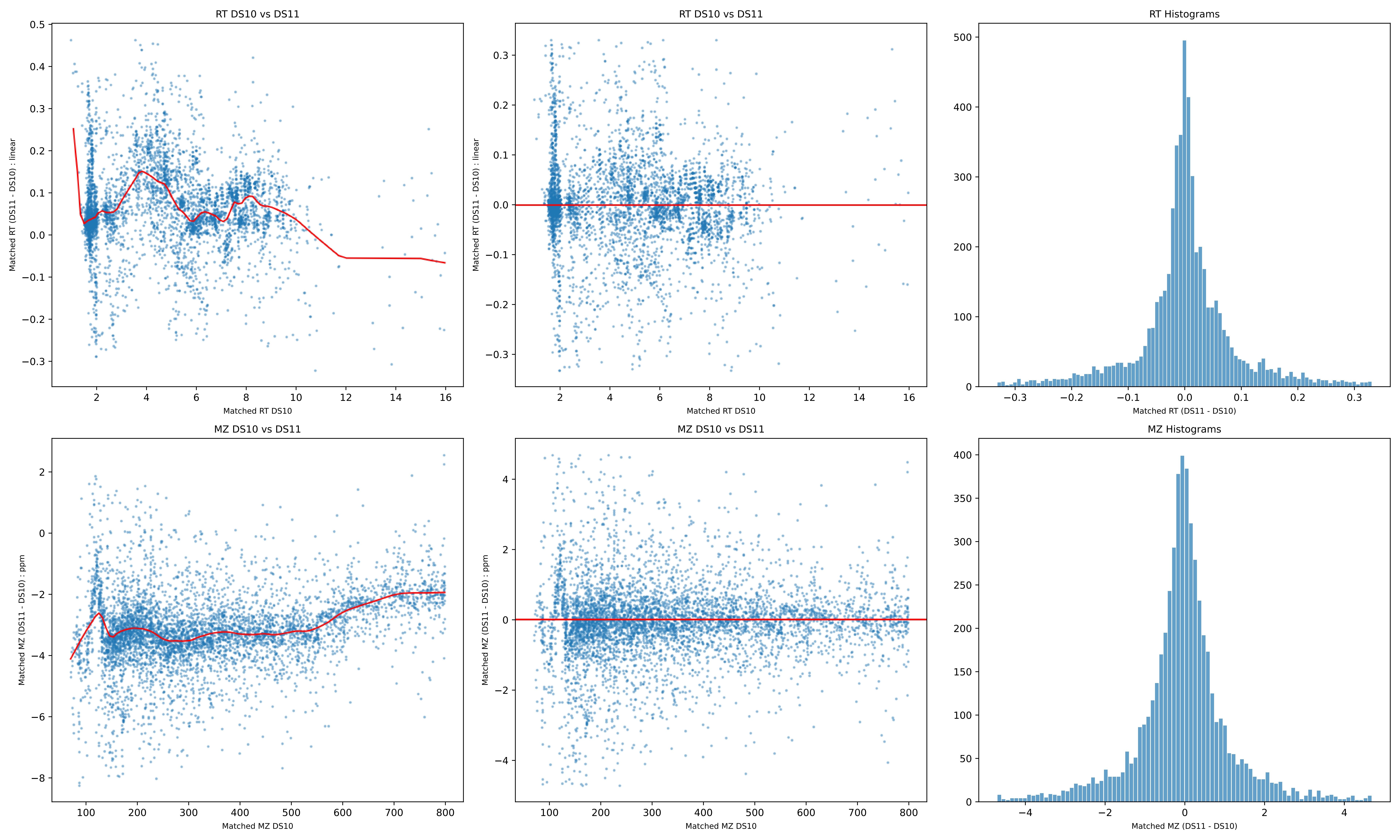
RT DS10 vs DS11
Matched RT (DS11 - DS10) : linear
Matched RT DS10
RT DS10 vs DS11
Matched RT (DS11 - DS10) : linear
Matched RT DS10
RT Histograms
Matched RT (DS11 - DS10)
MZ DS10 vs DS11
Matched MZ (DS11 - DS10) : ppm
Matched MZ DS10
MZ DS10 vs DS11
Matched MZ (DS11 - DS10) : ppm
Matched MZ DS10
MZ Histograms
Matched MZ (DS11 - DS10)

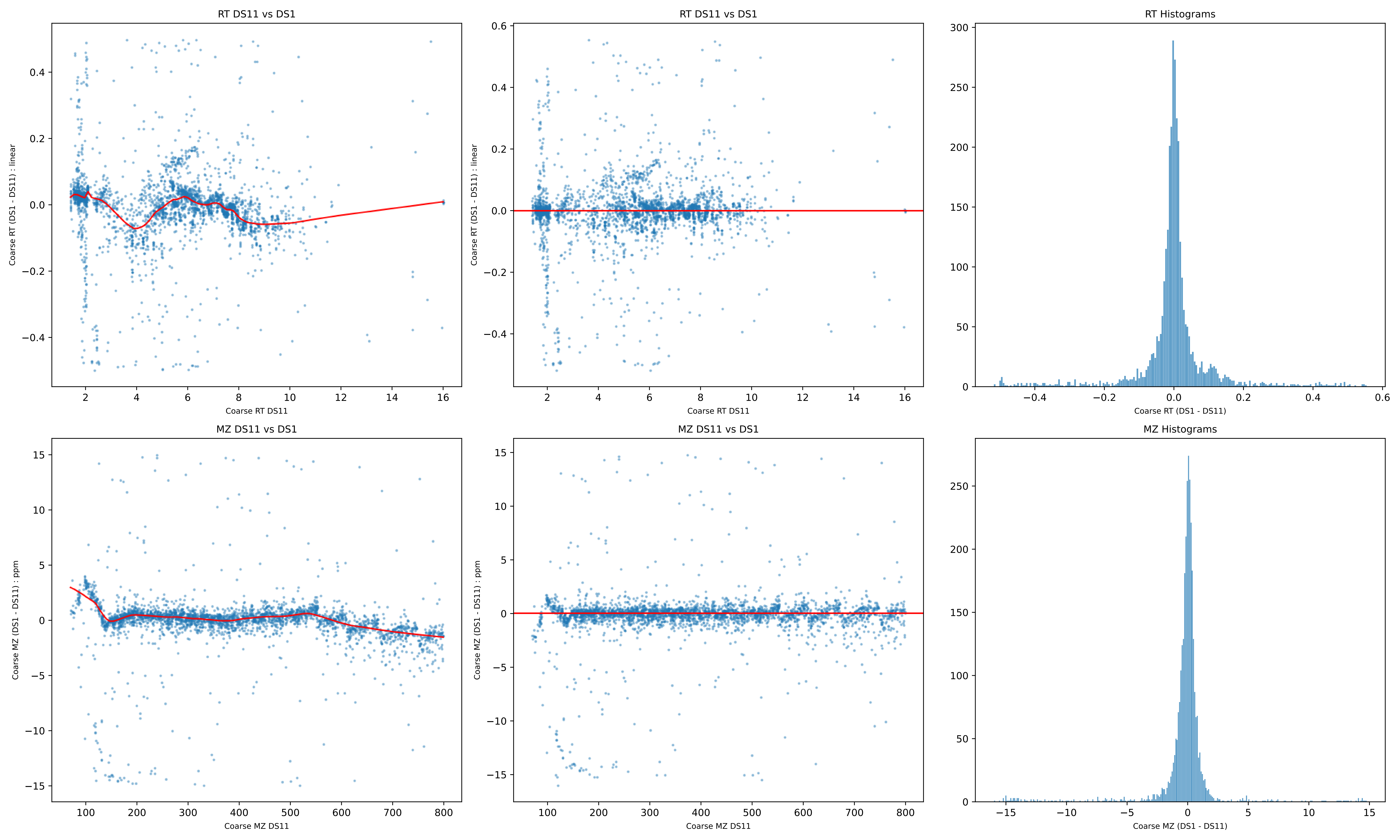
RT DS11 vs DS1
Coarse RT (DS1 - DS11) : linear
Coarse RT DS11
RT DS11 vs DS1
Coarse RT (DS1 - DS11) : linear
Coarse RT DS11
RT Histograms
Coarse RT (DS1 - DS11)
MZ DS11 vs DS1
Coarse MZ (DS1 - DS11) : ppm
Coarse MZ DS11
MZ DS11 vs DS1
Coarse MZ (DS1 - DS11) : ppm
Coarse MZ DS11
MZ Histograms
Coarse MZ (DS1 - DS11)

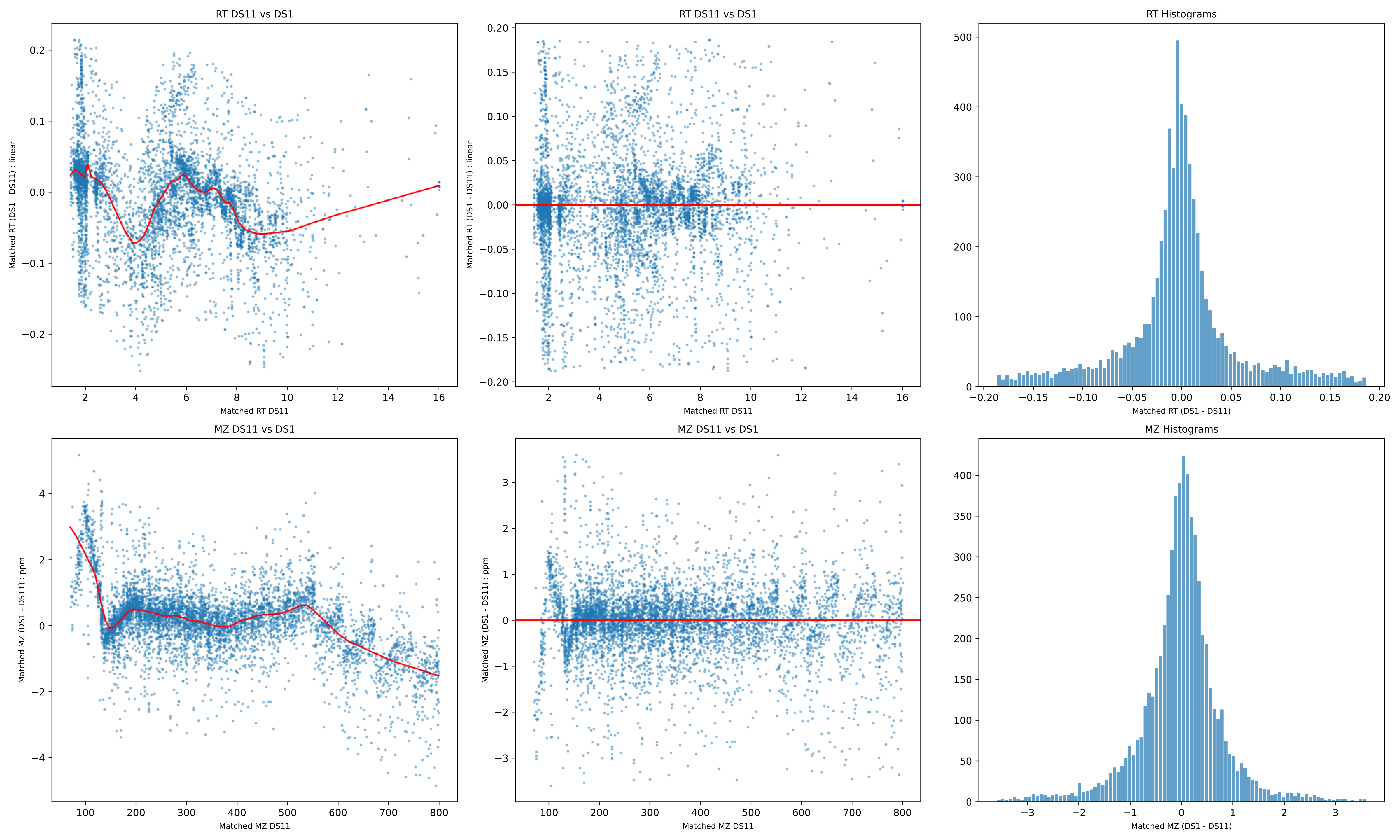

RT DS11 vs DS1
Matched RT (DS1 - DS11) : linear
Matched RT DS11
RT DS11 vs DS1
Matched RT (DS1 - DS11) : linear
Matched RT DS11
RT Histograms
Matched RT (DS1 - DS11)
MZ DS11 vs DS1
Matched MZ (DS1 - DS11) : ppm
Matched MZ DS11
MZ DS11 vs DS1
Matched MZ (DS1 - DS11) : ppm
Matched MZ DS11
MZ Histograms
Matched MZ (DS1 - DS11)

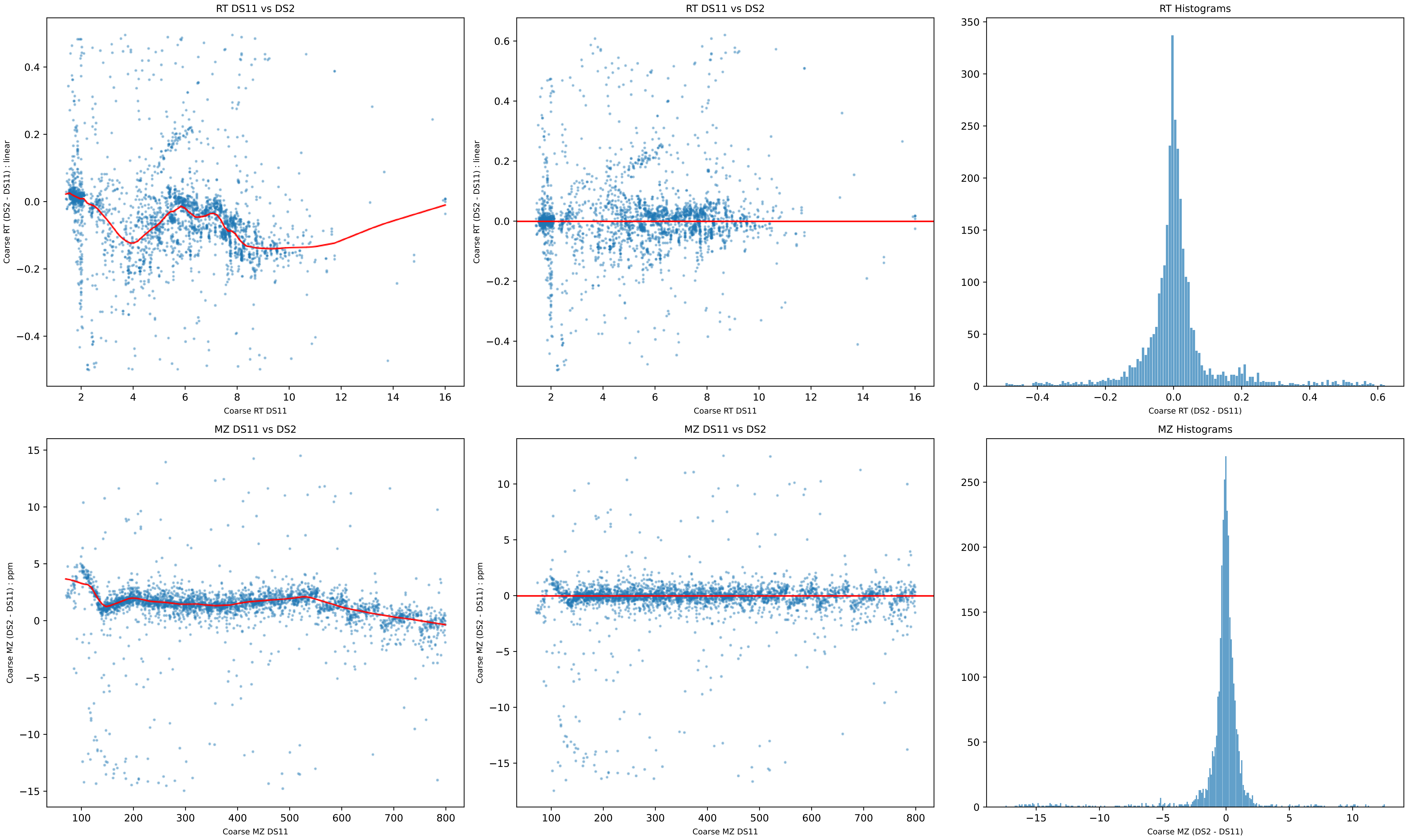
RT DS11 vs DS2
Coarse RT (DS2 - DS11) : linear
Coarse RT DS11
RT DS11 vs DS2
Coarse RT (DS2 - DS11) : linear
Coarse RT DS11
RT Histograms
Coarse RT (DS2 - DS11)
MZ DS11 vs DS2
Coarse MZ (DS2 - DS11) : ppm
Coarse MZ DS11
MZ DS11 vs DS2
Coarse MZ (DS2 - DS11) : ppm
Coarse MZ DS11
MZ Histograms
Coarse MZ (DS2 - DS11)

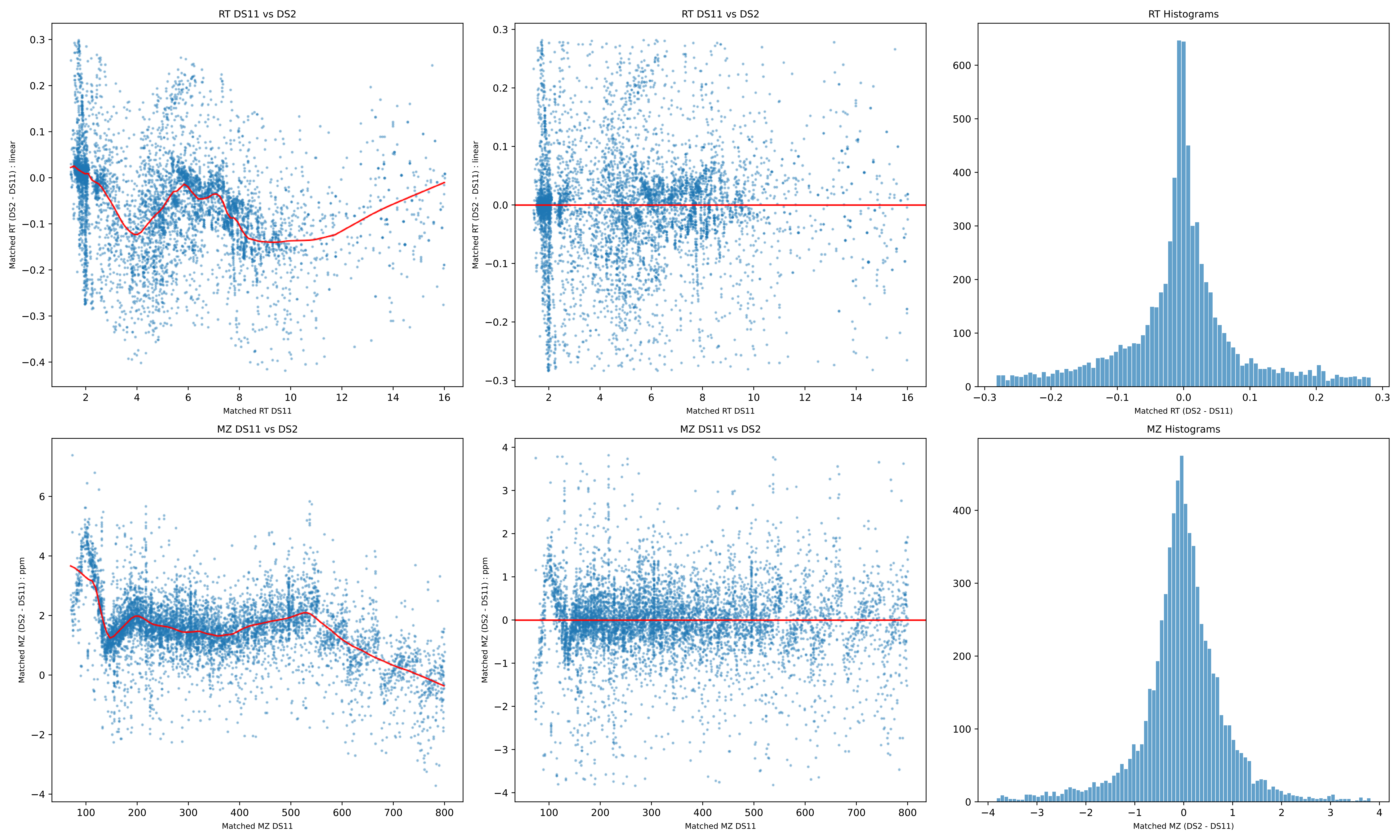
RT DS11 vs DS2
Matched RT (DS2 - DS11) : linear
Matched RT DS11
RT DS11 vs DS2
Matched RT (DS2 - DS11) : linear
Matched RT DS11
RT Histograms
Matched RT (DS2 - DS11)
MZ DS11 vs DS2
Matched MZ (DS2 - DS11) : ppm
Matched MZ DS11
MZ DS11 vs DS2
Matched MZ (DS2 - DS11) : ppm
Matched MZ DS11
MZ Histograms
Matched MZ (DS2 - DS11)

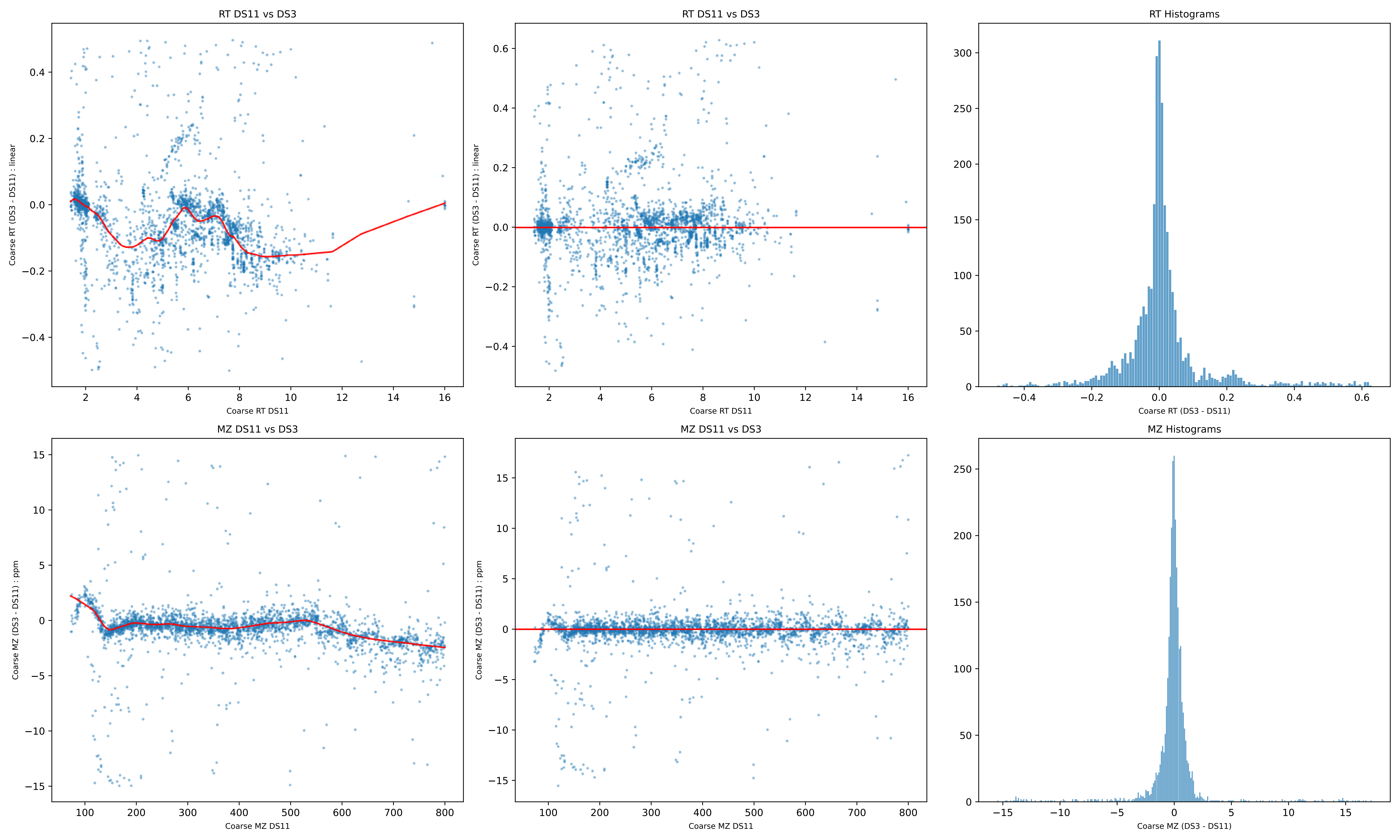
RT DS11 vs DS3
Coarse RT (DS3 - DS11) : linear
Coarse RT DS11
RT DS11 vs DS3
Coarse RT (DS3 - DS11) : linear
Coarse RT DS11
RT Histograms
Coarse RT (DS3 - DS11)
MZ DS11 vs DS3
Coarse MZ (DS3 - DS11) : ppm
Coarse MZ DS11
MZ DS11 vs DS3
Coarse MZ (DS3 - DS11) : ppm
Coarse MZ DS11
MZ Histograms
Coarse MZ (DS3 - DS11)

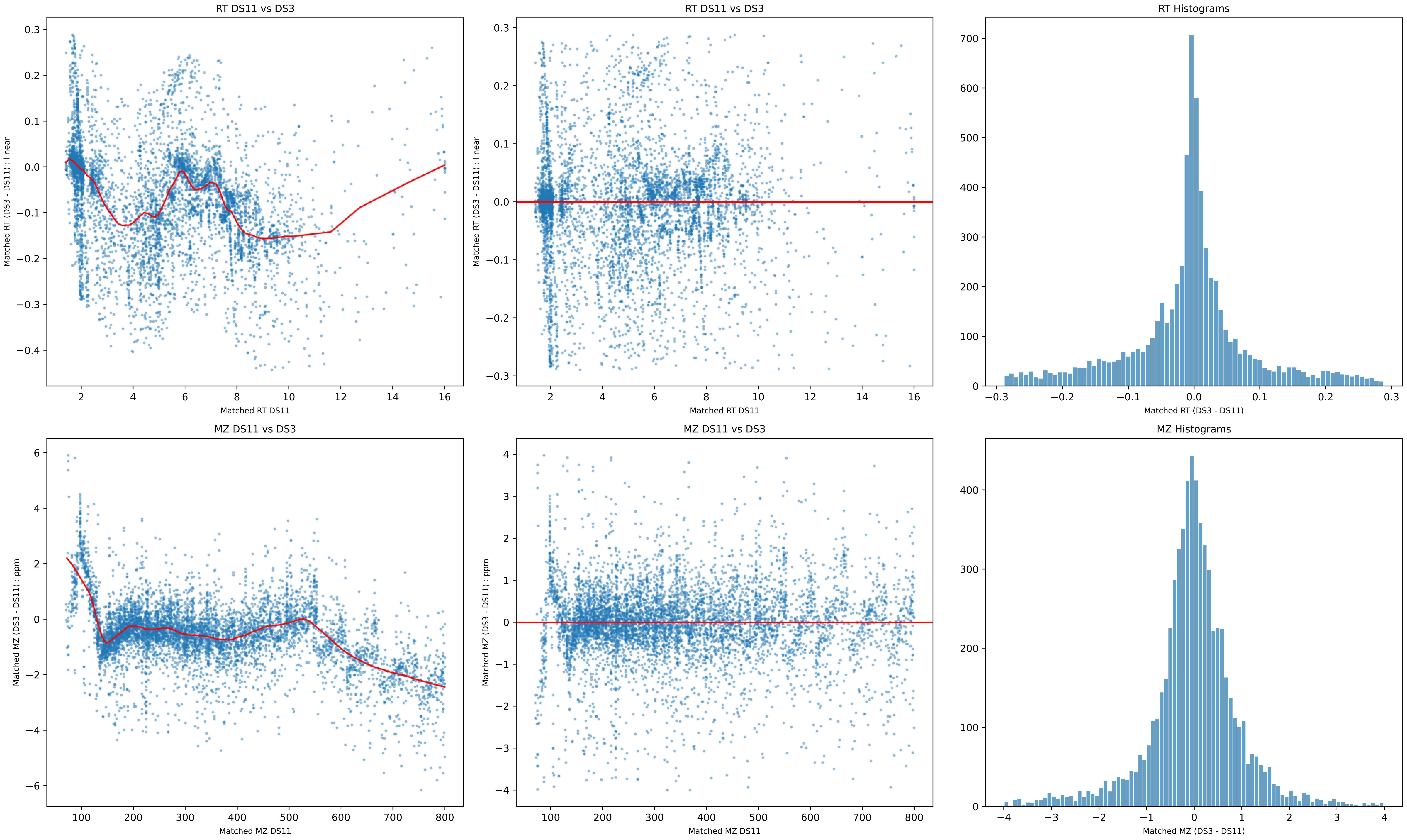
RT DS11 vs DS3
Matched RT (DS3 - DS11) : linear
0.3
0.2
0.1
0.0
−0.1
−0.2
−0.3
−0.4
2
4
6
8
10
12
14
16
Matched RT DS11
RT DS11 vs DS3
Matched RT (DS3 - DS11) : linear
0.3
0.2
0.1
0.0
−0.1
−0.2
−0.3
2
4
6
8
10
12
14
16
Matched RT DS11
RT Histograms
700
600
500
400
300
200
100
0
−0.3
−0.2
−0.1
0.0
0.1
0.2
0.3
Matched RT (DS3 - DS11)
MZ DS11 vs DS3
Matched MZ (DS3 - DS11) : ppm
6
4
2
0
−2
−4
−6
100
200
300
400
500
600
700
800
Matched MZ DS11
MZ DS11 vs DS3
Matched MZ (DS3 - DS11) : ppm
4
3
2
1
0
−1
−2
−3
−4
100
200
300
400
500
600
700
800
Matched MZ DS11
MZ Histograms
400
300
200
100
0
−4
−3
−2
−1
0
1
2
3
4
Matched MZ (DS3 - DS11)

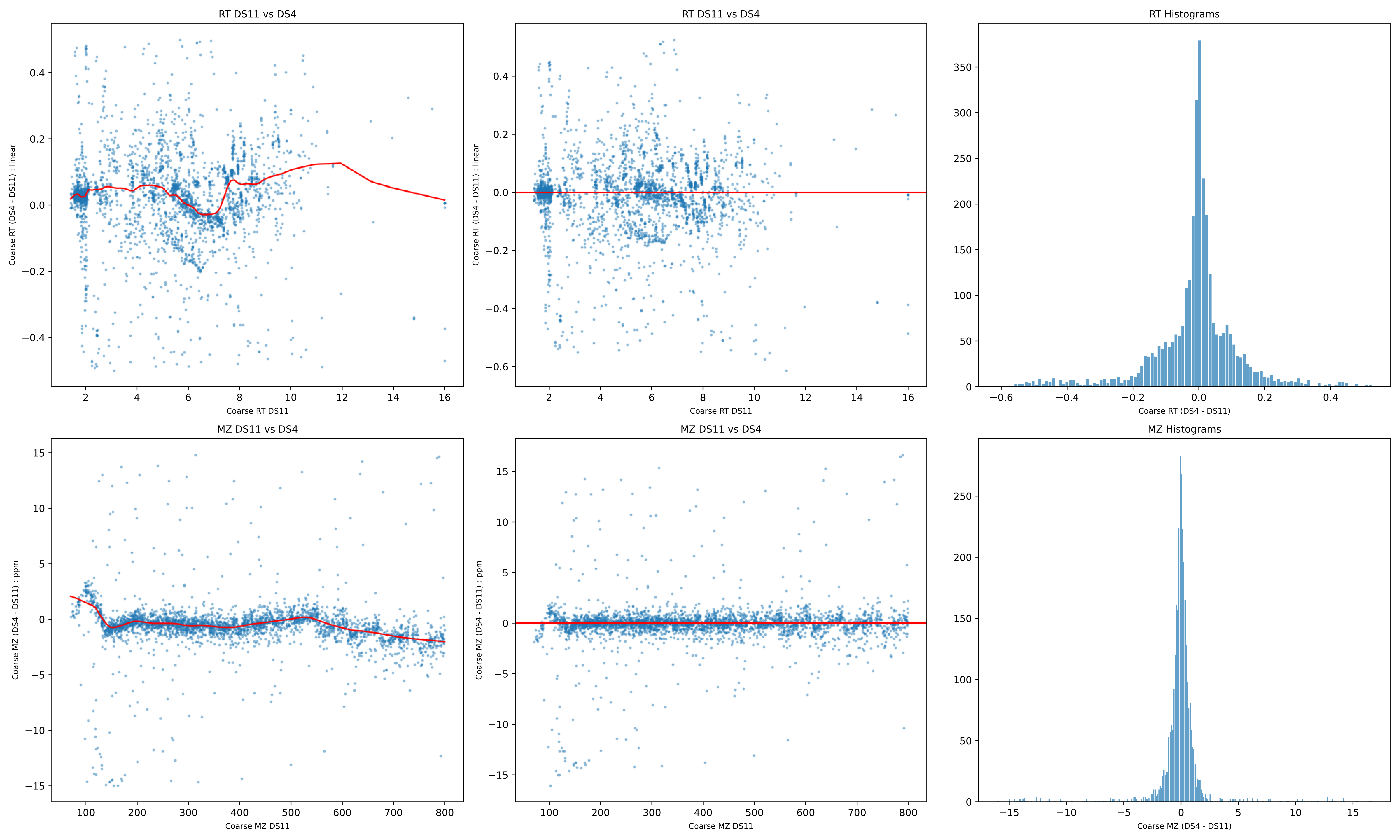
RT DS11 vs DS4
Coarse RT (DS4 - DS11) : linear
Coarse RT DS11
RT DS11 vs DS4
Coarse RT (DS4 - DS11) : linear
Coarse RT DS11
RT Histograms
Coarse RT (DS4 - DS11)
MZ DS11 vs DS4
Coarse MZ (DS4 - DS11) : ppm
Coarse MZ DS11
MZ DS11 vs DS4
Coarse MZ (DS4 - DS11) : ppm
Coarse MZ DS11
MZ Histograms
Coarse MZ (DS4 - DS11)

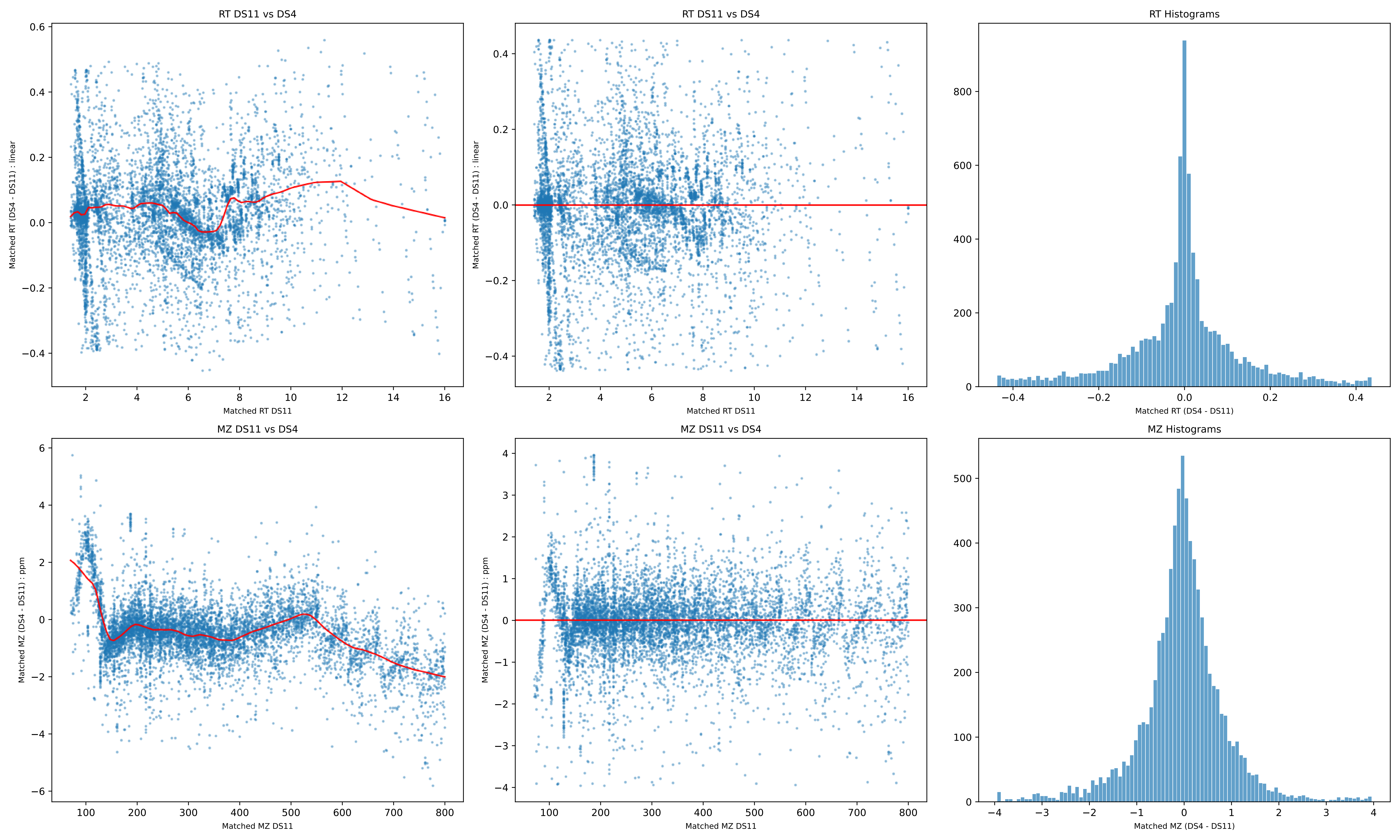
RT DS11 vs DS4
Matched RT (DS4 - DS11) : linear
Matched RT DS11
RT DS11 vs DS4
Matched RT (DS4 - DS11) : linear
Matched RT DS11
RT Histograms
Matched RT (DS4 - DS11)
MZ DS11 vs DS4
Matched MZ (DS4 - DS11) : ppm
Matched MZ DS11
MZ DS11 vs DS4
Matched MZ (DS4 - DS11) : ppm
Matched MZ DS11
MZ Histograms
Matched MZ (DS4 - DS11)

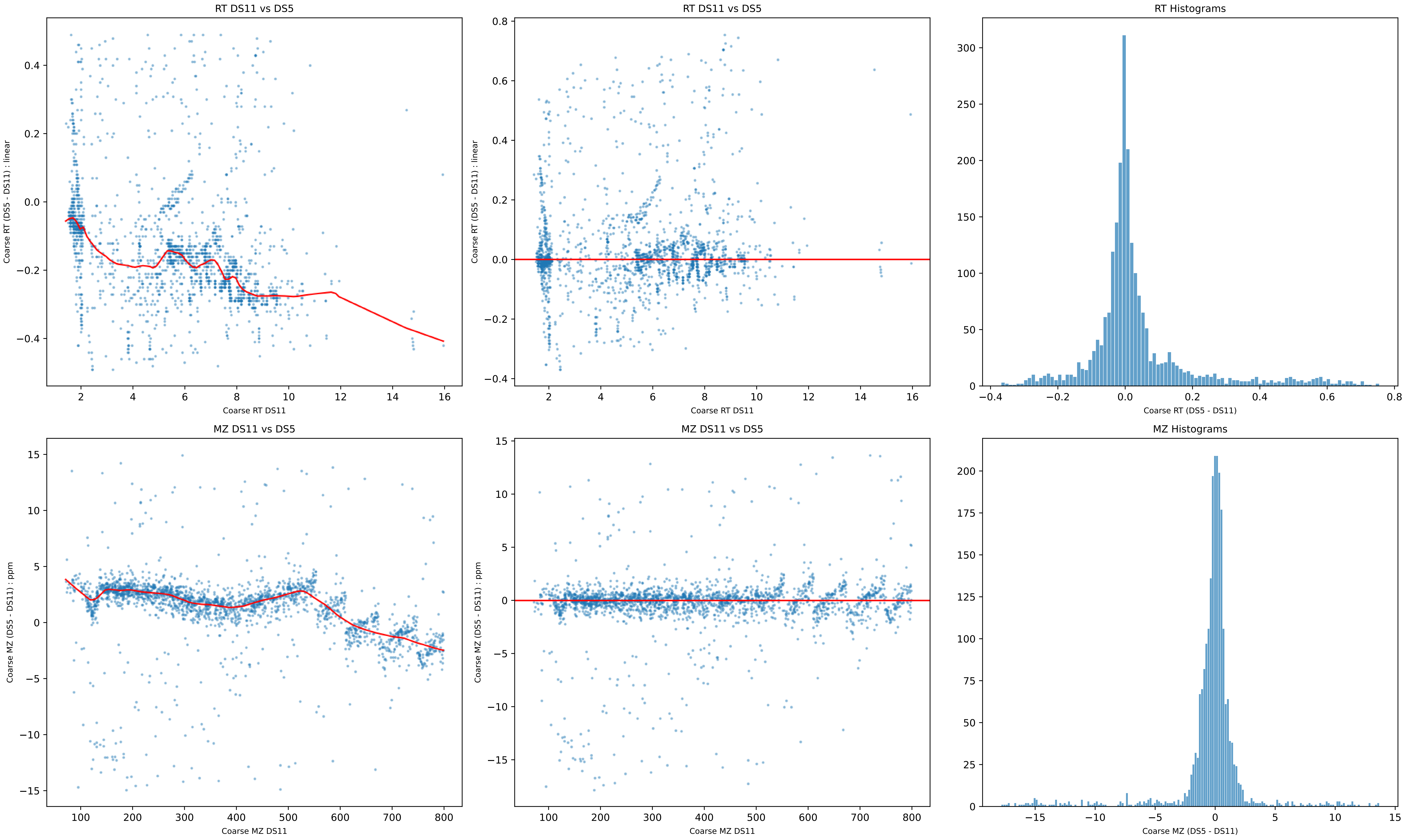
RT DS11 vs DS5
Coarse RT (DS5 - DS11) : linear
Coarse RT DS11
RT DS11 vs DS5
Coarse RT (DS5 - DS11) : linear
Coarse RT DS11
RT Histograms
Coarse RT (DS5 - DS11)
MZ DS11 vs DS5
Coarse MZ (DS5 - DS11) : ppm
Coarse MZ DS11
MZ DS11 vs DS5
Coarse MZ (DS5 - DS11) : ppm
Coarse MZ DS11
MZ Histograms
Coarse MZ (DS5 - DS11)

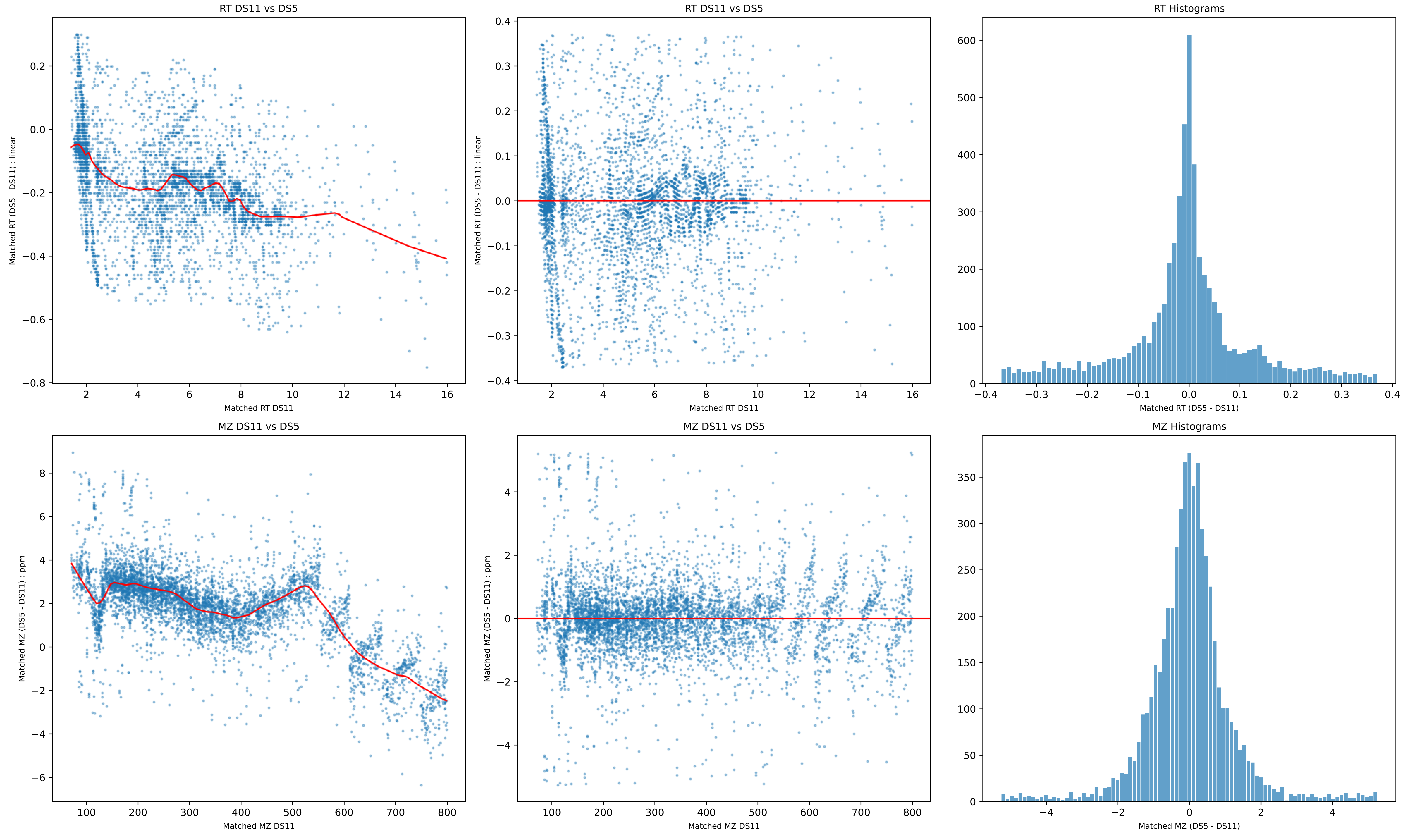

RT DS11 vs DS5
Matched RT (DS5 - DS11) : linear
Matched RT DS11
RT DS11 vs DS5
Matched RT (DS5 - DS11) : linear
Matched RT DS11
RT Histograms
Matched RT (DS5 - DS11)
MZ DS11 vs DS5
Matched MZ (DS5 - DS11) : ppm
Matched MZ DS11
MZ DS11 vs DS5
Matched MZ (DS5 - DS11) : ppm
Matched MZ DS11
MZ Histograms
Matched MZ (DS5 - DS11)

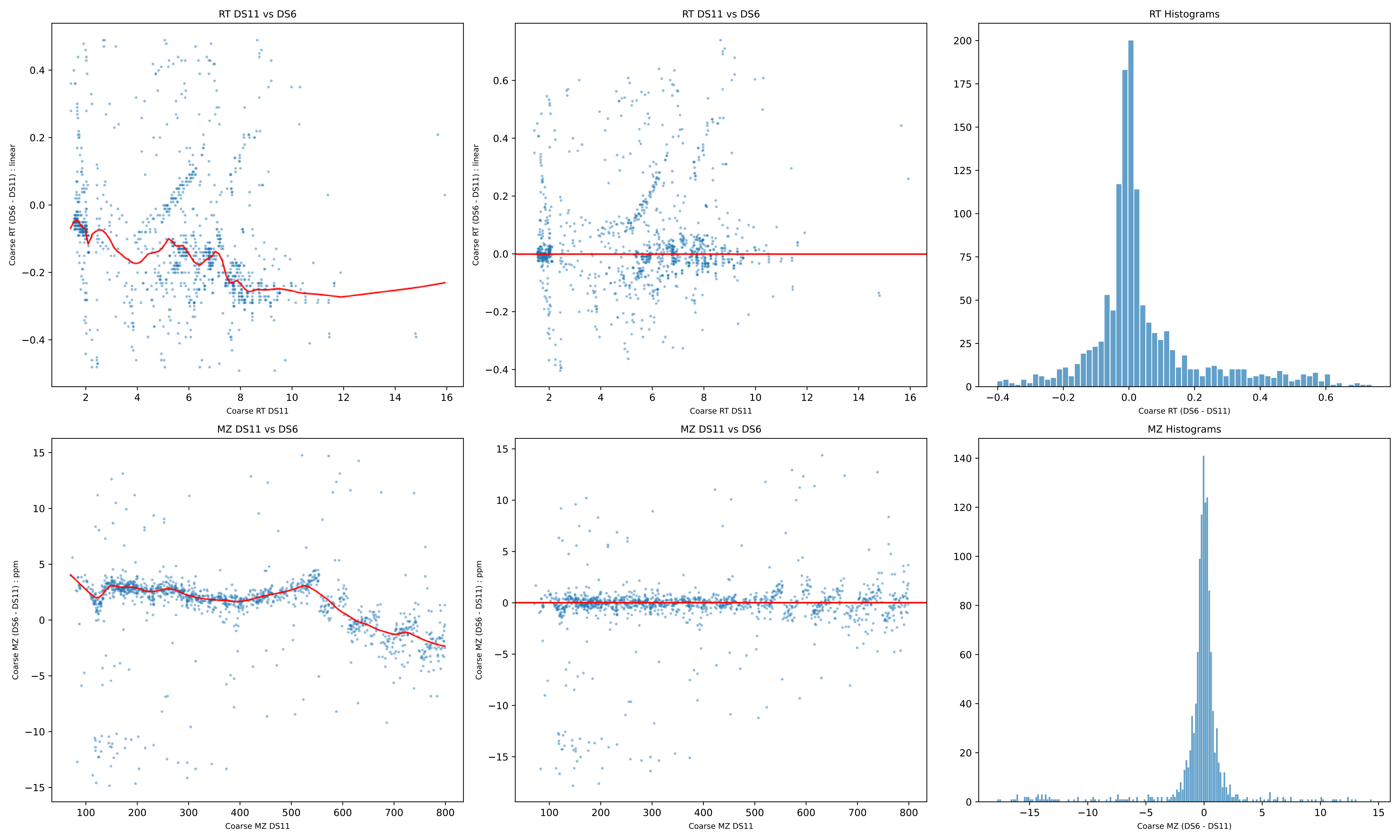

RT DS11 vs DS6
Coarse RT (DS6 - DS11) : linear
Coarse RT DS11
RT DS11 vs DS6
Coarse RT (DS6 - DS11) : linear
Coarse RT DS11
RT Histograms
Coarse RT (DS6 - DS11)
MZ DS11 vs DS6
Coarse MZ (DS6 - DS11) : ppm
Coarse MZ DS11
MZ DS11 vs DS6
Coarse MZ (DS6 - DS11) : ppm
Coarse MZ DS11
MZ Histograms
Coarse MZ (DS6 - DS11)

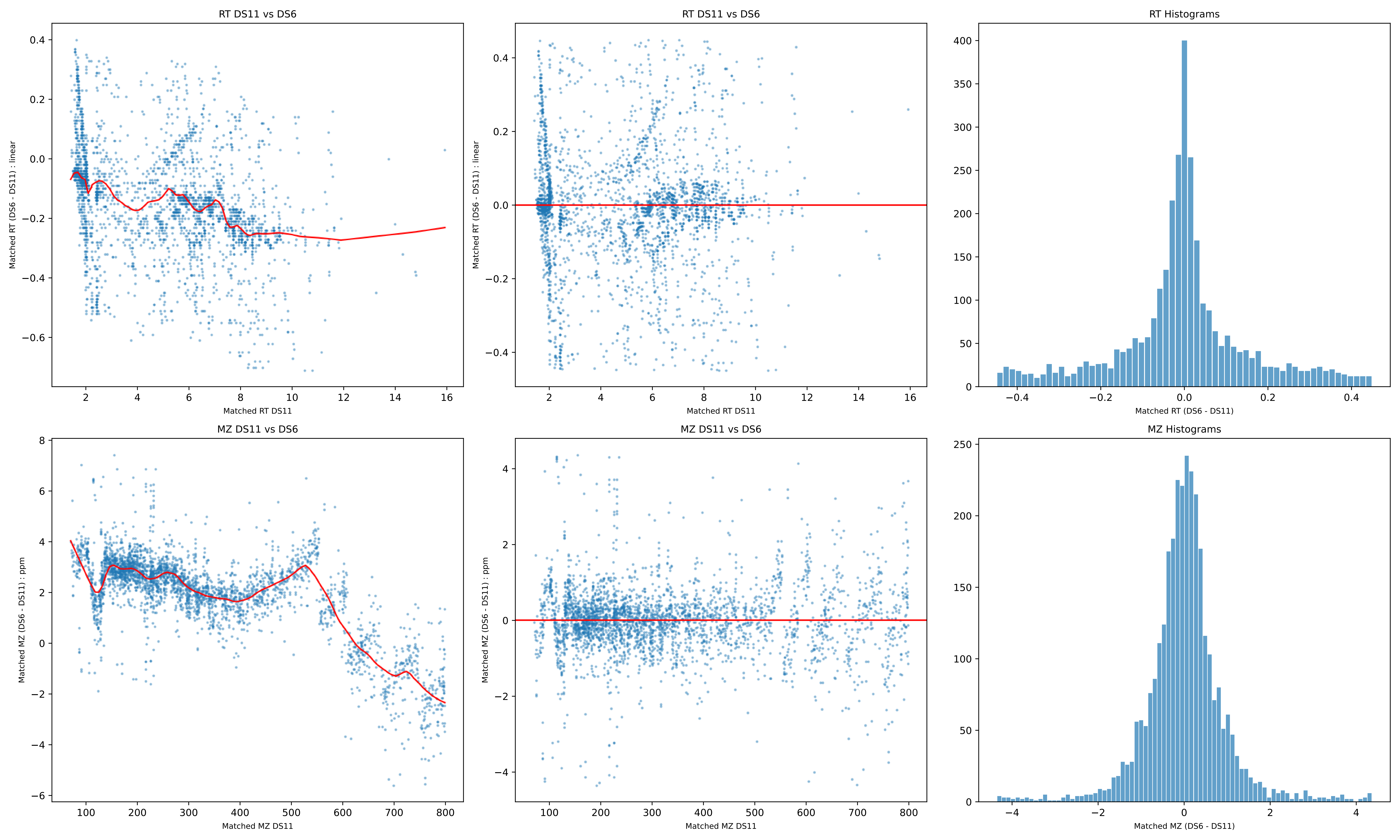
RT DS11 vs DS6
Matched RT (DS6 - DS11) : linear
Matched RT DS11
RT DS11 vs DS6
Matched RT (DS6 - DS11) : linear
Matched RT DS11
RT Histograms
Matched RT (DS6 - DS11)
MZ DS11 vs DS6
Matched MZ (DS6 - DS11) : ppm
Matched MZ DS11
MZ DS11 vs DS6
Matched MZ (DS6 - DS11) : ppm
Matched MZ DS11
MZ Histograms
Matched MZ (DS6 - DS11)

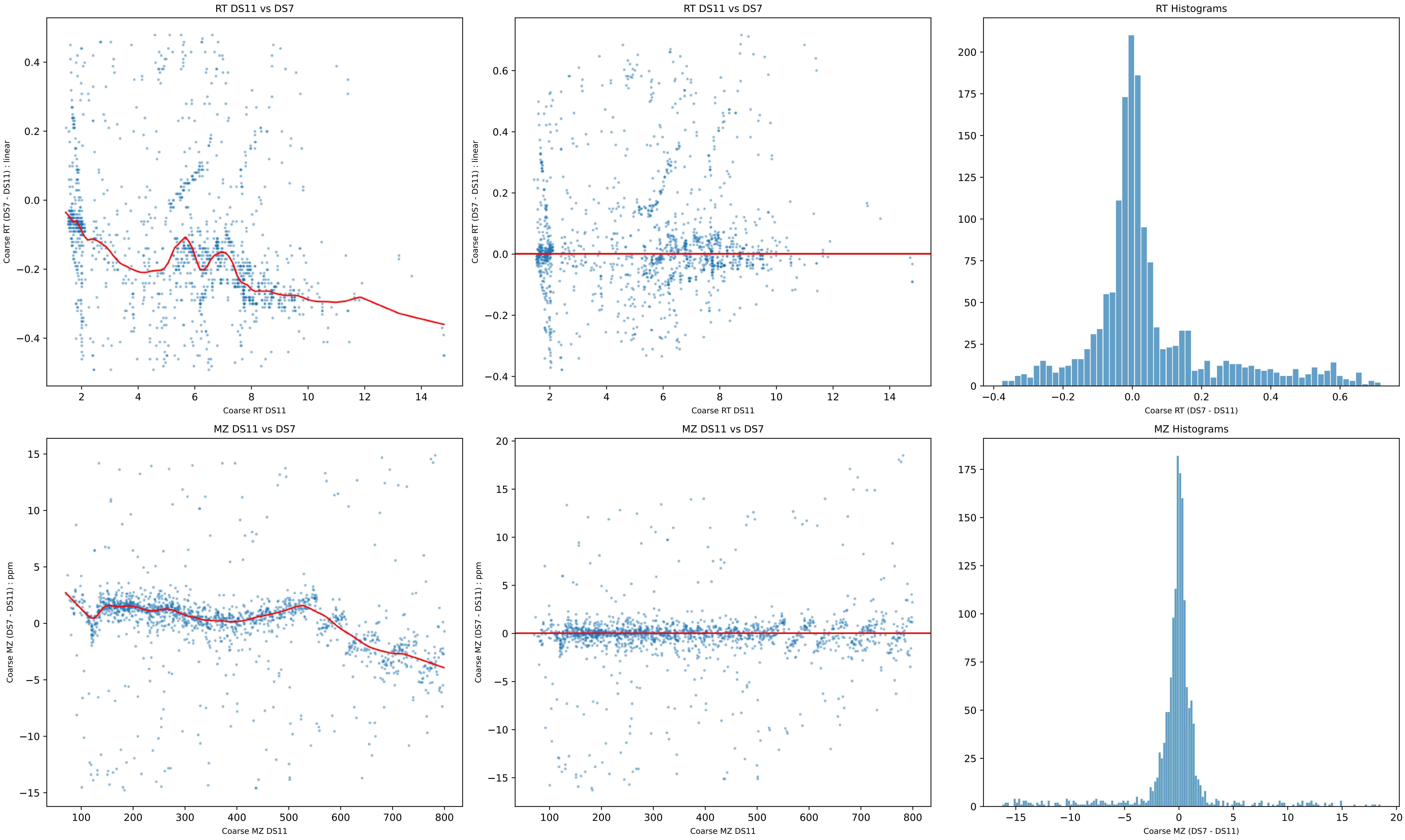

RT DS11 vs DS7
Coarse RT (DS7 - DS11) : linear
Coarse RT DS11
RT DS11 vs DS7
Coarse RT (DS7 - DS11) : linear
Coarse RT DS11
RT Histograms
Coarse RT (DS7 - DS11)
MZ DS11 vs DS7
Coarse MZ (DS7 - DS11) : ppm
Coarse MZ DS11
MZ DS11 vs DS7
Coarse MZ (DS7 - DS11) : ppm
Coarse MZ DS11
MZ Histograms
Coarse MZ (DS7 - DS11)

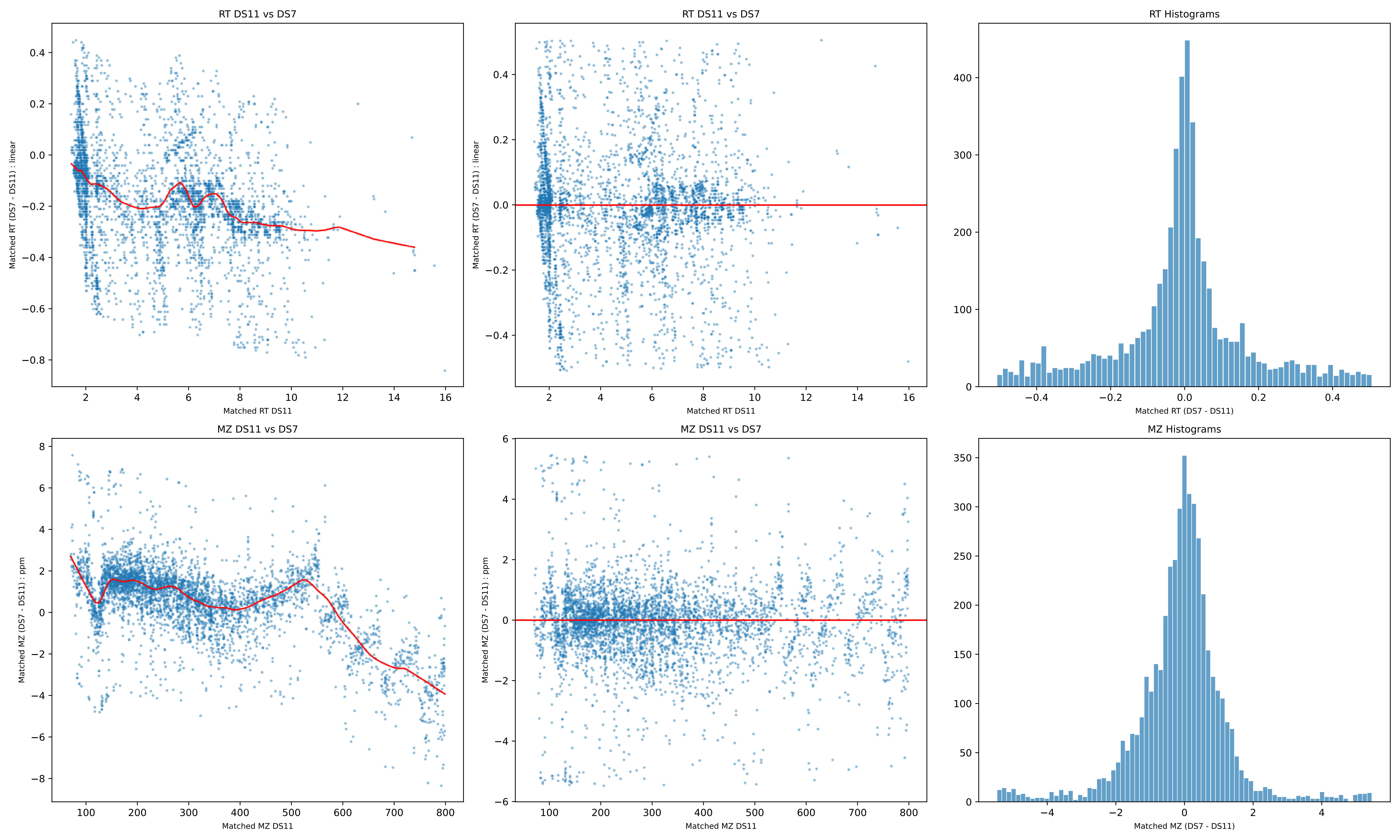
RT DS11 vs DS7
Matched RT (DS7 - DS11) : linear
Matched RT DS11
RT DS11 vs DS7
Matched RT (DS7 - DS11) : linear
Matched RT DS11
RT Histograms
Matched RT (DS7 - DS11)
MZ DS11 vs DS7
Matched MZ (DS7 - DS11) : ppm
Matched MZ DS11
MZ DS11 vs DS7
Matched MZ (DS7 - DS11) : ppm
Matched MZ DS11
MZ Histograms
Matched MZ (DS7 - DS11)

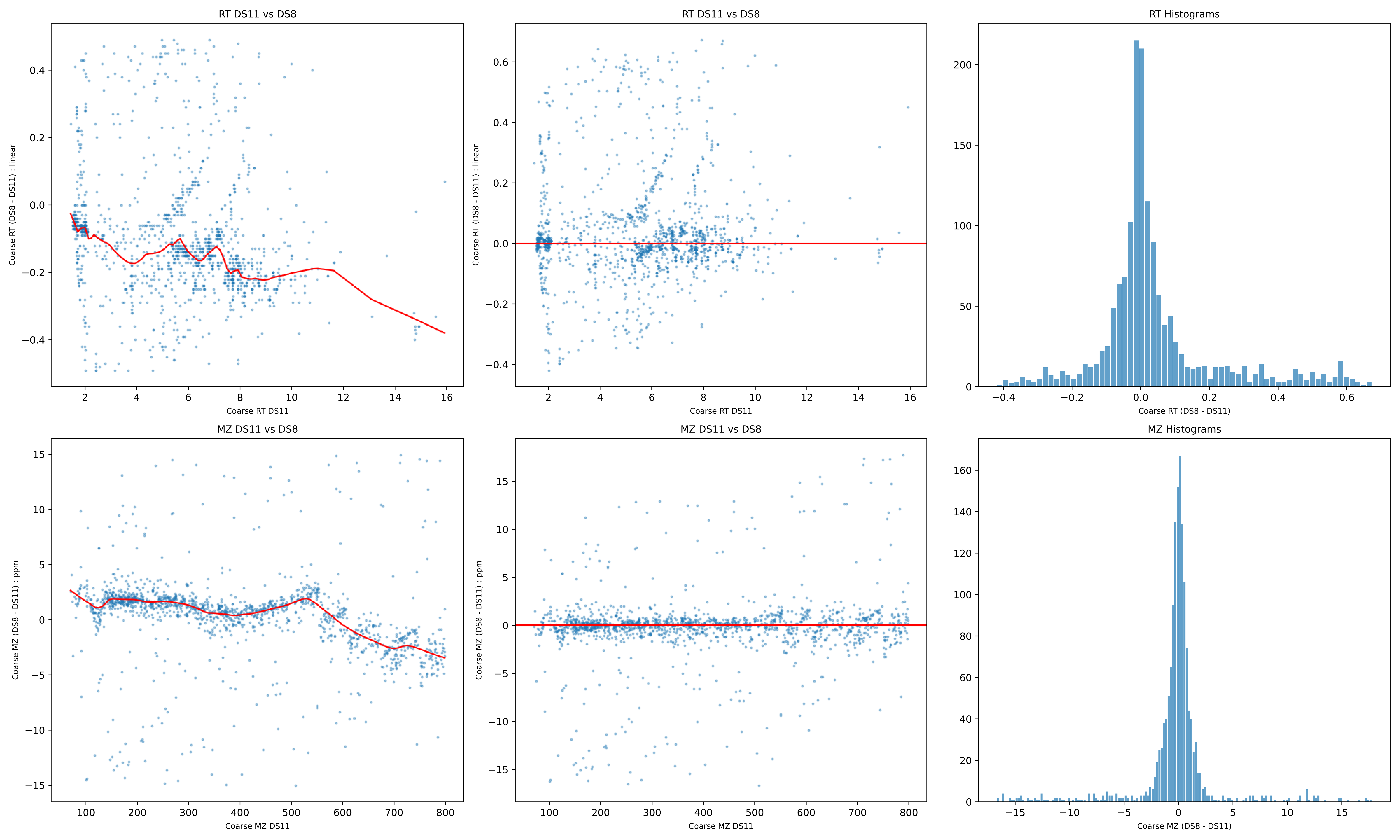
RT DS11 vs DS8
Coarse RT (DS8 - DS11) : linear
Coarse RT DS11
RT DS11 vs DS8
Coarse RT (DS8 - DS11) : linear
Coarse RT DS11
RT Histograms
Coarse RT (DS8 - DS11)
MZ DS11 vs DS8
Coarse MZ (DS8 - DS11) : ppm
Coarse MZ DS11
MZ DS11 vs DS8
Coarse MZ (DS8 - DS11) : ppm
Coarse MZ DS11
MZ Histograms
Coarse MZ (DS8 - DS11)

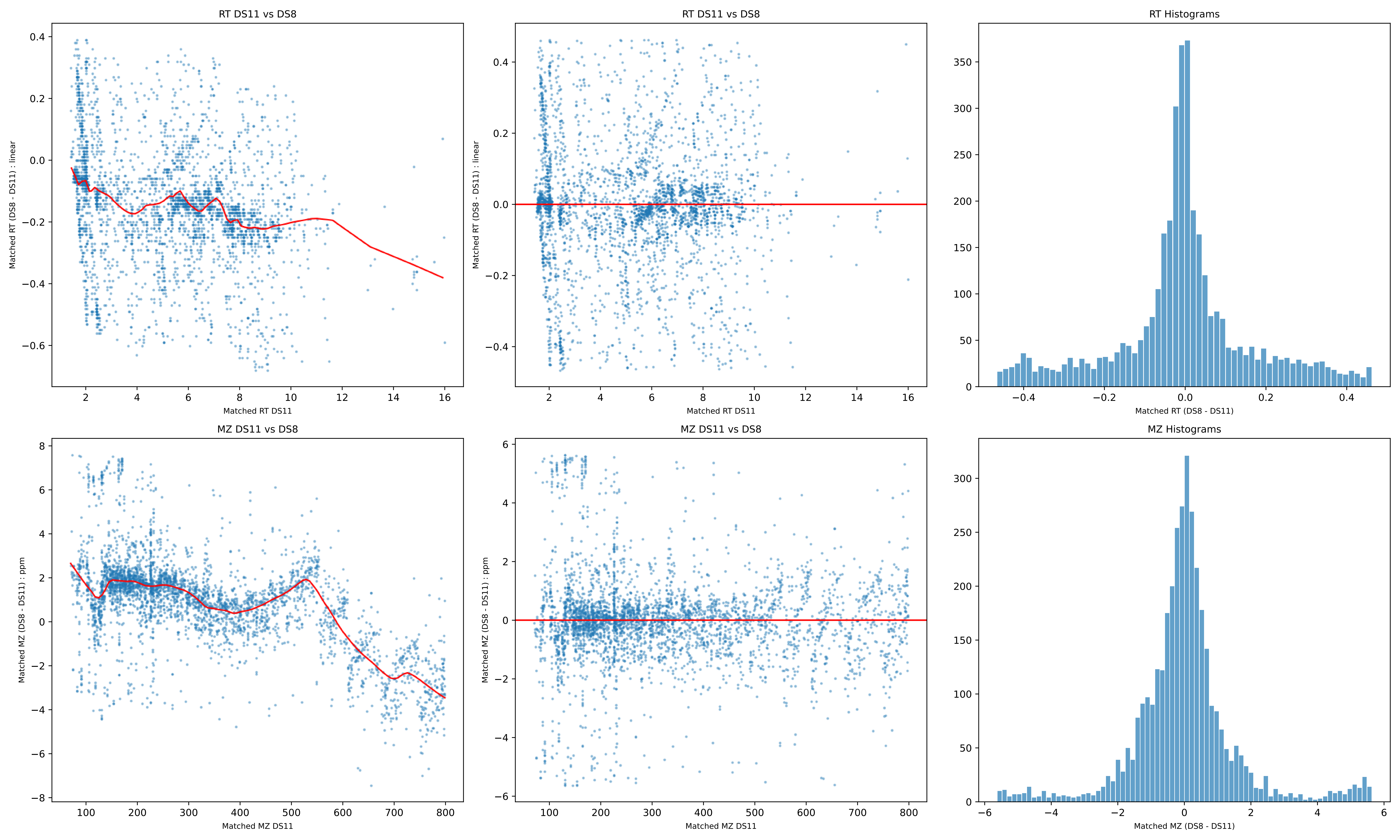

RT DS11 vs DS8
Matched RT (DS8 - DS11) : linear
Matched RT DS11
RT DS11 vs DS8
Matched RT (DS8 - DS11) : linear
Matched RT DS11
RT Histograms
Matched RT (DS8 - DS11)
MZ DS11 vs DS8
Matched MZ (DS8 - DS11) : ppm
Matched MZ DS11
MZ DS11 vs DS8
Matched MZ (DS8 - DS11) : ppm
Matched MZ DS11
MZ Histograms
Matched MZ (DS8 - DS11)

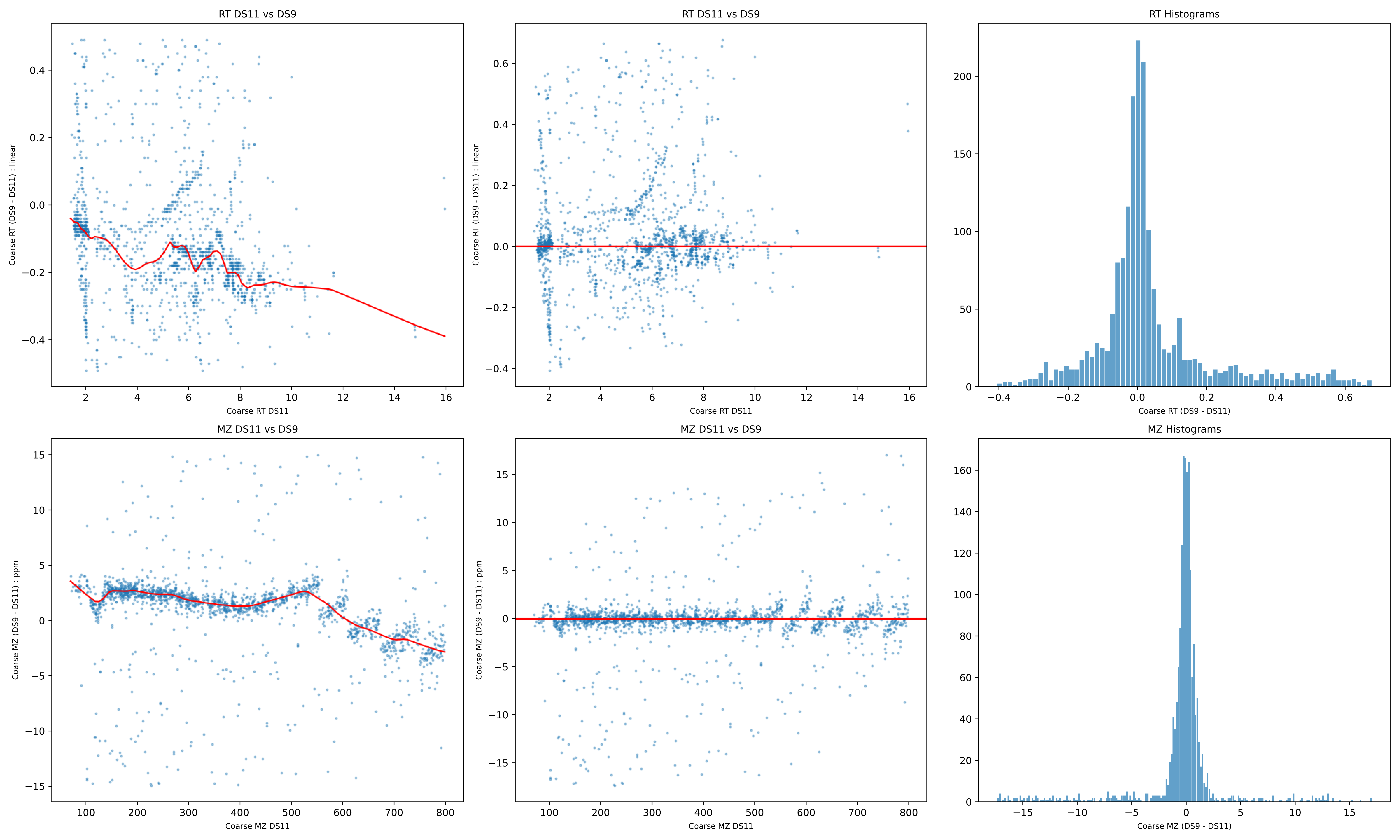

RT DS11 vs DS9
Coarse RT (DS9 - DS11) : linear
Coarse RT DS11
RT DS11 vs DS9
Coarse RT (DS9 - DS11) : linear
Coarse RT DS11
RT Histograms
Coarse RT (DS9 - DS11)
MZ DS11 vs DS9
Coarse MZ (DS9 - DS11) : ppm
Coarse MZ DS11
MZ DS11 vs DS9
Coarse MZ (DS9 - DS11) : ppm
Coarse MZ DS11
MZ Histograms
Coarse MZ (DS9 - DS11)

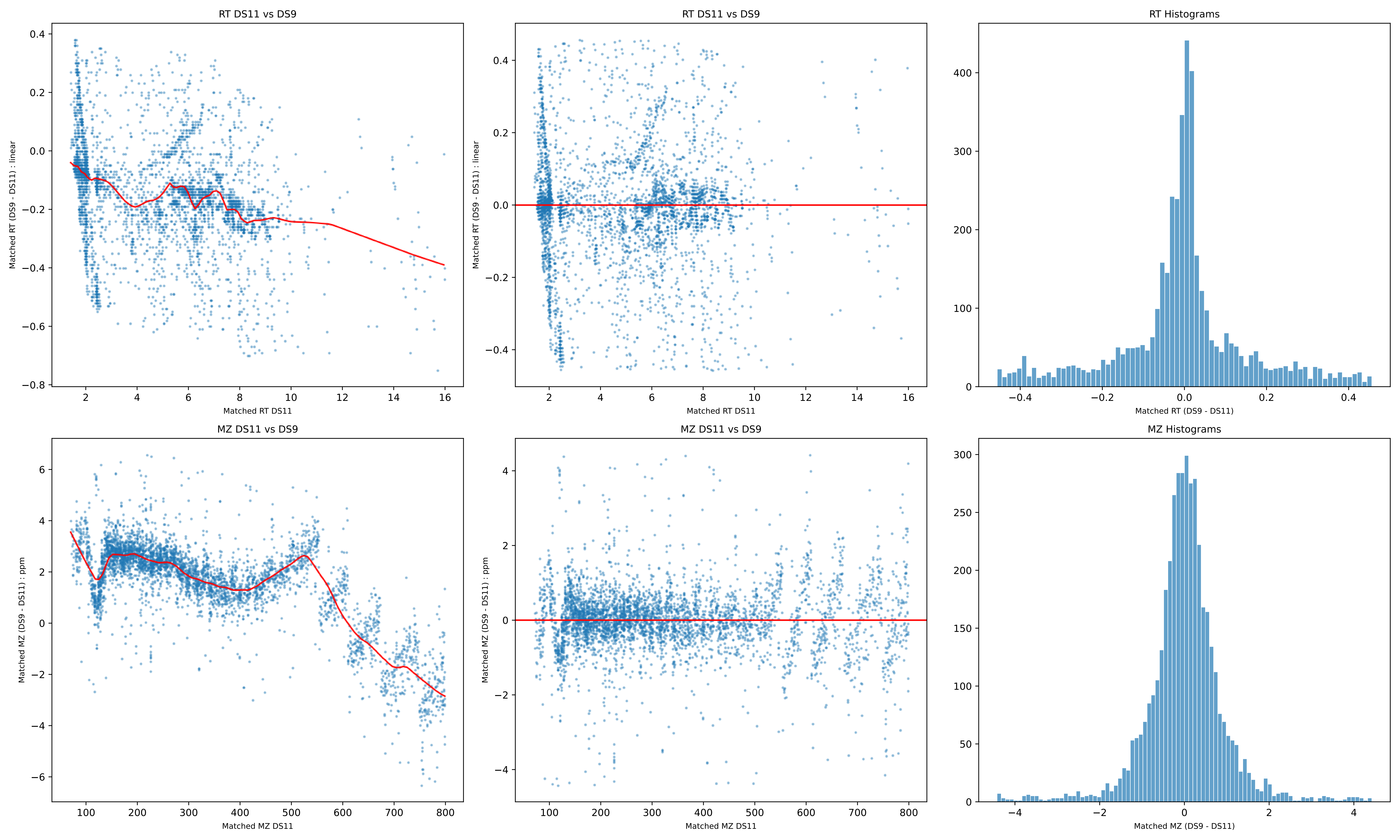
RT DS11 vs DS9
Matched RT (DS9 - DS11) : linear
Matched RT DS11
RT DS11 vs DS9
Matched RT (DS9 - DS11) : linear
Matched RT DS11
RT Histograms
Matched RT (DS9 - DS11)
MZ DS11 vs DS9
Matched MZ (DS9 - DS11) : ppm
Matched MZ DS11
MZ DS11 vs DS9
Matched MZ (DS9 - DS11) : ppm
Matched MZ DS11
MZ Histograms
Matched MZ (DS9 - DS11)

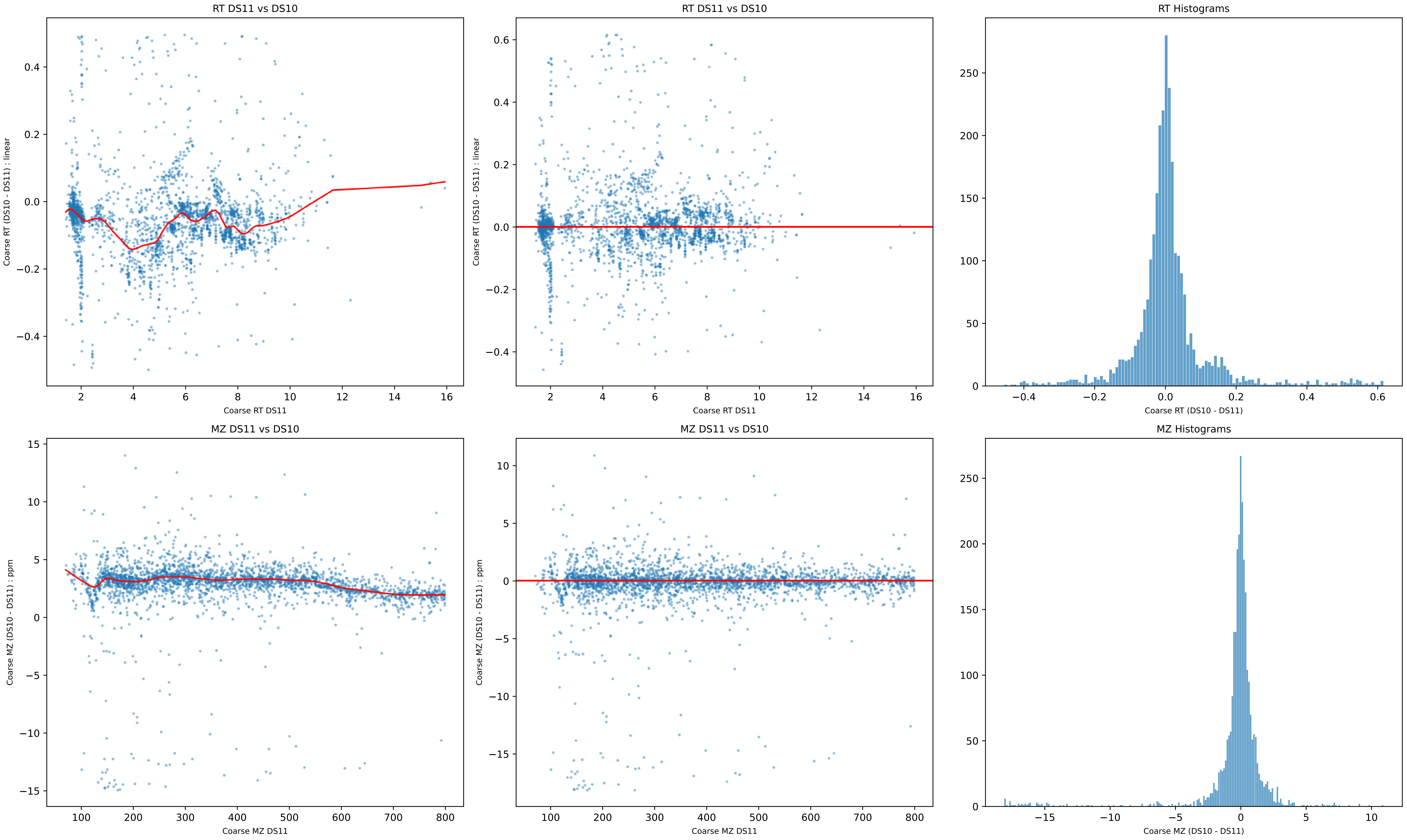
RT DS11 vs DS10
Coarse RT (DS10 - DS11) : linear
Coarse RT DS11
RT DS11 vs DS10
Coarse RT (DS10 - DS11) : linear
Coarse RT DS11
RT Histograms
Coarse RT (DS10 - DS11)
MZ DS11 vs DS10
Coarse MZ (DS10 - DS11) : ppm
Coarse MZ DS11
MZ DS11 vs DS10
Coarse MZ (DS10 - DS11) : ppm
Coarse MZ DS11
MZ Histograms
Coarse MZ (DS10 - DS11)

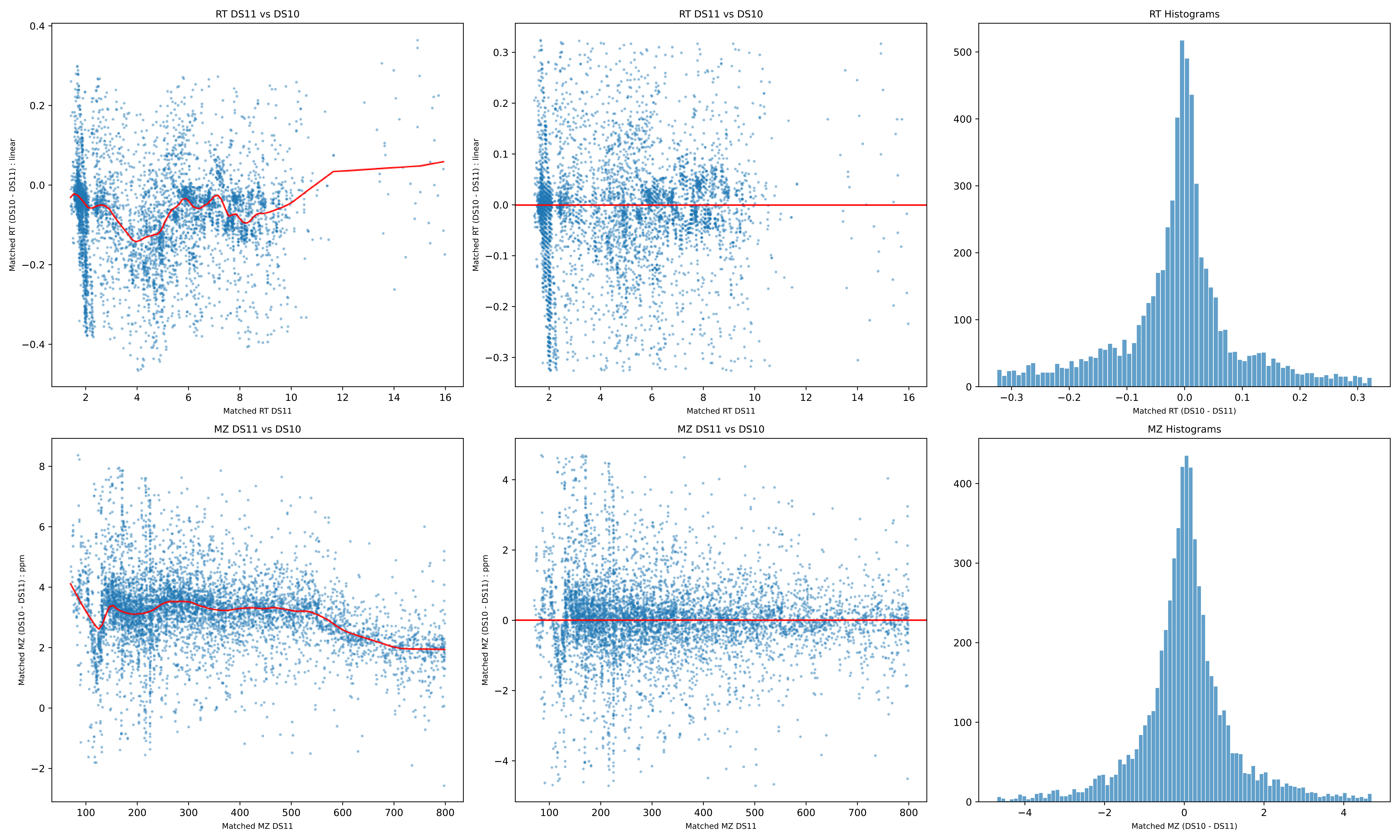
RT DS11 vs DS10
Matched RT (DS10 - DS11) : linear
Matched RT DS11
RT DS11 vs DS10
Matched RT (DS10 - DS11) : linear
Matched RT DS11
RT Histograms
Matched RT (DS10 - DS11)
MZ DS11 vs DS10
Matched MZ (DS10 - DS11) : ppm
Matched MZ DS11
MZ DS11 vs DS10
Matched MZ (DS10 - DS11) : ppm
Matched MZ DS11
MZ Histograms
Matched MZ (DS10 - DS11)
